# Altering microtubule dynamics is synergistically toxic with inhibition of the spindle checkpoint

**DOI:** 10.1101/706077

**Authors:** Klaske M. Schukken, Yi-Chih Lin, Michael Schubert, Stephanie F. Preuss, Judith E. Simon, Hilda van den Bos, Zuzana Storchova, Maria Colome-Tatche, Holger Bastians, Diana C.J. Spierings, Floris Foijer

**Affiliations:** European Research Institute for the Biology of Ageing, University of Groningen, University Medical Centre Groningen, A. Deusinglaan 1, Groningen, 9713 AV, The Netherlands; Goettingen Center for Molecular Biosciences and University Medical Center, Goettingen, Germany; Department of Molecular Genetics, University of Kaiserslautern, Germany; Institute of Computational Biology, Helmholtz Center Munich, German Research Center for Environmental Health, Neuherberg, Germany; TUM School of Life Sciences Weihenstephan, Technical University of Munich, Freising, Germany

## Abstract

Chromosome instability (CIN) and aneuploidy are hallmarks of cancer. As the majority of cancers are aneuploid, targeting aneuploidy or CIN may be an effective way to target a broad spectrum of cancers. Here, we perform two small molecule compound screens to identify drugs that selectively target cells that are aneuploid or exhibit a CIN phenotype. We find that aneuploid cells are much more sensitive to the energy metabolism regulating drug ZLN005 than their euploid counterparts. Furthermore, cells with an ongoing CIN phenotype, induced by spindle assembly checkpoint (SAC) alleviation, are significantly more sensitive to the Src kinase inhibitor SKI606. We show that inhibiting Src kinase increases microtubule polymerization rates and, more generally, that deregulating microtubule polymerization rates is particularly toxic to cells with a defective SAC. Our findings therefore suggest that tumors with a dysfunctional SAC are particularly sensitive to microtubule poisons and, vice versa, that compounds alleviating the SAC provide a powerful means to treat tumors with deregulated microtubule dynamics.

## Introduction

Chromosomal INstability (CIN) is the process through which chromosomes mis-segregate during mitosis. CIN leads to cells with an abnormal DNA content, a state known as aneuploidy. As 3 out of 4 cancers are aneuploid (Weaver & Cleveland, 2006; Foijer *et al*, 2008; Duijf *et al*, 2013), CIN is considered an important contributor to tumorigenesis. Indeed, CIN has been associated with metastasis (Bloomfield & Duesberg, 2016; Xu *et al*, 2016), increased probability of drug resistance (Sansregret & Swanton, 2017; Lee *et al*, 2011) and generally, a lowered patient survival (McGranahan *et al*, 2012; Carter *et al*, 2006; Walther *et al*, 2008). While the frequent occurrence of CIN and resulting aneuploidy in cancer is generally attributed to the acquired ability of cancer cells to adapt their palette of oncogenic features as the tumor evolves, ongoing chromosome missegregation also has negative effects on cancer cells. The downside of CIN for cancer cells is that most newly acquired karyotypes lead to reduced proliferation (Torres *et al*, 2007; Williams *et al*, 2008; Foijer *et al*, 2017) and induction of aneuploidy-imposed stresses (Torres *et al*, 2010). In addition to this, ongoing missegregation causes further structural DNA damage (Zhang *et al*, 2015; MacKenzie *et al*, 2017) that, together with unfavorable karyotypes, leads to cell death (Kops *et al*, 2004; Burds *et al*, 2005; Santaguida *et al*, 2017) or senescence (Andriani *et al*, 2016).

To protect from CIN, cells have mechanisms in place that maintain proper chromosome inheritance. The Spindle Assembly Checkpoint (SAC) is one such mechanism preventing CIN by inhibiting the onset of anaphase until all chromosomes are properly attached to the two opposing spindle poles, reviewed in detail by Musacchio and Salmon (Musacchio & Salmon, 2007). Interfering with the SAC, for instance by inactivating key components of the checkpoint, leads to frequent chromosome mis-segregation events, and is commonly used to study the consequences of CIN *in vitro* and *in vivo* (Foijer *et al*, 2013, 2014; Kops *et al*, 2004; Foijer *et al*, 2017).

While complete loss of SAC function is rare in human cancer (Gordon *et al*, 2012), many cancers show signs of a partially impaired SAC, for instance as a result of increased expression of proteins with a direct role in the SAC or their regulators, such as Rb mutations that lead to increased expression of Mad2 and thus provoke a CIN phenotype (Pfau & Amon, 2012). Furthermore, altered microtubule dynamics are another source of CIN (28) in many cancers (Stolz *et al*, 2015; Ertych *et al*, 2014) as restoring tubulin dynamics to normal levels can decrease CIN rates in many cancer cell lines (Ertych *et al*, 2014). Conversely, commonly-used cancer drugs such as Paclitaxel or Vincristine interfere with microtubule polymerization rates thus increasing CIN rates in cancer cells. This observation suggests that imposing CIN phenotypes onto cancer cells is a powerful strategy to eradicate tumors. However, it is not yet clear whether exacerbating CIN in cells with a preexisting CIN phenotype is wise or not.

As CIN and aneuploidy discriminate cancer cells from healthy cells, both make for attractive targets for cancer therapy. To reveal potential general vulnerabilities of aneuploid cells, Tang *et al.* performed a small molecule compound screen, which revealed the energy stress-inducing compound AICAR to be more toxic to aneuploid cells than euploid cells (Tang *et al*, 2011b). This aneuploidy-specific toxicity was shown to be true in cell culture experiments as well as in cancer mouse models, a promising result for future aneuploid cancer therapies.

While CIN and aneuploidy are intimately related, CIN has additional effects on cell physiology and growth in addition to those imposed by the resulting aneuploidy (Schukken & Foijer, 2017). Since CIN drives karyotype heterogeneity thus increasing the rate of evolution that cancer cells use to acquire new features and adapt (Giam & Rancati, 2015; McGranahan *et al*, 2012), targeting CIN would provide an even more powerful means to kill cancer cells than aneuploidy alone.

In this study we therefore performed two small-scale drug screens, one to identify small molecule compounds that target aneuploid cells and another to find compounds that are more toxic to CIN cells than to chromosomally stable cells. For this purpose, we selected a collection of drug-like molecules from a list of drugs already being used in the clinic, or in advanced stage clinical trials. Compounds were further selected for their potential role in targeting CIN or aneuploid cells, such as targeting cell survival (Dekanty *et al*, 2012; Foijer *et al*, 2013), proliferation (Gogendeau *et al*, 2015; Ben-david *et al*, 2014; Sheltzer *et al*, 2017; Williams *et al*, 2008), protein processing (Oromendia *et al*, 2012; Stingele *et al*, 2012), DNA repair (Bakhoum *et al*, 2014, 2018), transcriptional deregulation (Upender *et al*, 2004; Stingele *et al*, 2012), and cellular metabolism (Tang *et al*, 2011b; Williams *et al*, 2008) as these processes are typically deregulated in aneuploid cells. Indeed, our screen for aneuploidy-targeting compounds revealed a compound targeting cellular metabolism, validating earlier findings from the Amon lab (Tang *et al*, 2011a). Furthermore, the CIN screen revealed that the Src inhibitor Bosutinib is synergistically toxic to cells with an alleviated SAC. We find that the mechanism underlying the toxicity of Bosutinib in SAC-deficient cells results from deregulated tubulin polymerization rates imposed by Src inhibition. Our results therefore indicate that combining SAC inhibition with tubulin deregulation is synergistically toxic to cells and might provide a powerful means to target cancer cells with a CIN phenotype.

## Results

CIN and the resulting aneuploidy lead to a deregulated transcriptome and proteome (Stingele *et al*, 2012; Foijer *et al*, 2013, 2014; Tang *et al*, 2011b), and can provoke senescence (Andriani *et al*, 2016; Santaguida *et al*, 2017) or apoptosis (Giam & Rancati, 2015). Furthermore, ongoing CIN can lead to further DNA damage (Zhang *et al*, 2015; MacKenzie *et al*, 2017). We therefore reasoned that targeting RNA or protein processing, transcriptional regulation, apoptosis, or DNA repair might be particularly toxic to aneuploid cells and cells exhibiting a CIN phenotype. As CIN and aneuploidy are different concepts (Schukken & Foijer, 2017) and have different consequences for cells (Schukken & Foijer, 2017; Stingele *et al*, 2012; Andriani *et al*, 2016), aneuploidy and CIN might impose different therapeutic vulnerabilities. To test this, we performed two small-scale drug screens, one to identify compounds that selectively kill aneuploid cells and another to identify small molecules that selectively kill CIN cells.

### A small-scale drug screen to identify compounds that selectively kill aneuploid cells

We first selected 95 drug-like-molecules from a drug library composed of drugs that target processes that aneuploid or CIN cells might rely on and are already being used in the clinic, or being tested in clinical trials (**Sup. Table 1**). Next, we determined the initial drug concentration for each drug to be used in the screen. For this, we exposed wildtype RPE1 cells (a diploid non-cancer cell line derived from retinal epithelium (Soto *et al*, 2017)) to decreasing concentrations of the drugs, starting at 10 μM for all compounds, and compared cell proliferation of drug-exposed cells to proliferation of DMSO-treated cells over a period of 7 days. We purposely chose a non-transformed cell line, as this allows studying the combinational effect of CIN and drugs in an otherwise unperturbed setting.

Next, we subjected stable aneuploid RPE-1 cells, trisomic for chromosomes (chrs.) 5 and 12 (**Sup. Fig. 1A**, (Stingele *et al*, 2012)), to the same drug-treatment regime and compared proliferation between diploid and aneuploid RPE1 cells (Sup. Data 1) using an Incucyte high content imager. **Sup. Fig. 1B** schematically shows the experimental design and analysis approach. Note that aneuploid RPE1 cells showed a modestly reduced proliferation rate compared to control RPE1 cells (**Sup. Fig. 1C**) in line with earlier observations (Williams *et al*, 2008) for which we corrected when analyzing the growth curves. To quantify differences between diploid and aneuploid RPE1 cells, we compared the area under the curve (AUC) as a measure of cumulative cell growth (**Fig. 1A, B**) and the slope of the logarithmic growth as a measure for the proliferation rate (**Fig. 1C, D**), also see Materials and Methods. While this screen revealed some drugs for which aneuploid RPE1 cells were more sensitive (log2>0; p<0.05) or less sensitive (log2<0; p<0.05), we only found one compound (#2379 (ZLN005; a transcriptional regulator of PGC-1α) for which the effect was significant after Bonferroni multiple testing correction (**Fig. 1A**) in one of the two screens. The combined effects of aneuploidy and ZLN005 act synergistically as assessed by a Bliss independence test (50% stronger effect than additive, p=3.2E-3) (Zhao *et al*, 2014). Indeed, further validation confirmed the selective growth defect of aneuploid RPE1 cells imposed by ZLN005 (**Fig. 1E**). However, as ZLN005 targets energy metabolism, very similar to what others have found for AICAR (Tang *et al*, 2011b), we did not pursue this compound further. We therefore conclude that our aneuploidy screen did not uncover novel targetable vulnerabilities of aneuploid cells and next performed a screen for compounds that selectively kill CIN cells.

**Figure 1.**
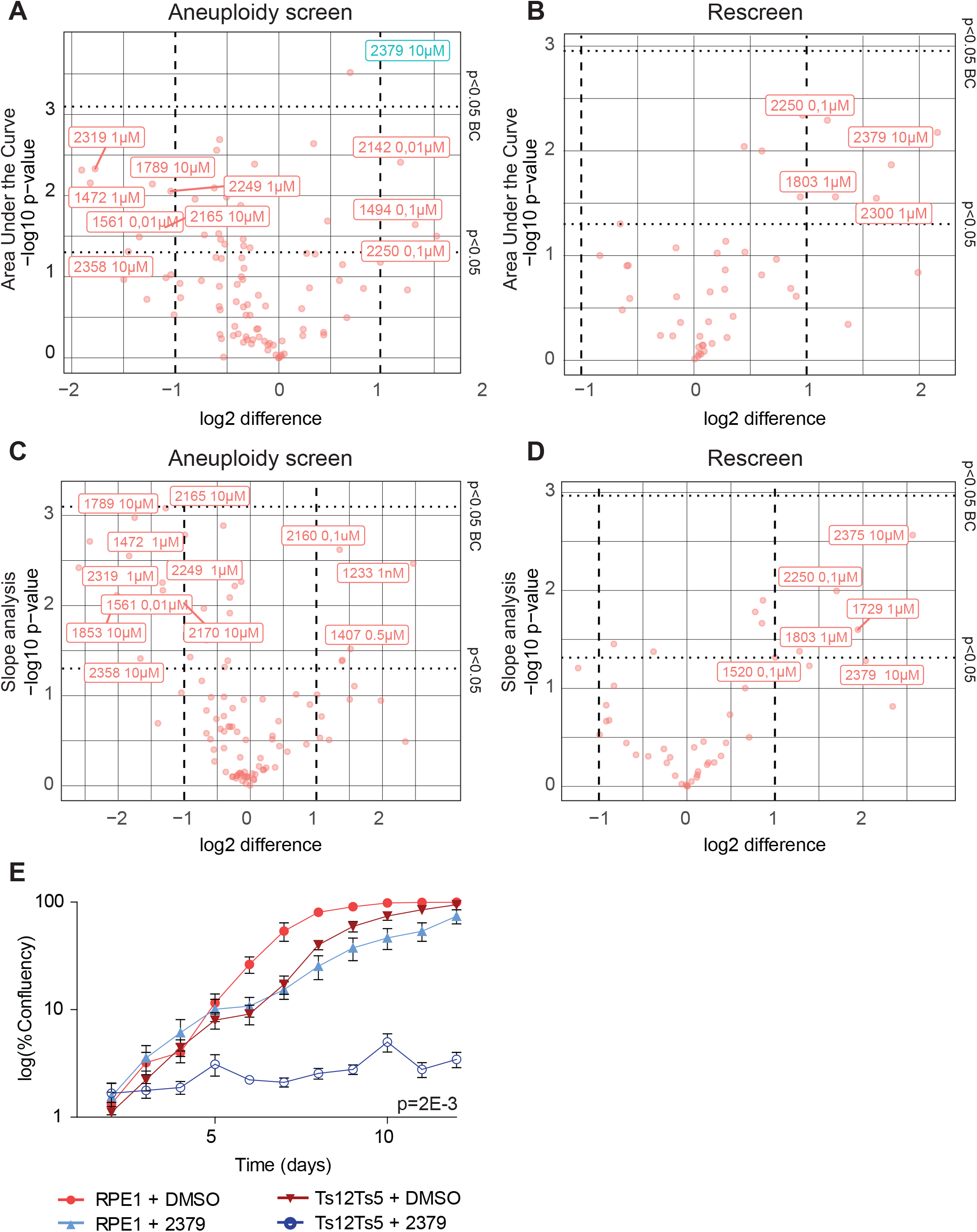
Aneuploid cells are sensitive to a metabolism-enhancing drug. **(A-D)** RPE1 control cells and stable aneuploid RPE1 Ts12 Ts5 cells were screened with 95 drugs, each drug screened in triplicate. 45 drugs were rescreened. The p-values and the log difference between a drug’s effect on RPE1 and RPE1 Ts12 Ts5 cells were plotted. Data was analyzed through quantification of Area under the Curve (AUC, **A, B**), and slope analysis (**C, D**) of both the initial screen (**A, C**) and rescreened drugs (**B, D**). Drugs with difference >1 and p-value<0,05 after Bonferroni correction are indicated in blue. (**E**) Validation growth curves of RPE1 control and RPE1 Ts12 Ts5 cells with and without 10 μM 2379. Data obtained by sequential daily microscope images, analyzed by FIJI-Phantast. All data involves at least 3 biological replicates, each with 3 technical replicates. Error bars indicate standard error of the mean (SEM). P-values are calculated in two-sided t-test for AUC, correcting for cell line control. DMSO control curves are shared with Sup. Fig. 1C & Sup. Fig. 3G.

### A conditional Mad2 knockdown cell line to model chromosomal instability

To screen for compounds that selectively kill cells with a CIN phenotype, we needed a cell line in which CIN can be provoked in an inducible fashion, as long-term CIN phenotypes are typically selected against in tissue culture (Kops *et al*, 2004; Foijer *et al*, 2014). For this, we engineered RPE1 hTert cells in which the SAC can be inhibited through expression of a Doxycycline (dox)-inducible Mad2 shRNA construct, from here on referred to as Mad2 conditional knockdown (Mad2^cKD^) RPE1 cells. Mad2 knockdown efficiency was quantified by quantitative PCR (**Fig. 2A**) and Western blot (**Fig. 2B**), which revealed that Mad2 levels were reduced by 90% within 3 days of dox treatment. To test whether Mad2 inhibition was sufficient to alleviate the SAC, we exposed cells to the microtubule poison nocodazole, determined accumulation in mitosis by quantifying phospho-histone H3 using flow cytometry and found that dox-treatment for 3 days or longer was sufficient to completely alleviate the SAC in Mad2^cKD^ RPE1 cells (**Fig. 2C**). Therefore, for all follow-up experiments involving Mad2^cKD^ RPE1 cells, cells were treated with doxycycline for a minimum of 3 days. As expected, we found that Mad2^cKD^ moderately decreased cell proliferation (~25%), which we corrected for in our downstream analyses (**Sup. Fig. 2A**). Next, we determined whether SAC inhibition in Mad2^cKD^ RPE1 cells indeed leads to a CIN phenotype. To this aim, we quantified interphase and mitotic abnormalities using live cell imaging (**Fig. 2D, 2E**). Indeed, Mad2^cKD^ cells displayed a significantly increased CIN-rate: 46% of the Mad2^cKD^ RPE1 cells displayed mitotic abnormalities compared to only 1% of control cells. Additionally, the fraction of cells with interphase remnants of mitotic aberrations such as micronuclei increased from 2% to 24%. Finally, we quantified aneuploidy by single cell whole genome sequencing (scWGS, (van den Bos *et al*, 2016; Bakker *et al*, 2016)). While control RPE1 cells show little aneuploidy (2 out of 114 cells sequenced) except for a known structural abnormality for chr. 10 (**Fig. 2F**, and (Worrall *et al*, 2018)), 45% of dox-treated Mad2^cKO^ cells displayed multiple aneuploidies per cell (76 out of 169 cells, **Fig. 2G**) within 5 days after induction of the Mad2 shRNA, confirming a substantial CIN phenotype. Together these features make the Mad2^cKD^ cells highly suitable to screen for compounds that kill CIN cells.

**Figure 2.**
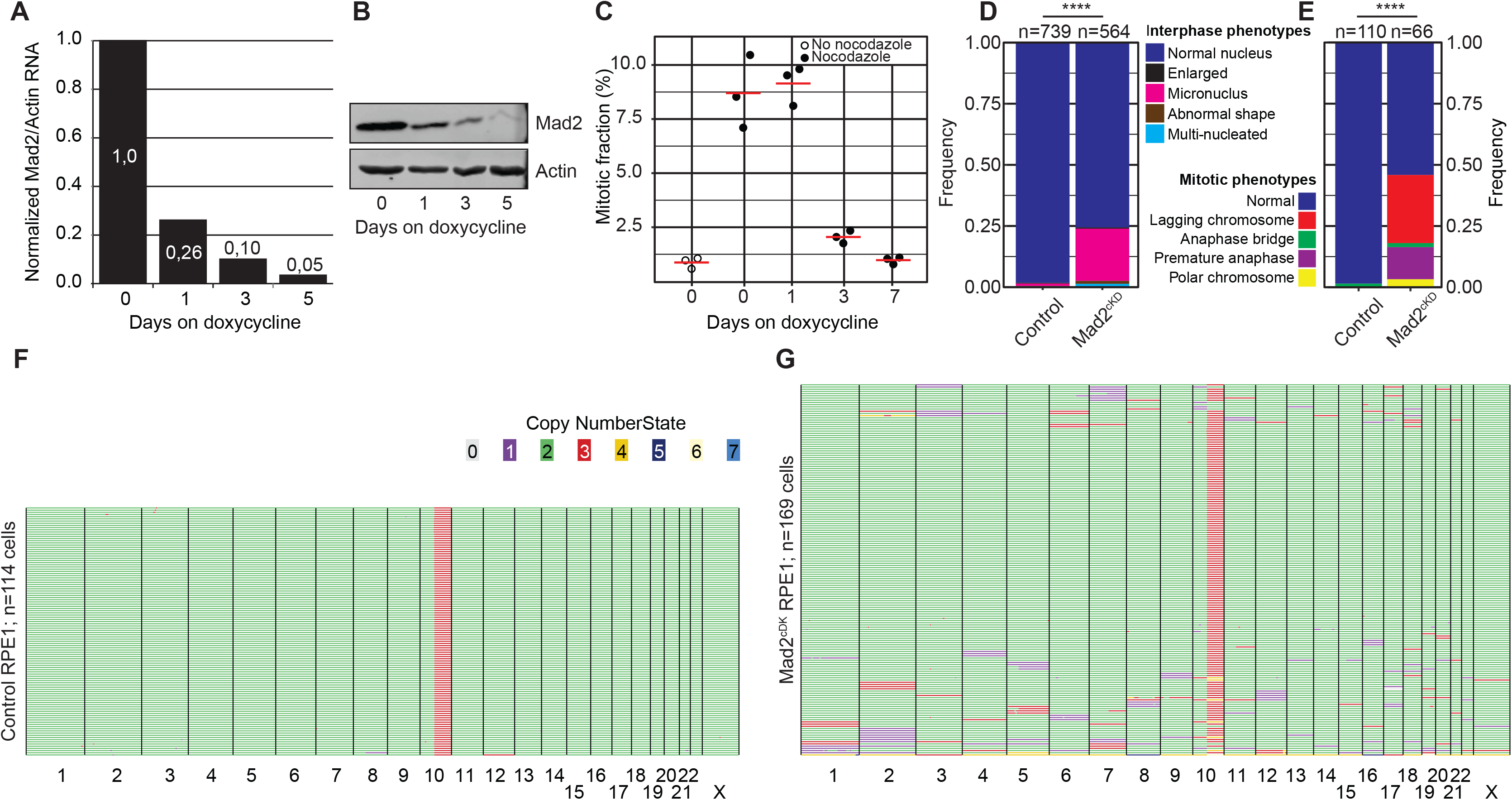
Engineering a cell line for conditional CIN. (**A**) Quantitative PCR for Mad2 RNA levels over time in Mad2^cKD^ RPE1 cells. (**B**) Western blot for Mad2 levels over time in RPE1 in Mad2^cKD^ RPE1 cells (**C**) Mitotic accumulation of nocodazole-challenged control and Mad2^cKD^ RPE1 cells measured by phosphorylated Histone H3. (**D, E**) Quantification of mitotic phenotypes of control and Mad2^cKD^ RPE1 cells assessed by time-lapse imaging for interphase cells (**D**) and mitotic cells (**E**). “n” refers to the number of cells analyzed, p-values from Chi-squared test. Data also displayed in Fig. 5H. (**F, G**) Single cell whole genome sequencing data quantified by AneuFinder for RPE1 control cells (**F**, 114 cells, 2 aneuploid) and Mad2^cKD^ RPE1 cells following 5 days of Doxycycline treatment (**G**, 169 cells, 76 aneuploid). Colors refer to the copy number state for each chromosome (fragment).

### The Src inhibitor SKI606 selectively kills cells Mad2^cKD^ cells

We next employed the Mad2^cKD^ RPE1 cells to screen for compounds that selectively kill CIN cells (**Sup. Fig 2B**). For this, we exposed control and Mad2^cKD^ RPE1 cells to 58 compounds (Sup. Table 2) and compared the maximum proliferation rate and cumulative cell number between Mad2^cKD^ RPE1 cells and control RPE1 cells using the same setup as for the aneuploidy screen described above. To assess both short term and longer-term effects of the drugs, we quantified proliferation and cumulative cell number over the first 4 days and over days 5-8 separately (**Sup. Fig. 2B**). Intriguingly, we found that the mTor inhibitor AZD8055 (compound #1561) at 0.1 μM acted synergistically with CIN in reducing cell numbers (31% greater than additive effect; p= 2.7E-4, Bliss independence test) during the first 4 days of the screen, but became fully toxic to both control and Mad2^cKD^ cells from day 5 onward (**Fig. 3A, B**, Sup. Data 2 for all growth curves). Conversely, we found that the Src inhibitor SKI606 (compound #1407) at 0.1 μM acted synergistically with CIN (48% greater effect than additive; p=7.3E-3, Bliss independence test) during the second half of the screen (**Fig. 3C, D**, Sup. Data 2) and less so during the first half of the screen. Note that the observed effects were not related to the doxycycline treatment required to induce Mad2 shRNA, as doxycycline alone had no effect on proliferation (**Sup. Fig. 3A**). Next, we wanted to validate our findings in independent growth assays. In addition to AZD8055 and SKI606, we also retested compounds #2180 (TMP195; HDAC inhibitor), #2250 (CHR6494 trifluoroacetate, Haspin inhibitor)), #2831 (EPZ015666, Prmt5 inhibitor) #1801 (pyroxamide, HDAC1 inhibitor), #1803 (MS 275, HDAC 1 and 3 inhibitor), and #2008 (Tenovin 1, SIRT 1 & 2 inhibitor) that also showed some effect in the primary CIN screen. For these validation experiments, proliferation was quantified by daily cell confluency measurements from microscope images as described in Materials and Methods. These experiments revealed that while SKI606 (#1407), AZD8055 (#1561) and EPZ015666 (#2831) reproducibly inhibited the growth of Mad2^cKD^ RPE1 cells more that the growth of control cells (**Fig. 4A-F**), this was not the case for TMP195, CHR6494, pytoxamide, MS 275 and Tenovin (**Sup. Fig. 3B-G**). Given that SKI606 gave the largest growth inhibitory effect on Mad2^cKD^ RPE1 cells, most notably at 0.5 μM (**Fig. 4E**), we decided to further pursue this compound. It is interesting to note that SKI606 had no significant effect on stable aneuploid cells (**Sup. Fig. 3G-J**), and vice versa, that ZLN005 (#2379), identified in the aneuploidy screen had no significant effect on Mad2^cKD^ CIN cells (**Sup. Fig. 3K**), suggesting that compounds that are selectively toxic to stable aneuploid cells are not necessarily toxic to CIN cells, and vice versa.

**Figure 3.**
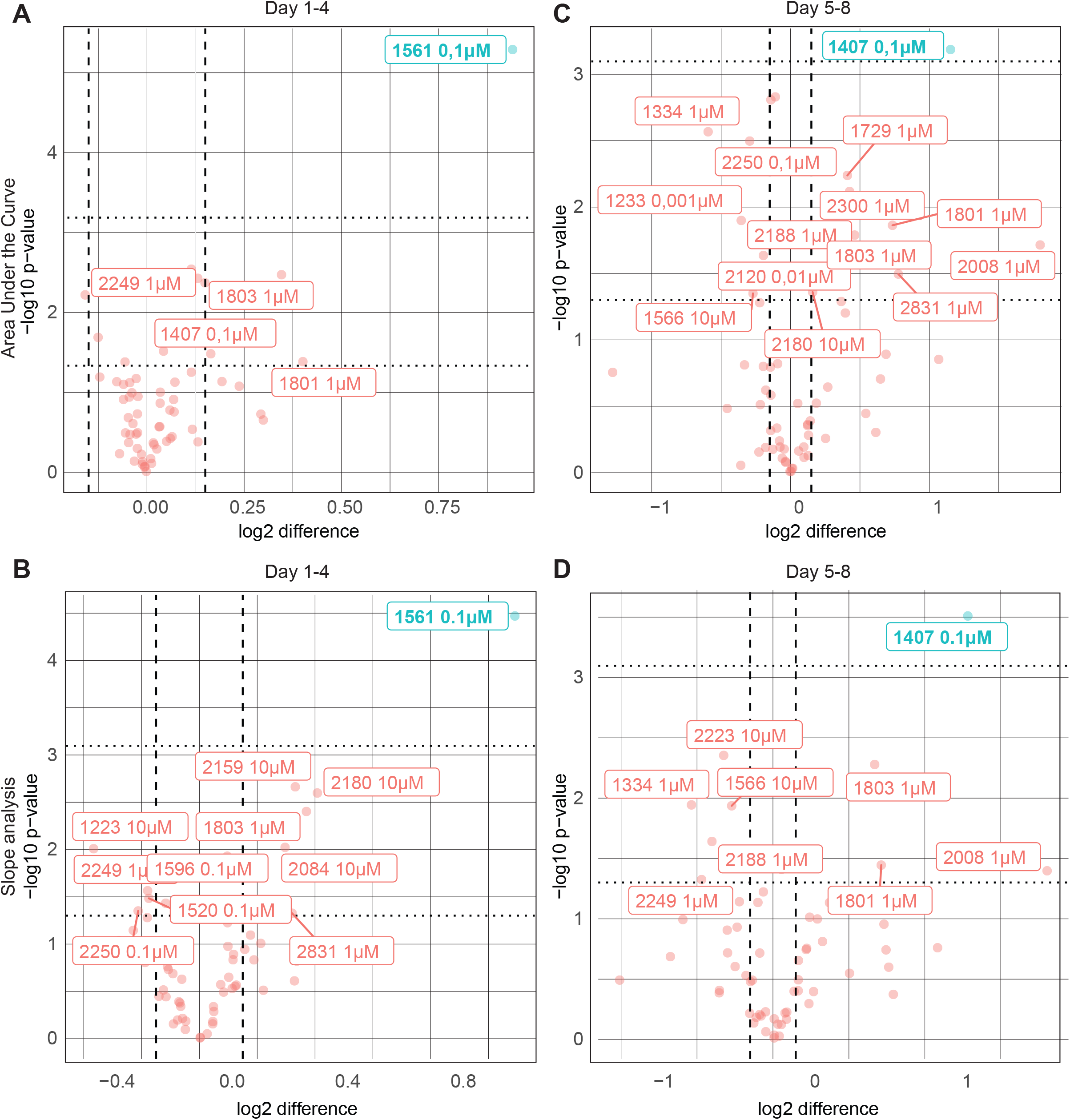
A screen for compounds that selectively kill CIN cells reveals several candidates. (**A-D**) Growth curves of control and Mad2^cKD^ RPE1 cells were analyzed during the first half (day 1-4) (**A. B**), and the second half (day 5-8) of the screen (**C, D**). Both AUC (**A, C**) and slope analysis (**B, D**) was used to quantify the data. The log (base 2) of the difference between CIN and control growth curves per drug was plotted against the negative log (base 10) of the p-value. Dashed vertical lines refer to a log difference of +/− 0.15. All drugs with log of difference >|0.15|, and p-value <0.05 are plotted; drugs with p-values<0.05 after Bonferroni correction are labeled blue.

**Figure 4.**
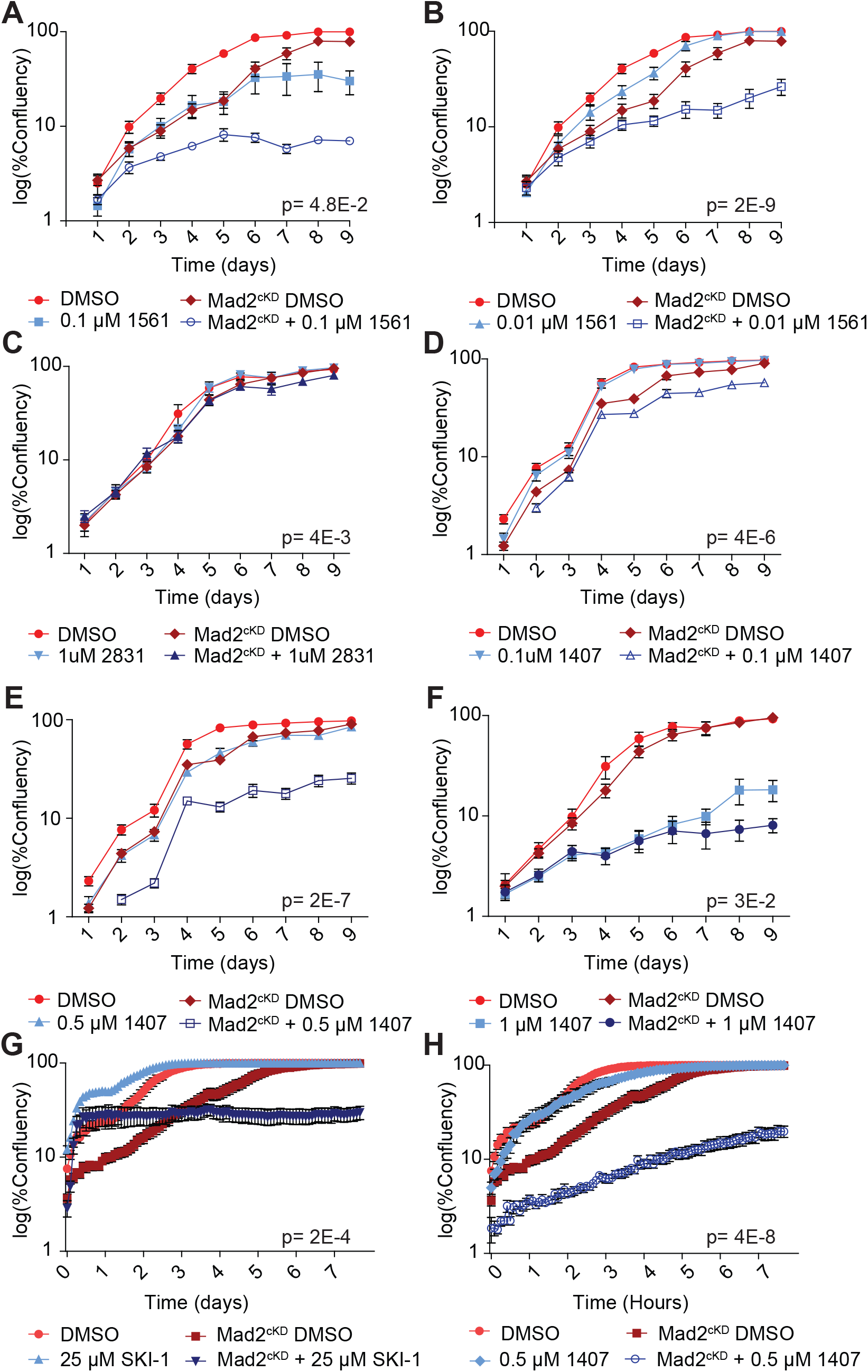
Validating candidate compounds that selectively target CIN cells. (**A-F**) Growth curves of control and Mad2^cKD^ RPE1 cells treated with 0.1 μM (**A**), and 0.01 μM (**B**) compound #1561, (**C**) 1 μM compound #2831, (**D-F**) 0.1 μM, 0.5 μM and 1 μM 1407, respectively. Data obtained by sequential daily microscope images, analyzed by FIJI-Phantast. Each point is a minimum of 3 biological replicates, each of which contains 3 technical replicates. Plotted is log scaled percentage confluency (cell coverage) over time. Error bars indicate SEM. P-values are calculated from paired, one sided t-tests of AUC corrected for cell line control. RPE1 DMSO and Mad2^cKD^ DMSO curves shared between A, B, and between C, F and **Sup. Fig. 3C**, and between D and E. (**G-H**) Incucyte growth curves of control and Mad2^cKD^ RPE1 cells treated with Src inhibitors SKI-1 (**G**) or compound #1407 (**H**, SKI606) for day 8-16. All points include data for six technical replicates. Error bars refer to SEM, p-values calculated from two-sided t-test of the AUC corrected for cell line controls. Data for DMSO control curves are shared between G, H and Fig. 5G.

SKI606 was designed as a tyrosine kinase inhibitor targeting Bcr-Abl (Golas *et al*, 2003) and Src (Boschelli *et al*, 2001). However, RPE1 cells do not have the Bcr-Abl fusion, making Src kinase the likely target. To test whether the observed effect of SKI606 on proliferation indeed acts through Src, we next compared the effect of SKI606 to the effect of another Src inhibitor, SKI-1. We found that SKI-1 displayed a similar synergy with CIN (**Fig. 4G**, **Sup. Fig. 3L**; 15-40% more effect than additive; Bliss independence test, p-values 1.5E-3 and 1.1E-3 for first 4 and last 4 days, respectively) as SKI606 (**Fig. 4H**, **Sup. Fig. 3M**) in inhibiting proliferation of Mad2^cKD^ RPE1 cells while having minimal effect on the proliferation of RPE1 control cells. We therefore conclude that Src inhibition is selectively toxic to cells with an impaired spindle assembly checkpoint.

### The synergy between Mad2 and Src inhibition does not involve impaired DNA damage signaling

Src is an oncogene, a key regulator of cell survival and mitosis (Thomas & Brugge, 1997), an activator of DNA-PK (Dittmann *et al*, 2008), and a regulator of actin organization (Destaing *et al*, 2008) and spindle orientation (Nakayama *et al*, 2012). We therefore next asked what the mechanism is between the observed synergy of Mad2 and Src inhibition in killing cells. As CIN leads to DNA damage (Janssen *et al*, 2011) and Src is involved in activating the DNA damage response via DNA-PK activation (Dittmann *et al*, 2008), we next tested whether DNA-PK inhibition would reproduce the results observed with Src inhibition. For this, we exposed cells to a DNA-PK inhibitor at a concentration that significantly increased γ-H2AX foci following gamma radiation, indicating impaired DNA repair (**Sup. Fig. 4A**). In this case, we found that DNA-PK inhibition was not synergistically toxic in dox-treated Mad2^cKD^ cells (**Sup. Fig. 4B**). In line with this, another DNA-PK inhibitor that was included in our screen (compound #1463; NU7441) did not show a differential effect between control and CIN RPE1 cells. Finally, we found that 4 Gray of irradiation and SKI606 both decreased proliferation of RPE1 cells as expected, but that SKI606 did not inhibit the growth of irradiated cells more than of controls, indicating that SKI606-invoked growth inhibition is independent of DNA damage (**Sup. Fig. 4C**). We therefore conclude that the observed synergy between Mad2 and Src inhibition is not caused by exacerbating DNA damage.

### SKI606 increases CIN in SAC deficient cells by deregulating microtubule polymerization rates

Since SKI606 does not appear to target aneuploidy-imposed stresses, nor DNA damage, we next investigated whether SKI606 affects chromosome missegregation rates. For this, we performed time-lapse imaging experiments with control and Mad2^cKD^ RPE1 cells expressing H2B-GFP, and quantified mitotic abnormalities in presence or absence of SKI606. Interestingly, we found that while SKI606 did not increase CIN in control cells, it did significantly increase CIN in Mad2^cKD^ cells (**Fig. 5A**), increasing the missegregation rates from 46% to 79%.

**Figure 5.**
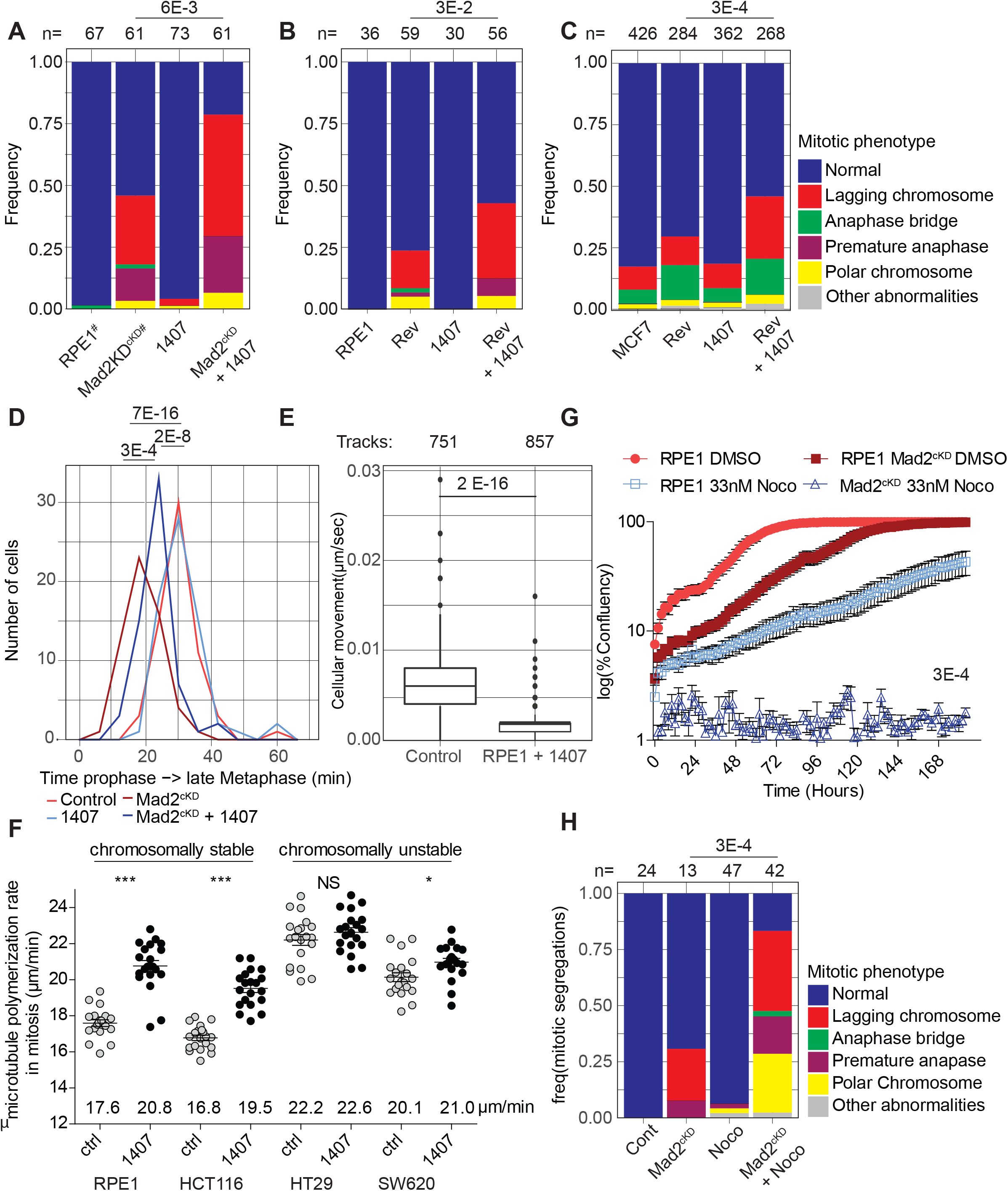
1407 significantly increases CIN in SAC-deficient cells by altering microtubule dynamics. (**A-C**) Frequency of mitotic abnormalities in control and Mad2^cKD^ RPE1 cells with and without 0.5 μM compound #1407 (**A**), RPE1 cells with 150 nM Reversine with and without 0.5 μM compound #1407 (**B**), and MCF7 cells treated with 15 nM Reversine and/or 0.5 μM compound #1407 (**C**). Data obtained by time-lapse microscopy imaging and includes at least three biological replicates. P-values are calculated from Chi-squared test. (**D**) Quantification of time from start prophase to late metaphase for control and Mad2^cKD^ RPE1 cells with and without 0.5 μM compound #1407. At least 29 mitoses were analyzed per condition from a minimum of 3 time-lapse microscopy experiments. (**E**) Boxplot showing mean cell migration speed (μm/second) of RPE1 cells with or without 0.5 μM 1407. Data include a minimum of 3 independent imaging experiments. P-values are calculated using a Wilcox test. (**F**) Microtubule plus end growth rate in mitosis with and without 0.5 μM compound #1407. Each dot represents the average of 20 microtubule movements within a cell, 20 cells per condition. (**G**) Incucyte-based growth curves of control and Mad2^cKD^ RPE1 in presence or absence of 33nM nocodazole at days 8-16. AUC is plotted relative to cell line controls, P-values are calculated using a Wilcoxon-Mann Whitney test. Data for DMSO control curves are also used in Fig. 4G & 4H. (**H**) Frequency of mitotic abnormalities in RPE1 cells with or without 0.5 μM compound #1407 and/or 33 nM nocodazole. Data obtained by time-lapse microscopy imaging and includes at least three biological replicates. P-values are calculated from Chi-squared test. “n” referrers to the number of mitotic events per condition. “#” refers to that the same data is also used in **Fig. 2E**.

To exclude the possibility that our observations were an artifact specific to Mad2^cKD^ cells, we also alleviated the SAC using the SAC inhibitor Reversine in RPE1 cells, and found that SKI606 indeed specifically increases chromosome missegregation rates in Reversine-treated cells (**Fig. 5B**). We also found that this phenotype persisted in other cell lines. For instance, SKI606 increased CIN rates of Reversine-treated MCF7 breast cancer cells from 30% to 46%, while SKI606 did not change CIN rates of MCF7 cells (17% to 19%) in the absence of Reversine (**Fig. 5C**). Together, these observations suggest that Src inhibition exacerbates a CIN phenotype specifically in cells with an impaired SAC.

To further investigate the mechanism underlying the effects of SKI606 on chromosome segregation, we determined whether Src inhibition had an effect on mitotic timing. For this, we compared mitotic length between control and Mad2^cKD^ RPE1 cells, with and without Src inhibition. While Mad2 alleviation decreased the time from prophase to metaphase as observed previously (Meraldi *et al*, 2004), mitotic length again increased when Mad2^cKD^ RPE1 cells were exposed to SKI606 (**Fig. 5D**). This suggests that the increased chromosome missegregation rates in SKI606-treated Mad2^cKD^ cells were not caused by further SAC inhibition and might be the result of altered microtubule dynamics. Mitotic timing of control RPE1 cells was unaffected by SKI606 treatment in line with the absence of a CIN phenotype in SKI606-treated control RPE1 cells. Furthermore, when analyzing the time-lapse data, we also noted that SKI606-treated cells (RPE1 (**Fig. 5E**) as well as MCF7 cells (**Sup. Fig. 5A**)) displayed reduced cell motility, also suggesting an effect of SKI606 on microtubule dynamics.

Given our results and a known role for Src in spindle orientation (Nakayama *et al*, 2012) and microtubule nucleation (Colello *et al*, 2010), we next investigated the effect of SKI606 on microtubule (MT) dynamics in a number of CIN and non-CIN (cancer) cell lines. For this, we quantified MT dynamics by time-lapse imaging in control- and SKI606-treated cells expressing EB3-GFP, which labels the plus-end tips of MTs and can therefore be used to quantify MT dynamics (Stepanova *et al*, 2003). Taking this approach, we found that SKI606 significantly increased MT polymerization rates in RPE1 as well as in diploid, non-CIN HCT116 cancer cells. Interestingly, we found that SKI606 increased the MT polymerization rates in these non-CIN cell lines to rates comparable to observed in the CIN cancer cell lines SW620 and HT29 (**Fig. 5F**). However, SKI606 treatment failed to further increase MT polymerization rates in HT29 cells, and only had a minor effect on MT polymerization rates in SW620 cells, suggesting that MT polymerization rates had reached their physiological maximum in these lines (**Fig. 5F**). Similar as observed for RPE1 and MCF7 cells, we found that SKI606 treatment did not increase chromosome missegregation rates in DMSO-treated HT29 cells (**Sup. Fig. 5B**). However, while Reversine treatment modestly increased CIN rates in HT29 cells as expected, combined SKI606 and Reversine treatment failed to increase CIN rates in HT29 cells further (**Sup. Fig. 5B**), providing additional proof that SKI606 acts through deregulating MT polymerization rates. Given these results, and as increased MT polymerization rates have previously been shown to drive CIN phenotypes (Ertych *et al*, 2014), we conclude that SKI606 contributes to a CIN phenotype by altering MT polymerization rates.

### Altering microtubule dynamics is synergistically toxic with SAC inhibition

To determine whether the synergy between altering MT dynamics and SAC inhibition was specific to SKI606 or would also apply to other MT poisons, we next tested the effect of SAC alleviation with low doses of nocodazole, which also increased MT polymerization rates (Ertych *et al*, 2014). For this, we first determined a non-toxic concentration for long-term (up to 8 days) treatment of nocodazole. While 250, 100, 50 and 25 ng/ml of nocodazole completely inhibited proliferation under these conditions, 10ng/ml (33nM) nocodazole was compatible with cell division. Indeed, while 33 nM nocodazole still reduced proliferation of RPE1 control cells, it was significantly more toxic to Mad2^cKD^ RPE1 cells, confirming the synthetic lethality between SAC inhibition and deregulating MT polymerization rates (**Fig. 5G**, **Sup. Fig. 5C**, 13% more than additive effect, p= 7.0E-3, Bliss independence test). Also in this setting, the observed synergy between low doses of nocodazole and SAC inhibition coincided with increased chromosome missegregation rates: while 33 nM nocodazole provoked mitotic abnormalities in only 6% of control RPE1 cells, 83% of nocodazole-exposed Mad2^cDK^ RPE1 suffered from defective mitoses, compared to 31% in the absence of nocodazole (**Fig. 5H**). Finally, when we combined SKI606 with SAC alleviation in the CIN cell line HT29, in which MT polymerization rates cannot further be increased (**Fig. 5F**, (Ertych *et al*, 2014)), we found that SKI606-imposed Src inhibition was no longer acting synergistically with SAC alleviation in killing cells (**Sup. Fig. 5D**), further indicating that altering MT dynamics is underlying the synergy observed between SKI606 and SAC inhibition. We conclude that altering MT polymerization rates synergizes with SAC inhibition in killing cells, thus providing new therapeutic opportunities for cancers in which either the SAC or MT dynamics are disturbed.

## Discussion

Chromosomal instability and the resulting aneuploidy are hallmark features of cancer cells. As both features discriminate cancer cells from healthy cells, they are promising therapeutic targets. In this study, we explored whether cells exhibiting CIN or stable aneuploidy displayed selective vulnerabilities to particular drugs. As CIN and aneuploidy trigger a number of responses in cells, including, but not limited to proteotoxic stress (Oromendia *et al*, 2012; Stingele *et al*, 2012), a deregulated cellular metabolism (Tang *et al*, 2011b; Williams *et al*, 2008), a DNA damage response (Zhang *et al*, 2015; MacKenzie *et al*, 2017), senescence (Andriani *et al*, 2016) and apoptosis (Ohashi *et al*, 2015), we selected a small library of FDA-approved drugs or compounds in clinical trials targeting aneuploidy/CIN-related responses.

### Aneuploid cells are sensitive to compounds that hyperactivate the cellular metabolism

When we screened for compounds that selectively kill aneuploid cells, we found that ZLN005, a transcriptional stimulator of PGC-1α, was significantly more toxic to double-trisomic RPE1 Ts12 Ts5 cells (Stingele *et al*, 2012) than control cells. PGC-1α is a master regulator of mitochondrial biogenesis and energy metabolism and its activation is thus expected to increase the cellular metabolism. None of the other tested compounds showed reproducible toxicity specific to aneuploid cells. While somewhat disappointing, it is important to note that we only tested a limited number of compounds (95 in total) in this screen and that large-scale future screens can still reveal new therapeutic vulnerabilities of aneuploid cells. Our findings in aneuploid cells correspond well with an earlier study by Tang *et al* (Tang *et al*, 2011b), who identified the energy stress-inducing drug AICAR as a compound that selectively targets aneuploid cells. Our findings form an independent confirmation of these findings and warrant further research on the molecular mechanism underlying this sensitivity.

Notably, ZLN005 did not emerge as a compound selectively targeting cells with an ongoing CIN phenotype (Mad2^cKD^ RPE1 cells; **Sup. Fig. 3K**) even though the CIN phenotype in these cells was shown to lead to substantial aneuploidy (**Fig. 2G**). Possibly, aneuploid cells need to adapt to the aneuploid state before becoming sensitive to drugs that exacerbate the cellular metabolisms (Mad2^cKD^ RPE1 cells were only exposed for up to 12 days to a CIN phenotype). Alternatively, as ongoing CIN and aneuploidy trigger (partially) different responses in cells (Bakker *et al*, 2018) they might also display differential vulnerabilities.

### Synthetic lethal interaction between inhibition of the SAC and Src activity

In addition to screening for compounds that selectively kill aneuploid cells, we also screened for compounds that selectively kill cells with a CIN phenotype. For the latter, we engineered RPE1 cells in which we could alleviate the SAC in an inducible fashion (Mad2^cKD^ RPE1 cells). This screen identified SKI606, a Src inhibitor as a compound that selectively kills cells with an alleviated SAC. We tested the phenotype with another Src inhibitor, which yielded the same phenotype. We find that Src inhibition increases the chromosome missegregation rate specifically in cells with an impaired mitotic checkpoint. Although, to our knowledge, Src has not been directly implicated with maintaining mitotic fidelity, Src has been shown to promote microtubule nucleation and regrowth (Colello *et al*, 2010) by binding to gamma-tubulin complexes (Kukharskyy *et al*, 2004). Additionally, Src was found to facilitate spindle orientation (Nakayama *et al*, 2012), and oncogenic v-Src has been associated with cytokinesis failure (Nakayama *et al*, 2017). This study revealed that Src inhibition results in increased MT polymerization rates, thus exacerbating the CIN phenotype imposed by Mad2 loss. Interestingly, while several Src inhibitors were found to affect tubulin polymerization, this was typically labeled as “dual mechanism of action” rather than a downstream effect of Src inhibition (Smolinski *et al*, 2018; Liu *et al*, 2013). Our study suggests that increased MT polymerization rates are a direct consequence of Src inhibition and that therefore these effects should be taken into consideration when treating patients with Src inhibitors.

### Mechanism underlying synthetic lethal interaction between SAC alleviation and Src inhibition

What can explain the synergy between SAC alleviation and Src inhibition in killing cells? Our data indicates that Src inhibition leads to increased MT polymerization rates. Importantly, we show that other MT destabilizing drugs display the same lethal interaction with SAC inhibition, further supporting our hypothesis that the synthetic lethal interaction between SAC and Src inhibition is explained by the role that Src has in regulating MT polymerization rates. But why are SAC-deficient cells specifically vulnerable to deregulated MT dynamics and why does Src inhibition not even impose a modest CIN phenotype upon control cells? When the SAC is operational, cells will arrest in metaphase until all chromosomes are properly aligned and attached. Therefore, if MT polymerization rates are increased, thus leading to decreased kinetochore-MT stability, the SAC will still delay mitosis until all chromosomes are properly attached. However, when the SAC is alleviated, the SAC can no longer compensate for deregulated MT activity, thus increasing the frequency of chromosome missegregation events in the presence of Src inhibitors (**Fig. 6**).

**Figure 6.**
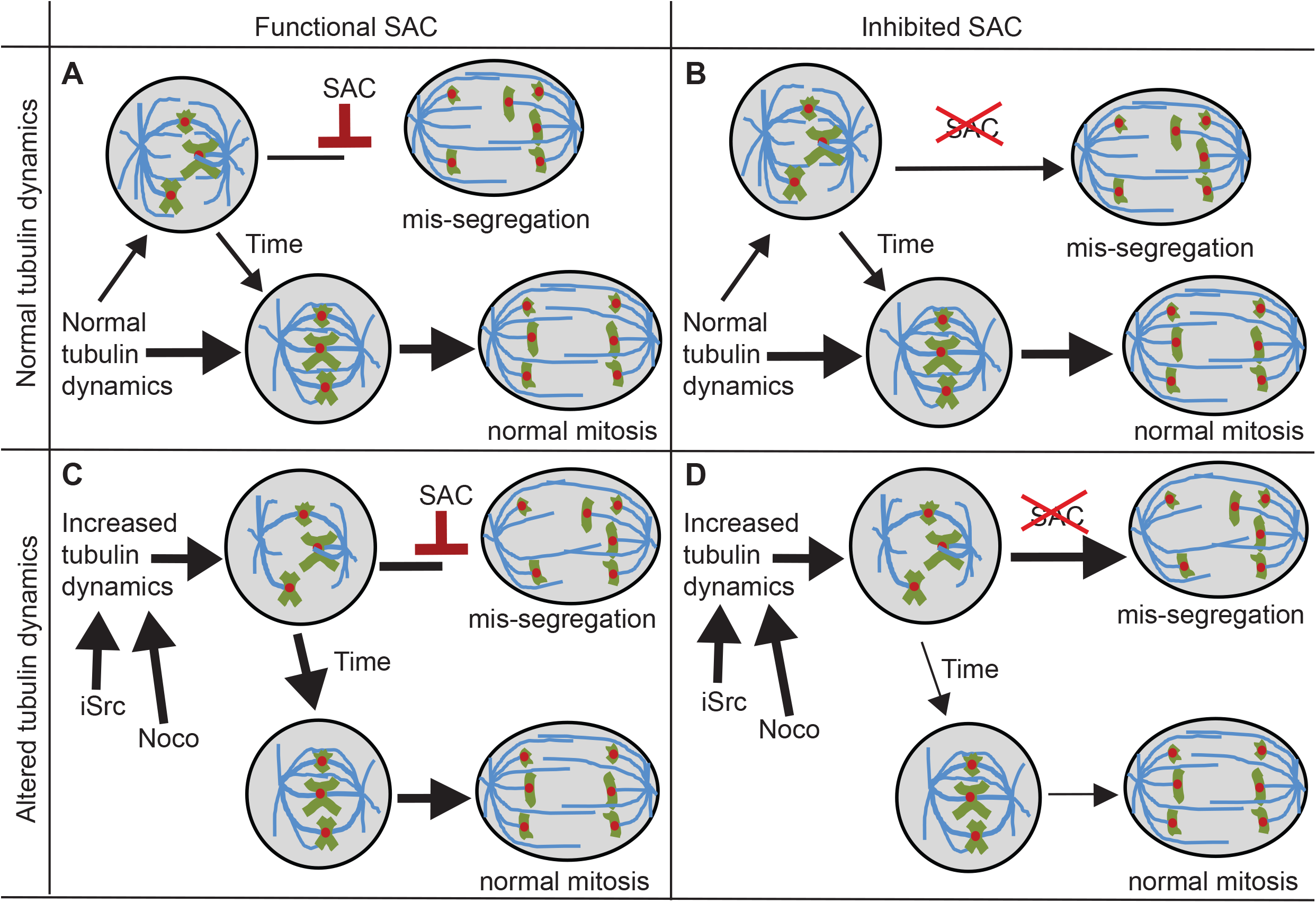
Proposed mechanism of how increased microtubule dynamics and SAC inhibition lead to synergistic toxicity by exacerbating the CIN phenotype. (**A**) Cells with normal tubulin dynamics and a functional SAC have very low chromosome mis-segregation rates. (**B**) Cells with normal tubulin dynamics but with an alleviated SAC display intermediate chromosome mis-segregation. (**C**) Cells with high tubulin dynamics, but a functioning SAC correct unattached kinetochores before entering anaphase. (**D**) Cells with increased microtubule dynamics and an alleviated SAC suffer from increased numbers of unaligned chromosomes that are not signaled by the SAC leading to increased rates of chromosome mis-segregation.

### Implications for cancer therapy

Chromosomal instability and aneuploidy are hallmarks of cancer cells, affecting ~70% of all solid cancers (Duijf *et al*, 2013). Therefore, therapies that exploit this feature might have a broad applicability. Our study suggests that exacerbating CIN in cells with a pre-existing CIN phenotype is a powerful strategy to selectively kill CIN cells. Indeed, that increasing CIN is a powerful method to target genome instable cancers has been reported by others as well (Silk *et al*, 2013; Zasadil *et al*, 2016; Thompson *et al*, 2010; Kops *et al*, 2004). One possible explanation for this is that cancer cells tolerate low levels of CIN, until CIN rates exceed a threshold, after which it becomes too toxic for cell survival. Our results indicate that altering MT dynamics to a level that does not affect mitotic fidelity can already act synergistically with spindle assembly checkpoint defects or inhibitors. This is particularly relevant for cancers in which either the SAC or MT polymerization rates are affected, as a defect in one process would render the cancer cells extremely sensitive to the interference with the other process, making the therapy much more specific to the cancers cells and thus reducing side effects and long-term toxicity.

Alternatively, the synthetic lethal interaction can be exploited to kill dividing cancer cells in combination therapy using both drugs at much lower concentrations than when using the drugs as individual agents. Indeed, others have also reported that SAC inhibitors act synergistically with taxanes in killing cells in tissue culture (Janssen *et al*, 2009; Maia *et al*, 2018; Tannous *et al*, 2013; Jemaà *et al*, 2013; Bargiela-Iparraguirre *et al*, 2014) and in vivo in mouse studies (Wengner *et al*, 2016; Maia *et al*, 2015, 2018; Tannous *et al*, 2013; Jemaà *et al*, 2013). The observed synergy was shown to result from increased CIN (Thompson & Compton, 2008; Janssen *et al*, 2009). In fact, three clinical trials combining Mps1 inhibitors with Paclitaxel to target human cancers are currently ongoing (Boston-Pharmaceuticals, 2017; Servier, 2018; Bayer, 2015). In this study, we show that this synergy is not limited to Mps1 and taxanes, and that alternative approaches to alter MT dynamics (such as MT-destabilizing drugs like Vincristine or Src inhibitors) in combination SAC inhibition can be used to synergistically target CIN cells by significantly increasing CIN. However, before our findings can be taken into the clinic, further validation experiments are required, which should reveal whether Src inhibitors are as effective as other MT polymerization deregulating drugs and whether such drugs indeed act synergistically with SAC inhibitors in killing cancer cells in vivo.

## Material and Methods

### Cell culture and compounds

RPE1 and MCF7 (ATCC) cells were grown in DMEM supplemented with 10% FBS and 100 Units/ml Penicillin and 100μg/ml Streptomycin. HT29 cells were growth in McCoy media supplemented with 10% FBS and Pen/Strep as above. Aneuploid cells RPE1 with trisomy 12 and trisomy 5 (Ts12 Ts5) and RPE1 hTert cells were kindly provided by the Storchova lab (Stingele *et al*, 2012). Most drugs used in this study were synthesized by Syncom B.V. (Groningen, NL), except for AZD8055 (Sigma), EPZ015666 (Sigma), and SKI-1 (Abcam). All drugs were dissolved in DMSO (Sigma) and diluted in tissue culture medium as indicated. Used drug concentrations were titrated before the actual screen, starting from an initial drug concentration of 10 μM. If the initial drug concentration of 10 μM was (near-)toxic to wildtype RPE1 cells, cells were next exposed to 1 μM, 0.1 μM, 10 nM or 1 nM of the compound, until a concentration was found that was no longer toxic; see Sup. Data 3 for initial drug titration growth curves.

### Generation Mad2^cKD^ cells

Mad2^cKD^ RPE1 cells were generated by transducing RPE1 cells with a lentiviral construct targeting human Mad2l1 (5’ - GGAAAGAATCAAGGAGG - 3’) in a pTRIPZ backbone (Open Biosystems, Catalog # RHS4696-200677332). Cells were selected in 2 μg/ml puromycine for 48 hours and single cell clones picked. Knockdown efficiency was determined for several clones by Western blot and the clone with the largest Mad2 reduction was used for further experiments.

### Western blot

Cells were harvested during logarithmic growth phase and lysed in EBL buffer (150 mM NaCl, 50 mM Hepes pH 7.5, 5 mM EDTA, 0.1% NP-40). Protein quantification was done using Quickstart Bradford assay, the Quickstart BSA standard kit (BioRad Inc.) or the MultiScan Go (Thermo Scientific). Antibodies used were Actin (#4970, Cell Signaling), Tubulin (#ab7291, Abcam), and Mad2 (#610678, BD Bioscience). Secondary antibodies were anti-Mouse IRDye 680RD (ab216778, Abcam) and anti-Rabbit IRDye 800CW (ab216773, Abcam). Blots were visualized and quantified using on the Odyssey imaging system (Li-Cor).

### Single cell sequencing

For single cell sequencing, cells were harvested, nuclei isolated and stained with Hoechst using nuclear isolation buffer. Single nuclei were then sorted into 96-well plates using a FACSJazz sorter (BD Bioscience Ltd.). Single-cell DNA libraries were prepared and sequenced (NextSeq 500, Illumina) with approximately 1% genomic DNA coverage as described previously (van den Bos *et al*, 2016). Sequence BAM files were analyzed with AneuFinder version 1.10.2 using the eDivisive analysis model at 1MB as described elsewhere (Bakker *et al*, 2016). Sequence BAM files and R script used for analysis are available upon request. Single cell sequencing data has been deposited at the European Nucleotide Archive (ENA) under accession number PRJEB33217.

### Metaphase spreads

Cells were cultured with 100 ng/ml Colcemid for 3 hours, harvested, incubated in 75 mM KCl for 15 minutes and fixated in 3:1 Methanol: acetic acid. Fixated cells were dropped on glass slides and nuclei visualized using DAPI staining. Metaphase figures were inspected on an Olympus BX43 microscope using a 63x lens. A minimum of 50 karyotyping spreads were counted per condition.

### Incucyte growth curves

For aneuploid drug screens, 200 RPE1 cells were sorted into each well of 96-well plates by flow cytometry (FACSJazz, BD Bioscience). For the CIN screen and follow up screens, 1600 cells were seeded per well in a 96 well plate. For the latter, RPE1 Mad2^cKD^ cells were treated with 1 μg/ml doxycycline for a minimum of 3 days before the start of any screen and sorted into wells with 1μg/ml doxycycline. Each well contained media with drugs at the concentrations listed, and all measurements were performed with technical triplicates for each plate. Cell growth was monitored every 2 hours using an IncuCyte Zoom (Essen BioScience Ltd.). Drug-containing media were refreshed every 4 days, and for the CIN screen, cells were passaged 1:8 on day 4. Cell density was quantified using IncuCyte ZOOM 2018A software. Cell confluency of control and CIN cells with drugs were normalized to DMSO-treated cells (RPE1 + DMSO and Mad2^cKD^ RPE cells + DMSO, respectively) and calculated using two different approaches: area under the curve (AUC) and Slope Analysis. As for the screens multiple drugs were tested per plate, the same DMSO control was used (at least one per plate) to compare the effects of the drug-treated cells. The figure legends indicate in which panels the same DMSO control was used.

### Area under the curve (AUC)

The AUC was estimated by taking the sum of the confluences per time-point. These values were then set relative to the cell line control area under the curve (each AUC value was divided by the mean cell line control AUC), RPE1 control, RPE1 double trisomy (Ts12 Ts5) and RPE1 Mad2^cKD^ cells were analyzed as different cell lines. These relative AUC values were then compared between cell lines per drug using a two-sided t-test. P-values in the screen were corrected for multiple testing using Bonferroni correction (Haynes, 2013).

### Slope analysis

An existing R script to analyze Incucyte data (Chapman *et al*, 2016) was modified to find the cutoff point for logarithmic growth. The logarithmic growth cutoff was determined for each drug and cell line combination, and the confluency at the cutoff was taken and divided by the average confluency of the cell line control. Since the slope is defined as the height (confluency) divided by width (time at cutoff), and the cutoff time-point was set to be the same for all DMSO and drug pairs, the cutoff time-points cancel out when setting slope relative to the cell line control. The resulting relative slope values were compared between cell lines per drug type using a t-test. The modified IncuCyteDRC R package for screen slope analysis is available upon request.

### Bliss Independence test for synergistic toxicity between drugs

To validate that the observed effects in the comound screens were synergistic and not only additive, we made use of the Bliss Independence test. For this, we calculated the fractional growth inhibition, defined as 1-(AUC^drug-treated cells^/AUC^control cells^). Next, we calculated the expected inhibitions, assuming drugs and CIN/aneuploid conditions were additive, using the Bliss Independence equation: Expected Inhibition = Fa + Fb − (Fa*Fb), where Fa is fractional growth inhibition of drug A, and Fb was the fractional growth inhibition of either CIN or aneuploid cell conditions. The expected inhibition was compared to the actual growth inhibition; the p-value was determined using a t-test, and the greater than additive toxicity was found by taking the difference between expected and actual fractional inhibition.

### Growth curves analyzed by Fiji Phantast

For validation experiments, growth curves were determined from daily microscope images using a Olympus IX51 microscope. For these experiments 5,300 cells were seeded per well of a 24-well plate. Each well was imaged once a day for a minimum of 7 days. The FIJI package PHANTAST (Jaccard *et al*, 2014) was used to estimate confluency per well per time-point. PHANTAST settings were epsilon=5, and sigma ranging between 0.01 and 0.03 depending on cell coverage and confluency accuracy. All measurements include at least 3 technical replicates and three biological replicates (i.e. a minimum of 9 measurements per time point). Growth was plotted using Prism software (Graphpad). As multiple drugs were tested per plate, the same DMSO control was used to compare the effects of the drug-treated cells. The figure legends indicate in which panels the same DMSO control was used. Growth kinetics were normalized identical to IncuCyte measurements.

### Area under the Curve

The AUC was calculated by taking the sum of the confluency at all time-points per condition. This was set relative to cell line growth by dividing each AUC value by the average AUC for the DMSO cell line control for each plate. These values were compared between cell lines per drug with a two-sided t-test.

### Slope analysis FIJI growth curves

Growth curves were plotted on a log scale and the logarithmic growth cutoff point was estimated manually for each condition. To calculate the slope, the negative log of the confluency at that time-point was divided by the cutoff day: –log(confluency_cutoff_)/T_cutoff_. This was divided by the average slope of the cell line control to compensate for cell line growth differences. The replicates of the relative slope values were compared between cell lines per condition.

### Live cell Imaging and CIN analysis

RPE1, Mad2^cKD^ RPE1, MCF7 and HT29 cells expressing H2B-GFP were treated as indicated and imaged on a DeltaVision microscope (Applied Precision Ltd.). Interphase phenotypes were analyzed by quantifying nuclear morphology. Mitotic abnormalities were manually quantified from overnight live cell imaging movies. Measurements include at least three biological replicates and numbers of cells quantified are indicated in the text. A chi-squared test was used to test whether differences between conditions were significant. Mitotic time was analyzed by calculating the time-points between the first sign of DNA condensation to the last point before anaphase, and from the first anaphase time-point to the time-point at complete DNA de-condensation from time-lapse imaging data.

### Cell motility assay

To quantify cell motility, time-lapse movies were analyzed using FIJI Trackmate (Tinevez *et al*, 2017). A minimum of fifteen overnight imaging movies were used per condition including at least three biological replicates. Trackmate input conditions were optimized for each cell type. RPE1 nuclear diameter was set at 20 μM, while MCF7 nuclear diameter was set at 15 μM. Track statistics per condition were combined and Track speed was plotted in R ggPlot2 (Wickham, 2016). Differences were calculated with two-sided t-tests.

### Microtubule movement analysis

Microtubule plus end assembly rates were determined by tracking EB3-GFP protein (vector kindly provided by Linda Wordeman, USA) in live cell microscopy experiments as in Ertych *et al.* (Ertych *et al*, 2014). Average assembly rates (μm/min) were calculated based on data retrieved for 20 individual microtubules per cell that were randomly selected. A total of 20 cells were analyzed from 3 independent experiments. Significance was assessed using a two-sided, unpaired t-test.

## Author contributions

KMS and FF conceived and designed the study. KMS performed most of the experiments. YCL and HB performed and supervised the MT dynamics measurements, MS and MCT assisted with data analyses and statistical tests. SFP and JES provided technical support. ZS provided the aneuploid RPE1 cells. HVDB and DCJS performed single cell sequencing experiments. KMS and FF wrote the manuscript with input from all authors. FF supervised the work and provided funding.

## Acknowledgements

We are grateful to the members of the Foijer and Bruggeman labs and to the members of the PloidyNet consortium. This work was supported by the European Union FP7 Marie Curie Innovative Training Network grant PloidyNet (607722) and a Dutch Cancer Society project grant (2015-RUG-7833) to Foijer.

---

**Supplementary data 1**. Growth curves of aneuploid RPE1 cells exposed to small molecule compounds. Raw data used to produce Fig. 1.

**Supplementary data 2**. Growth curves of Mad2^cKD^ RPE1 cells exposed to small molecule compounds. Raw data used to produce Fig. 2.

**Supplementary data 3**. Drug titration curves used to determine starting drug concentrations for aneuploidy and CIN screens as described in Material and Methods.

**Supplementary Table 1**. List of compounds used in the drug screen to identify compounds that selectively kill aneuploid cells.

**Supplementary Table 2**. List of compounds used in the drug screen to identify compounds that selectively kill cells with a CIN phenotype.

**Supplementary Figure 1.**
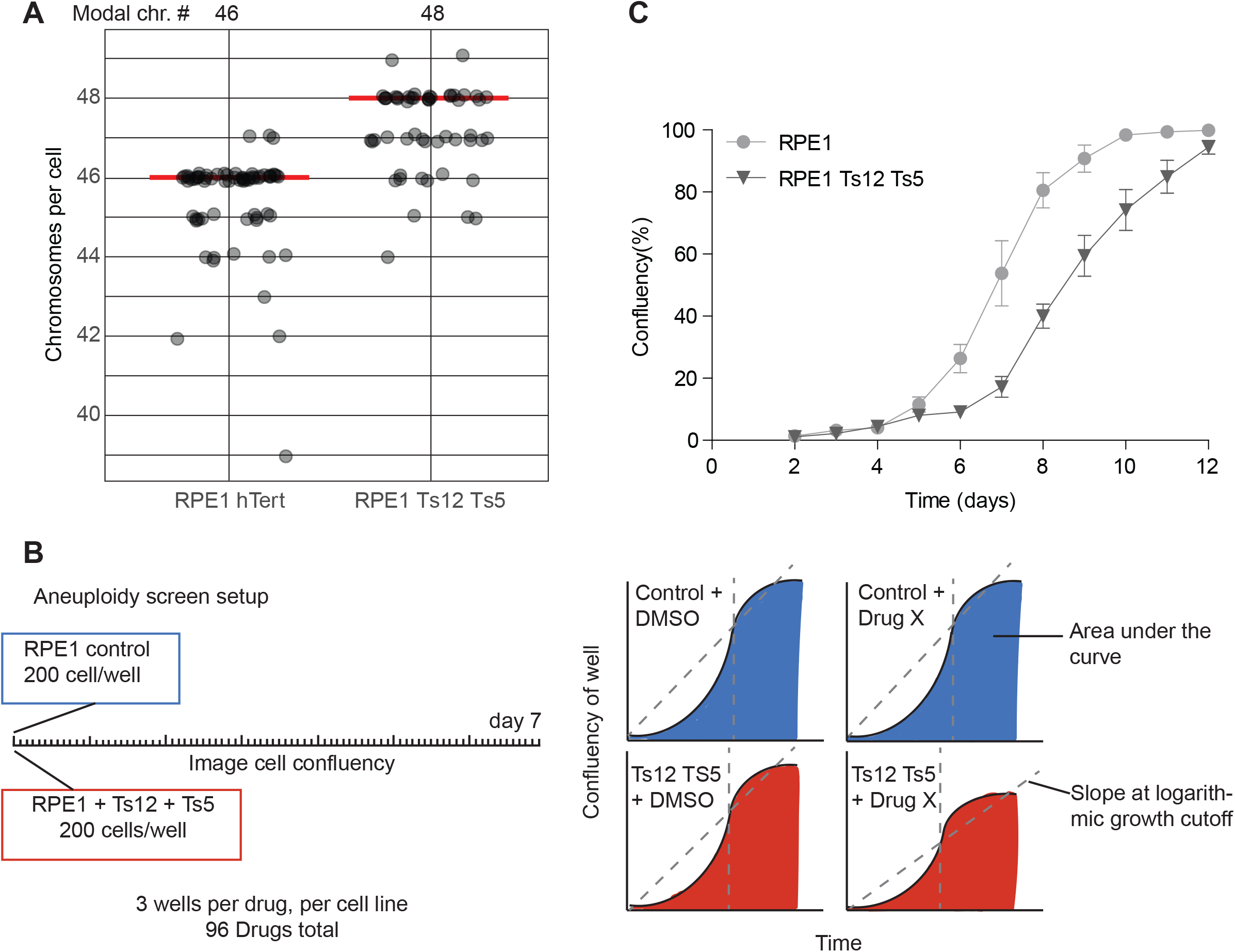
Aneuploid cell line validation and screen setup. (**A**) Chromosomes numbers assessed by metaphase spread in control RPE1 and RPE1 + Trisomy 12 + Trisomy 5 (Ts12 TS5) cells. A minimum of 50 metaphases were counted per condition. Modal chromosome counts are shown above, and labeled with a red line. (**B**) Aneuploid screen setup and data types used for analysis. (**C**) Growth curve of RPE1 control and RPE1 Ts12 Ts5 cells over time. Data obtained by sequential daily microscope images, analyzed by FIJI-Phantast. Data include at least three technical replicates each with three biological replicates. Error bars indicate SEM. Data for DMSO control curves are shared with Fig. 1E and **Sup. Fig. 3G**.

**Supplementary Figure 2.**
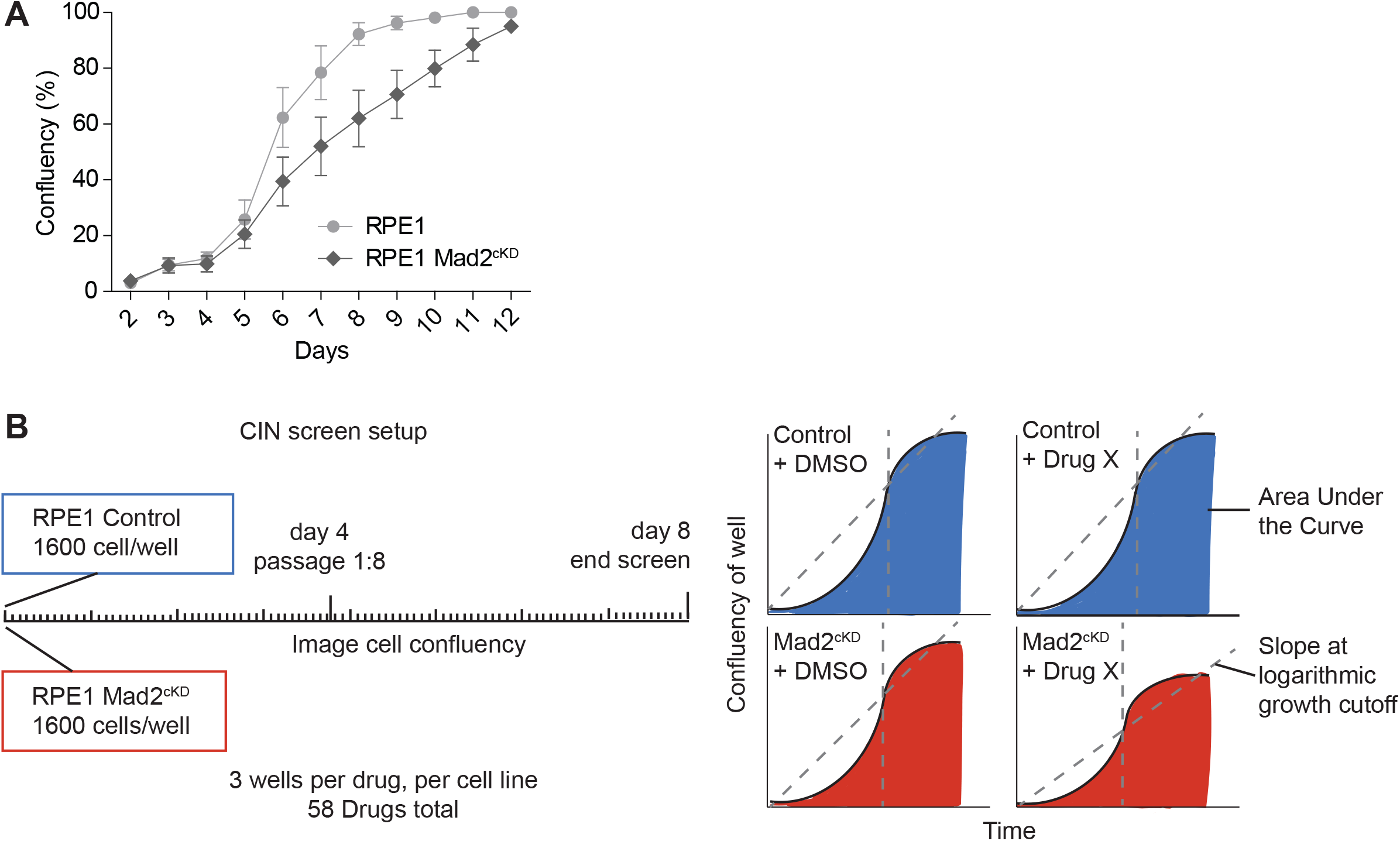
CIN screen setup. (**A**) Growth curve of control RPE1 and RPE1 Mad2^cKD^ cells over time. Data obtained by sequential daily microscope images, analyzed by FIJI-Phantast. Error bars indicate SEM of 3 biological replicates, each with 3 technical replicates. (**B**) CIN Screen setup and data types used for analysis.

**Supplementary Figure 3.**
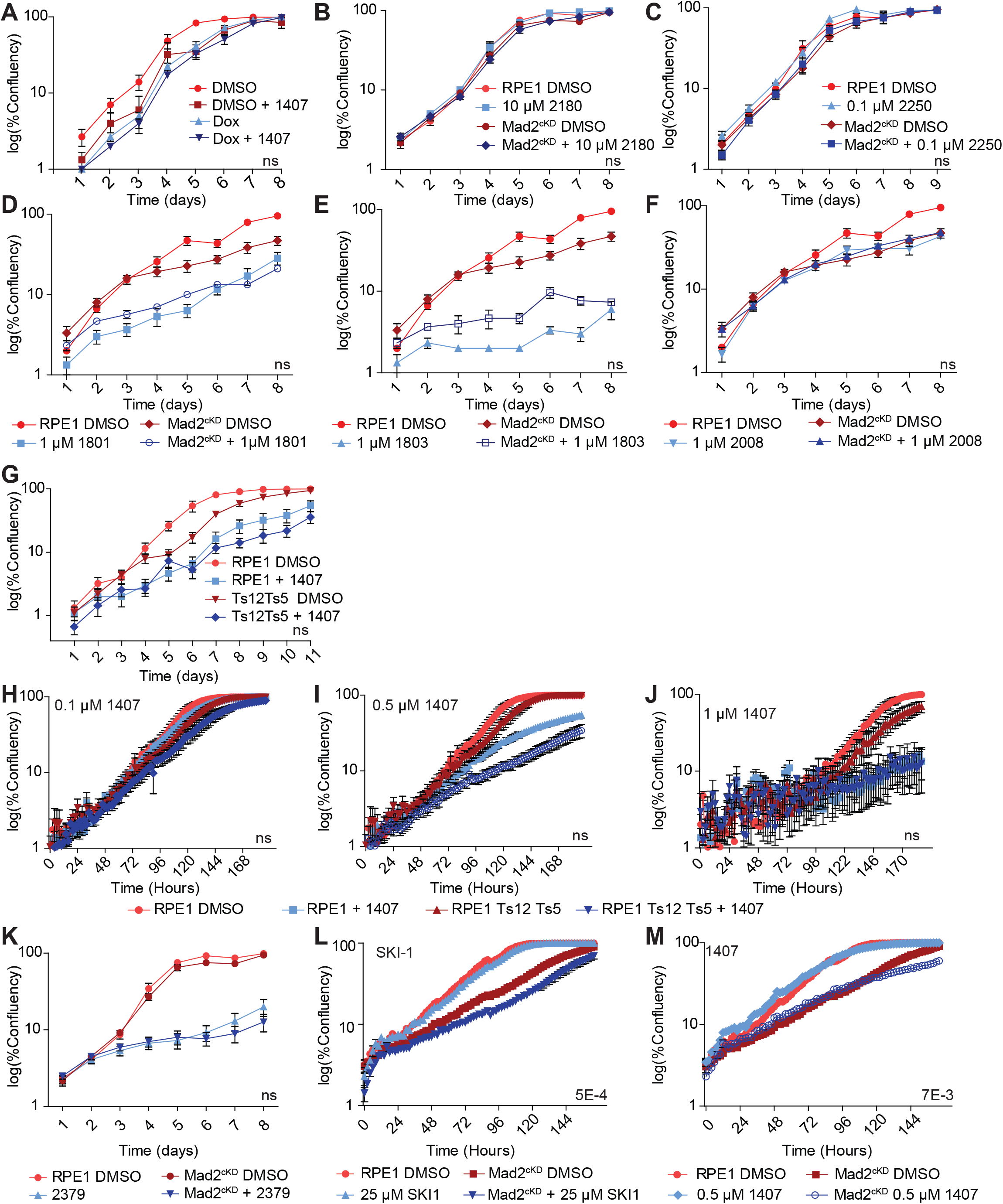
Validating CIN screen setup and hits. (**A**) Growth curve of doxycycline-treated RPE1 cells with or without 0.5 μM compound #1407. Data obtained by sequential daily microscope images, analyzed by FIJI-Phantast. Each data point includes 3 technical triplicates. P-values are calculated from two-sided t-test. (B-F) Growth curves of control and Mad2^cKD^ RPE1 cells treated with compounds #2180 (**B**), #2250 (**C**), #1801 (**D**), #1803 (**E**), and #2008 (**F**), respectively. Data obtained by sequential daily microscope images, analyzed by FIJI-Phantast. Each data point includes 3 technical triplicates. P-values are calculated from one-sided t-test. Data for DMSO control curves are shared between B & **Sup. Fig. 3K**, between D, E & F, and between C & Figure 4C & 4F. (**G**) Growth curves of RPE1 control and RPE1 Ts12 Ts5 cells with or without 0.5 μM compound #1407. Data obtained by sequential daily microscope images, analyzed by FIJI-Phantast. Data includes 3 biological replicates, each with 3 technical replicates. P-values are calculated from two-sided t-test. Data for DMSO control curves are shared with Fig. 1E & **Sup. Fig. 1C**. (H-J) Incucyte-derived growth curves of RPE1 and RPE1 (Ts12 Ts5) cells treated with 0.1 μM (**H**), 0.5 μM (**I**) and 0.1μM compound #1407 (**J**). Data represents 3 technical replicates per condition. P-values are calculated by two-sided t-test. (**K**) Growth curve of RPE1 control and RPE1 Mad2^cKD^ cells with or without 10 μM compound #2379. Data obtained by sequential daily microscope images, analyzed by FIJI-Phantast. Data includes three biological replicates, each with 3 technical replicates. P-values are calculated from two-sided t-test. Data for DMSO control curves are shared with Fig. 3B. (L-M) Incucyte-derived growth curves (days 1-4) of control and Mad2^cKD^ RPE1 cells treated with 25 μM SKI-1 (**L**) or 0.5uM compound #1407 (**M**) with 6 replicates each. P-values are calculated from two-sided t-test. Data for DMSO control curves are shared between L, M and **Sup. Fig. 5C**. Error bars indicate the SEM between measurements. T-tests are based on the AUC corrected for cell line controls.

**Supplementary Figure 4.**
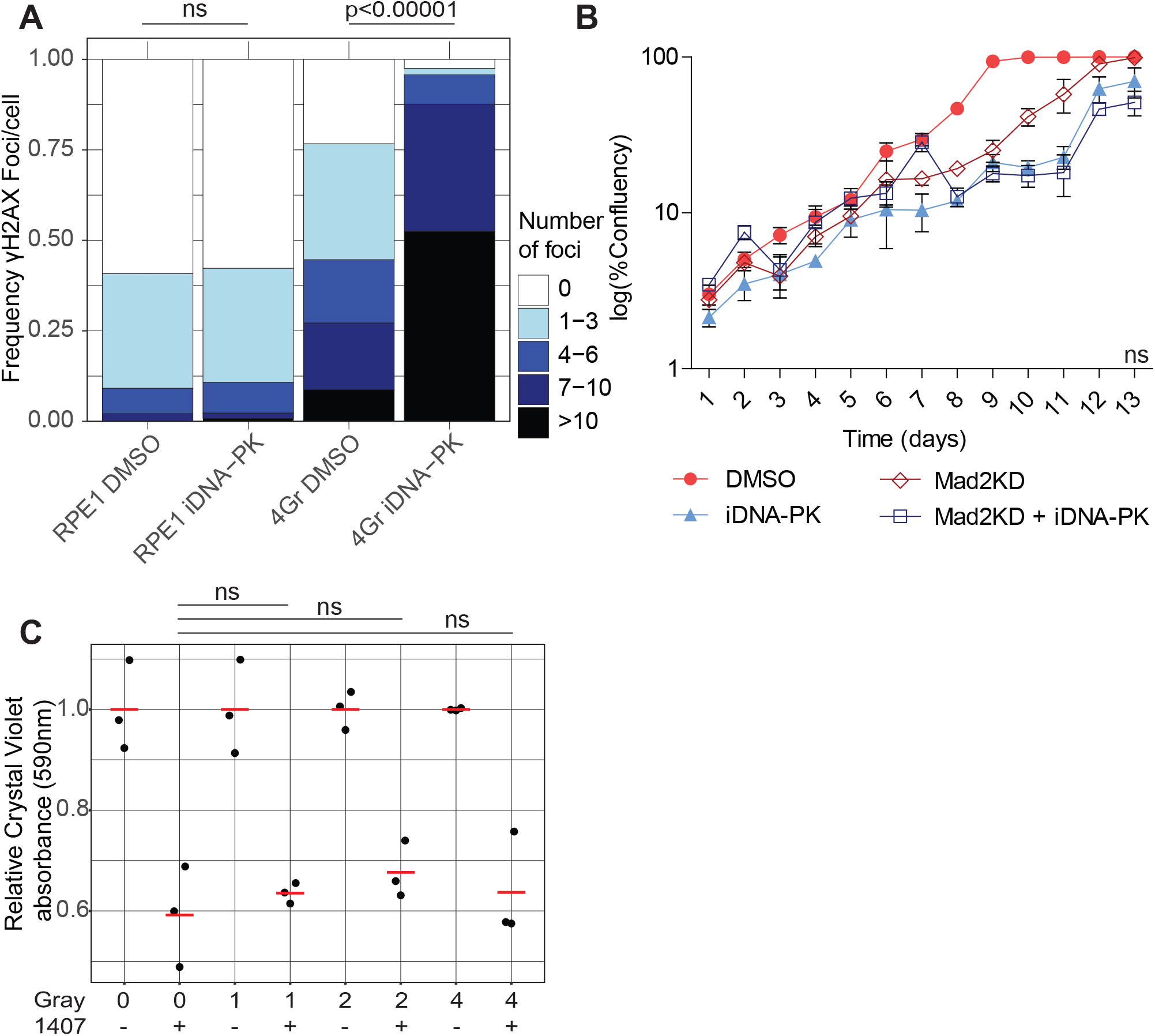
DNA damage is not synergistically toxic with Src inhibition in Mad2^cKD^ RPE1 cells. (**A**) DNA double strand breaks assessed by γH2AX labeling in control and irradiated (4 Gray, 15 hours post-irradiation) RPE1 cells with or without 250nM of the DNA-PK inhibitor KU0060648. Each category includes a minimum of 100 cells. P-values assessed by a Chi-squared test. (**B**) Growth curve of control and Mad2^cKD^ RPE1 cells with or without 250nM KU0060648. Data obtained by sequential daily microscope images, analyzed by FIJI-Phantast. Data includes three technical replicates. Error bars refer to the SEM. P-value is calculated for the AUC relative to the cell line control using a two-sided t-test. (**C**) Proliferation assessed by crystal violet staining of RPE1 cells 48 hours after 0, 1, 2 or 4 Gray irradiation with or without 0.5 μM 1407. All data was acquired in technical triplicates and corrected to radiation only control. P-value was assessed using a two-sided t-test.

**Supplementary Figure 5.**
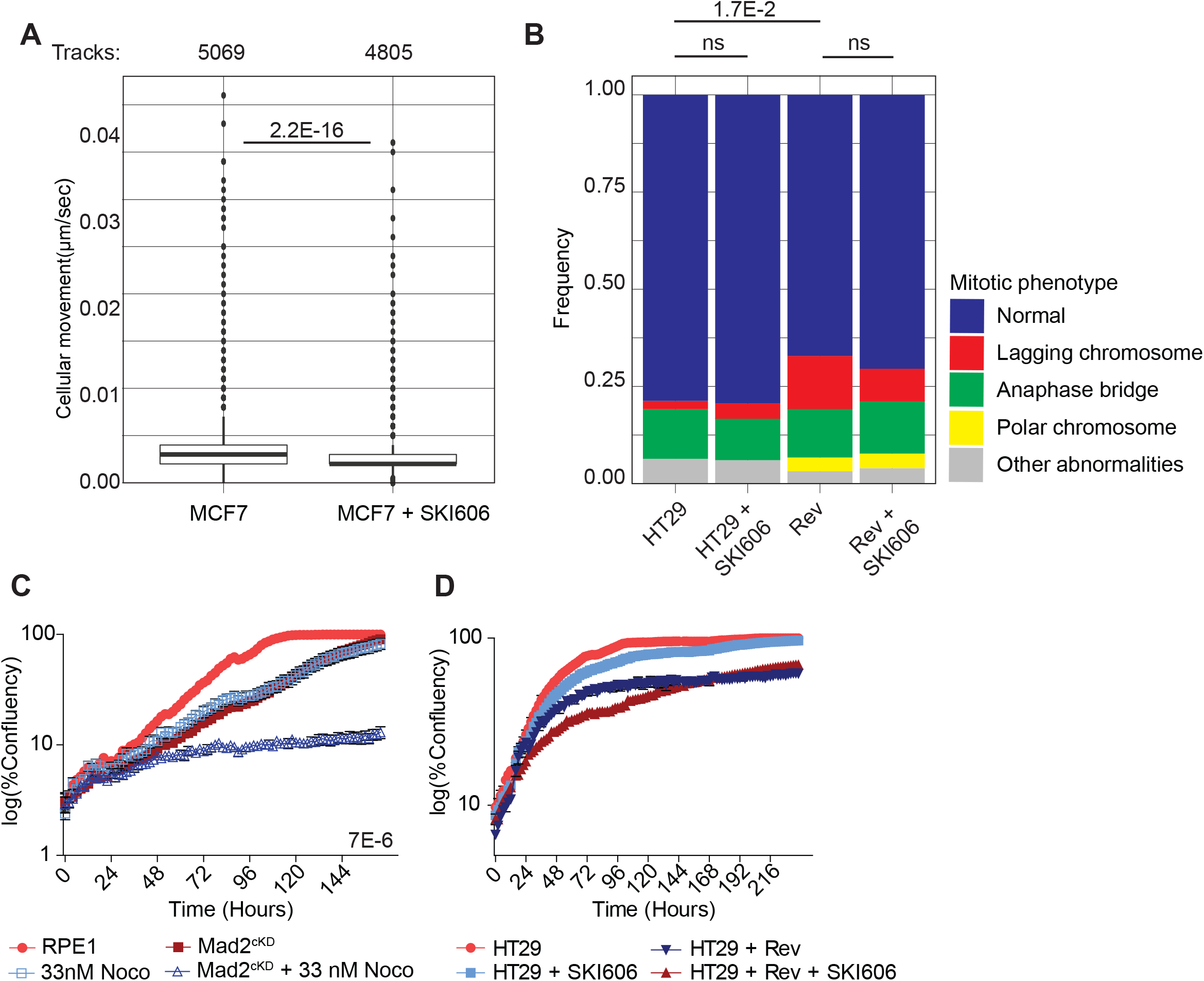
Src inhibition leads to increased microtubule polymerization rates. (**A**) Mean cell movement speed [μm/sec] of MCF7 cells with or without 0.5 μM of SKI606. Data include a minimum of 3 independent imaging experiments. P-values are calculated through a Wilcox Mann U test. (**B**) Frequency of mitotic abnormalities observed in HT29 cells incubated with or without 150nM Reversine and/or 0.5 μM SKI606. Data includes a minimum of 3 independent time-lapse imaging experiments, and a minimum of 90 mitotic events per condition. P-value was calculated from a Chi-squared test. (**C**) IncuCyte-derived growth curves (day 1-7) of control and Mad2^cKD^ RPE1 cells with or without 10ng/ml nocodazole. Data includes 6 replicates each. P-value was calculated from a two-sided t-test on the AUCs relative to the cell line’s control. Data for DMSO control curves are shared with **Sup. Fig. 3L** & **Sup. Fig. 3M**. (**D**) IncuCyte-derived growth curves of HT29 cells with or without 150nM Reversine and/or 0.5 μM SKI606. Data includes six replicates for each condition. P-value was calculated from a one-sided t-test of the AUC relative to the cell line’s control.

