## Supplementary material for "Altering microtubule dynamics is synergistically toxic with inhibition of the spindle checkpoint": Sup data 1

#### Supplementary data 1: Aneuploid screen growth curves

Aneuploid screen growth curves shown per cell line and per drug used. The top left corner lists the drug ID, the concentration and which cell line these growth curves belong to. RPE hTert are euploid control cells, and RPE1 Ts12 Ts5 are aneuploid double-trisomy cells.

One growth curve is displayed with the cell line DMSO control. If the drug was rescreened, a second cuve is also shown. All concentrations are in micromolar (uM).

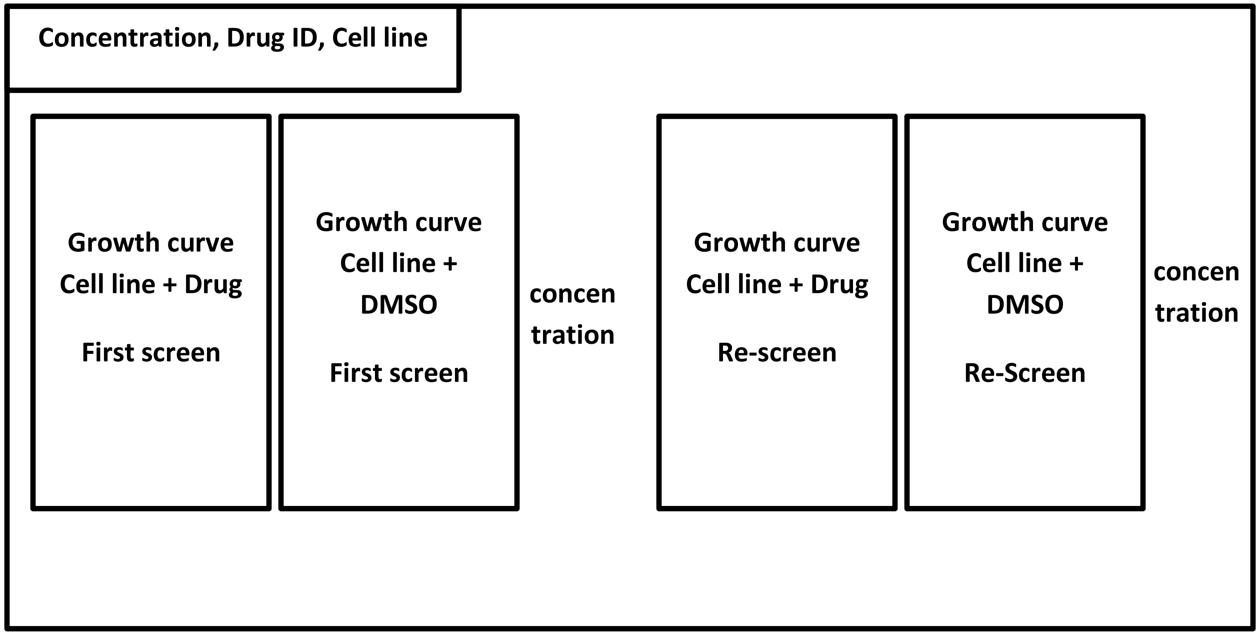

**10 uM 1122 RPE1 hTert**

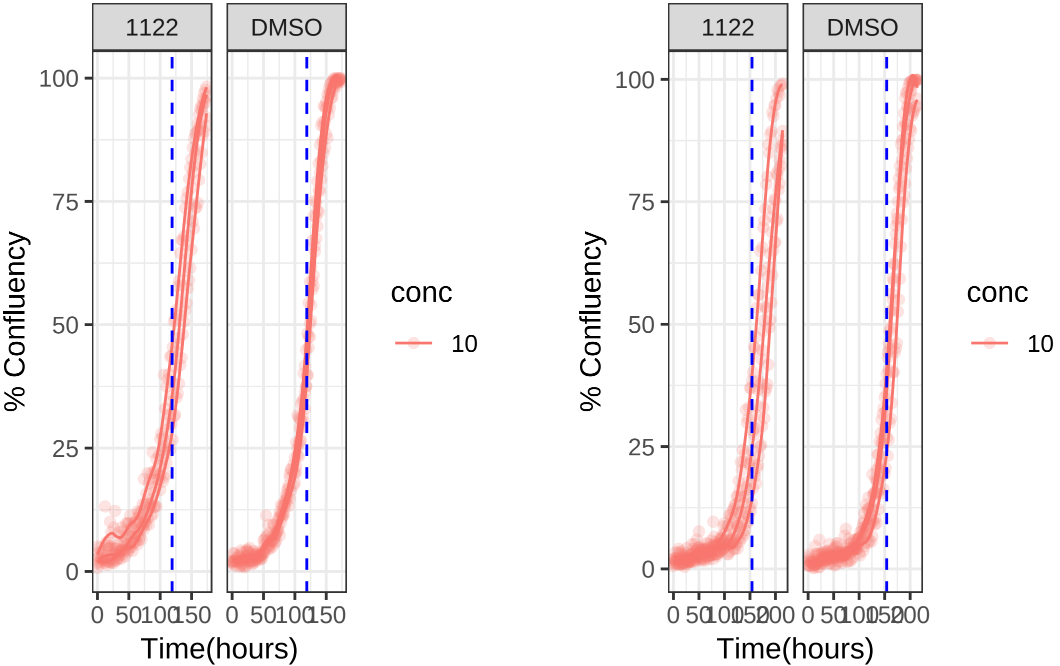

**10 uM 1122 RPE1 Ts12 Ts5**

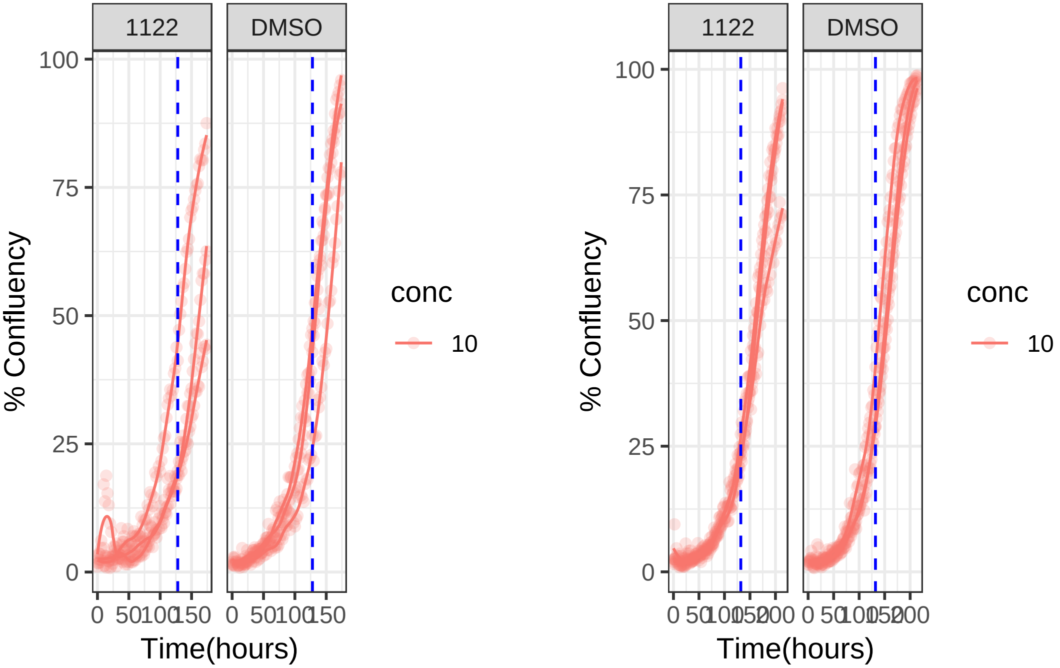

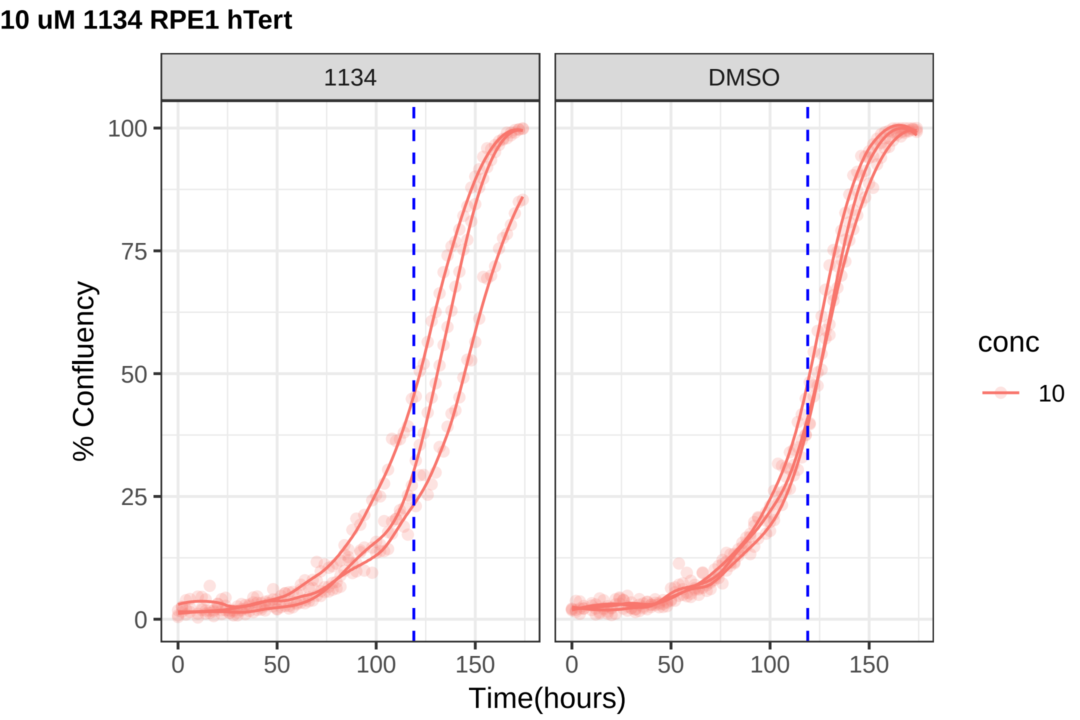

10 uM 1134 RPE1 hTert
1134
DMSO
% Confluency
0
25
50
75
100
0
50
100
150
Time(hours)
conc
10

**10 uM 1134 RPE1 Ts12 Ts5**

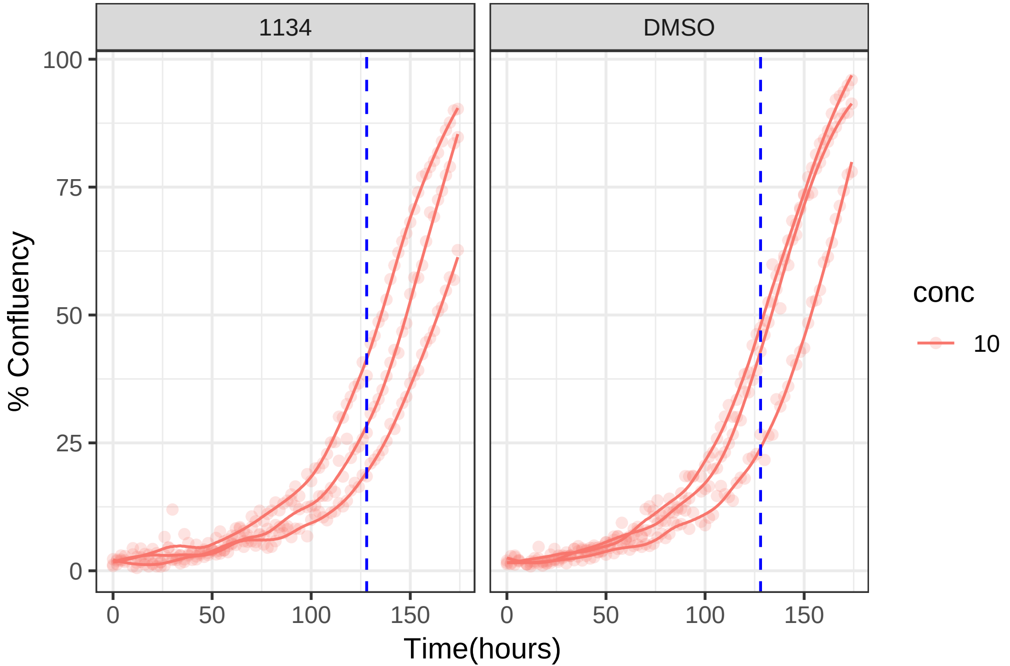

**10 uM 1223 RPE1 hTert**

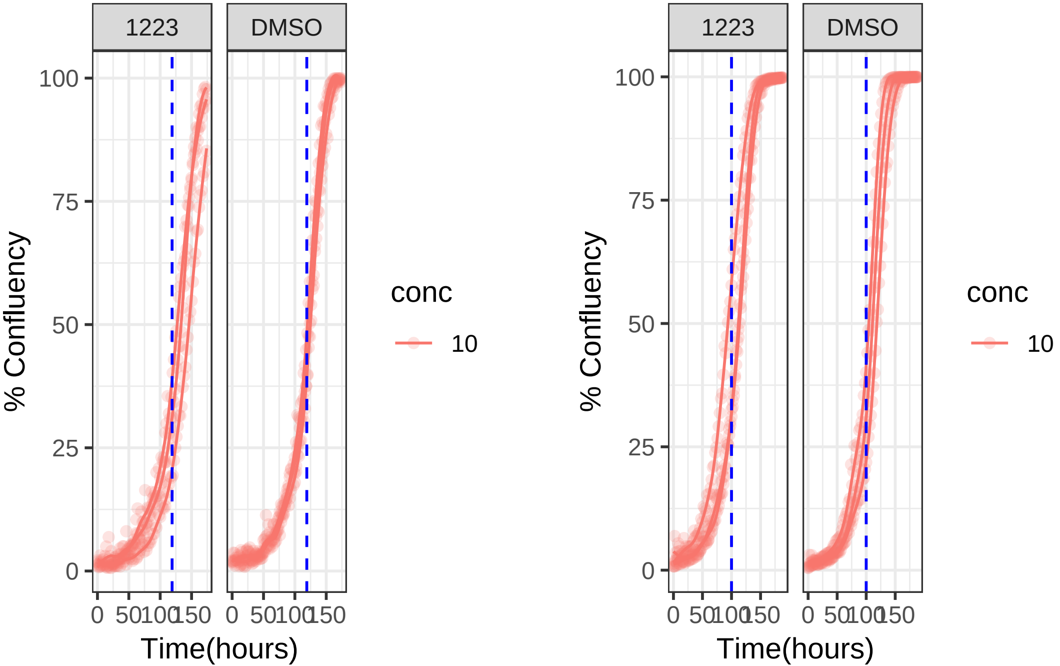

**10 uM 1223 RPE1 Ts12 Ts5**

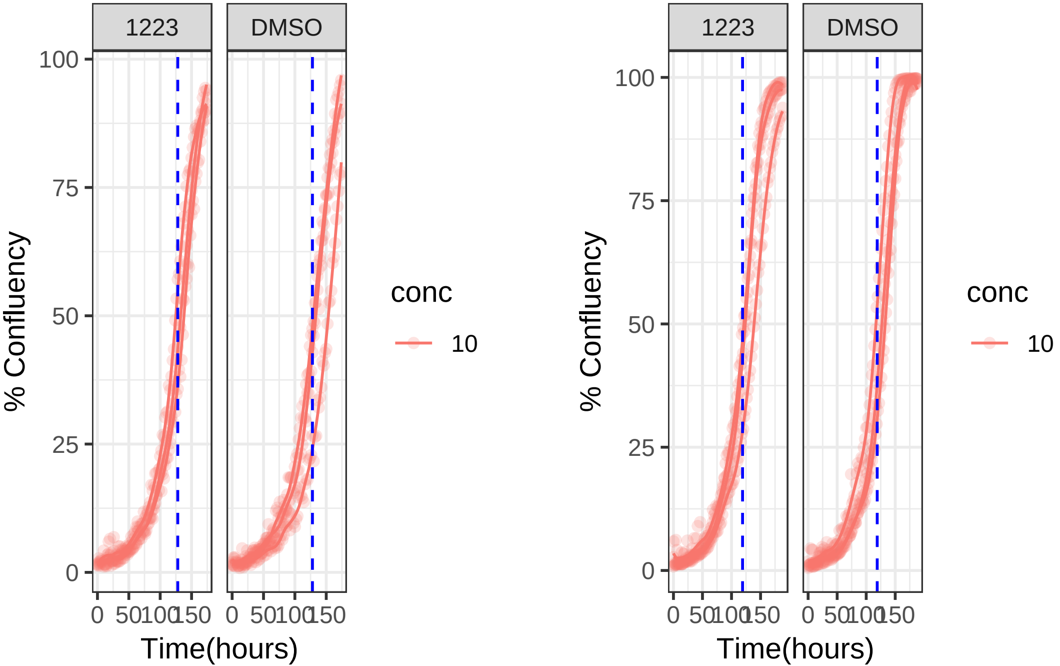

**0.001 uM 1233 RPE1 hTert**

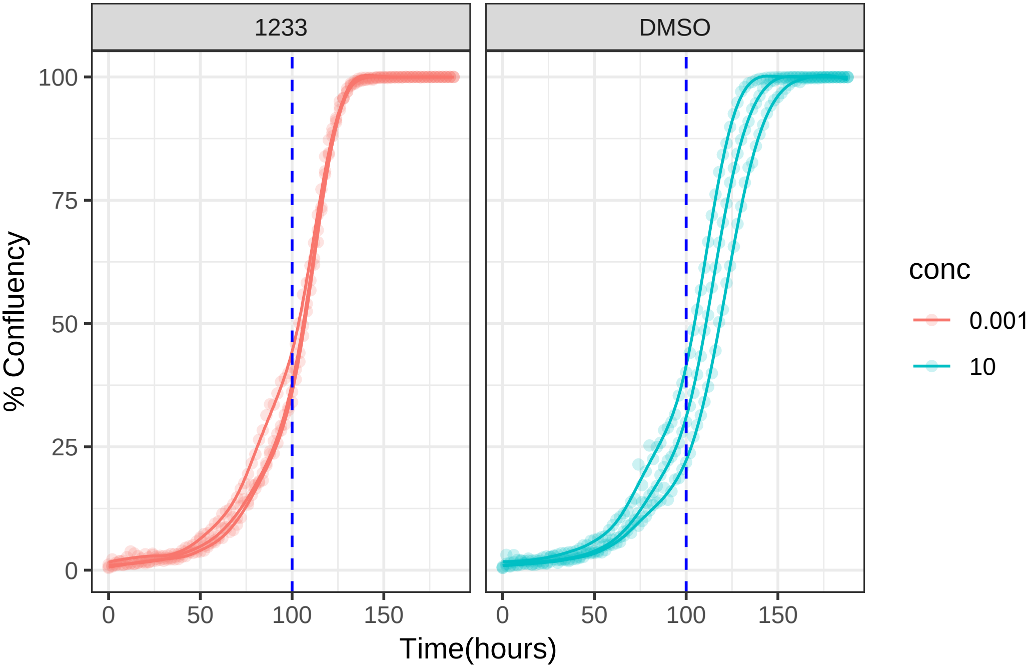

**0.001 uM 1233 RPE1 Ts12 Ts5**

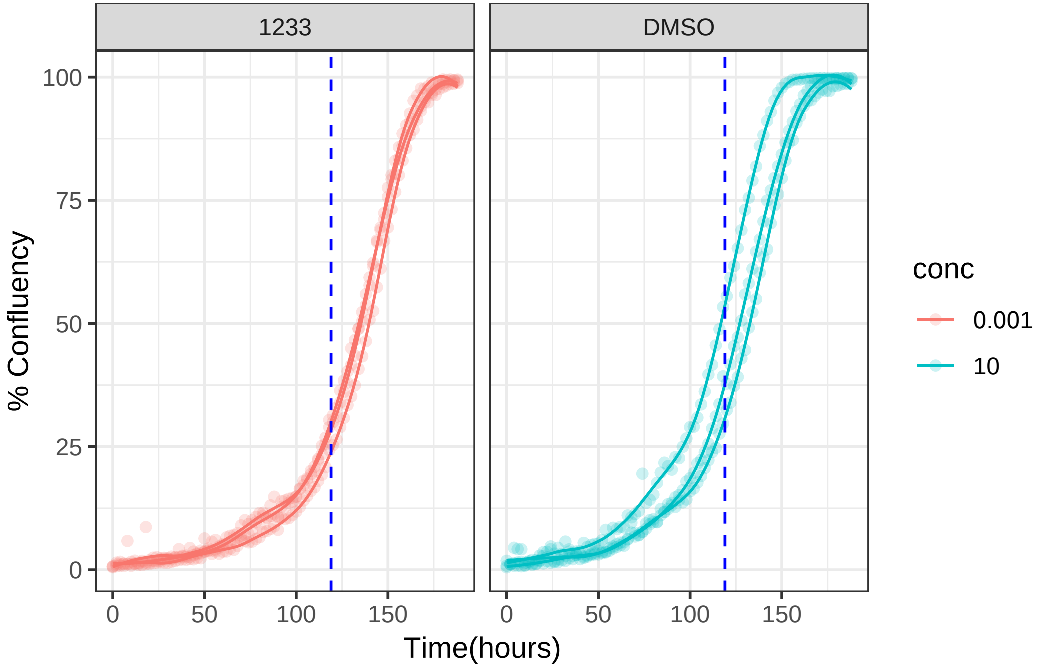

**10 uM 1237 RPE1 hTert**

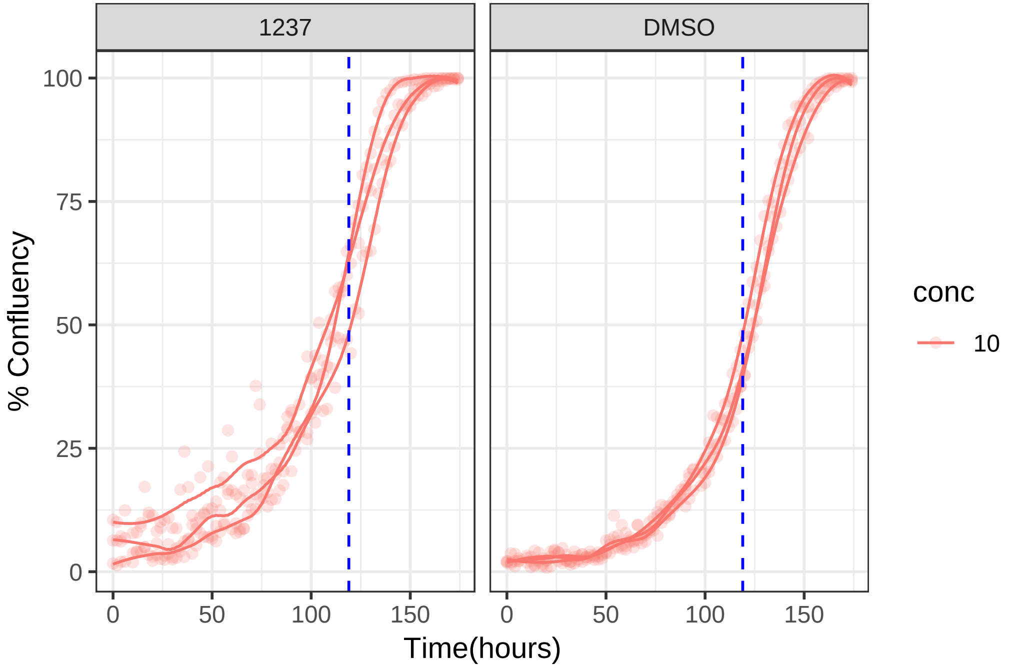

**10 uM 1237 RPE1 Ts12 Ts5**

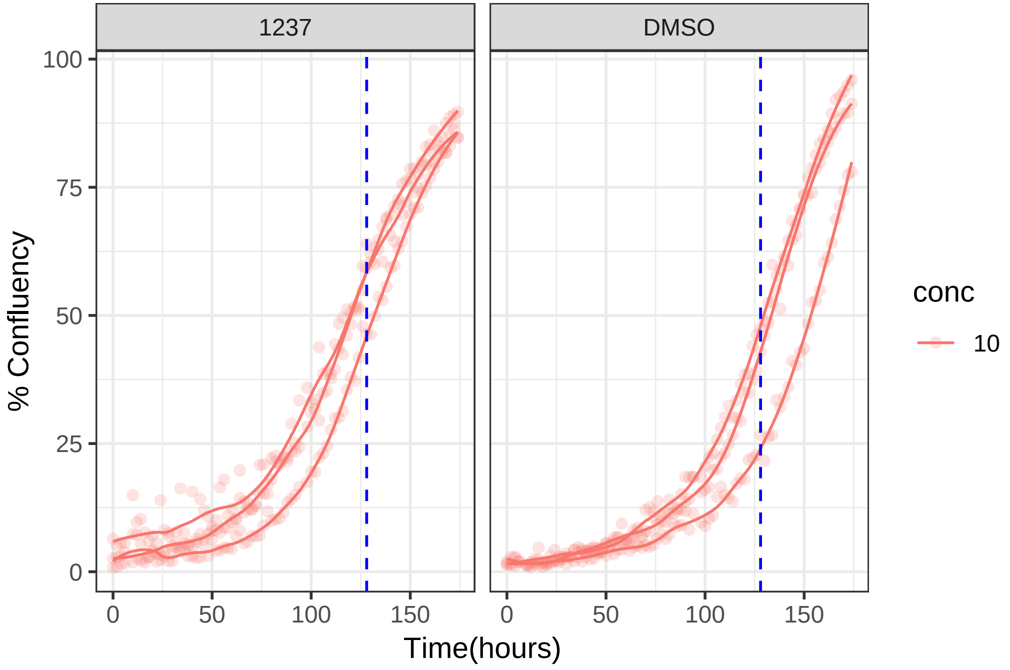

**10 uM 1239 RPE1 hTert**

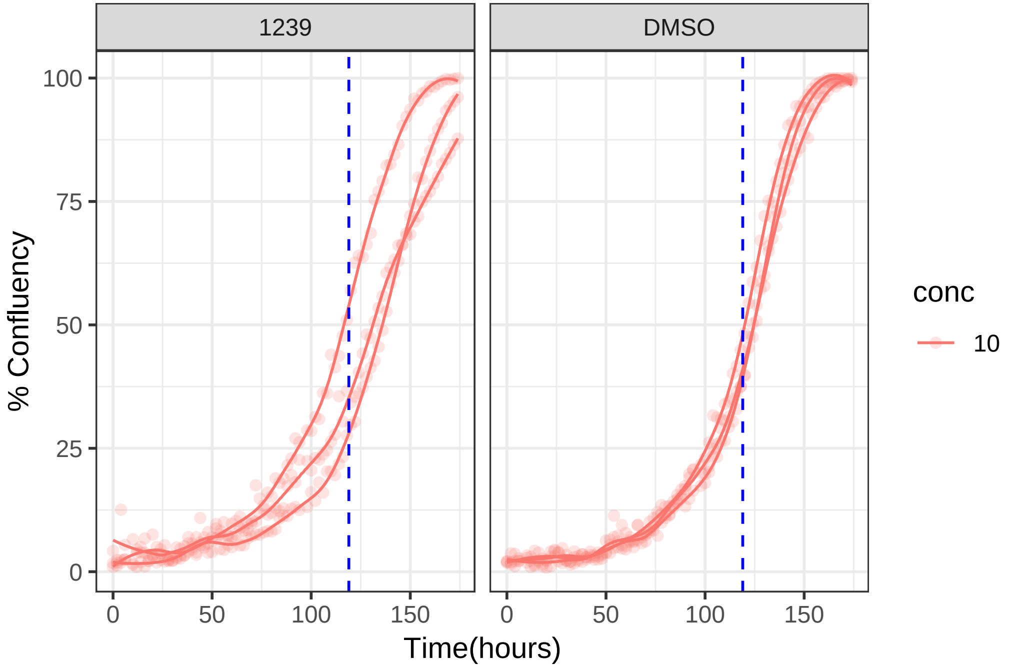

**10 uM 1239 RPE1 Ts12 Ts5**

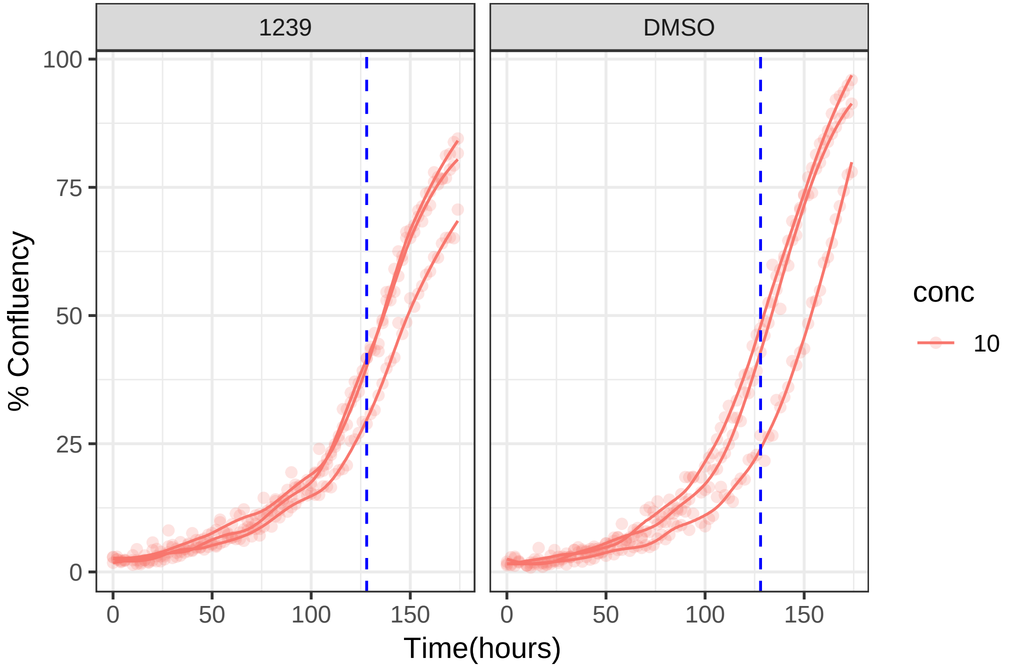

**10 uM 1243 RPE1 hTert**

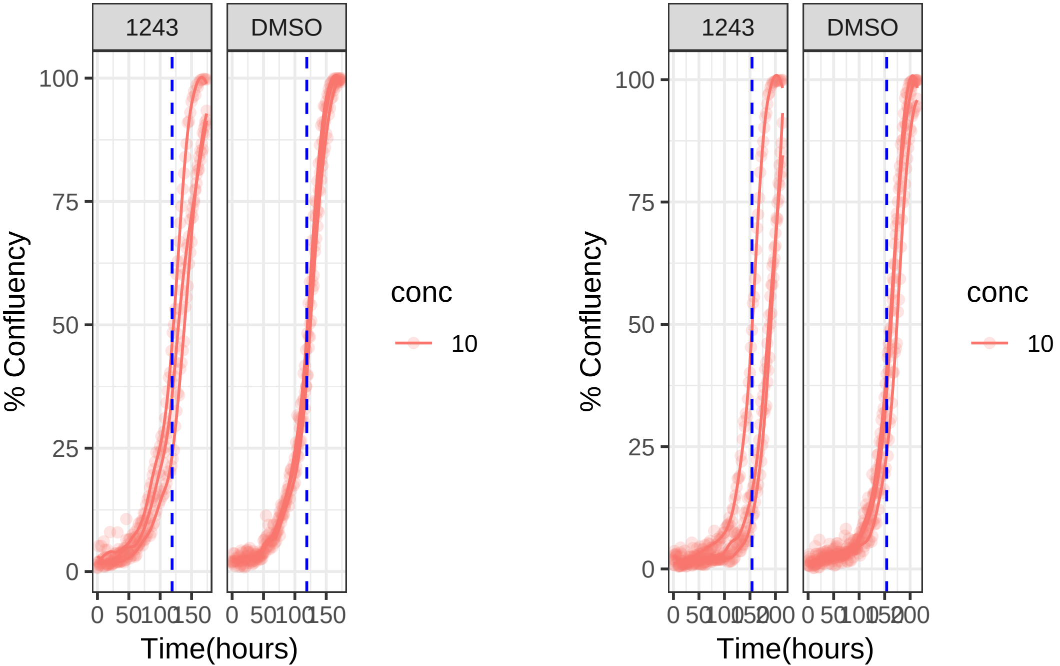

**10 uM 1243 RPE1 Ts12 Ts5**

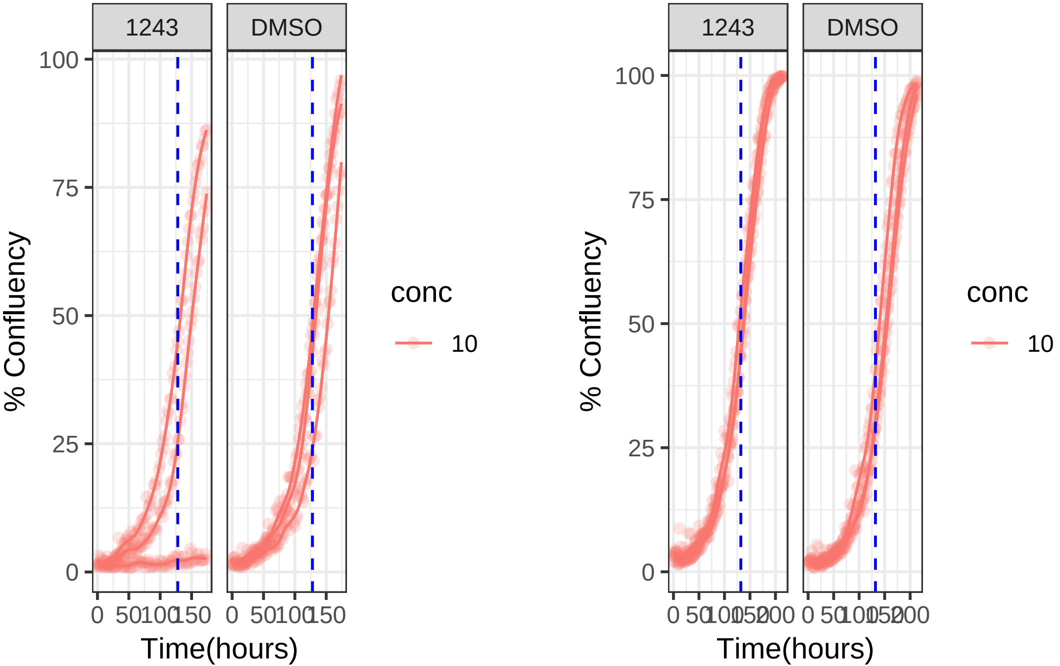

**10 uM 1254 RPE1 hTert**

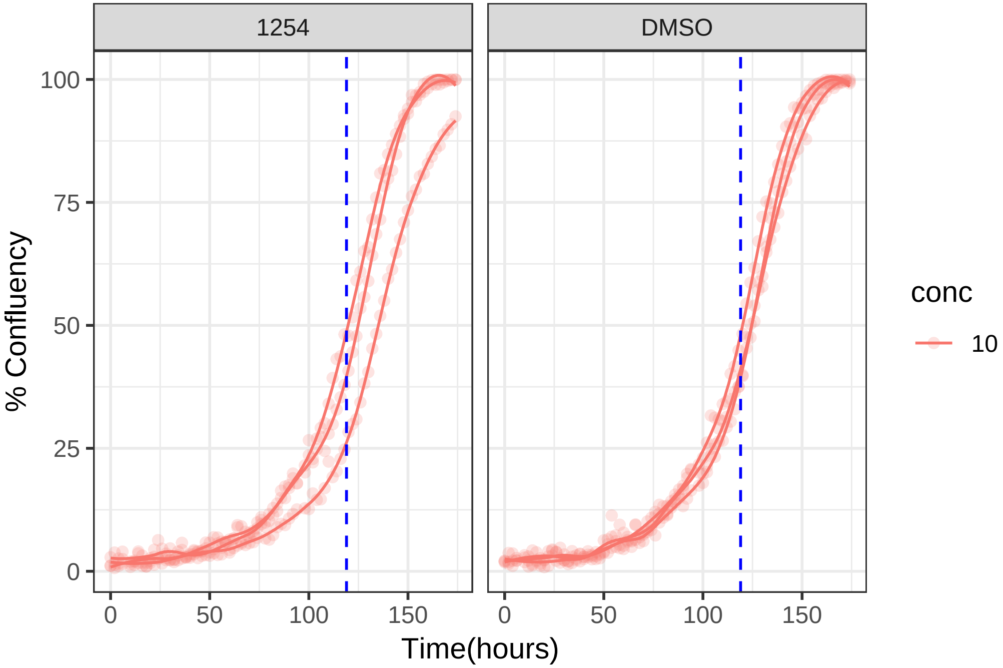

**10 uM 1254 RPE1 Ts12 Ts5**

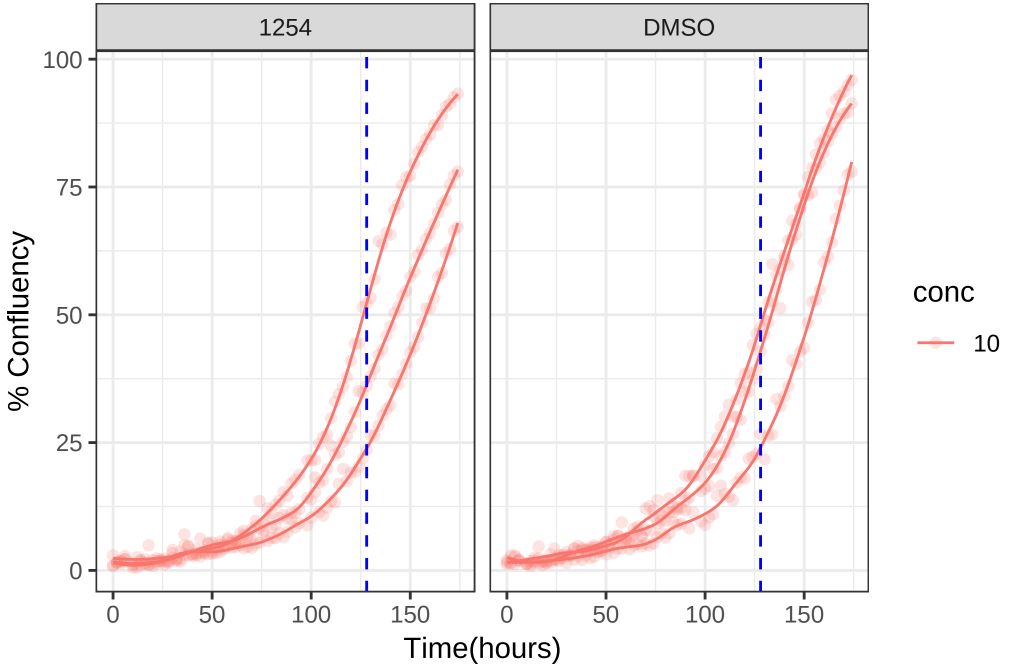

**10 uM 1268 RPE1 hTert**

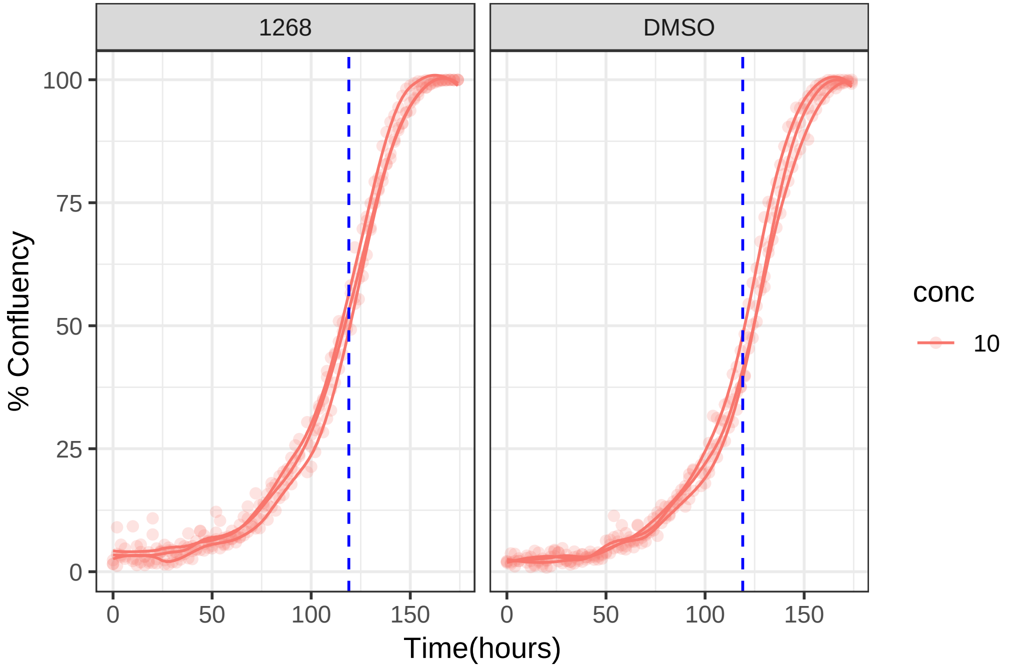

**10 uM 1268 RPE1 Ts12 Ts5**

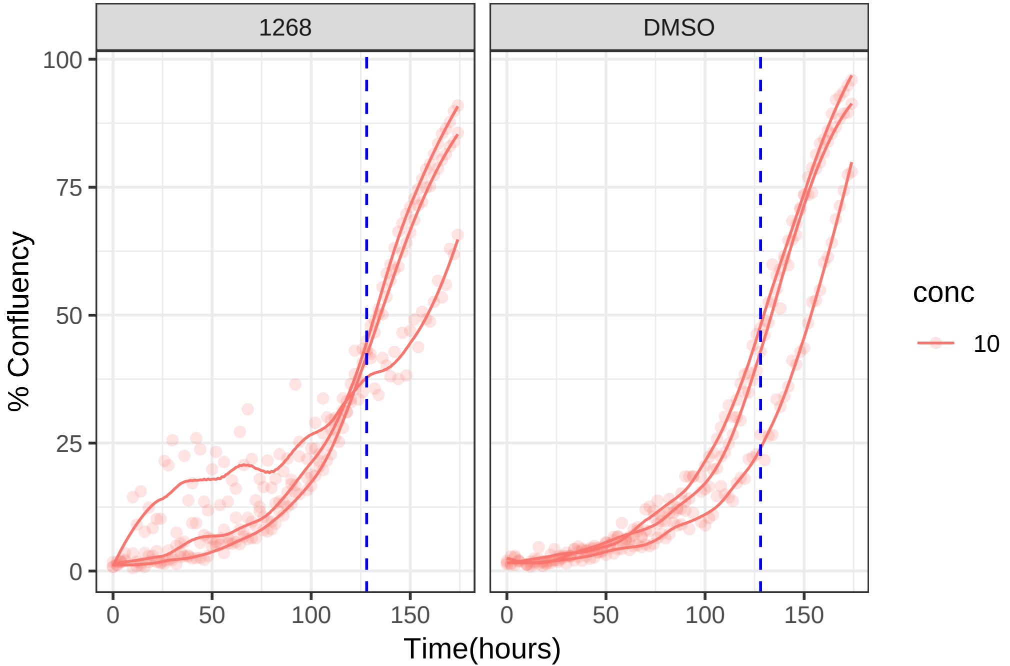

**0.1 uM 1275 RPE1 hTert**

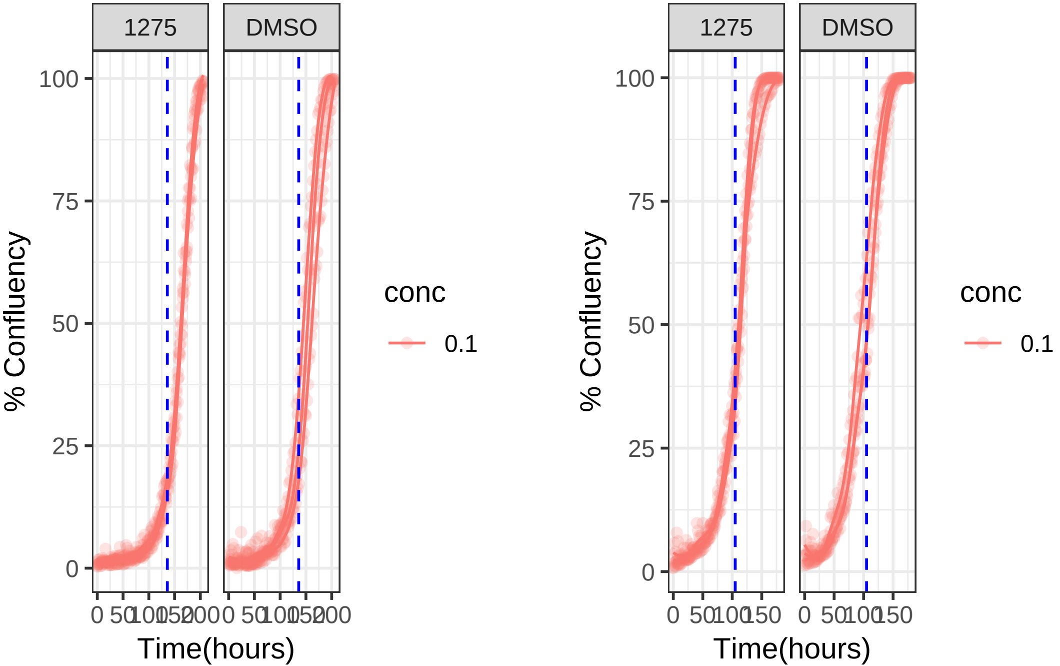

**0.1 uM 1275 RPE1 Ts12 Ts5**

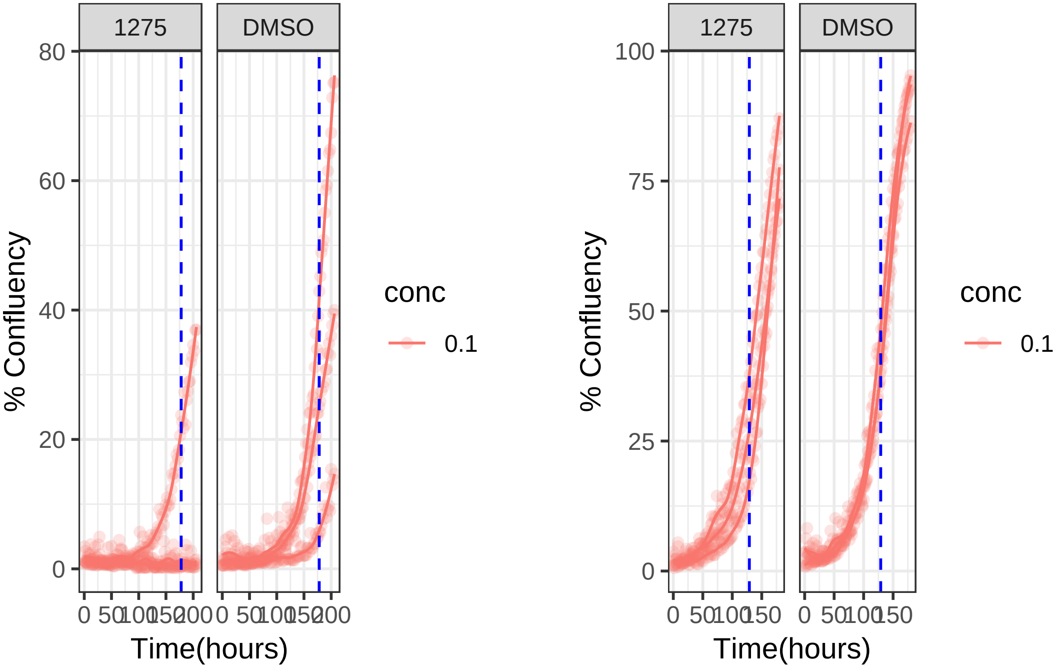

**1 uM 1310 RPE1 hTert**

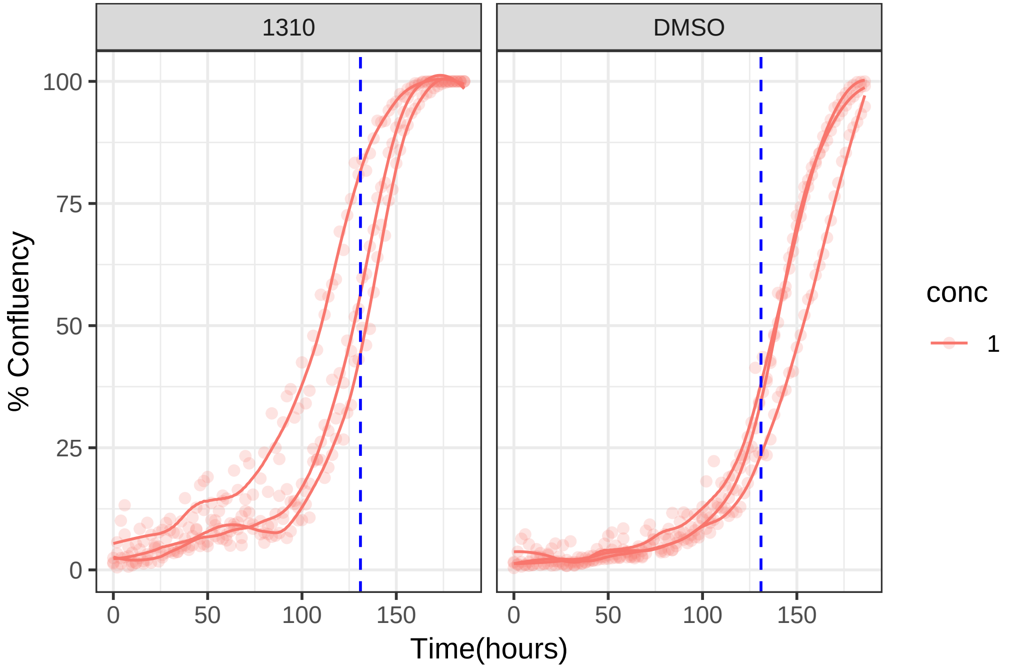

**1 uM 1310 RPE1 Ts12 Ts5**

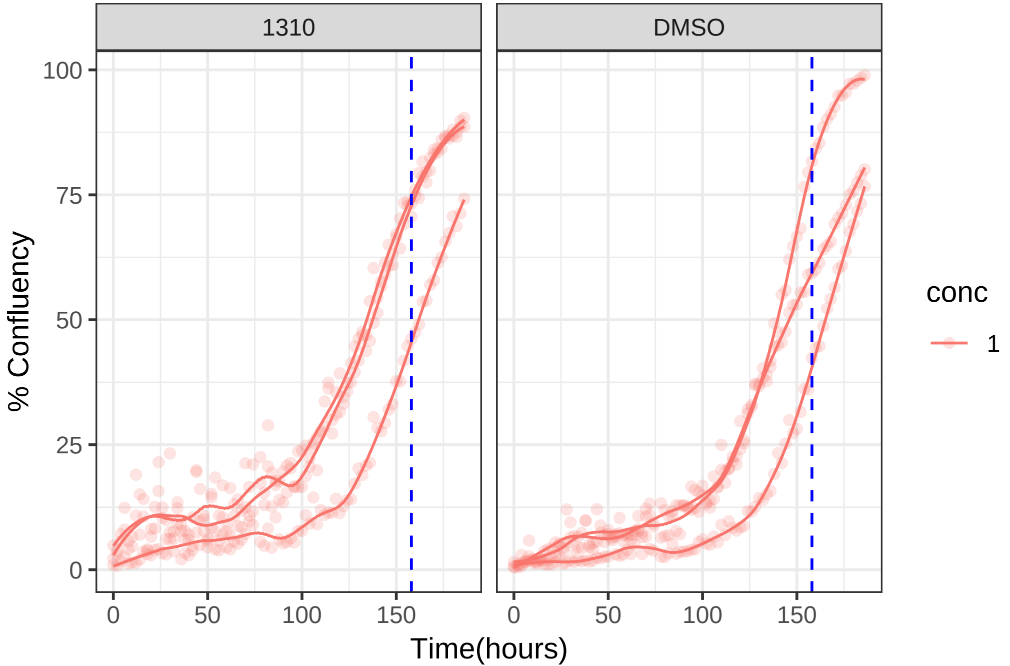

**1 uM 1334 RPE1 hTert**

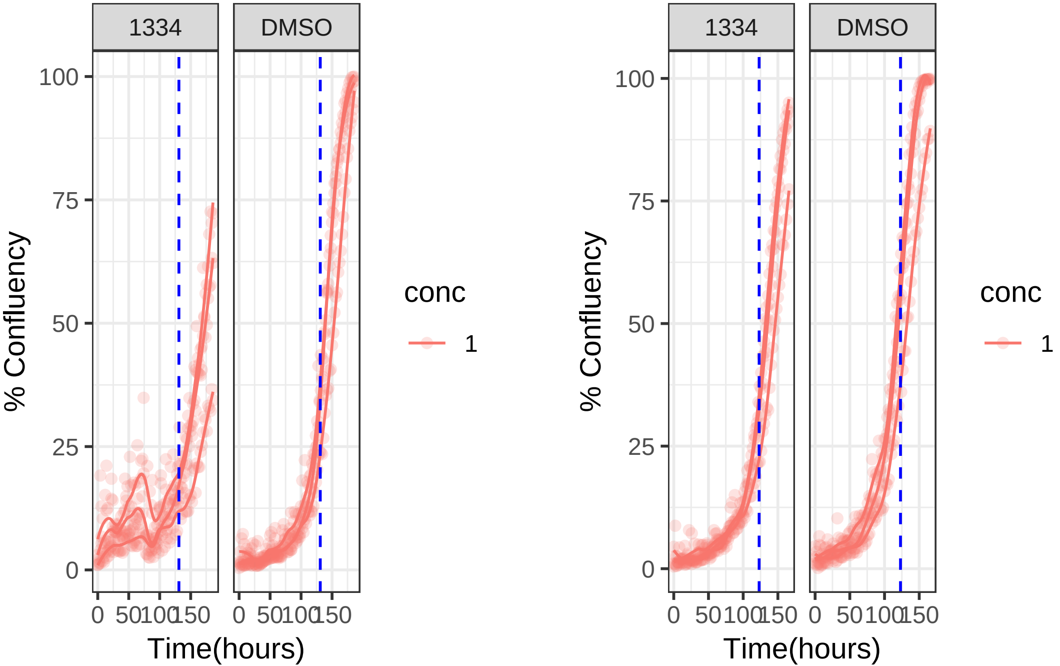

**1 uM 1334 RPE1 Ts12 Ts5**

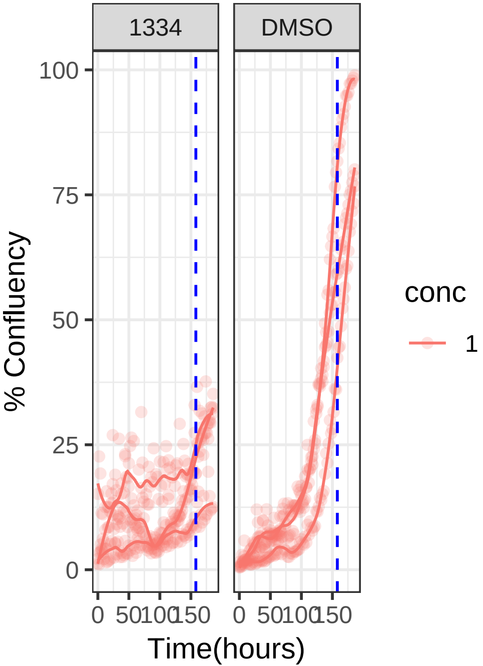


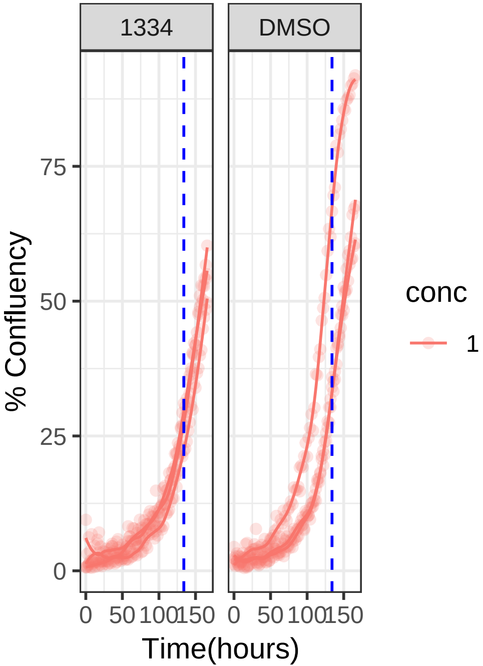

**10 uM 1355 RPE1 hTert**

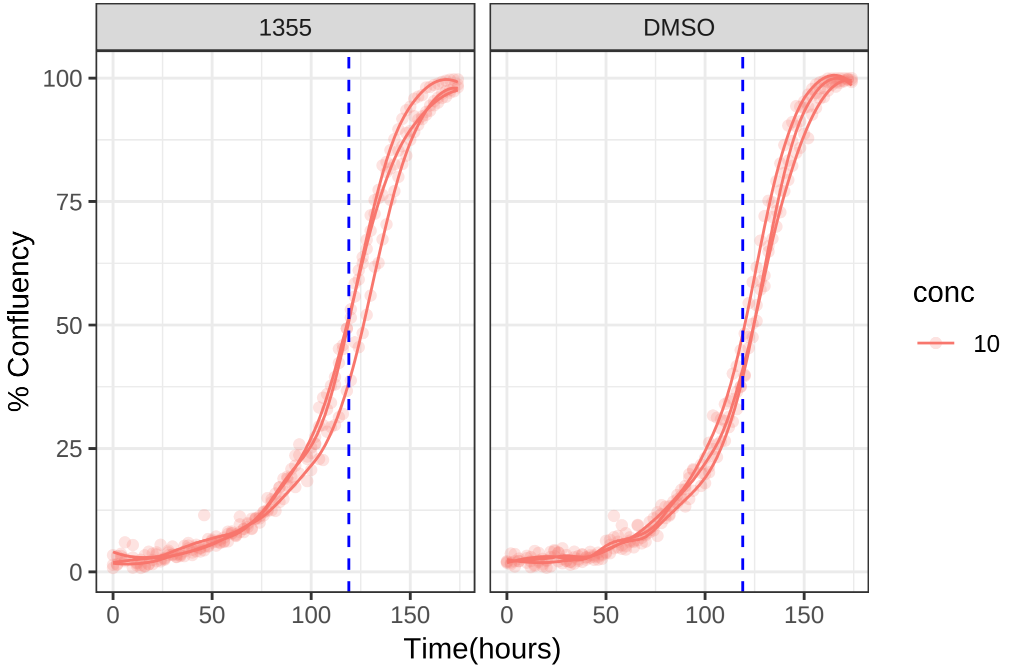

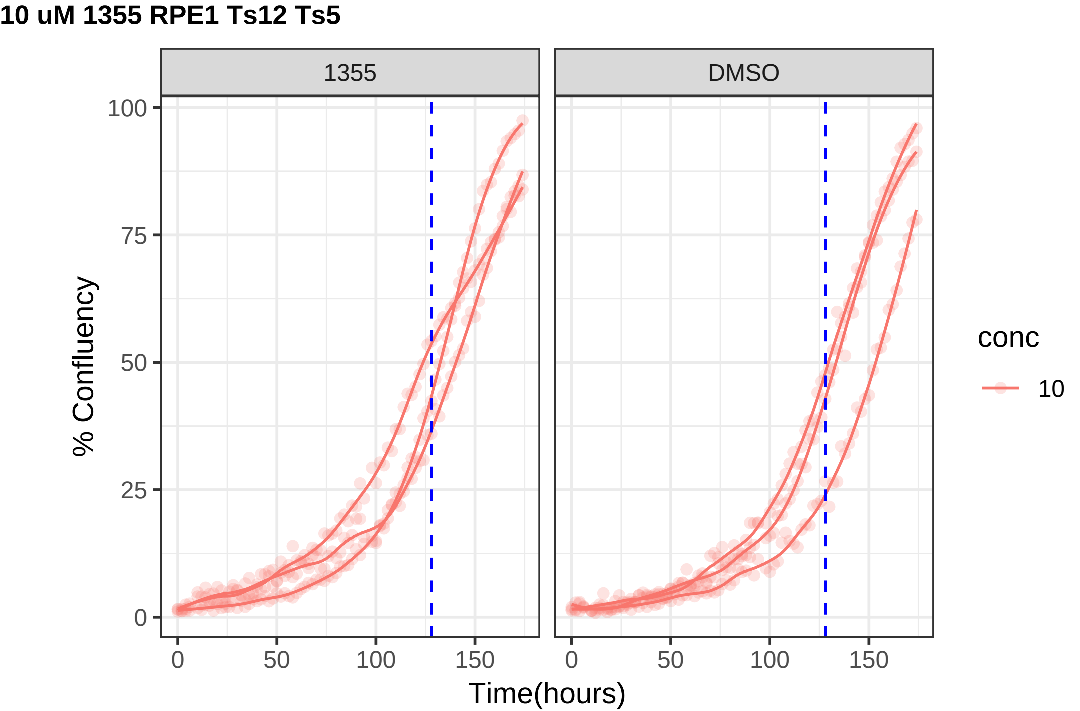
10 uM 1355 RPE1 Ts12 Ts5
1355
DMSO
% Confluency
100
75
50
25
0
0
50
100
150
Time(hours)
conc
10

**10 uM 1356 RPE1 hTert**

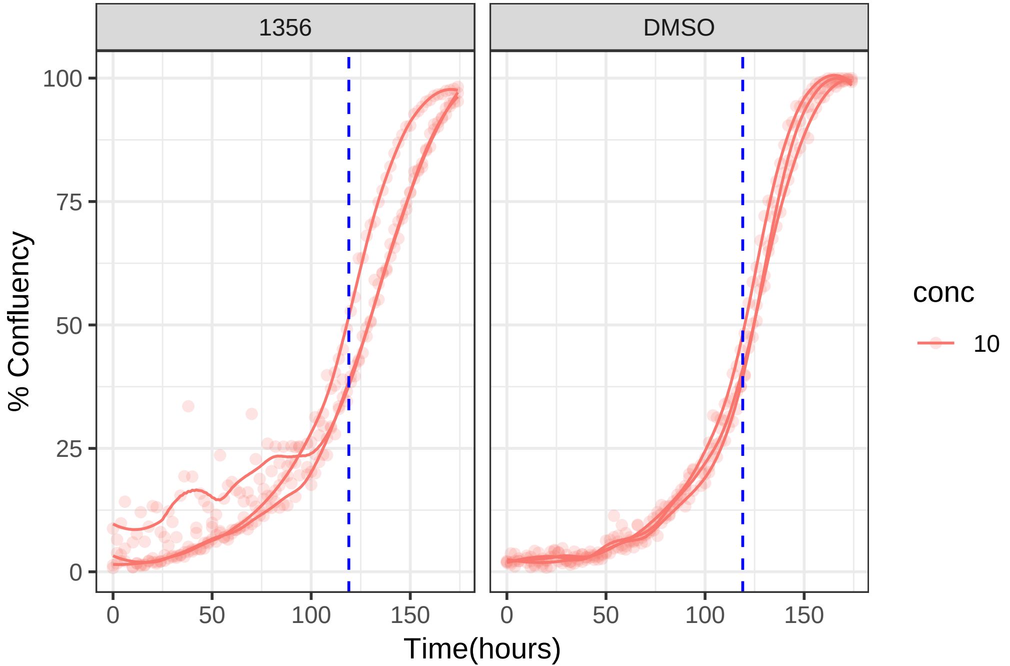

**10 uM 1356 RPE1 Ts12 Ts5**

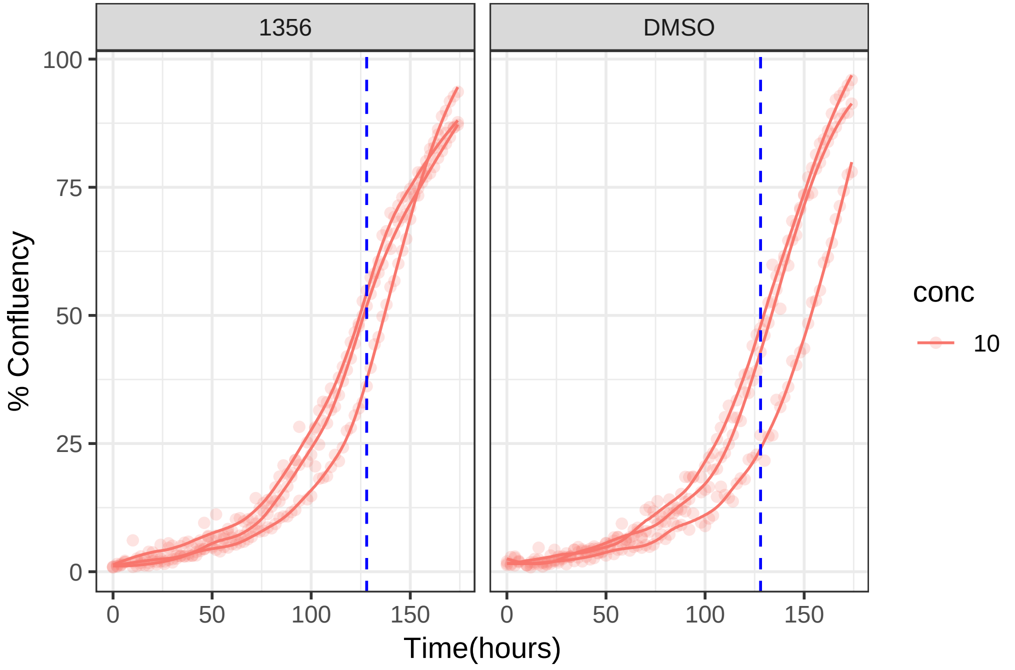

**1 uM 1362 RPE1 hTert**

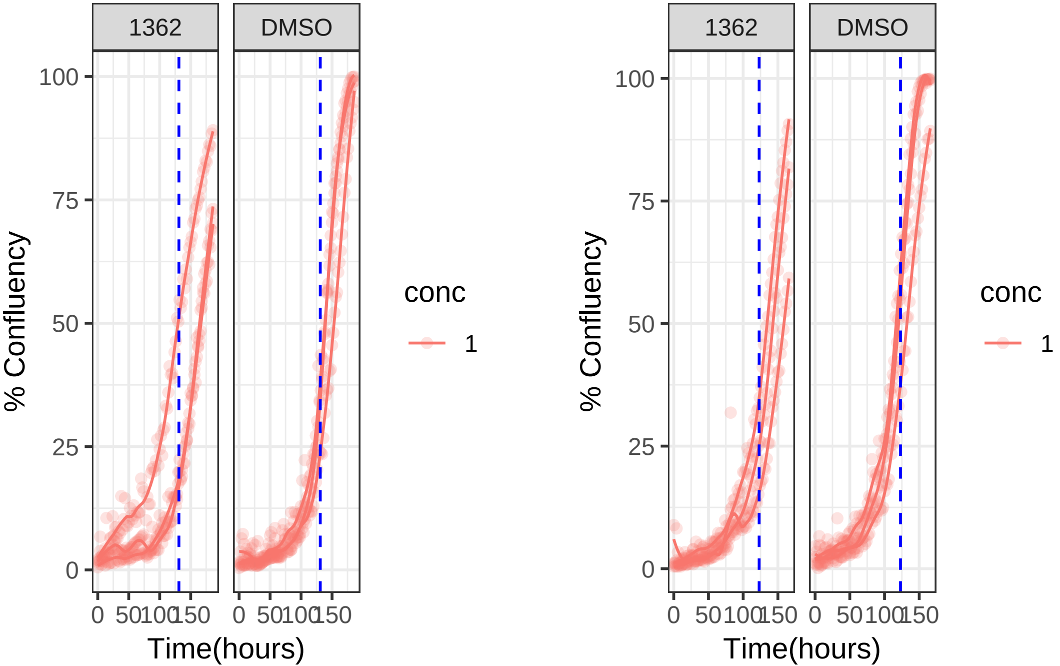

**1 uM 1362 RPE1 Ts12 Ts5**

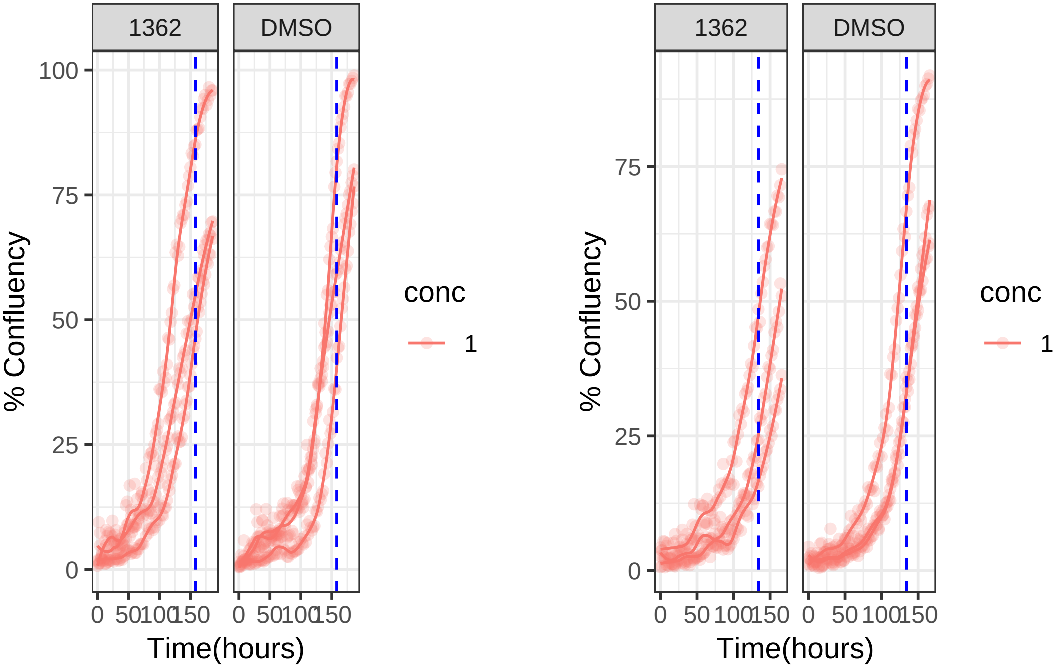

**10 uM 1375 RPE1 hTert**

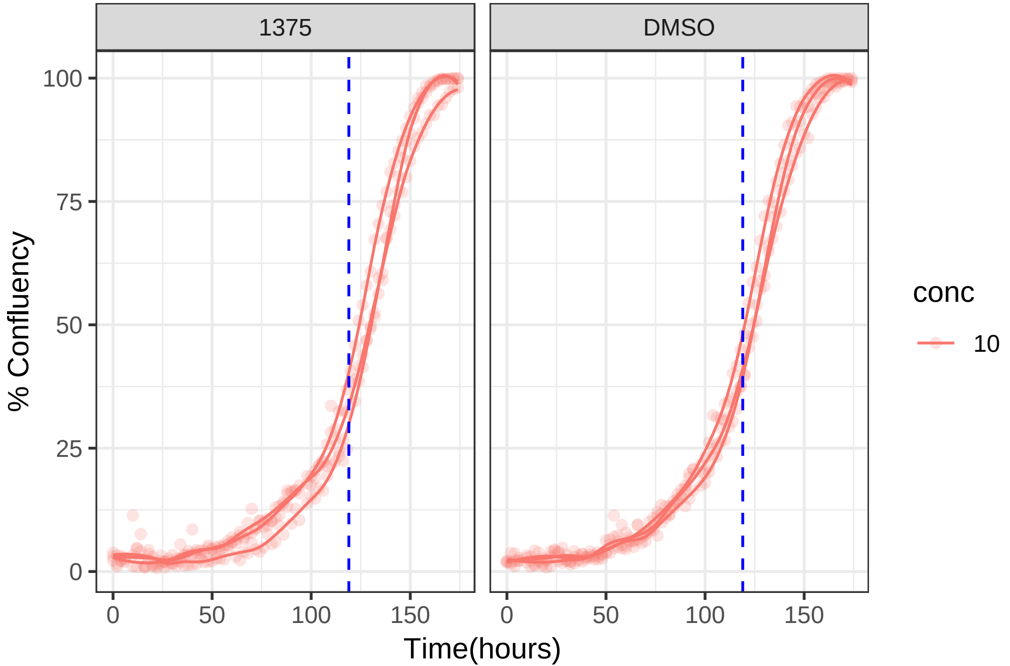

**10 uM 1375 RPE1 Ts12 Ts5**

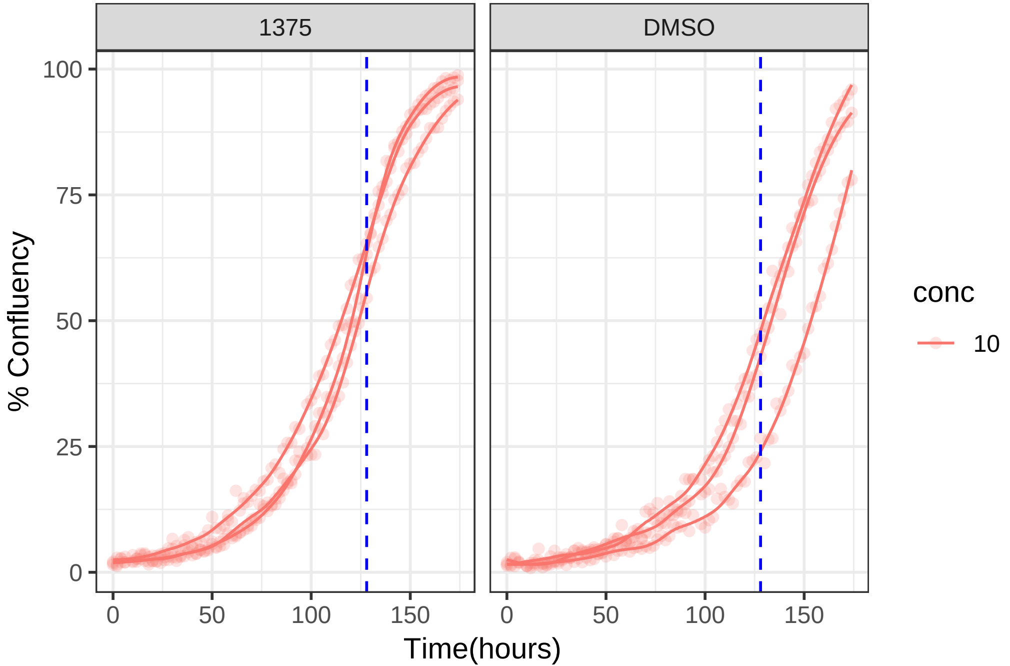

1 uM 1377 RPE1 hTert

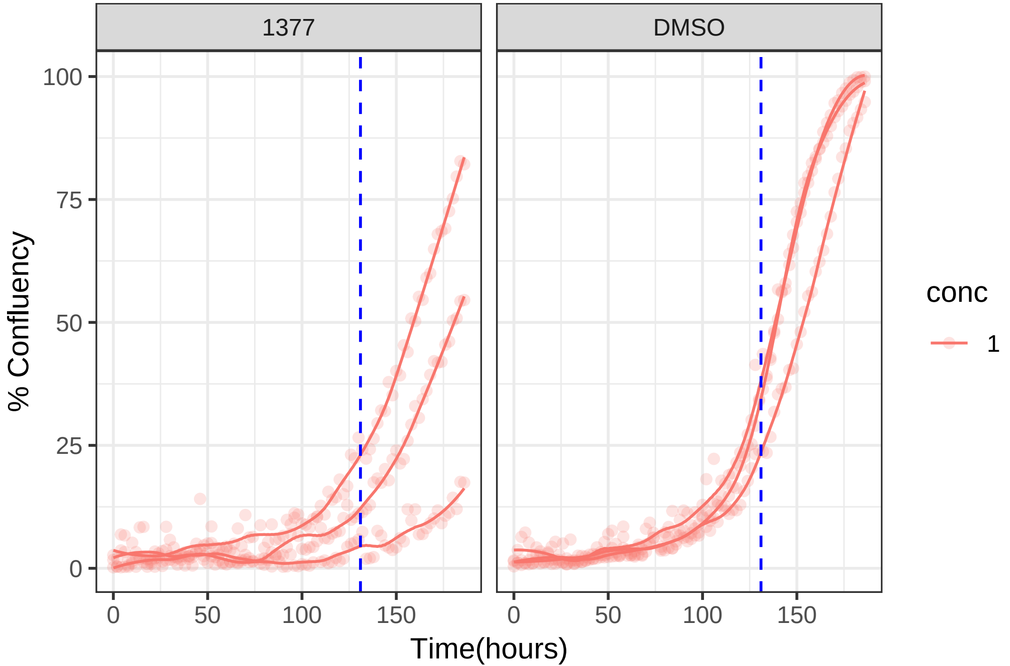

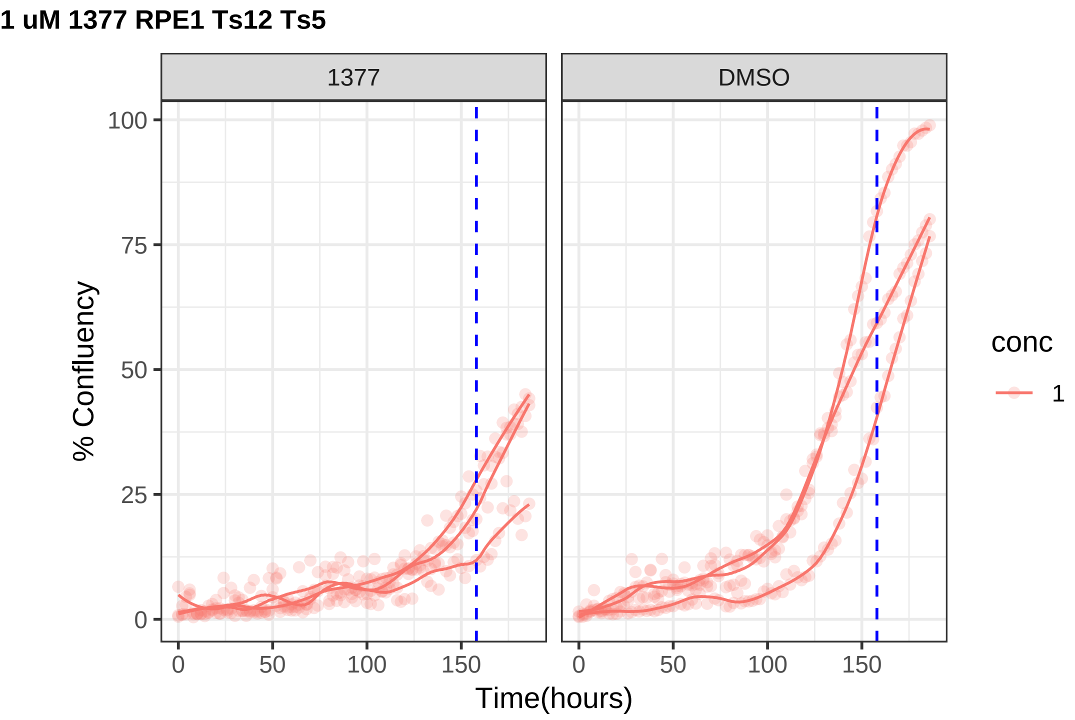

1 uM 1377 RPE1 Ts12 Ts5
1377
DMSO
% Confluency
100
75
50
25
0
0
50
100
150
Time(hours)
conc
1

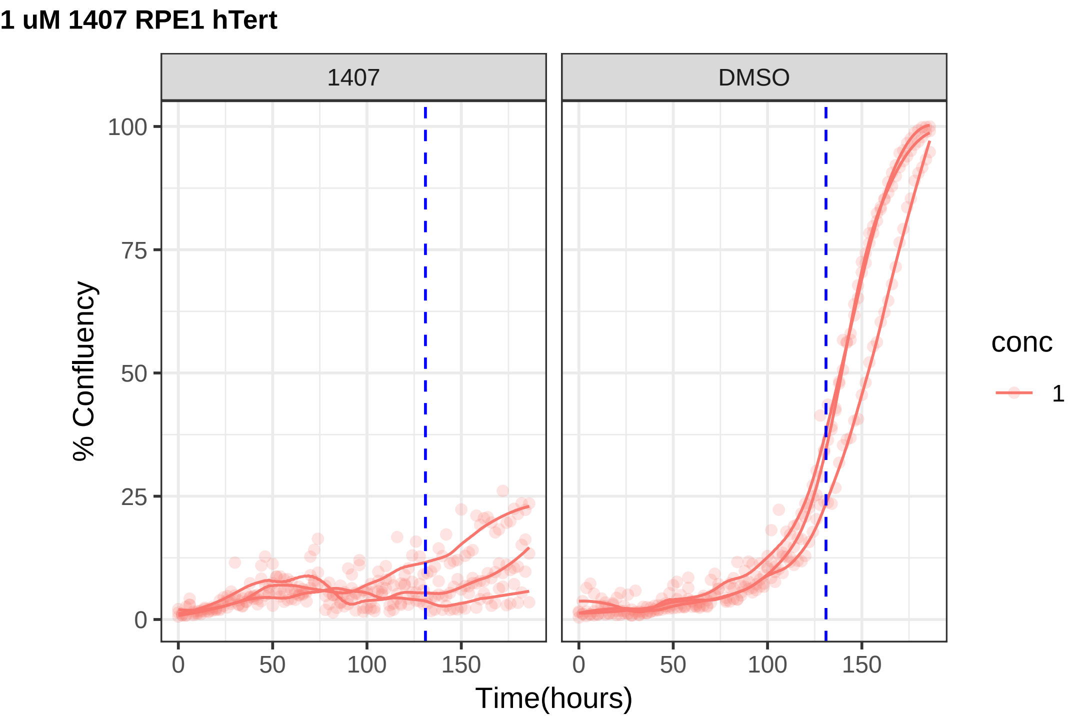
1 uM 1407 RPE1 hTert
1407
DMSO
% Confluency
100
75
50
25
0
0
50
100
150
Time(hours)
conc
1

**0.1 uM 1407 RPE1 hTert**

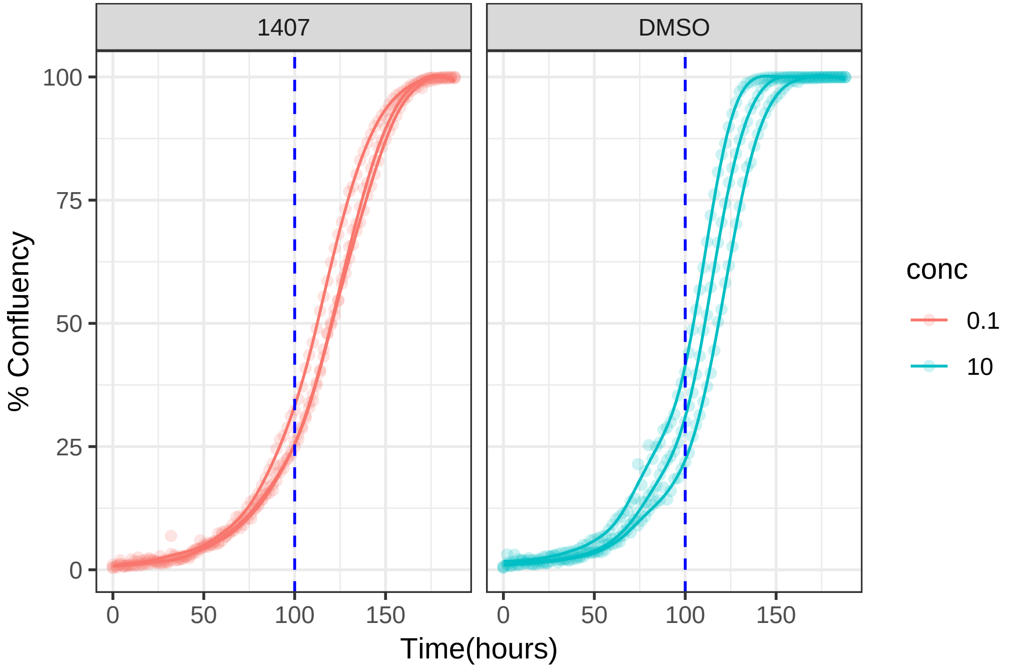

**0.5 uM 1407 RPE1 hTert**

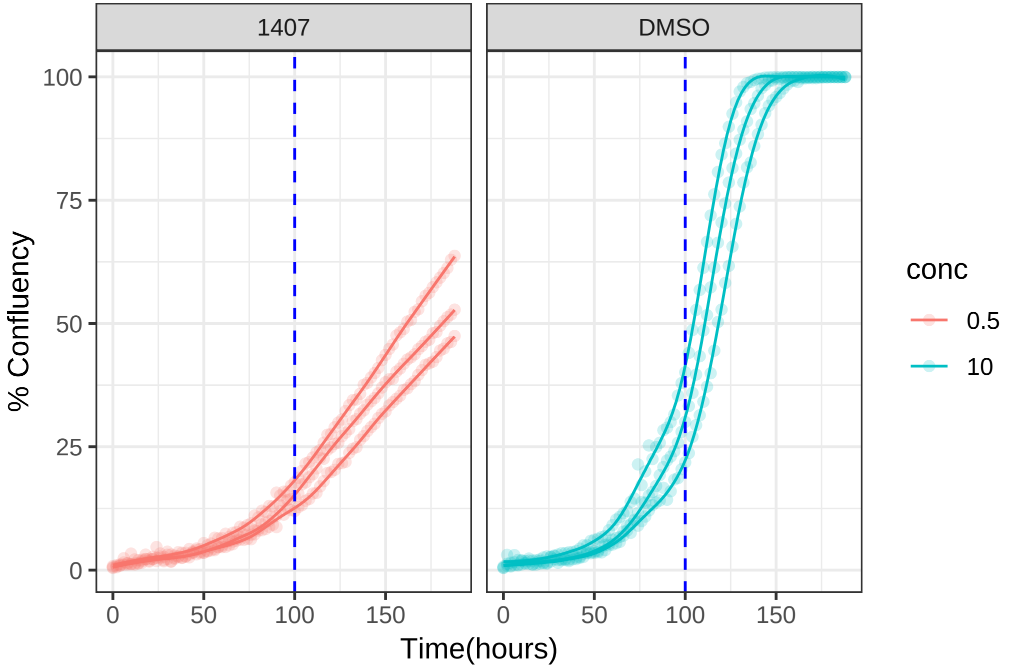

1 uM 1407 RPE1 Ts12 Ts5

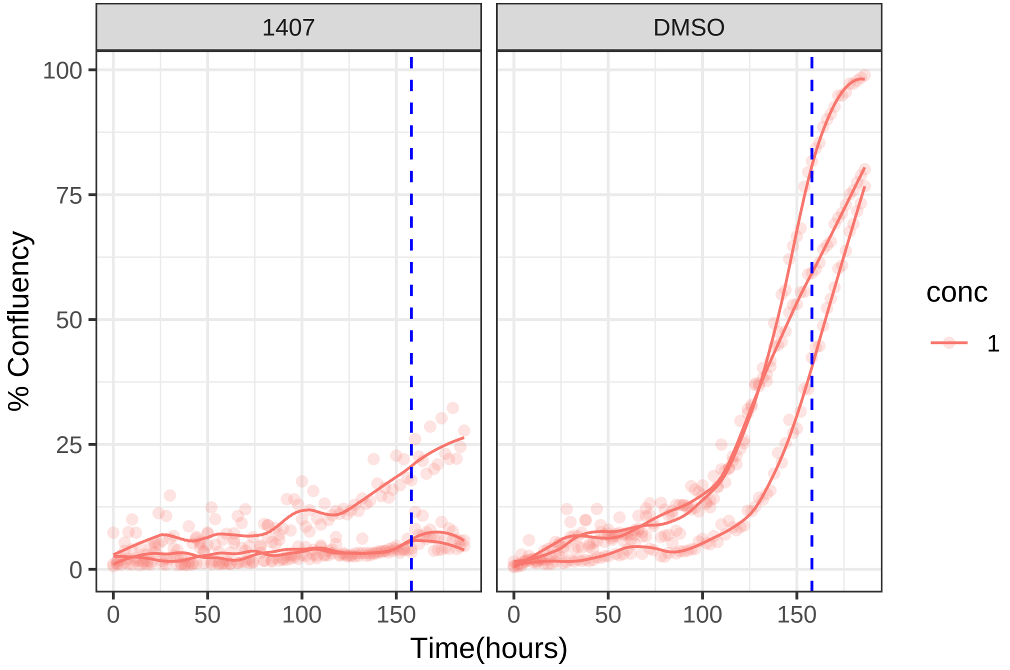

### 0.1 uM 1407 RPE1 Ts12 Ts5

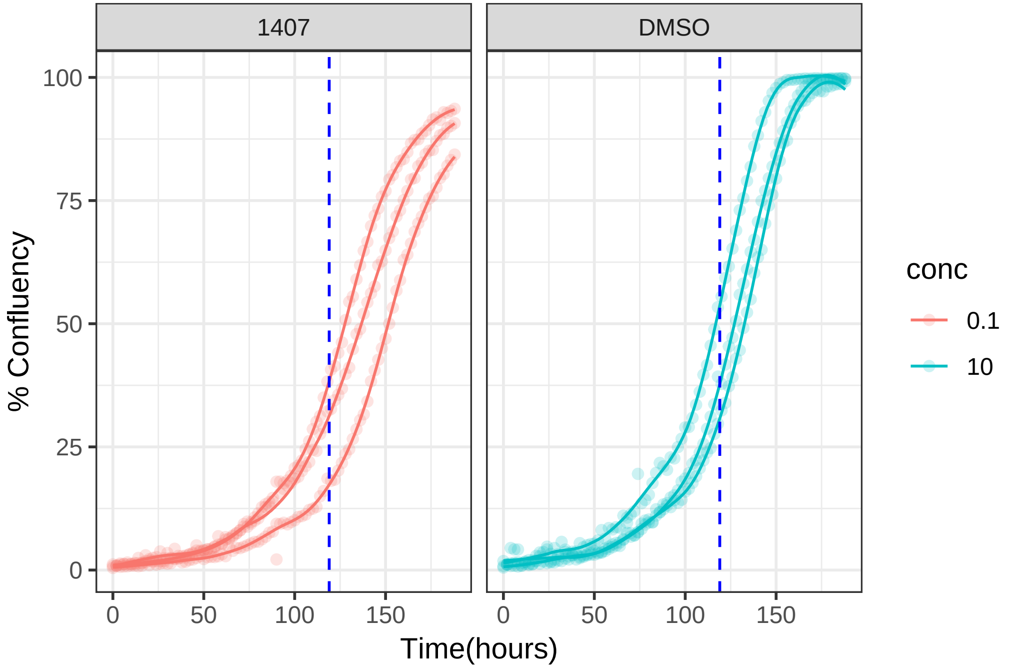

**0.5 uM 1407 RPE1 Ts12 Ts5**

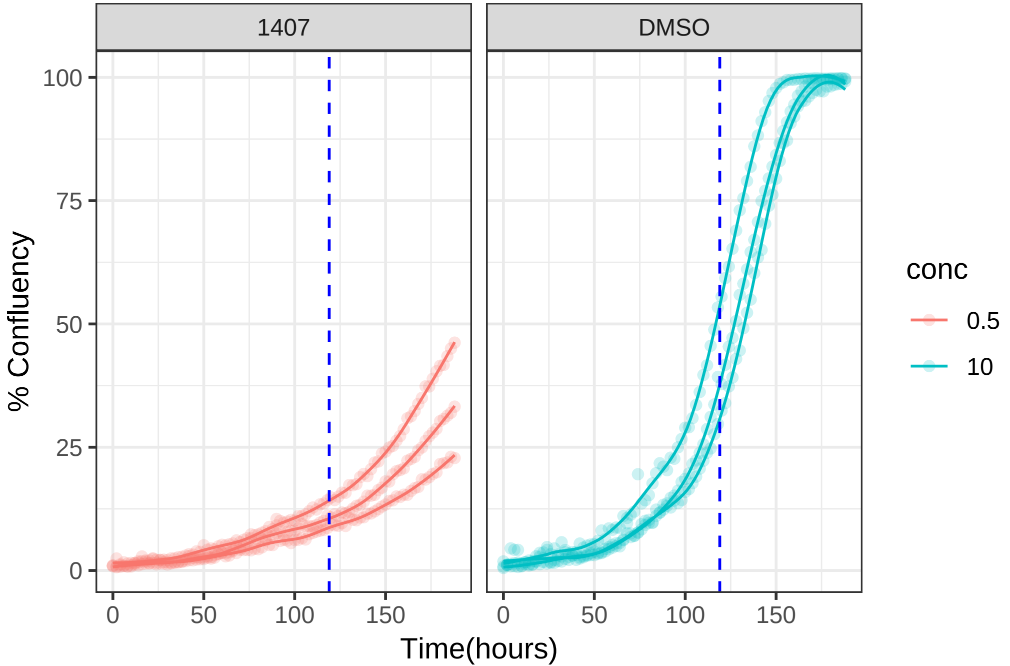

**10 uM 1424 RPE1 hTert**

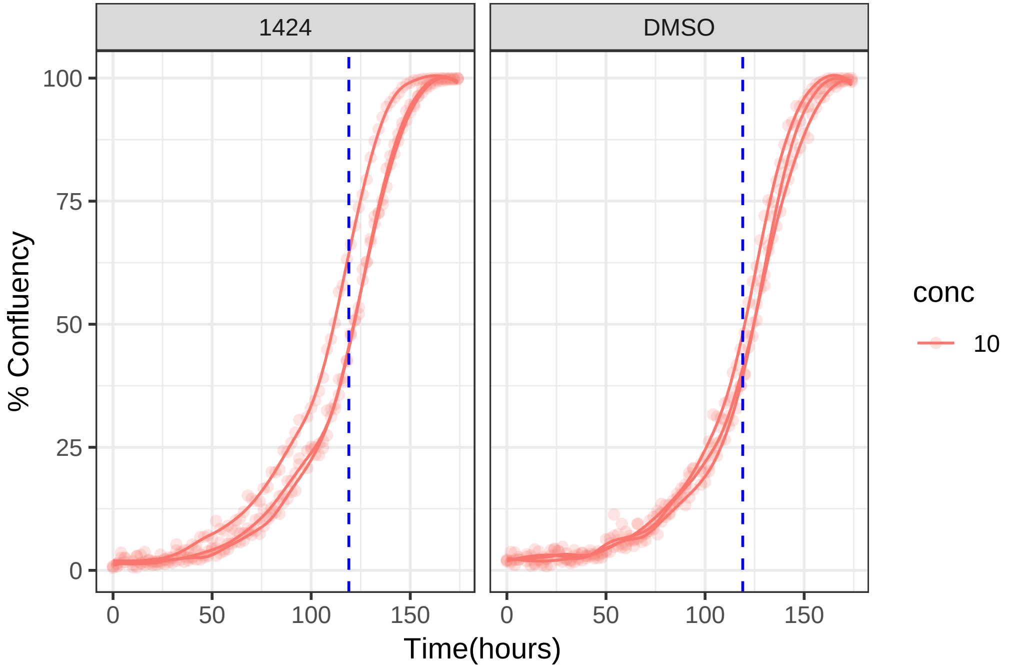

10 uM 1424 RPE1 Ts12 Ts5

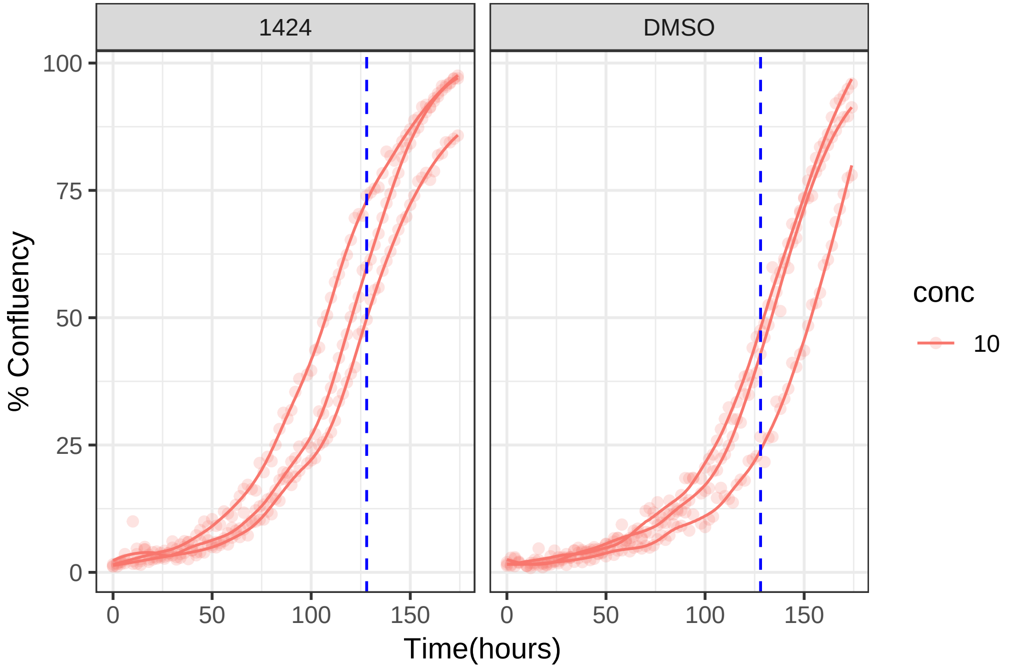

#### 10 uM 1459 RPE1 hTert

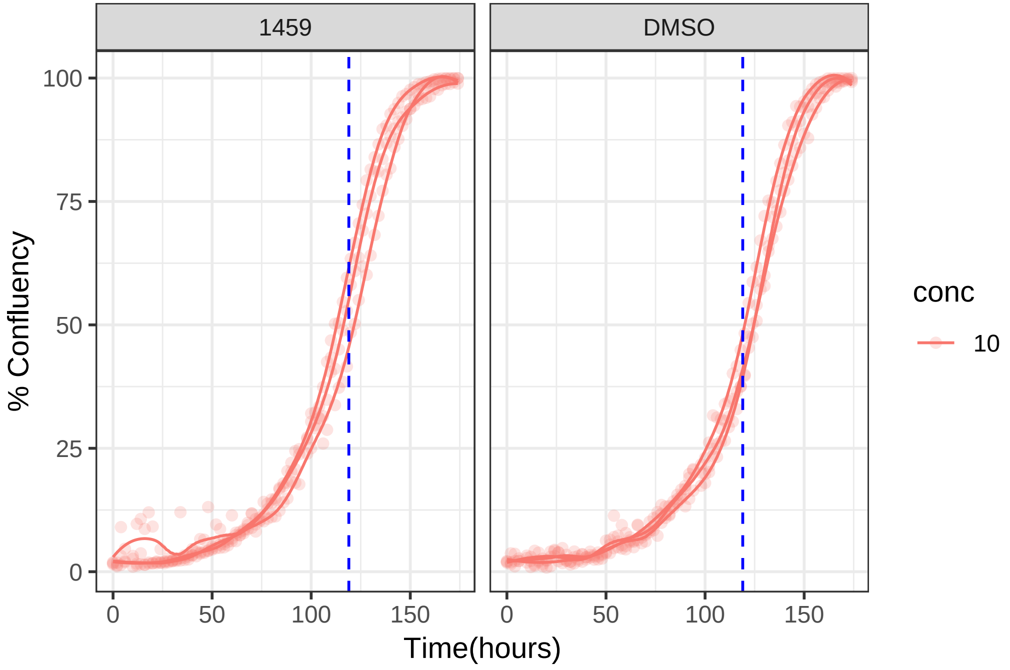

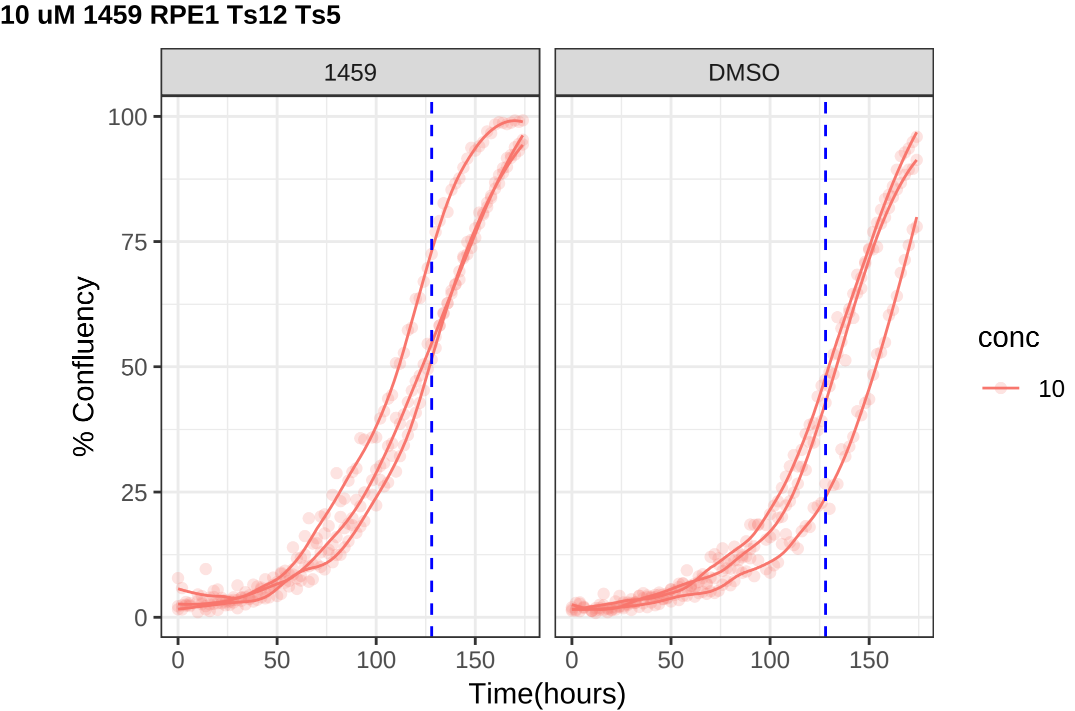
10 uM 1459 RPE1 Ts12 Ts5
1459
DMSO
% Confluency
100
75
50
25
0
0
50
100
150
Time(hours)
conc
10

**1 uM 1463 RPE1 hTert**

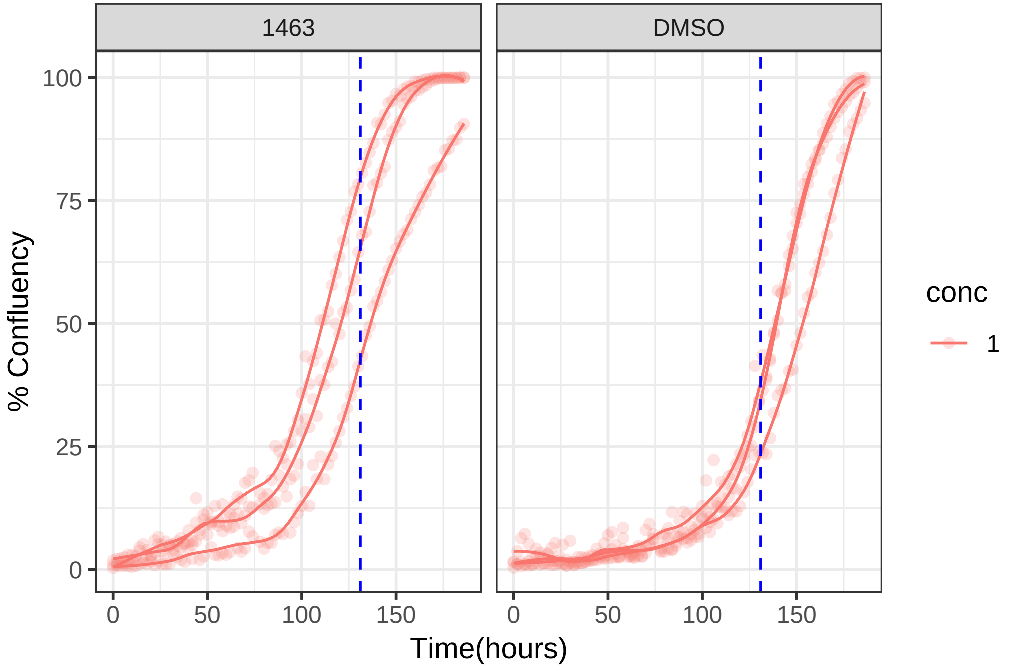

**1 uM 1463 RPE1 Ts12 Ts5**

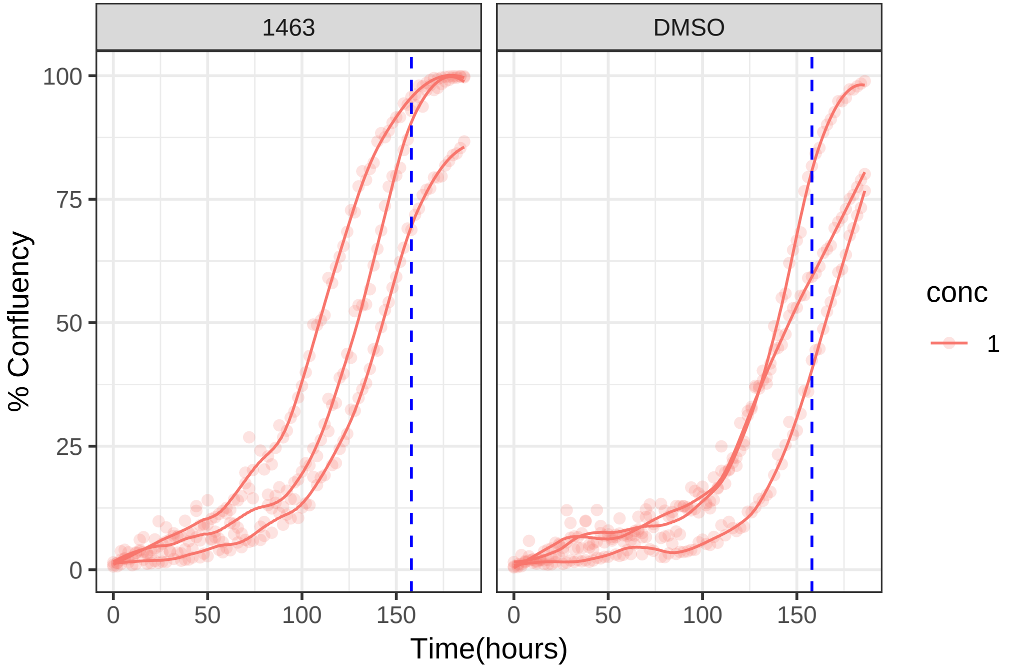

**1 uM 1472 RPE1 hTert**

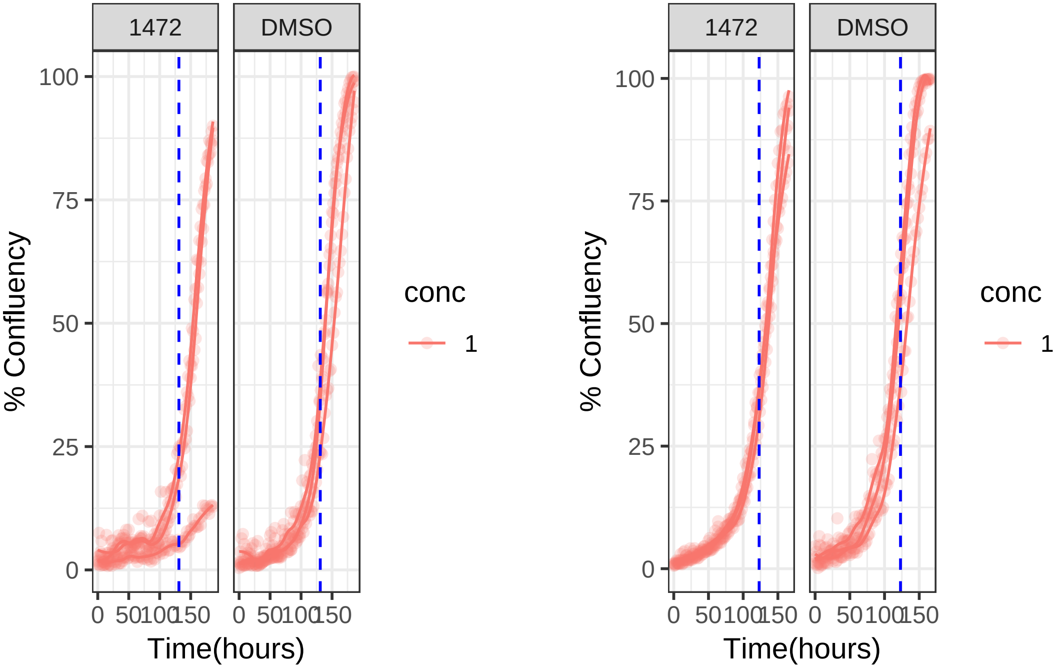

### 1 uM 1472 RPE1 Ts12 Ts5

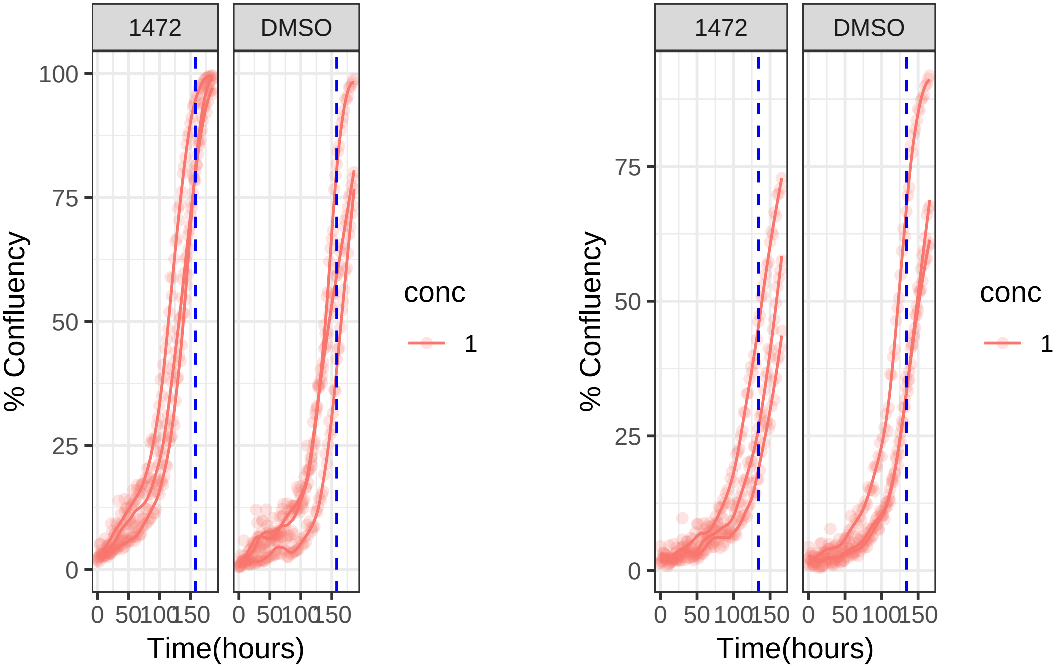

**0.1 uM 1494 RPE1 hTert**

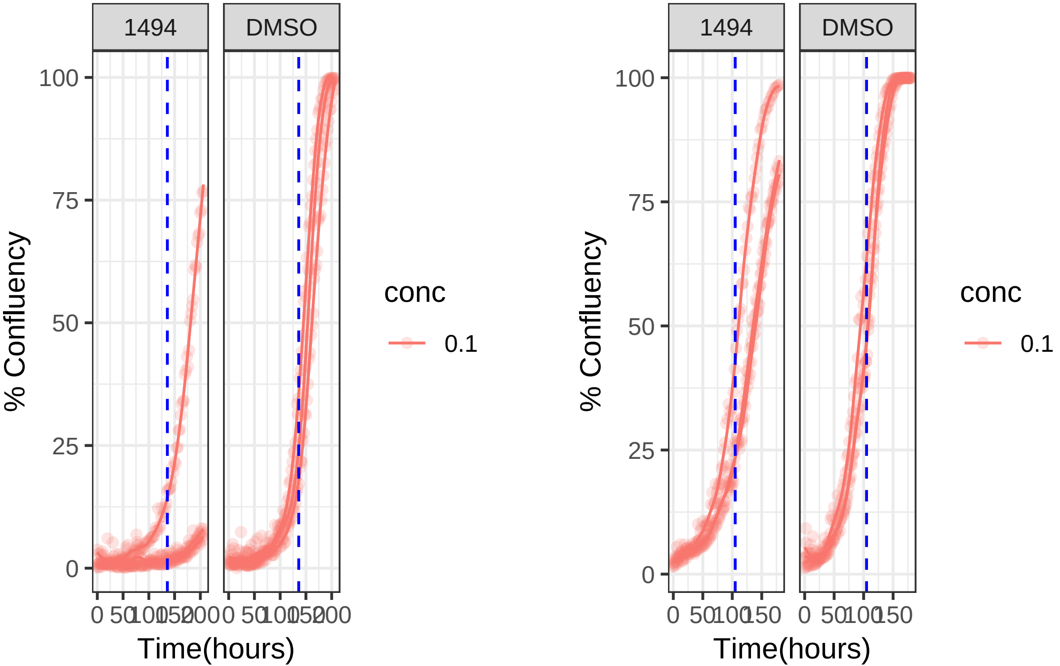

**0.1 uM 1494 RPE1 Ts12 Ts5**

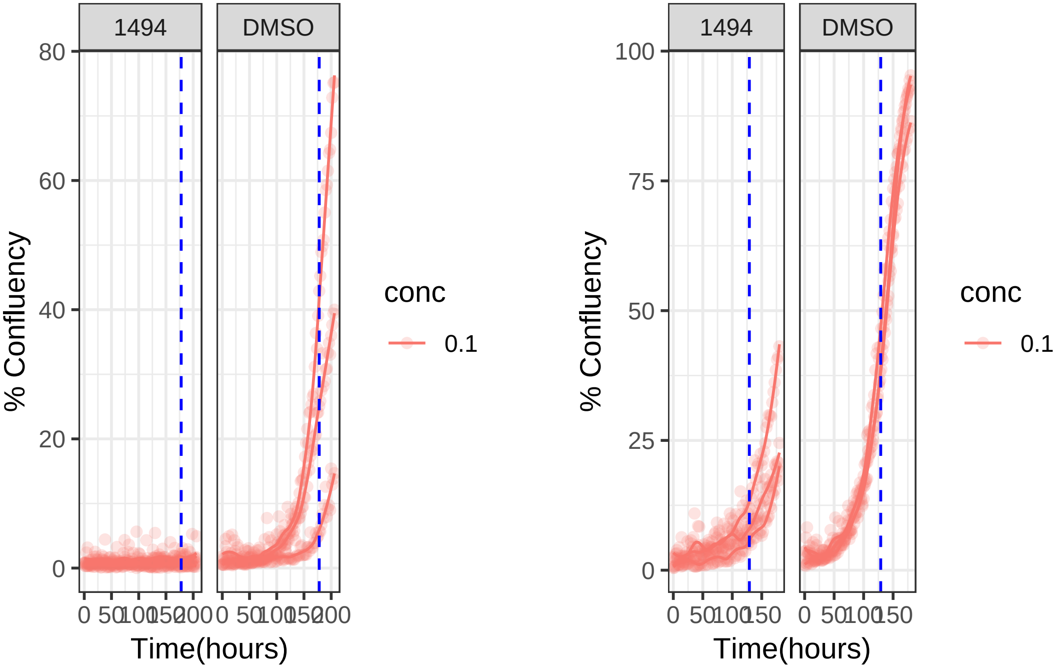

**1 uM 1516 RPE1 hTert**

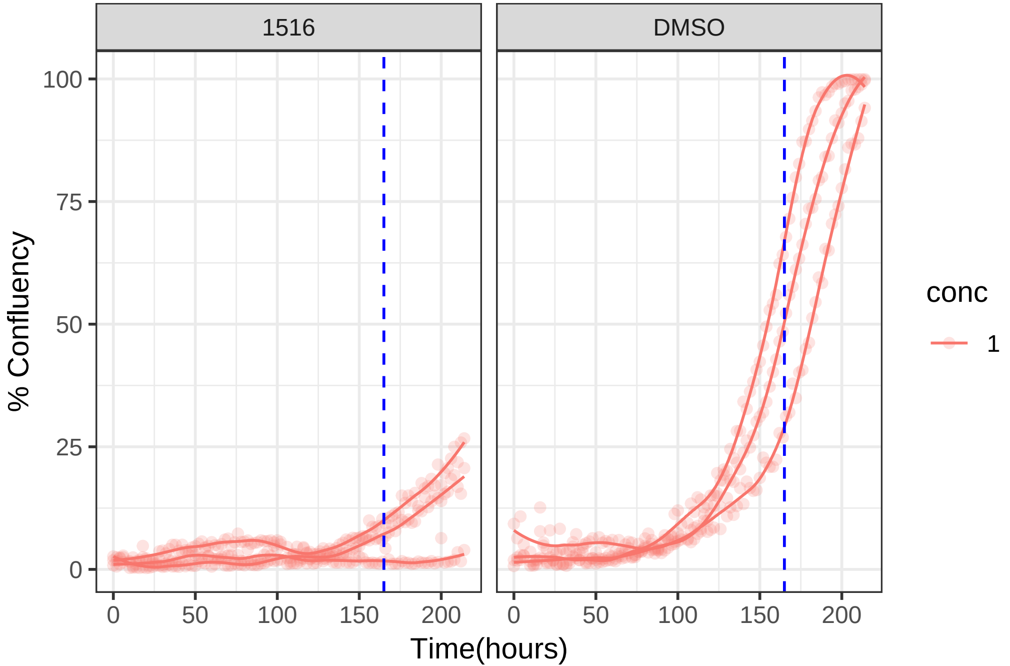

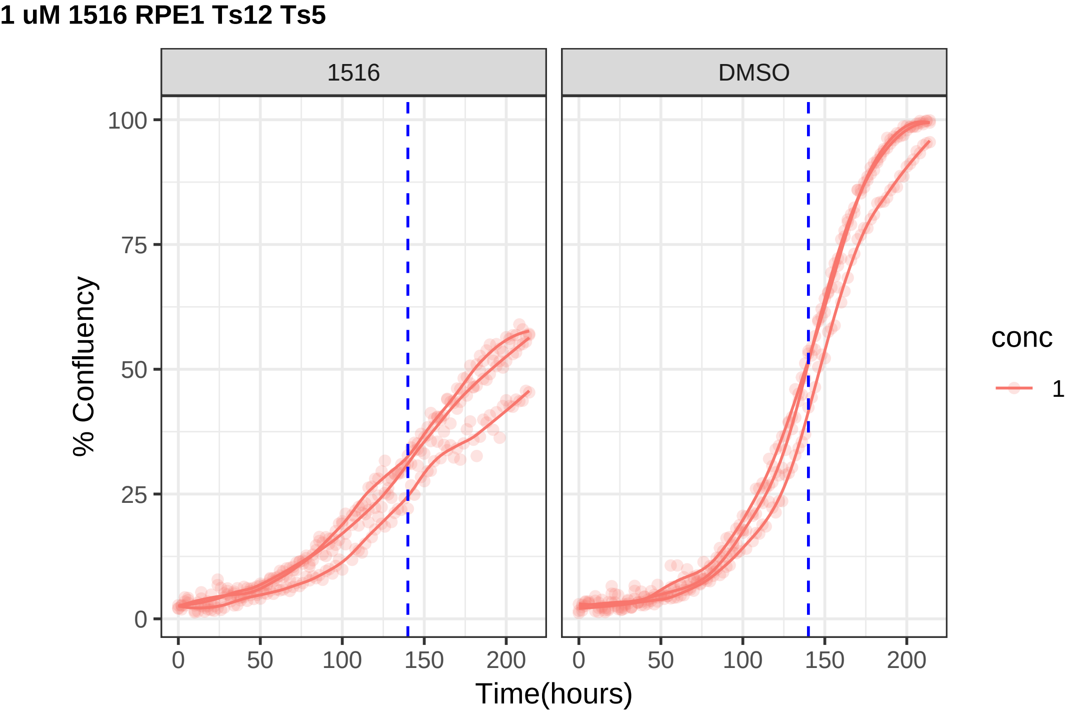

1 uM 1516 RPE1 Ts12 Ts5
1516
DMSO
% Confluency
Time(hours)
0
25
50
75
100
0
50
100
150
200
conc
1

**0.1 uM 1520 RPE1 hTert**

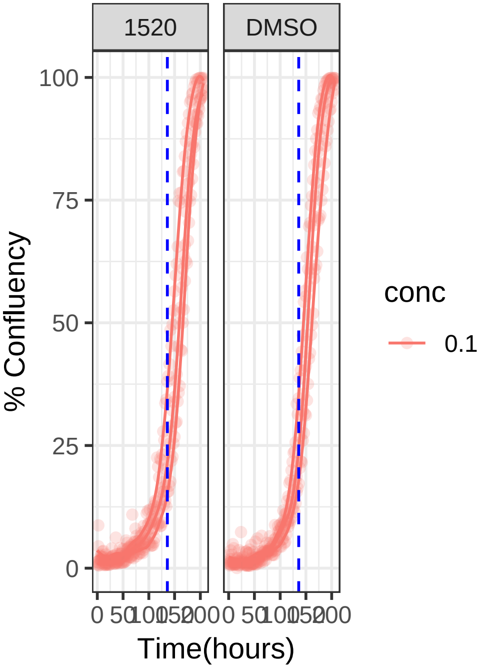


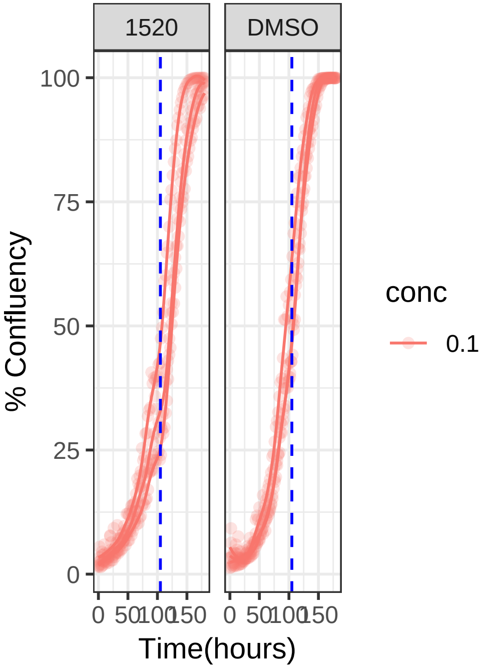

**0.1 uM 1520 RPE1 Ts12 Ts5**

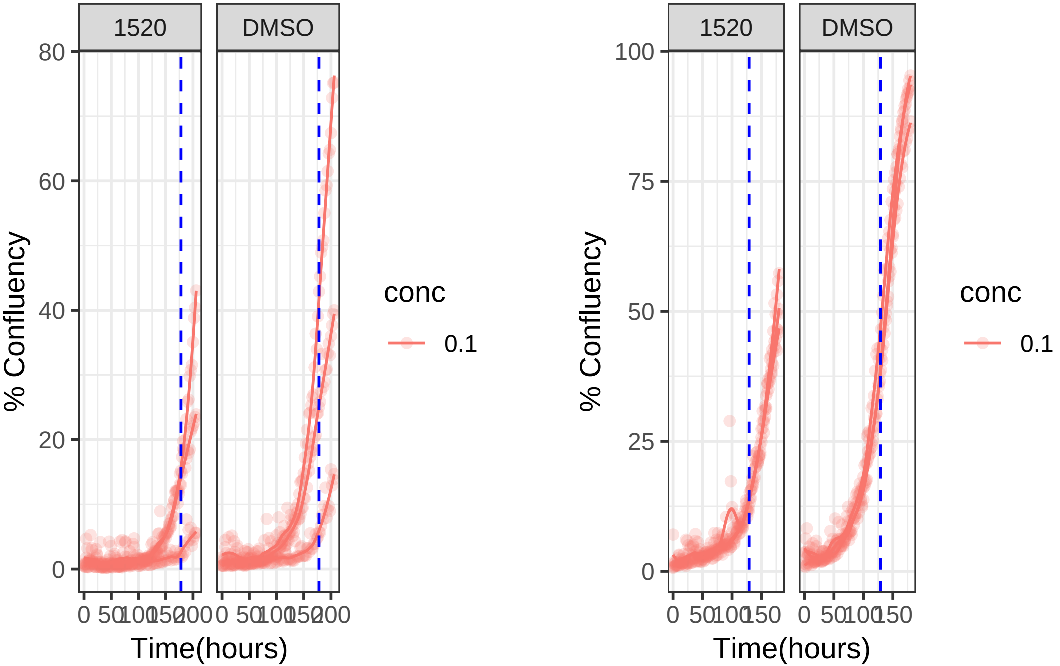

**0.01 uM 1561 RPE1 hTert**

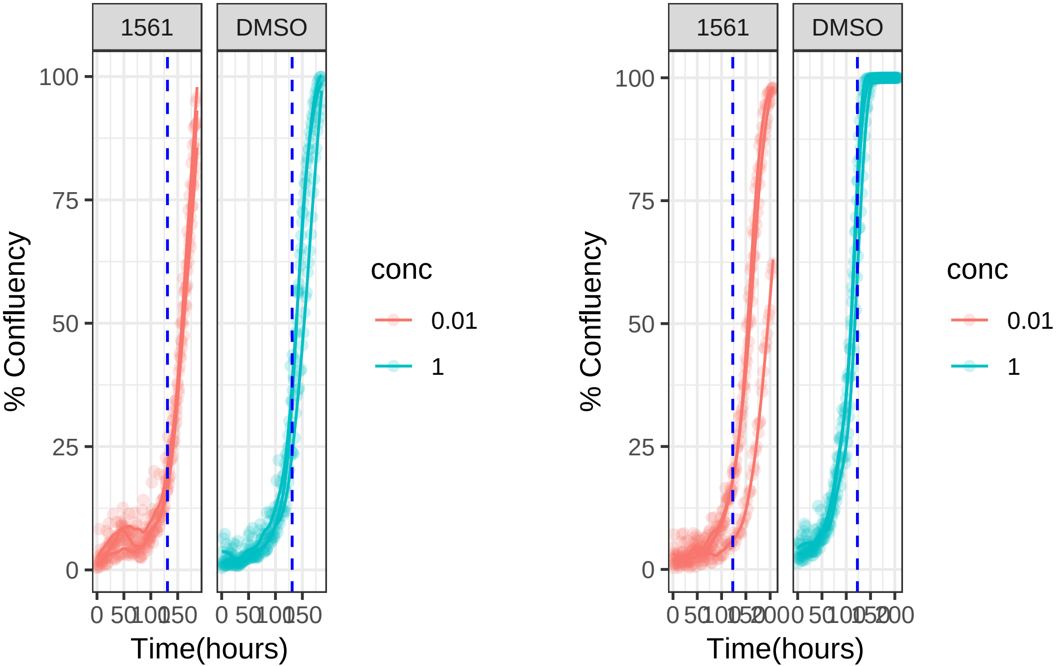

**0.1 uM 1561 RPE1 hTert**

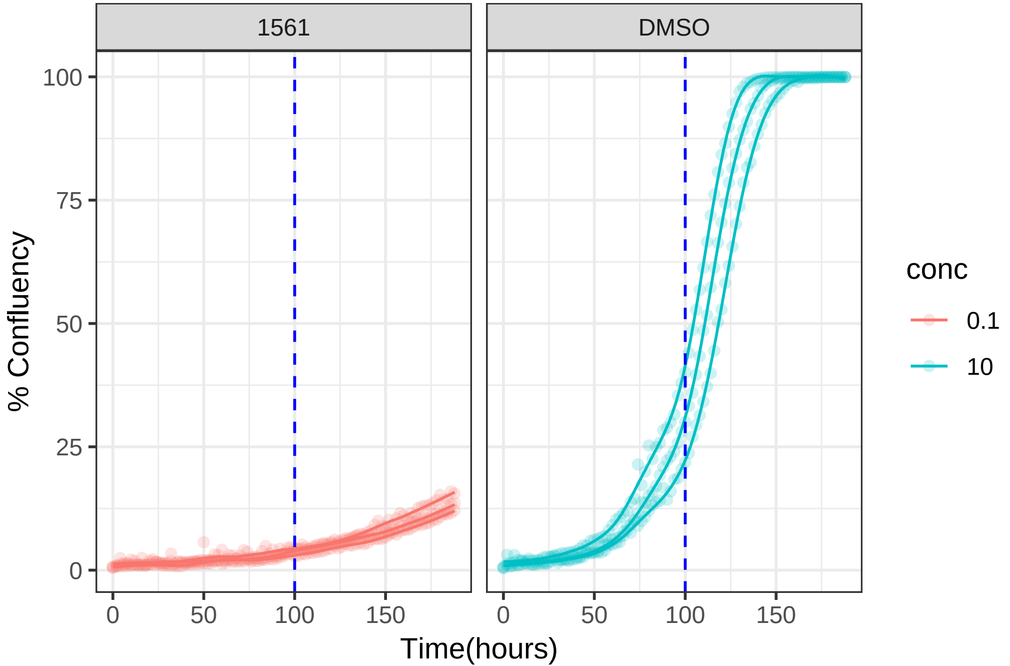

**0.01 uM 1561 RPE1 Ts12 Ts5**

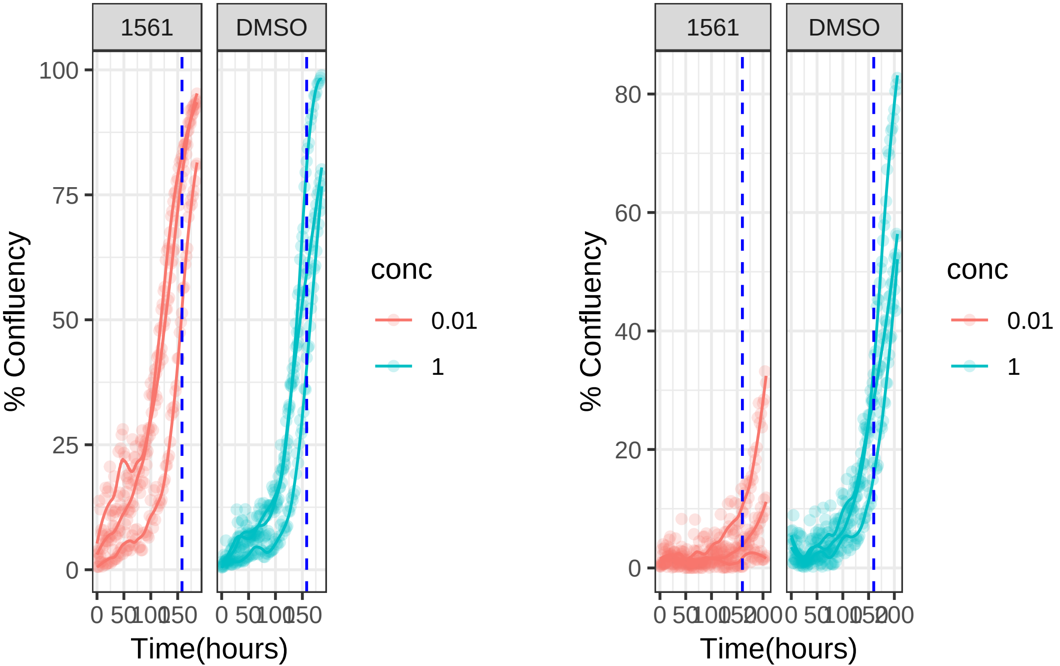

**0.1 uM 1561 RPE1 Ts12 Ts5**

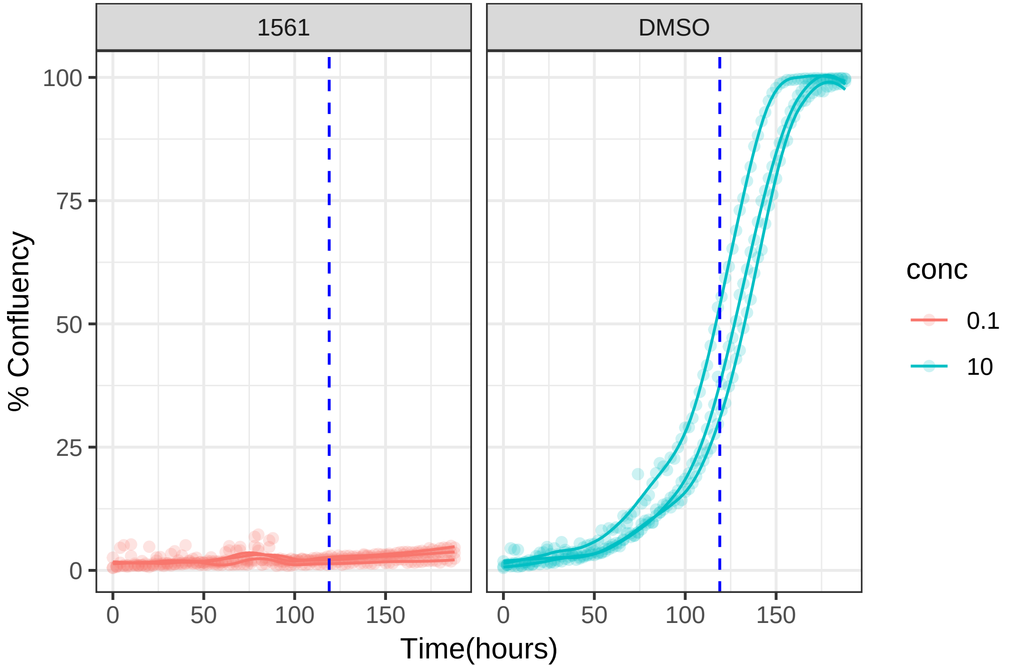

**10 uM 1566 RPE1 hTert**

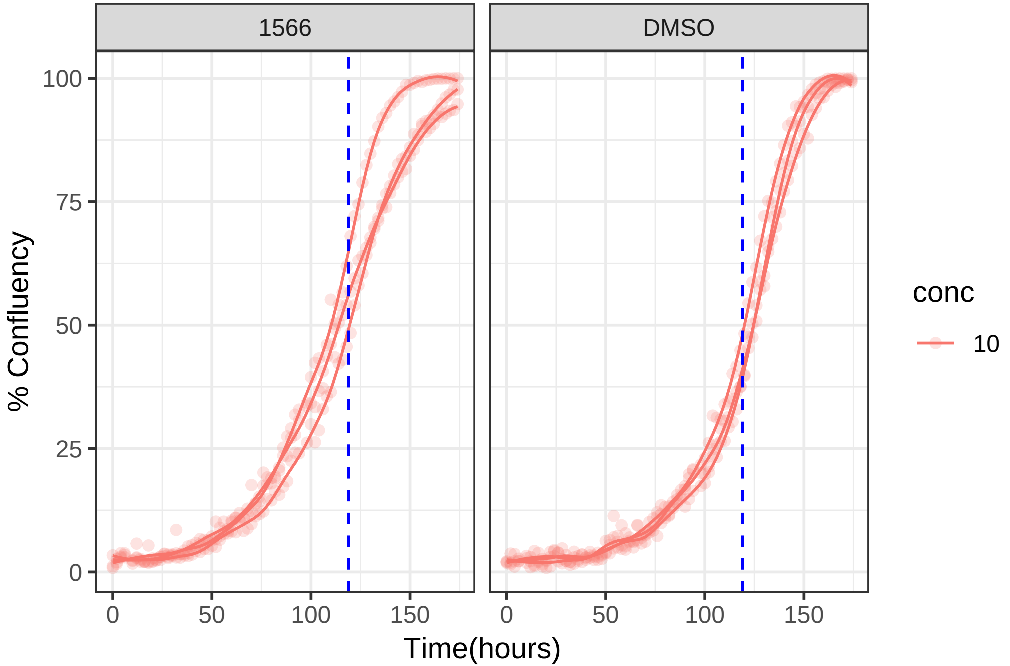

**10 uM 1566 RPE1 Ts12 Ts5**

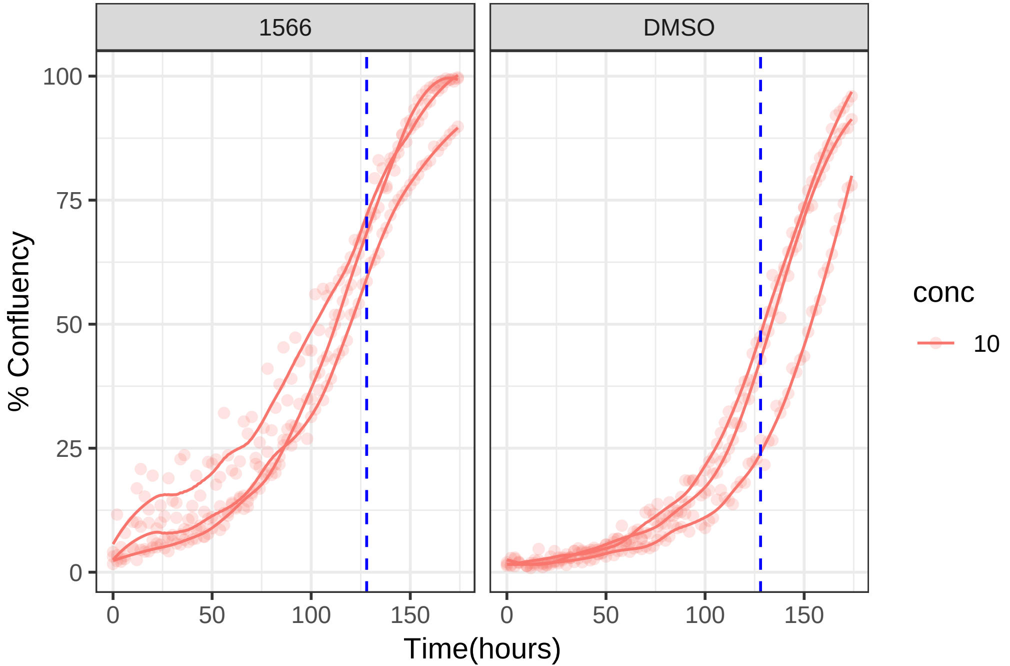

**0.1 uM 1596 RPE1 hTert**

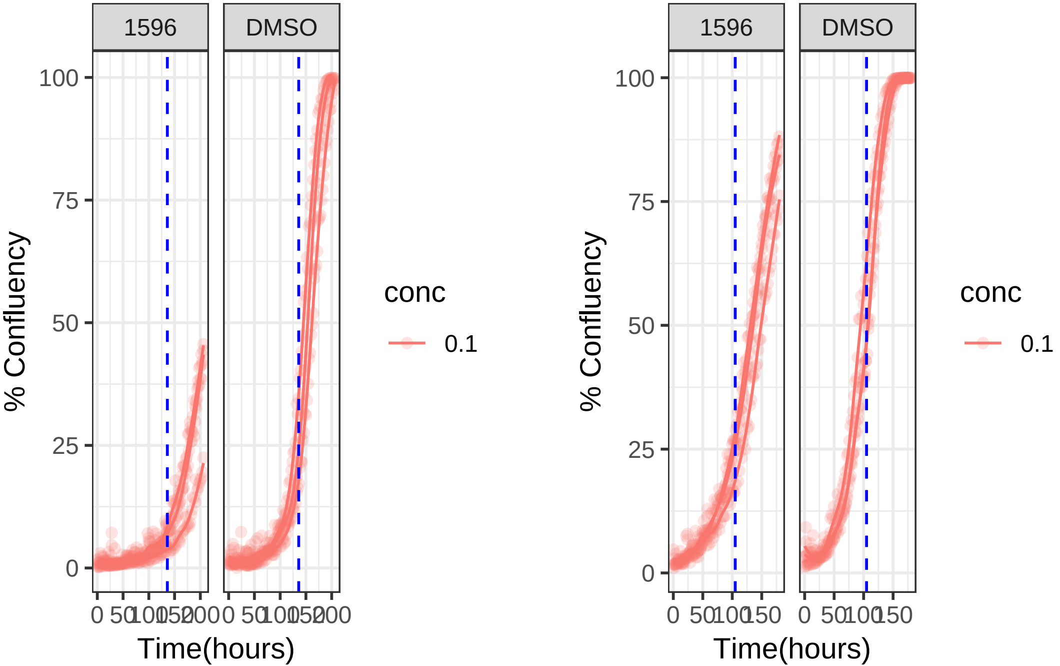

**0.1 uM 1596 RPE1 Ts12 Ts5**

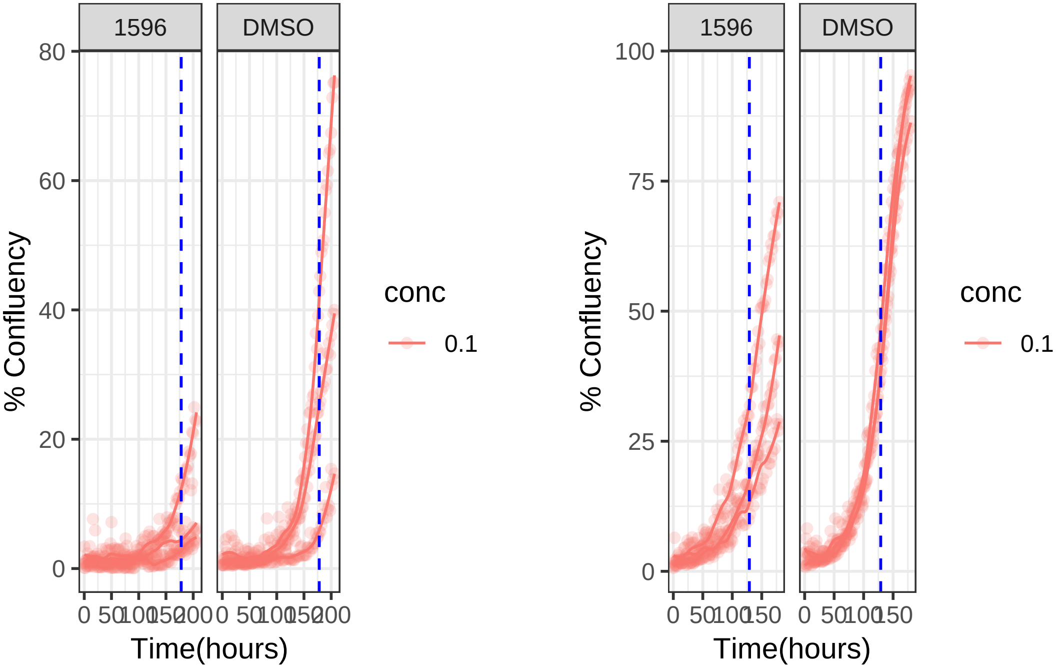

**0.1 uM 1629 RPE1 hTert**

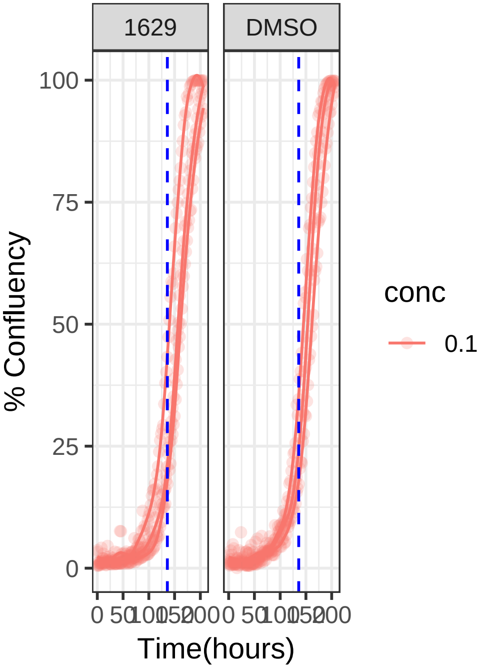


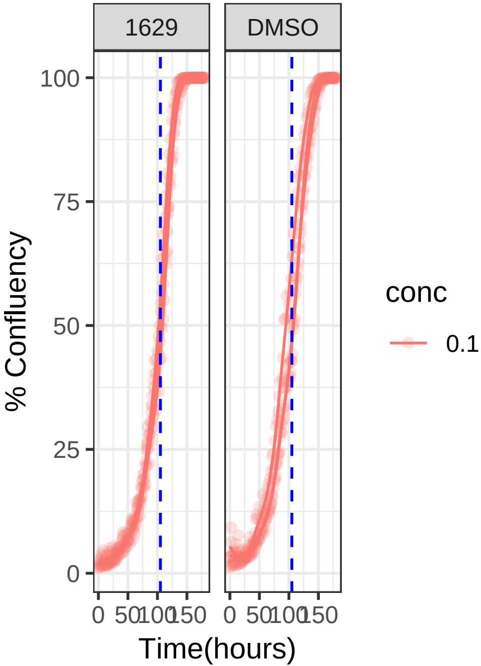

**0.1 uM 1629 RPE1 Ts12 Ts5**

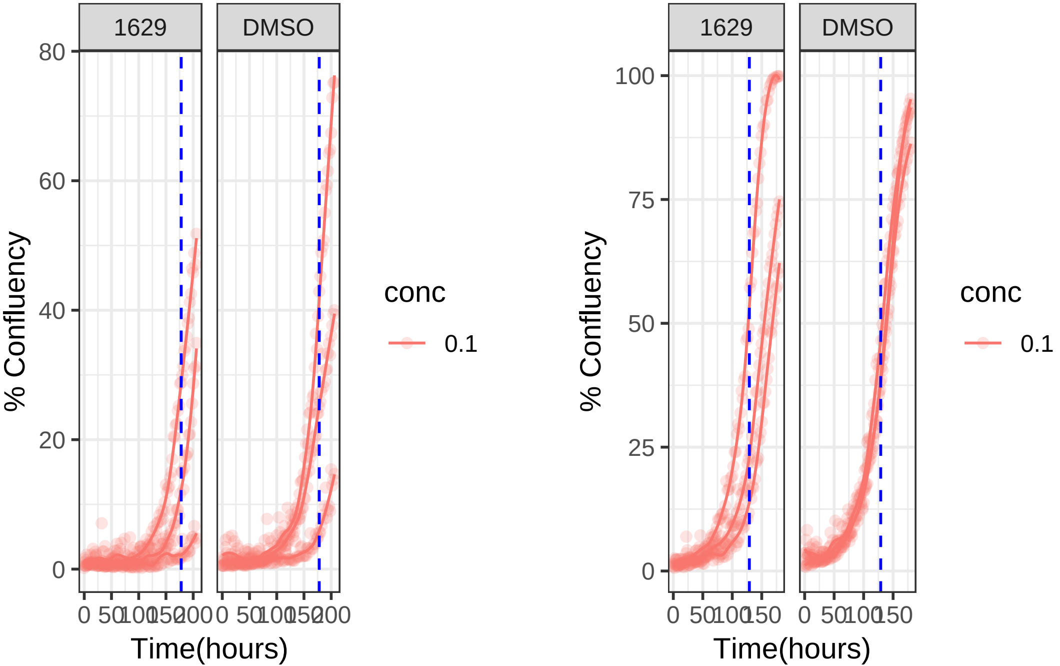

**1 uM 1645 RPE1 hTert**

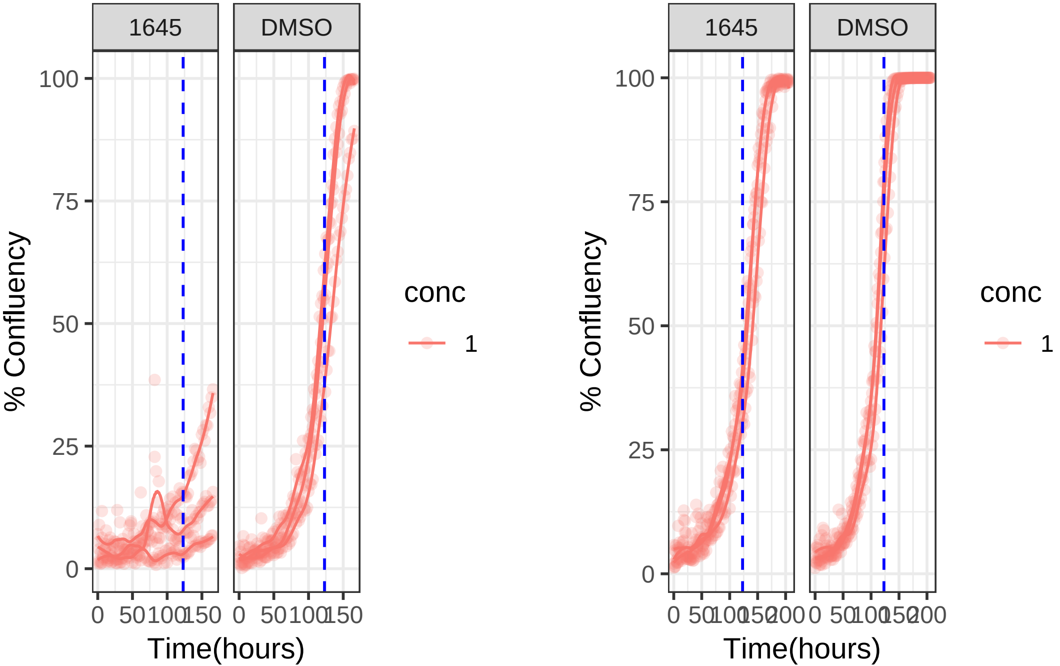

**0.1 uM 1645 RPE1 hTert**

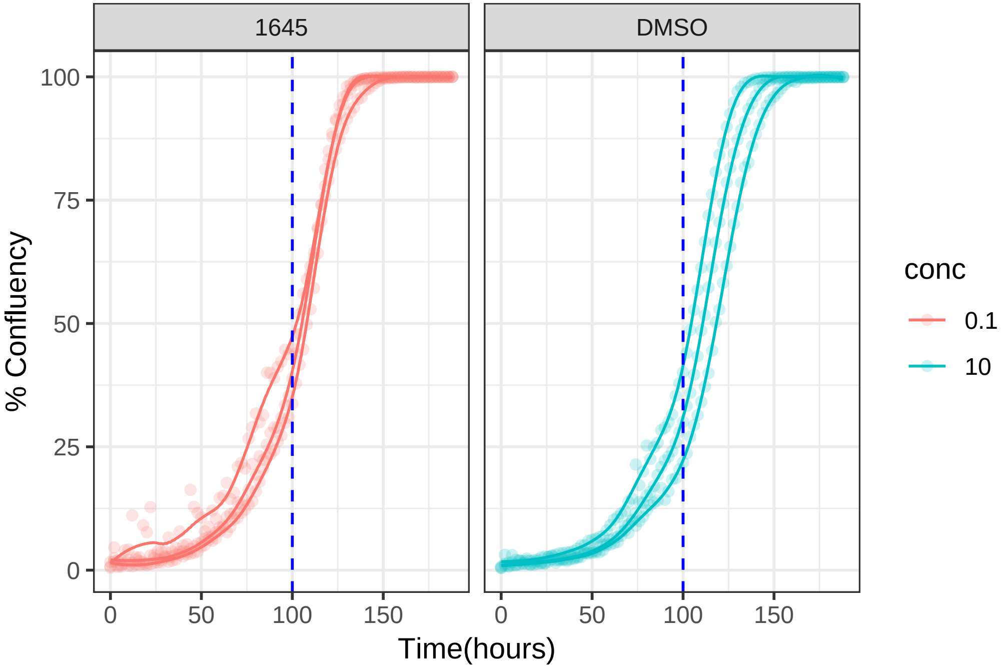

**1 uM 1645 RPE1 Ts12 Ts5**

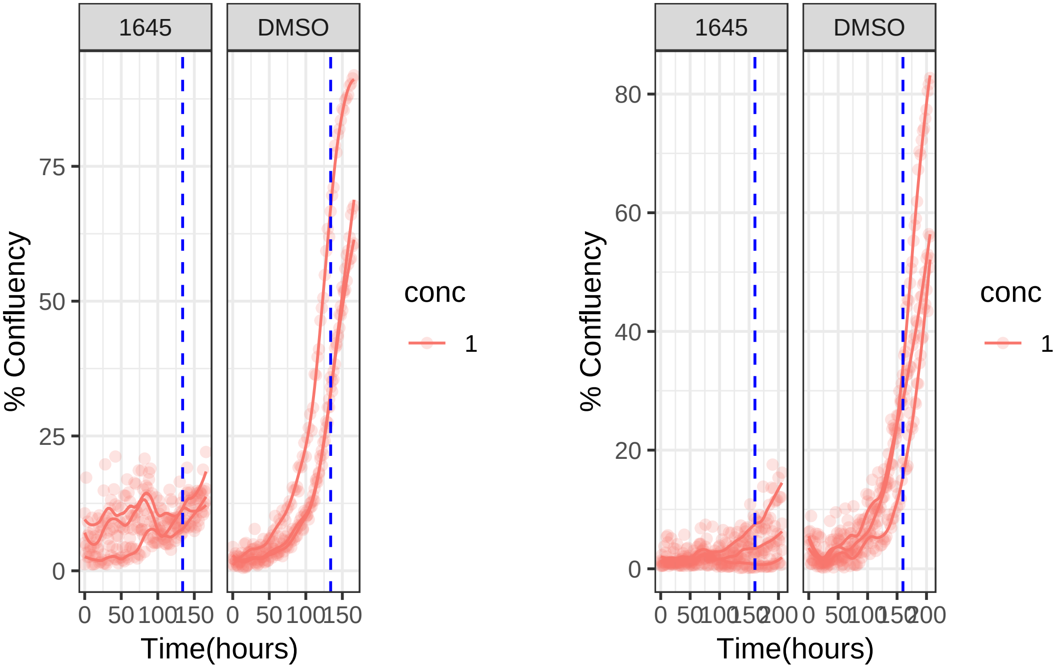

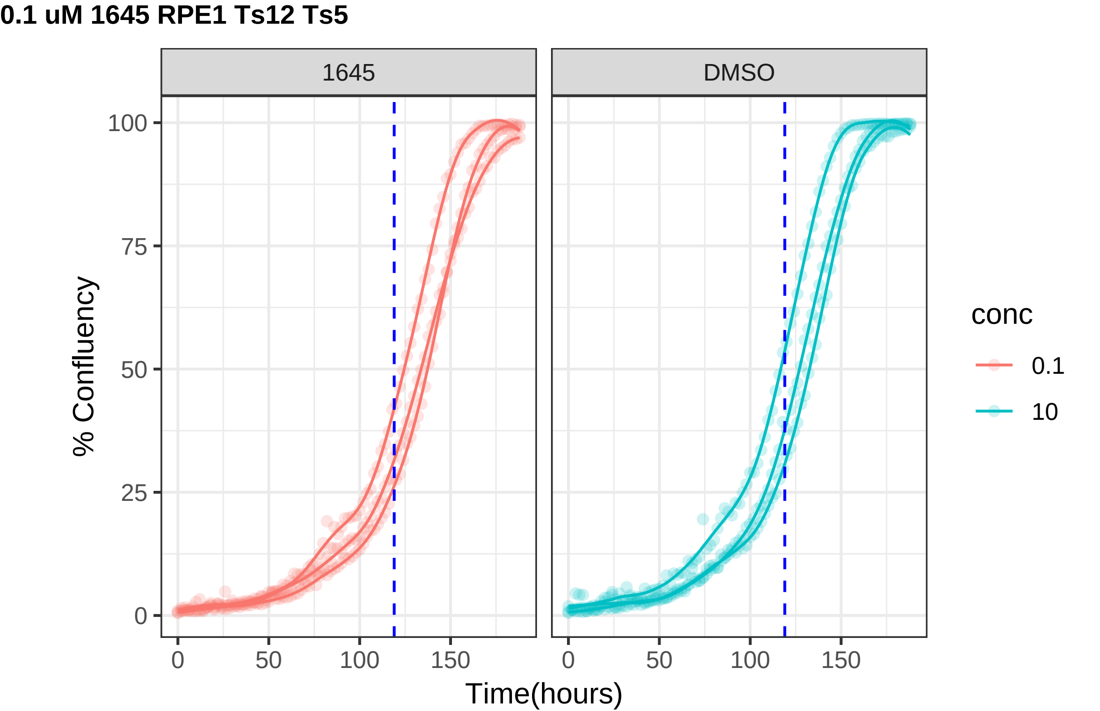

0.1 uM 1645 RPE1 Ts12 Ts5
1645
DMSO
% Confluency
100
75
50
25
0
0
50
100
150
Time(hours)
conc
0.1
10

**0.1 uM 1687 RPE1 hTert**

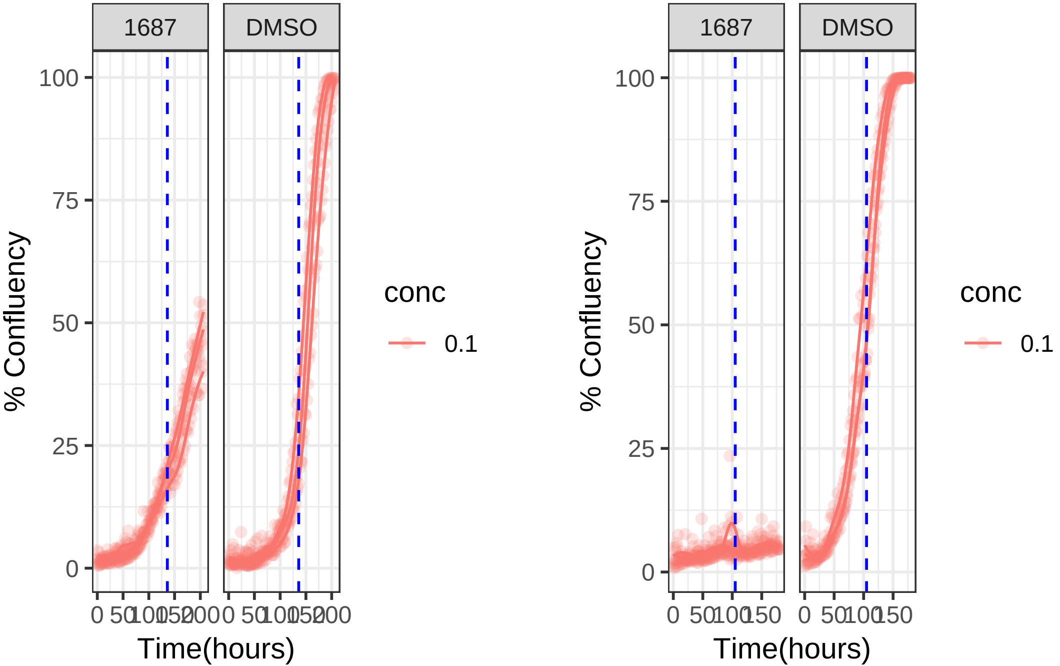

**0.01 uM 1687 RPE1 hTert**

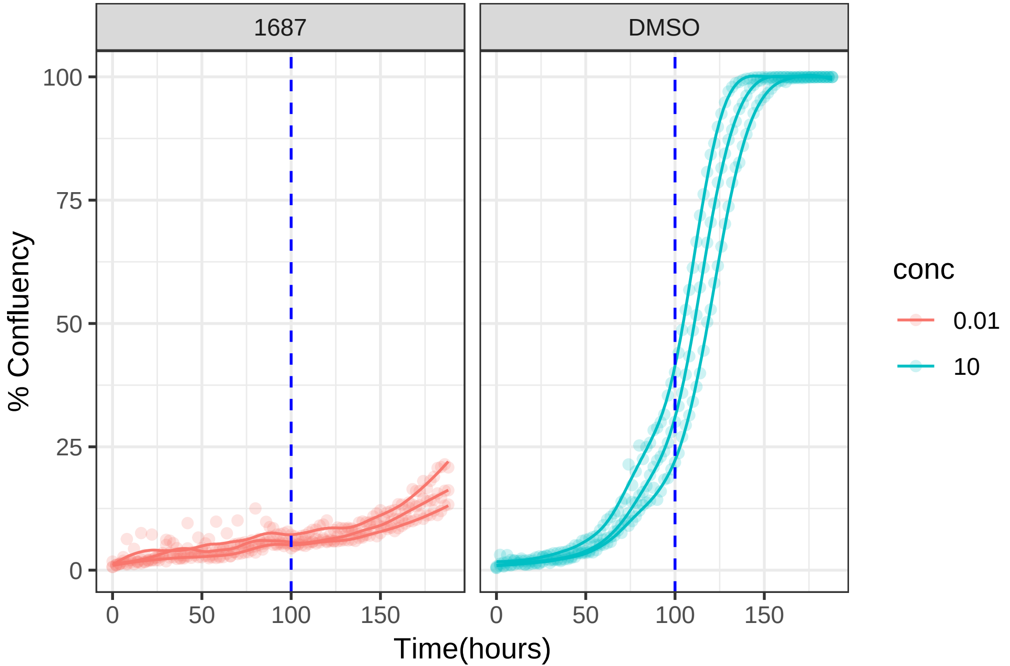

**0.1 uM 1687 RPE1 Ts12 Ts5**

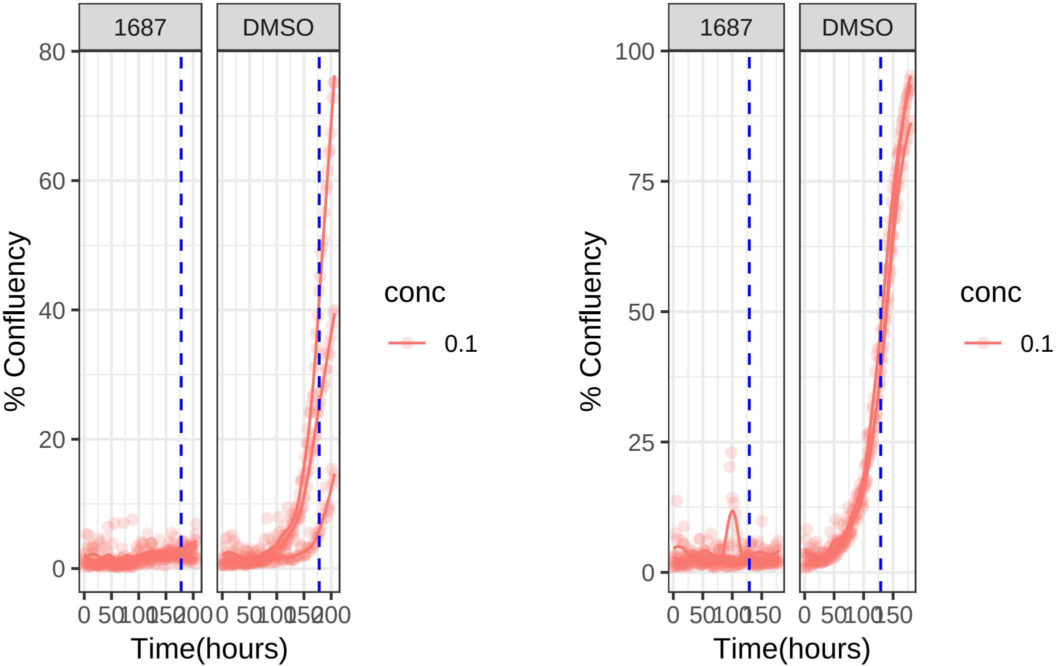

**0.01 uM 1687 RPE1 Ts12 Ts5**

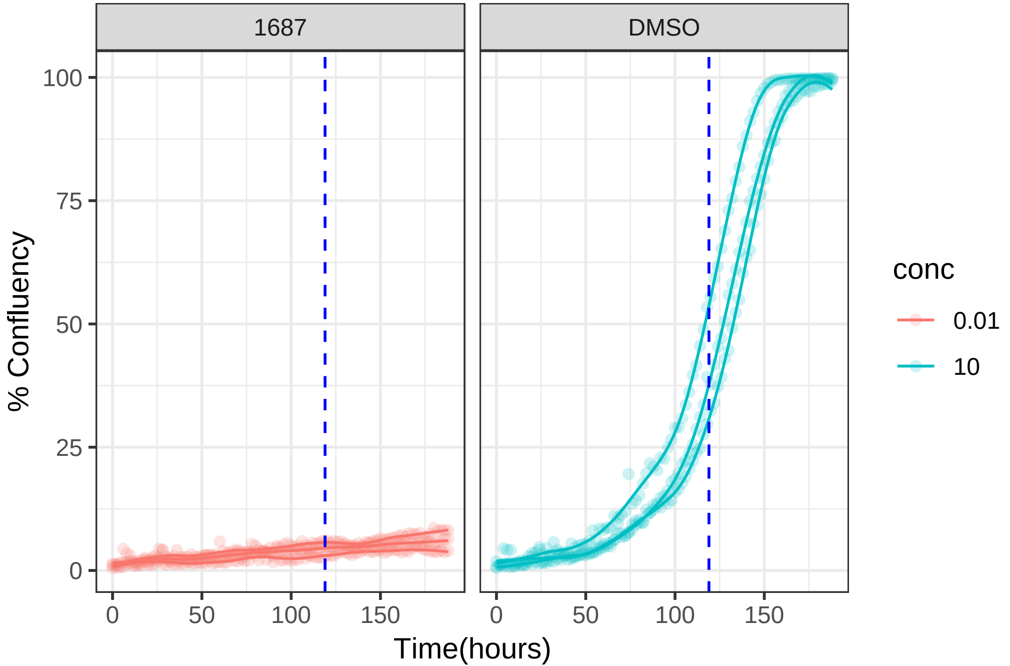

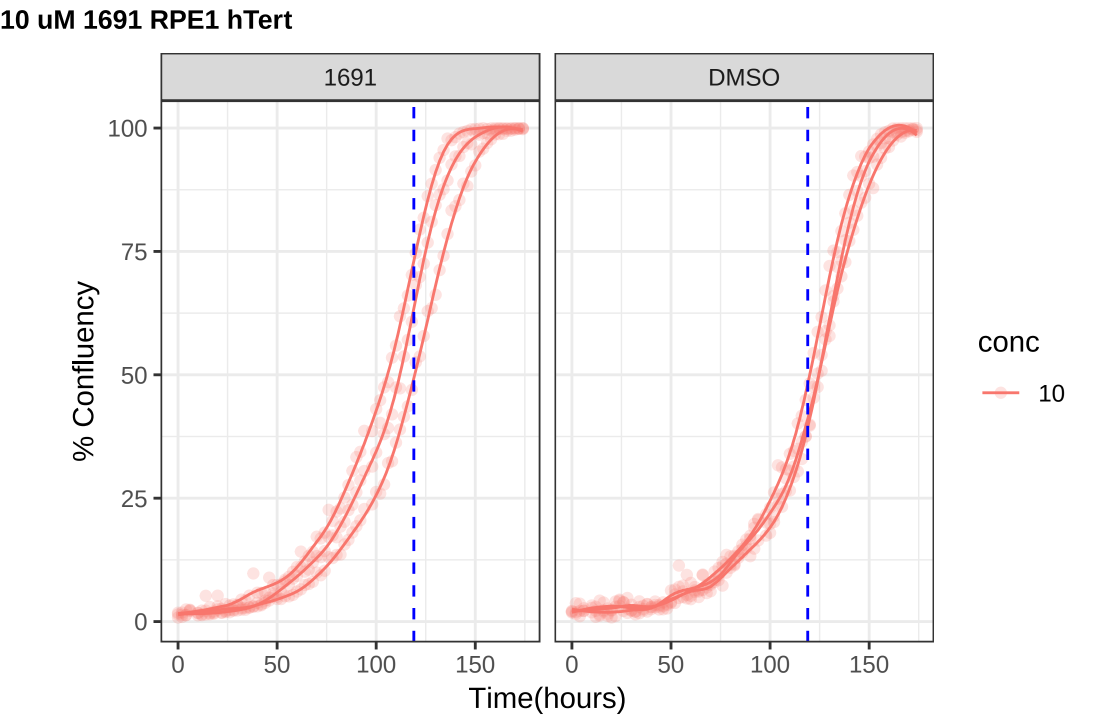
10 uM 1691 RPE1 hTert
1691
DMSO
% Confluency
0
25
50
75
100
0
50
100
150
Time(hours)
conc
10

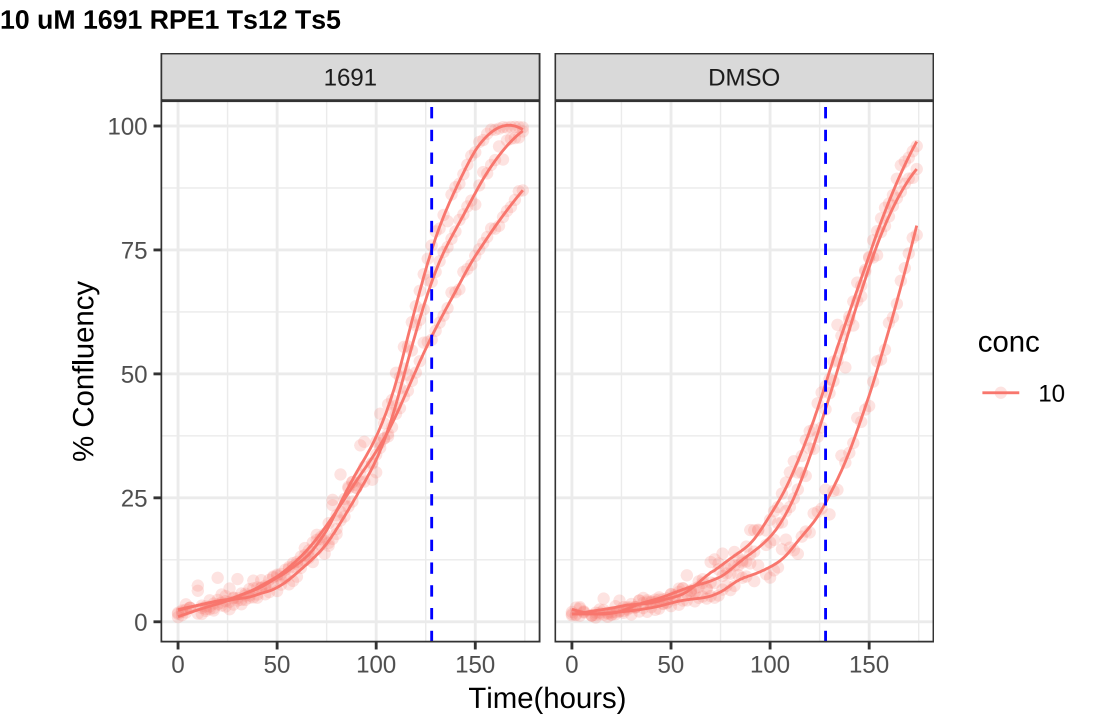
10 uM 1691 RPE1 Ts12 Ts5
1691
DMSO
% Confluency
Time(hours)
0
25
50
75
100
0
50
100
150
conc
10

**1 uM 1729 RPE1 hTert**

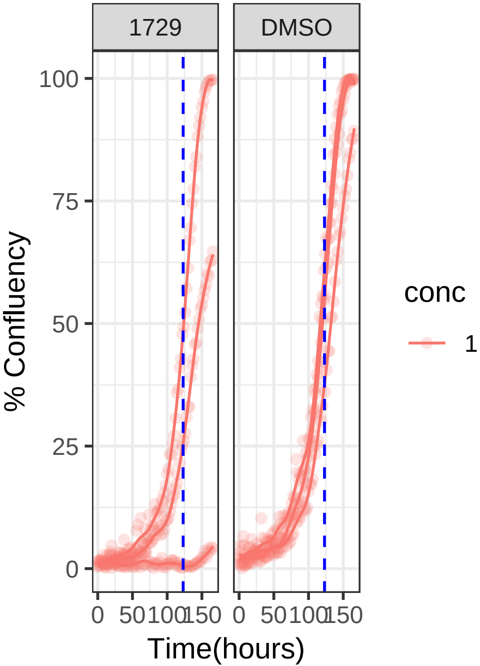


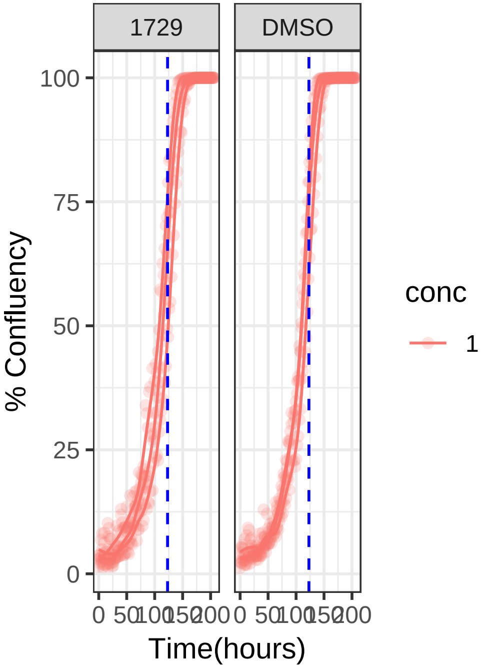

**1 uM 1729 RPE1 Ts12 Ts5**

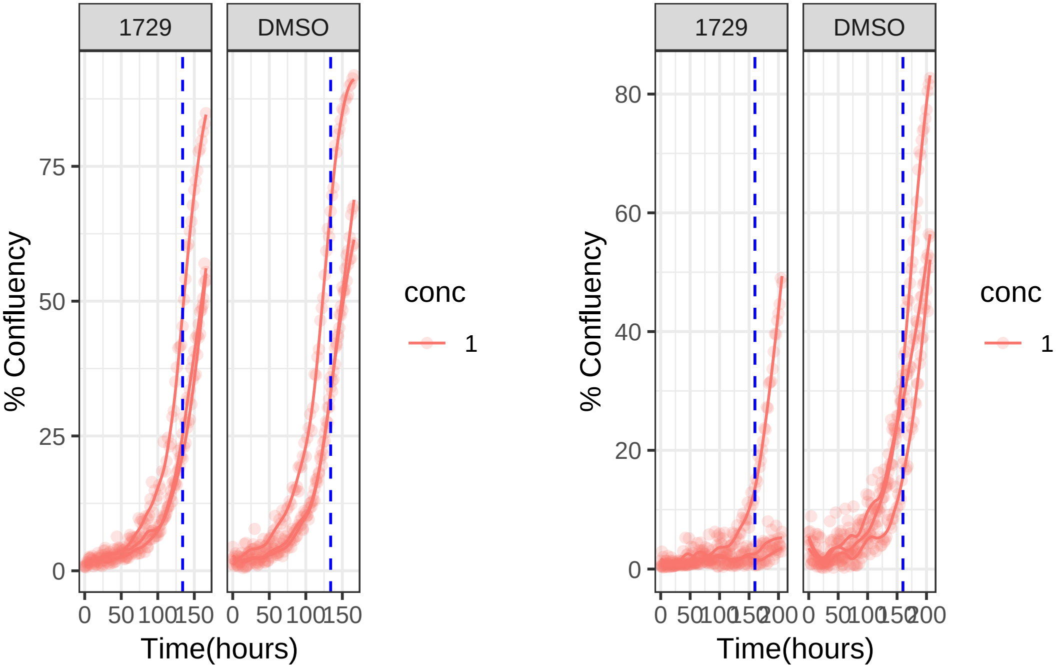

1 uM 1743 RPE1 hTert

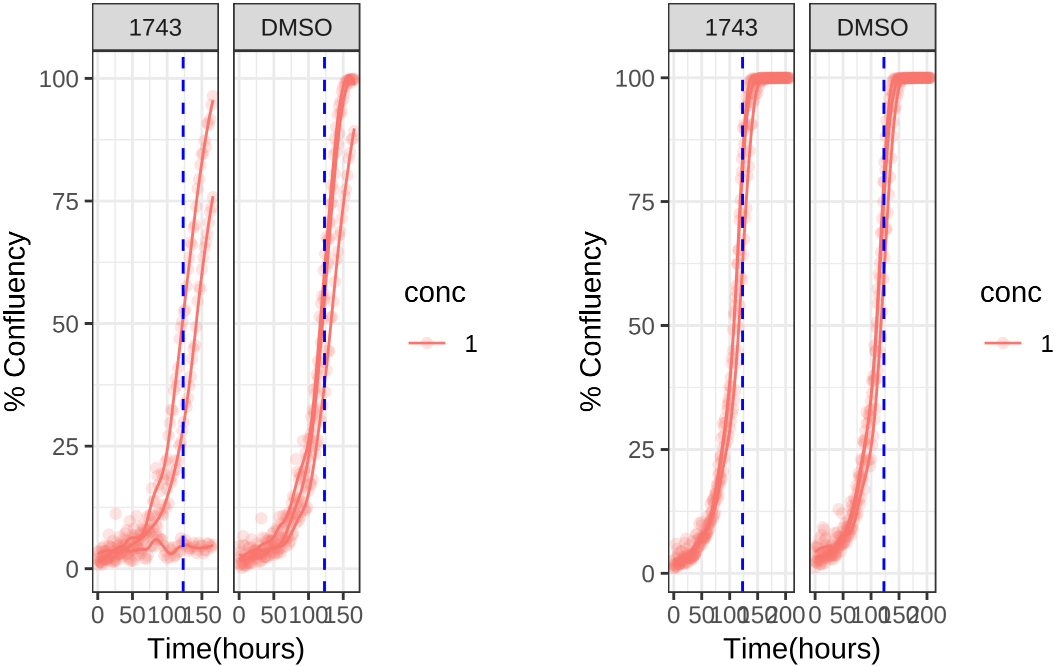

1 uM 1743 RPE1 Ts12 Ts5

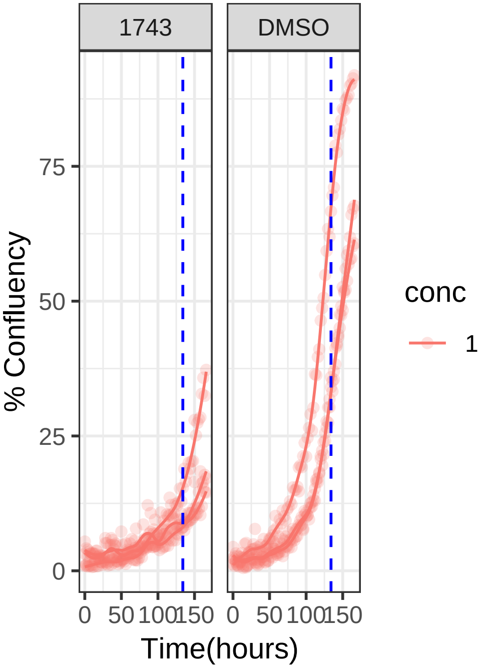


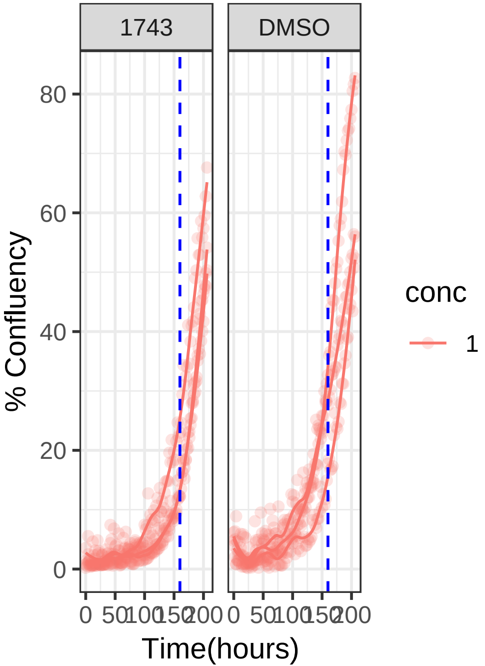

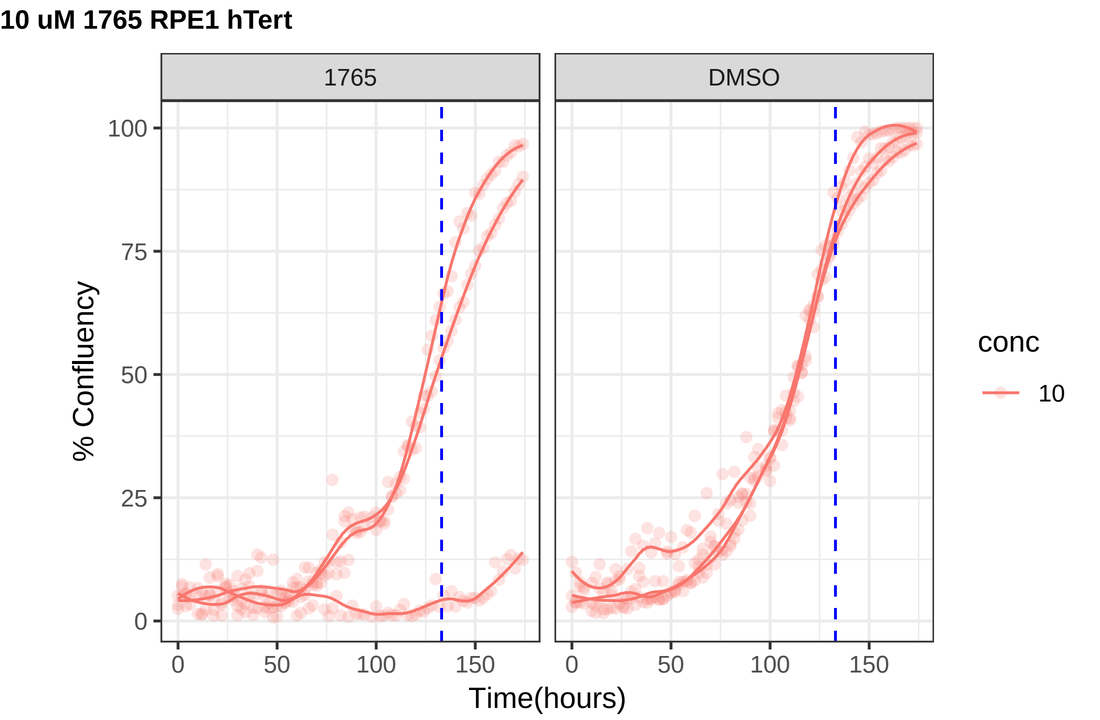
10 uM 1765 RPE1 hTert
1765
DMSO
% Confluency
100
75
50
25
0
0
50
100
150
Time(hours)
conc
10

**10 uM 1765 RPE1 Ts12 Ts5**

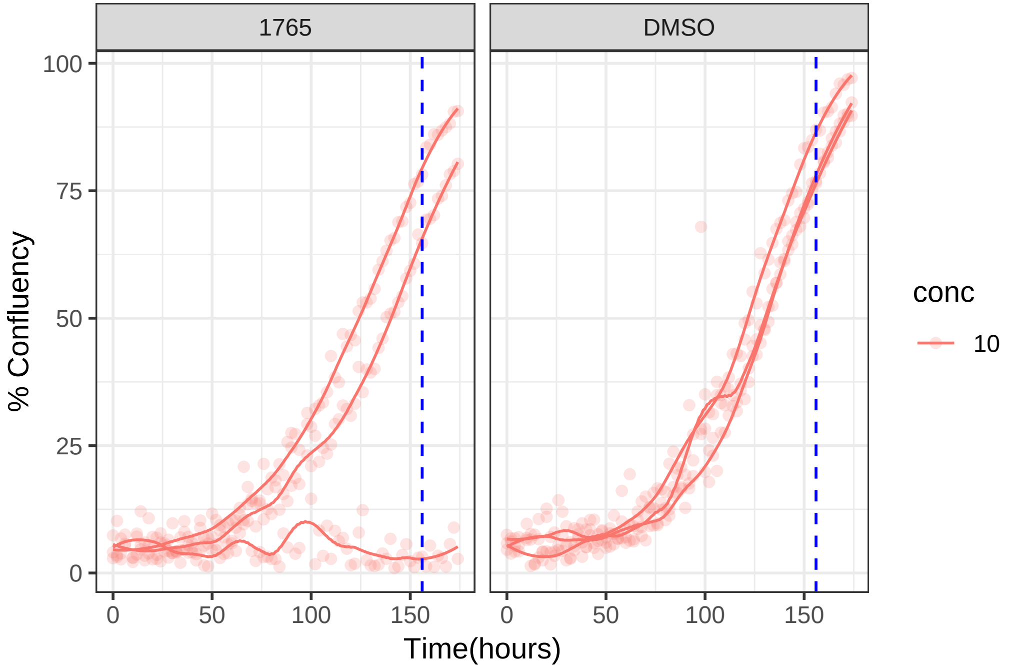

**10 uM 1789 RPE1 hTert**

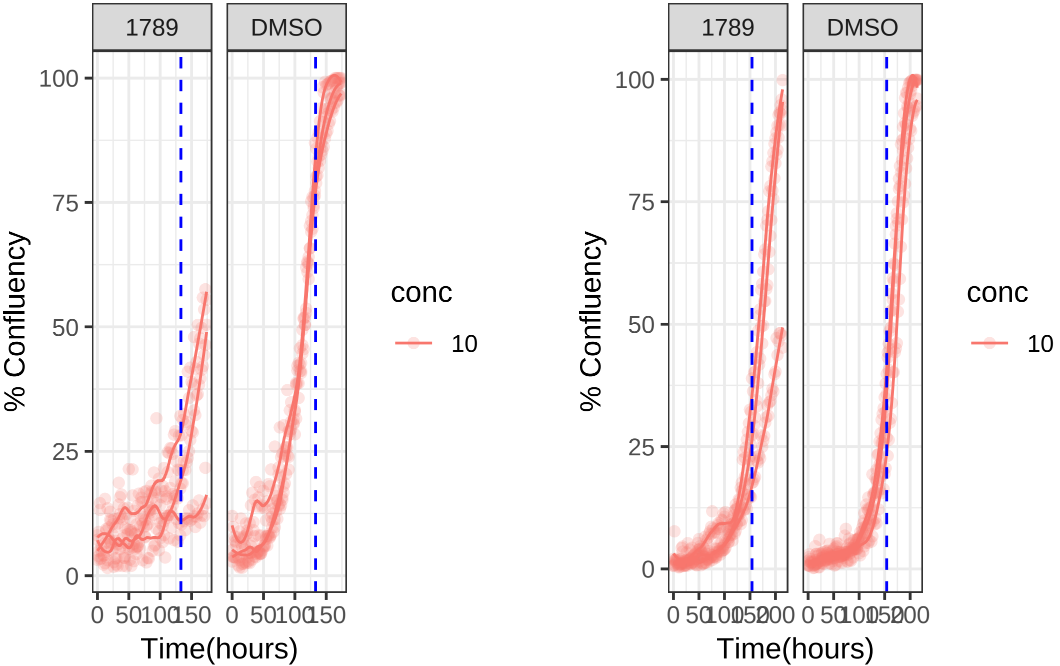

**10 uM 1789 RPE1 Ts12 Ts5**

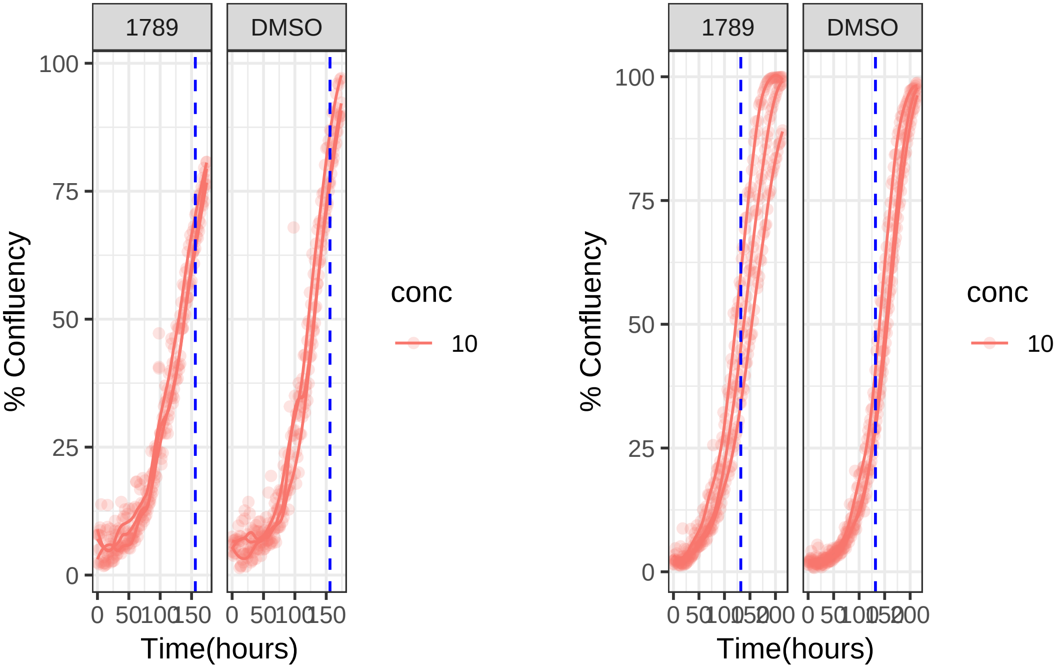

**1 uM 1797 RPE1 hTert**

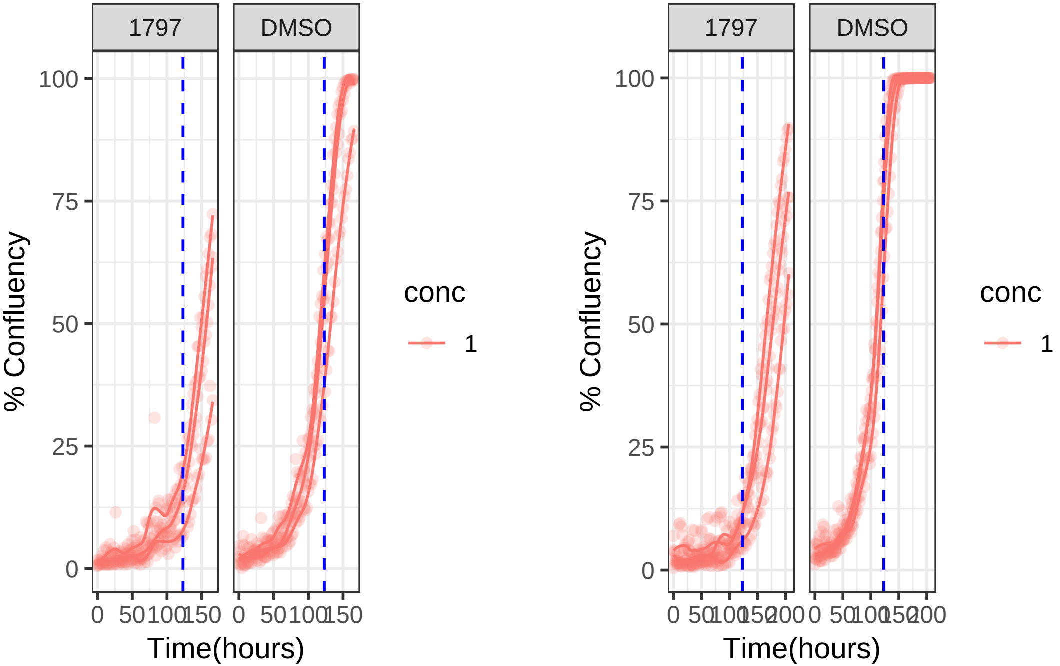

**1 uM 1797 RPE1 Ts12 Ts5**

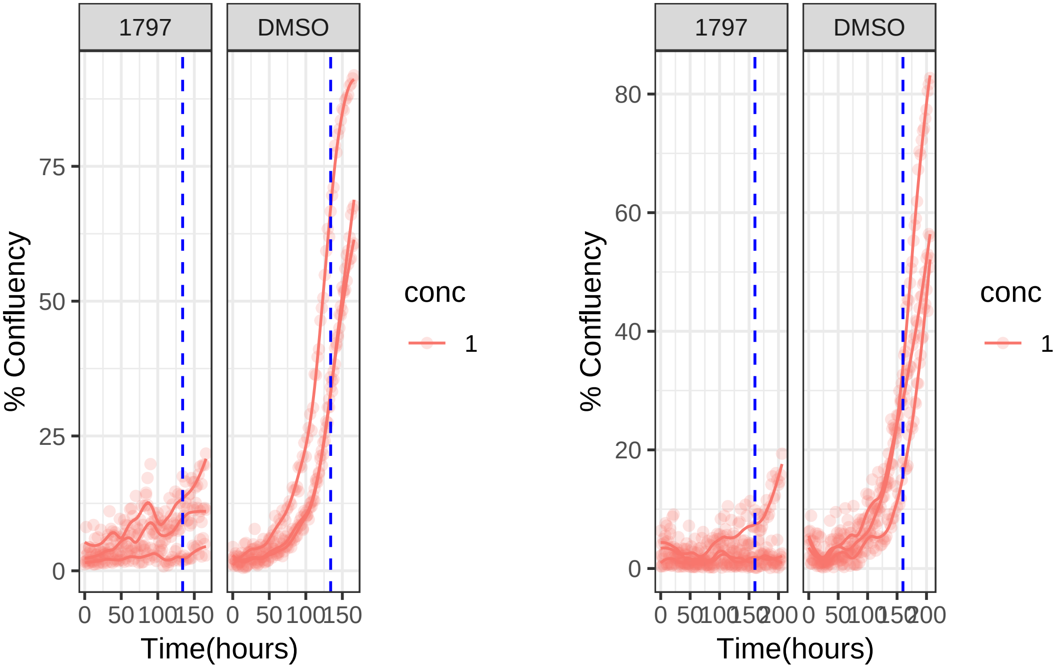

**1 uM 1801 RPE1 hTert**

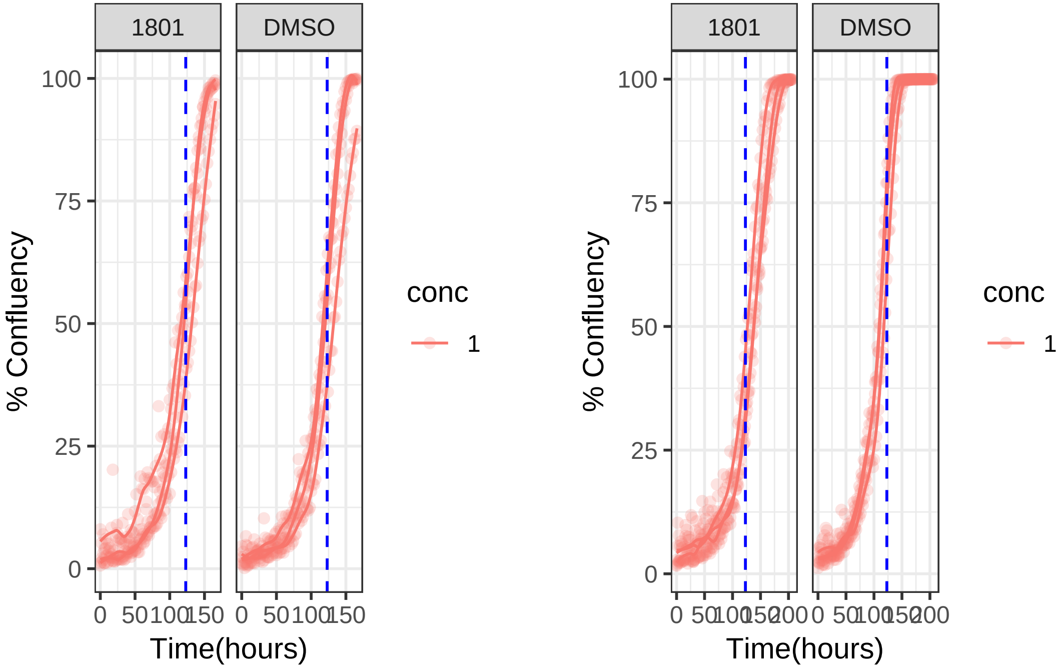

**1 uM 1801 RPE1 Ts12 Ts5**

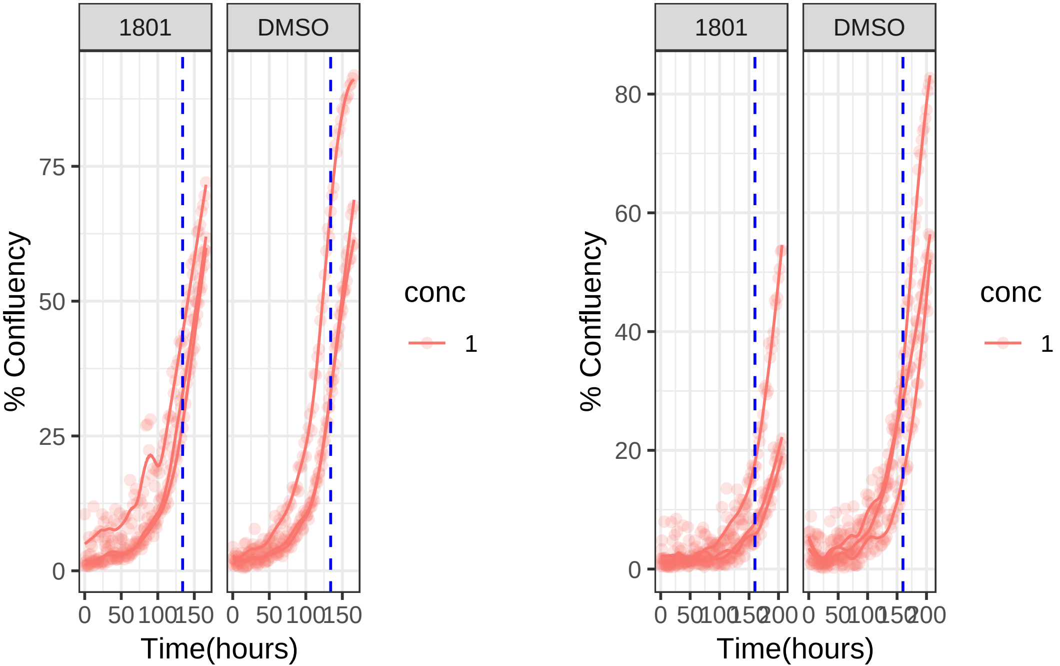

**1 uM 1803 RPE1 hTert**

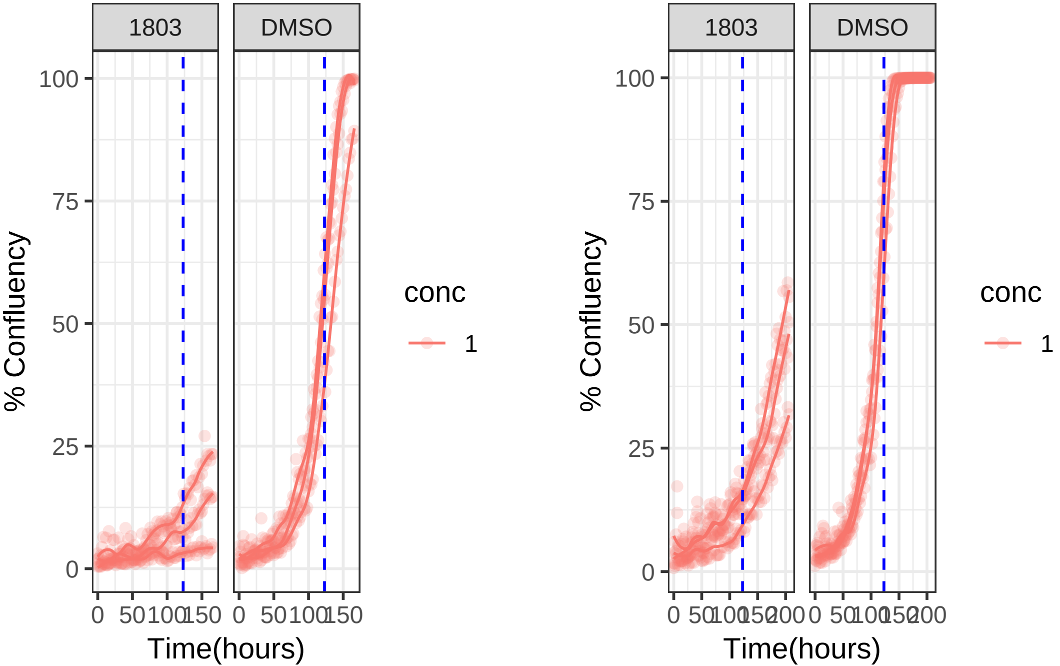

**1 uM 1803 RPE1 Ts12 Ts5**

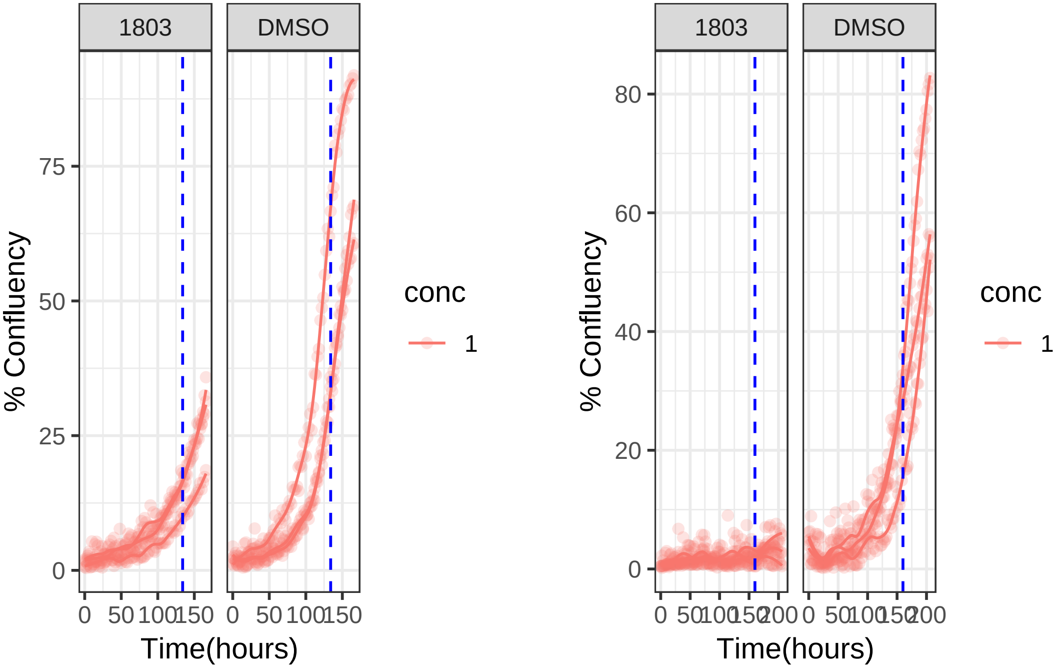

**0.01 uM 1807 RPE1 hTert**

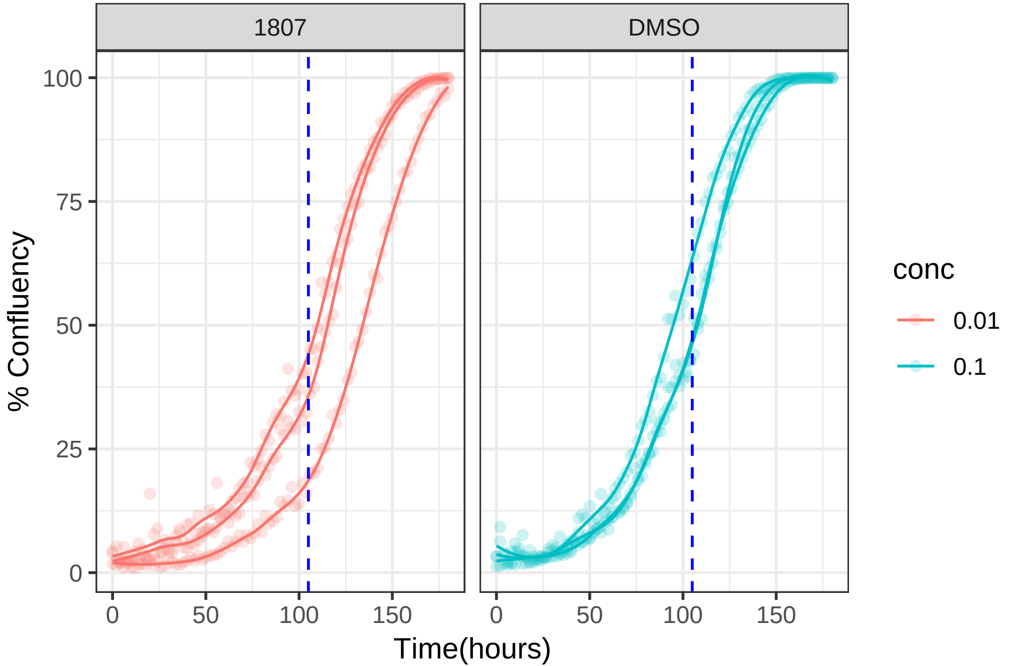

0.01 uM 1807 RPE1 Ts12 Ts5

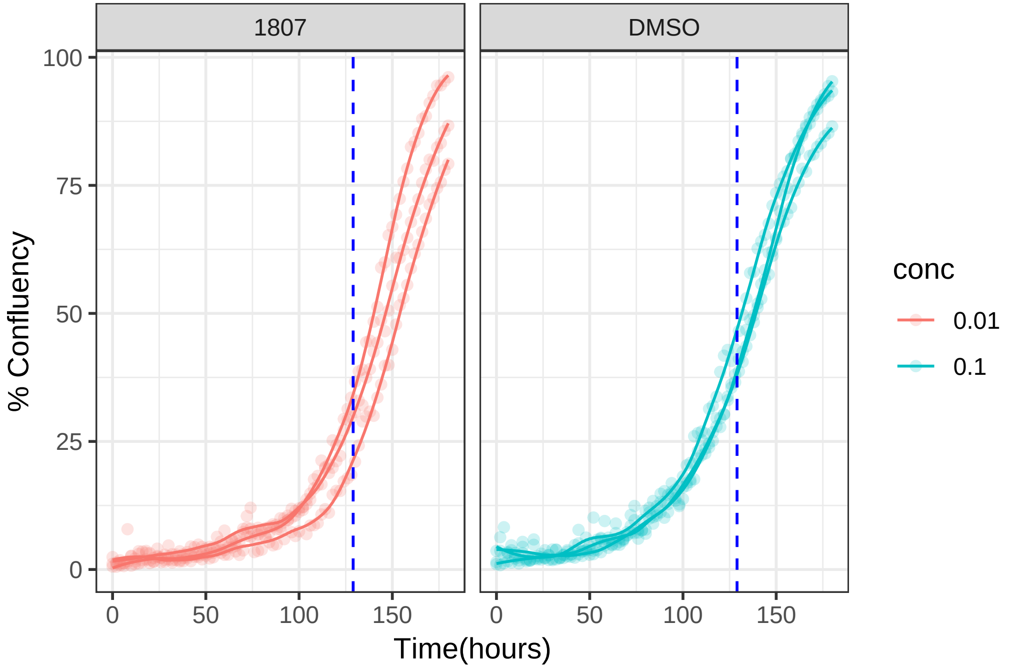

**10 uM 1814 RPE1 hTert**

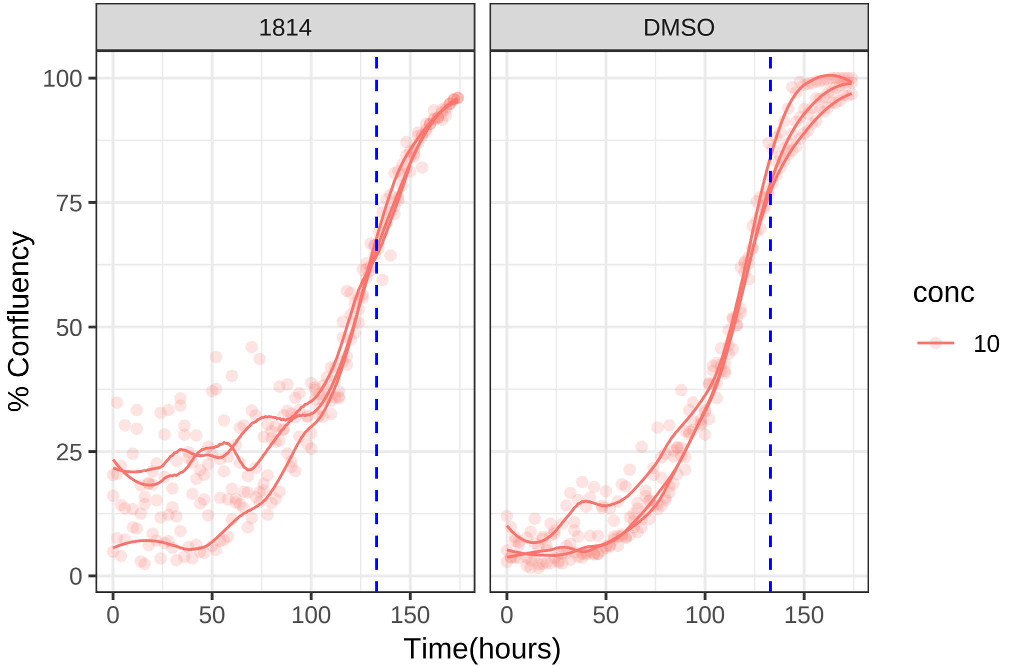

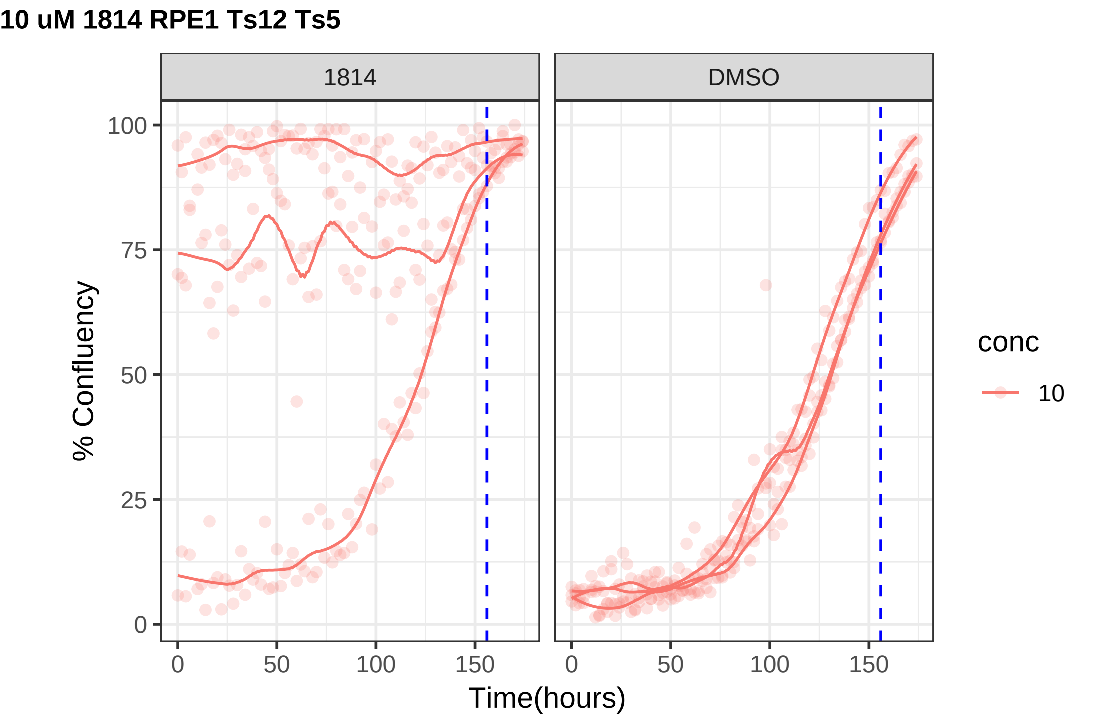
10 uM 1814 RPE1 Ts12 Ts5
1814
DMSO
% Confluency
Time(hours)
conc
10
0
25
50
75
100
0
50
100
150

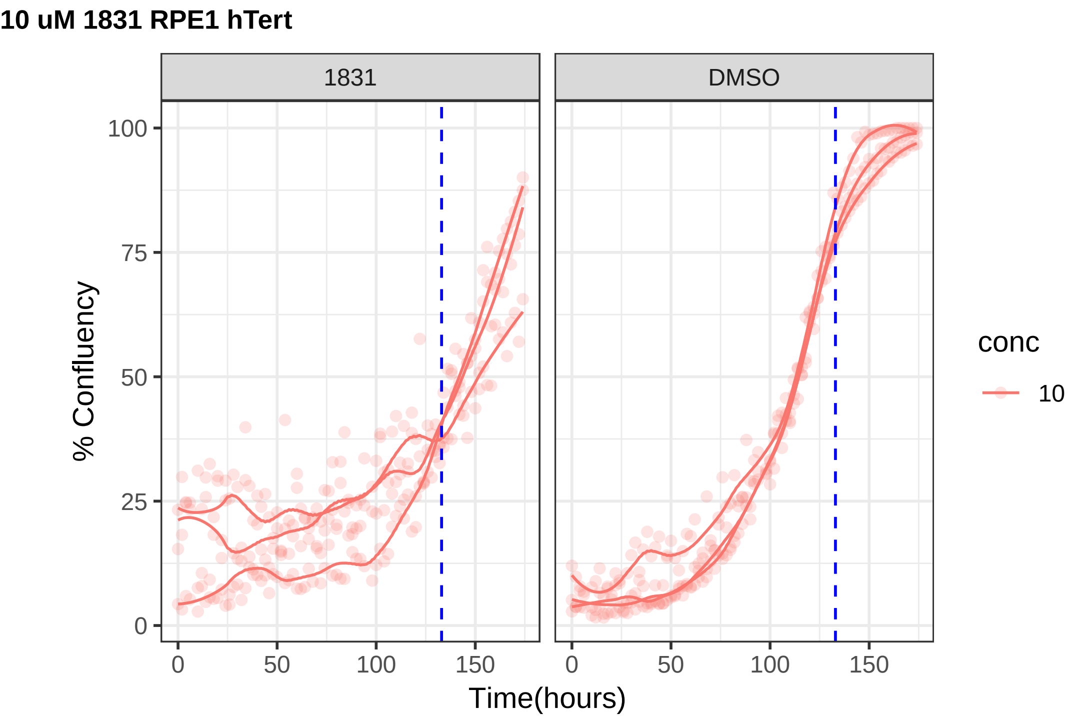
10 uM 1831 RPE1 hTert
1831
DMSO
% Confluency
Time(hours)
0
25
50
75
100
0
50
100
150
conc
10

**10 uM 1831 RPE1 Ts12 Ts5**

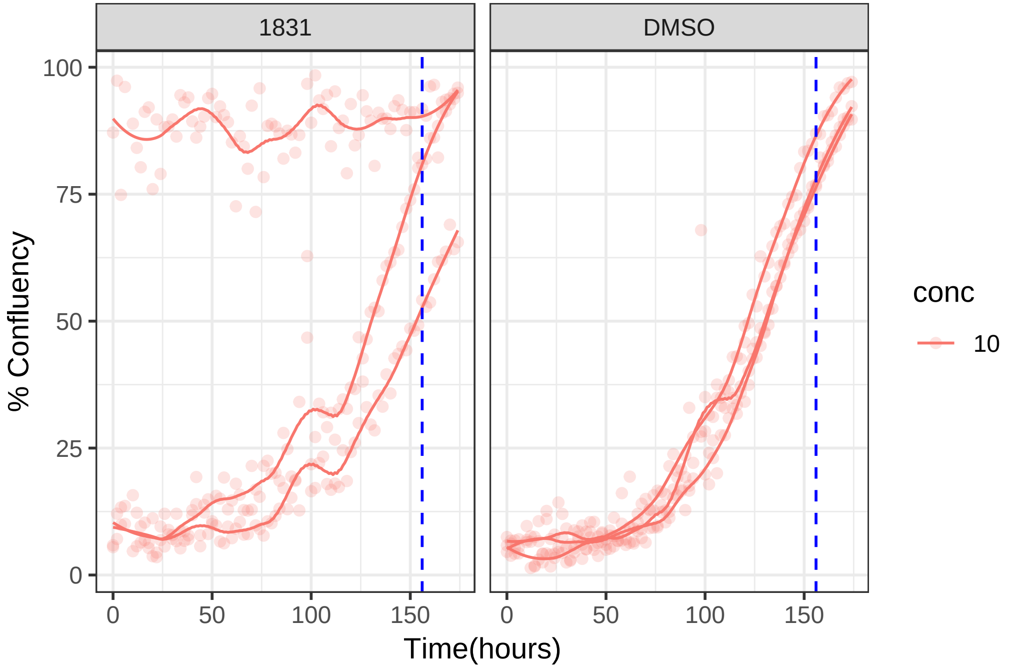

**10 uM 1853 RPE1 hTert**

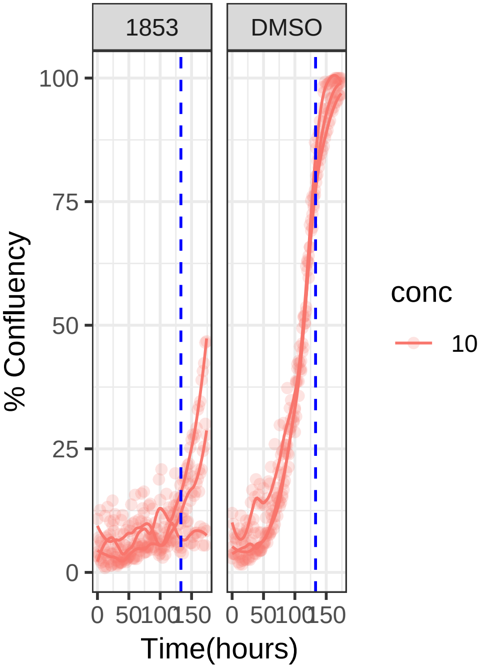


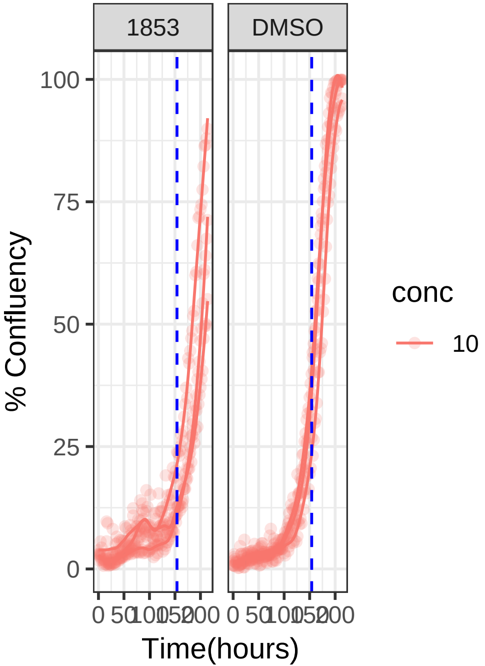

**10 uM 1853 RPE1 Ts12 Ts5**

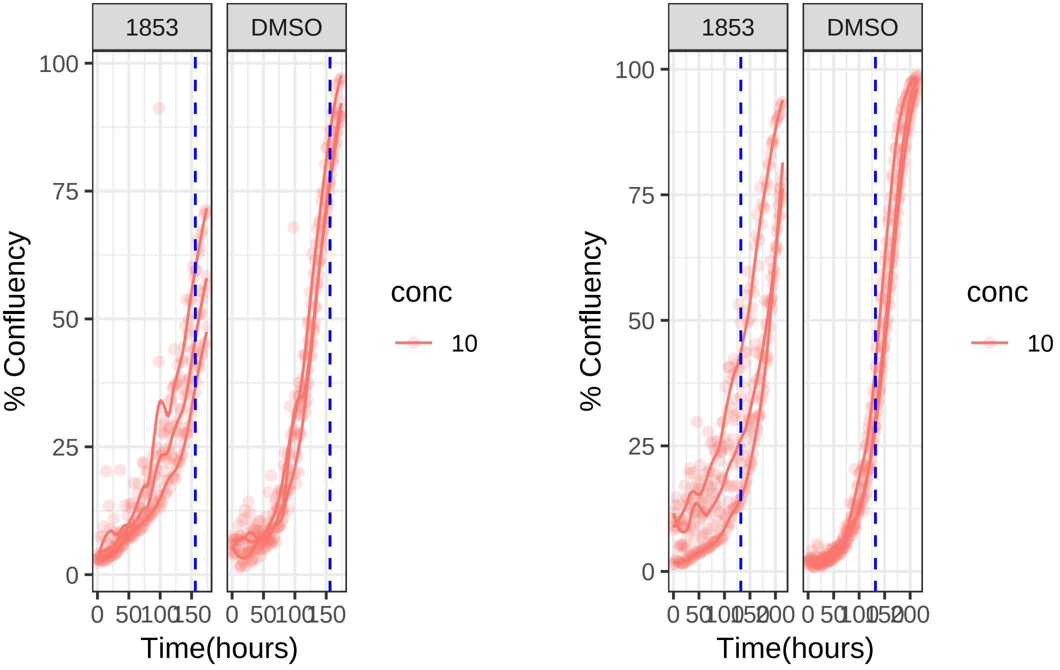

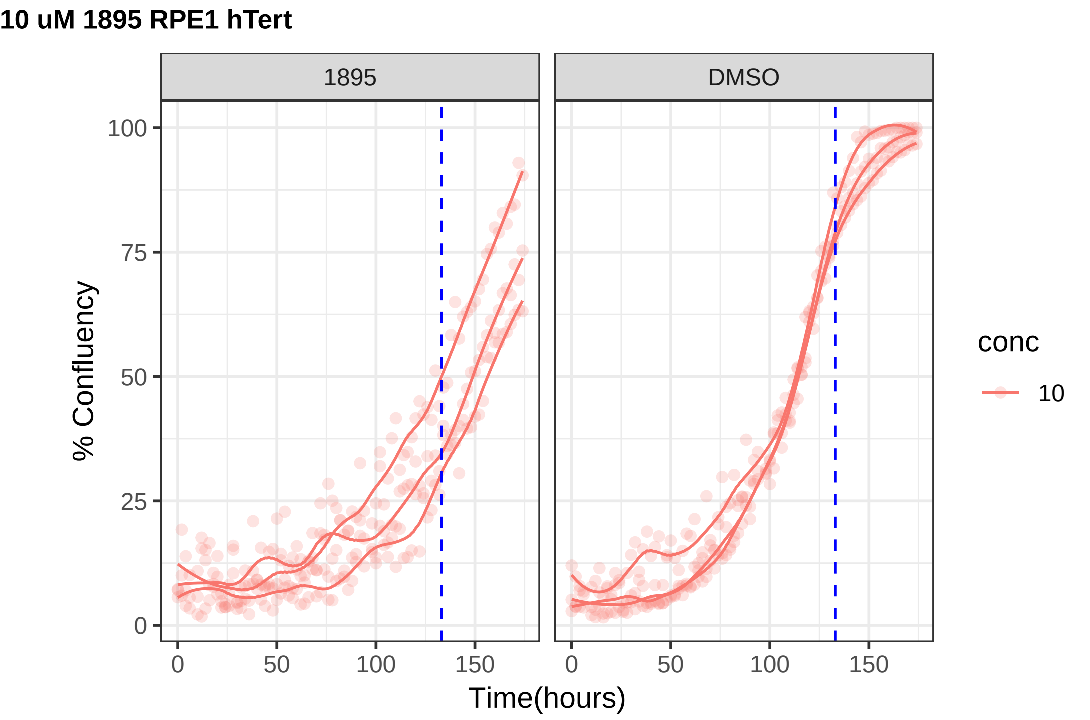

10 uM 1895 RPE1 hTert
1895
DMSO
% Confluency
0
25
50
75
100
0
50
100
150
Time(hours)
conc
10

#### 10 uM 1895 RPE1 Ts12 Ts5

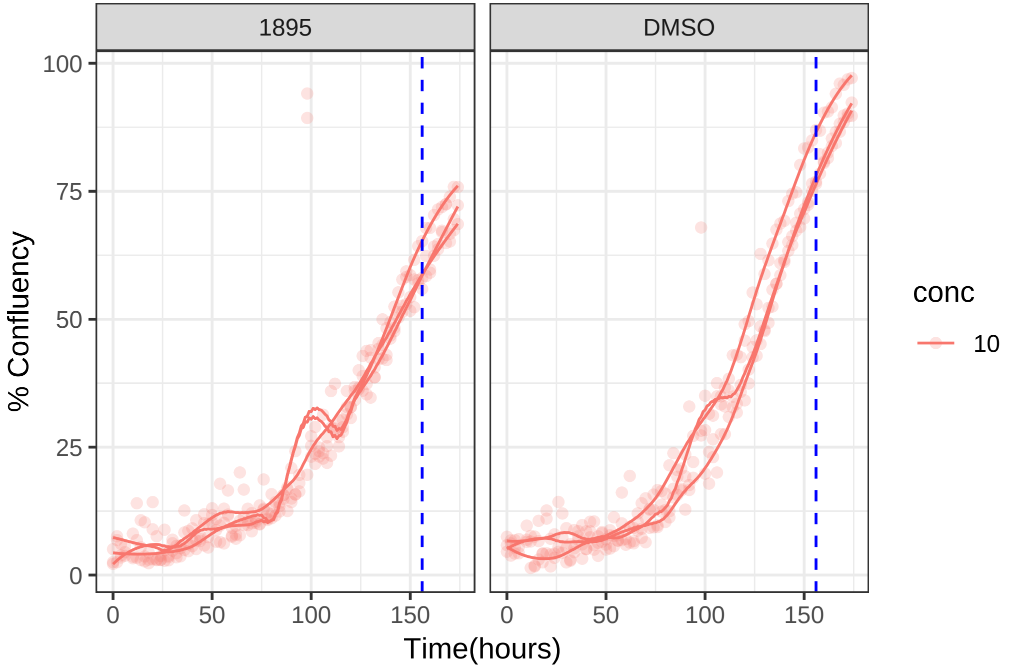

**10 uM 1928 RPE1 hTert**

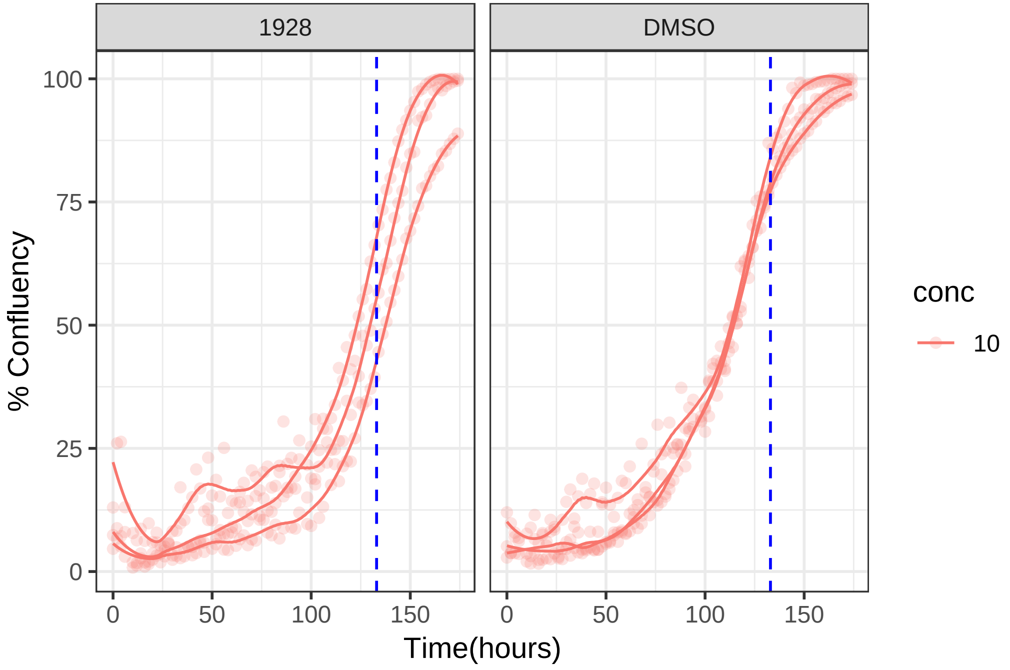

**10 uM 1928 RPE1 Ts12 Ts5**

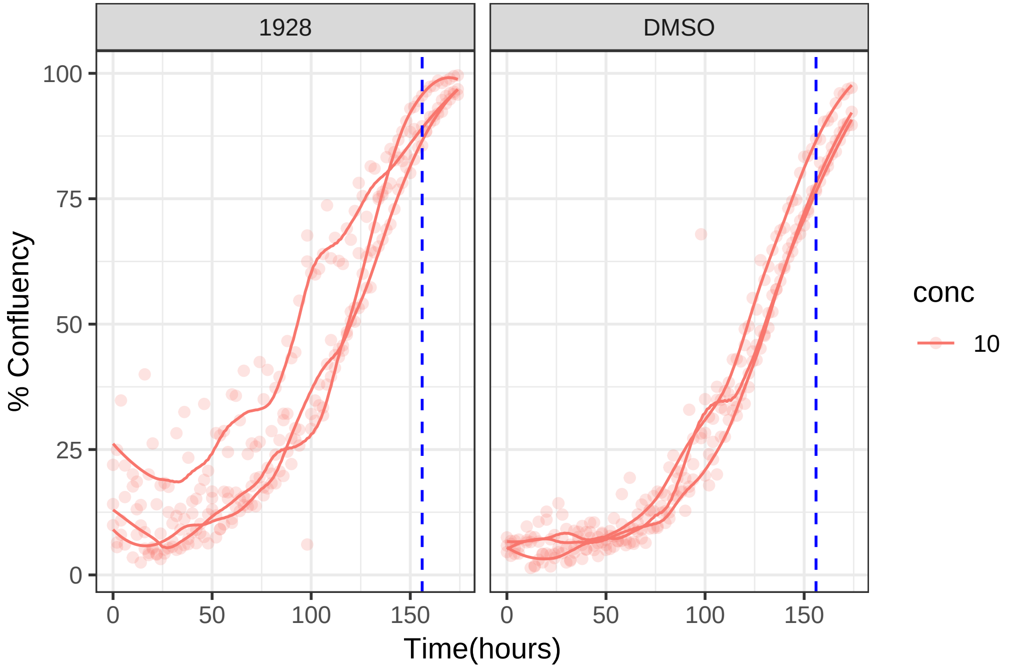

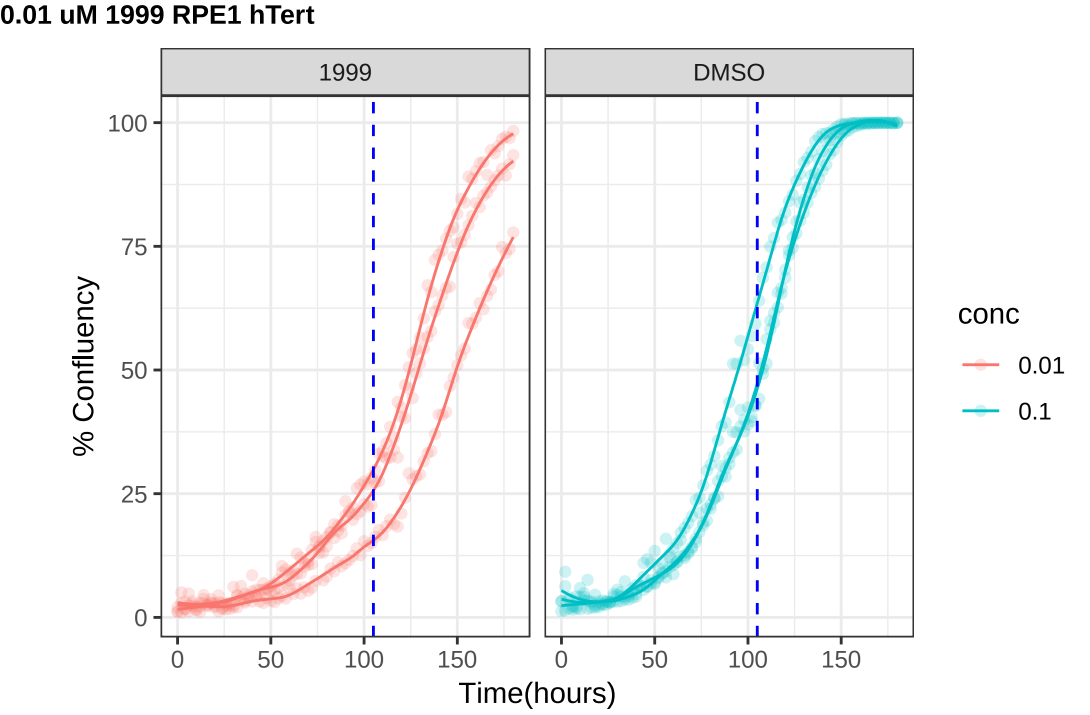
0.01 uM 1999 RPE1 hTert
1999
DMSO
% Confluency
100
75
50
25
0
0
50
100
150
Time(hours)
conc
0.01
0.1

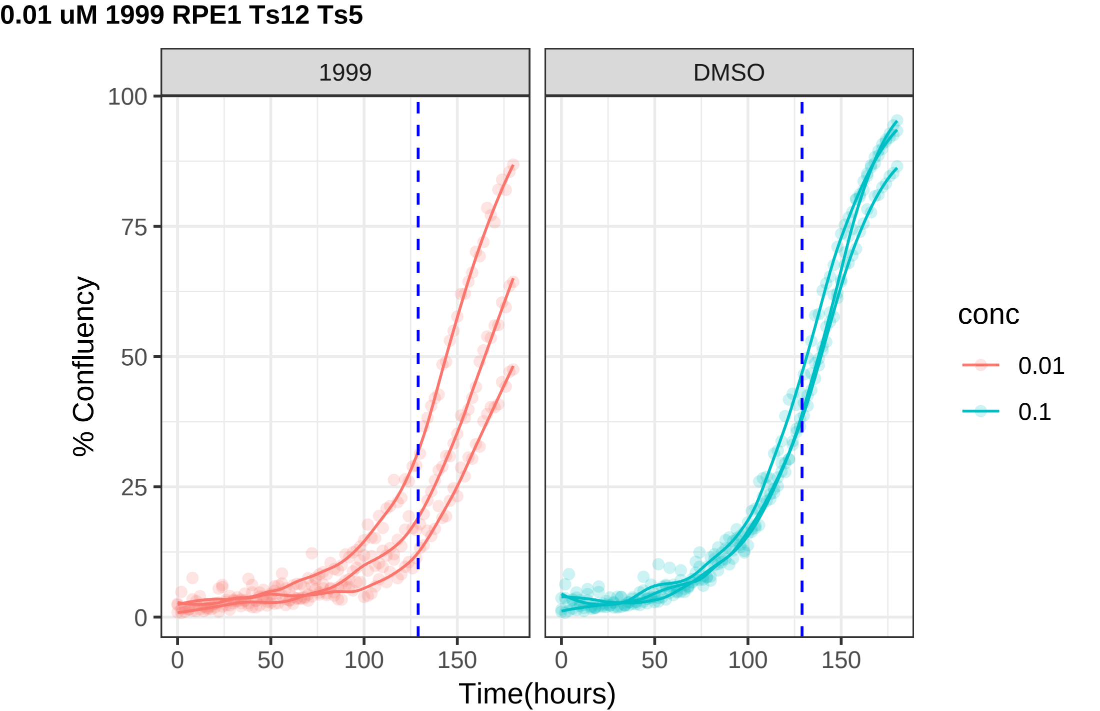

0.01 uM 1999 RPE1 Ts12 Ts5
1999
DMSO
100
75
50
25
0
% Confluency
0
50
100
150
Time(hours)
conc
0.01
0.1

**1 uM 2008 RPE1 hTert**

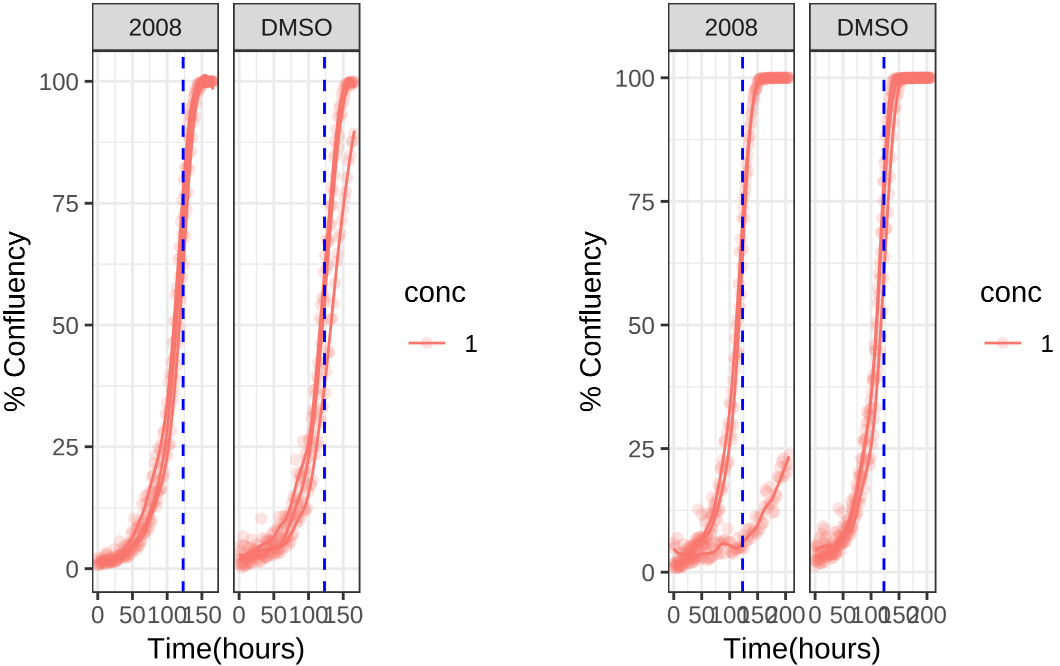

**1 uM 2008 RPE1 Ts12 Ts5**

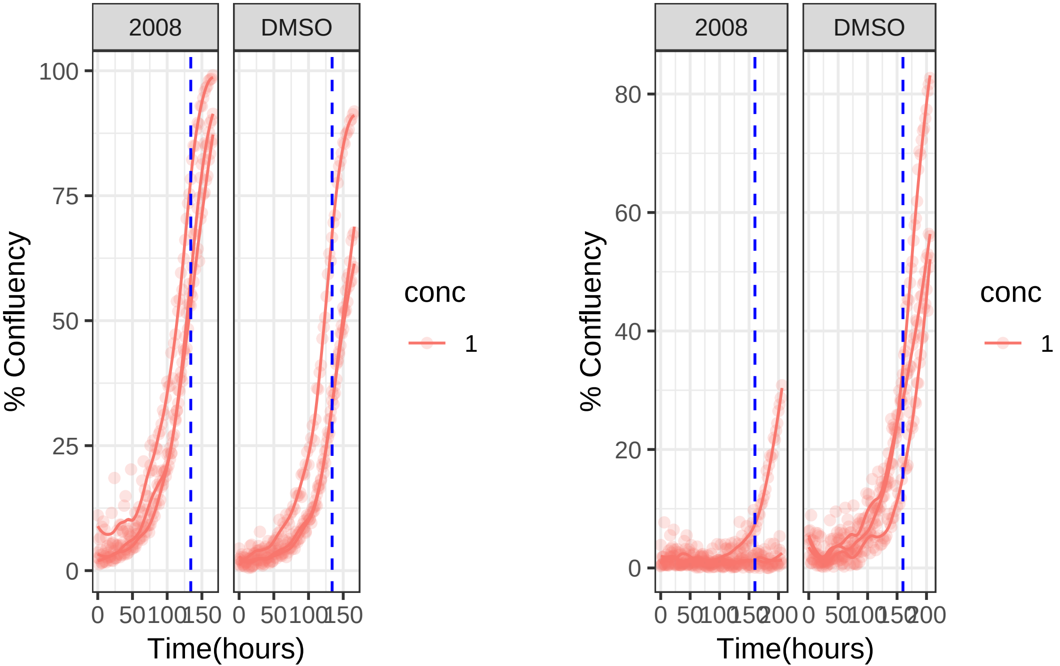

**1 uM 2014 RPE1 hTert**

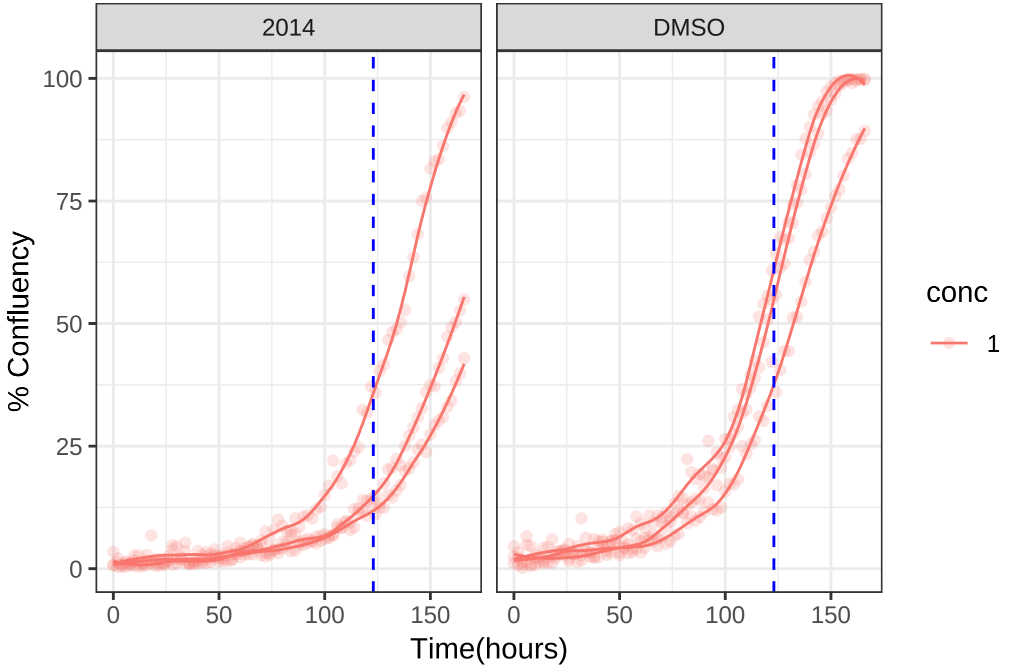

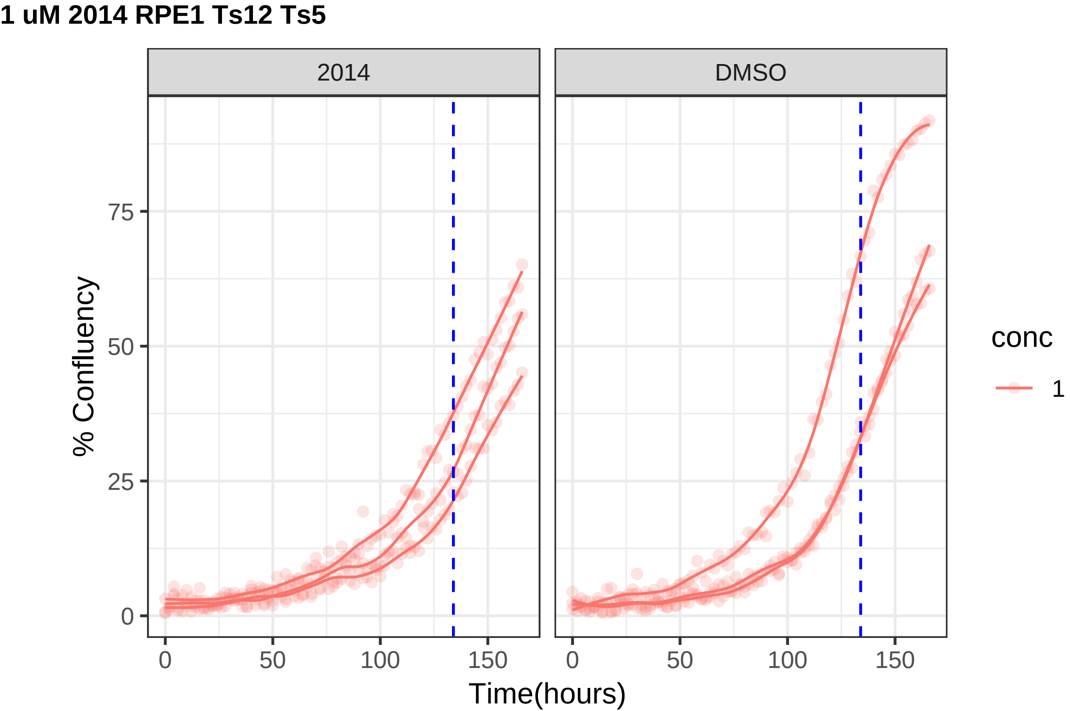
1 uM 2014 RPE1 Ts12 Ts5
2014
DMSO
% Confluency
0
25
50
75
0
50
100
150
0
50
100
150
Time(hours)
conc
1

### 10 uM 2084 RPE1 hTert

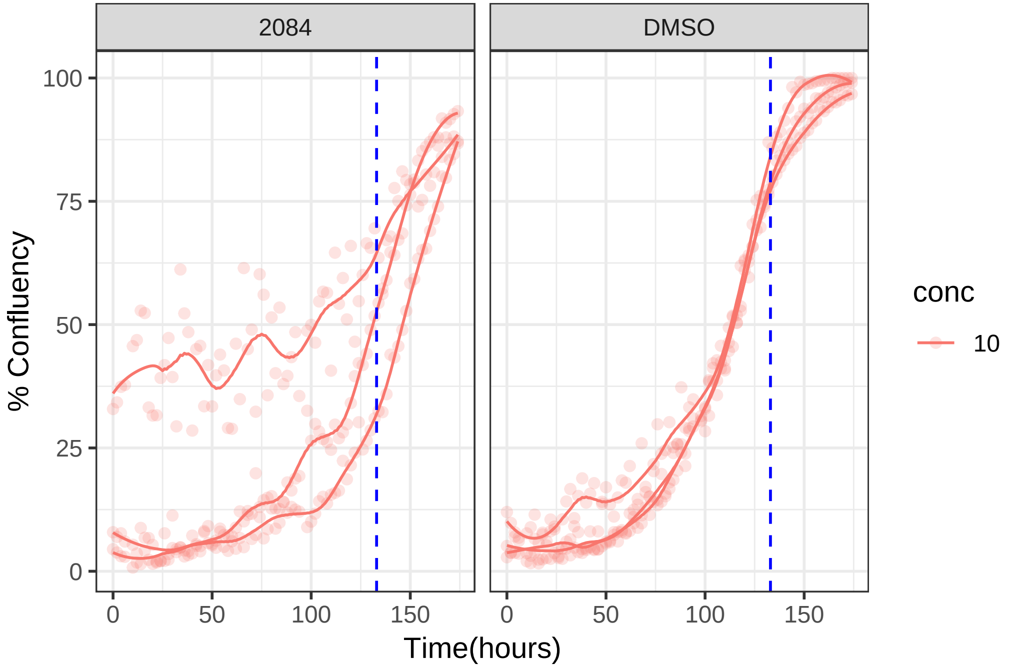

#### 10 uM 2084 RPE1 Ts12 Ts5

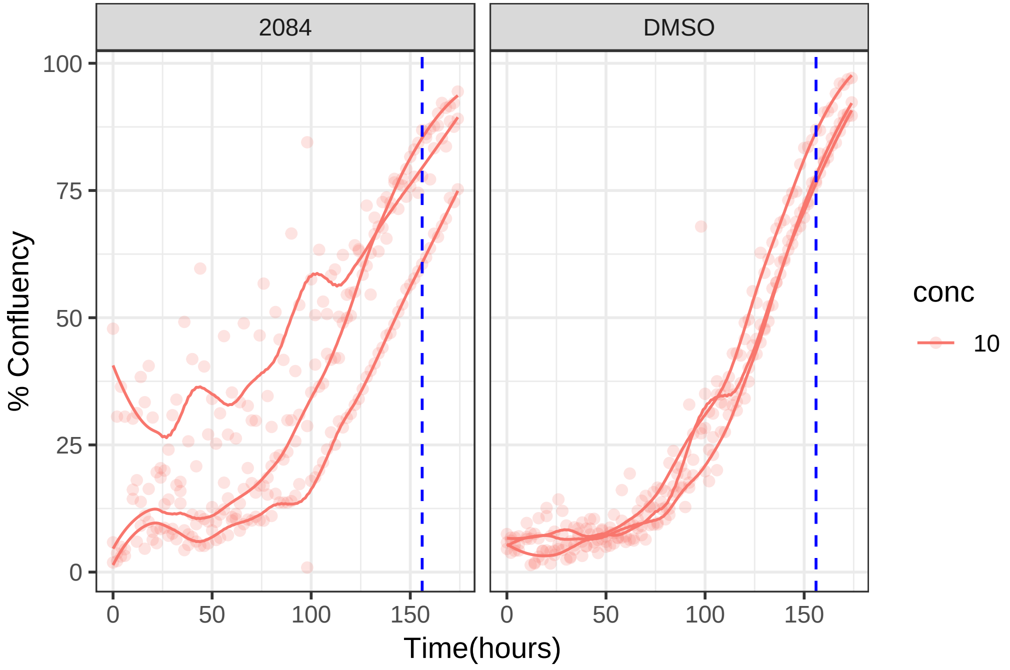

**10 uM 2100 RPE1 hTert**

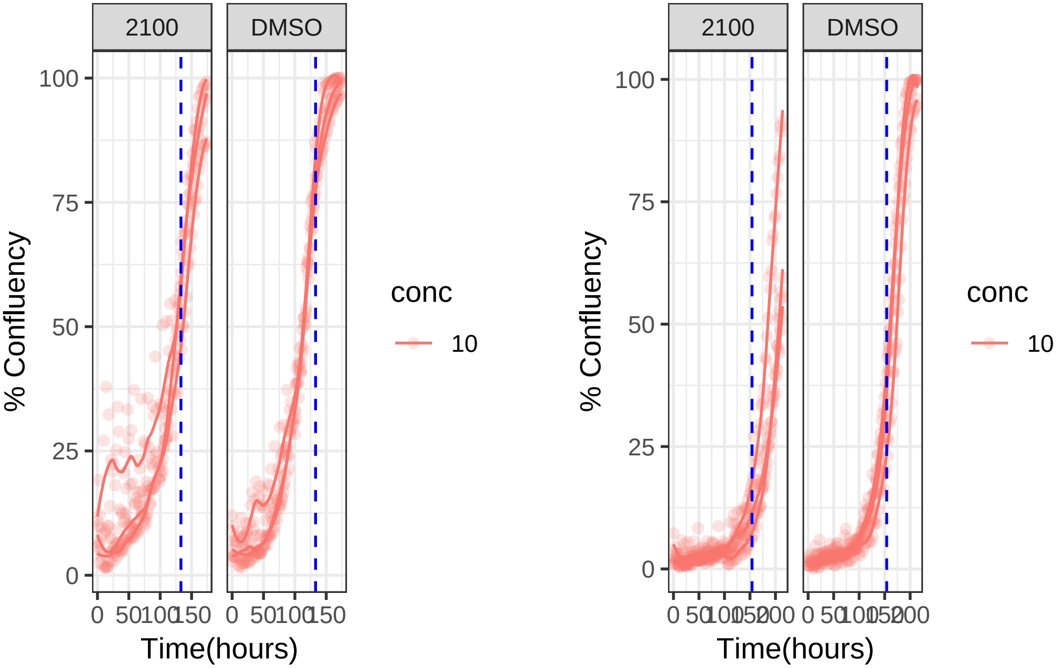

**10 uM 2100 RPE1 Ts12 Ts5**

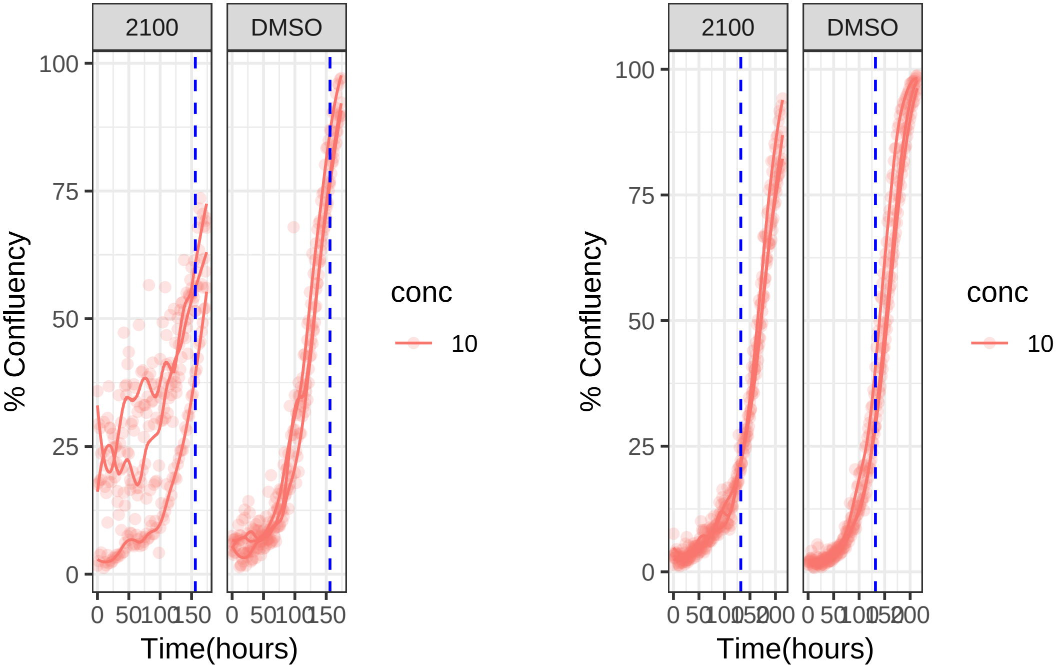

**10 uM 2114 RPE1 hTert**

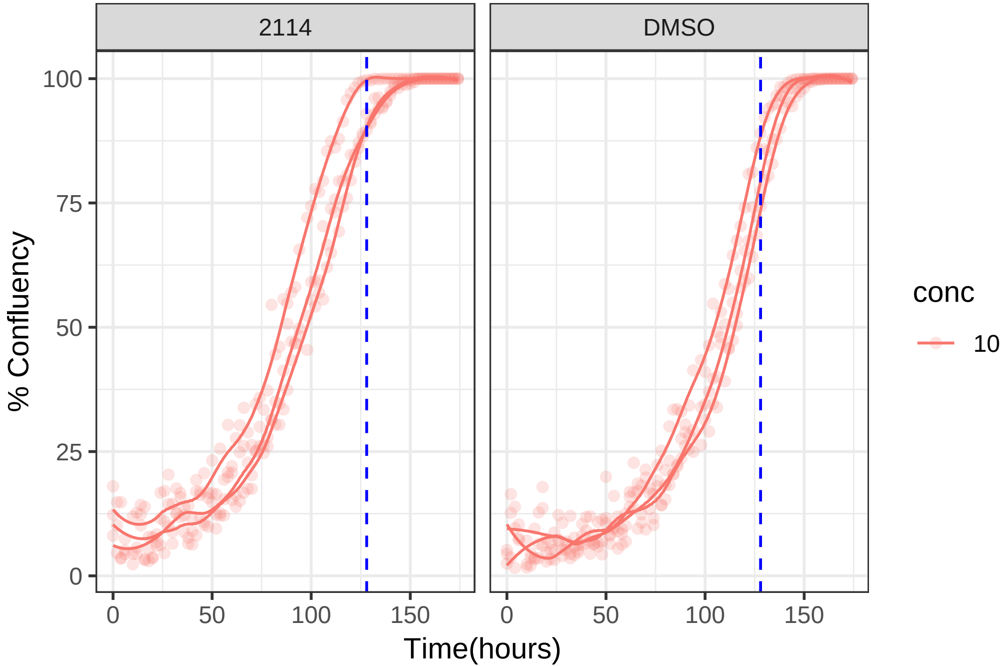

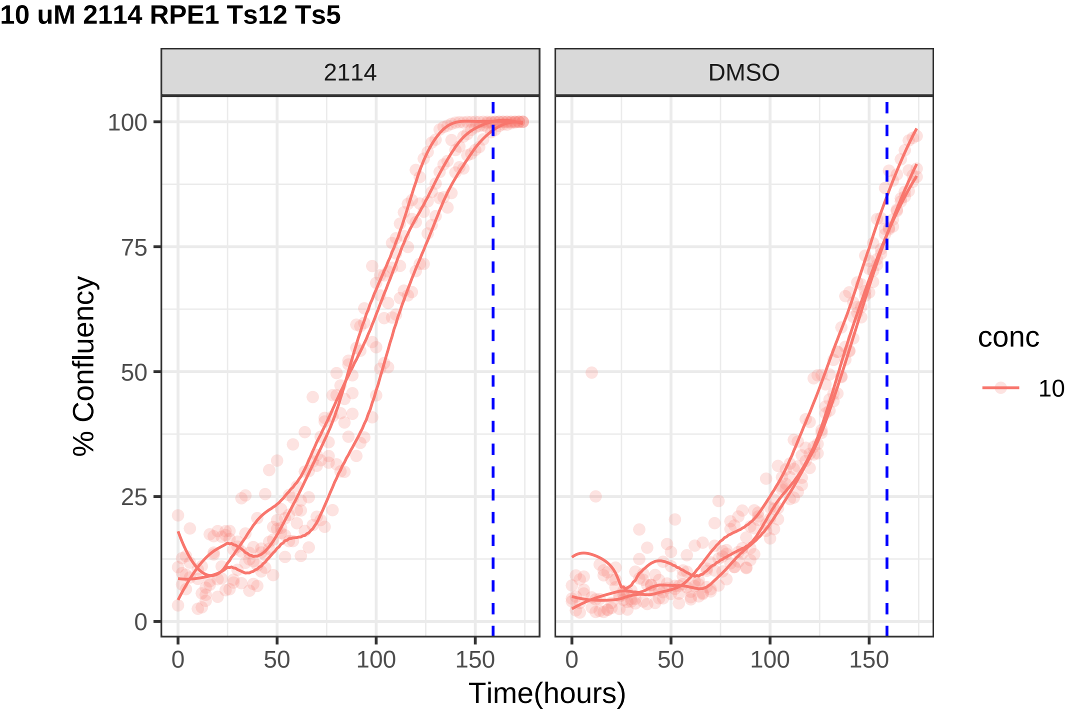

10 uM 2114 RPE1 Ts12 Ts5
2114
DMSO
% Confluency
Time(hours)
0
25
50
75
100
0
50
100
150
conc
10

**0.01 uM 2120 RPE1 hTert**

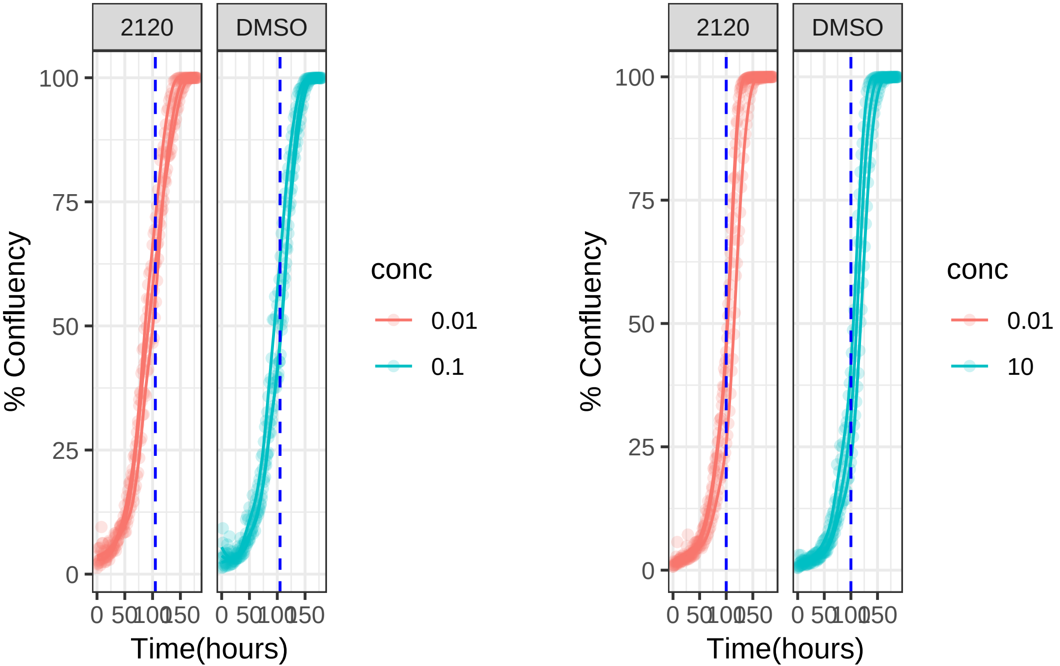

**0.01 uM 2120 RPE1 Ts12 Ts5**

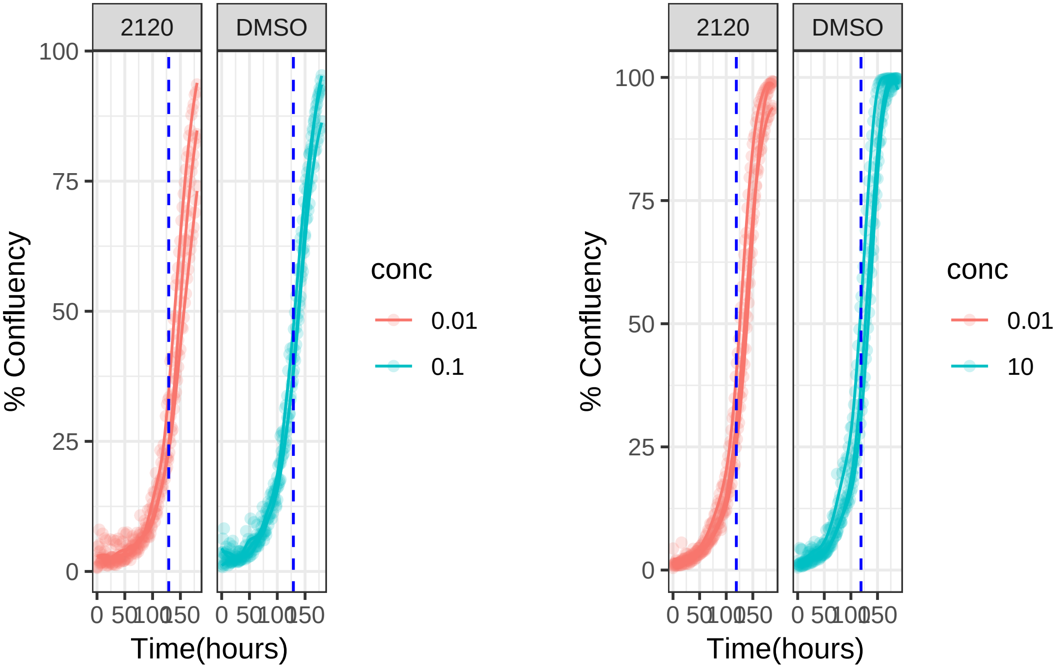

### 10 uM 2138 RPE1 hTert

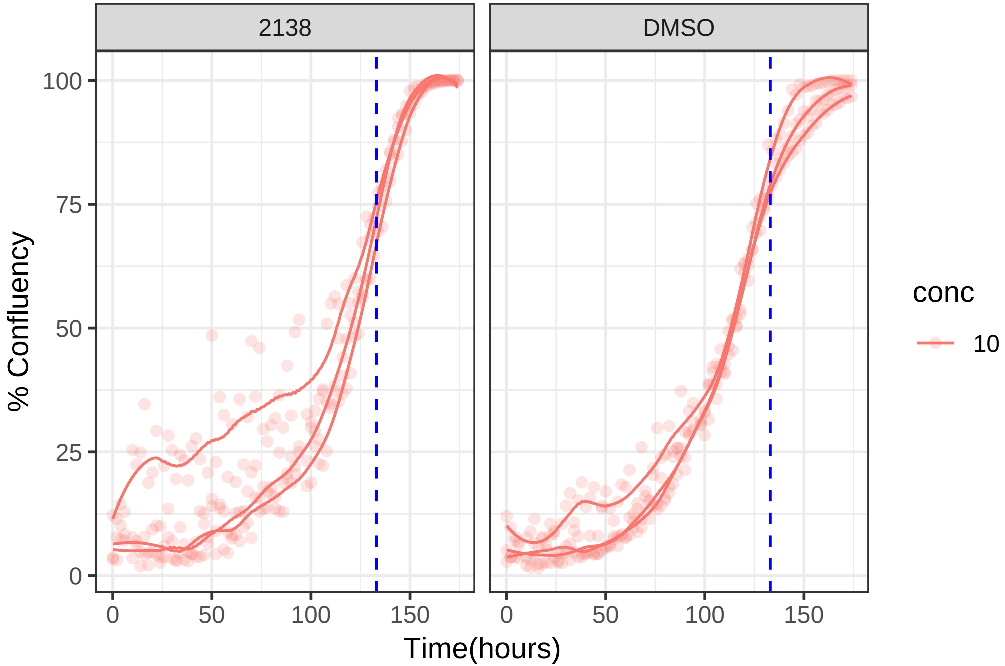

**10 uM 2138 RPE1 Ts12 Ts5**

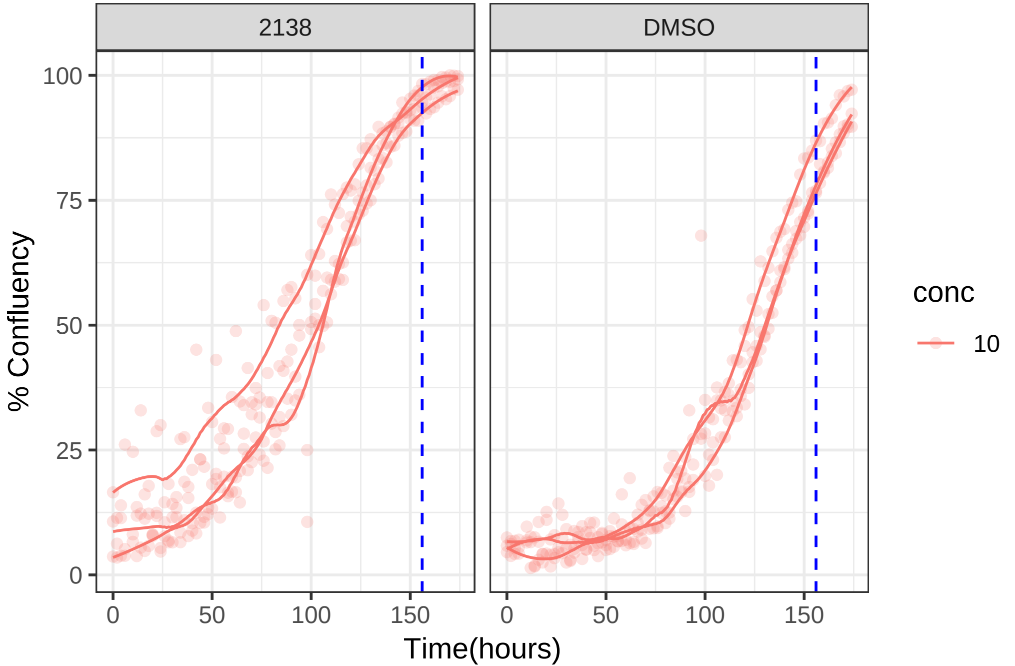

**10 uM 2140 RPE1 hTert**

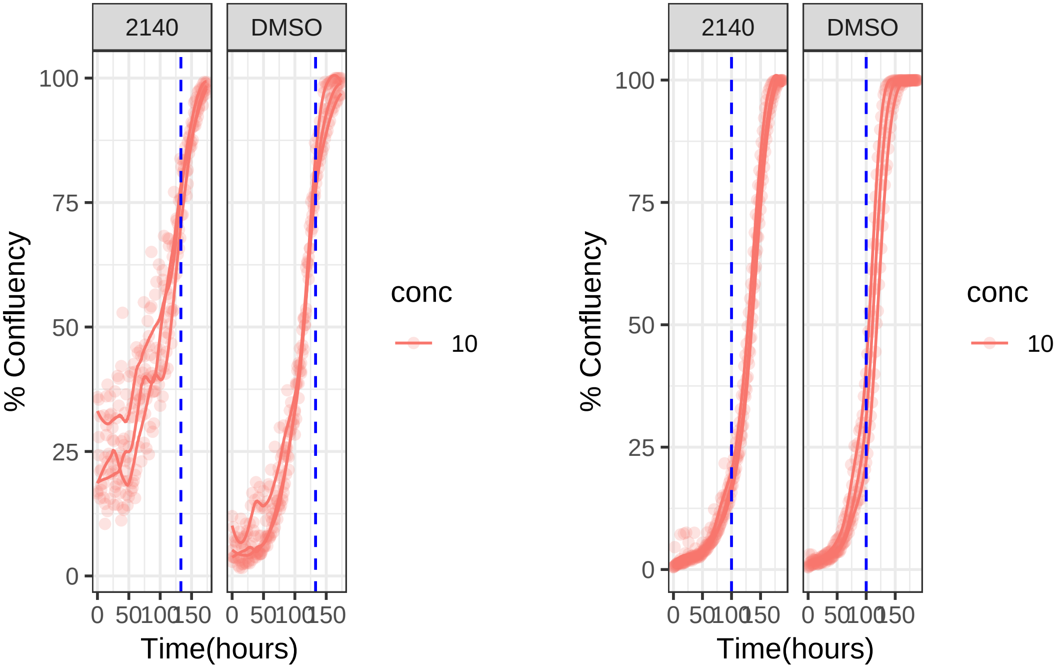

**10 uM 2140 RPE1 Ts12 Ts5**

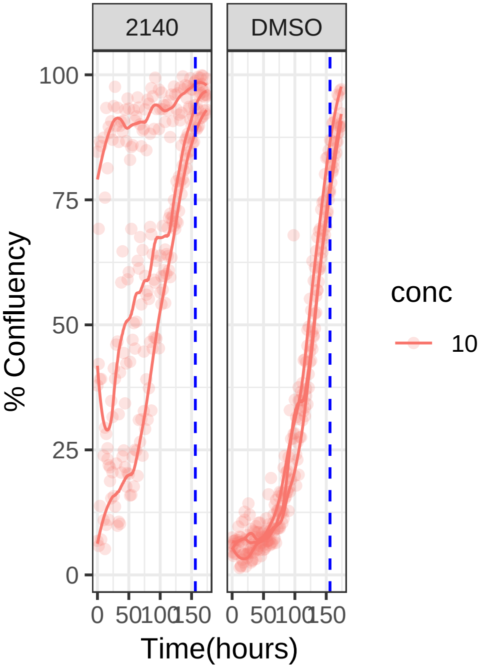


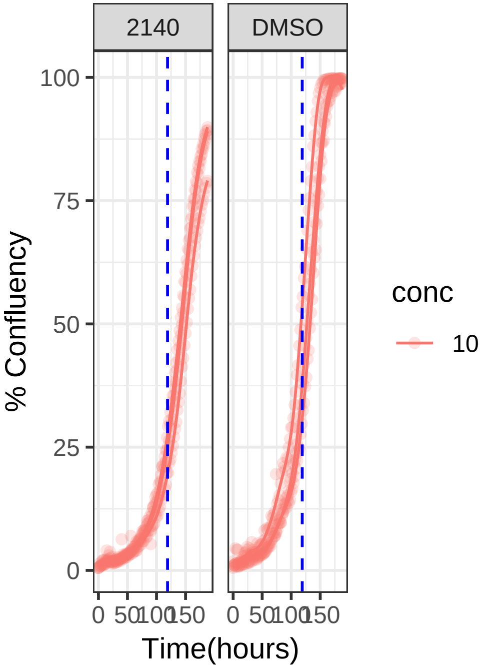

**0.01 uM 2142 RPE1 hTert**

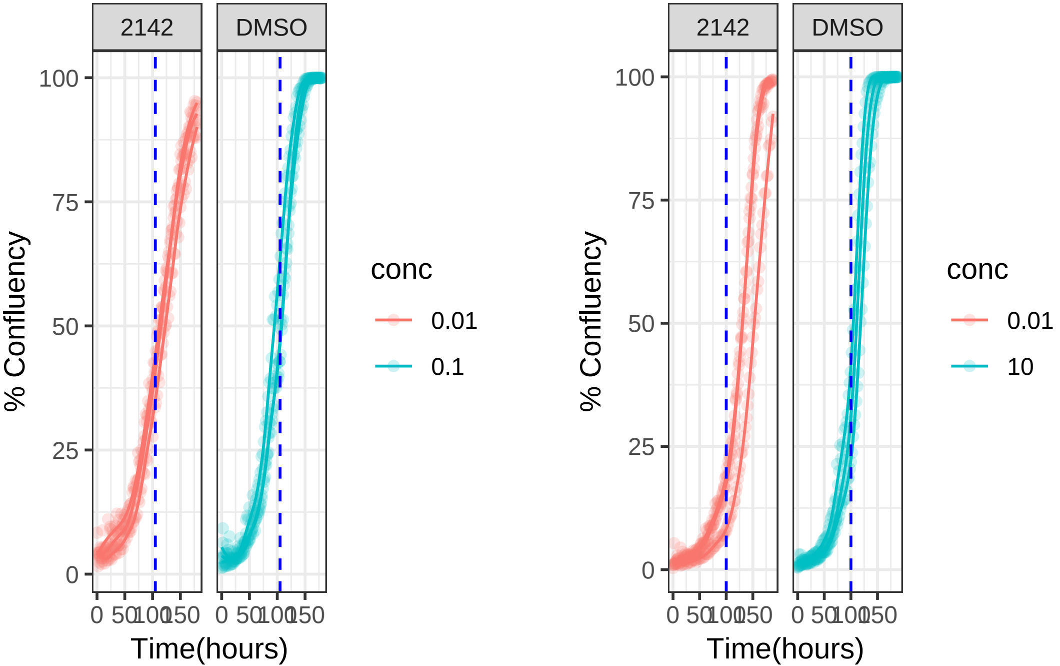

**0.01 uM 2142 RPE1 Ts12 Ts5**

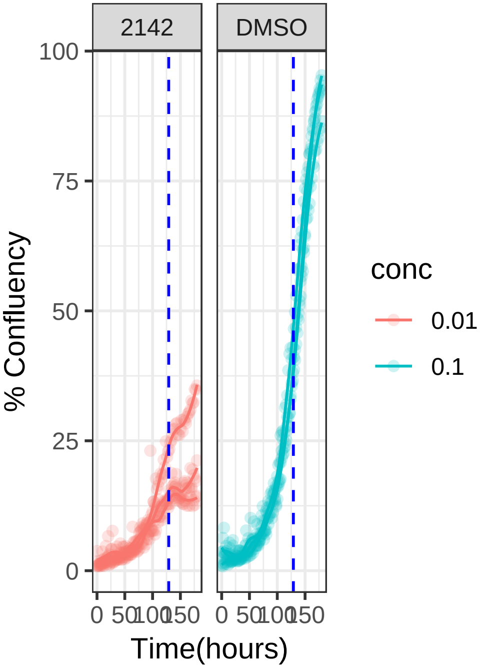


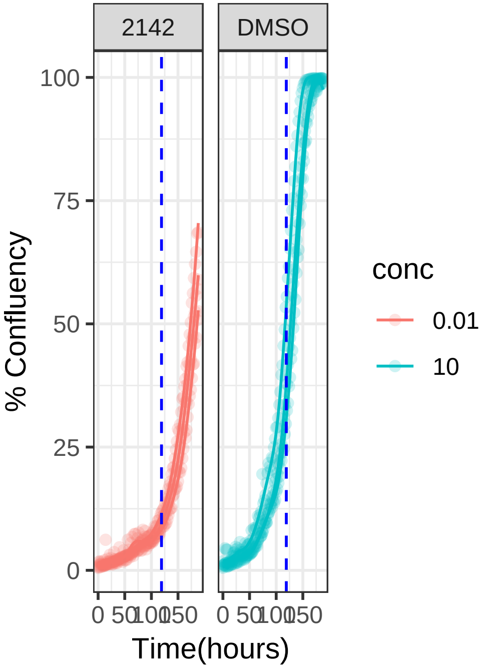

**10 uM 2158 RPE1 hTert**

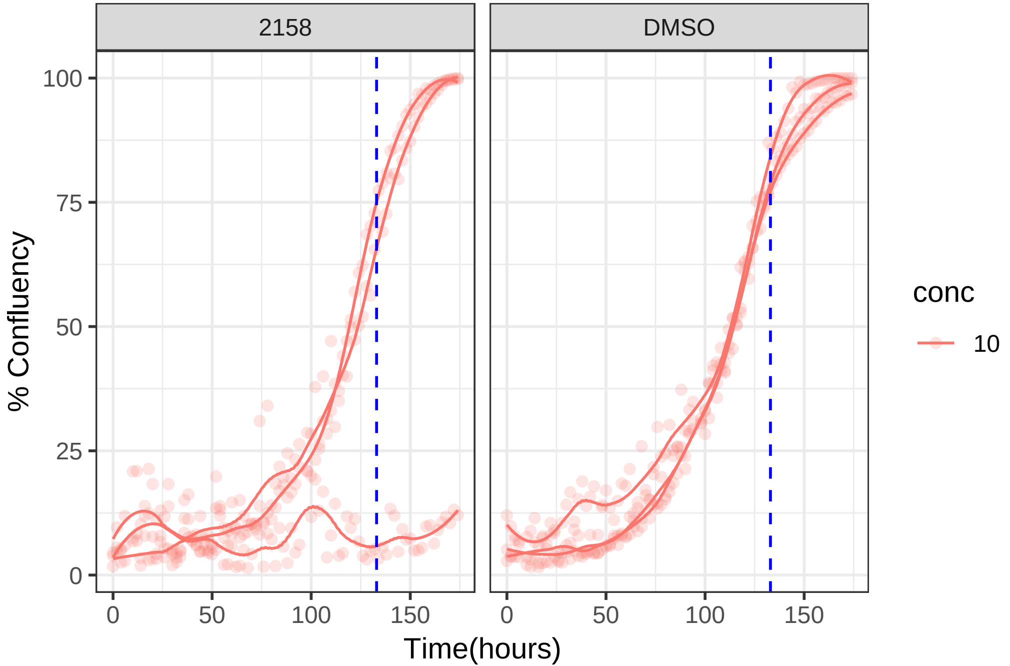

**10 uM 2158 RPE1 Ts12 Ts5**

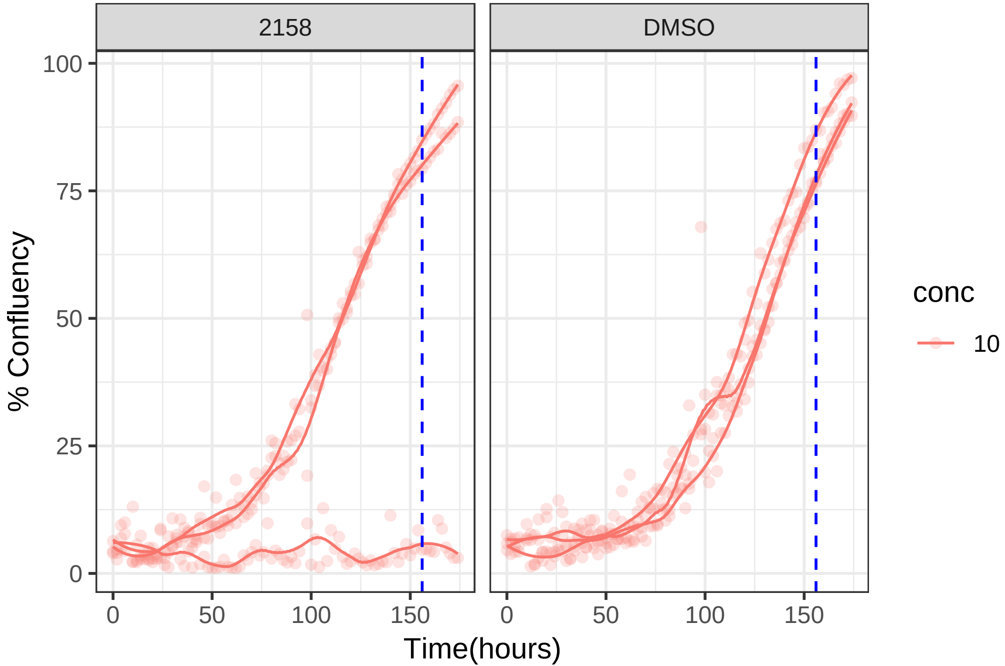

**10 uM 2159 RPE1 hTert**

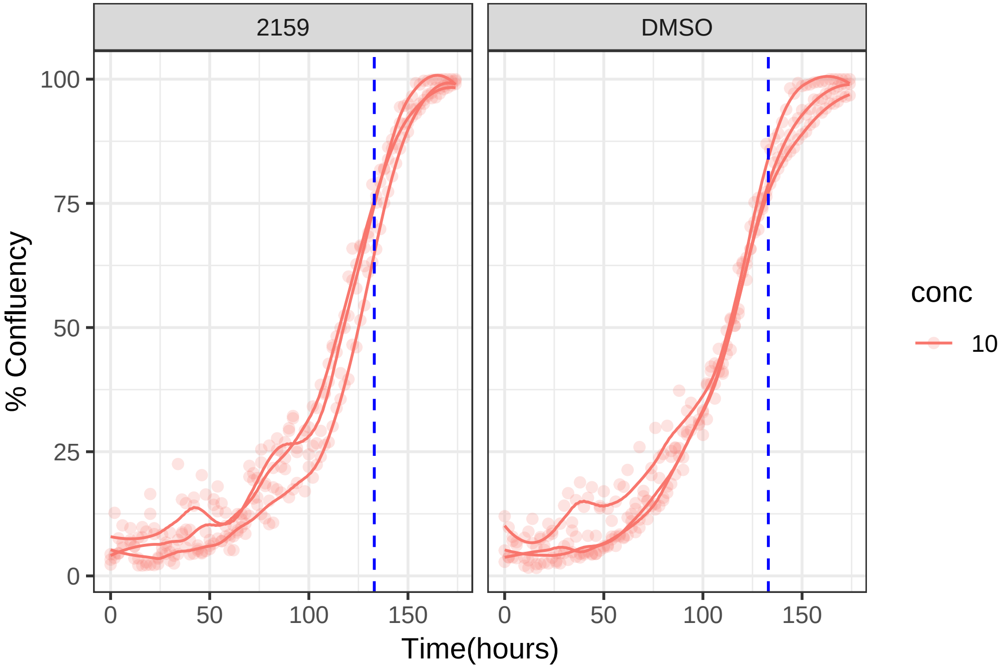

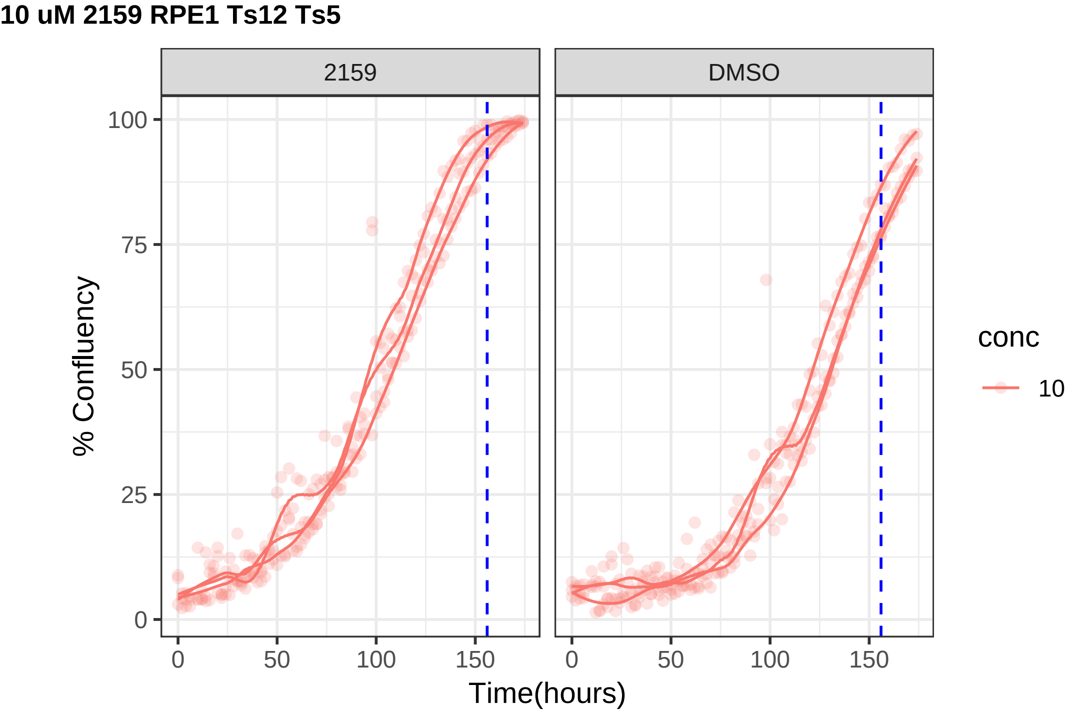
10 uM 2159 RPE1 Ts12 Ts5
2159
DMSO
% Confluency
0
25
50
75
100
0
50
100
150
Time(hours)
conc
10

**0.1 uM 2160 RPE1 hTert**

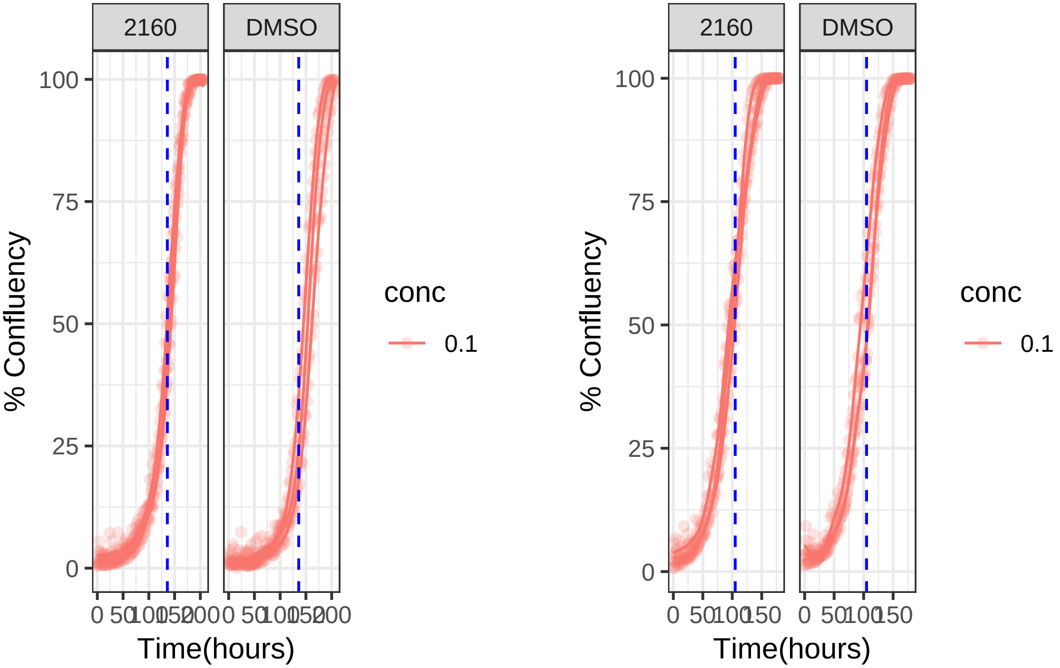

**0.1 uM 2160 RPE1 Ts12 Ts5**

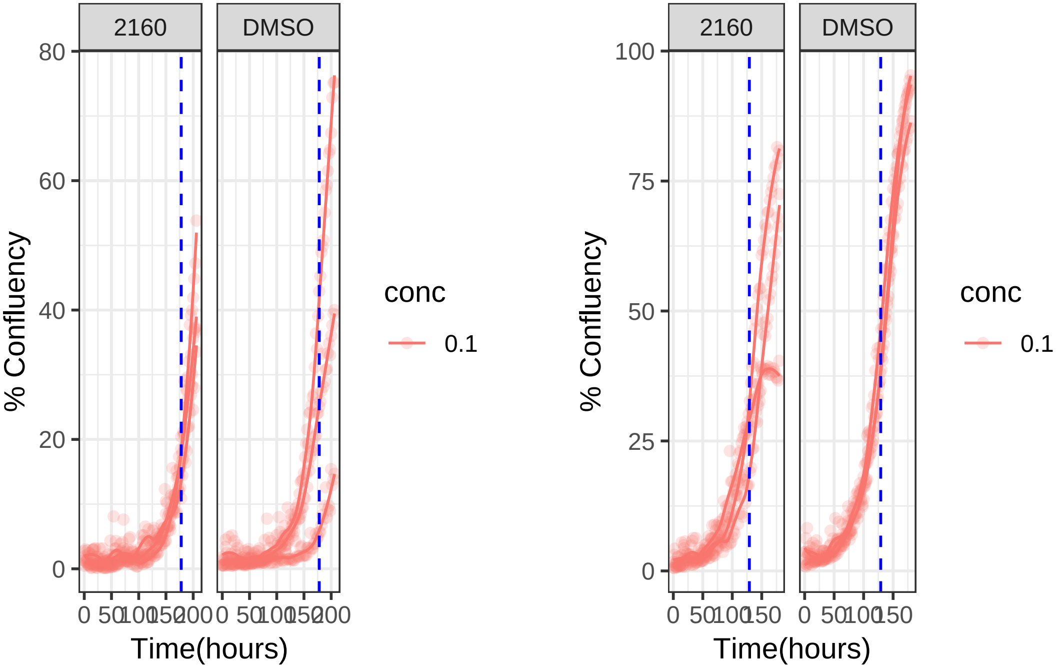

**10 uM 2165 RPE1 hTert**

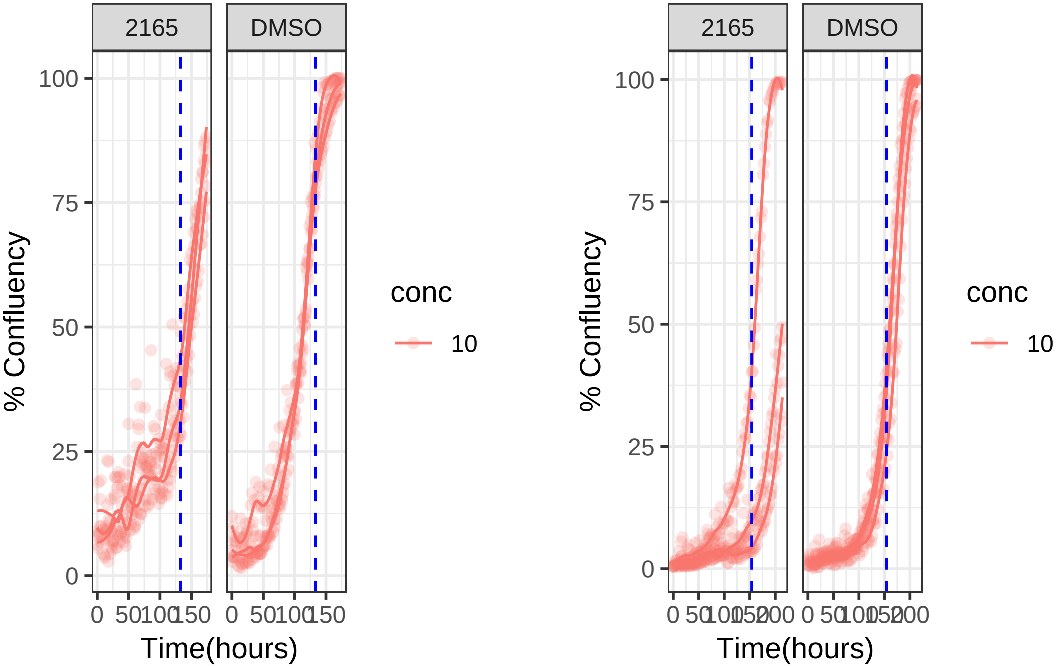

**10 uM 2165 RPE1 Ts12 Ts5**

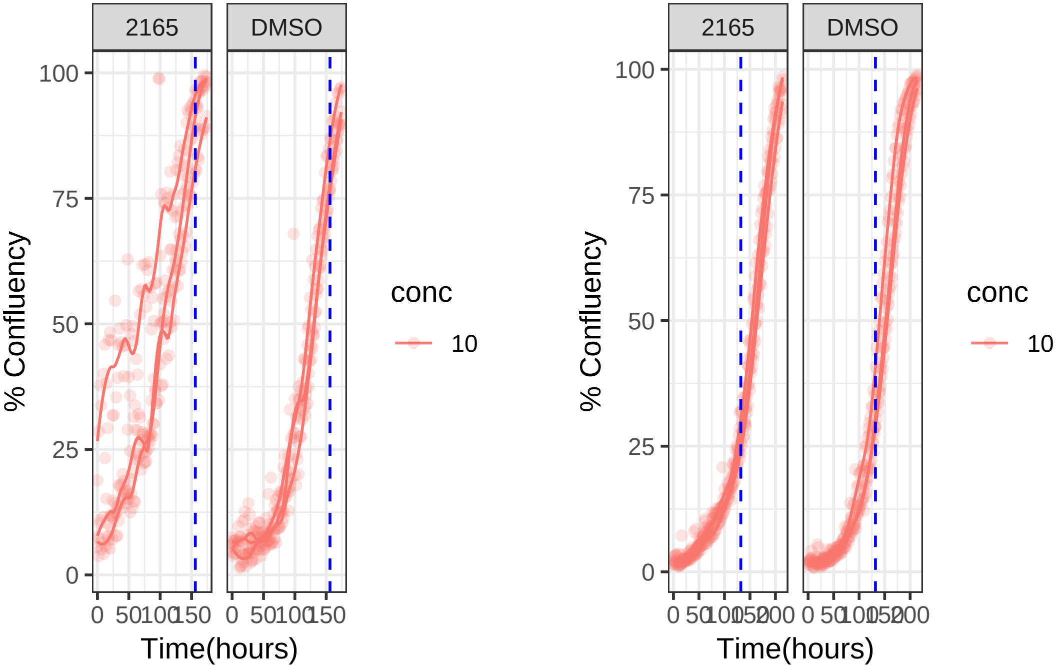

**10 uM 2168 RPE1 hTert**

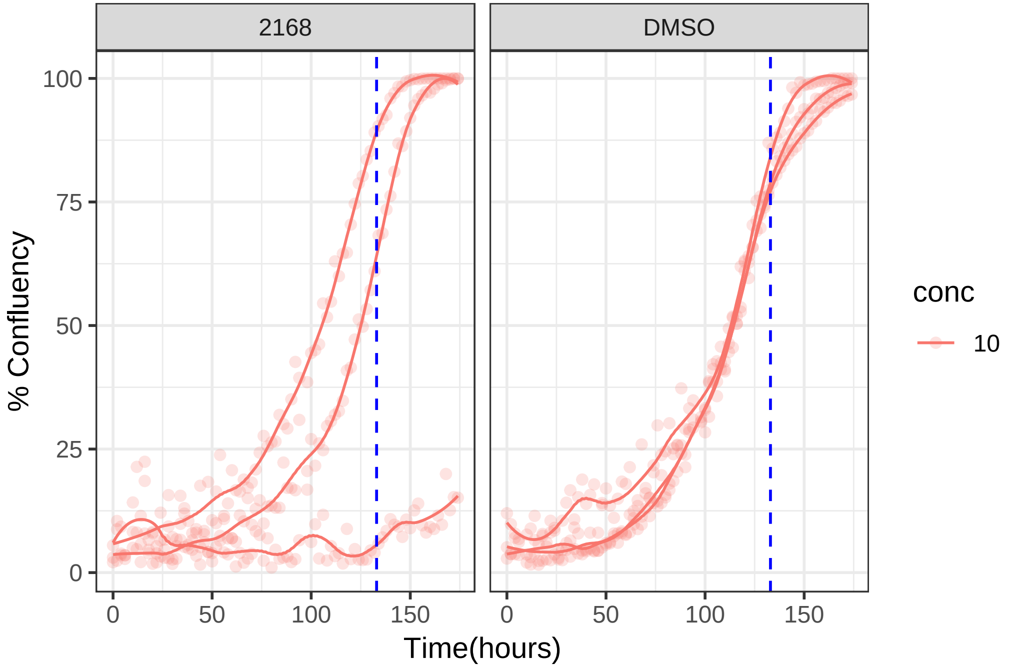

**10 uM 2168 RPE1 Ts12 Ts5**

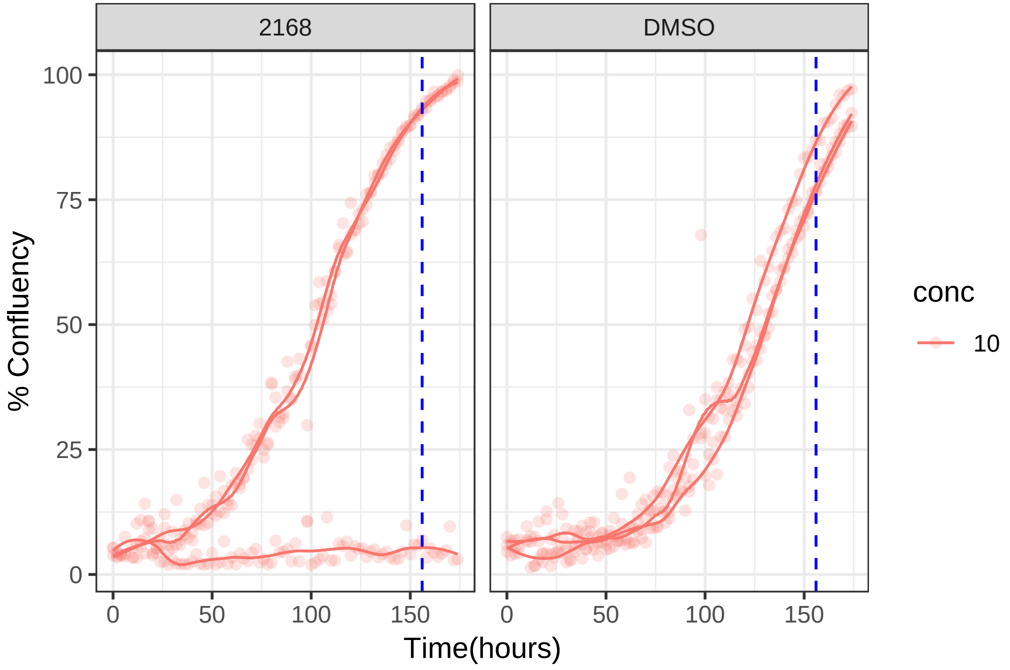

**10 uM 2170 RPE1 hTert**

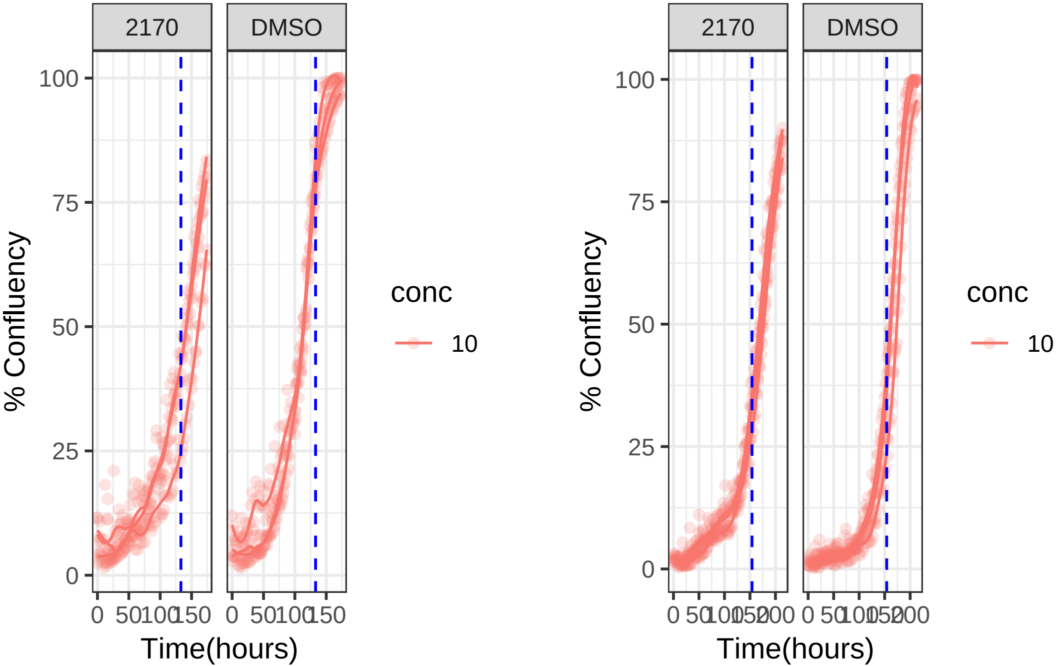

**10 uM 2170 RPE1 Ts12 Ts5**

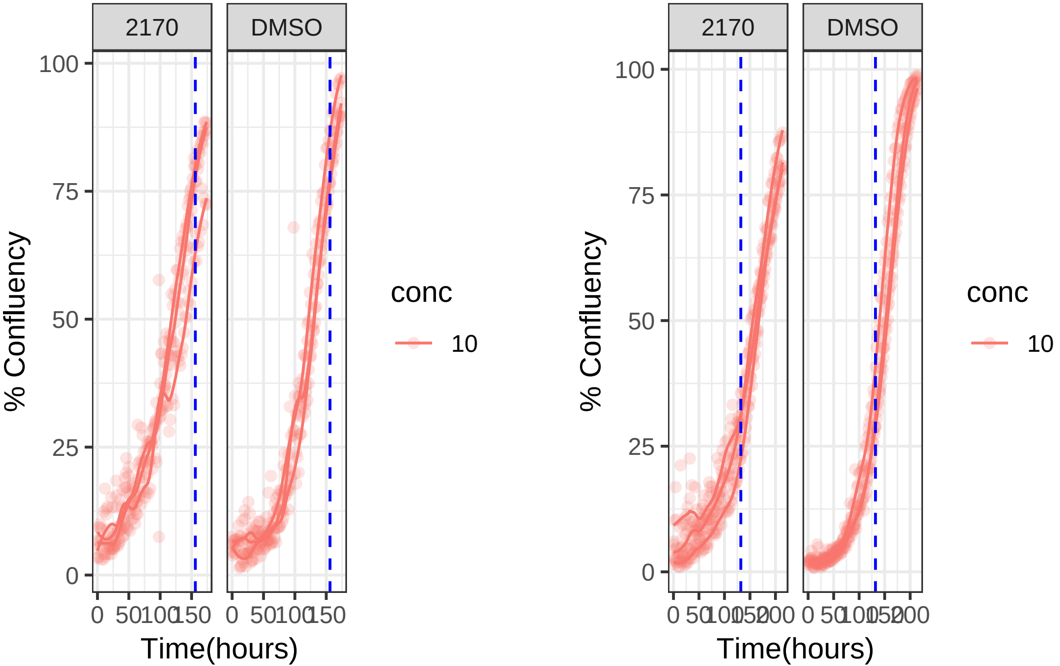

**10 uM 2180 RPE1 hTert**

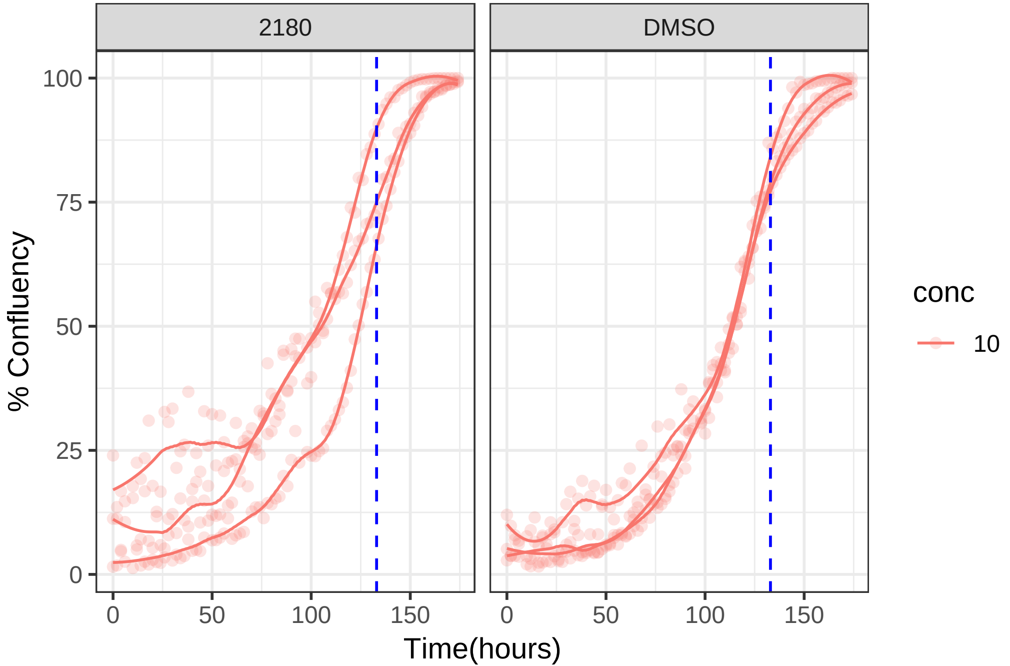

**10 uM 2180 RPE1 Ts12 Ts5**

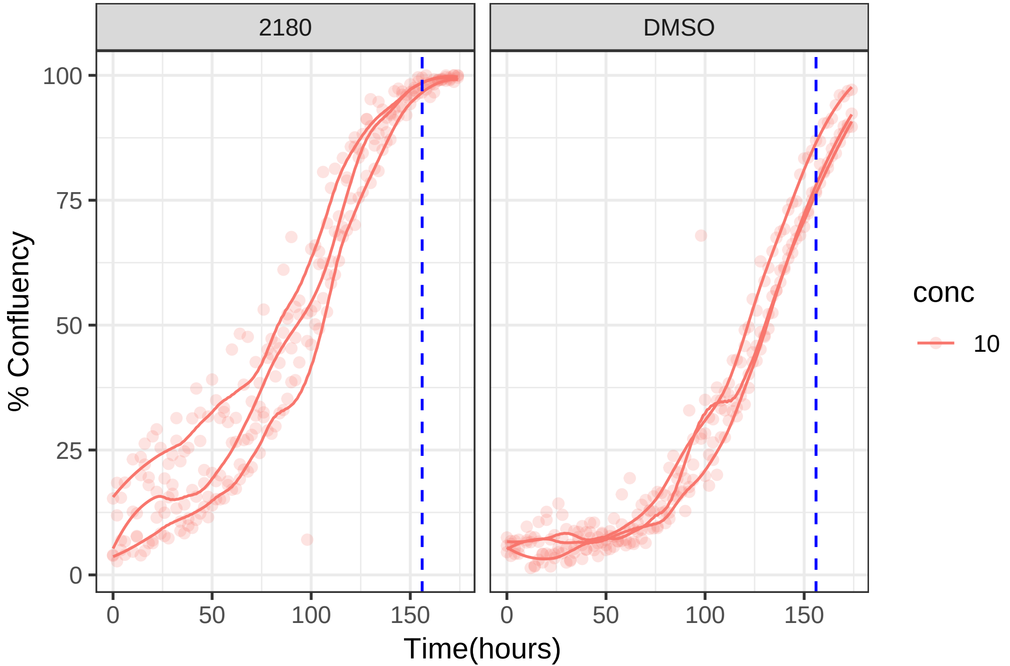

**10 uM 2185 RPE1 hTert**

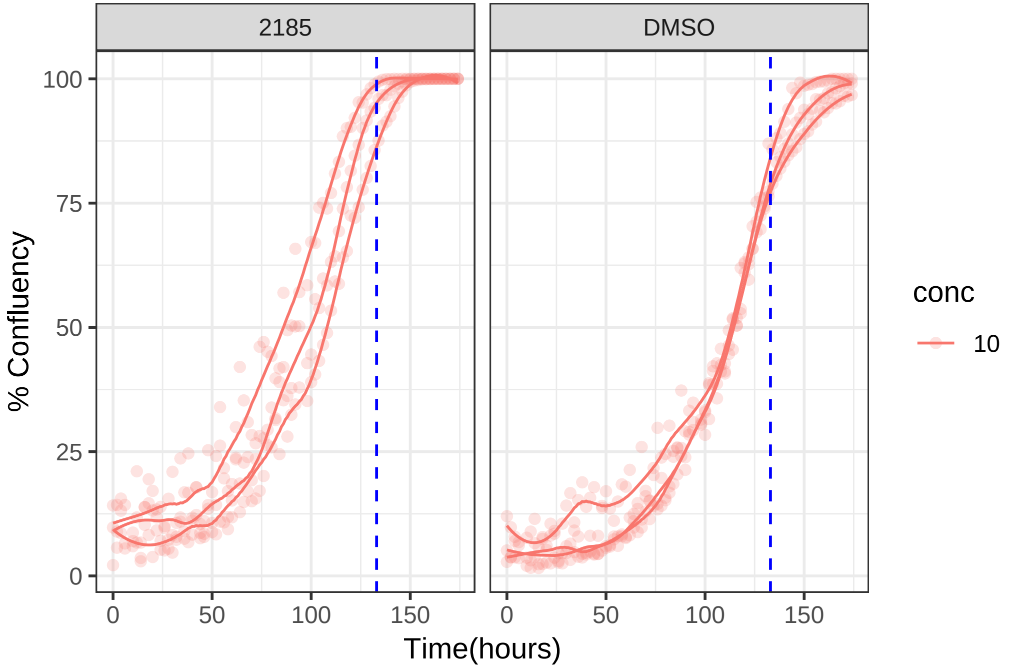

**10 uM 2185 RPE1 Ts12 Ts5**

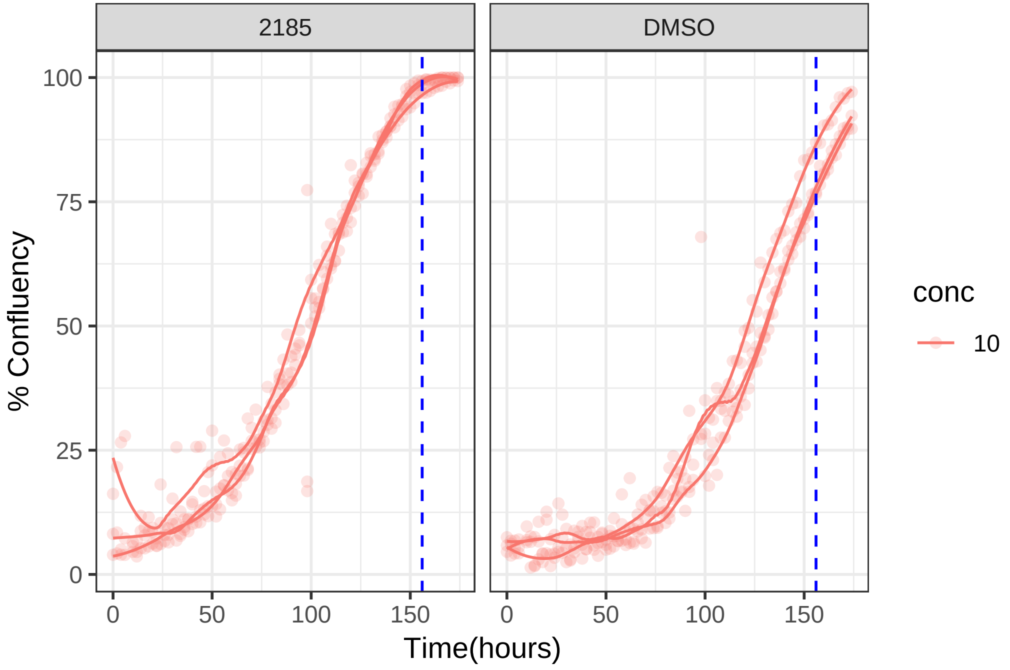

**1 uM 2188 RPE1 hTert**

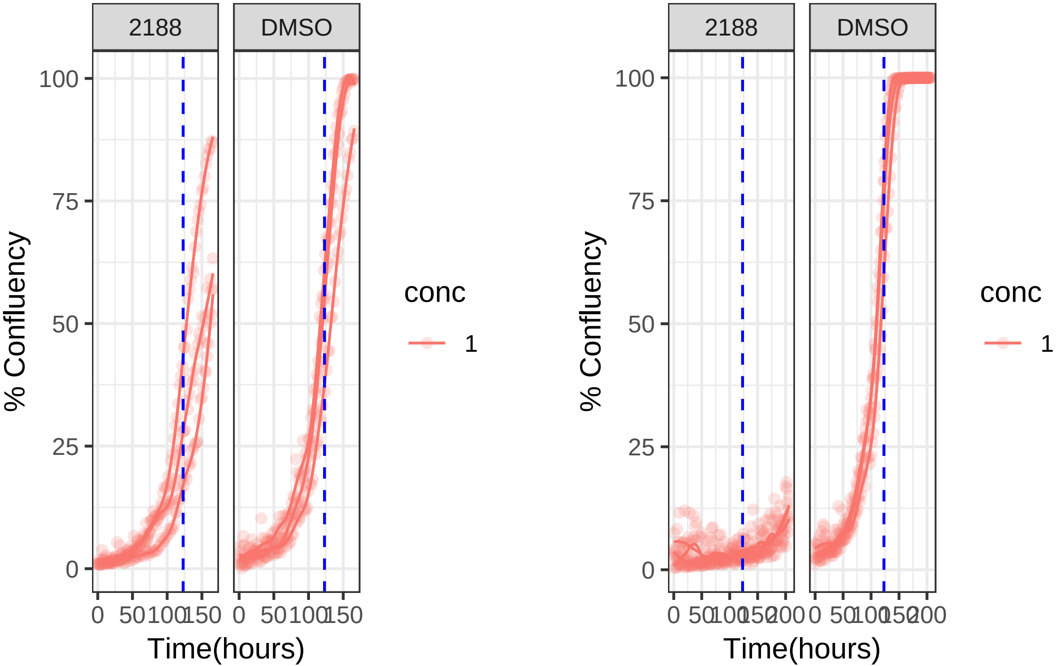

**1 uM 2188 RPE1 Ts12 Ts5**

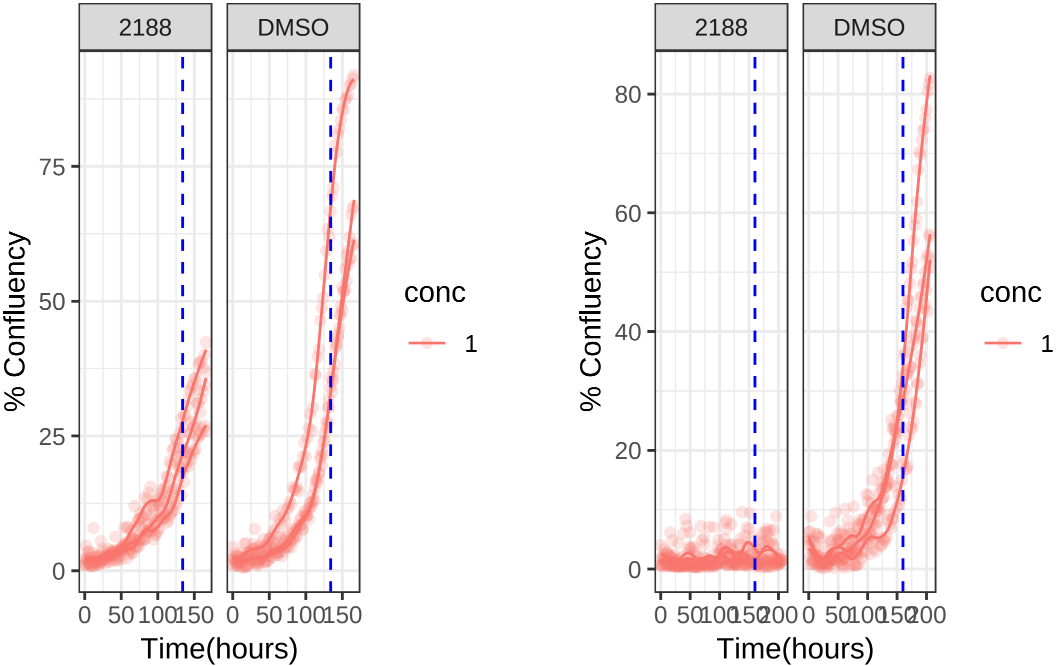

1 uM 2195 RPE1 hTert

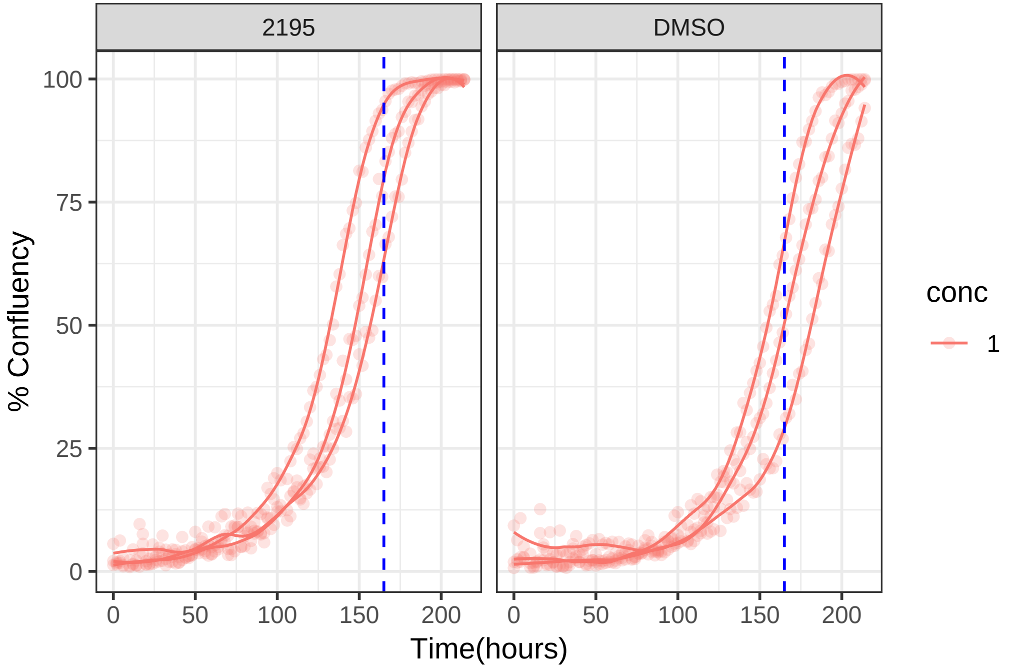

**1 uM 2195 RPE1 Ts12 Ts5**

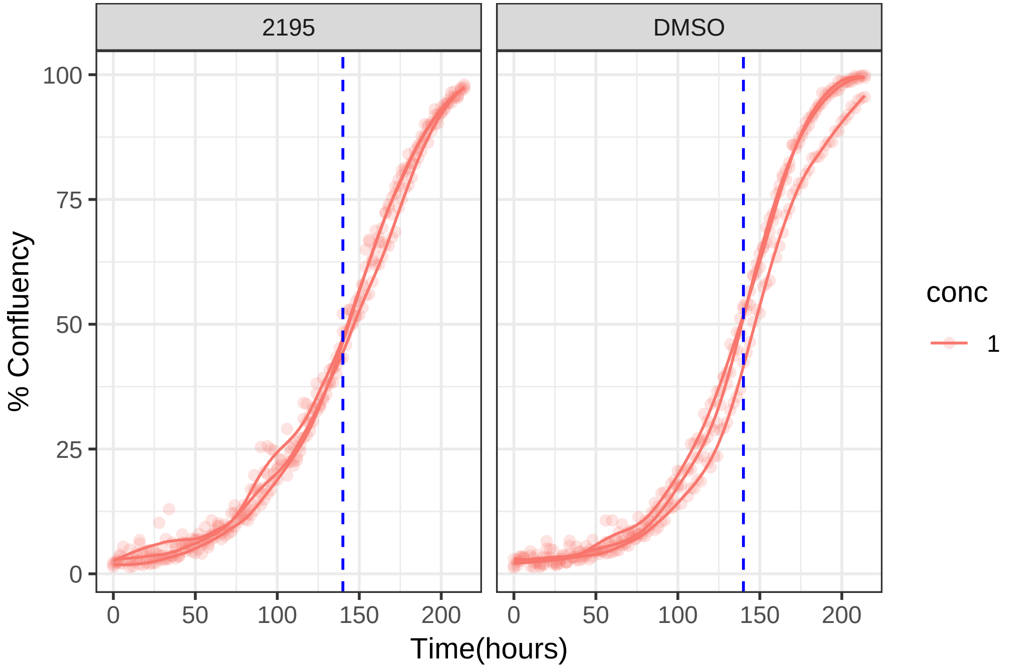

**0.001 uM 2198 RPE1 hTert**

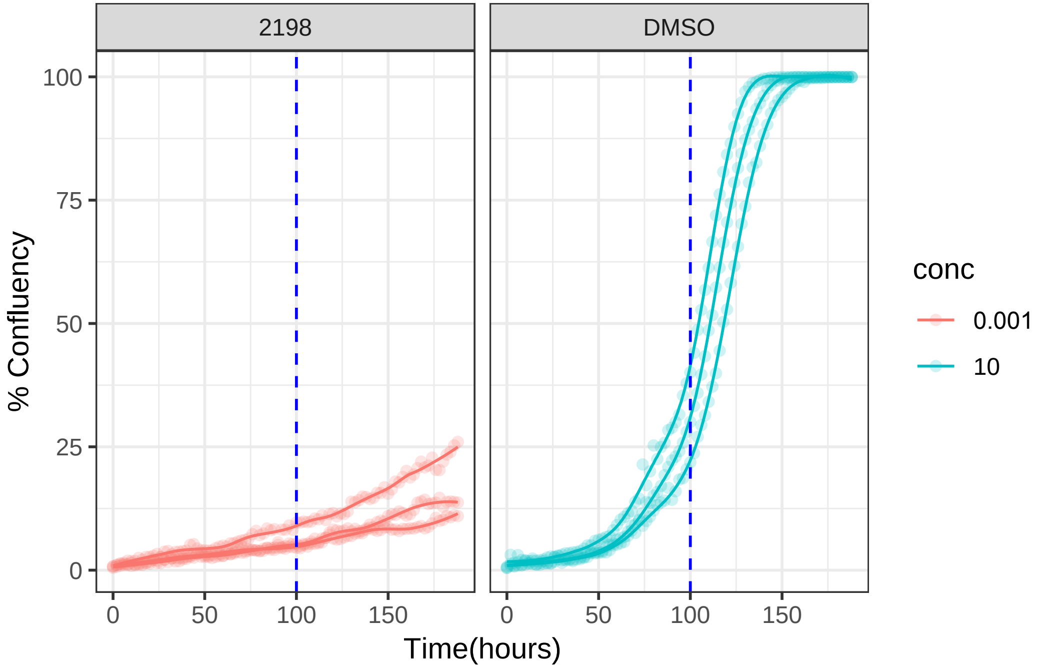

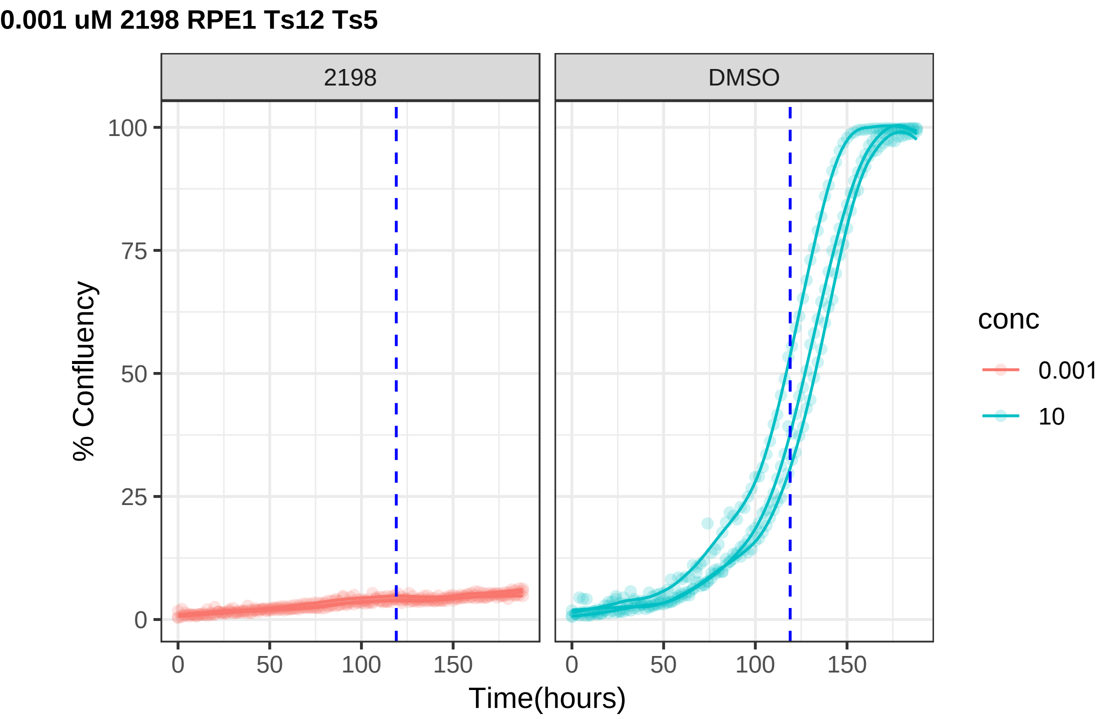
0.001 uM 2198 RPE1 Ts12 Ts5
2198
DMSO
% Confluency
100
75
50
25
0
0
50
100
150
Time(hours)
conc
0.001
10

**10 uM 2209 RPE1 hTert**

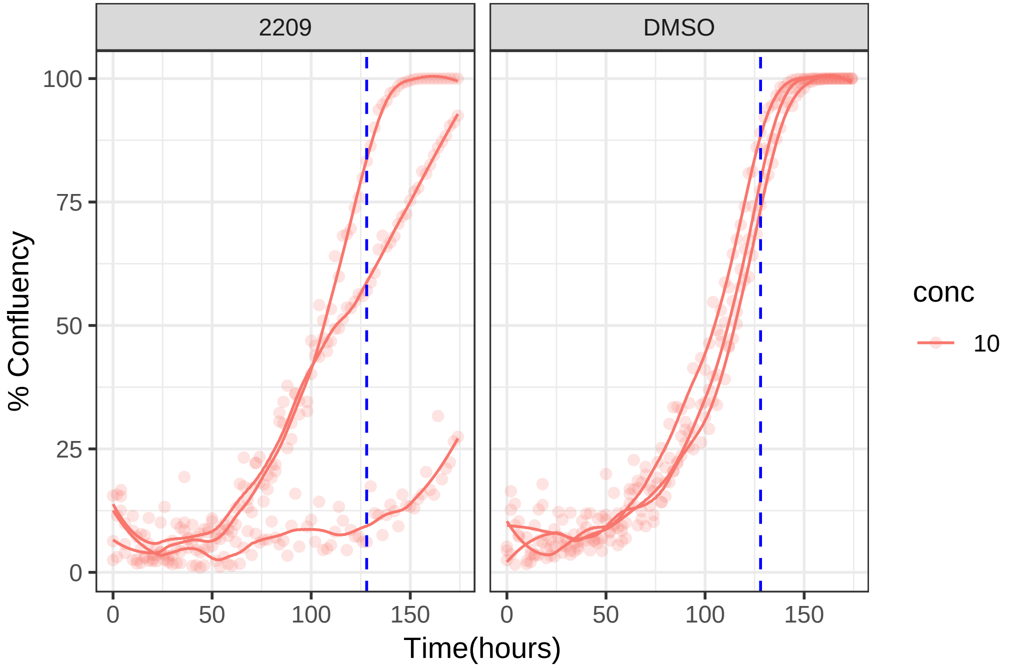

**10 uM 2209 RPE1 Ts12 Ts5**

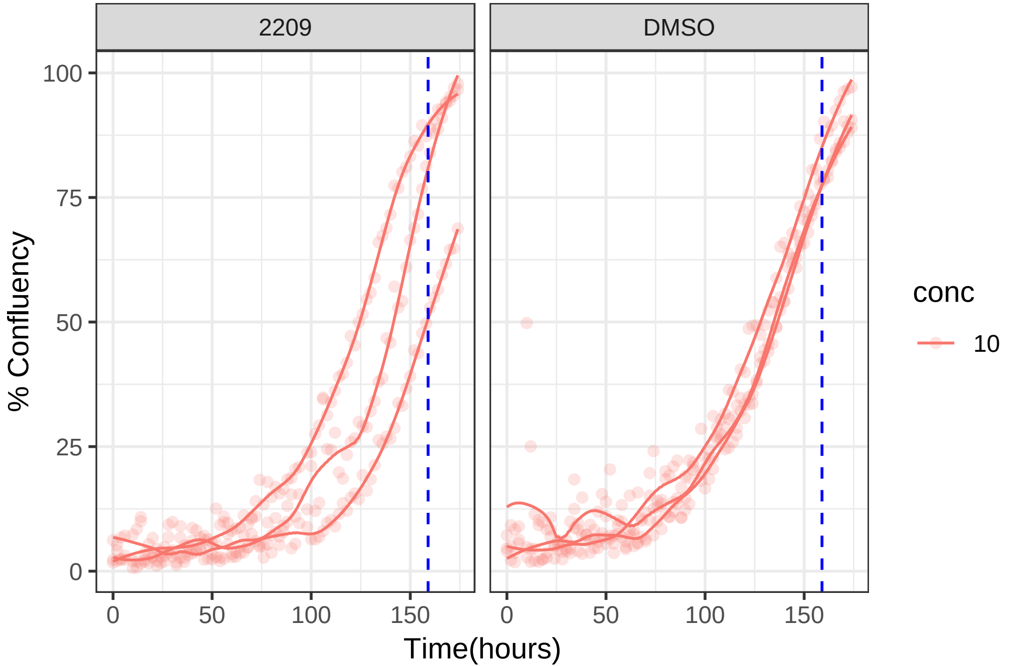

**10 uM 2210 RPE1 hTert**

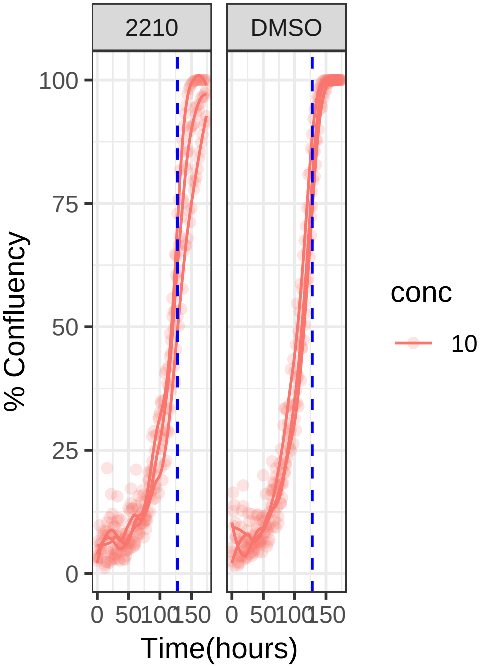


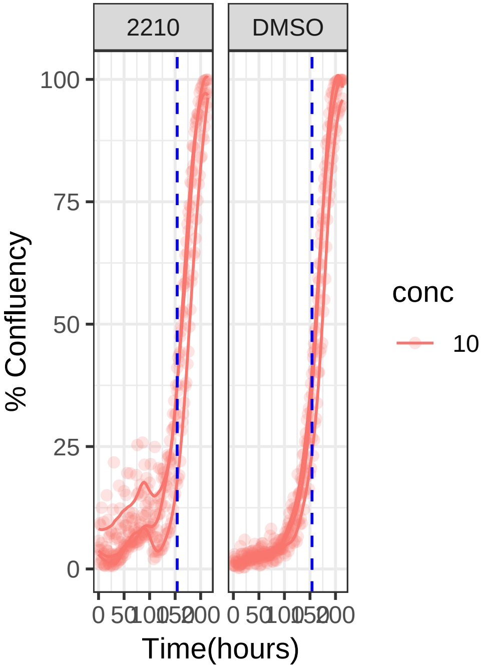

**10 uM 2210 RPE1 Ts12 Ts5**

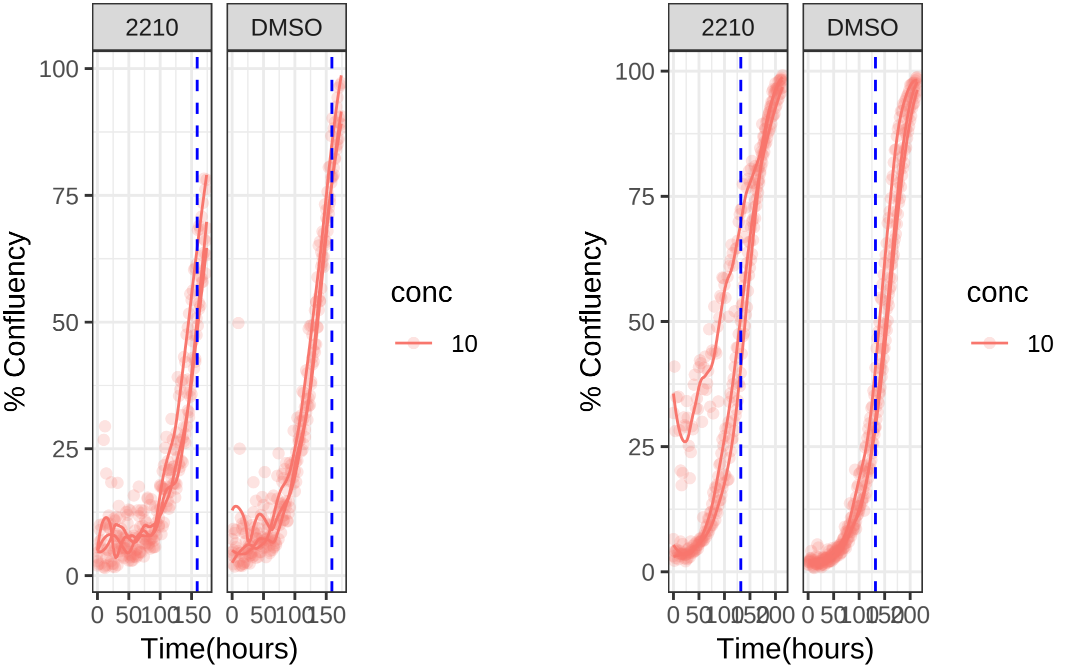

**10 uM 2223 RPE1 hTert**

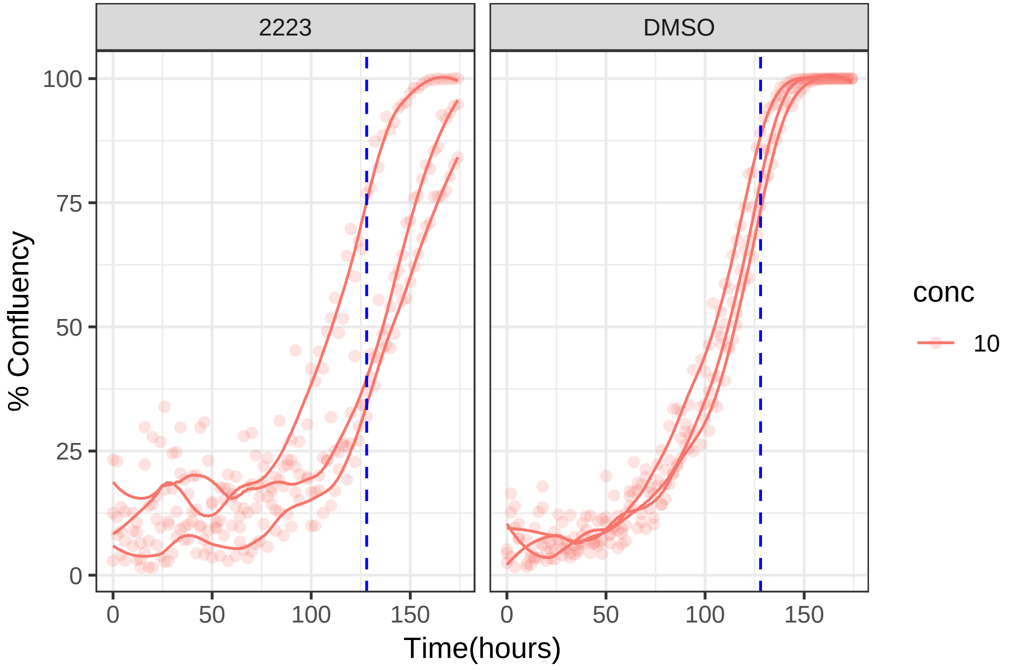

**10 uM 2223 RPE1 Ts12 Ts5**

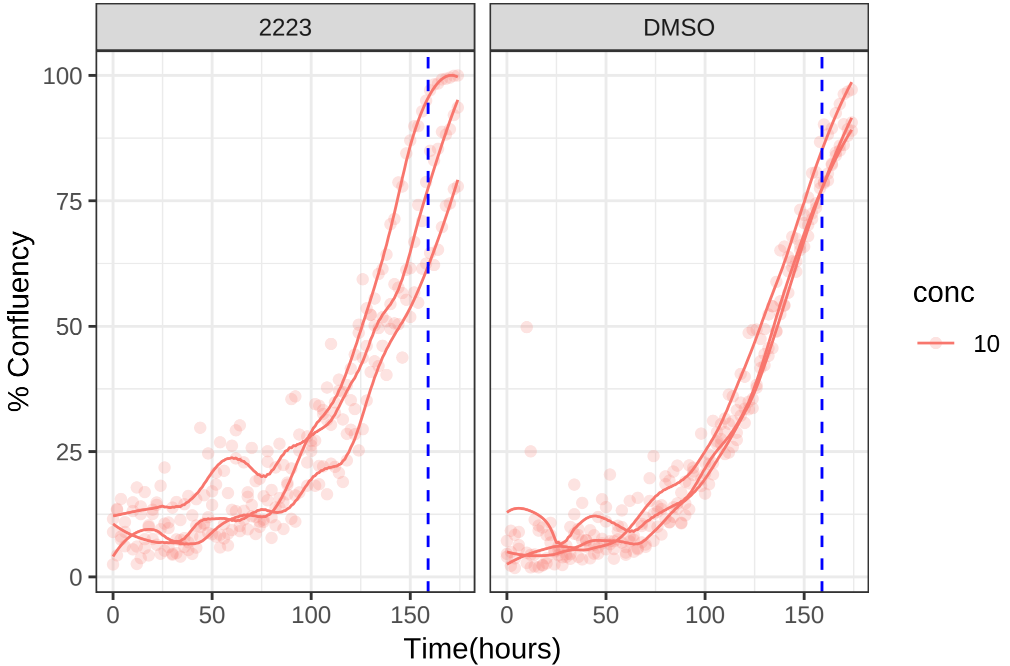

**10 uM 2227 RPE1 hTert**

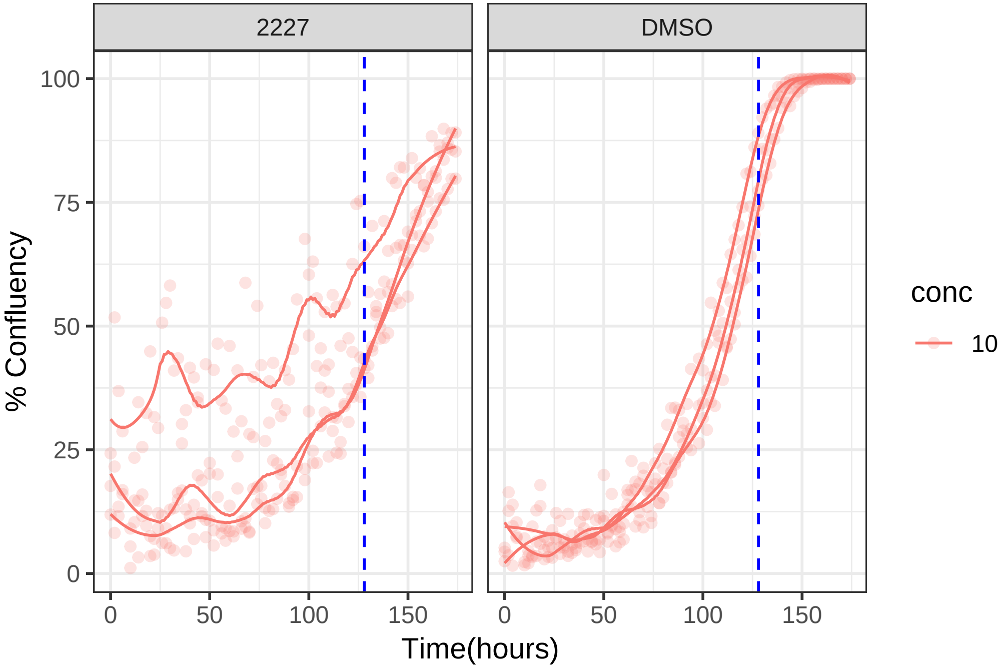

**10 uM 2227 RPE1 Ts12 Ts5**

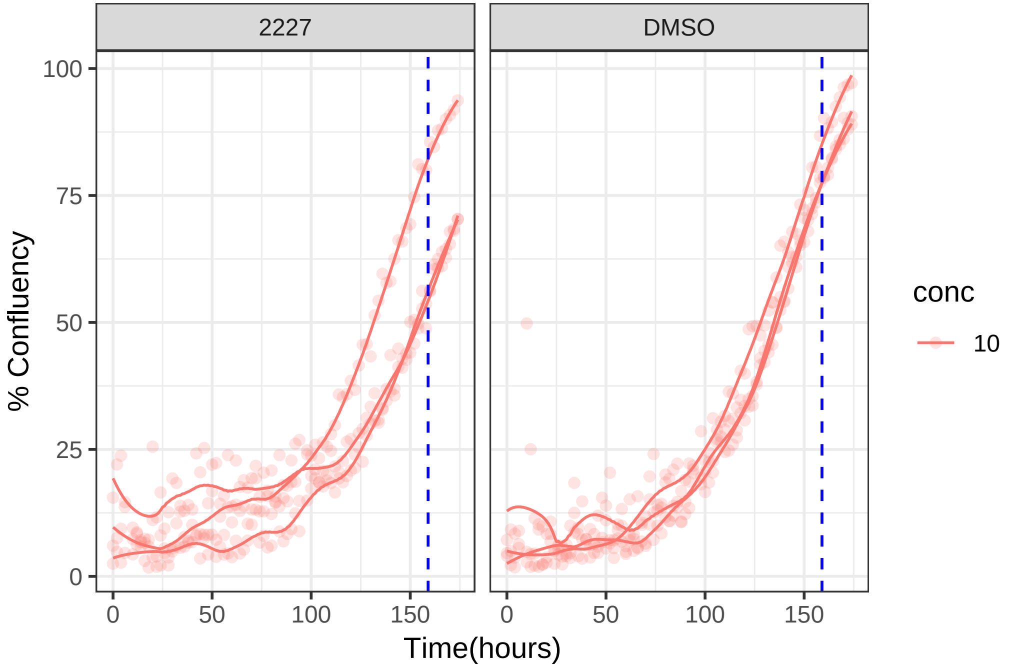

**1 uM 2241 RPE1 hTert**

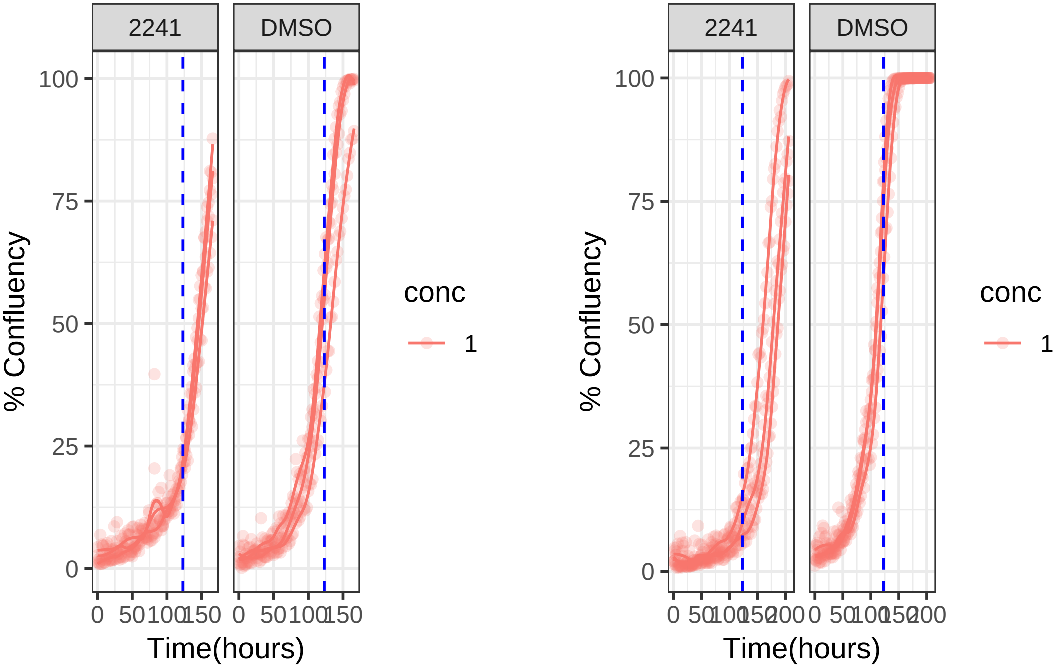

**1 uM 2241 RPE1 Ts12 Ts5**

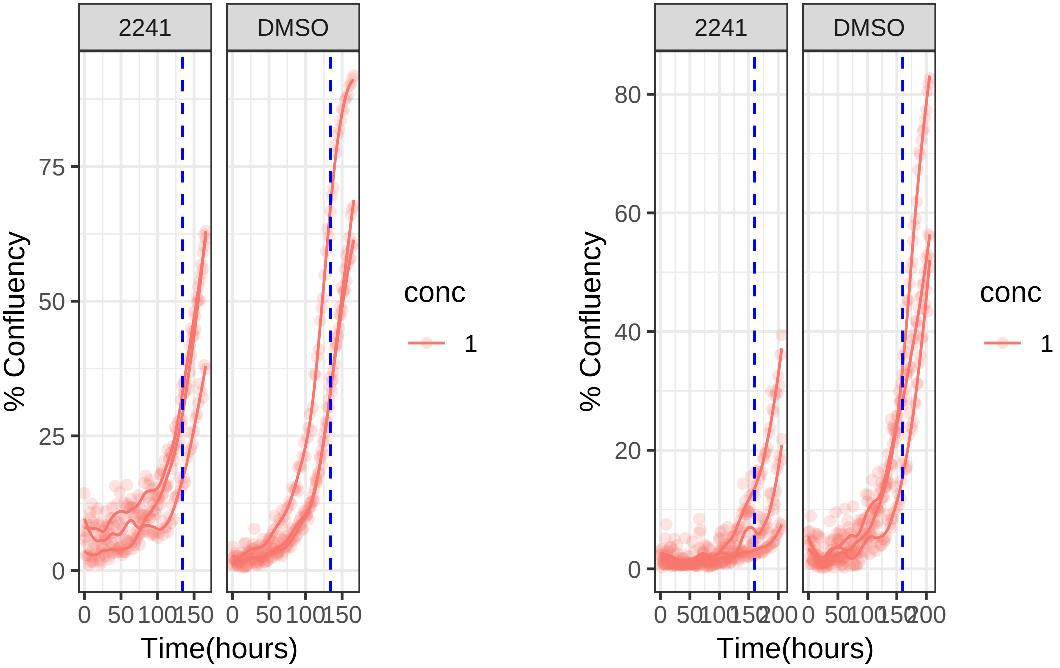

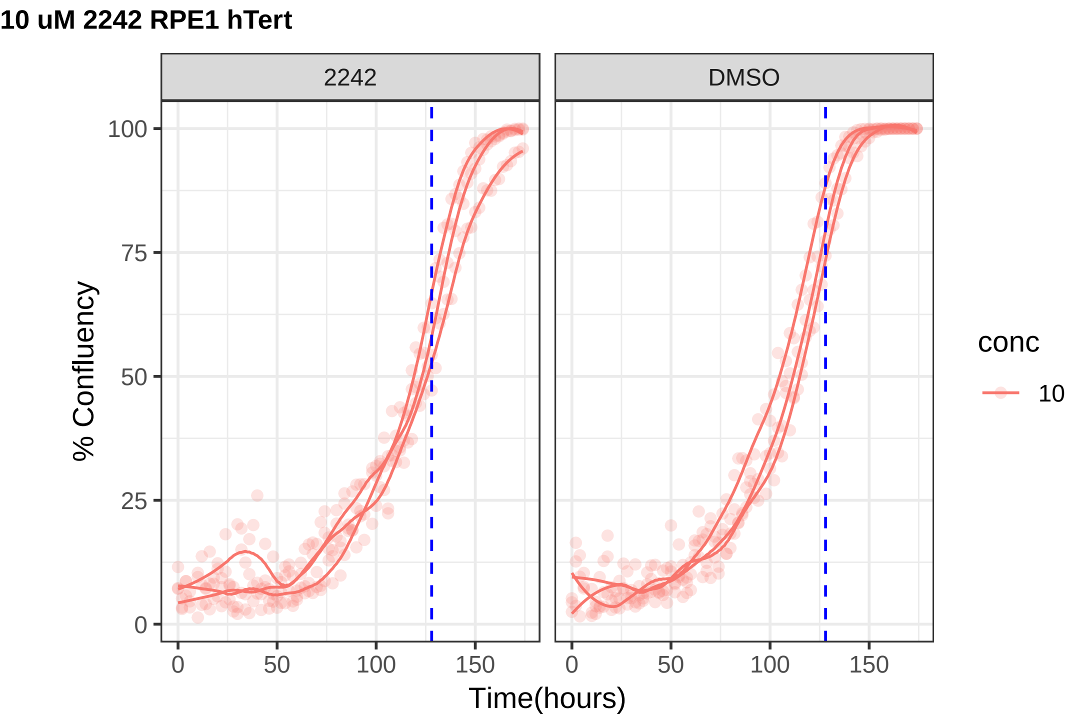
10 uM 2242 RPE1 hTert
2242
DMSO
% Confluency
0
25
50
75
100
0
50
100
150
Time(hours)
conc
10

**10 uM 2242 RPE1 Ts12 Ts5**

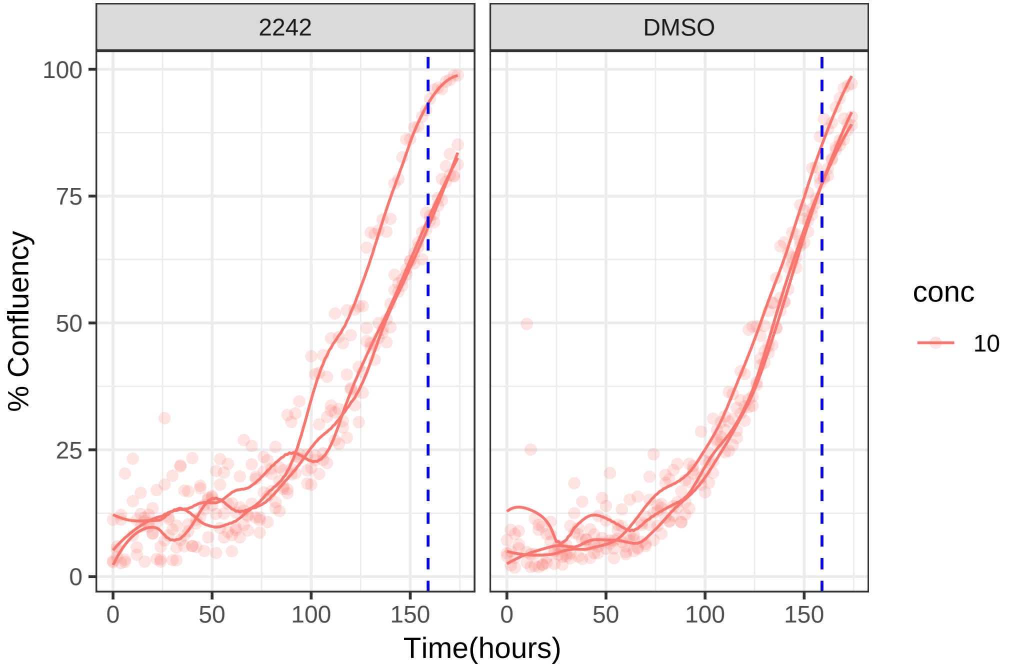

#### 1 uM 2244 RPE1 hTert

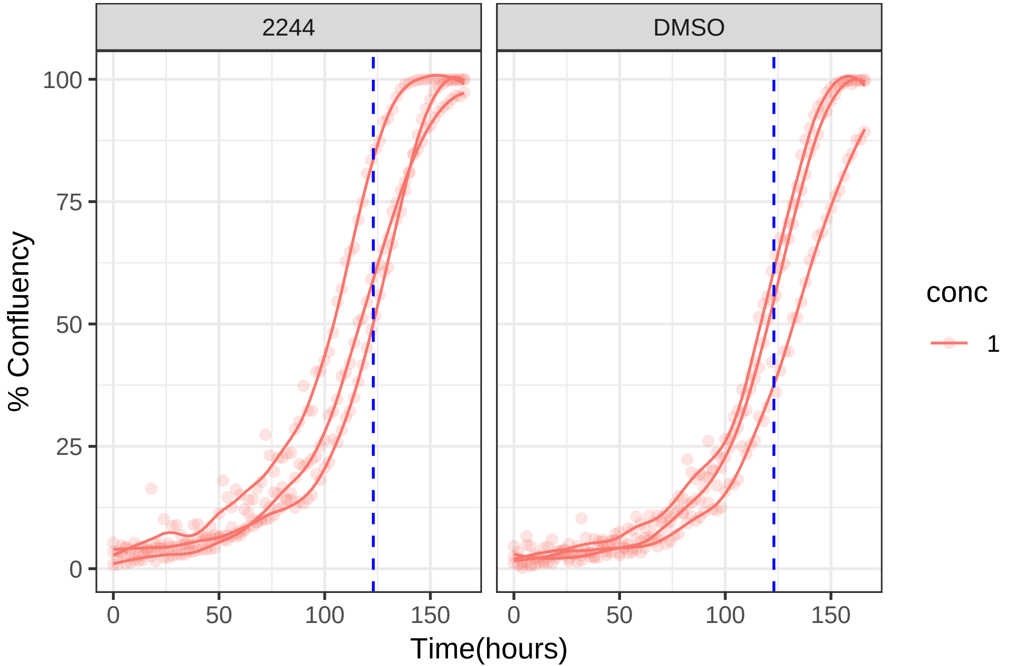

**1 uM 2244 RPE1 Ts12 Ts5**

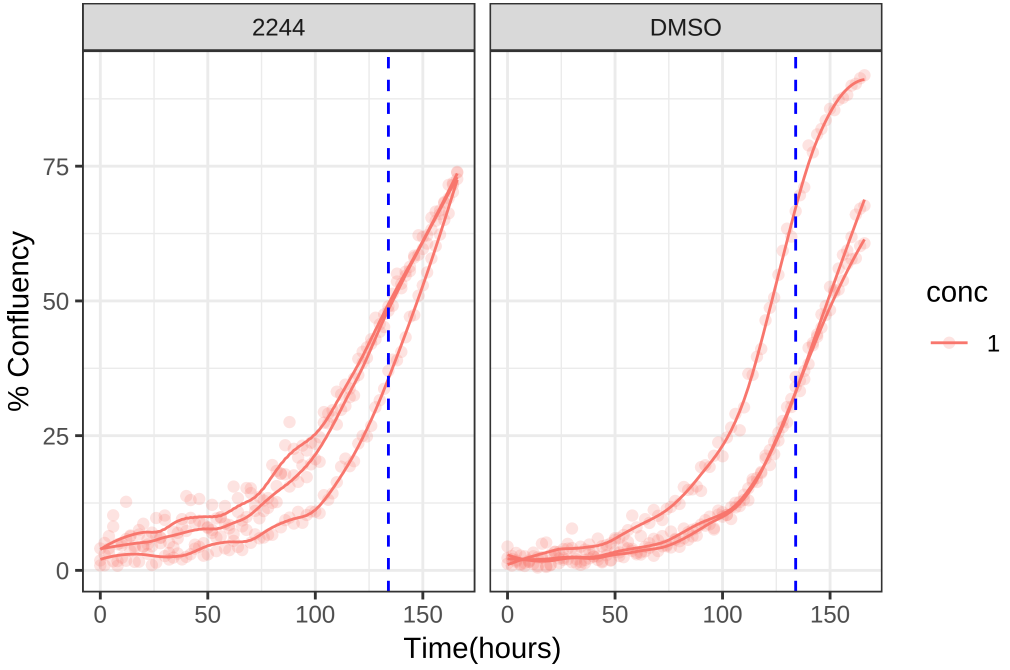

**1 uM 2249 RPE1 hTert**

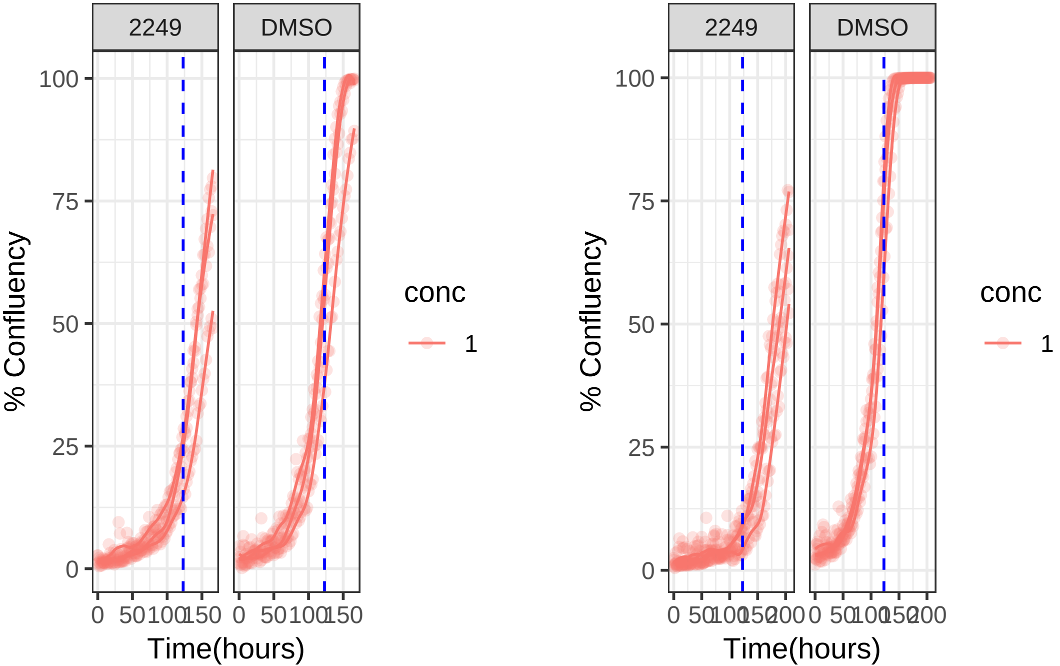

**1 uM 2249 RPE1 Ts12 Ts5**

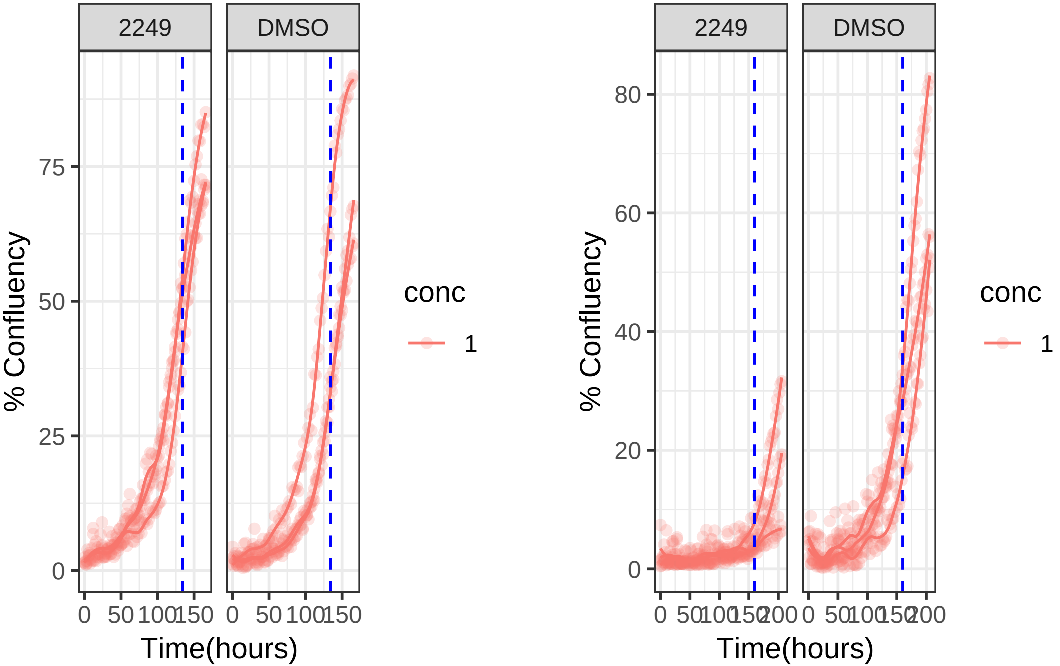

**0.1 uM 2250 RPE1 hTert**

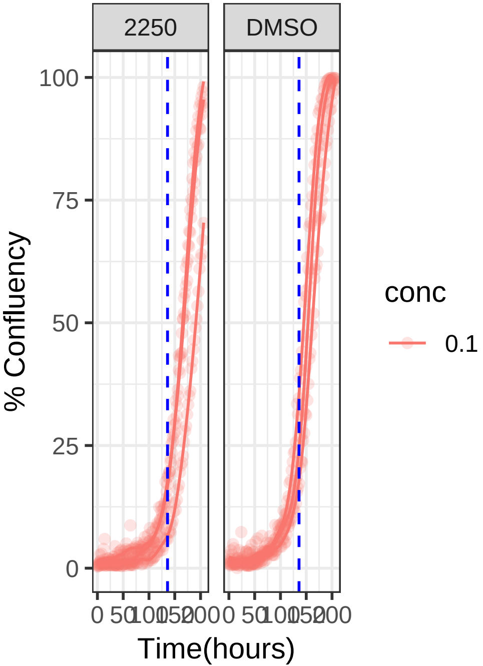


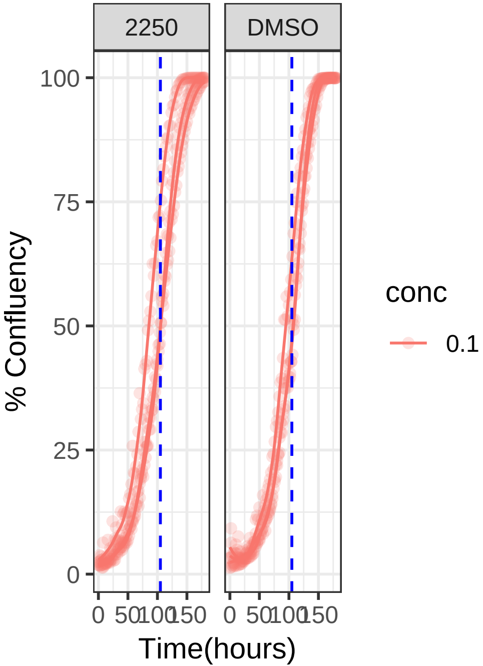

**0.1 uM 2250 RPE1 Ts12 Ts5**

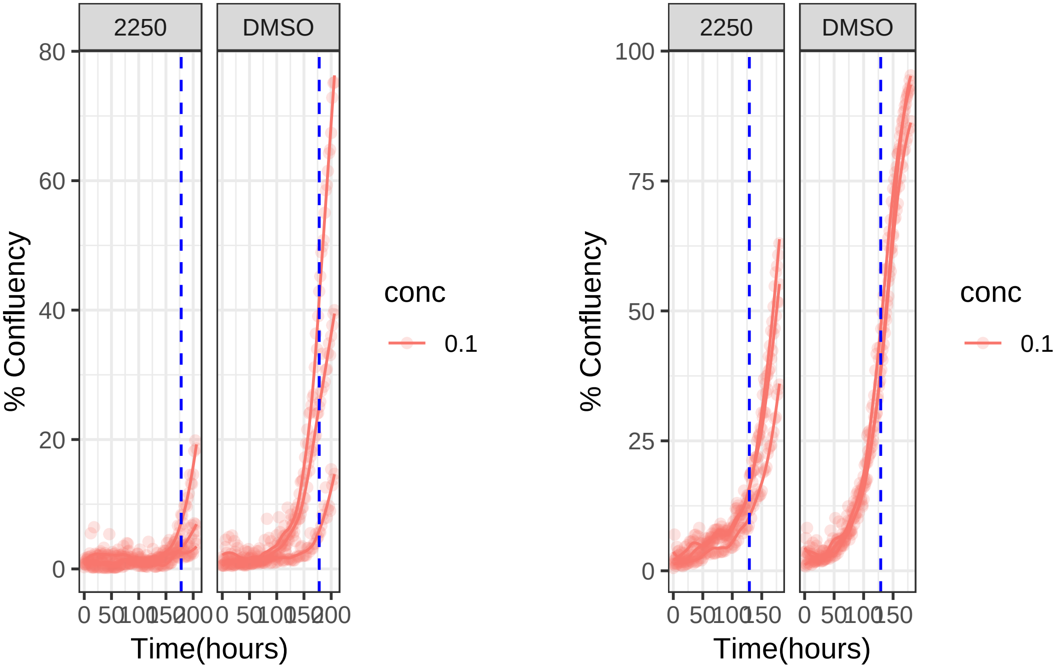

**10 uM 2262 RPE1 hTert**

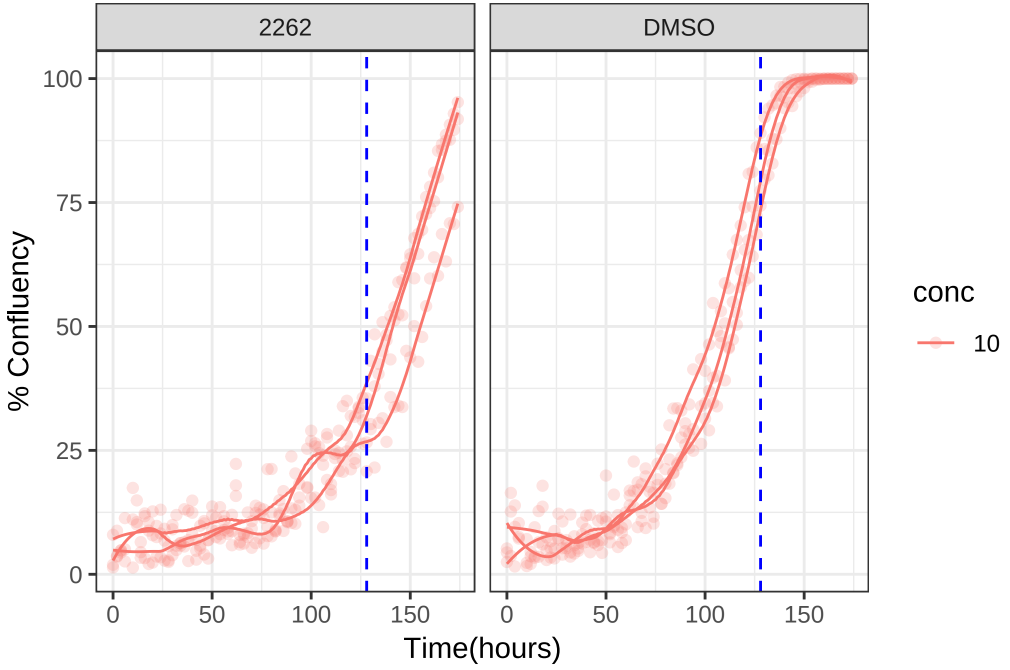

**10 uM 2262 RPE1 Ts12 Ts5**

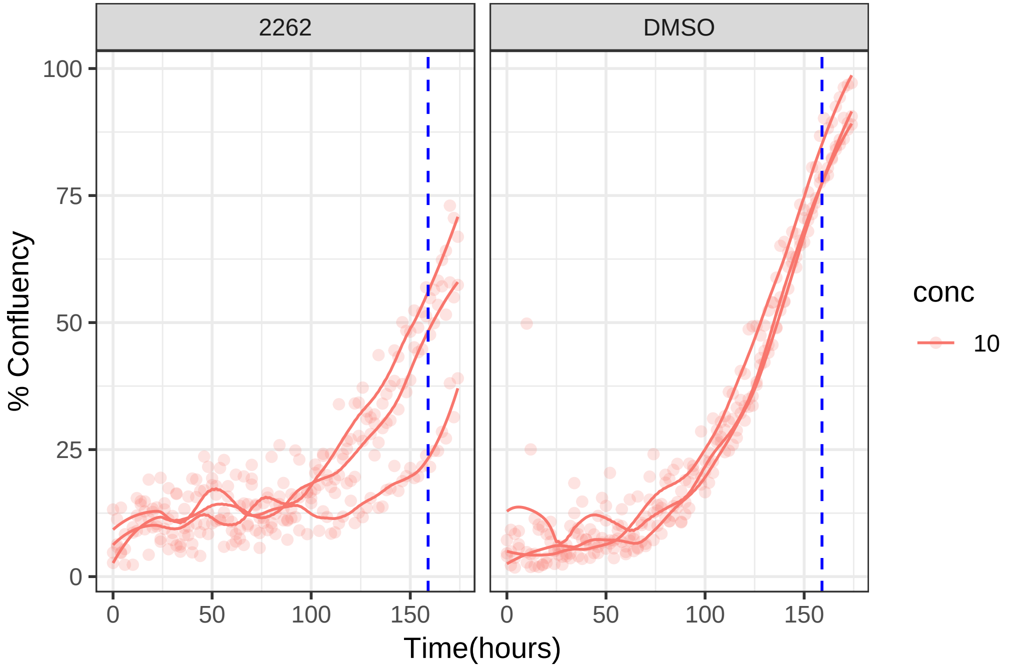

**10 uM 2268 RPE1 hTert**

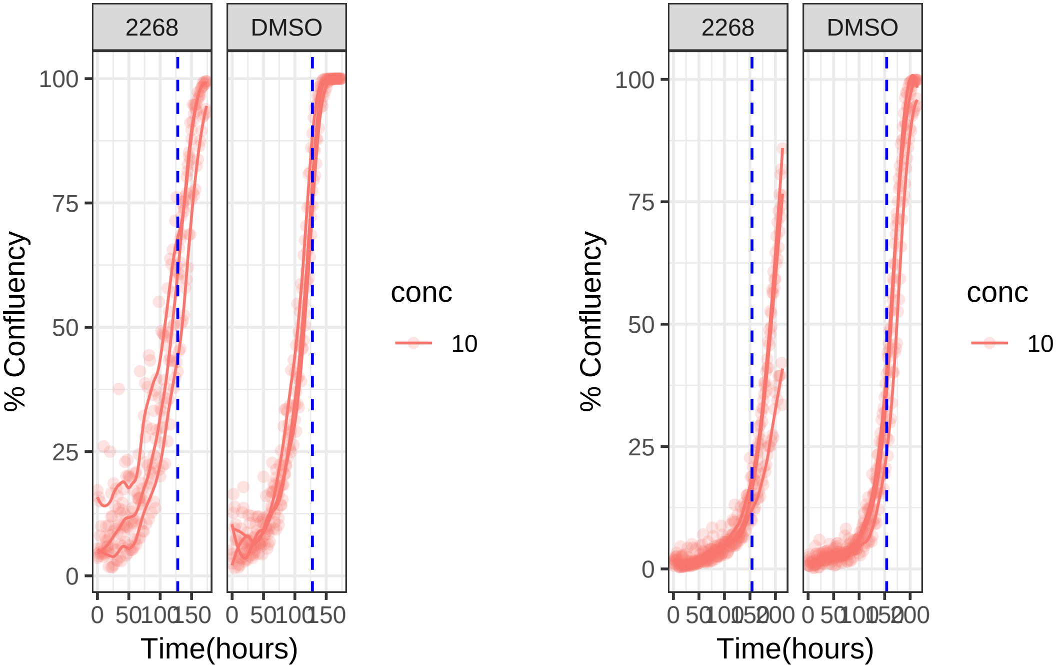

### 10 uM 2268 RPE1 Ts12 Ts5

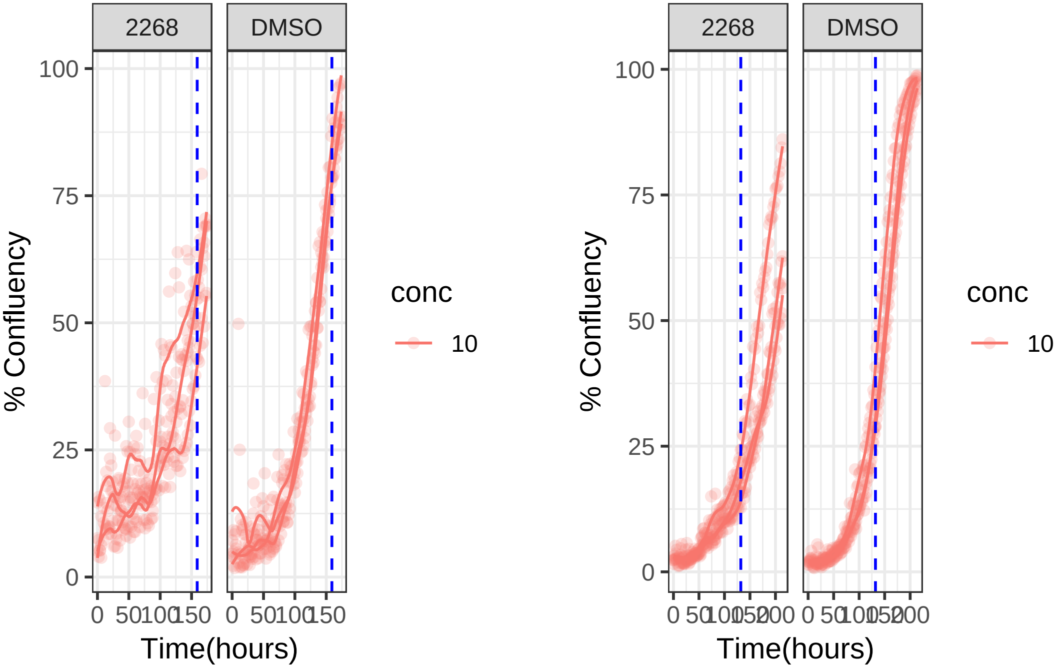

**10 uM 2278 RPE1 hTert**

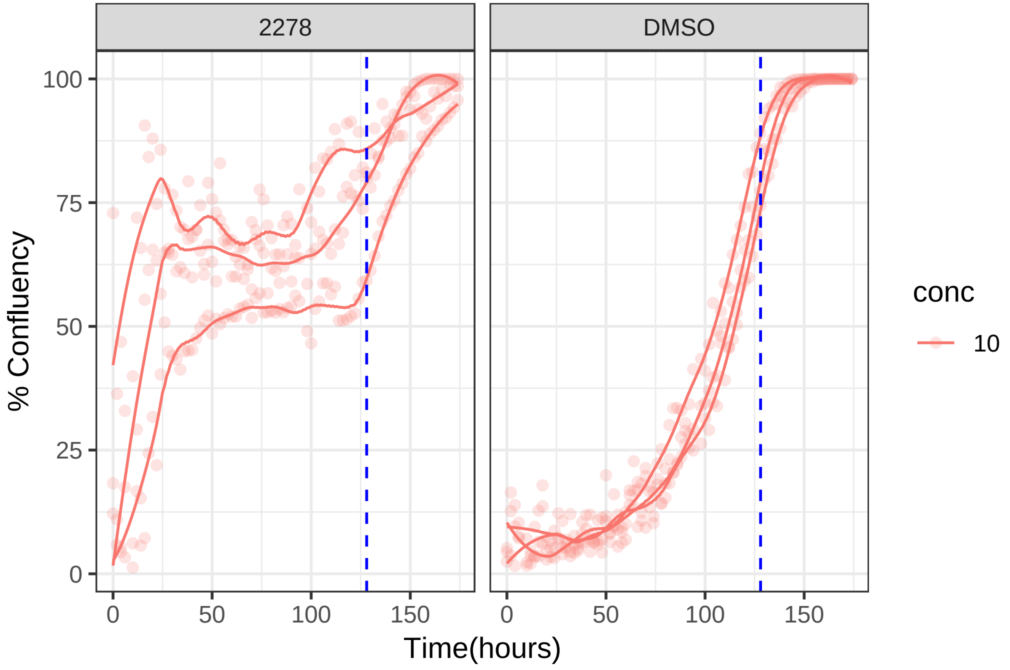

**10 uM 2278 RPE1 Ts12 Ts5**

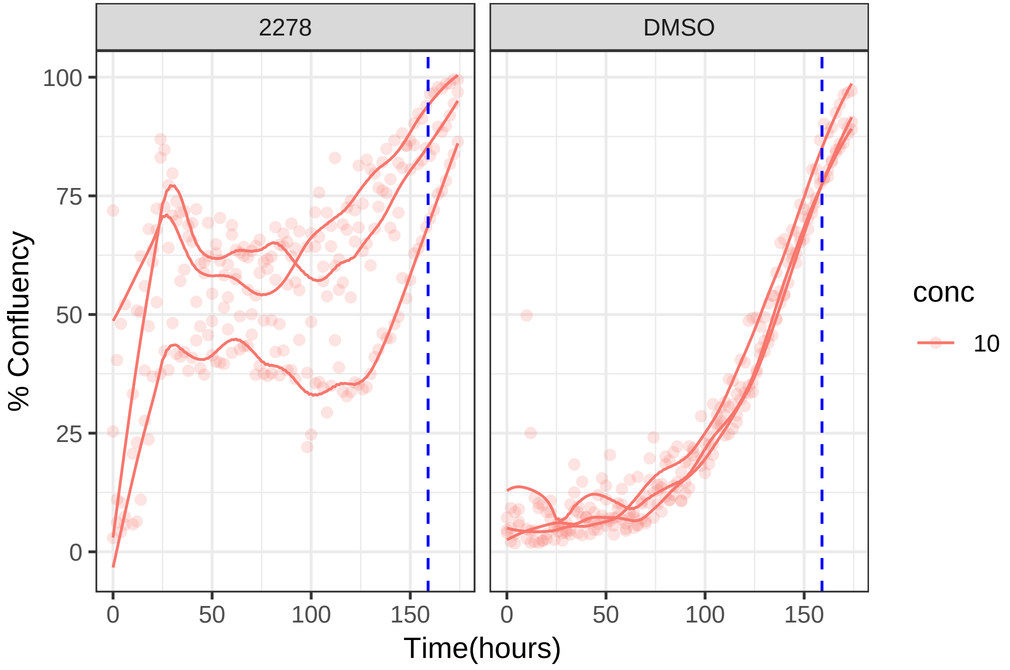

**10 uM 2293 RPE1 hTert**

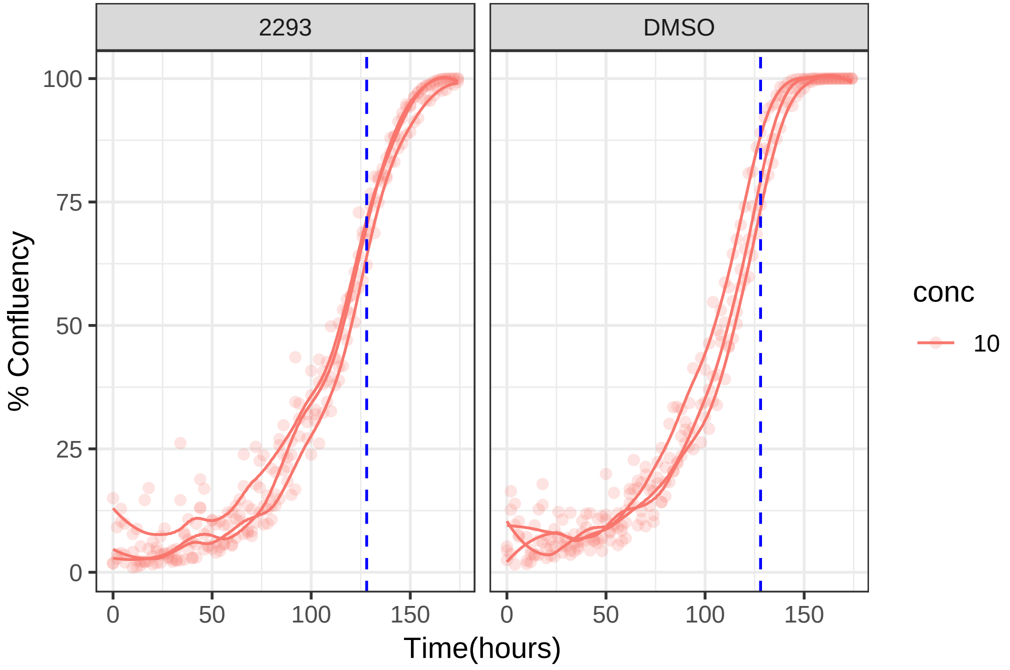

**10 uM 2293 RPE1 Ts12 Ts5**

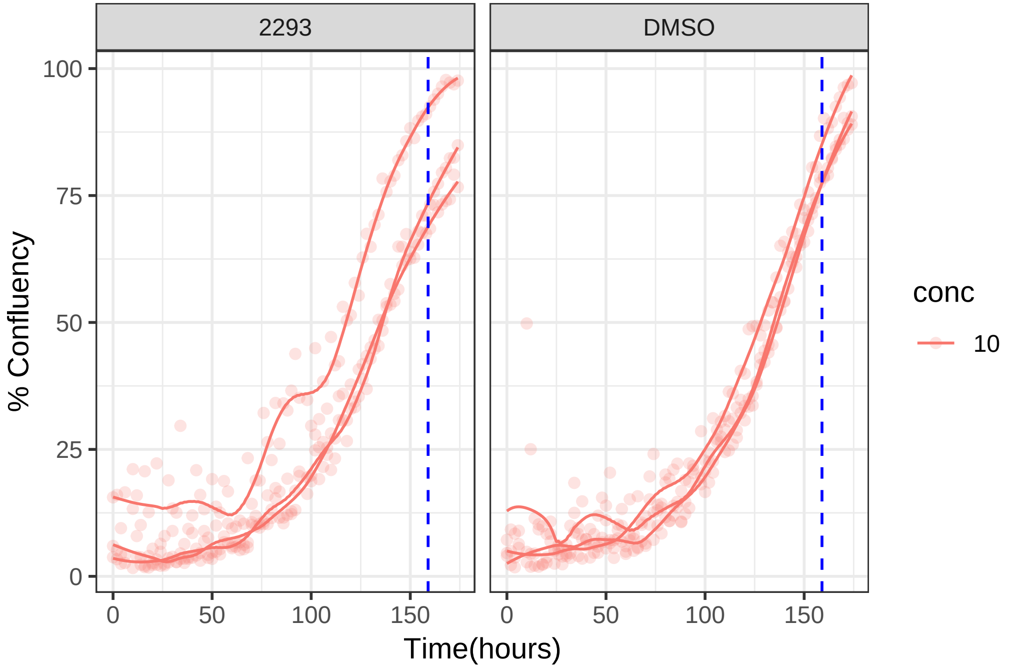

**1 uM 2300 RPE1 hTert**

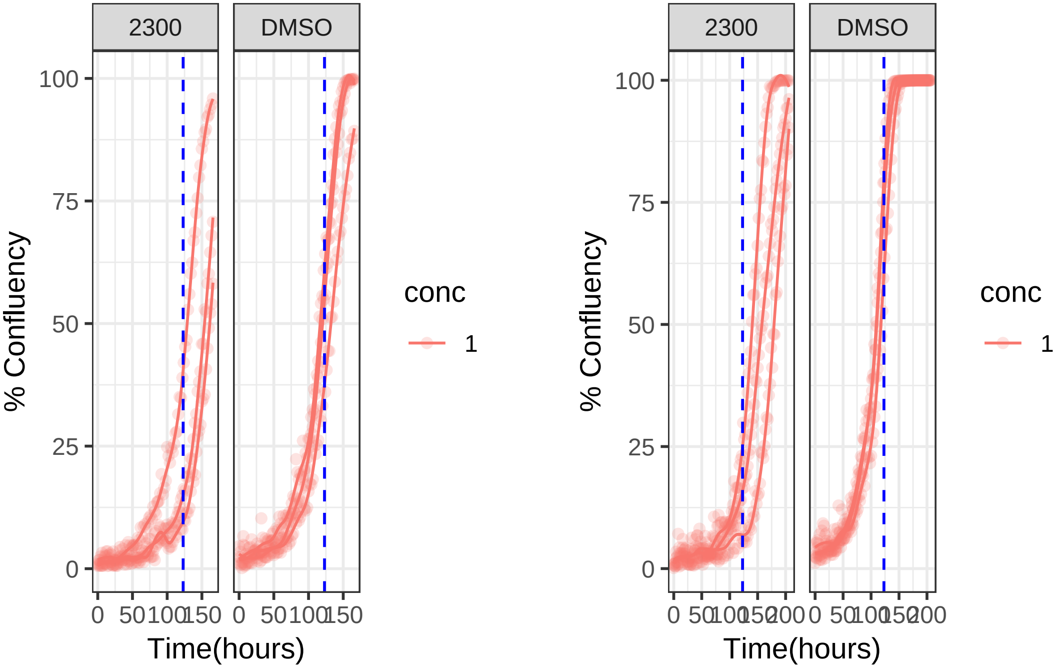

1 uM 2300 RPE1 Ts12 Ts5

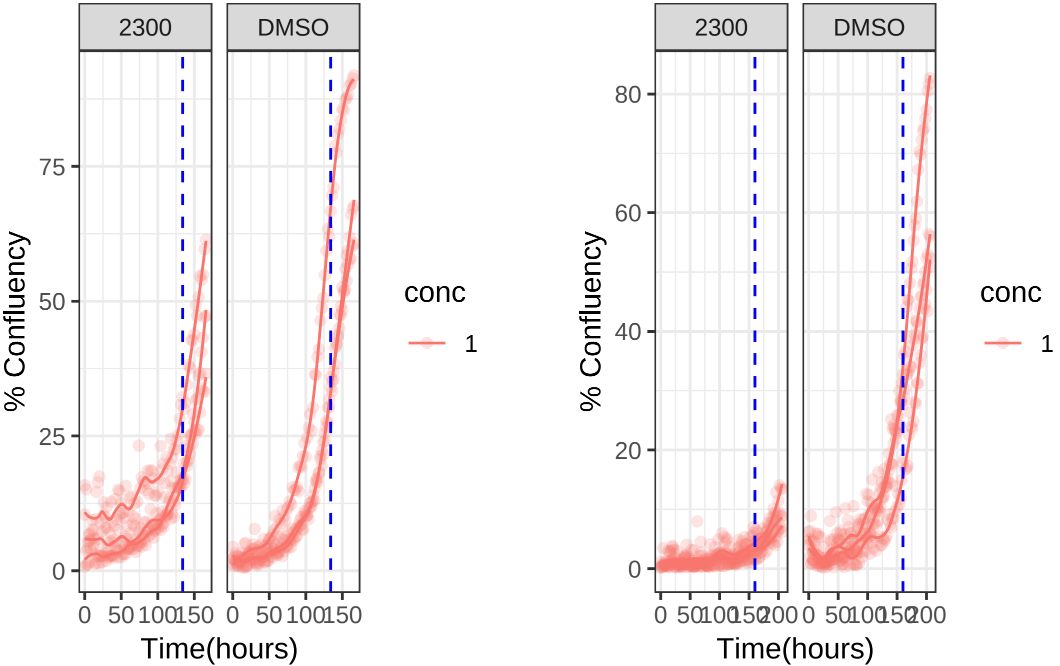

**10 uM 2319 RPE1 hTert**

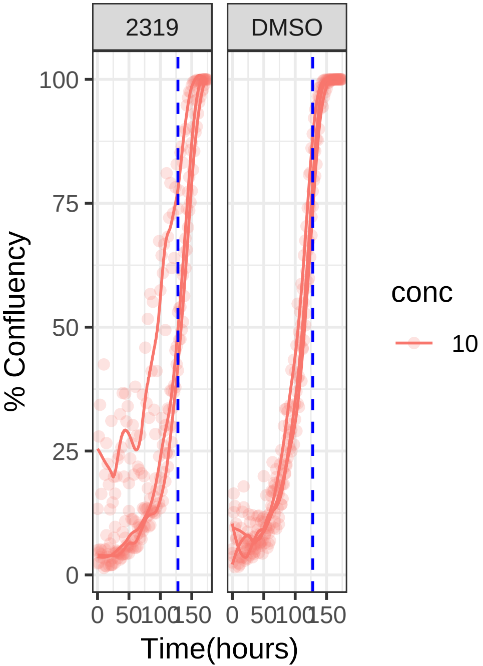


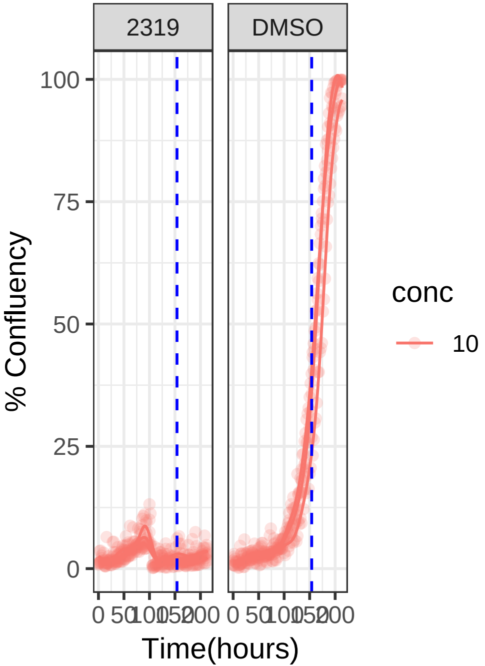

**1 uM 2319 RPE1 hTert**

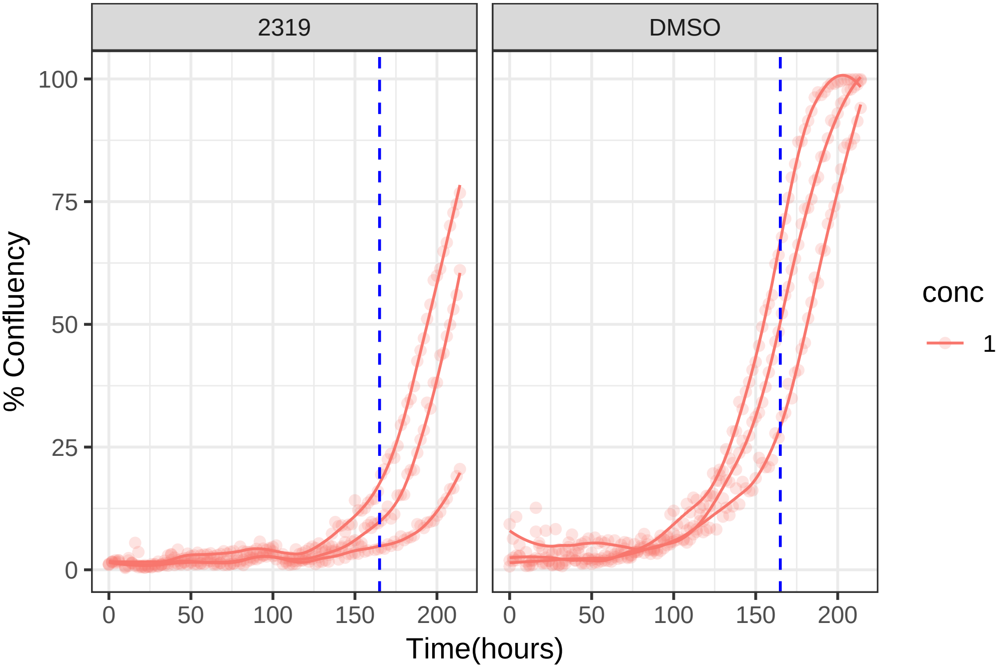

**10 uM 2319 RPE1 Ts12 Ts5**

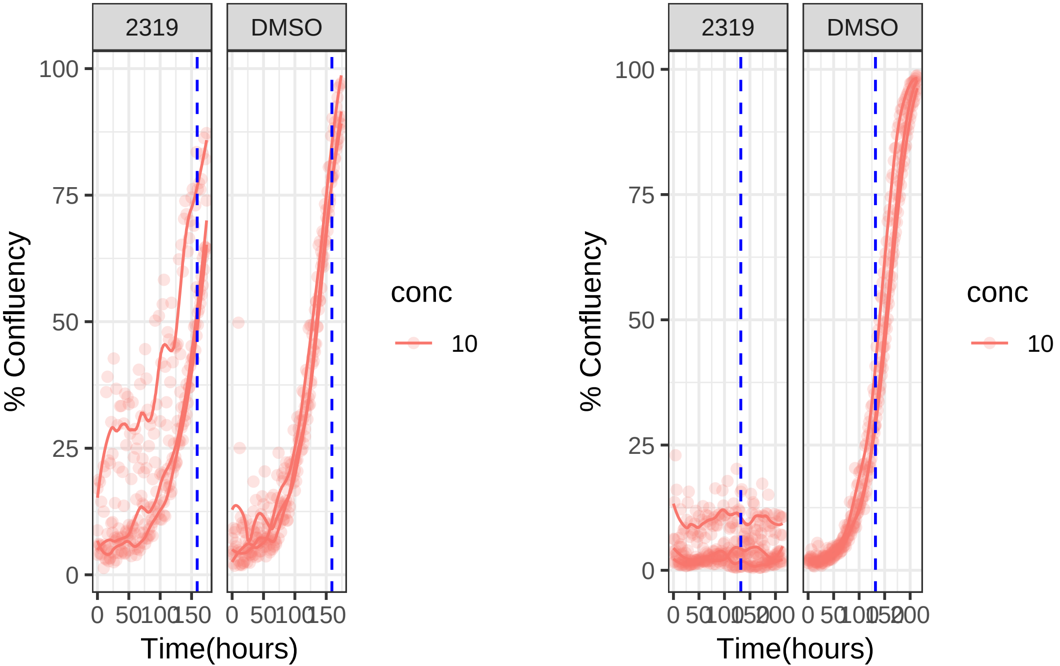

**1 uM 2319 RPE1 Ts12 Ts5**

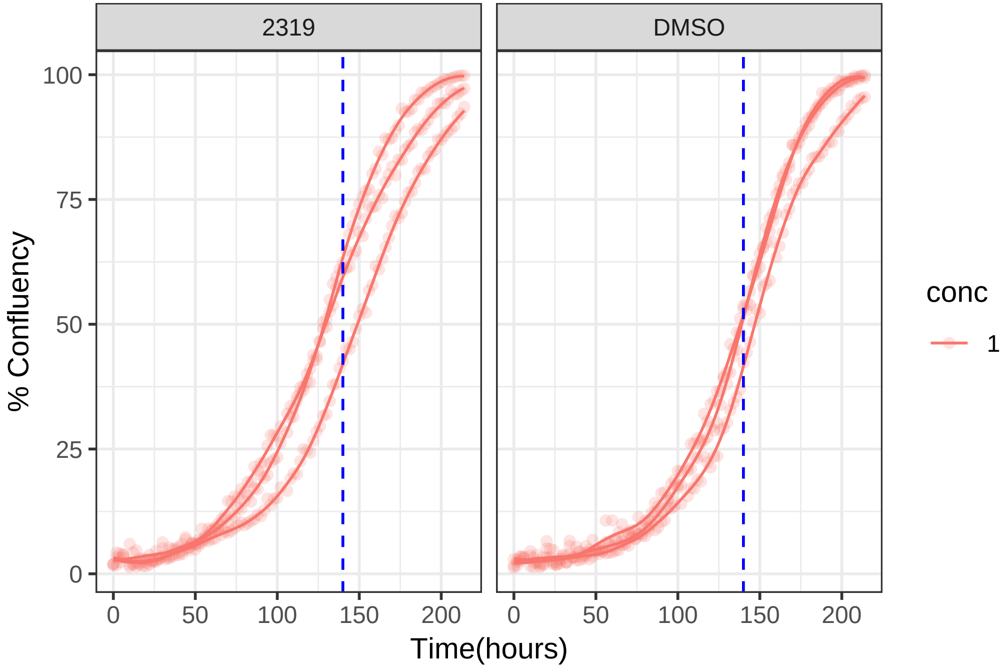

**1 uM 2323 RPE1 hTert**

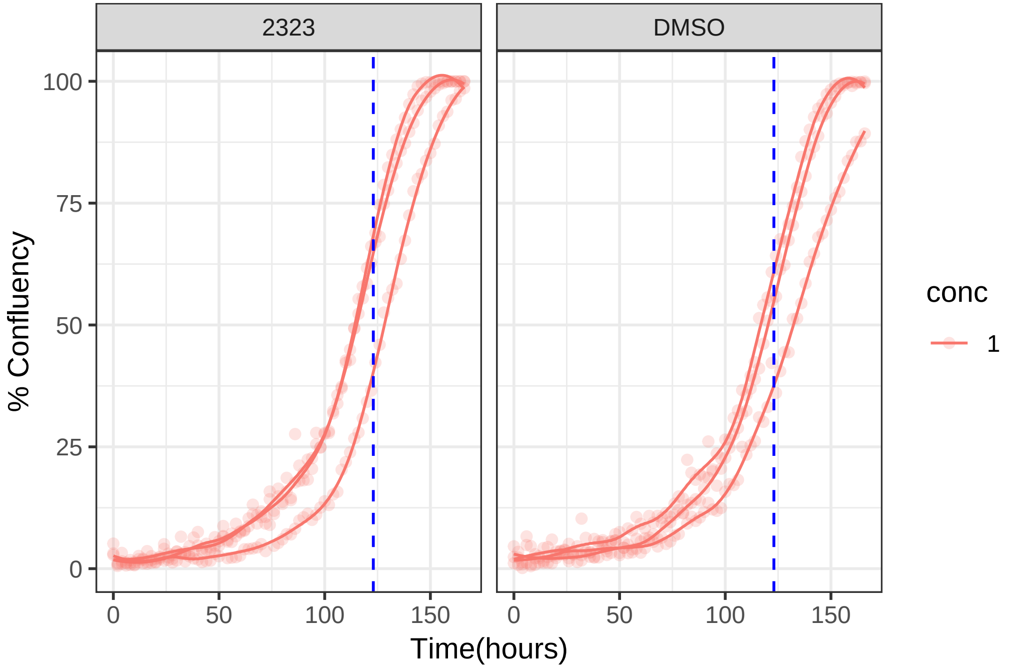

1 uM 2323 RPE1 Ts12 Ts5

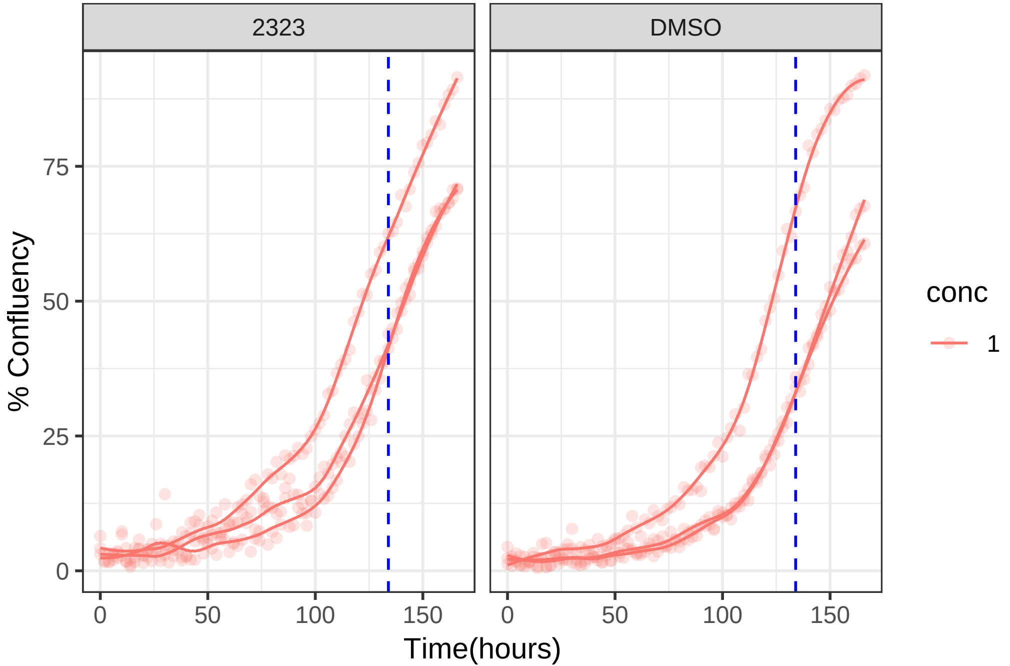

**10 uM 2325 RPE1 hTert**

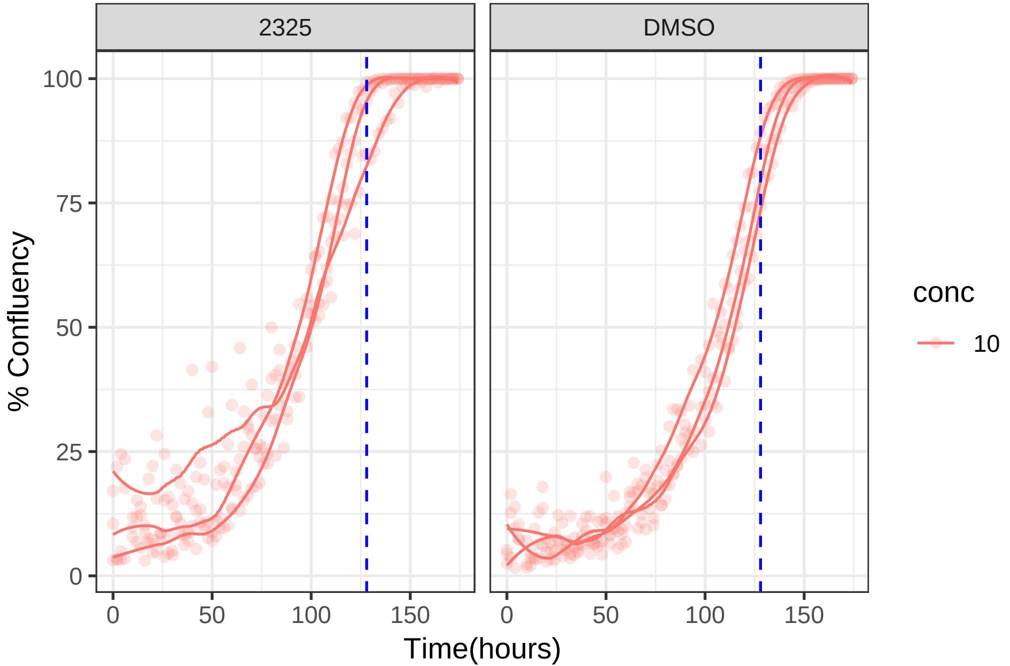

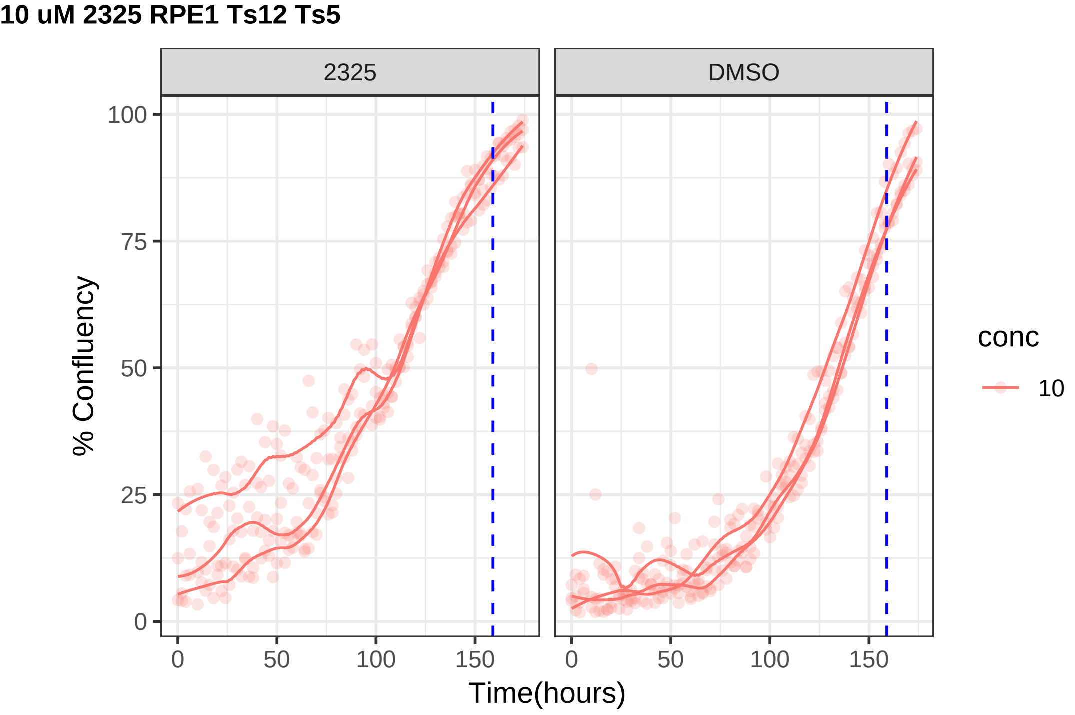
10 uM 2325 RPE1 Ts12 Ts5
2325
DMSO
% Confluency
0
25
50
75
100
0
50
100
150
Time(hours)
conc
10

**10 uM 2326 RPE1 hTert**

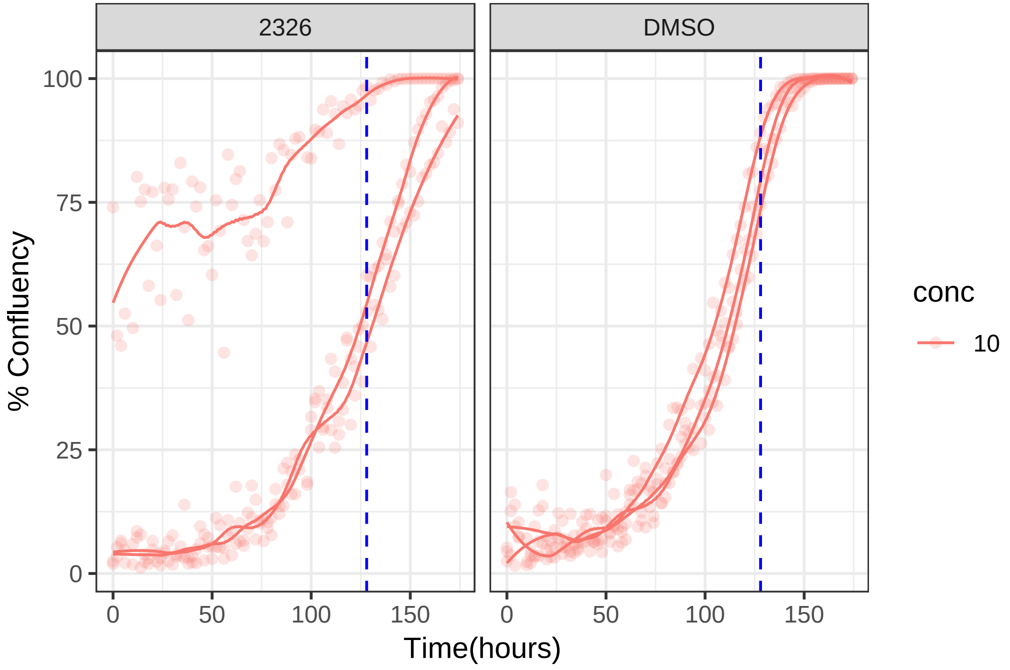

**10 uM 2326 RPE1 Ts12 Ts5**

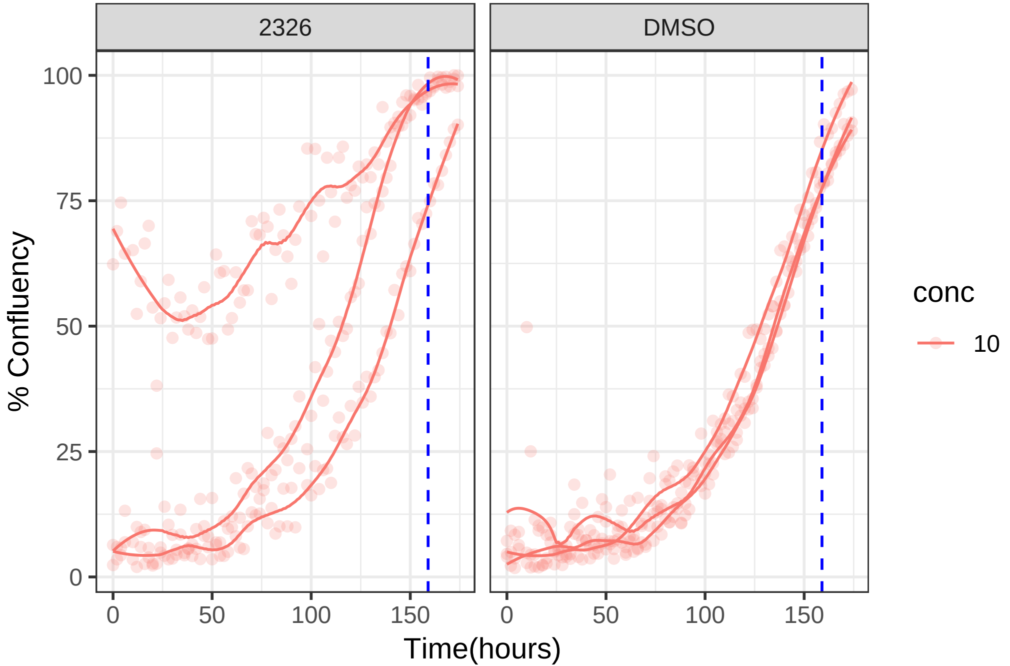

**10 uM 2327 RPE1 hTert**

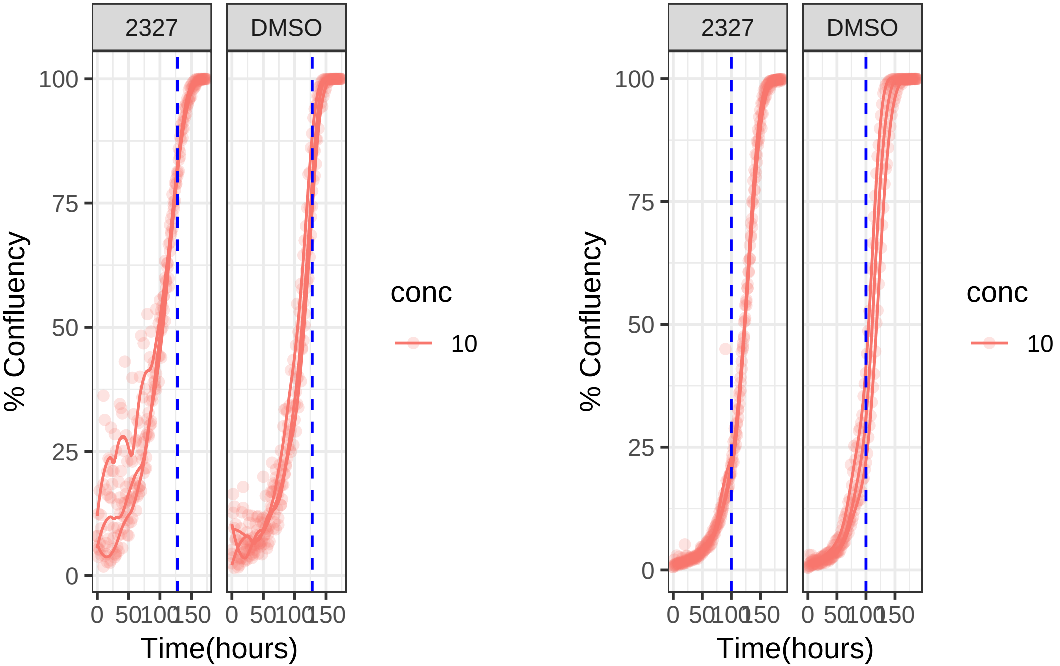

**10 uM 2327 RPE1 Ts12 Ts5**

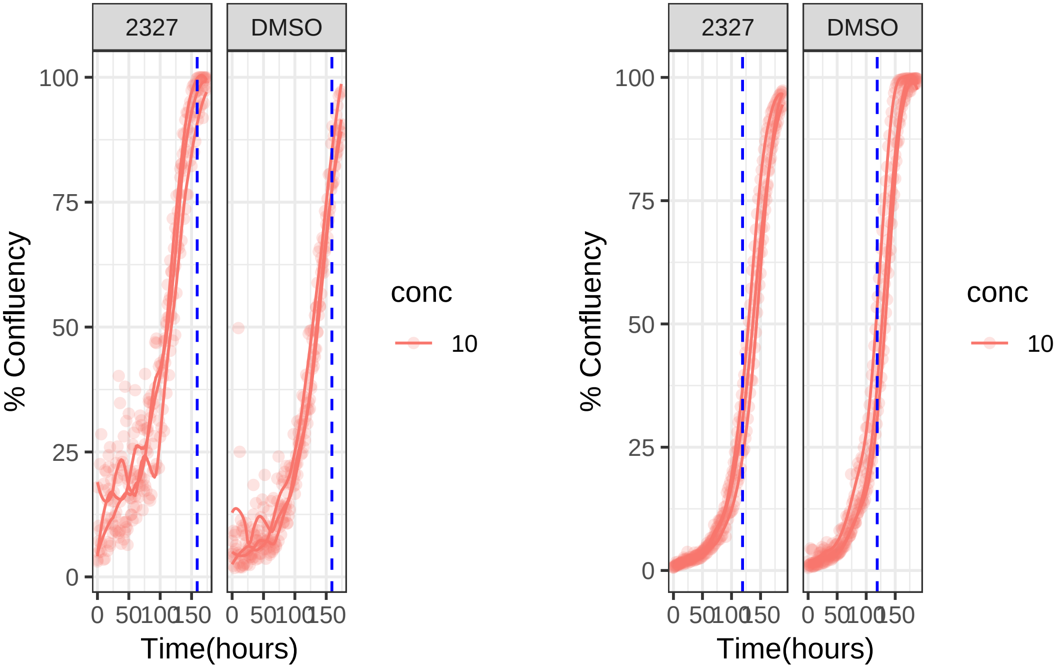

**10 uM 2339 RPE1 hTert**

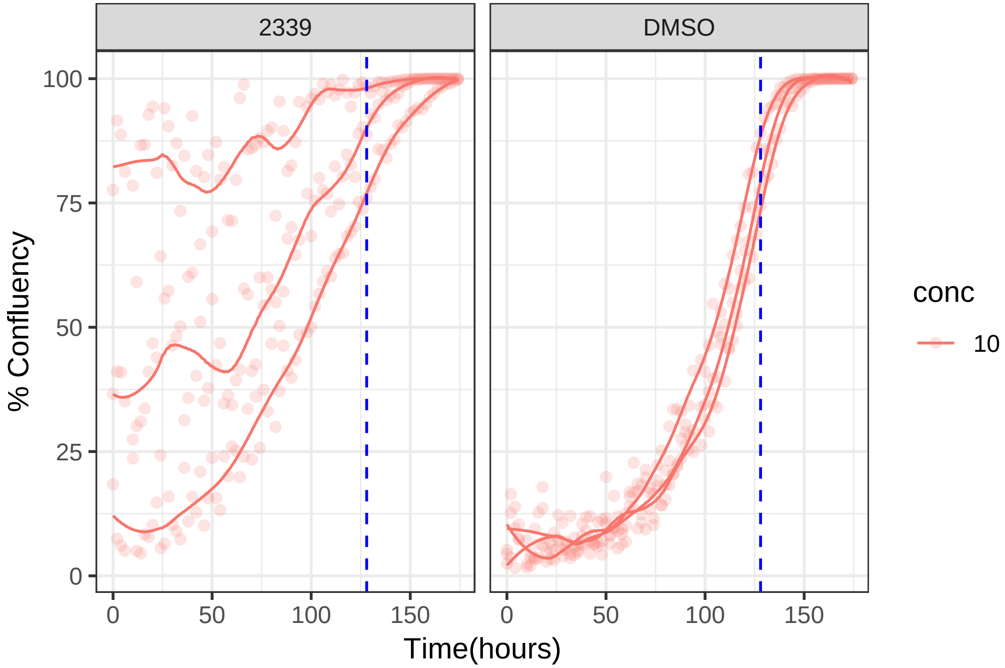

10 uM 2339 RPE1 Ts12 Ts5

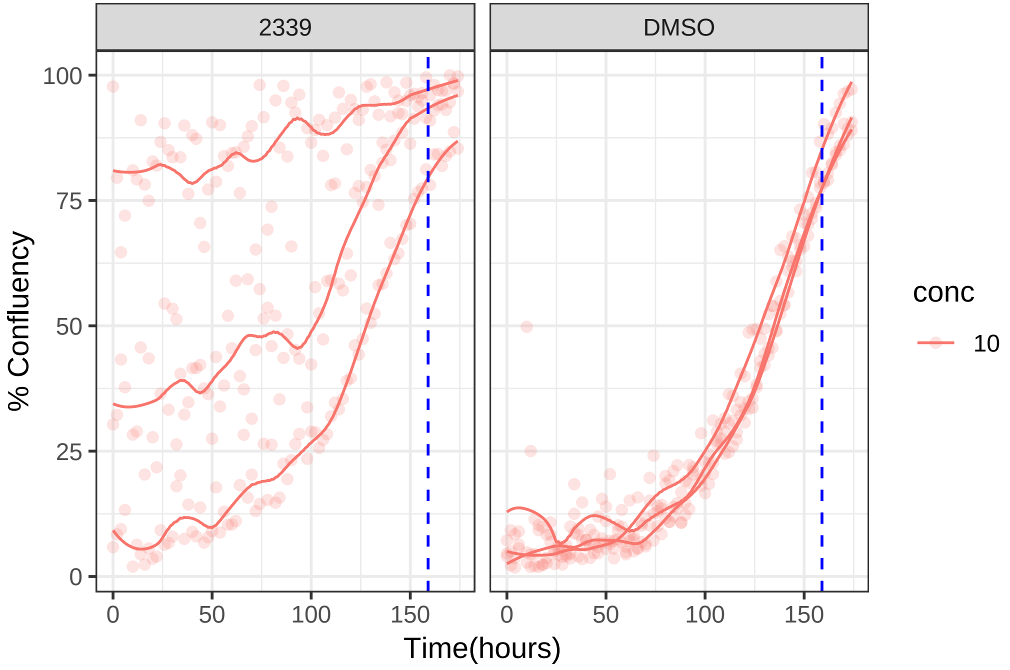

**0.1 uM 2347 RPE1 hTert**

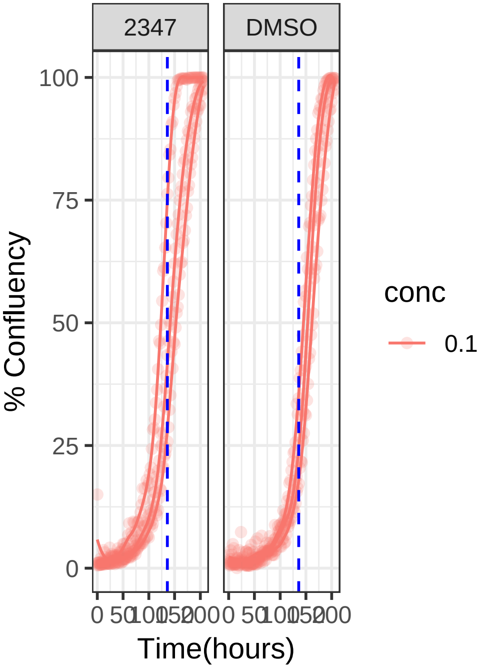


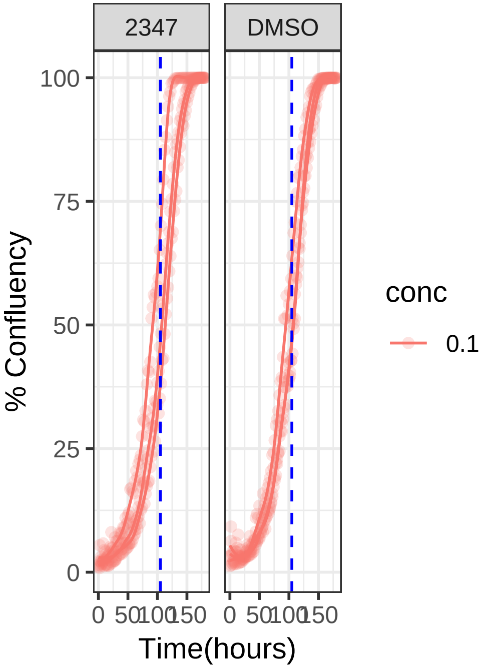

**0.1 uM 2347 RPE1 Ts12 Ts5**

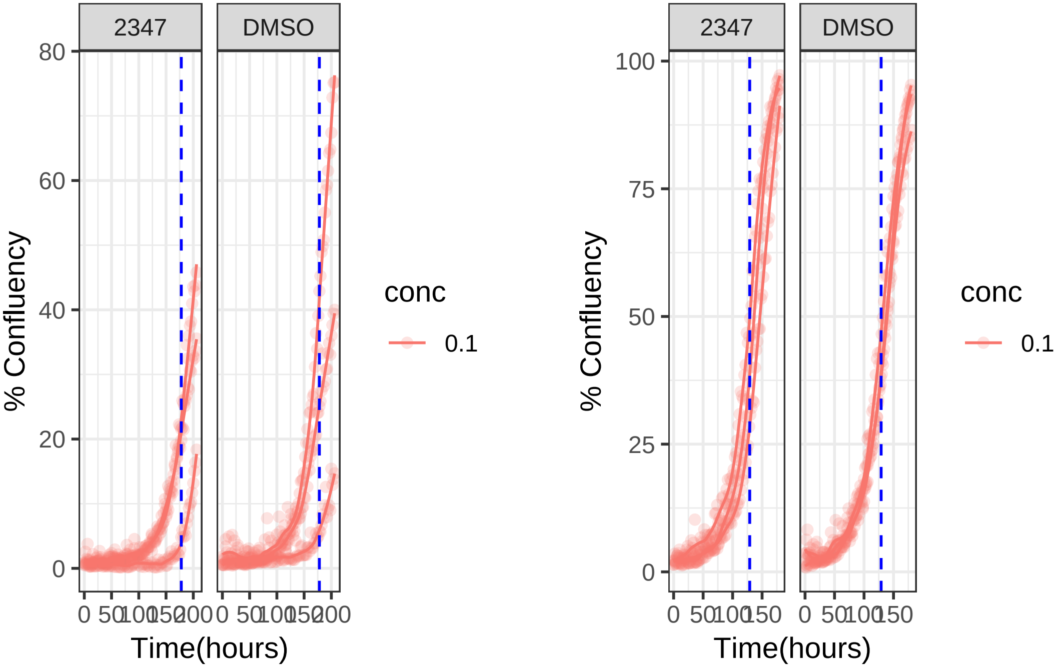

**10 uM 2358 RPE1 hTert**

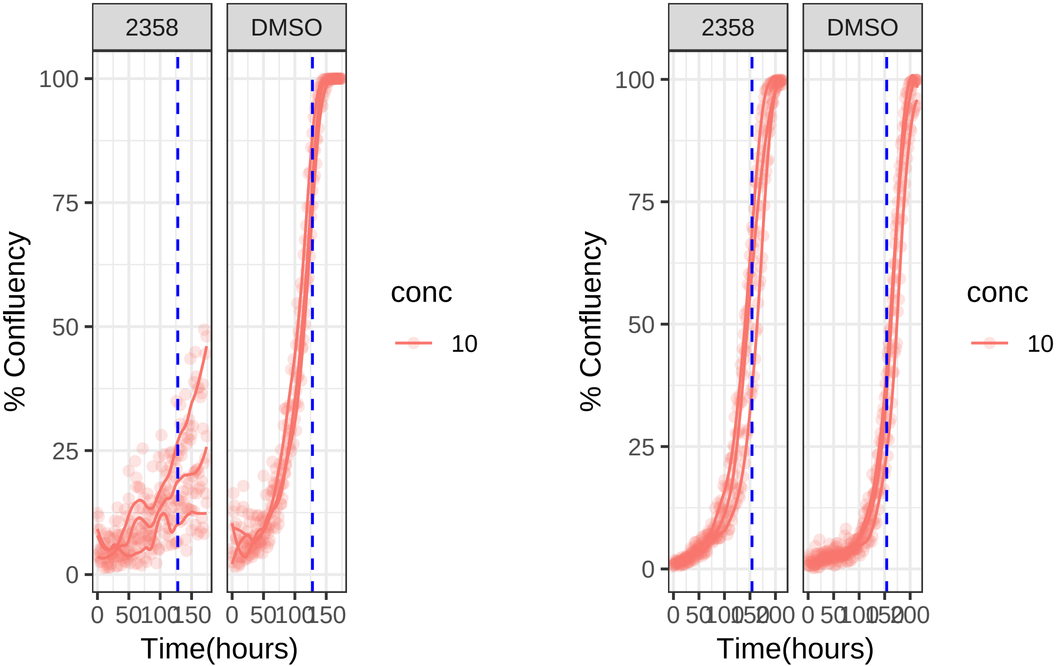

**10 uM 2358 RPE1 Ts12 Ts5**

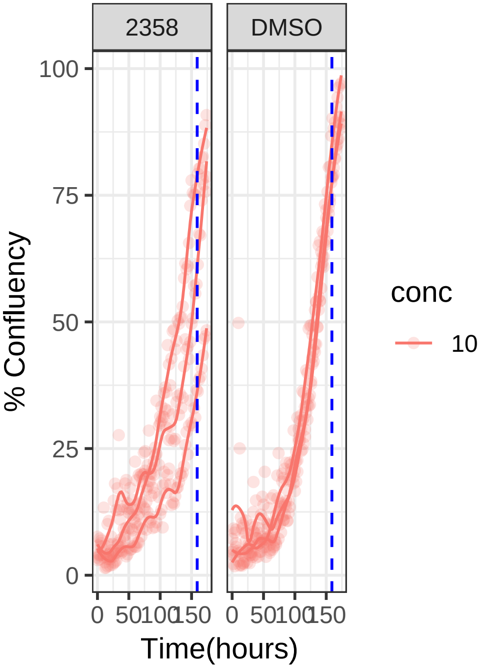


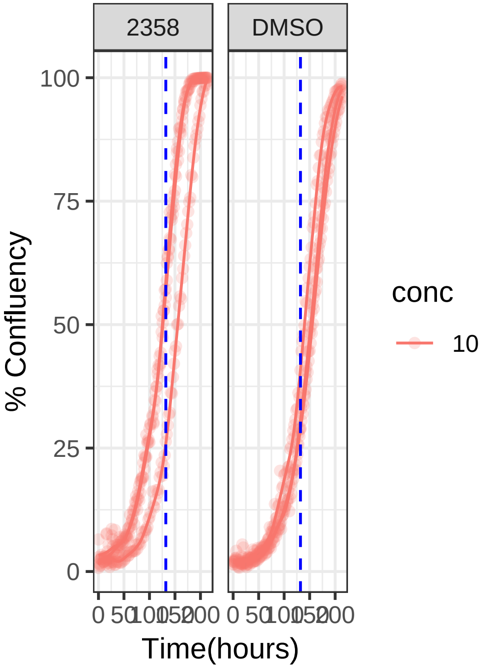

**1 uM 2359 RPE1 hTert**

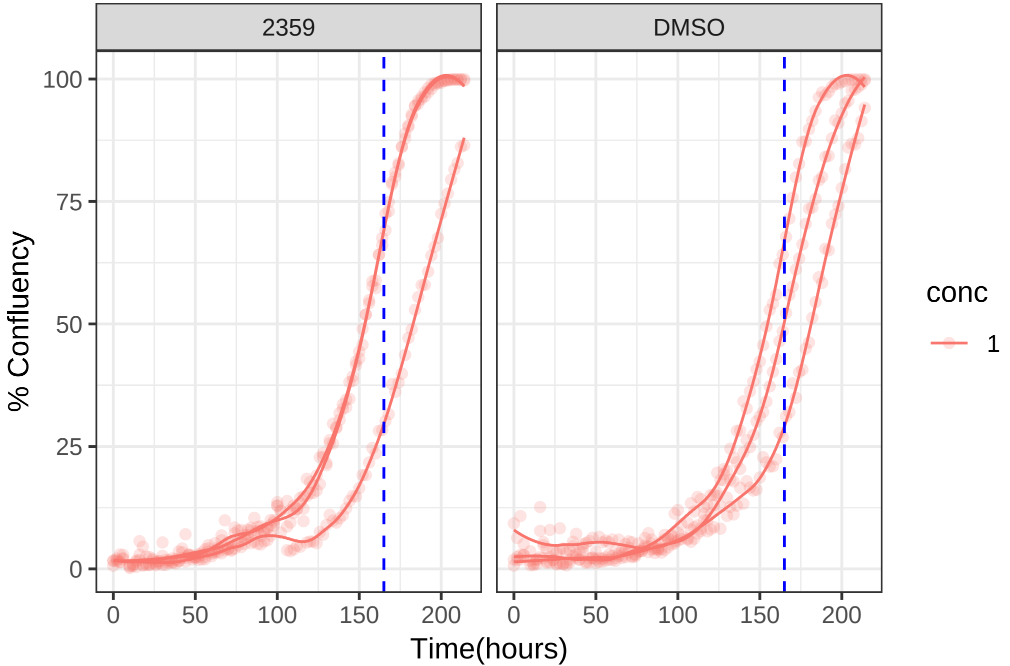

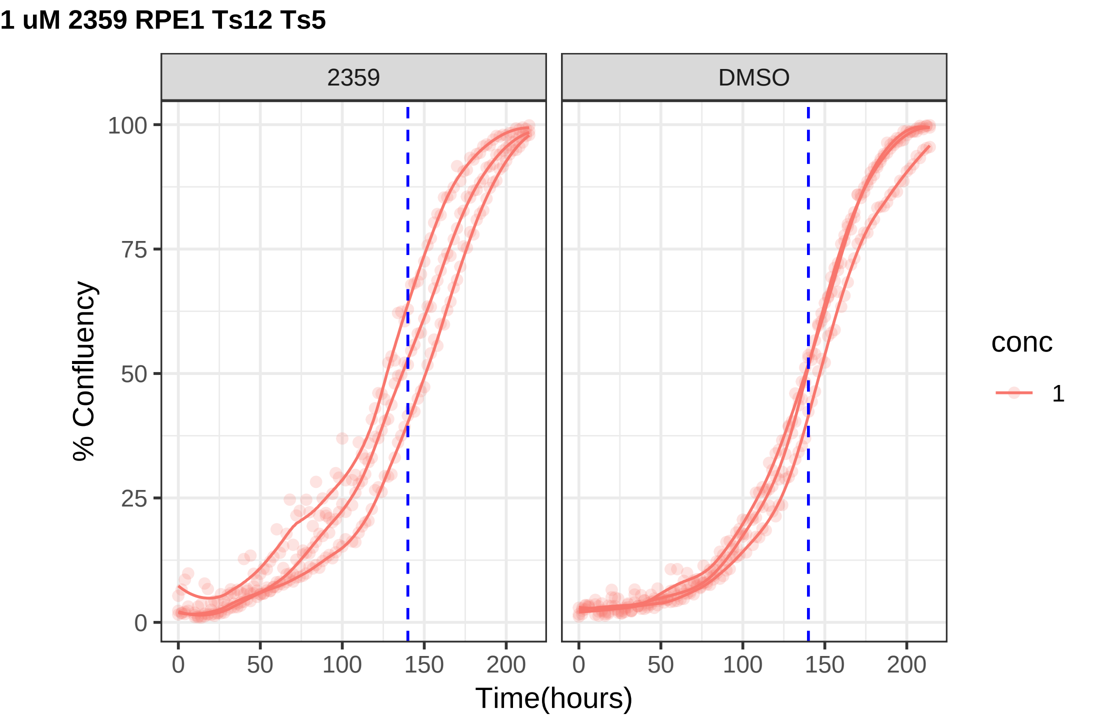
1 uM 2359 RPE1 Ts12 Ts5
2359
DMSO
% Confluency
Time(hours)
0
25
50
75
100
0
50
100
150
200
conc
1

**10 uM 2369 RPE1 hTert**

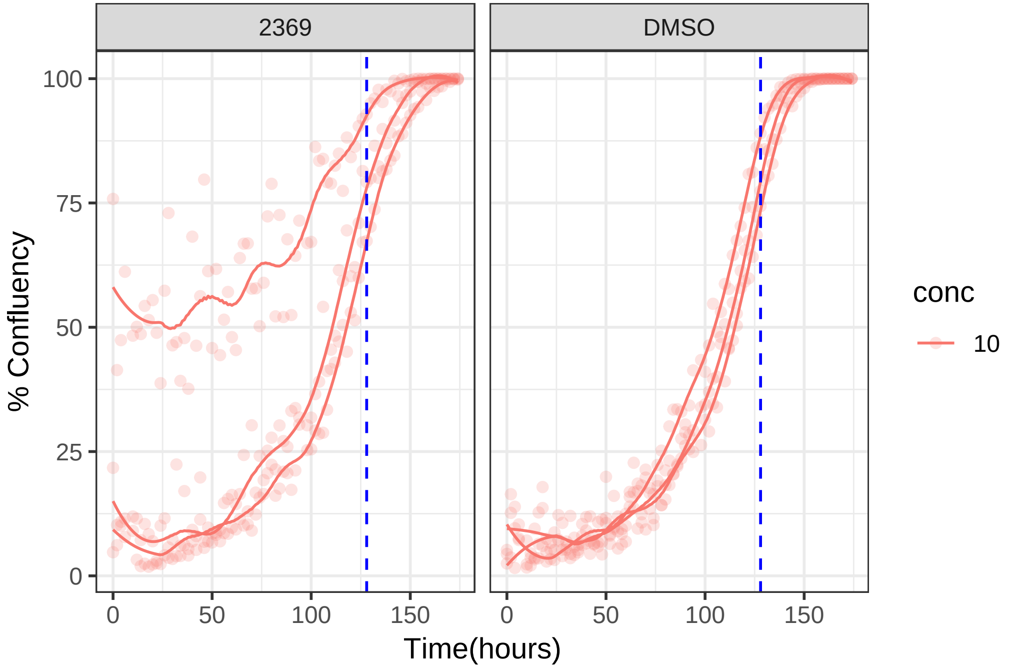

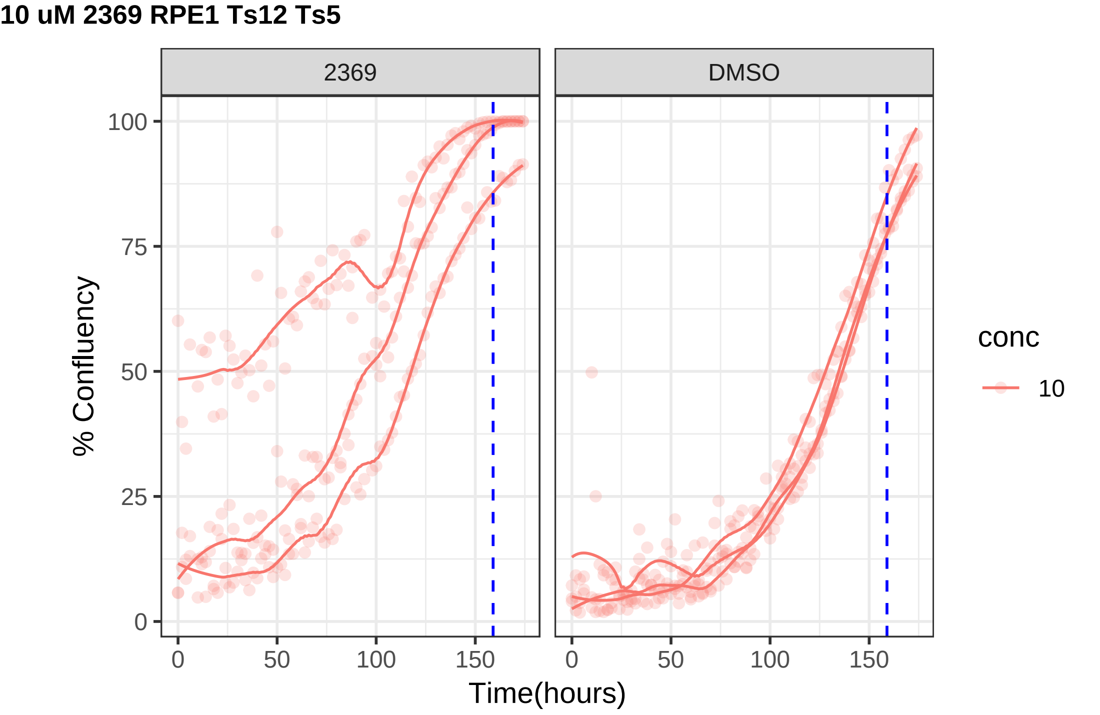

10 uM 2369 RPE1 Ts12 Ts5
2369
DMSO
% Confluency
0
25
50
75
100
0
50
100
150
Time(hours)
conc
10

**10 uM 2375 RPE1 hTert**

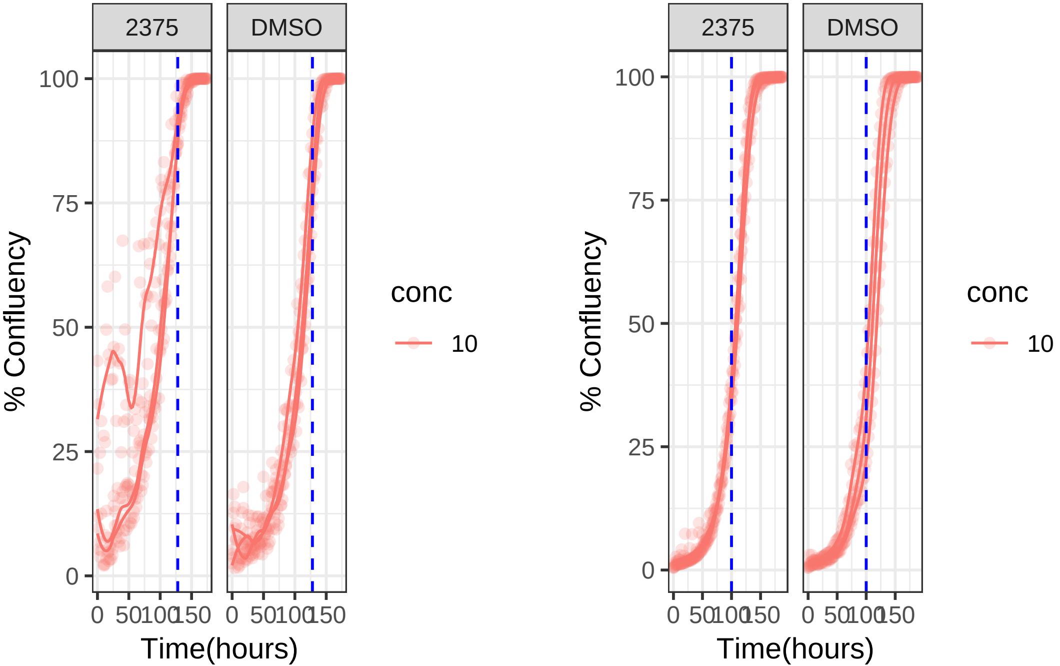

**10 uM 2375 RPE1 Ts12 Ts5**

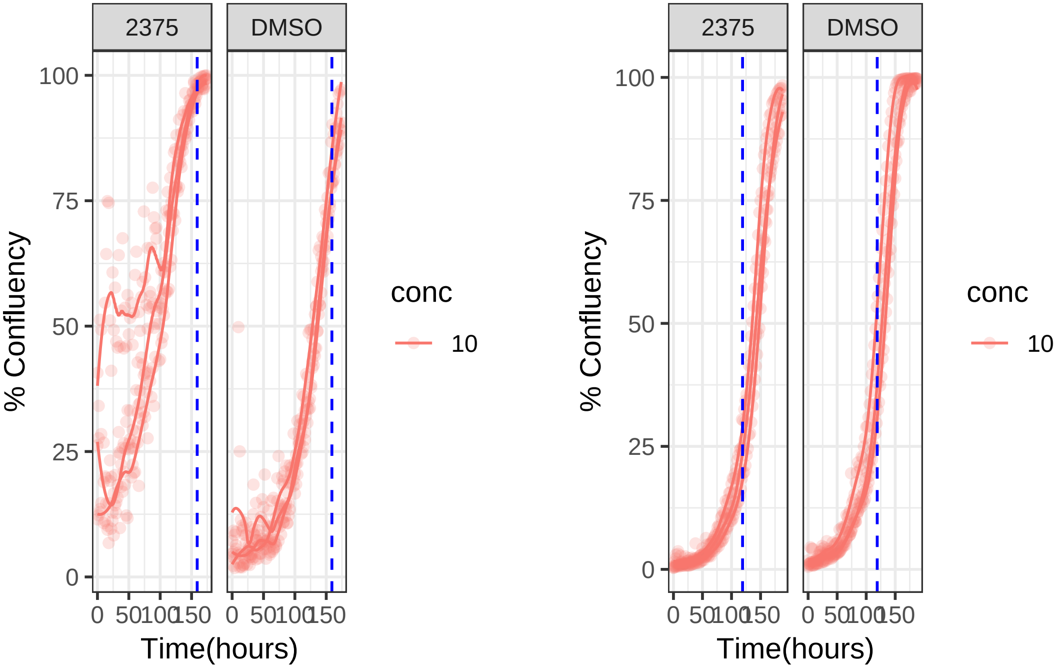

**10 uM 2379 RPE1 hTert**

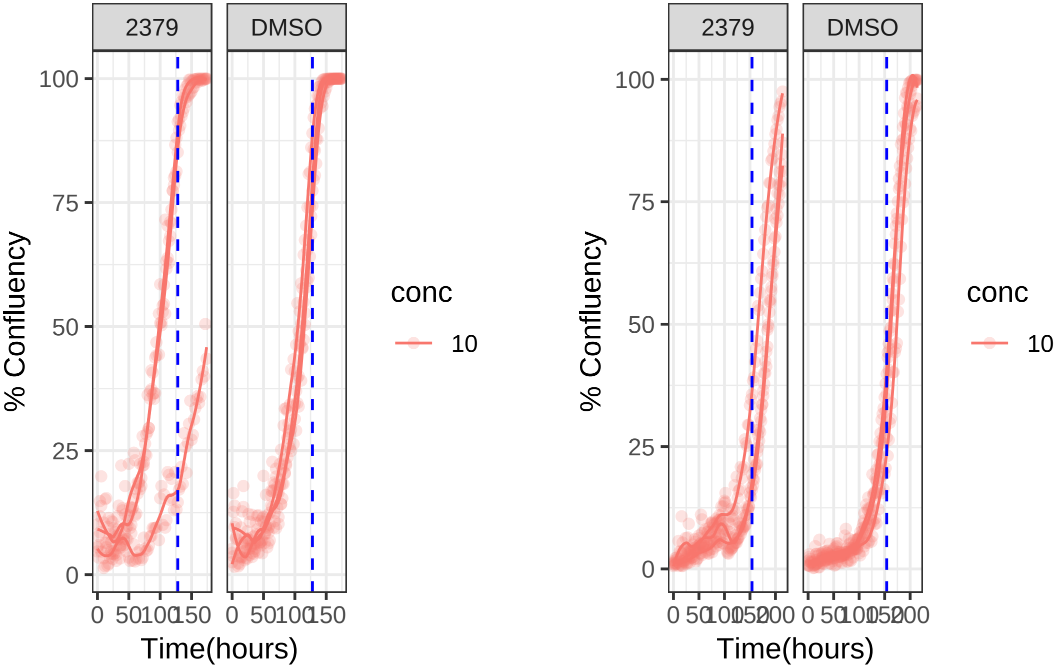

**10 uM 2379 RPE1 Ts12 Ts5**

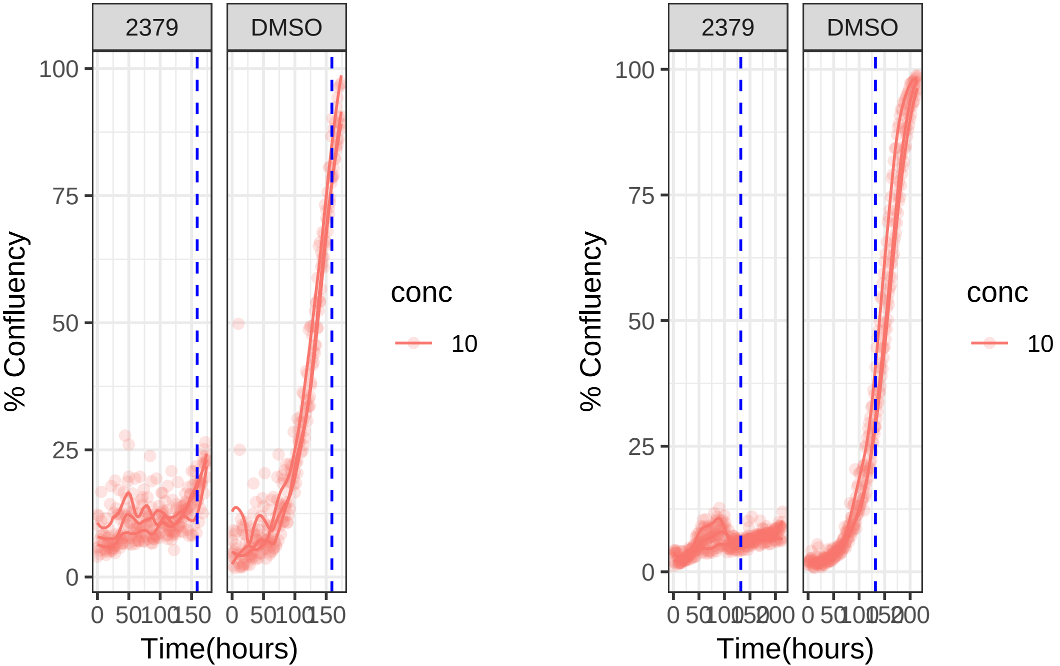

**10 uM 2402 RPE1 hTert**

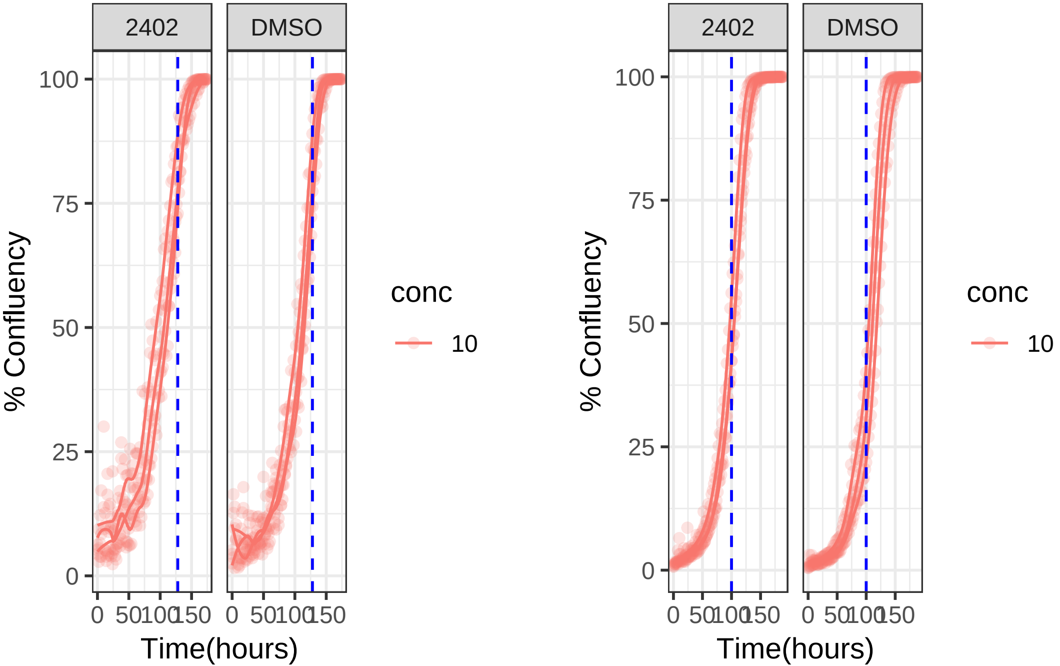

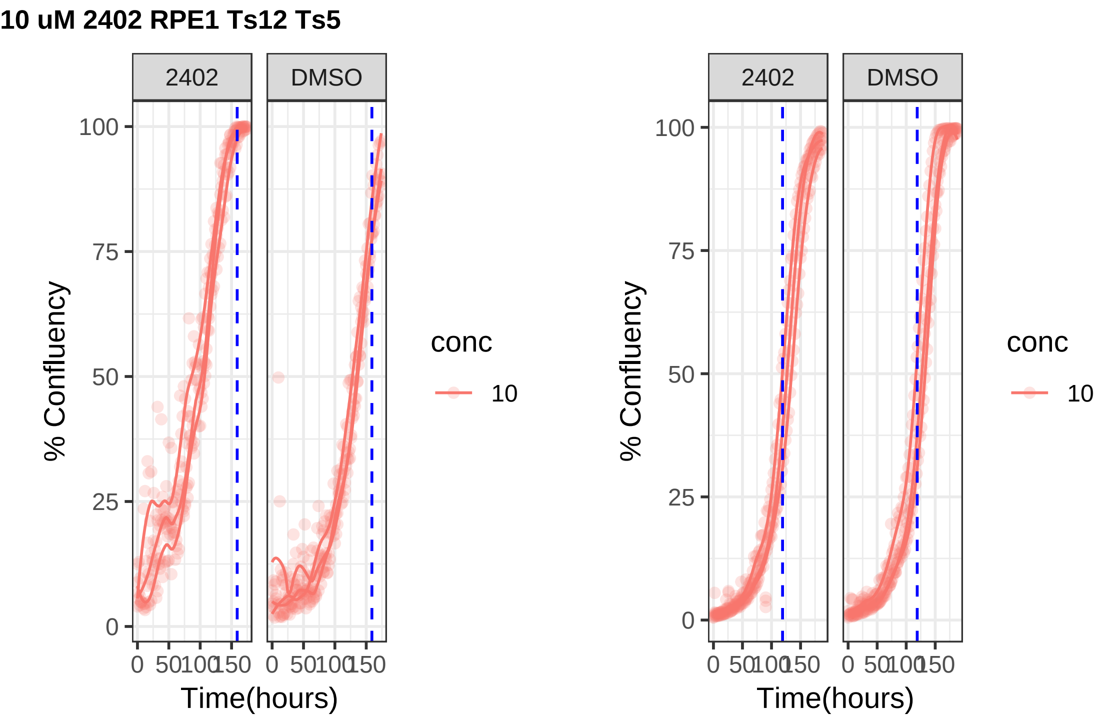

10 uM 2402 RPE1 Ts12 Ts5
2402
DMSO
% Confluency
100
75
50
25
0
0
50
100
150
Time(hours)
conc
10

**1 uM 2831 RPE1 hTert**

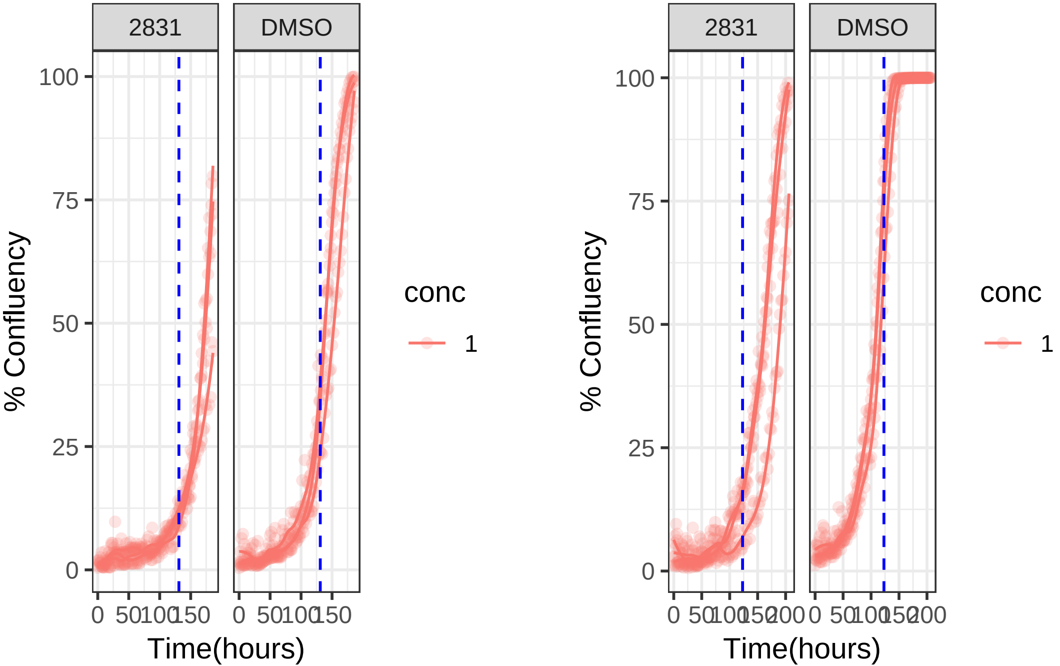

**10 uM 2831 RPE1 hTert**

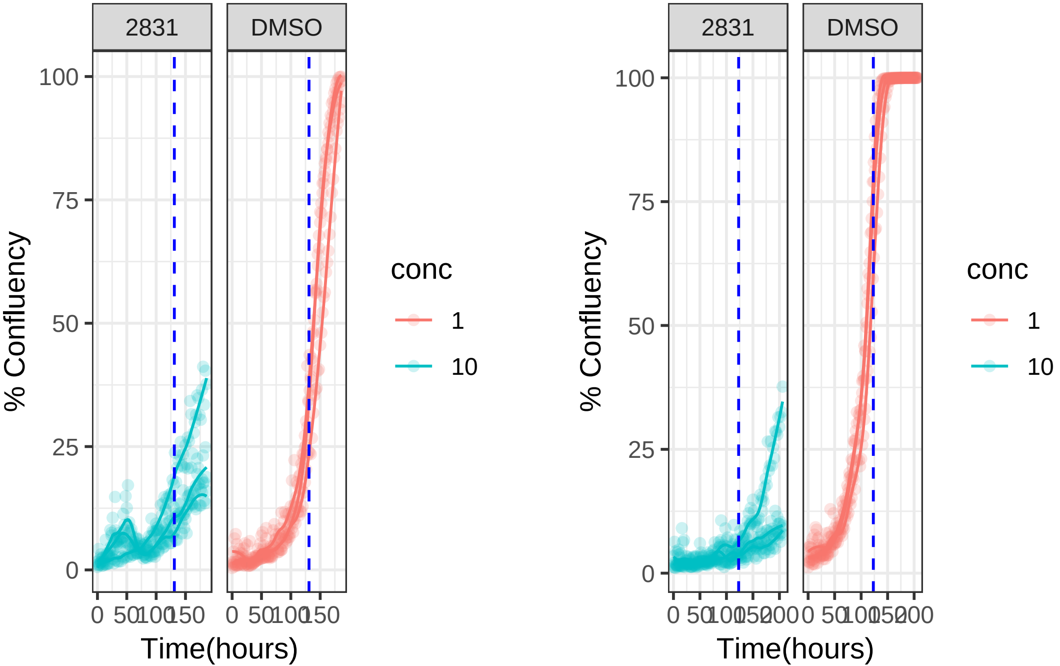

**1 uM 2831 RPE1 Ts12 Ts5**

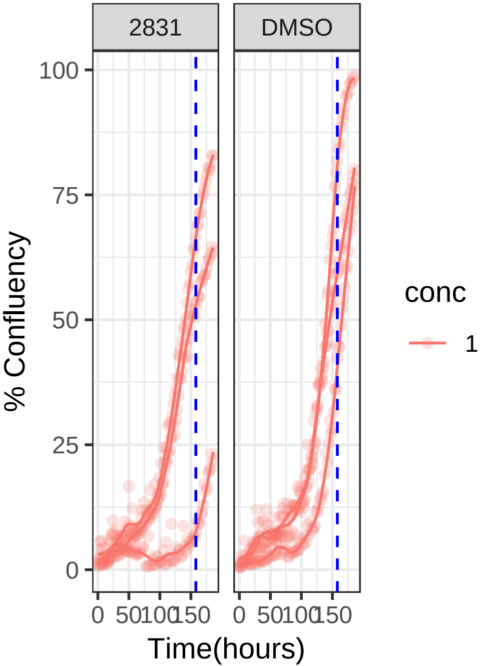


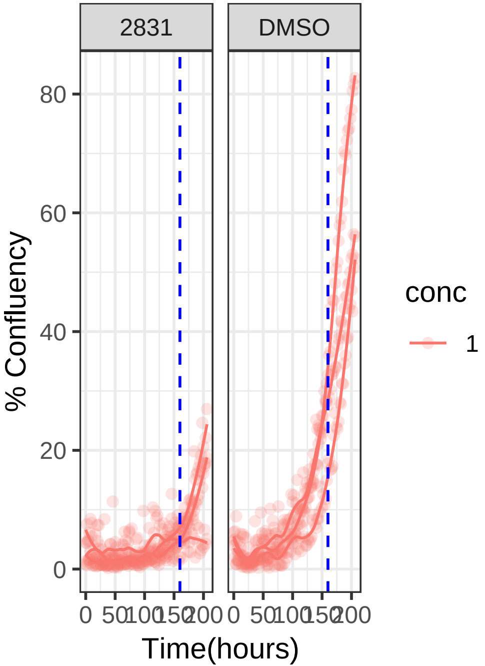

**10 uM 2831 RPE1 Ts12 Ts5**

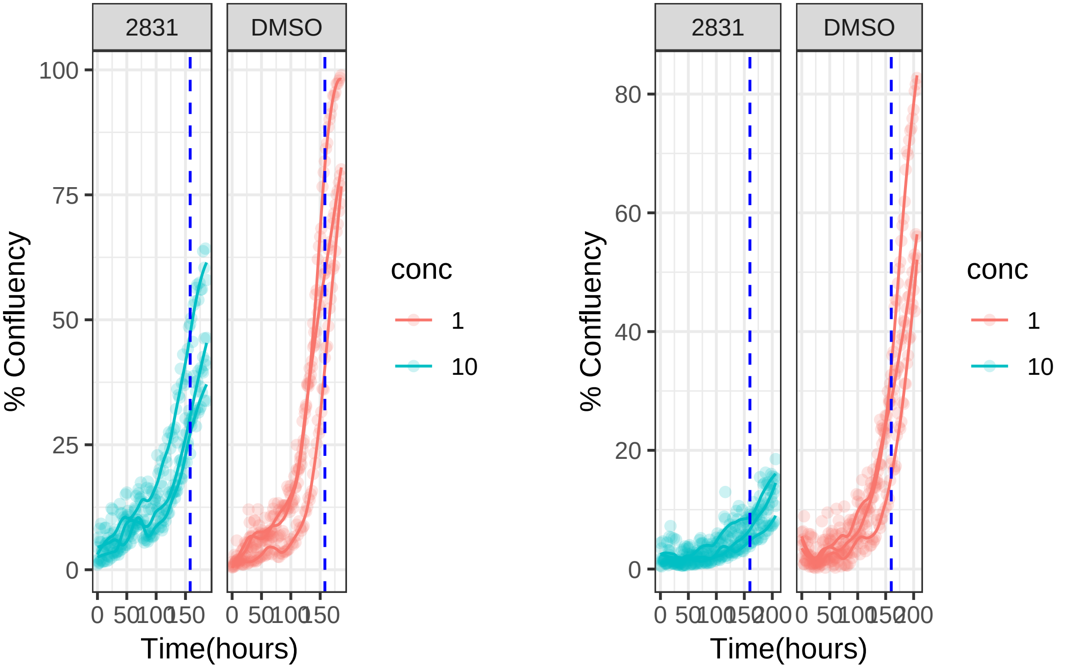

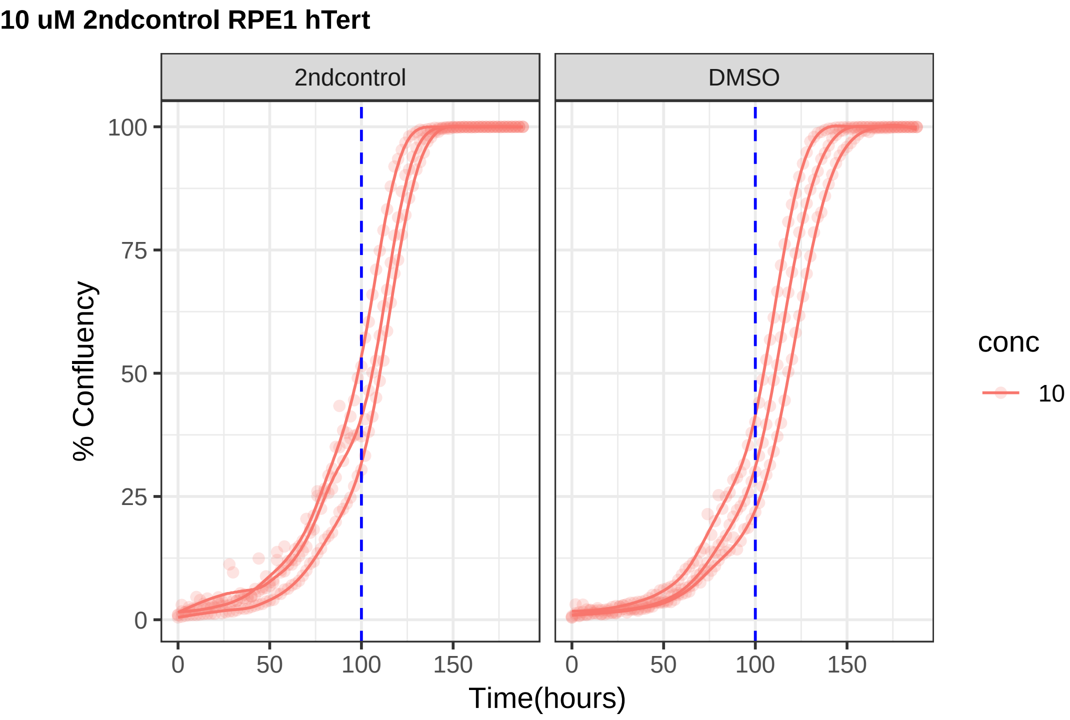
10 uM 2ndcontrol RPE1 hTert
2ndcontrol
DMSO
% Confluency
100
75
50
25
0
0
50
100
150
Time(hours)
conc
10

**10 uM 2ndcontrol RPE1 Ts12 Ts5**

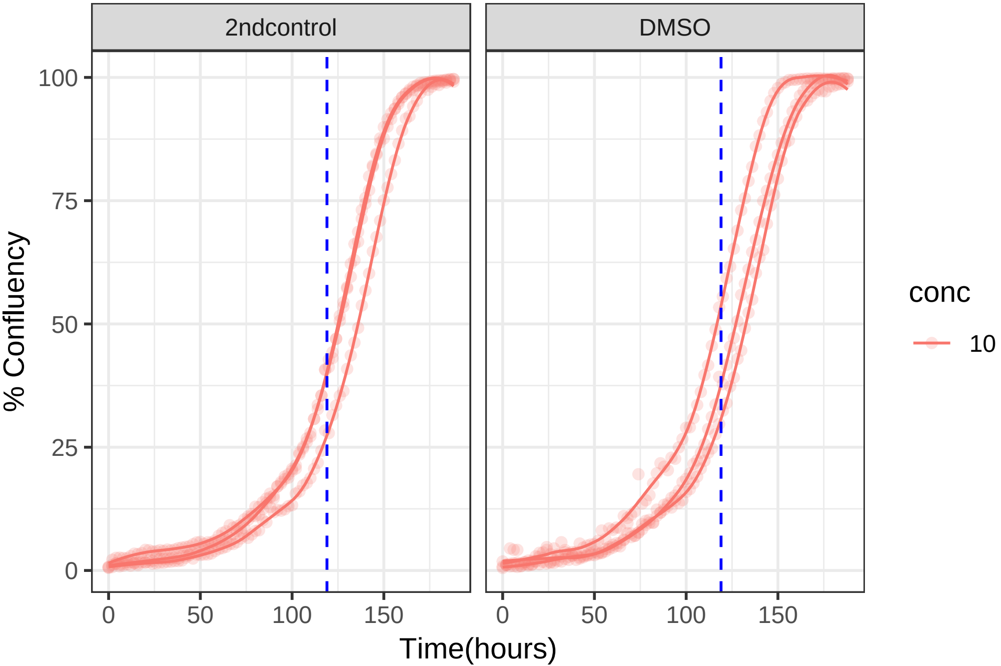
