## Supplementary material for "Altering microtubule dynamics is synergistically toxic with inhibition of the spindle checkpoint": Sup data 2

#### Supplementary data 2: CIN screen growth curves

CIN screen growth curves shown per cell line and per drug used. The top left corner lists the drug ID, the concentration and which cell line these growth curves belong to. RPE non-CIN control cells, and RPE1 Mad2KD are CIN cells due to the Mad2 knockdown.

The first (left) growth curve displayed is the first half of the CIN screen (day 1-4), then cells were passaged 1:8. The second growth curve (right) is the second half of the screen (day 5-8). Growth curves are shown for cell lines with the drug and with DMSO as a cell line control. All concentrations are in micromolar (uM).

Some drugs may have been screened at multiple concentrations.

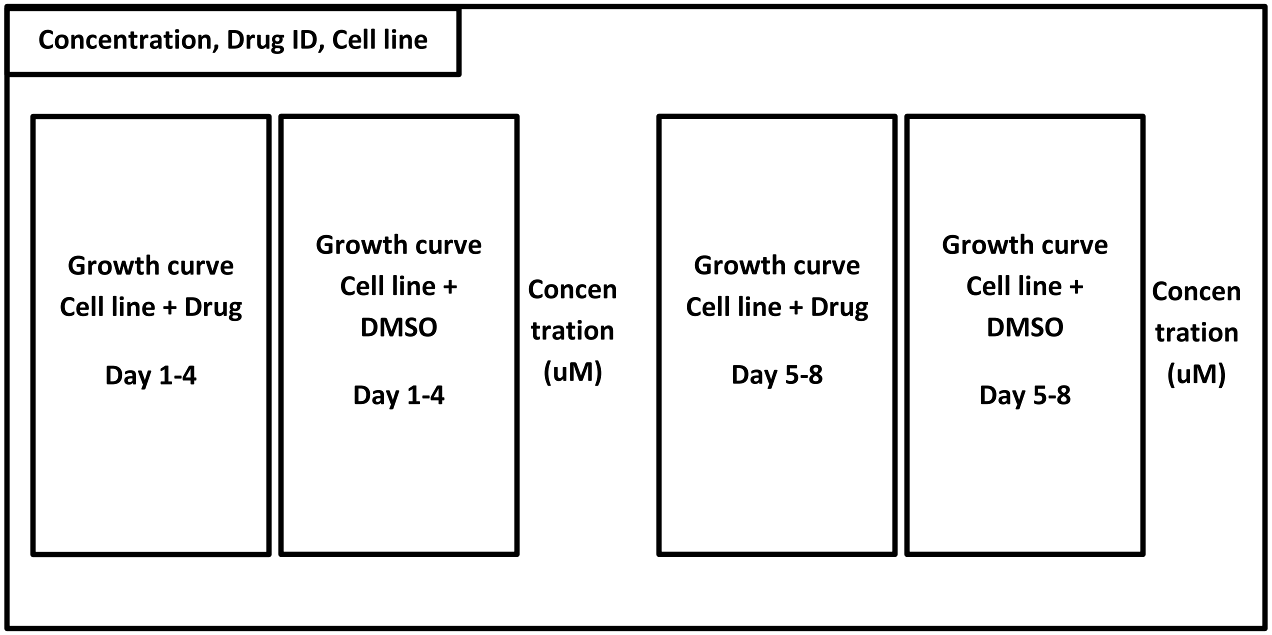

**10 uM 1134 RPE1 Mad2KD**

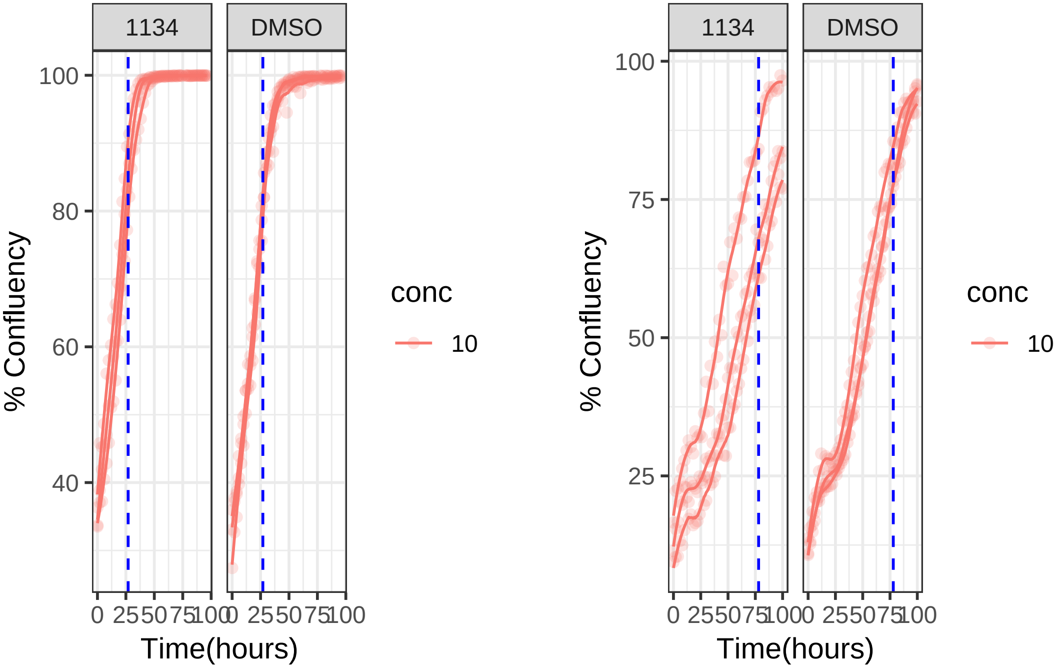

**10 uM 1134 RPE1**

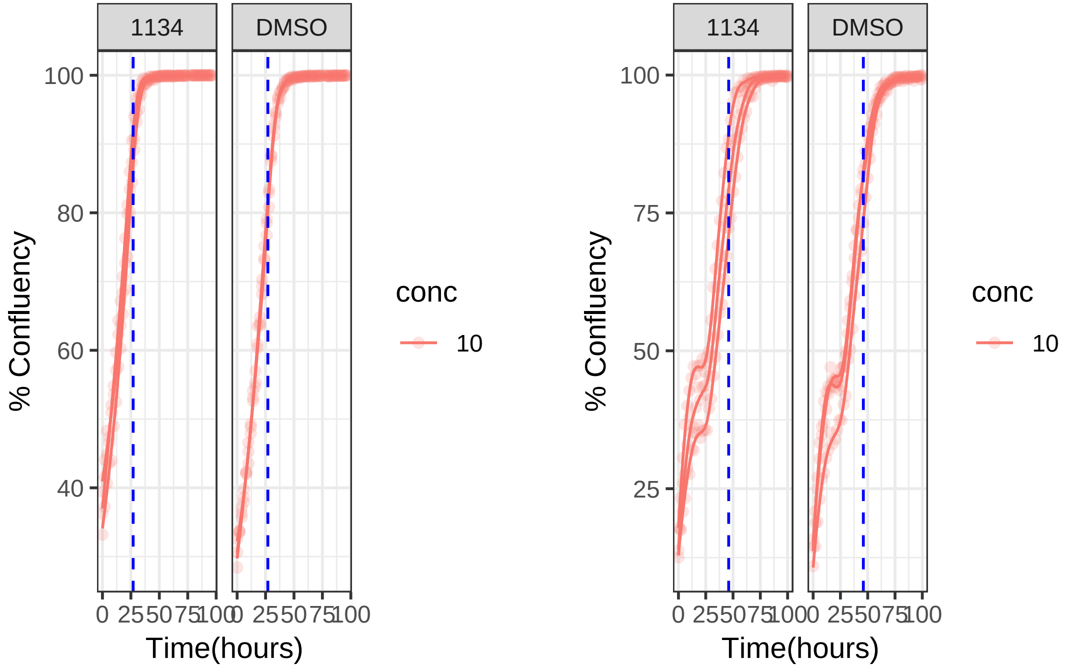

**10 uM 1223 RPE1 Mad2KD**

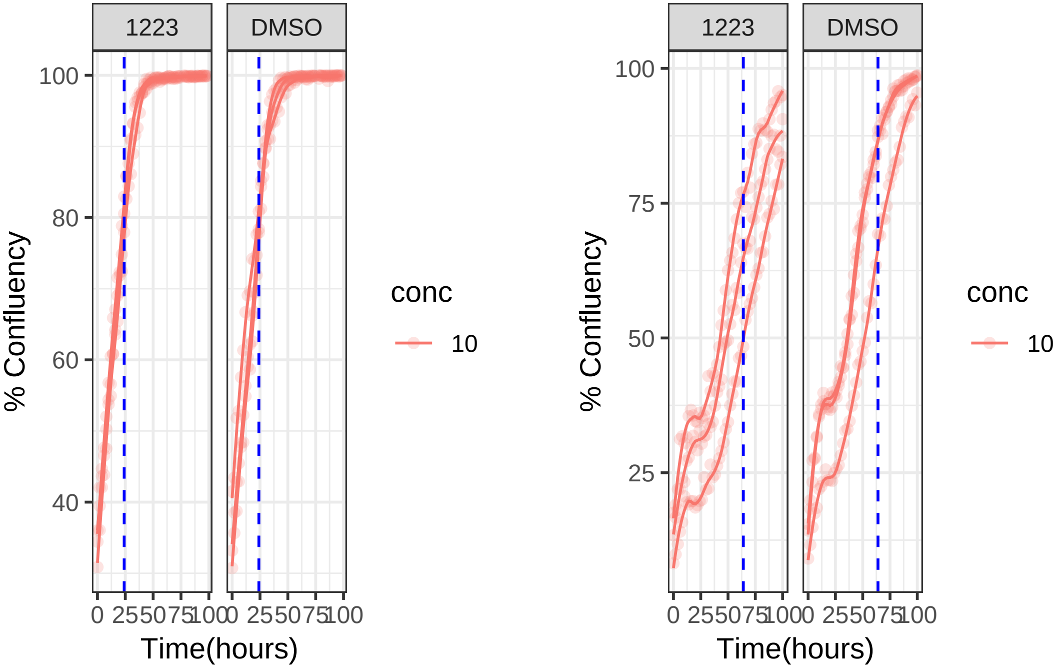

**10 uM 1223 RPE1**

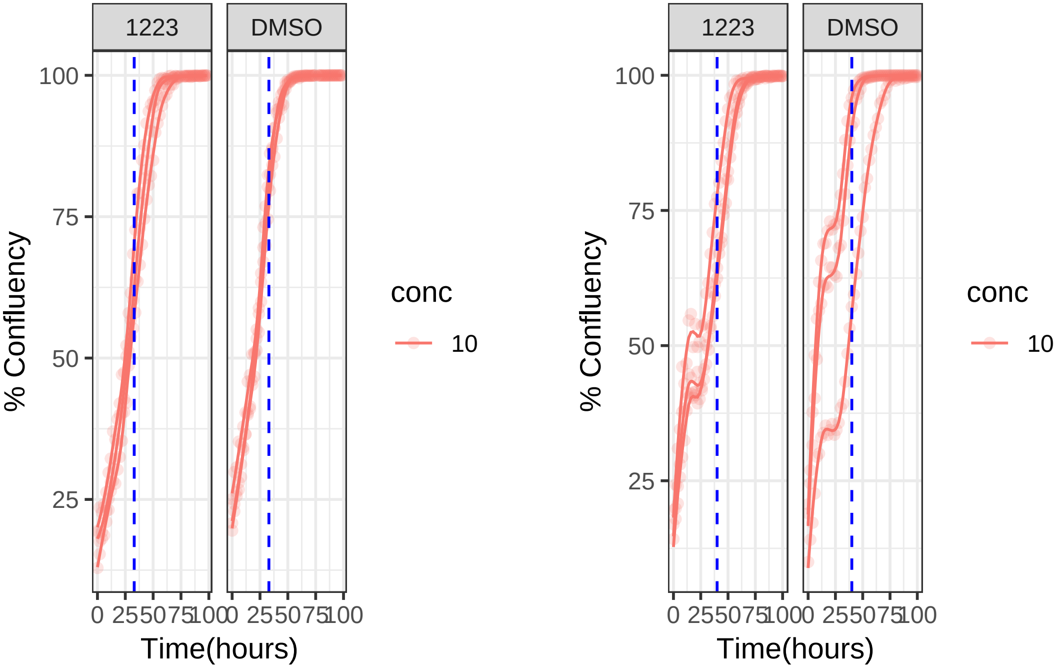

**0.001 uM 1233 RPE1 Mad2KD**

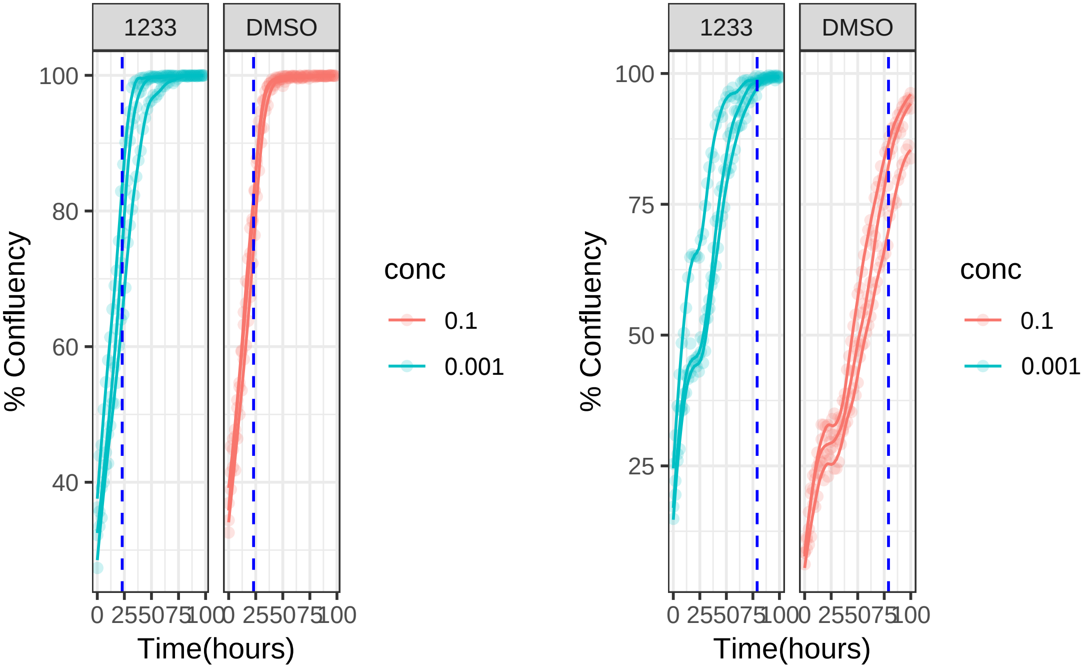

**0.001 uM 1233 RPE1**

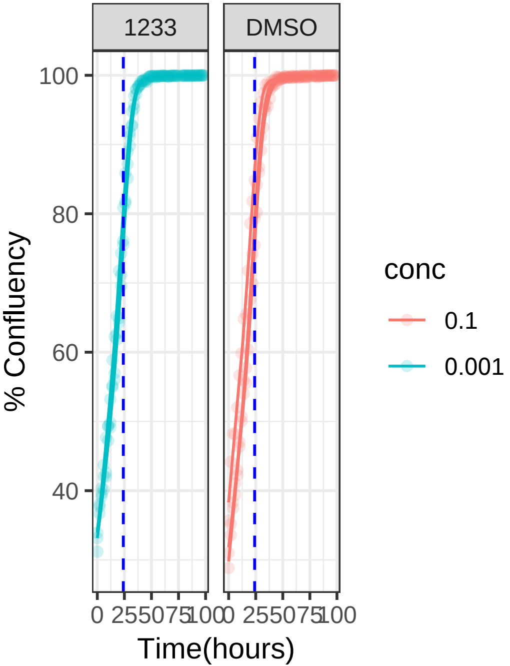


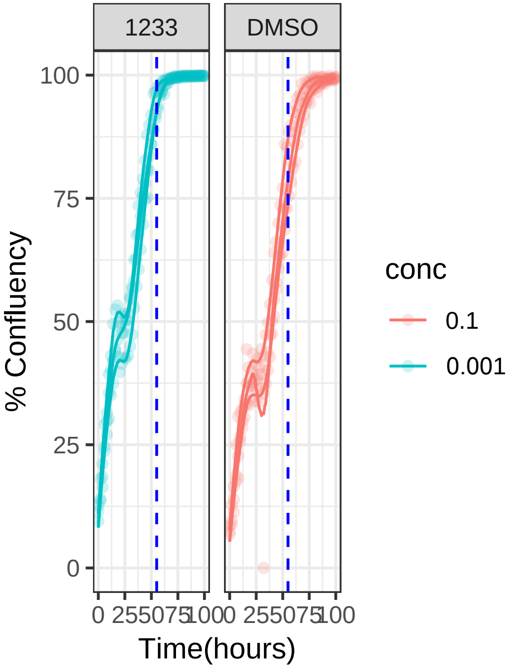

**10 uM 1239 RPE1 Mad2KD**

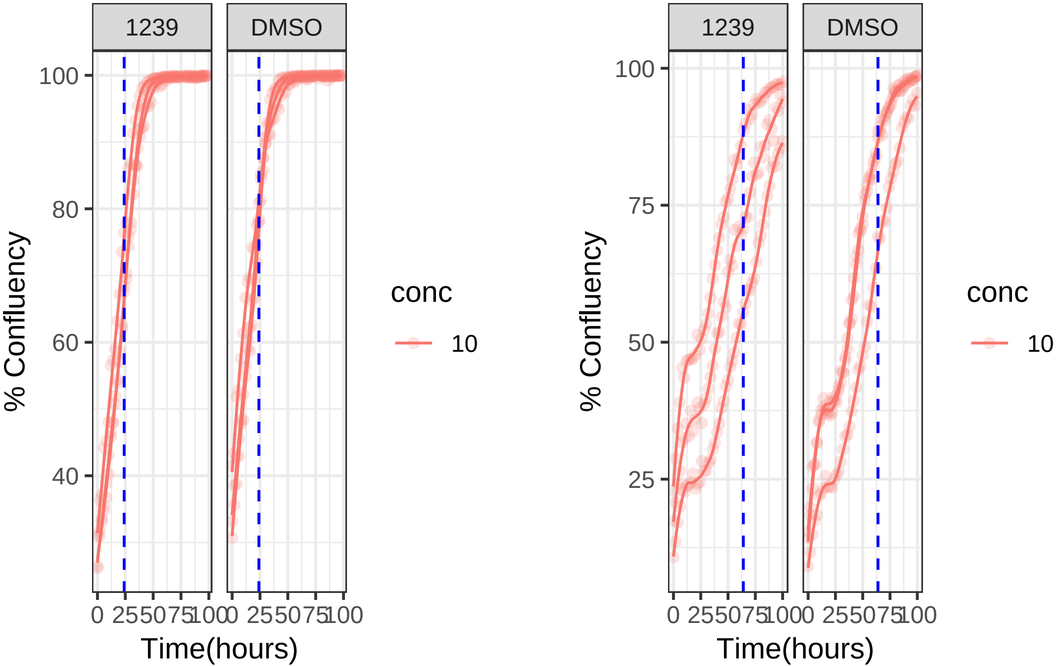

**10 uM 1239 RPE1**

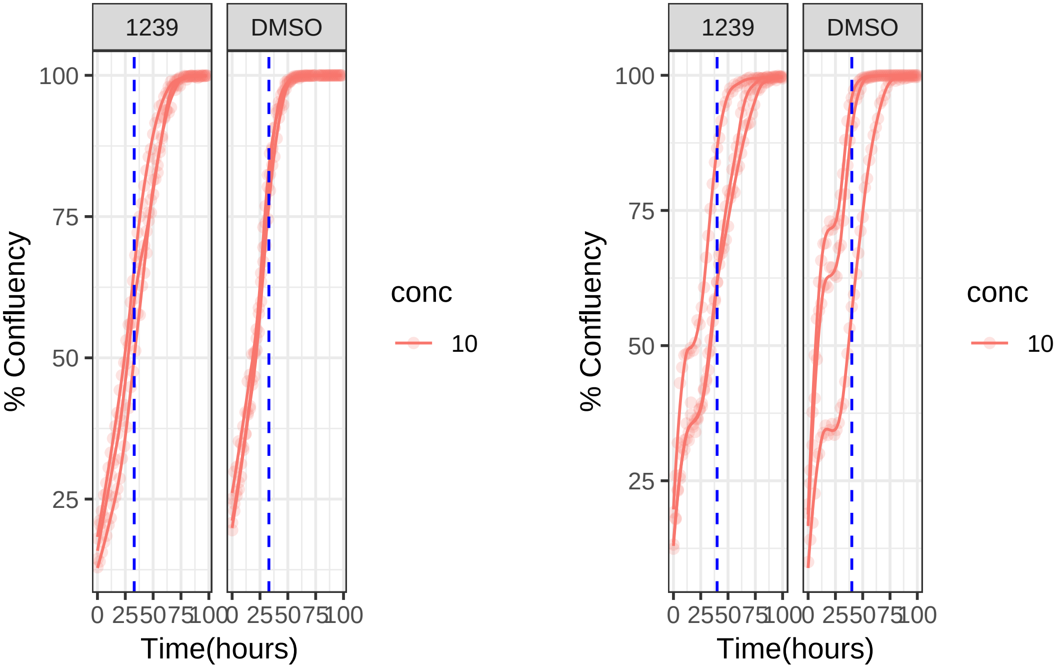

**10 uM 1243 RPE1 Mad2KD**

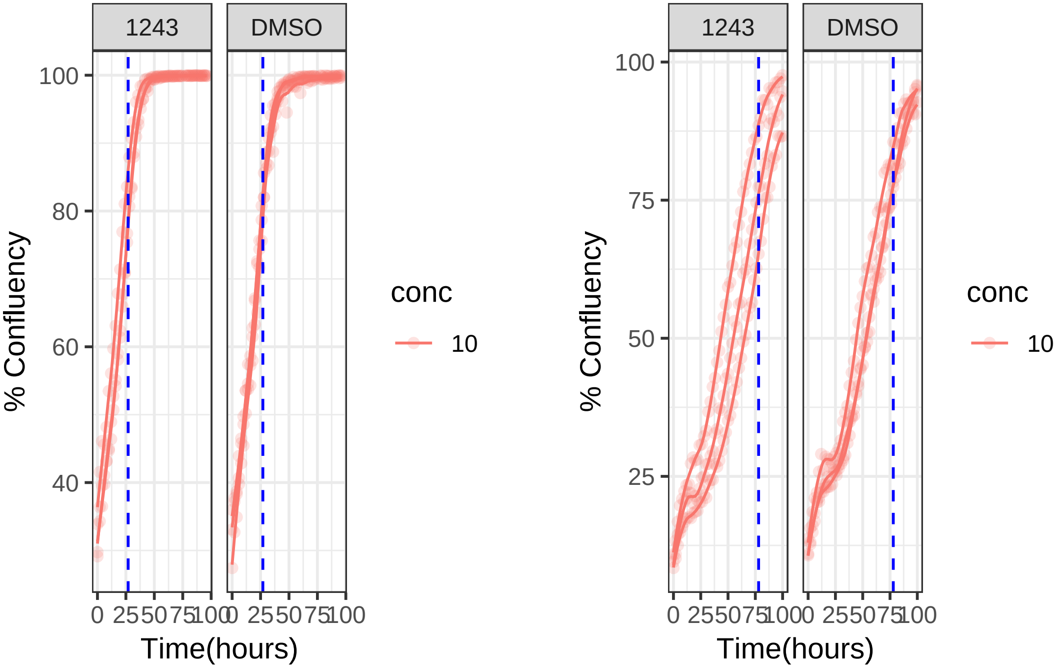

**10 uM 1243 RPE1**

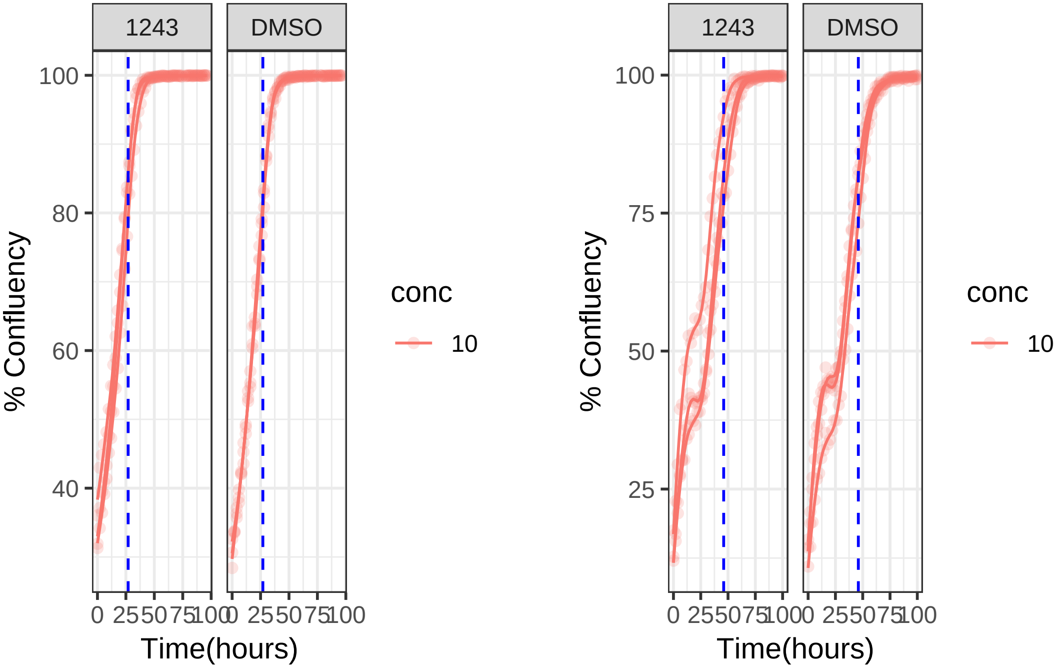

**0.1 uM 1275 RPE1 Mad2KD**

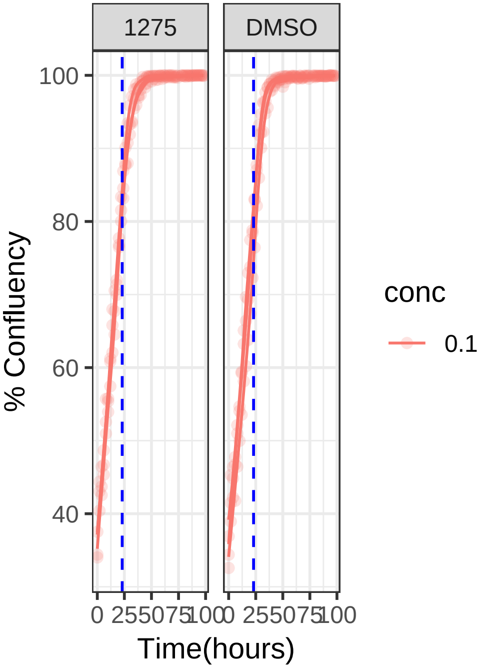


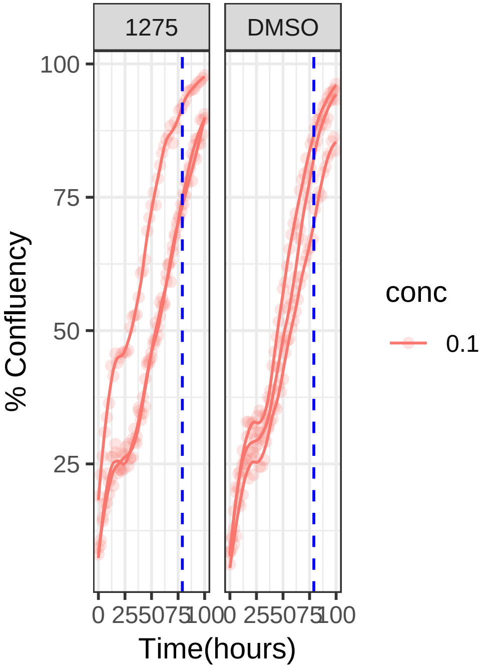

**0.1 uM 1275 RPE1**

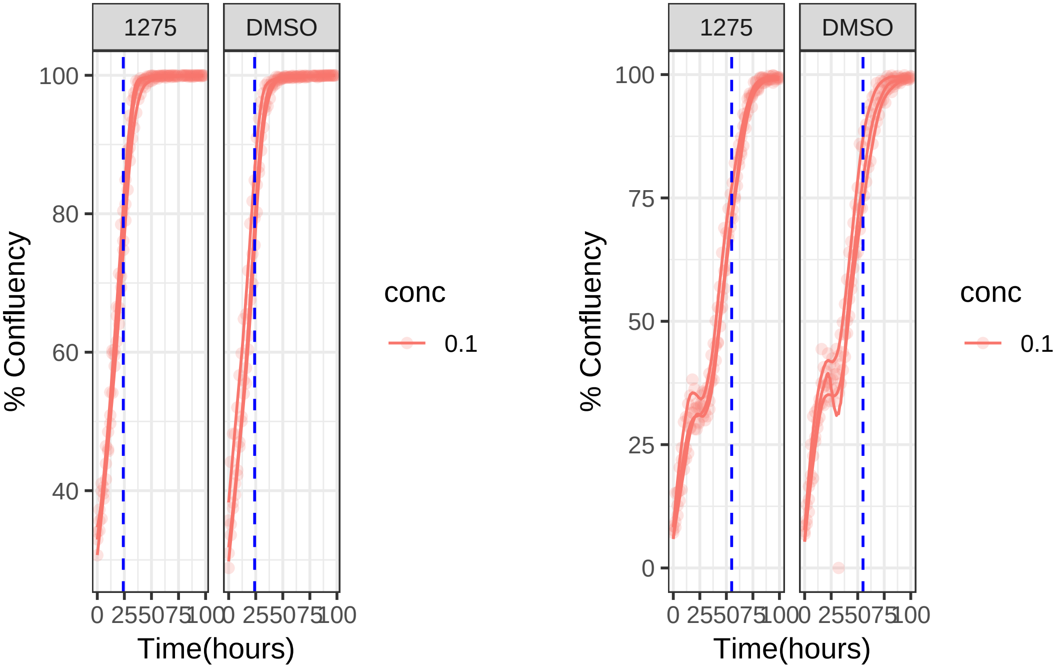

1 uM 1310 RPE1 Mad2KD

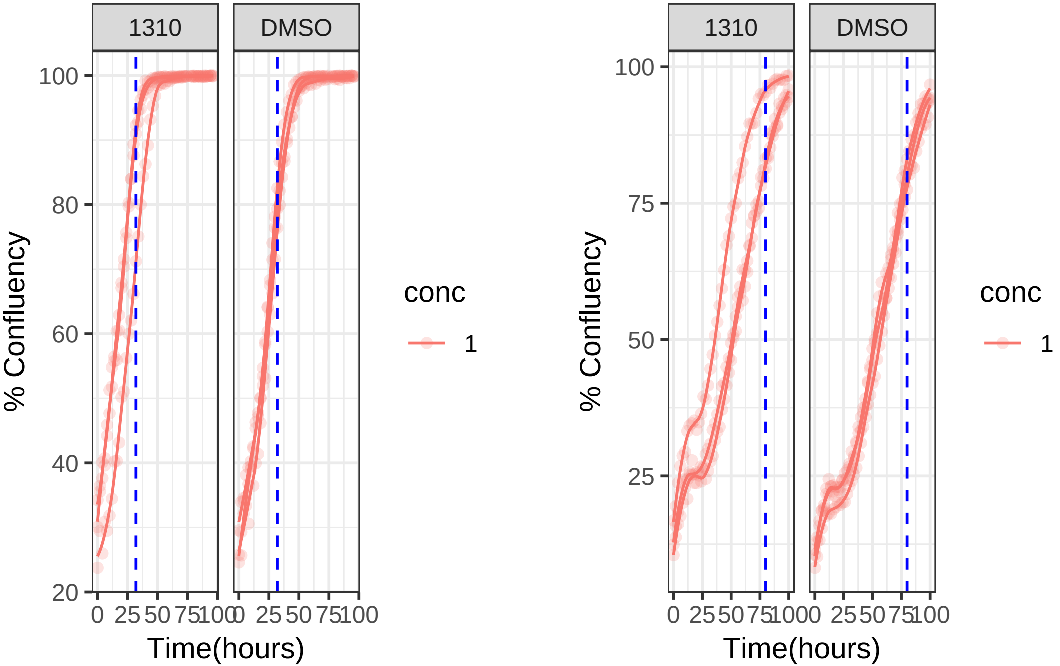

**1 uM 1310 RPE1**

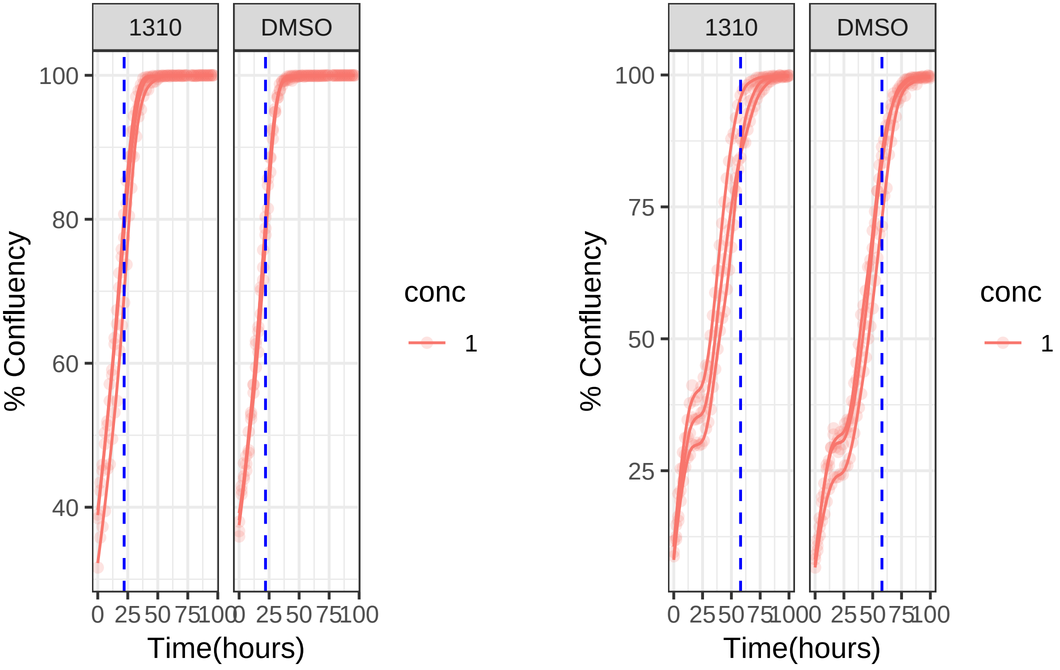

### 1 uM 1334 RPE1 Mad2KD

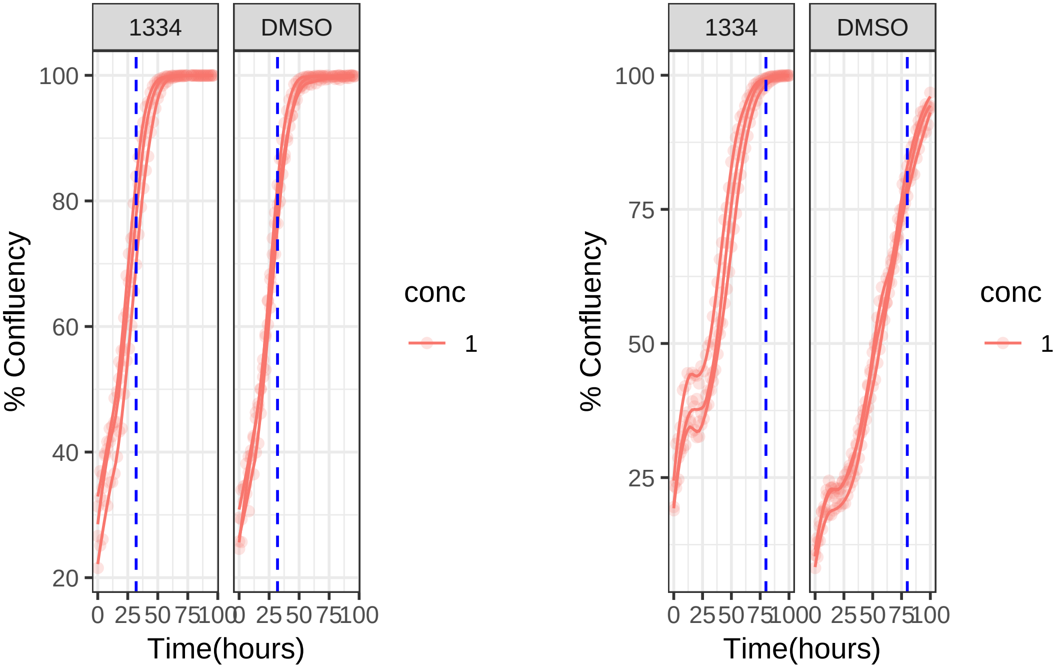

**1 uM 1334 RPE1**

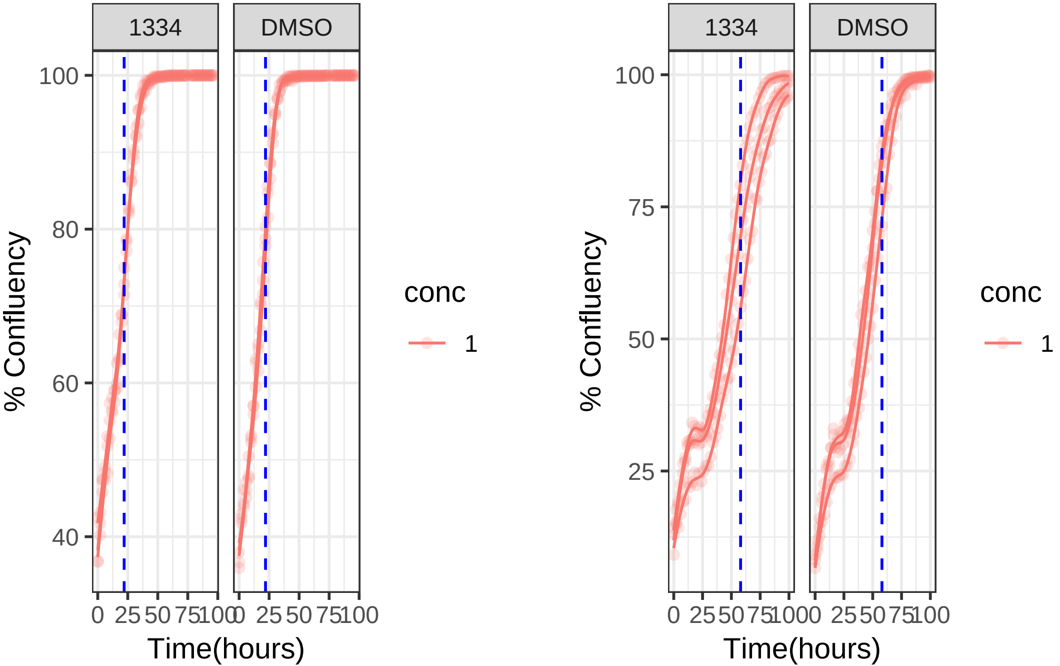

**1 uM 1362 RPE1 Mad2KD**

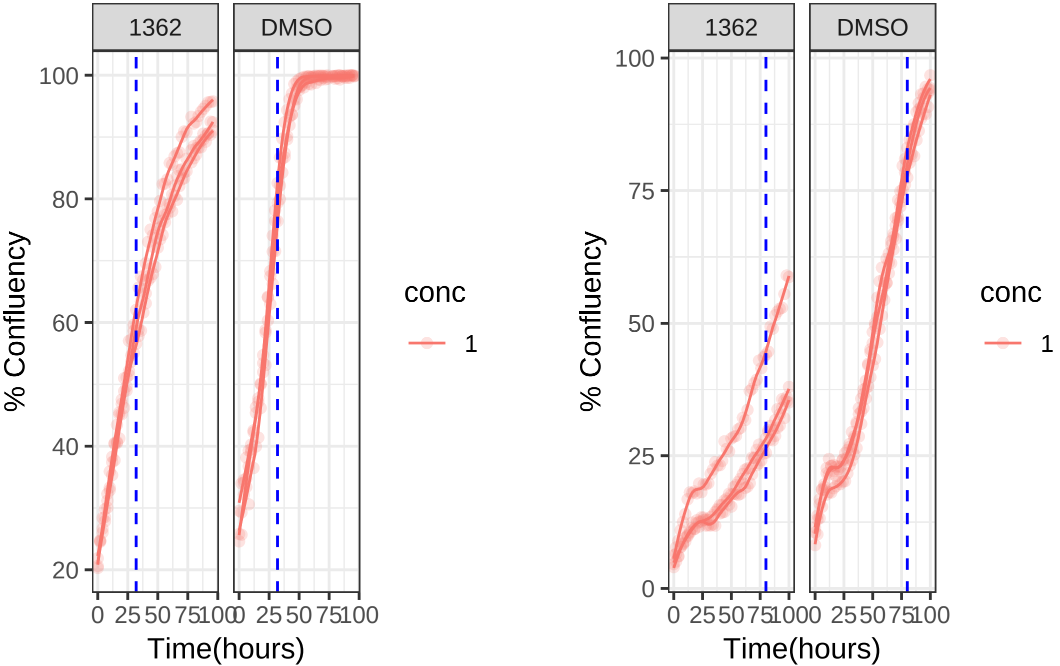

**1 uM 1362 RPE1**

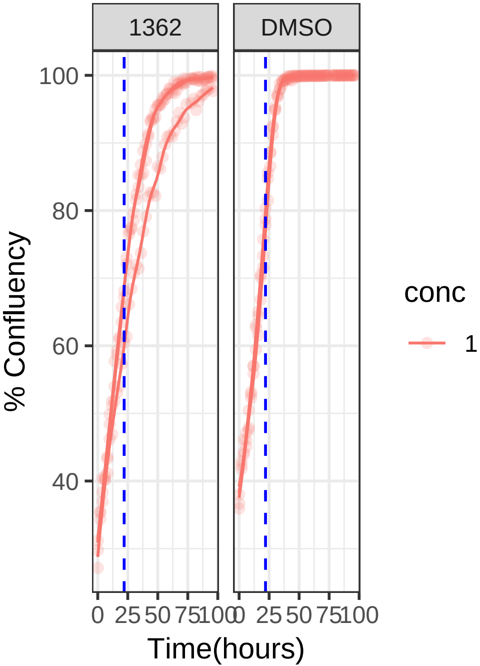


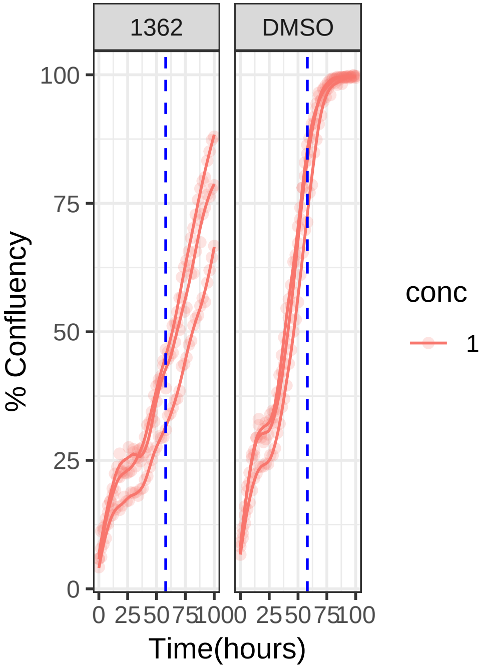

**1 uM 1377 RPE1 Mad2KD**

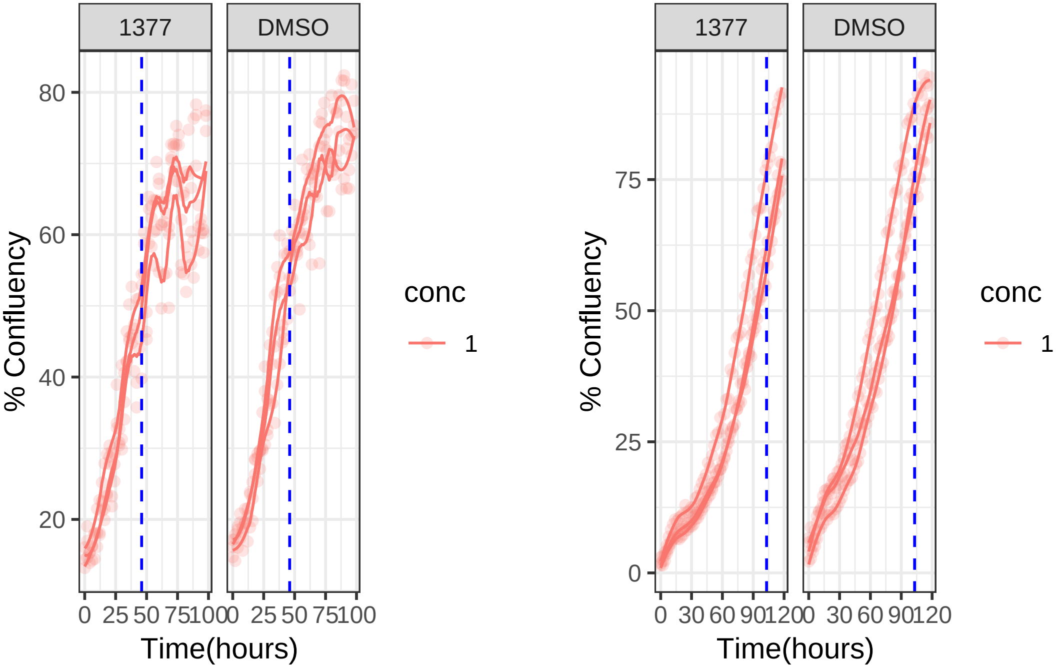

**1 uM 1377 RPE1**

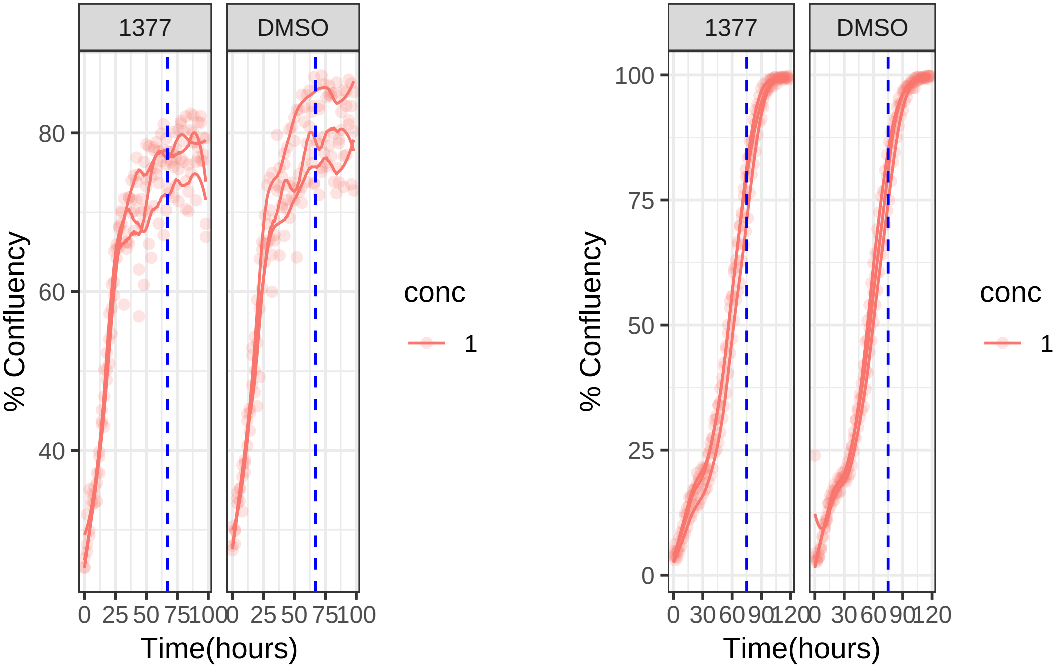

**0.1 uM 1407 RPE1 Mad2KD**

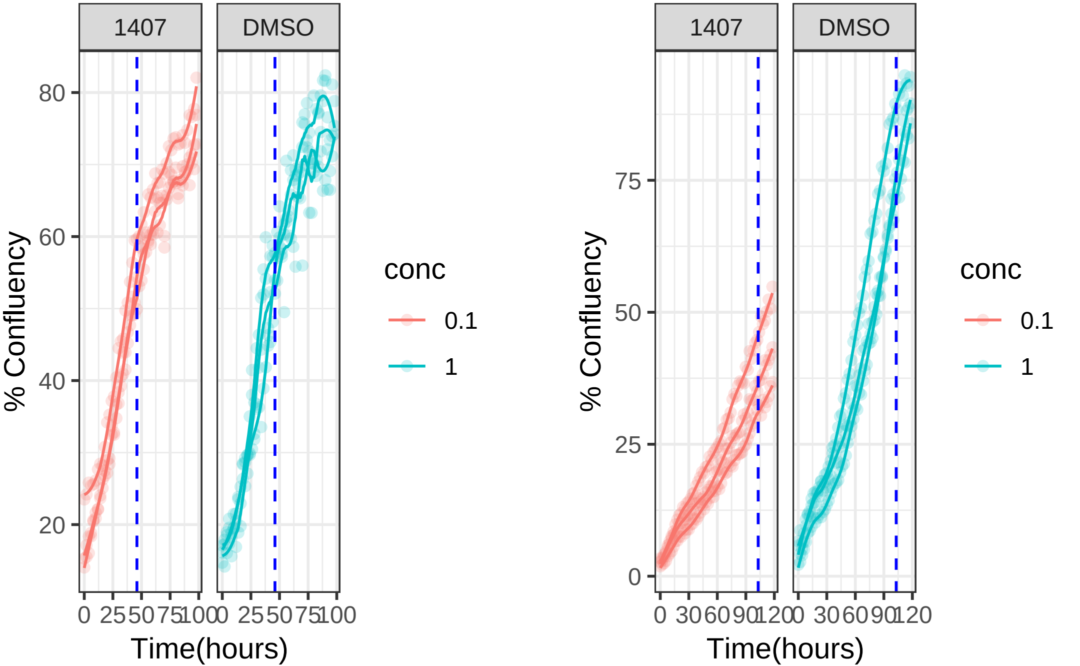

**1 uM 1407 RPE1 Mad2KD**

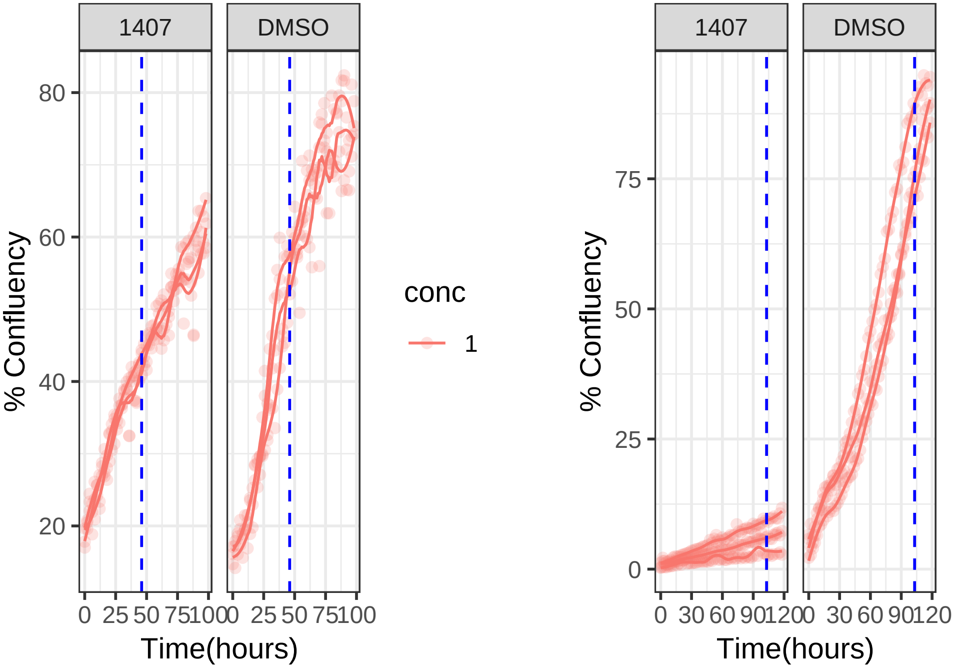

**0.1 uM 1407 RPE1**

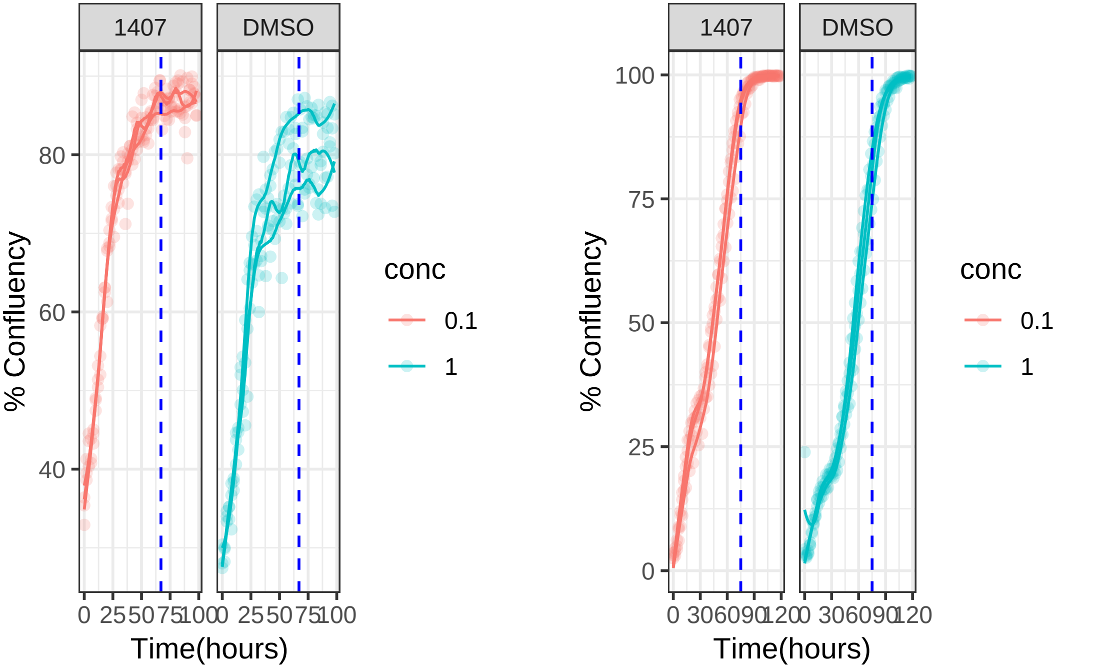

**1 uM 1407 RPE1**

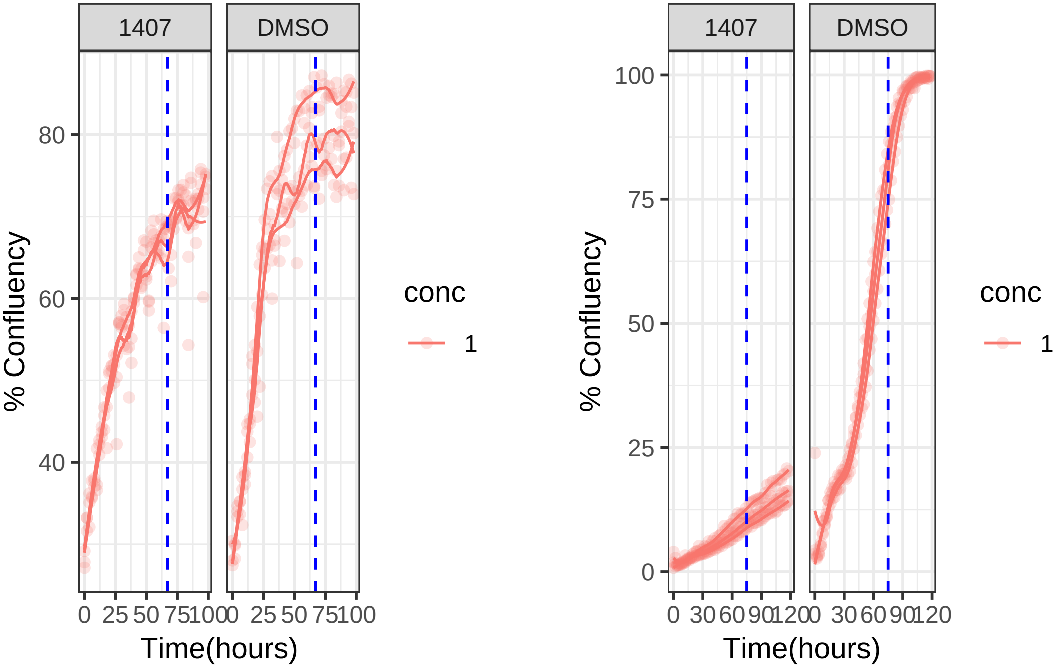

### 1 uM 1463 RPE1 Mad2KD

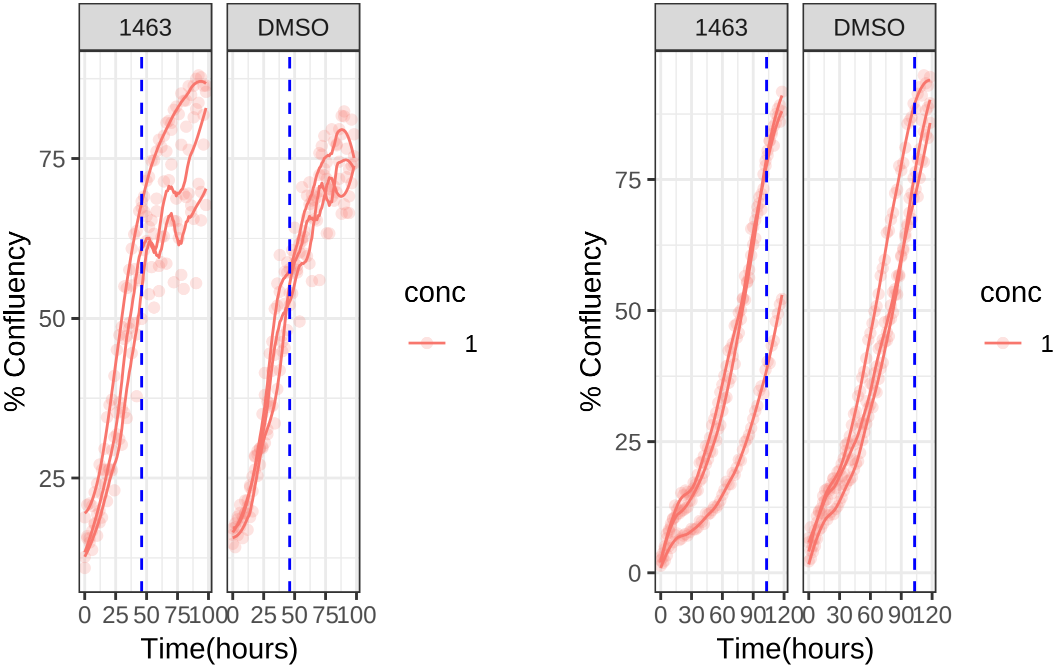

**1 uM 1463 RPE1**

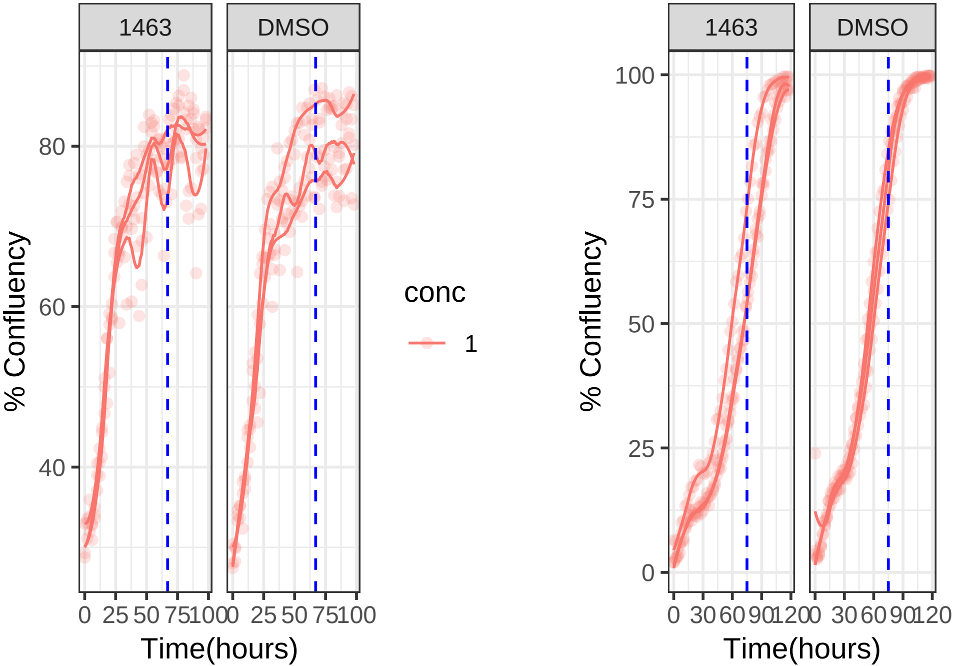

**1 uM 1472 RPE1 Mad2KD**

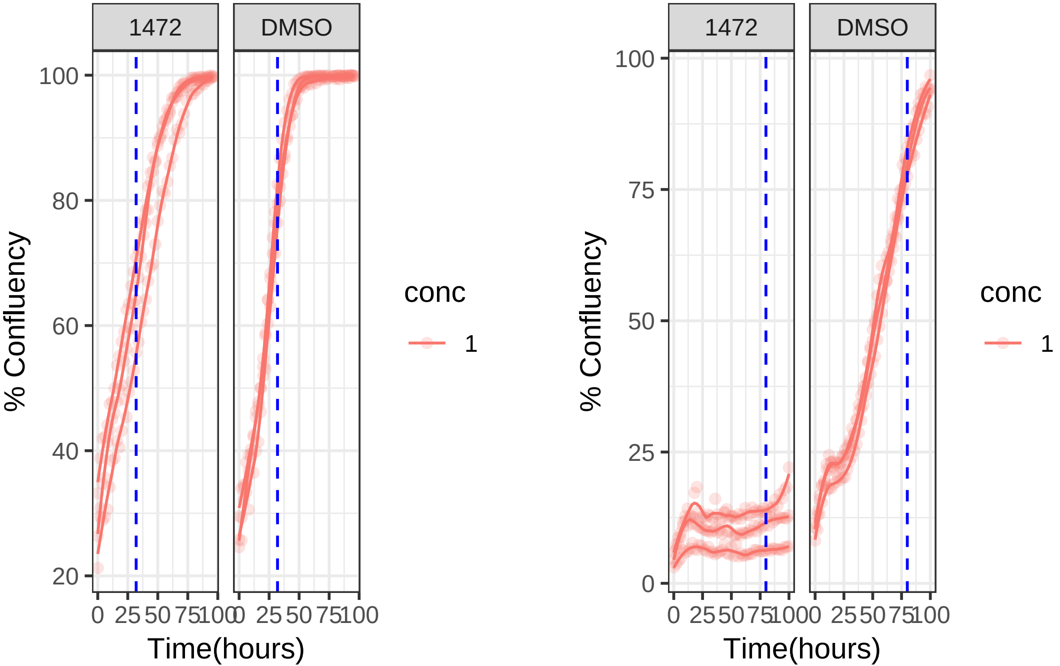

**1 uM 1472 RPE1**

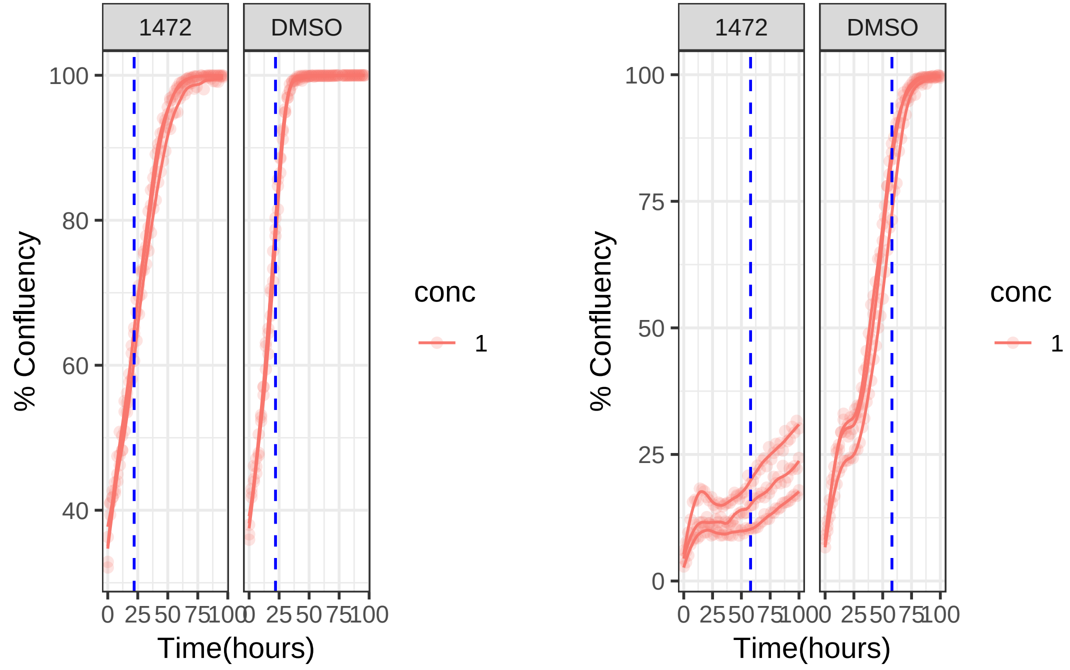

**0.1 uM 1494 RPE1 Mad2KD**

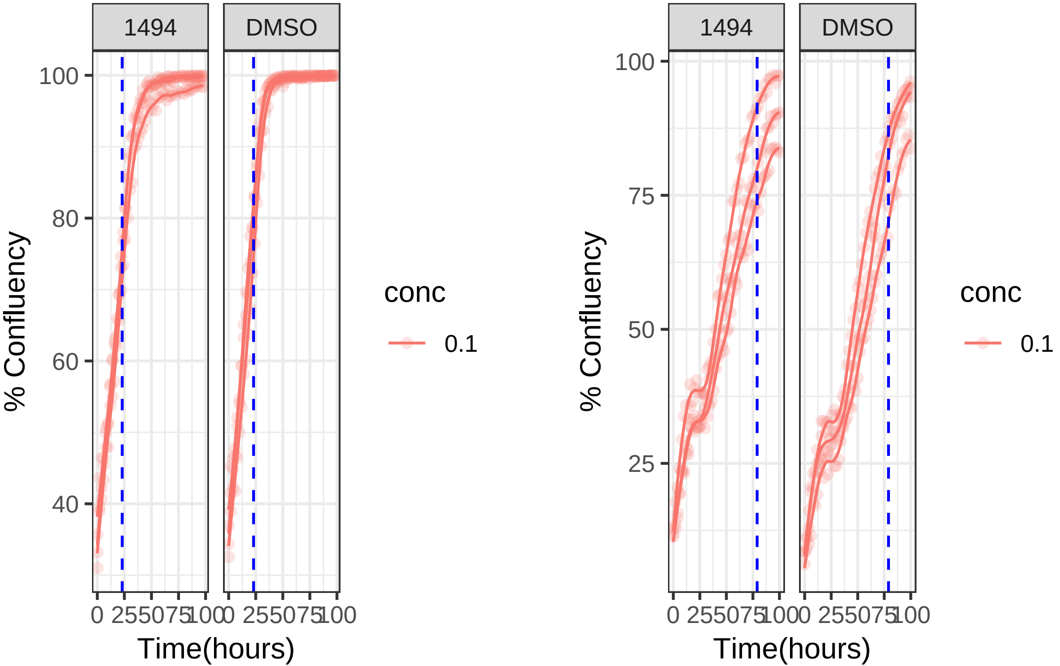

**0.1 uM 1494 RPE1**

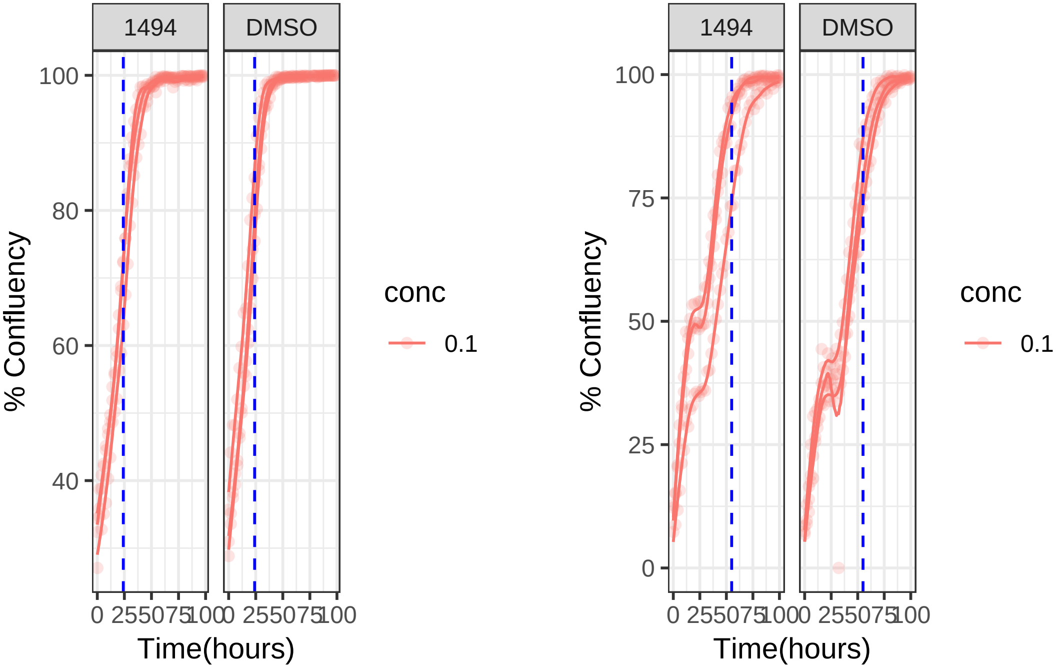

**10 uM 1520 RPE1 Mad2KD**

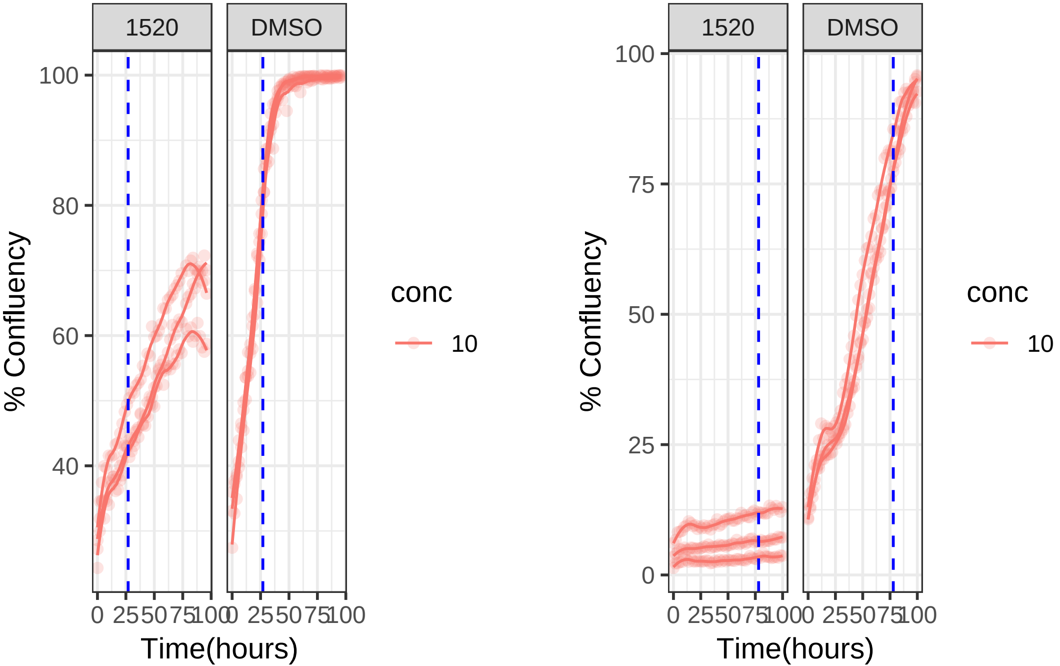

**0.1 uM 1520 RPE1 Mad2KD**

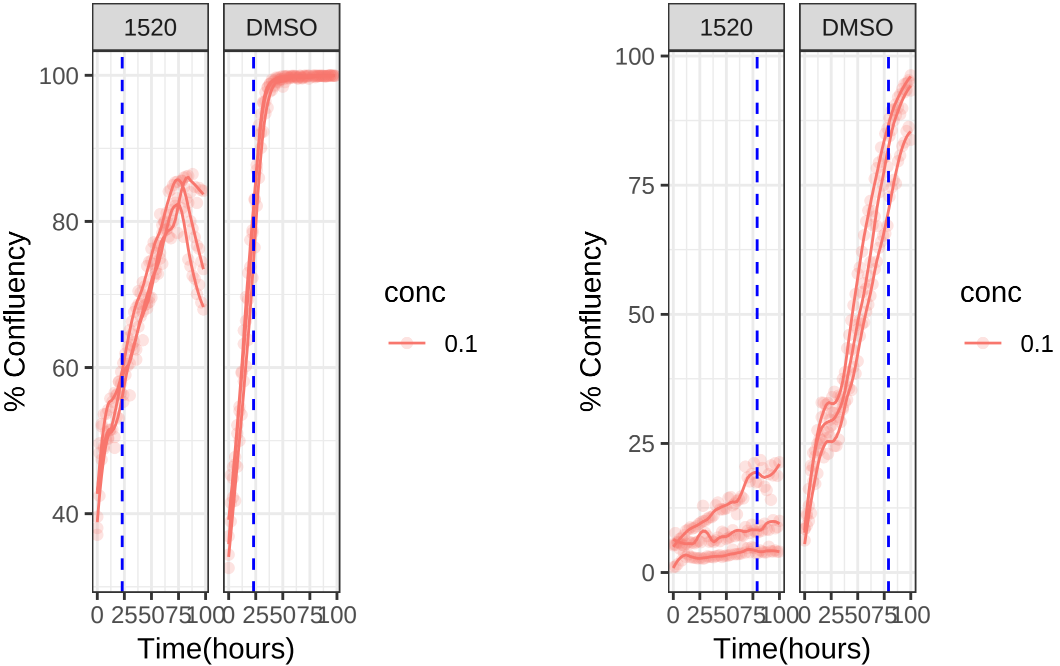

**10 uM 1520 RPE1**

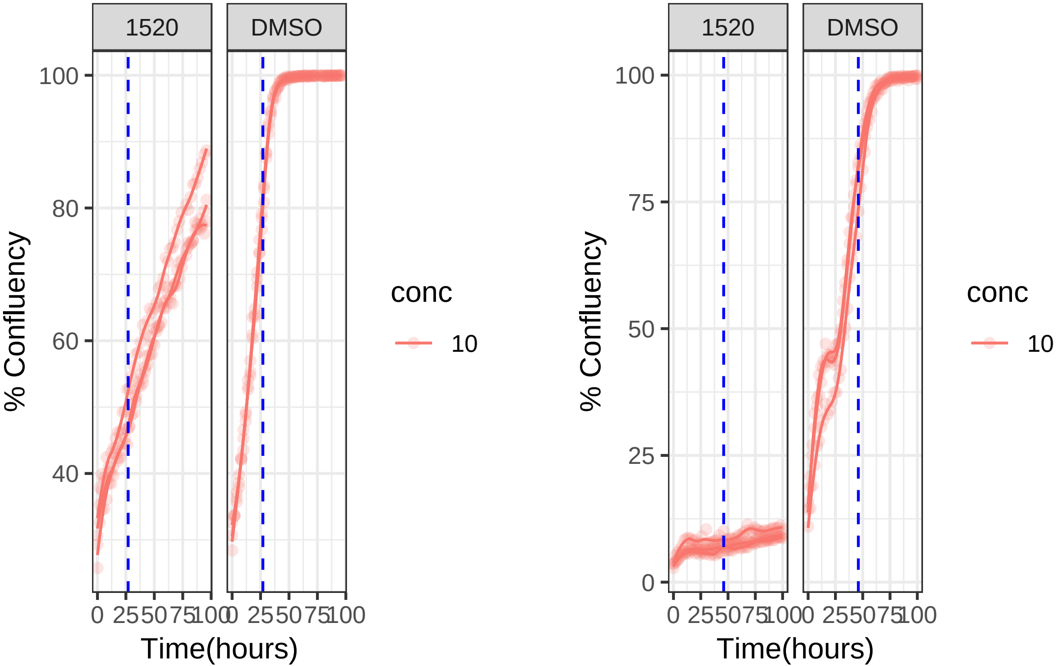

**0.1 uM 1520 RPE1**

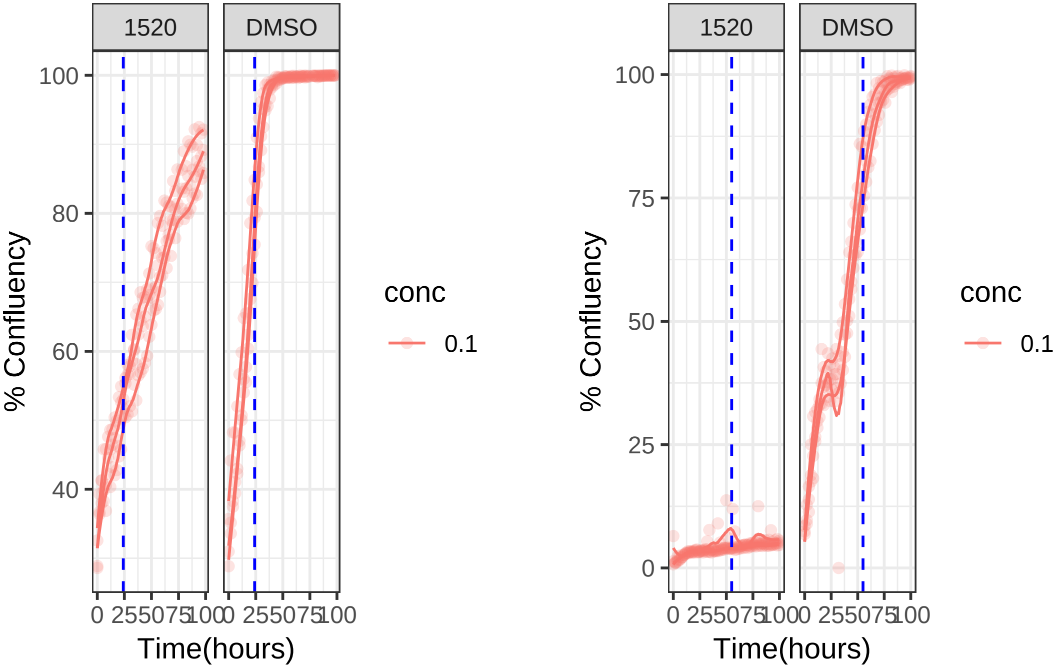

**0.01 uM 1561 RPE1 Mad2KD**

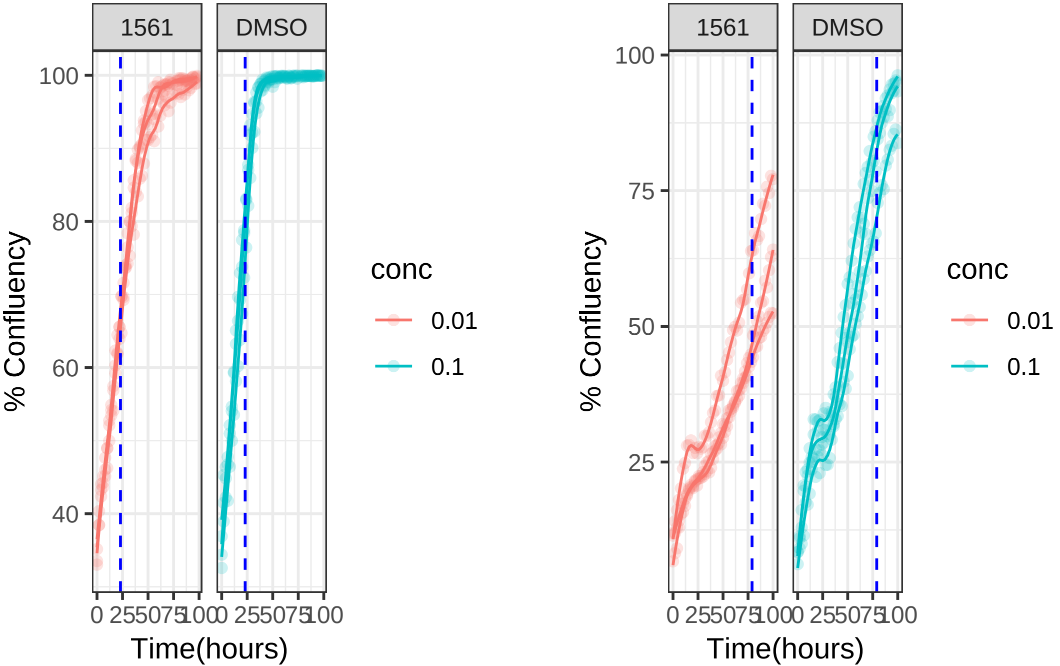

**0.1 uM 1561 RPE1 Mad2KD**

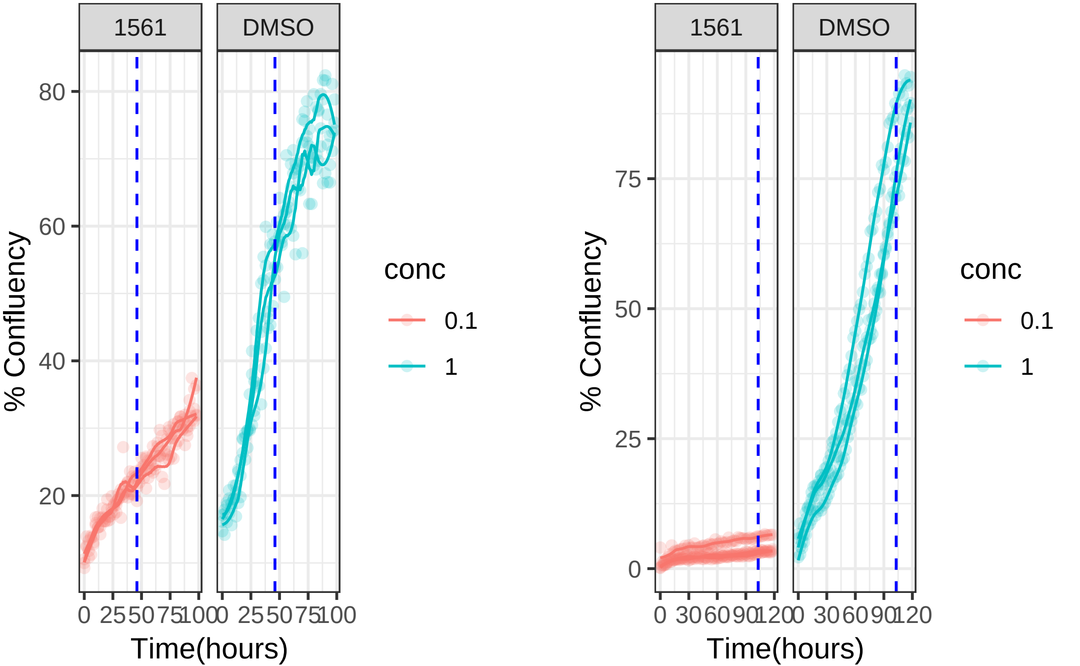

**0.01 uM 1561 RPE1**

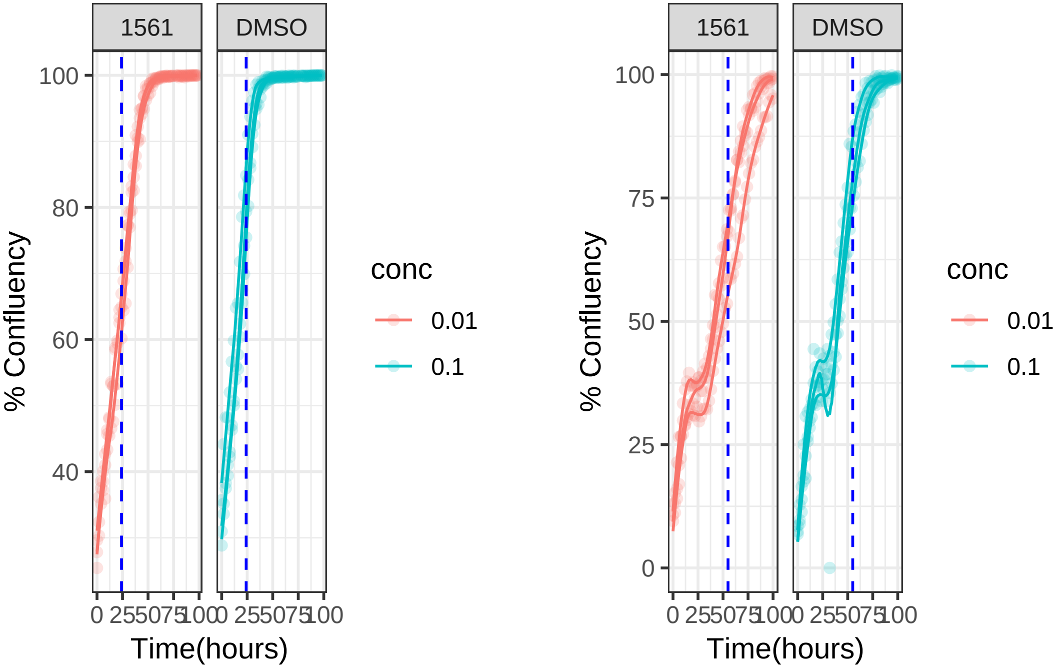

**0.1 uM 1561 RPE1**

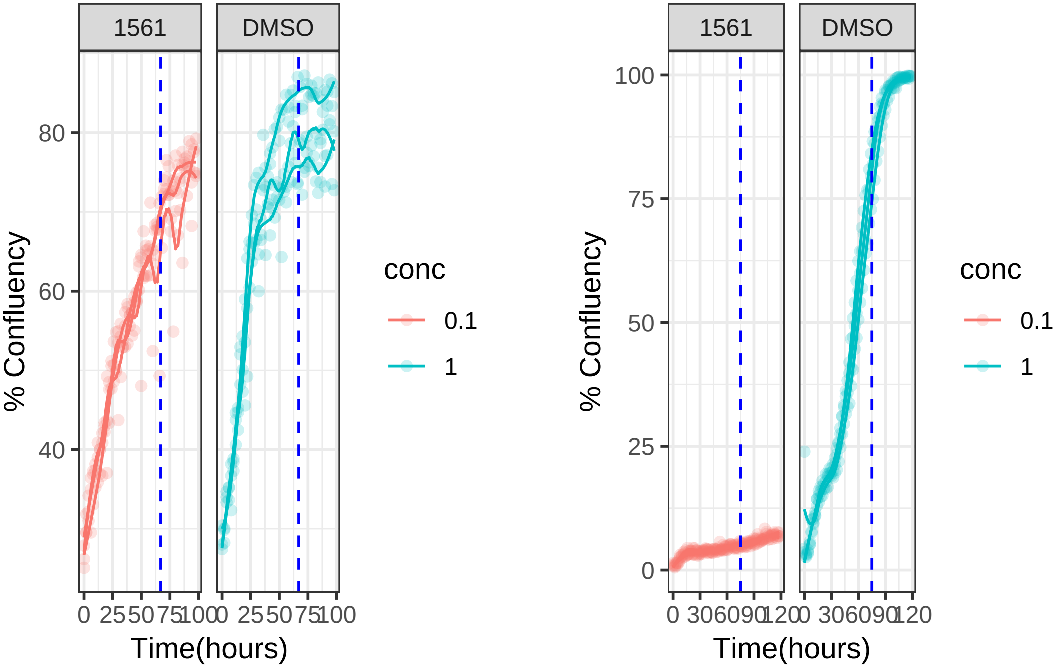

**10 uM 1566 RPE1 Mad2KD**

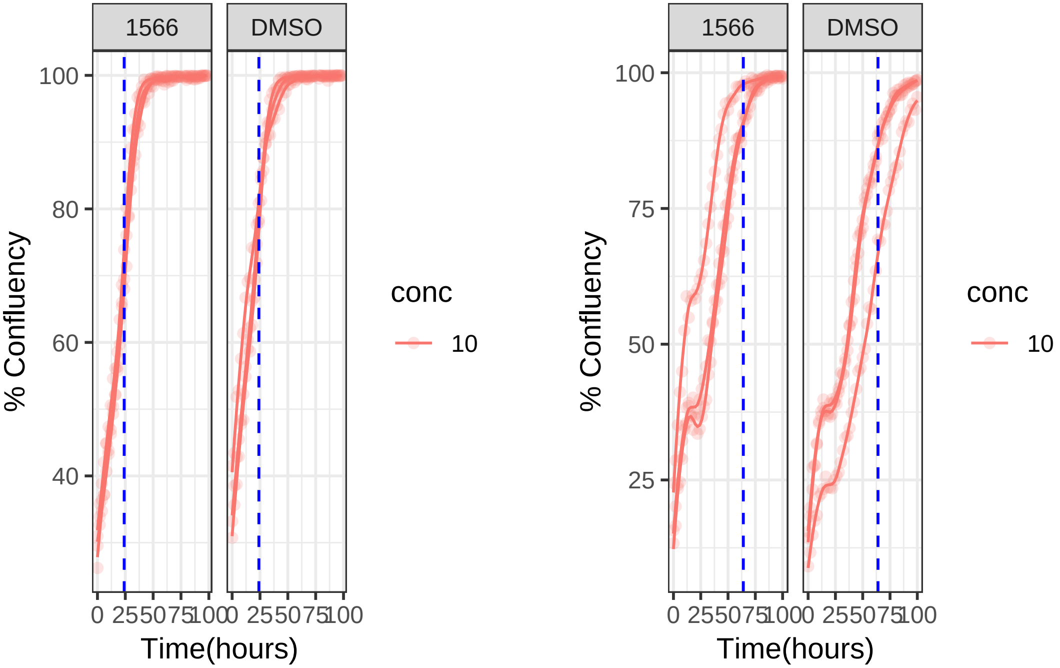

**10 uM 1566 RPE1**

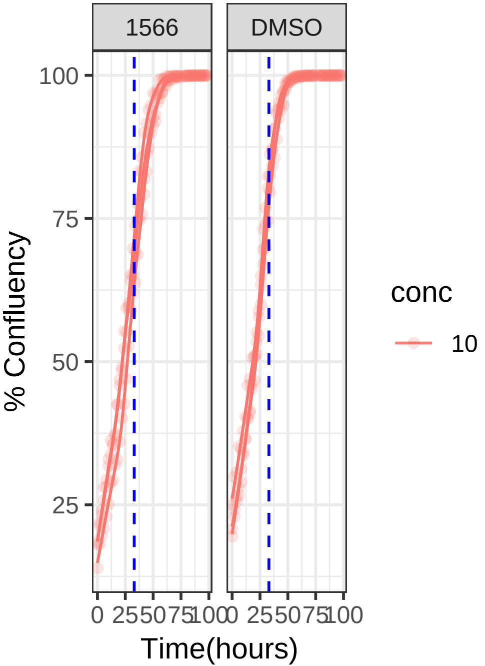


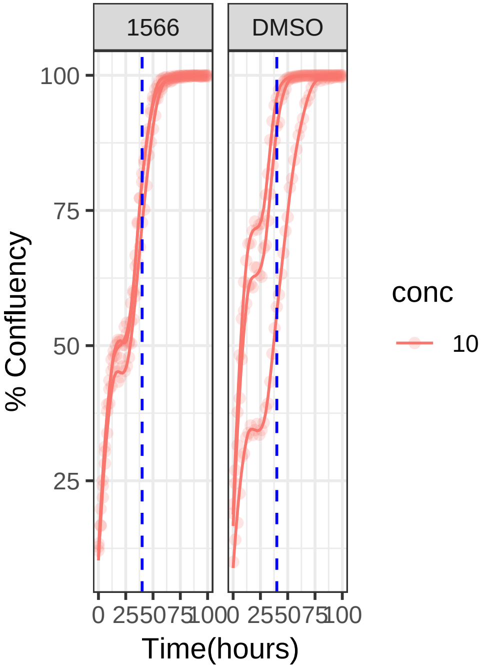

**0.1 uM 1596 RPE1 Mad2KD**

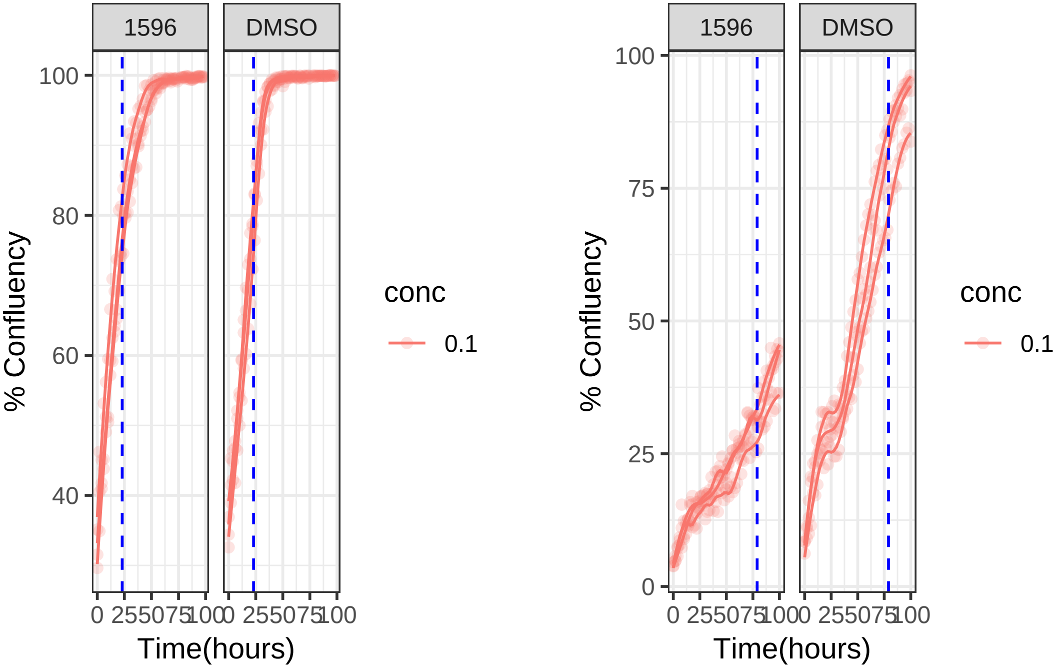

**0.1 uM 1596 RPE1**

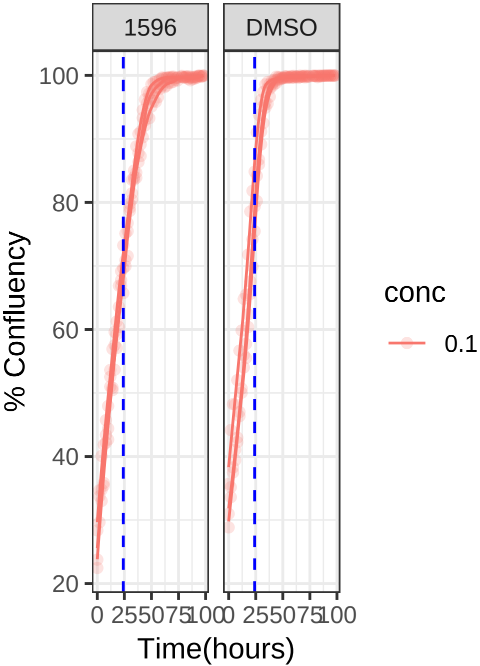


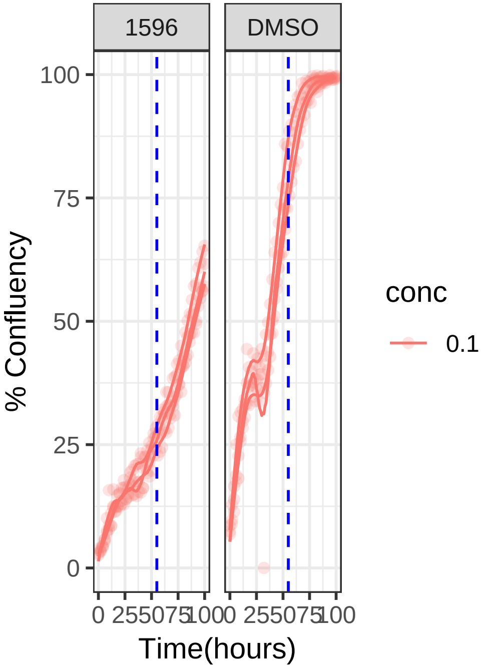

**0.1 uM 1629 RPE1 Mad2KD**

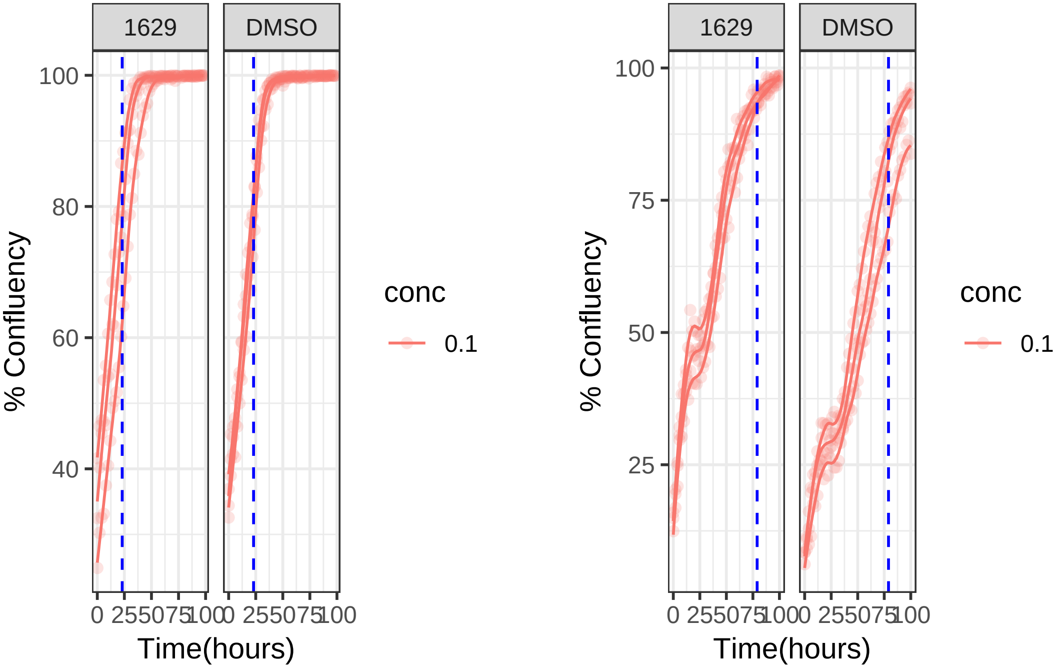

**0.1 uM 1629 RPE1**

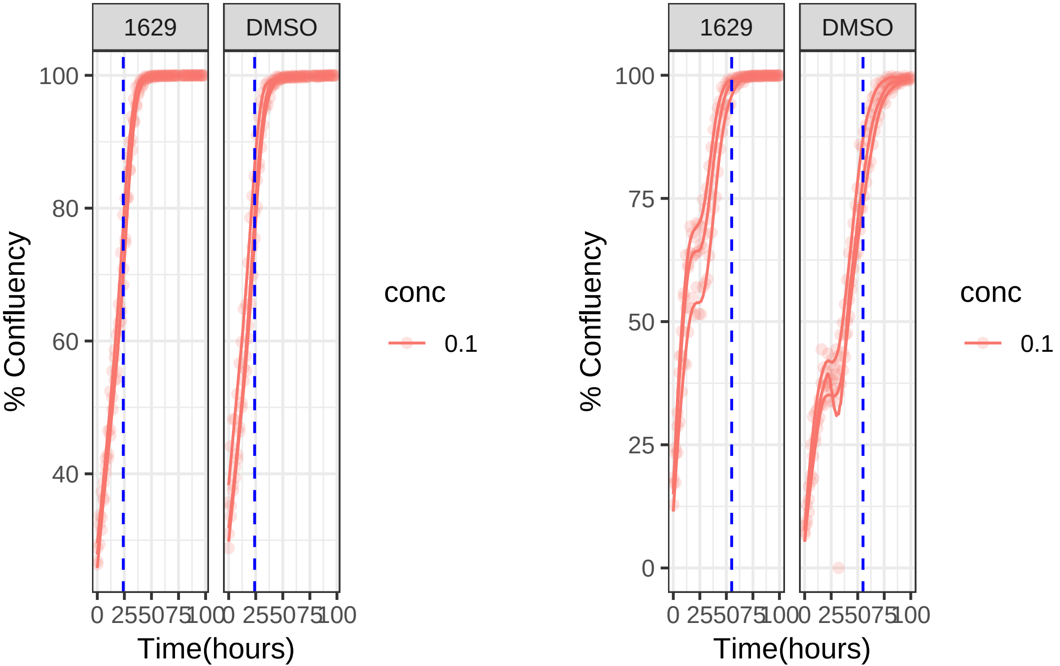

**1 uM 1645 RPE1 Mad2KD**

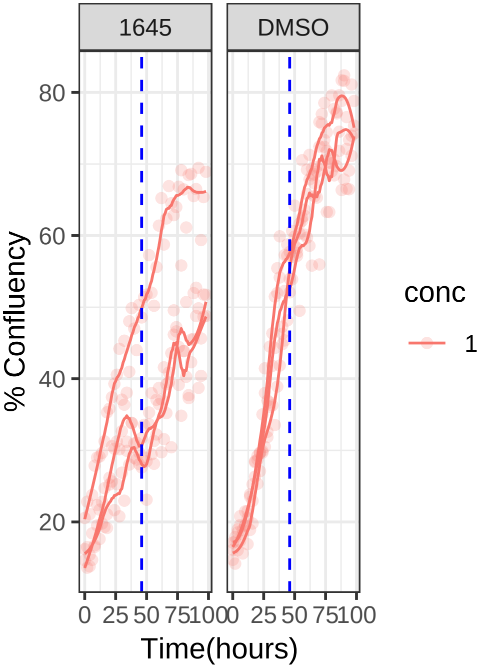


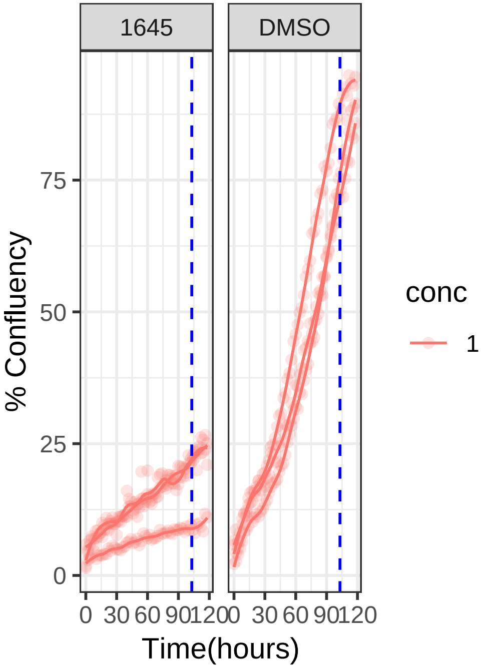

**1 uM 1645 RPE1**

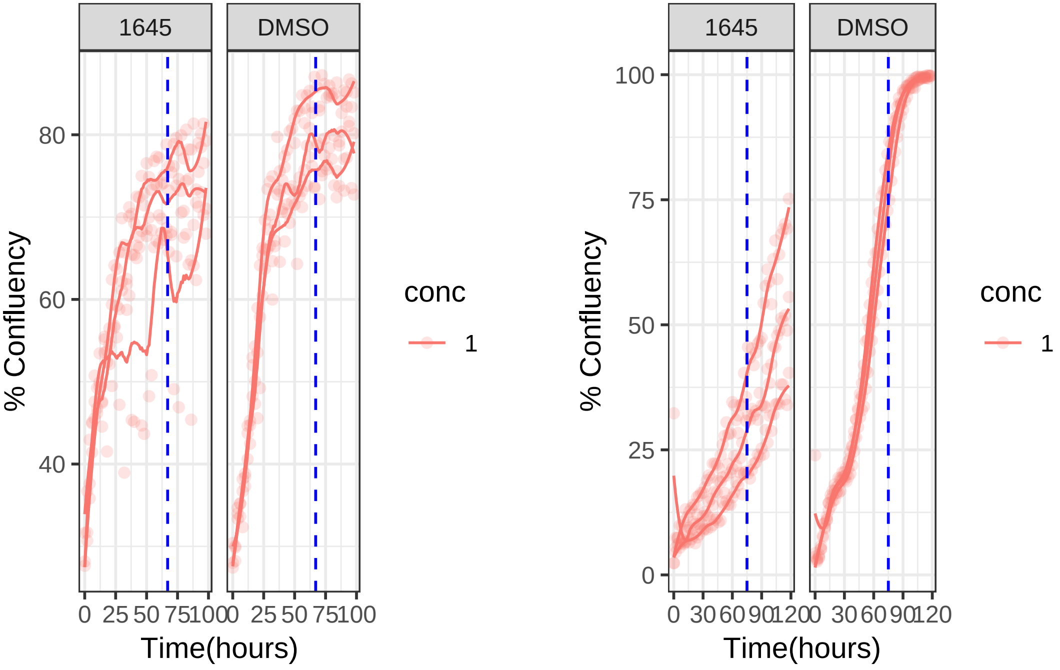

**0.01 uM 1687 RPE1 Mad2KD**

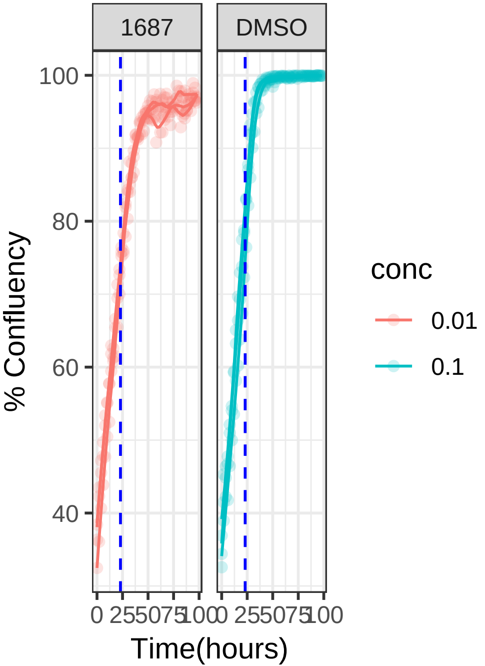


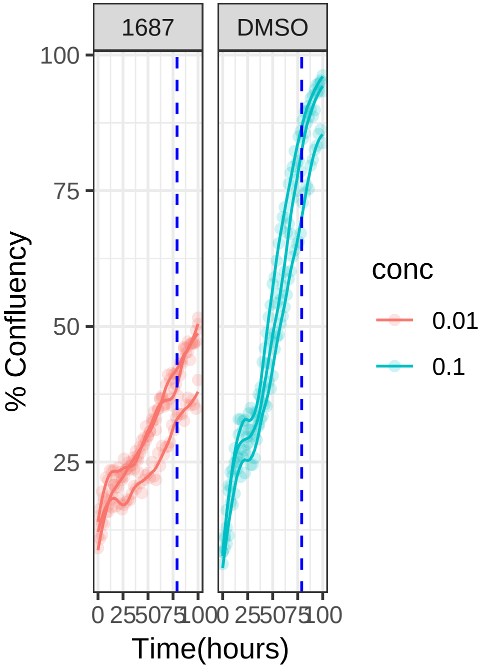

**0.01 uM 1687 RPE1**

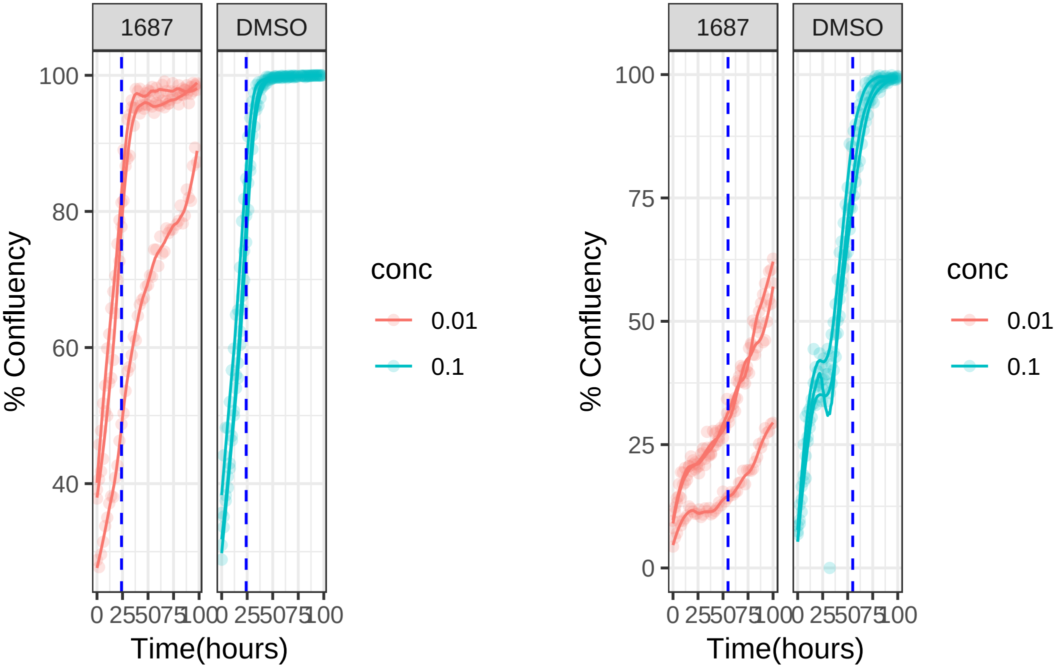

**1 uM 1729 RPE1 Mad2KD**

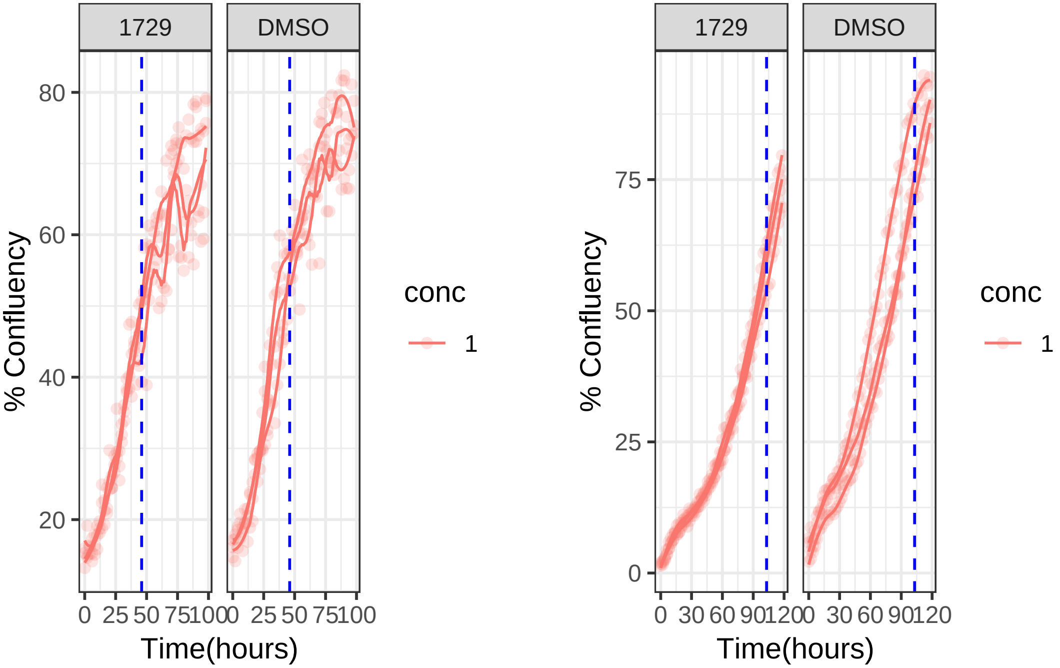

**1 uM 1729 RPE1**

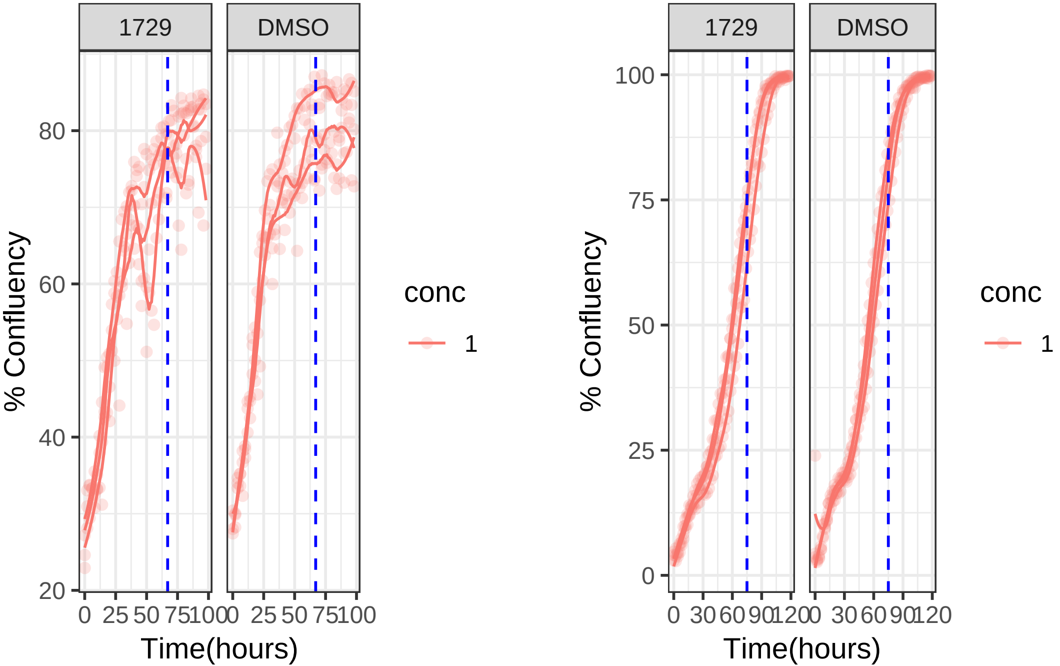

**1 uM 1743 RPE1 Mad2KD**

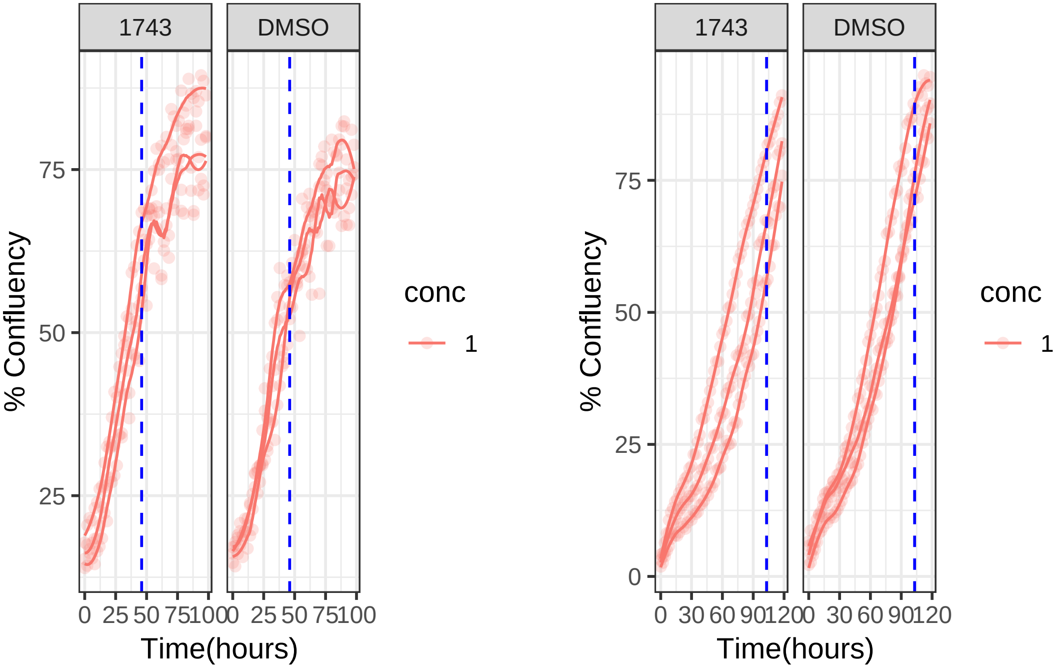

**1 uM 1743 RPE1**

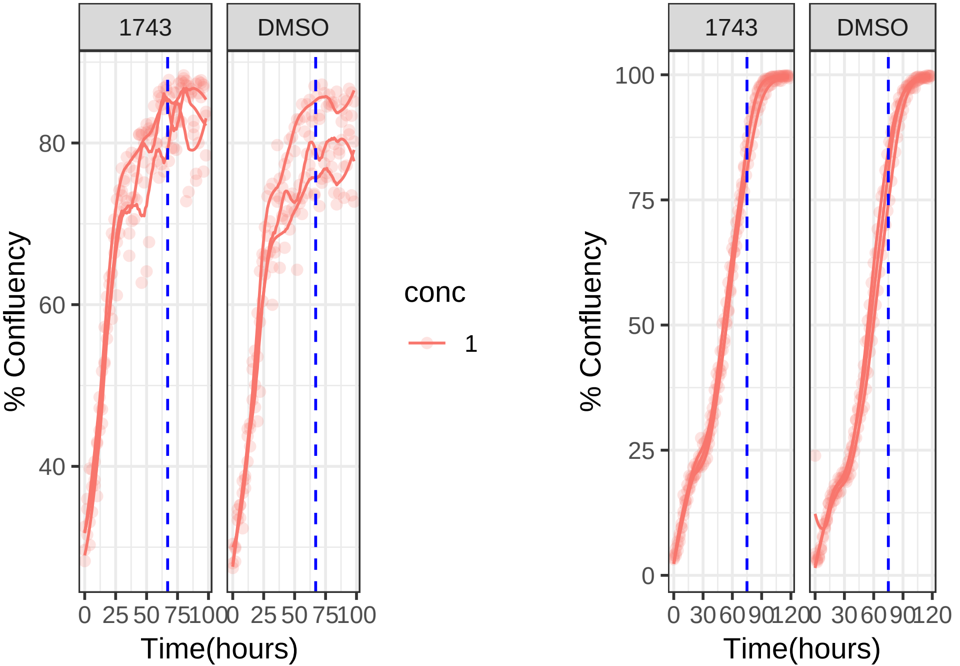

1 uM 1797 RPE1 Mad2KD

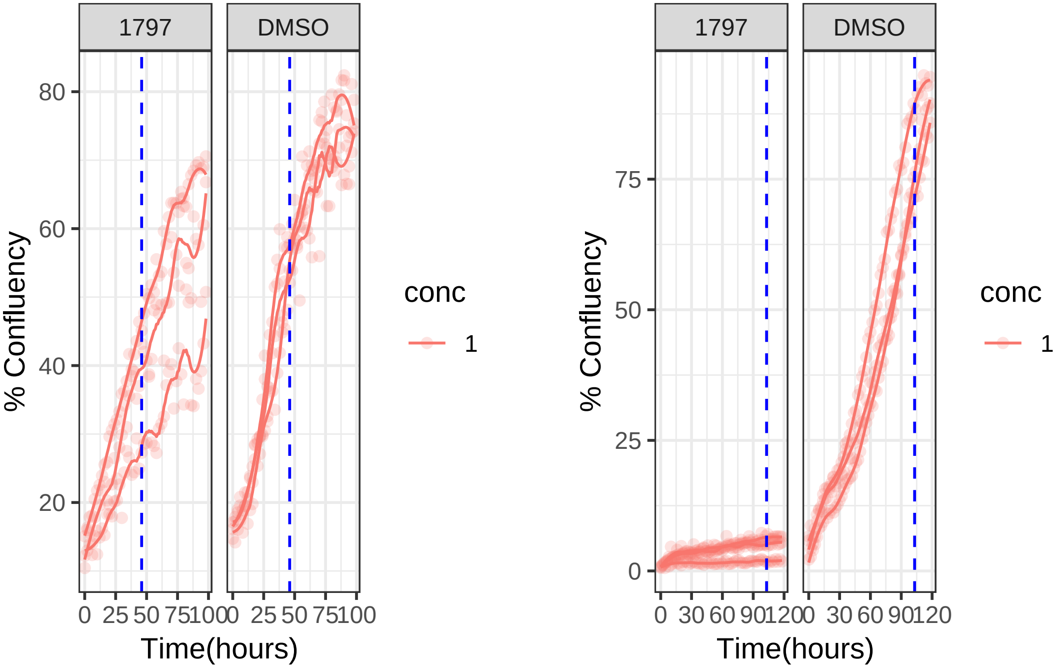

**1 uM 1797 RPE1**

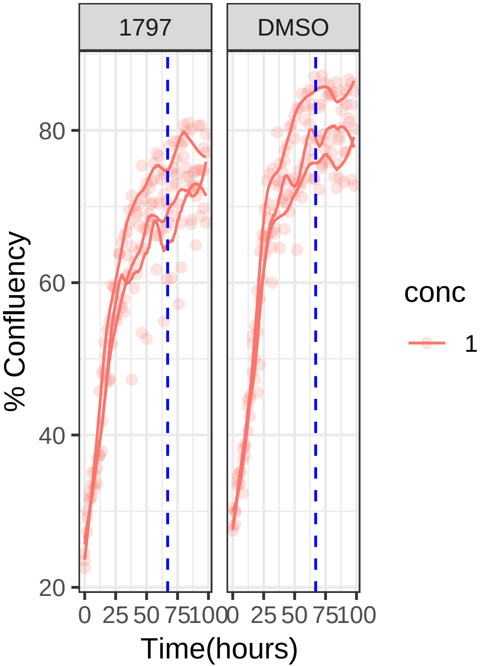


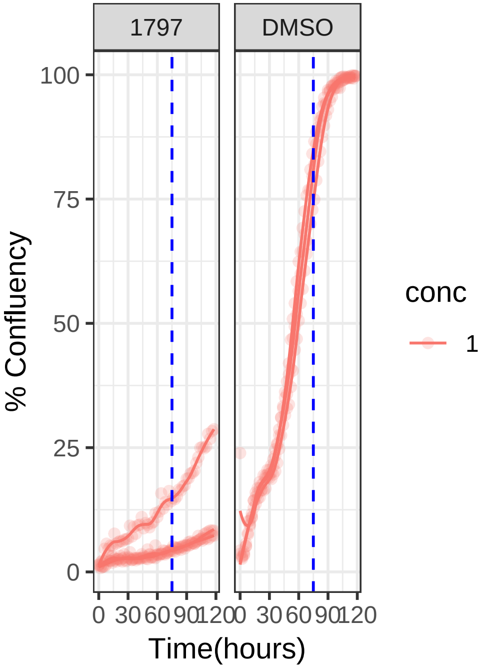

**1 uM 1801 RPE1 Mad2KD**

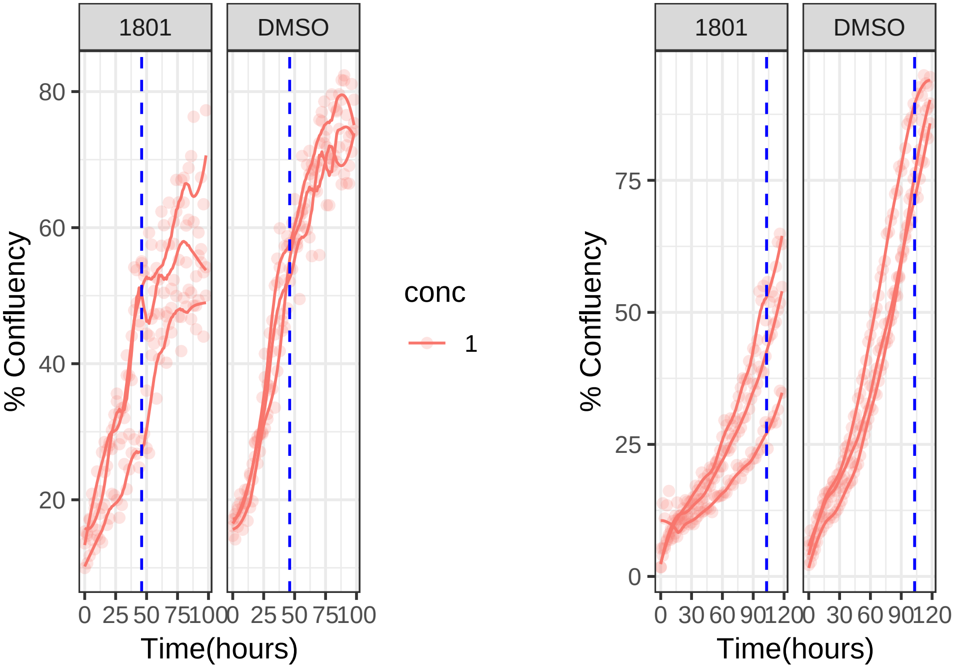

**1 uM 1801 RPE1**

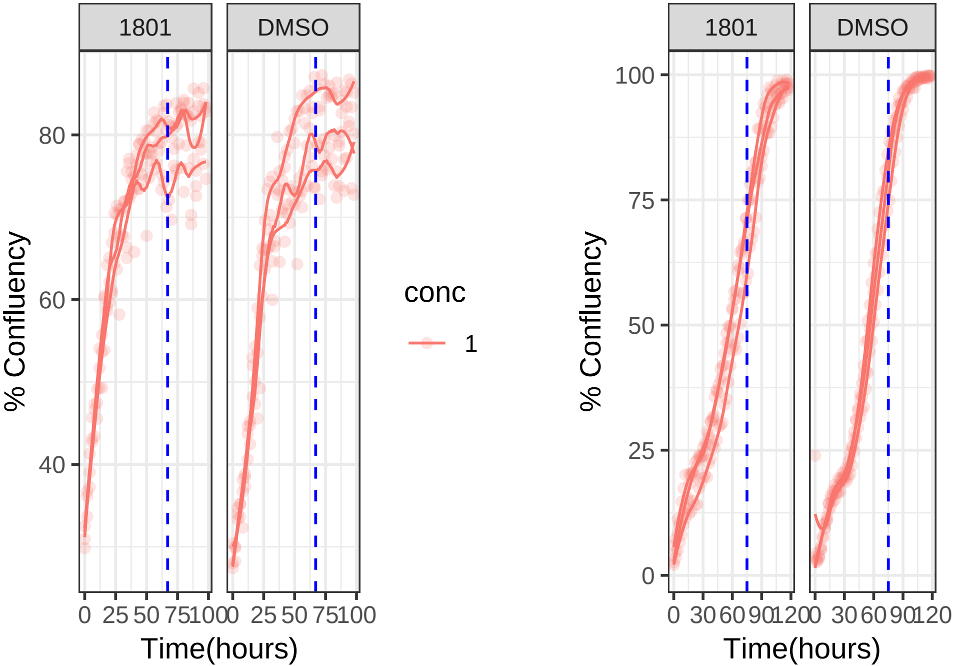

**1 uM 1803 RPE1 Mad2KD**

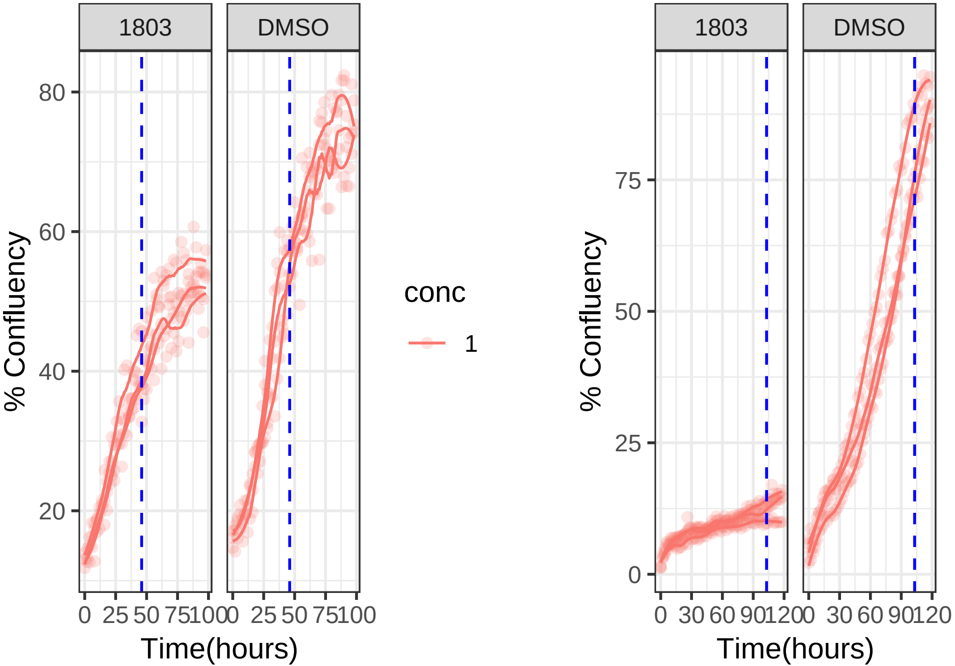

1 uM 1803 RPE1

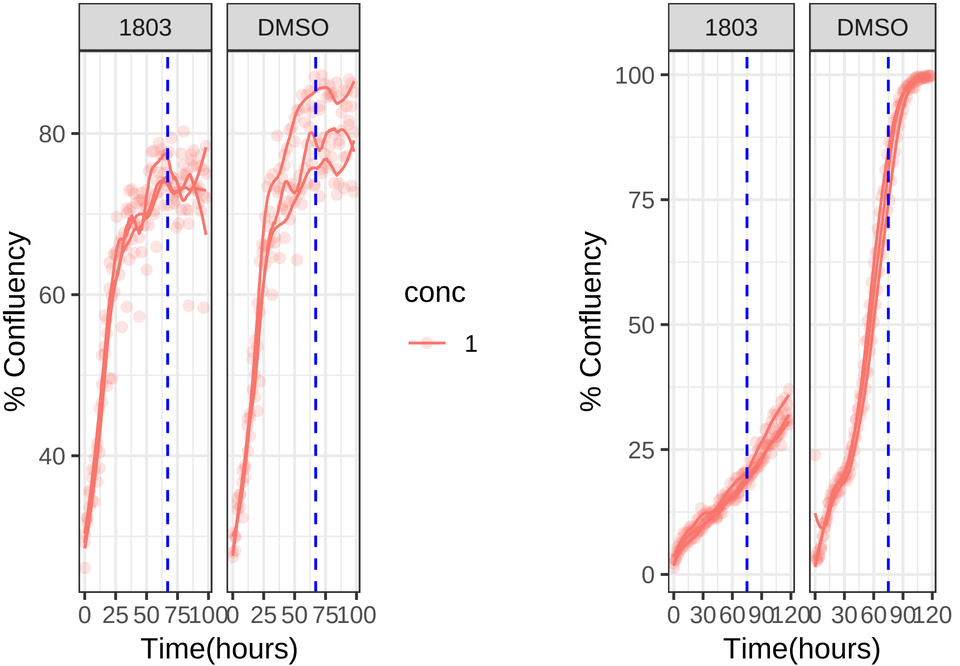

**0.01 uM 1807 RPE1 Mad2KD**

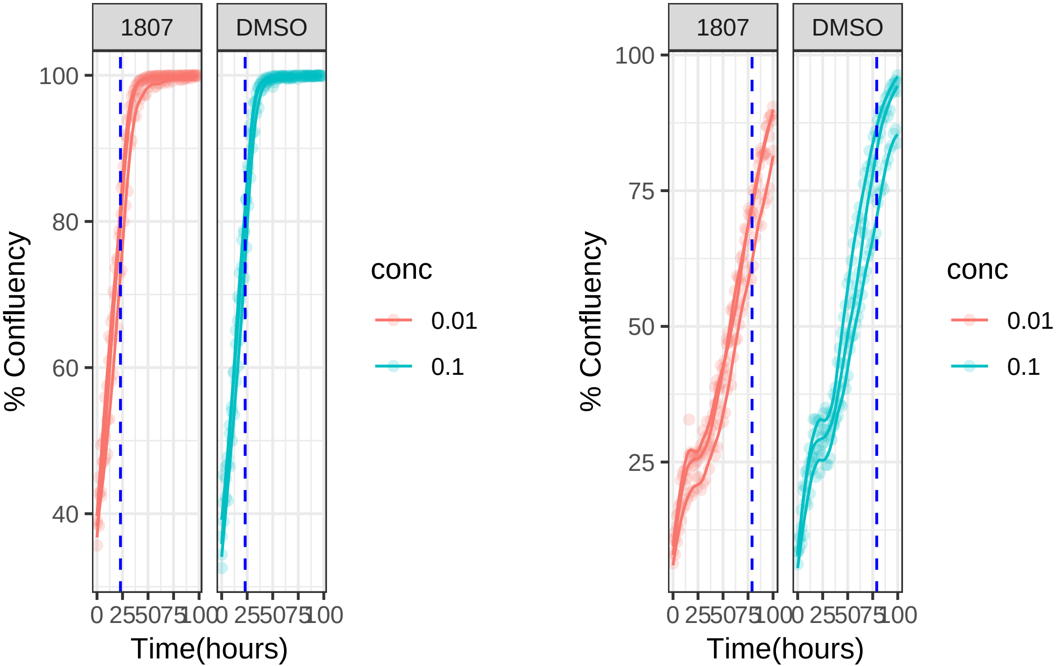

**0.01 uM 1807 RPE1**

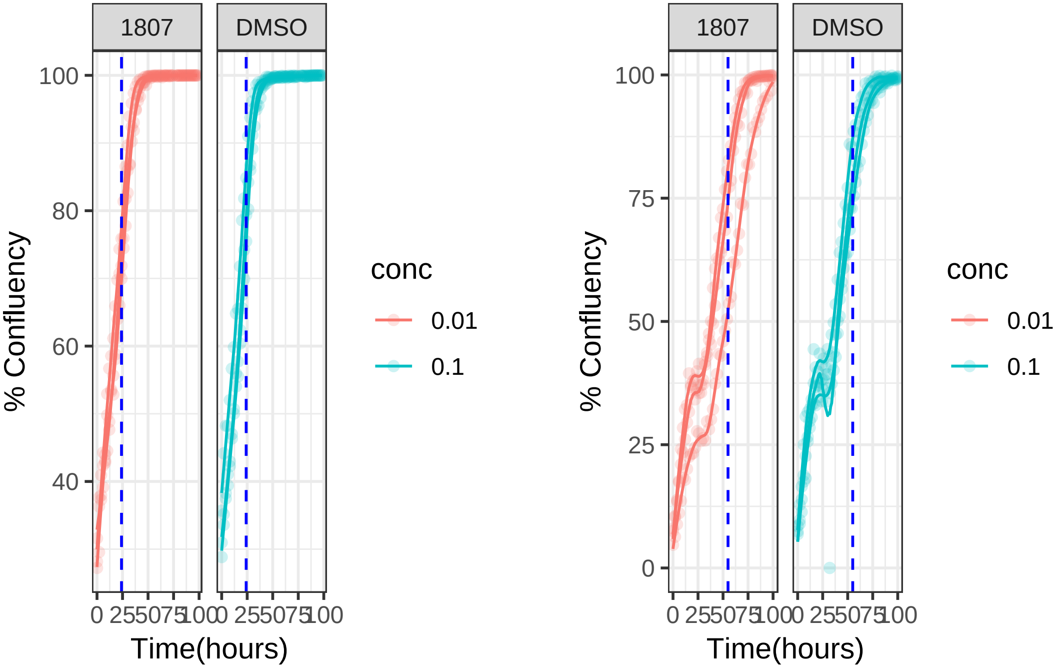

**10 uM 1831 RPE1 Mad2KD**

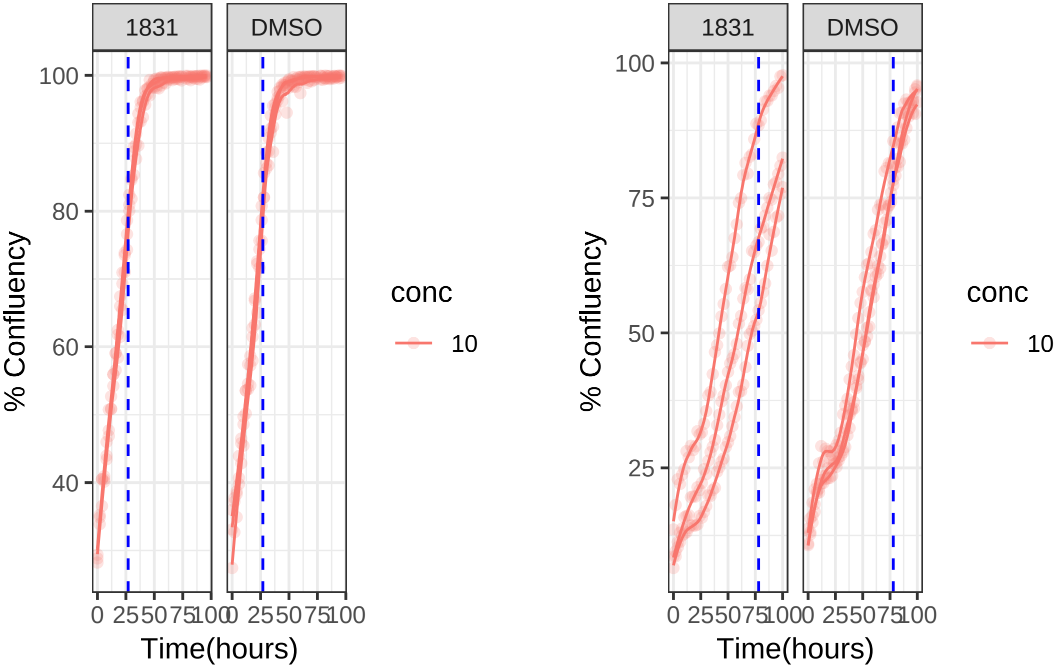

**10 uM 1831 RPE1**

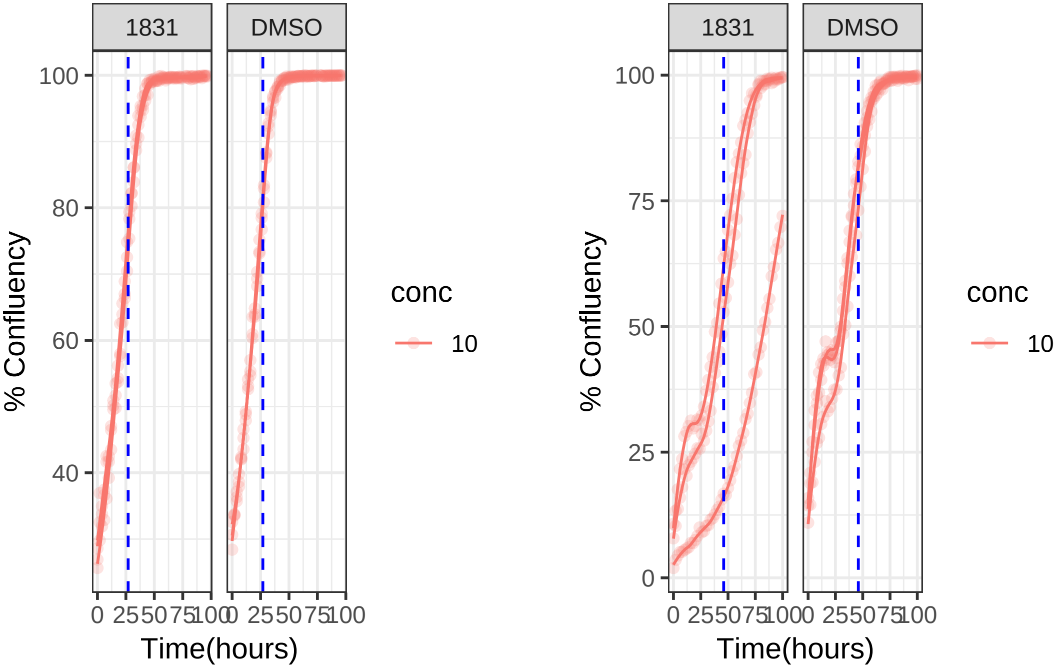

**10 uM 1895 RPE1 Mad2KD**

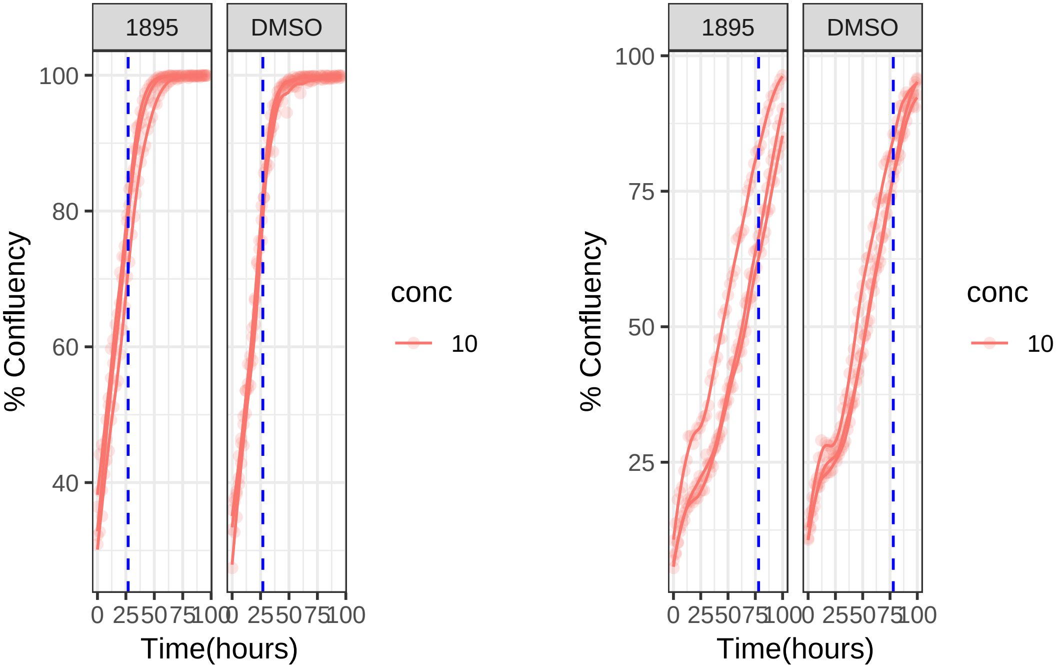

**10 uM 1895 RPE1**

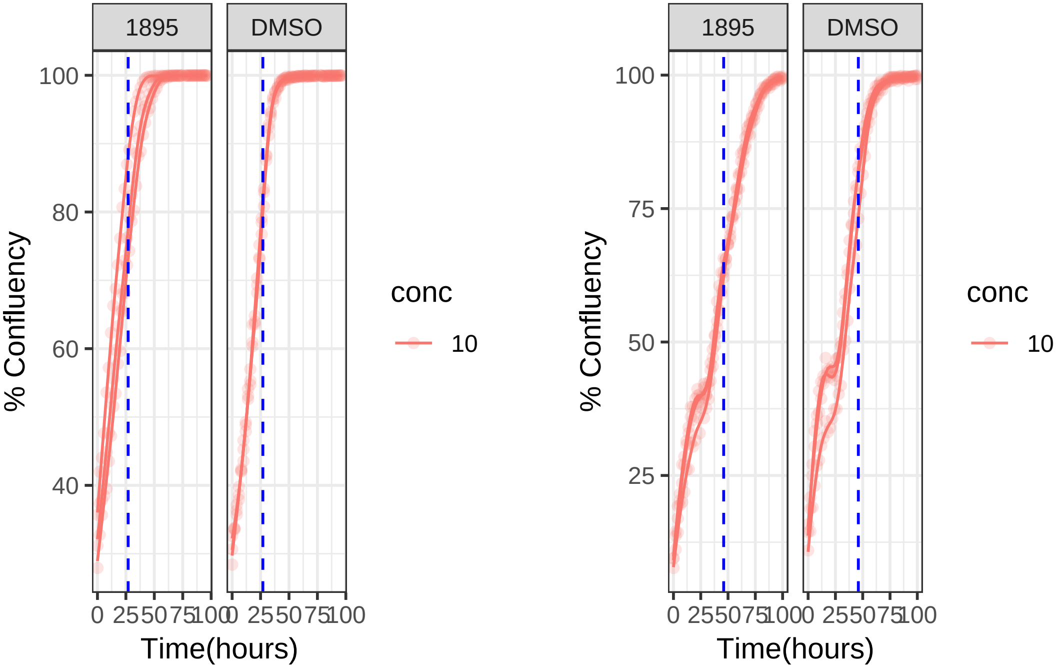

**0.01 uM 1999 RPE1 Mad2KD**

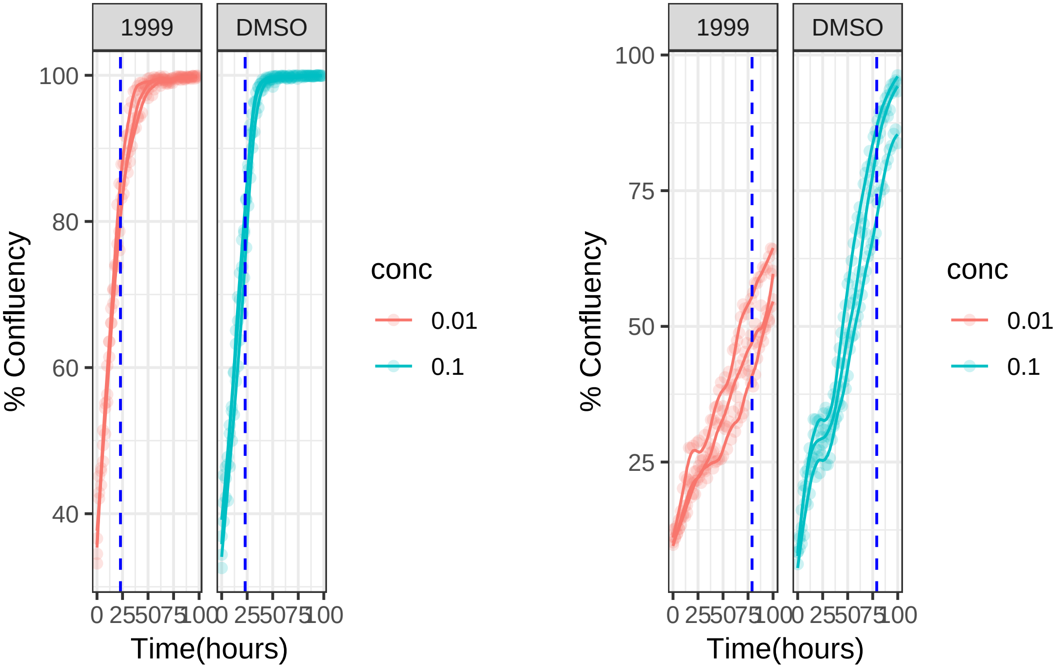

**0.01 uM 1999 RPE1**

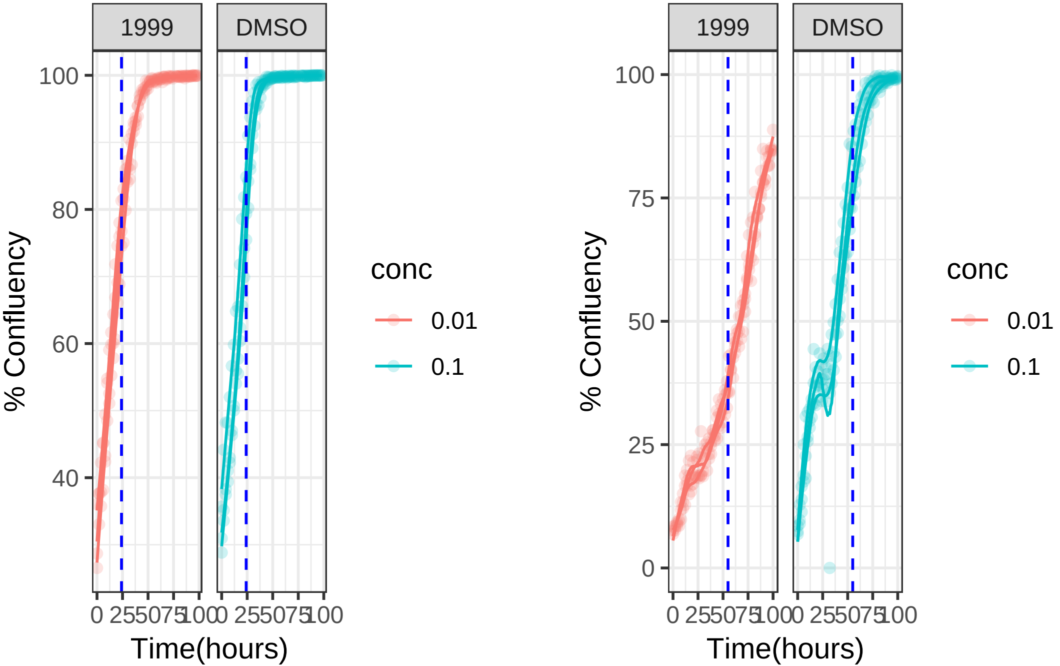

**1 uM 2008 RPE1 Mad2KD**

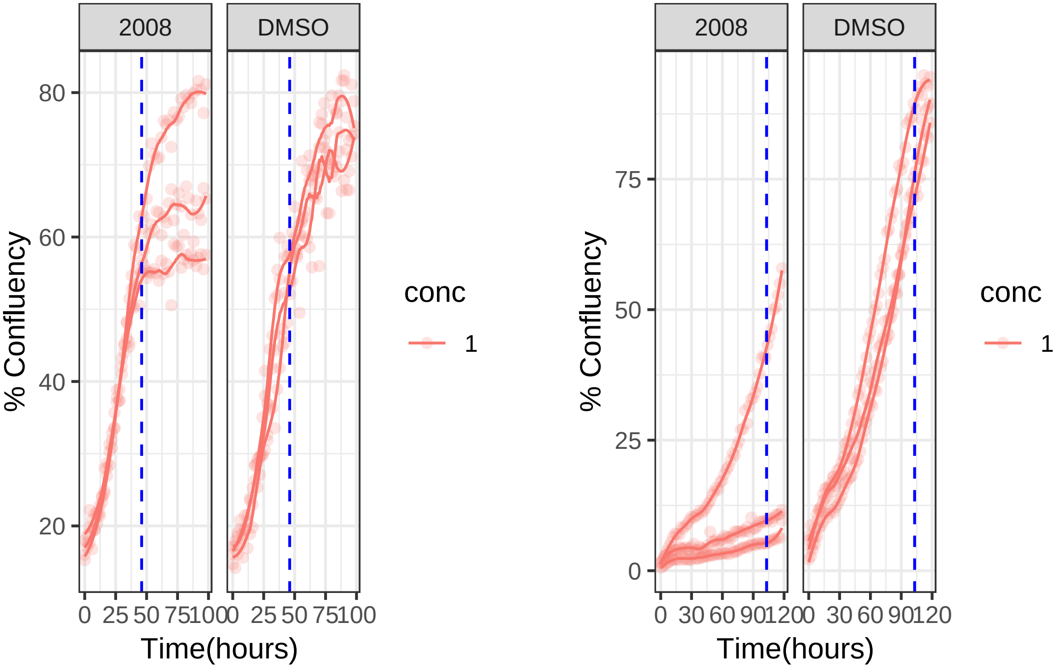

**1 uM 2008 RPE1**

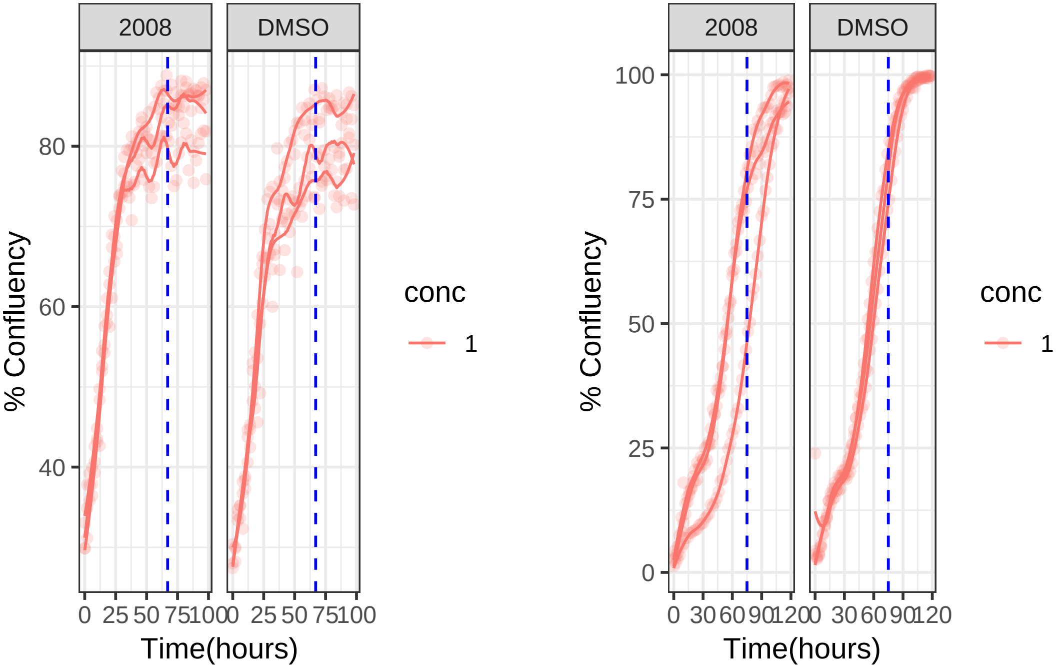

1 uM 2014 RPE1 Mad2KD

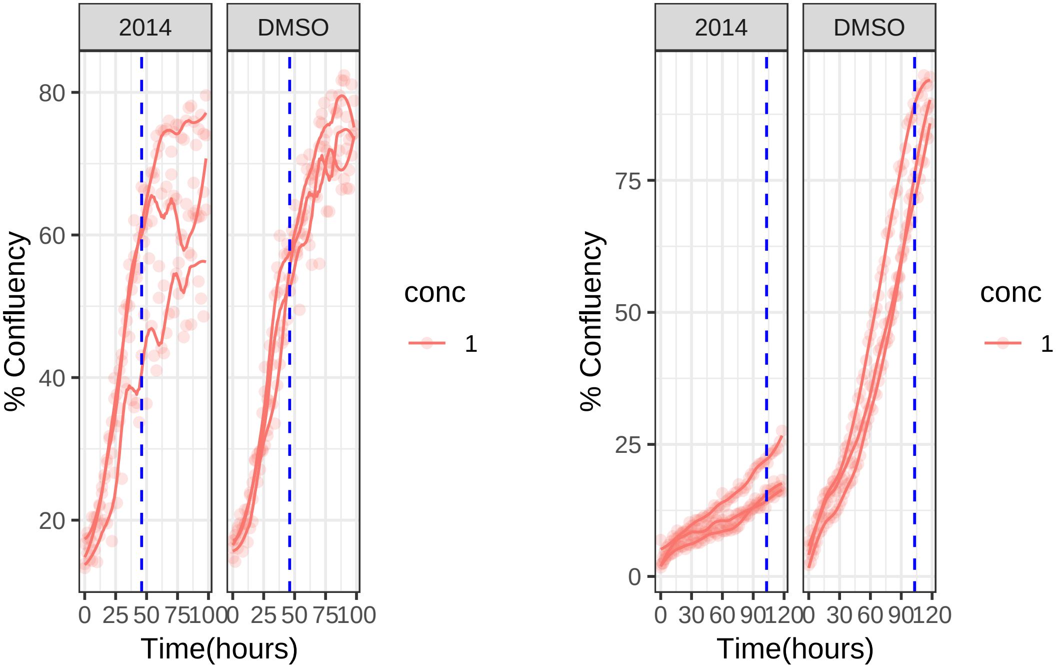

1 uM 2014 RPE1

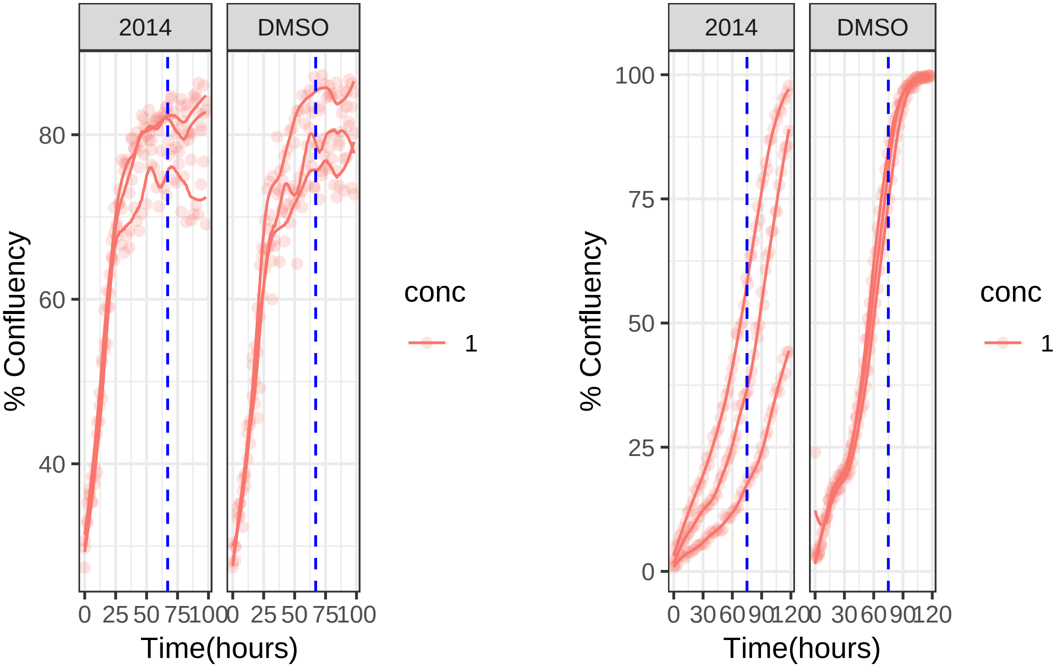

**10 uM 2084 RPE1 Mad2KD**

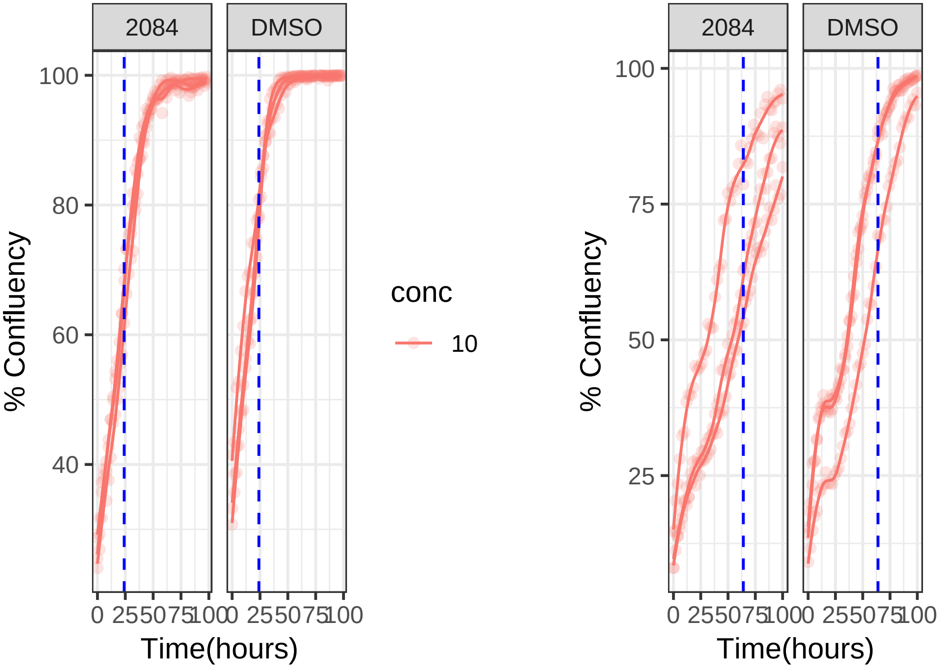

**10 uM 2084 RPE1**

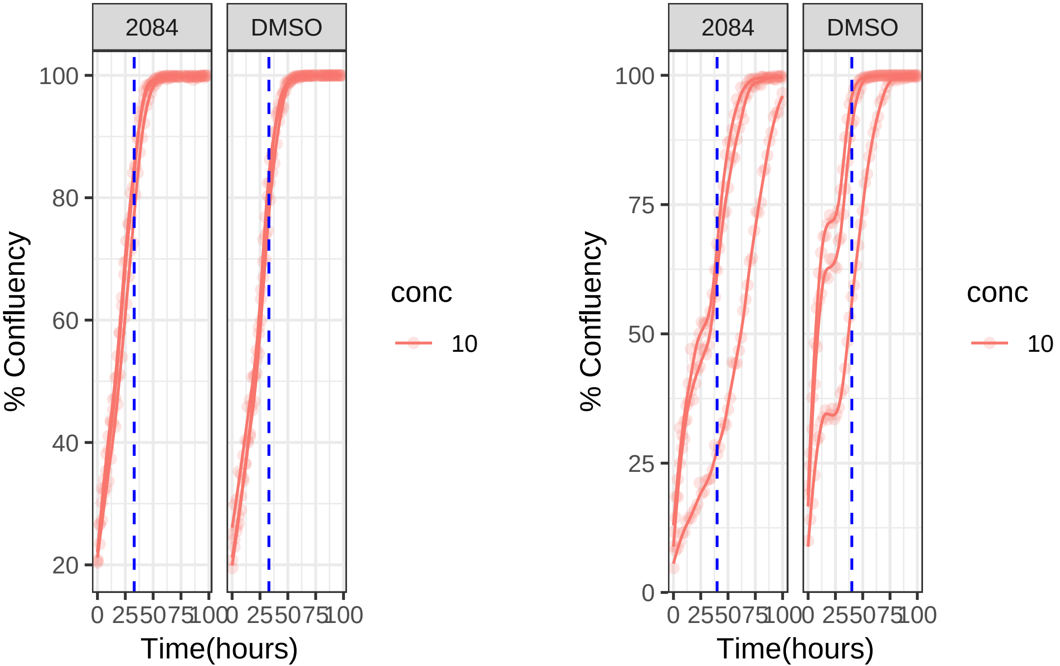

**0.01 uM 2120 RPE1 Mad2KD**

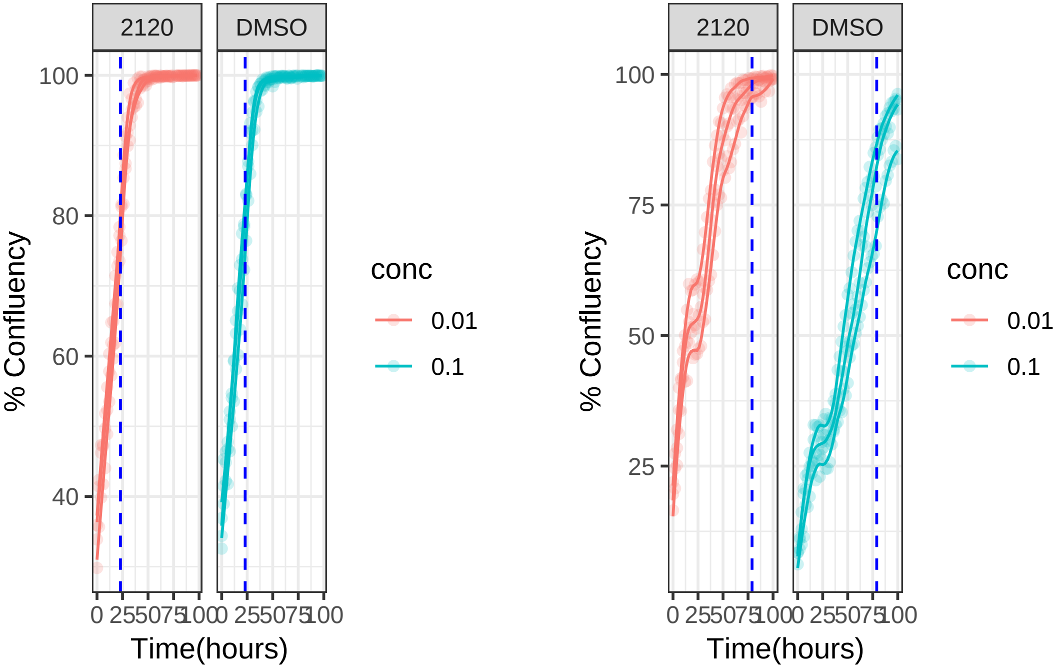

**0.01 uM 2120 RPE1**

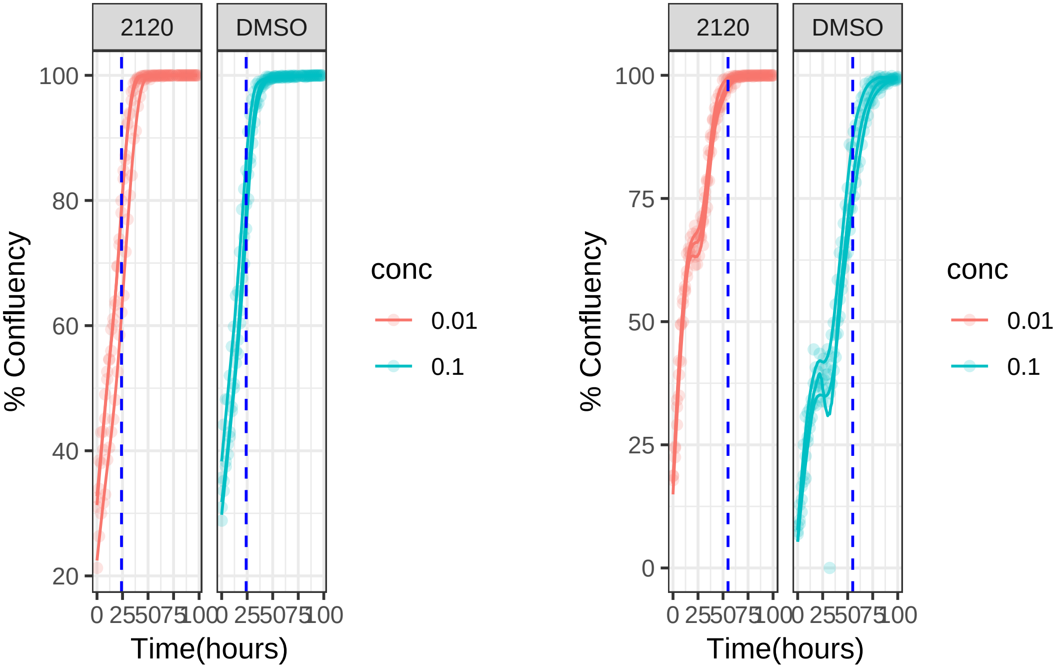

**0.01 uM 2142 RPE1 Mad2KD**

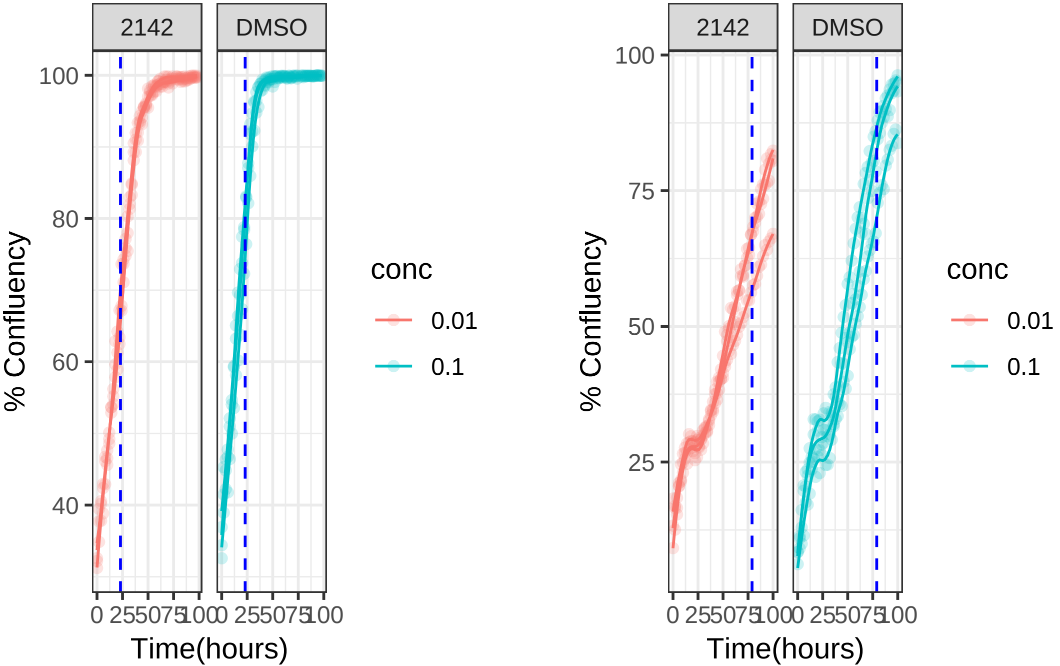

**0.01 uM 2142 RPE1**

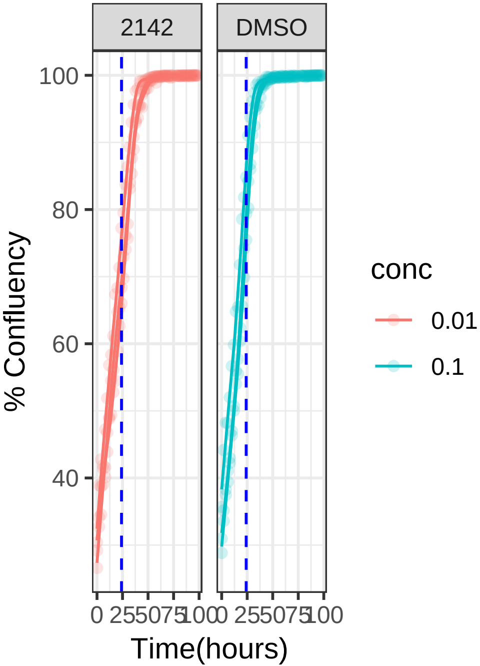


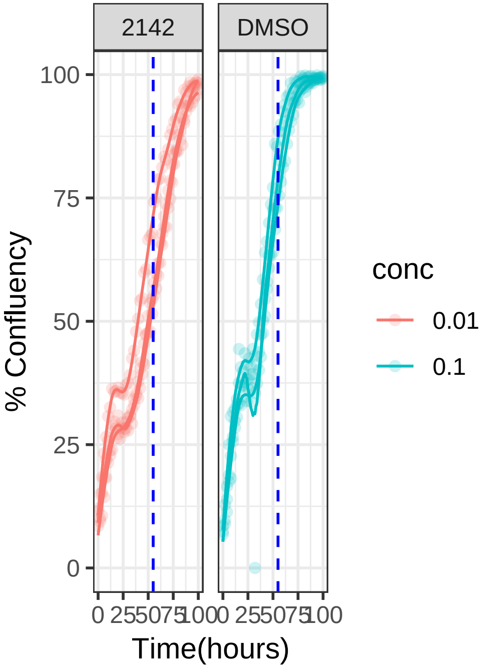

**10 uM 2159 RPE1 Mad2KD**

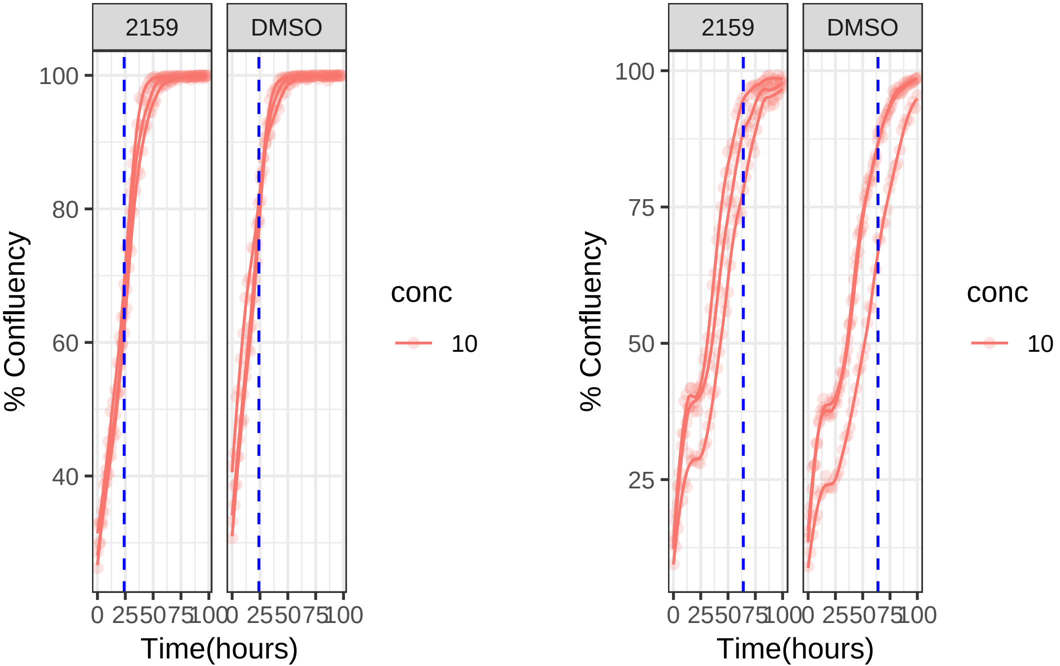

**10 uM 2159 RPE1**

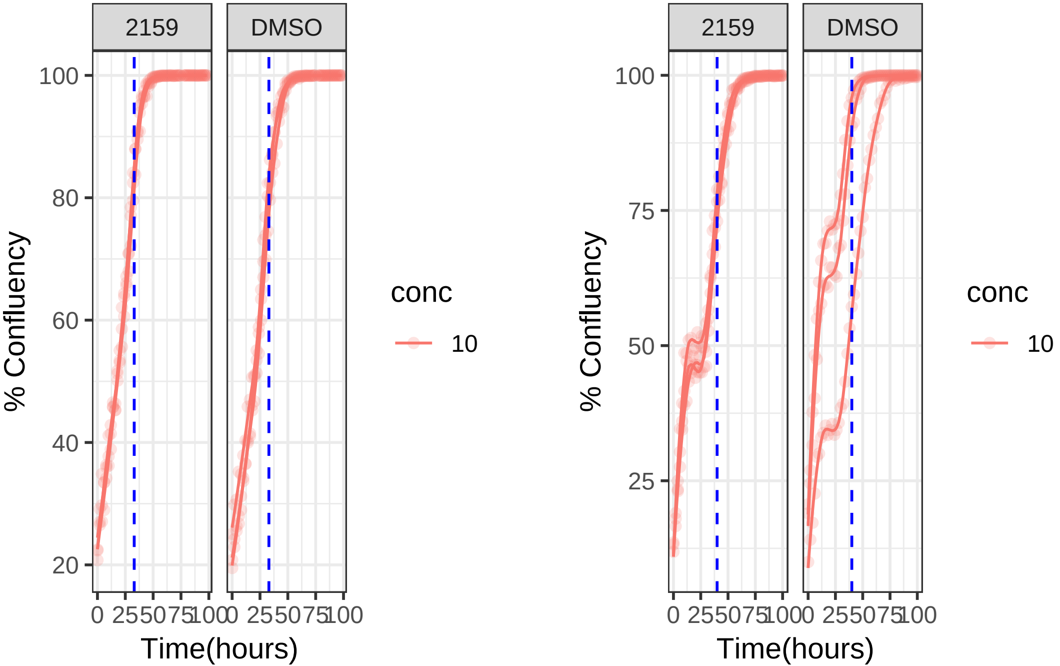

**0.1 uM 2160 RPE1 Mad2KD**

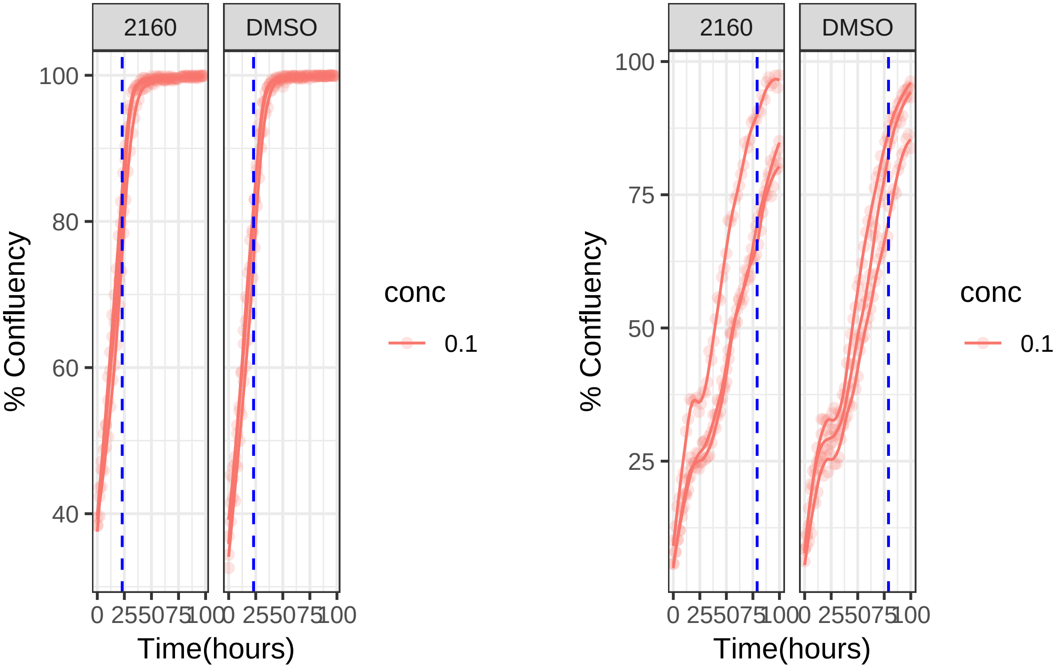

**0.1 uM 2160 RPE1**

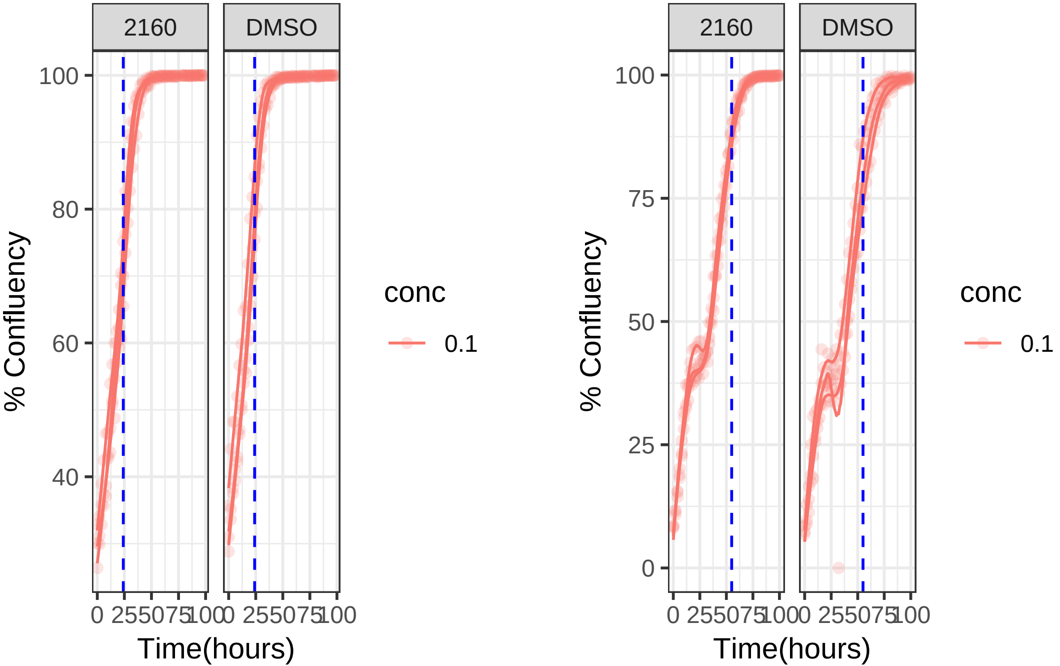

**10 uM 2170 RPE1 Mad2KD**

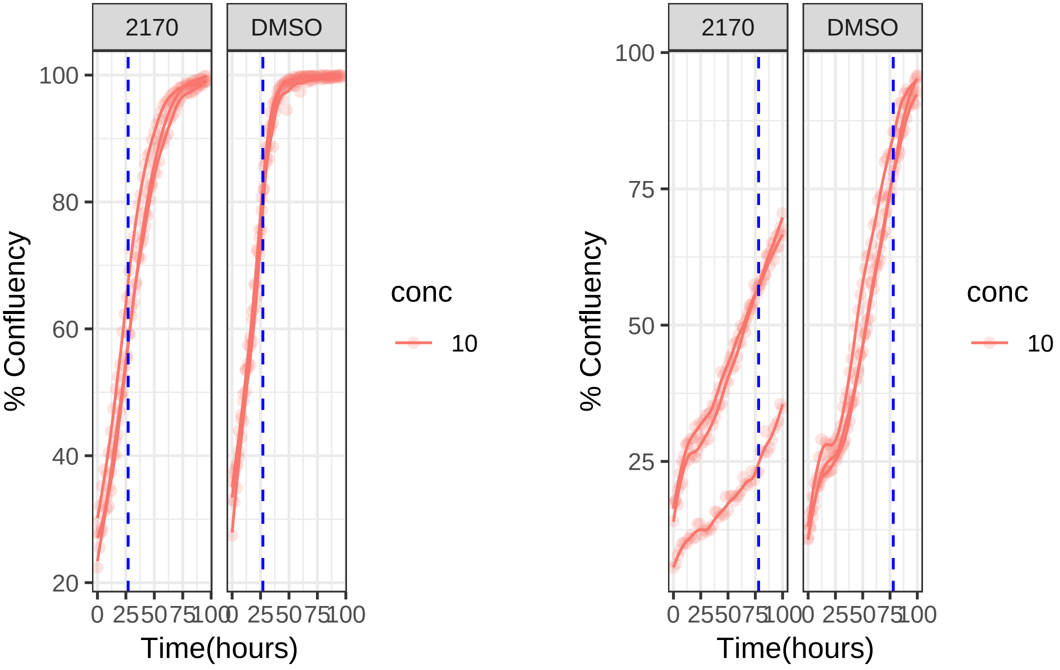

**10 uM 2170 RPE1**

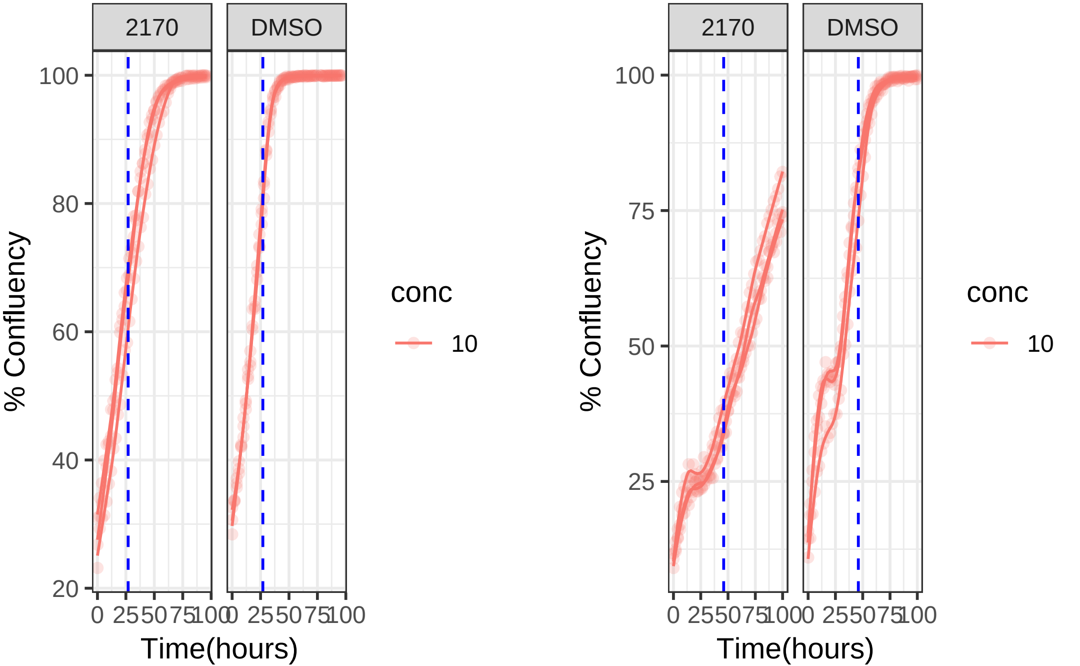

**10 uM 2180 RPE1 Mad2KD**

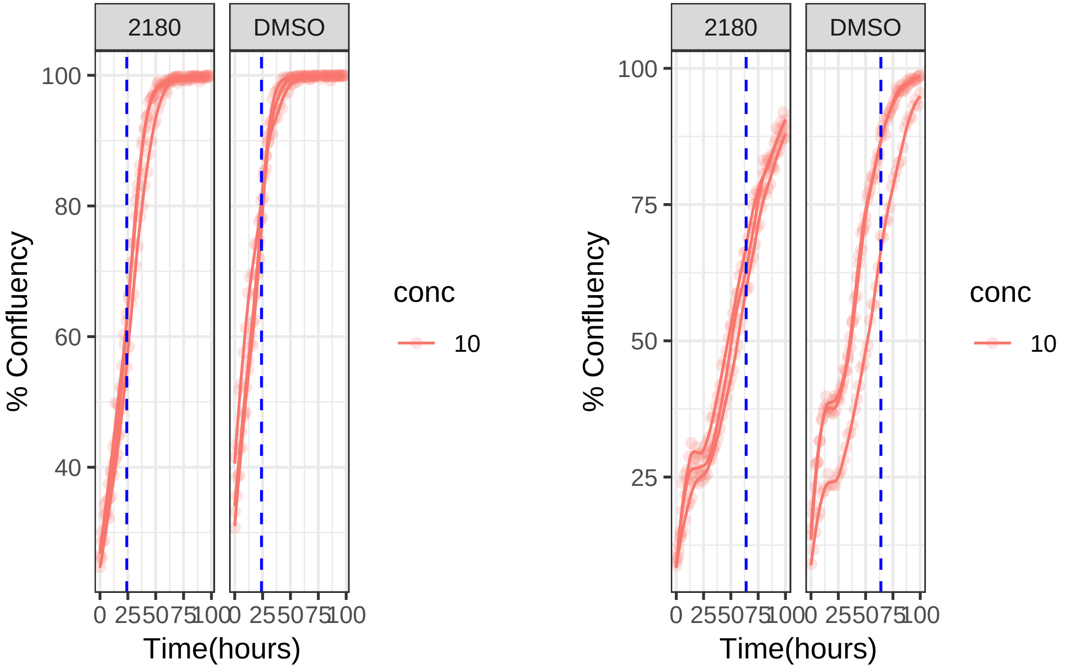

**10 uM 2180 RPE1**

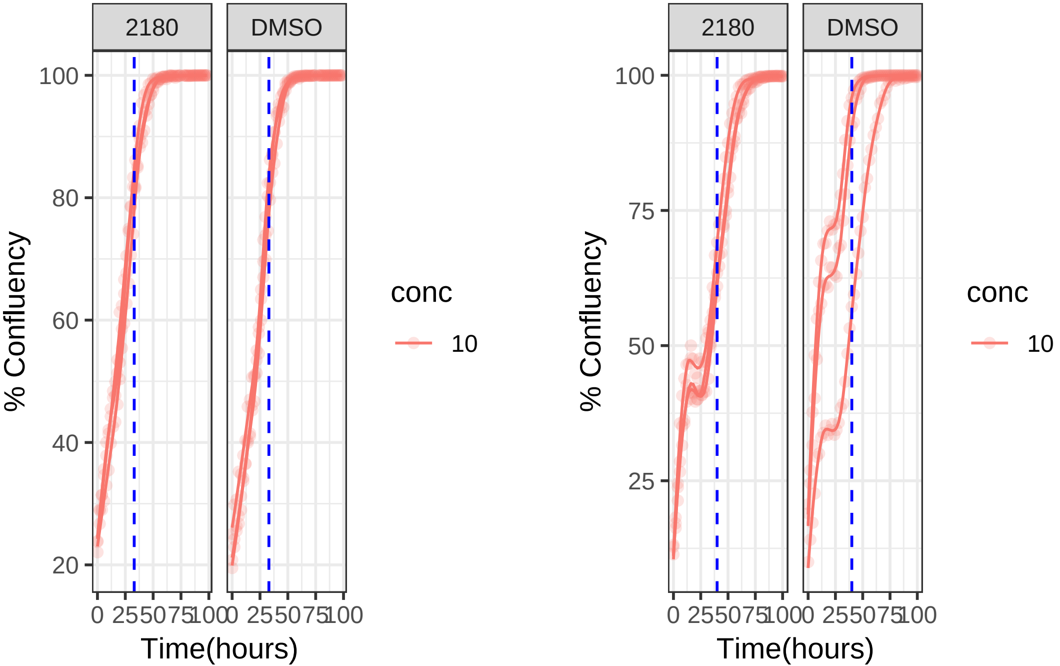

**1 uM 2188 RPE1 Mad2KD**

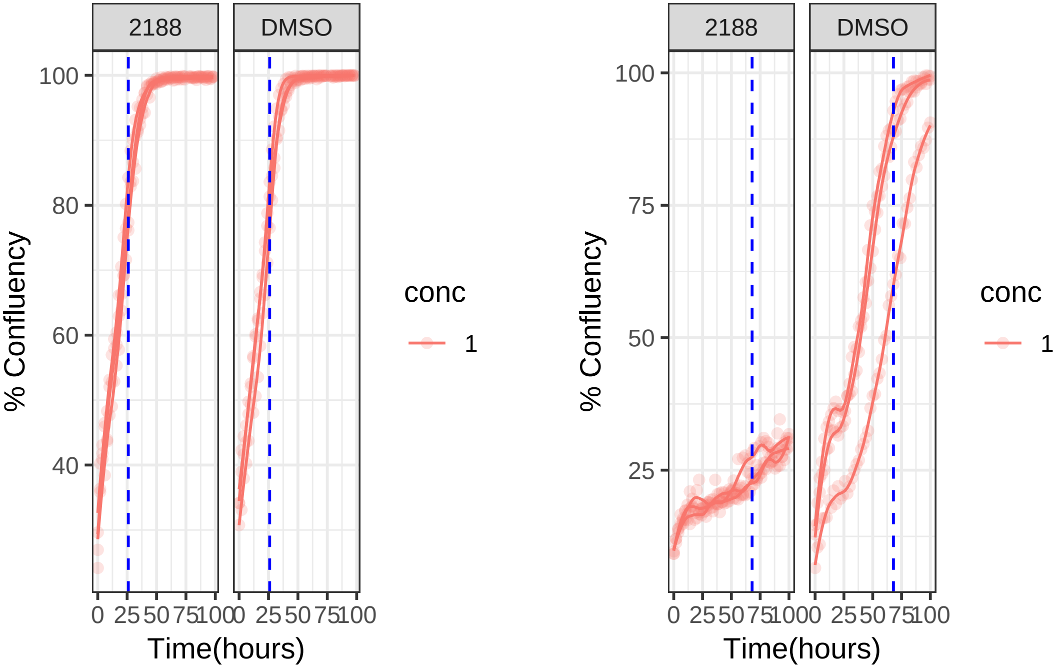

**1 uM 2188 RPE1**

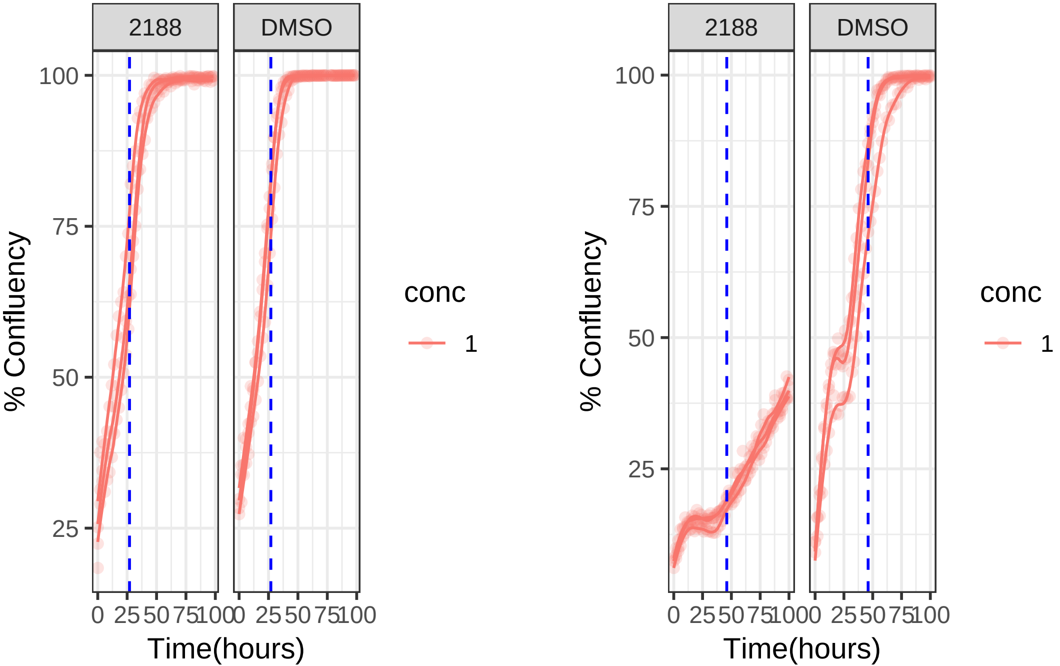

**0.001 uM 2198 RPE1 Mad2KD**

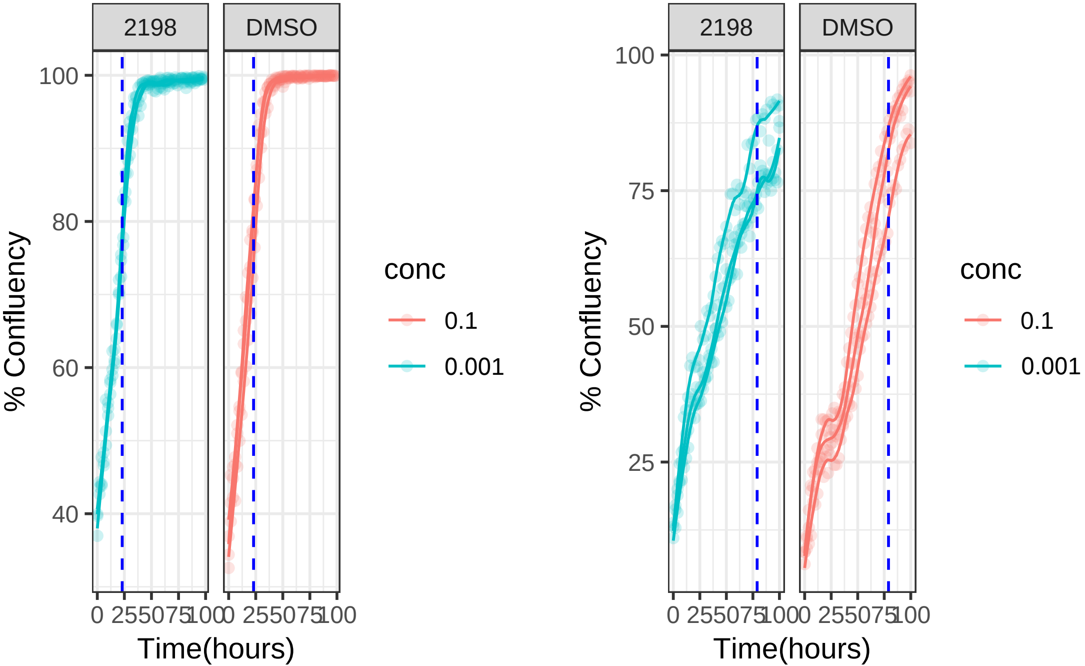

**0.001 uM 2198 RPE1**

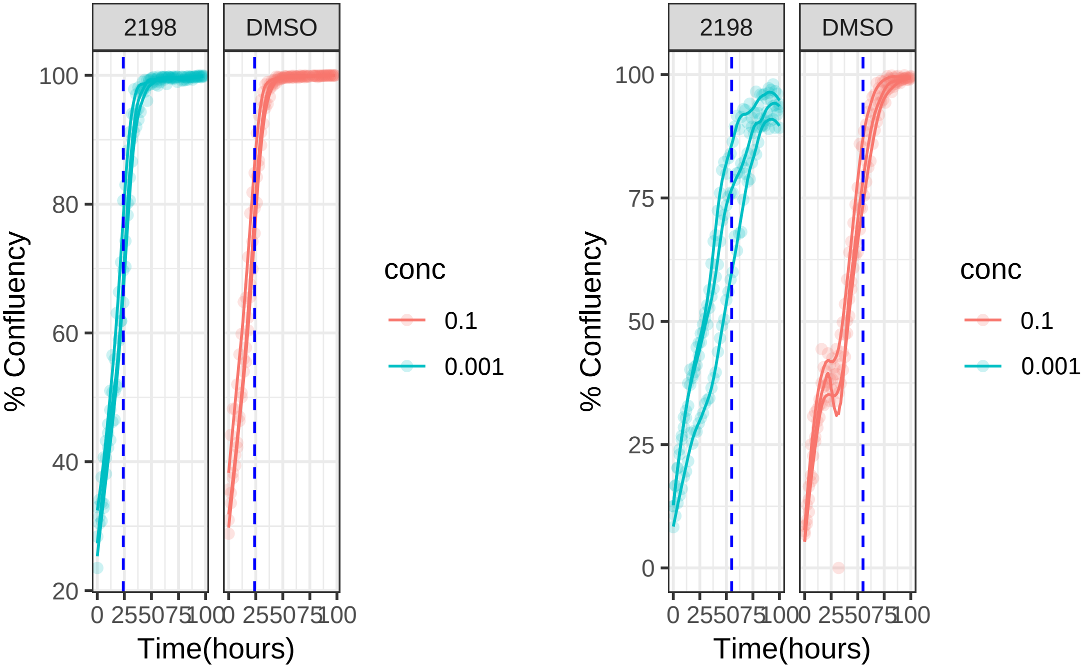

**10 uM 2210 RPE1 Mad2KD**

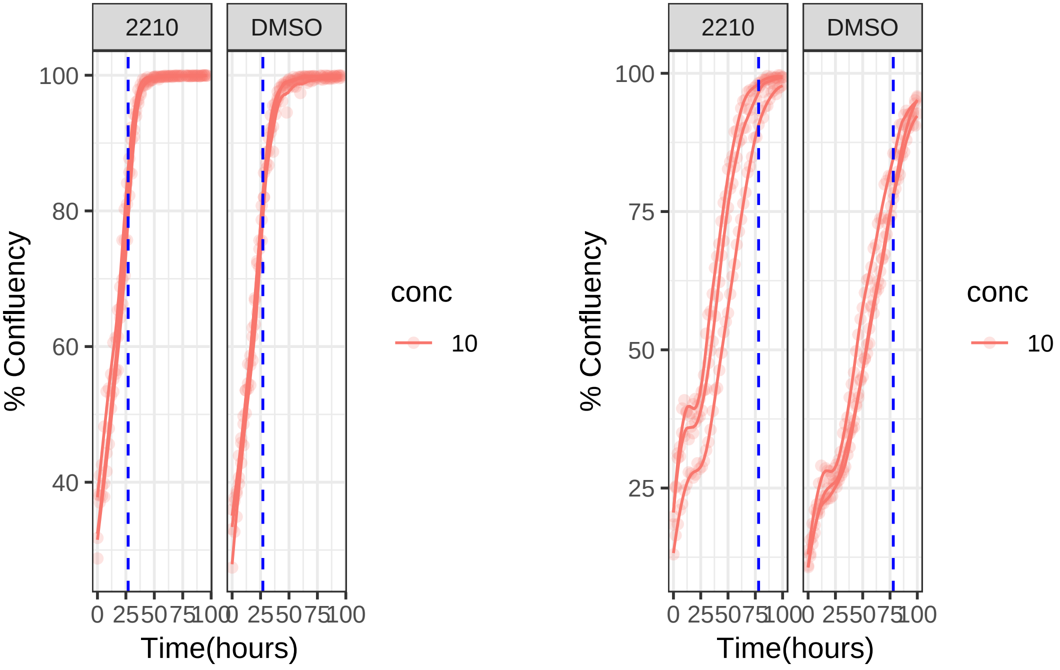

**10 uM 2210 RPE1**

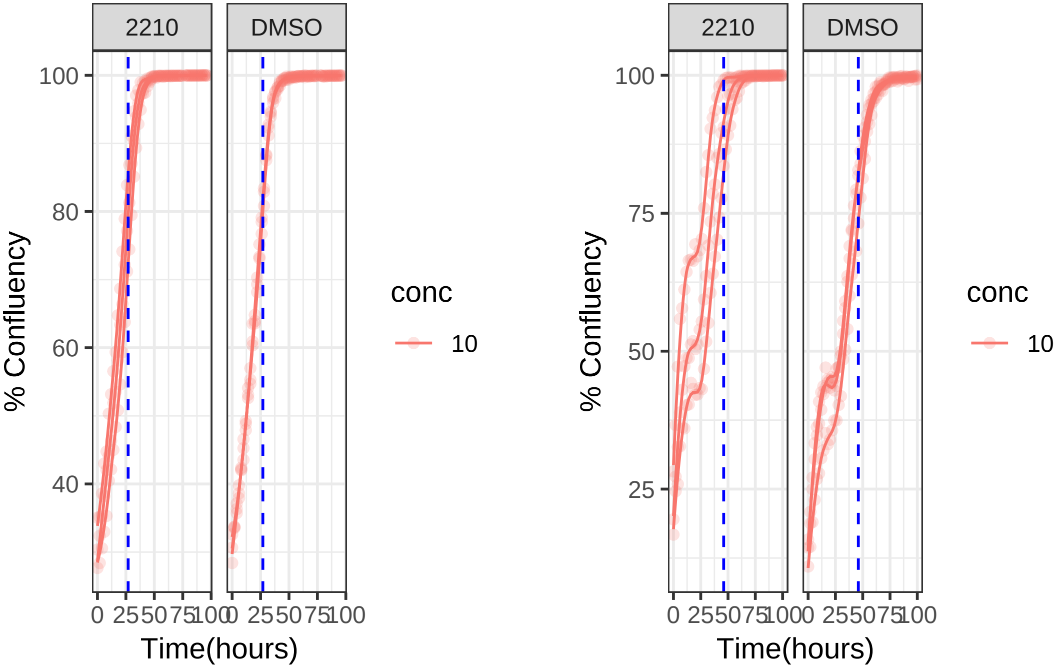

**10 uM 2223 RPE1 Mad2KD**

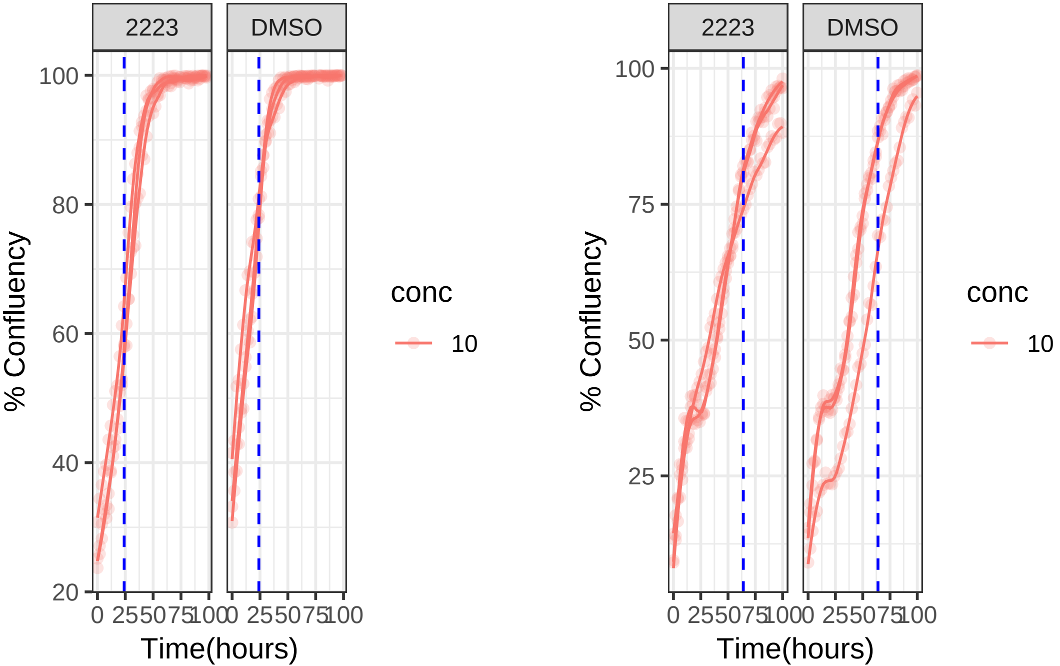

**10 uM 2223 RPE1**

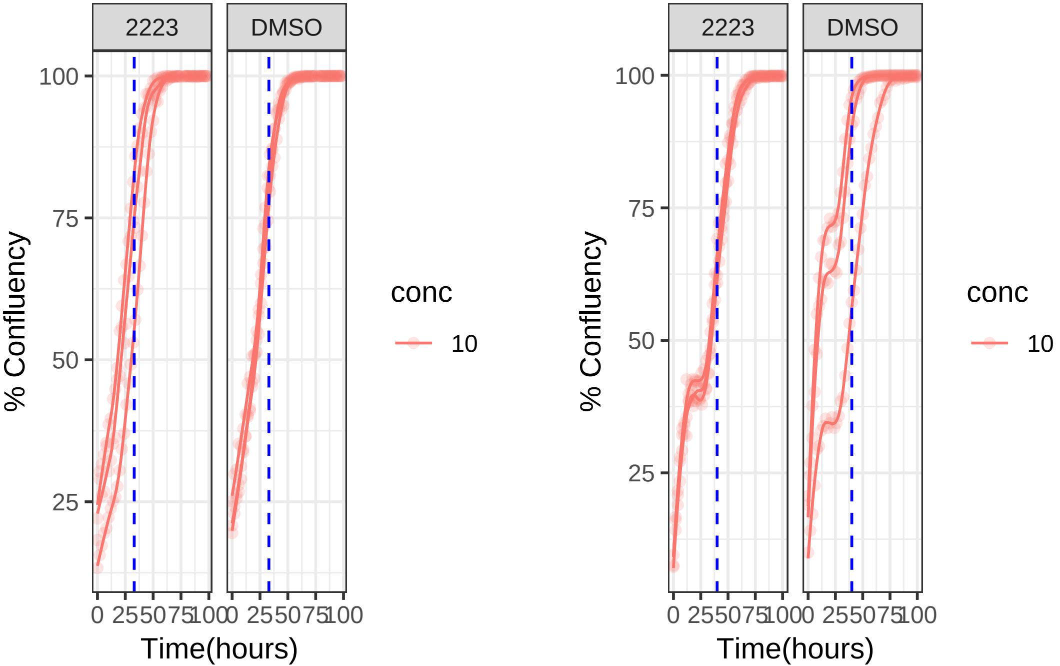

**10 uM 2227 RPE1 Mad2KD**

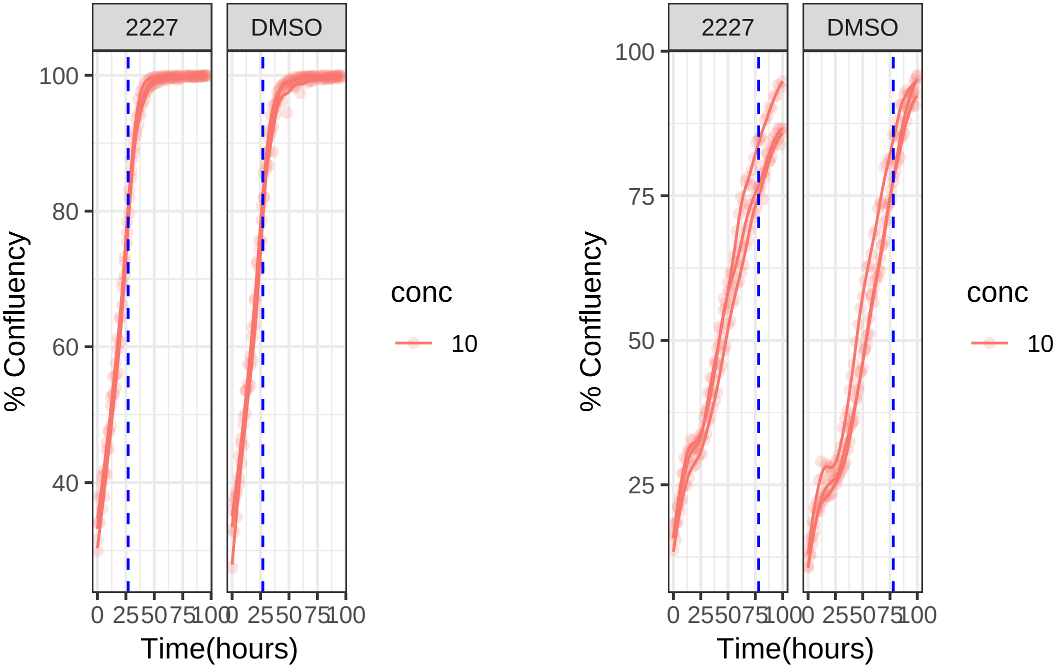

**10 uM 2227 RPE1**

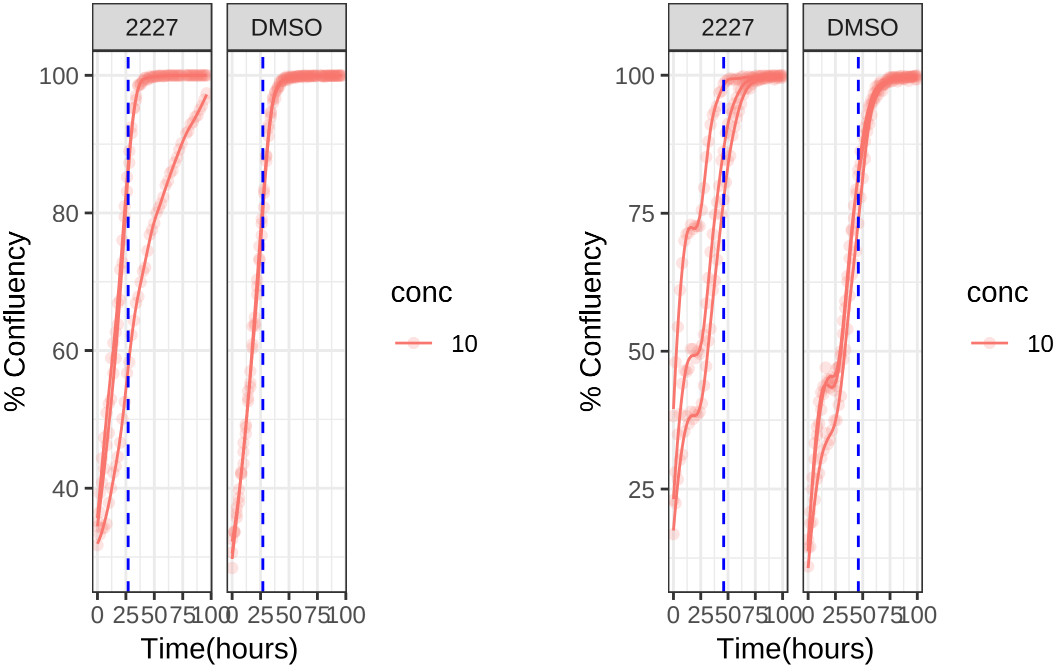

1 uM 2241 RPE1 Mad2KD

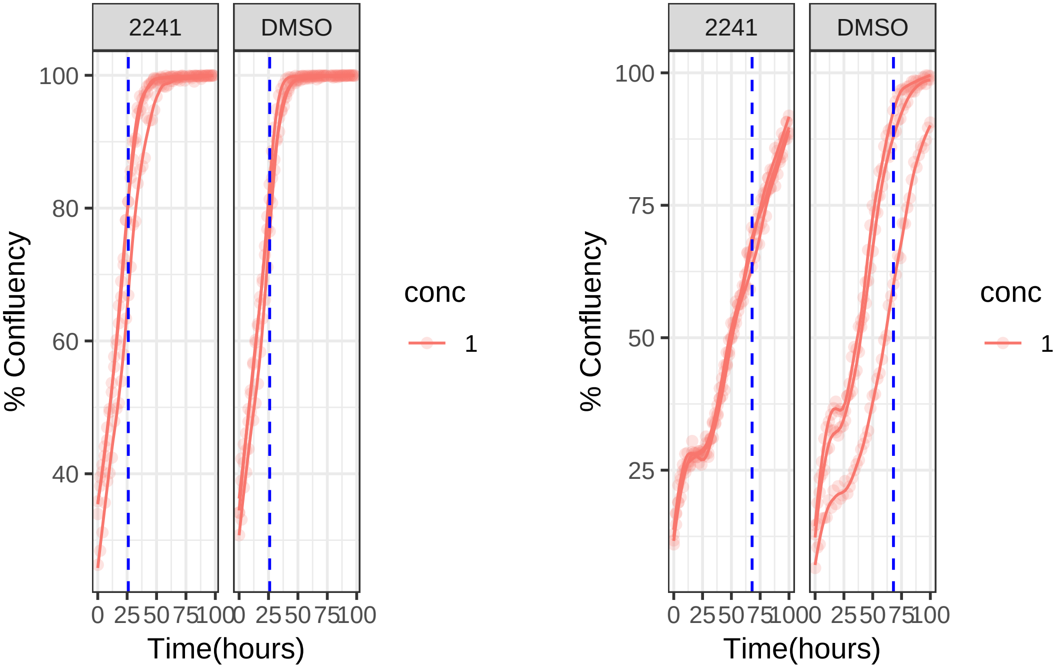

**1 uM 2241 RPE1**

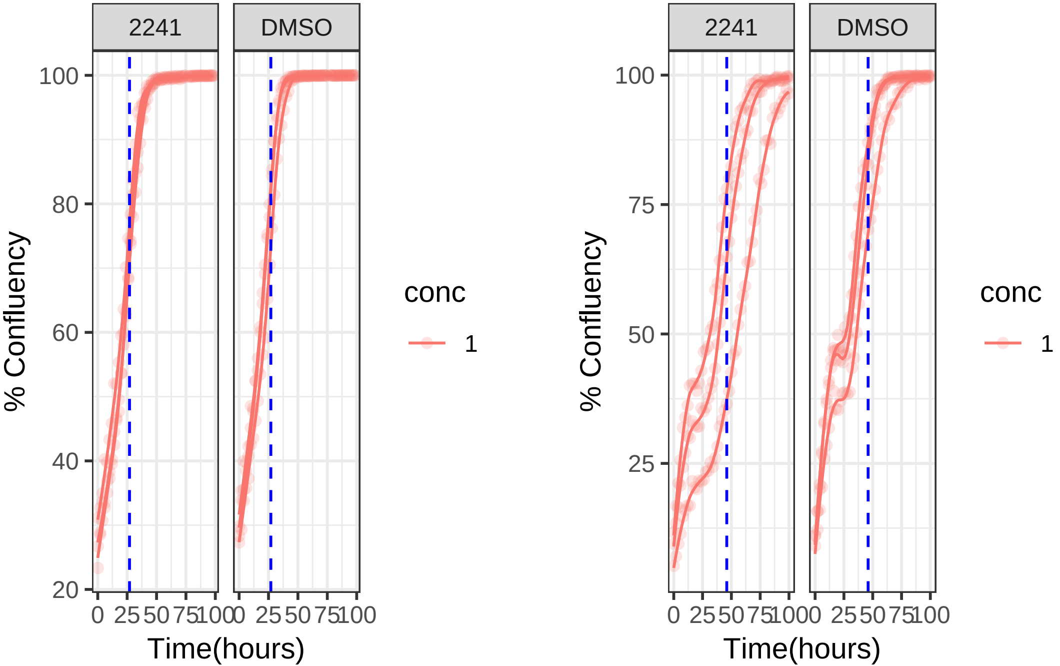

**1 uM 2244 RPE1 Mad2KD**

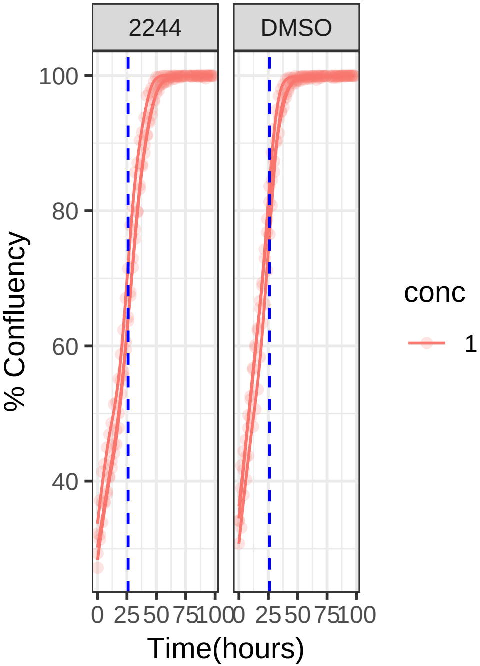


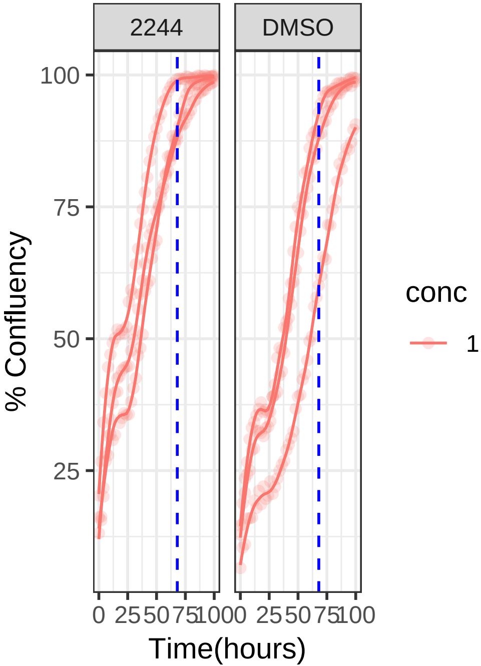

**1 uM 2244 RPE1**

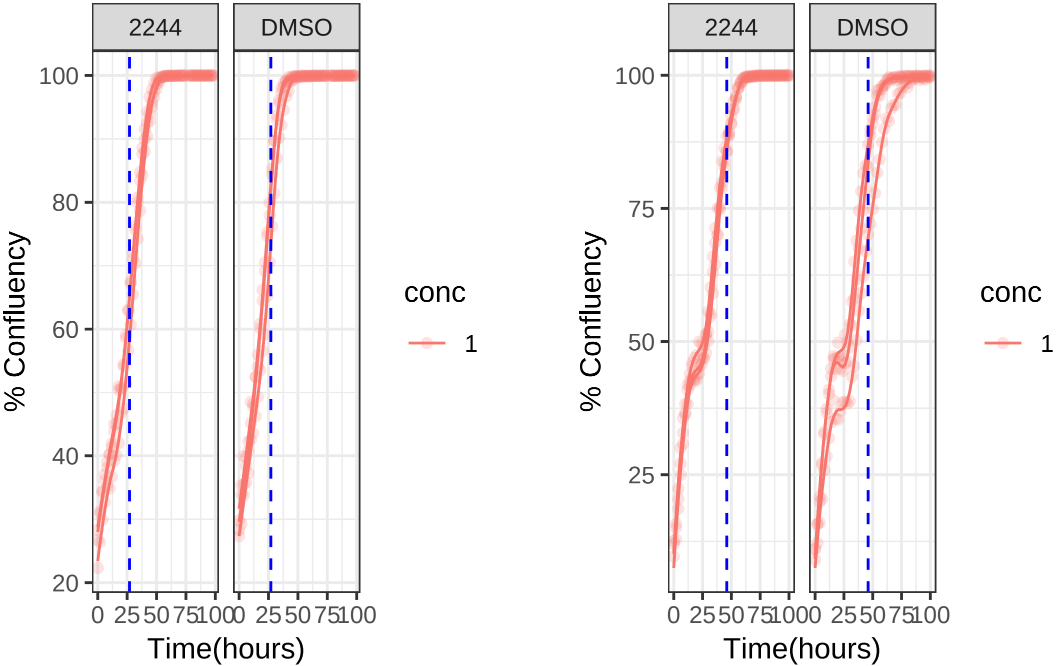

**1 uM 2249 RPE1 Mad2KD**

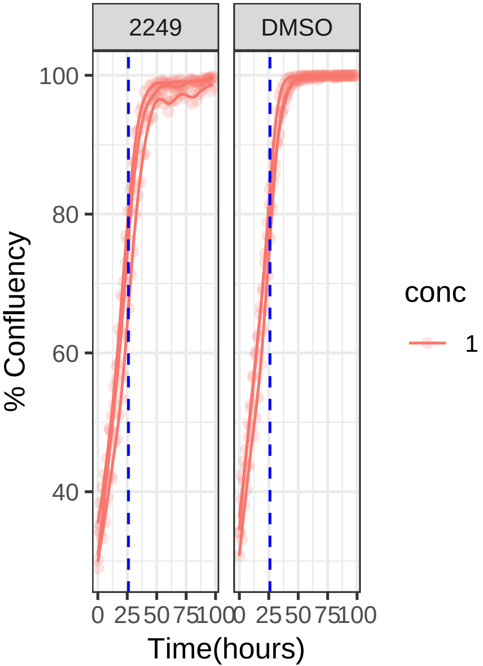


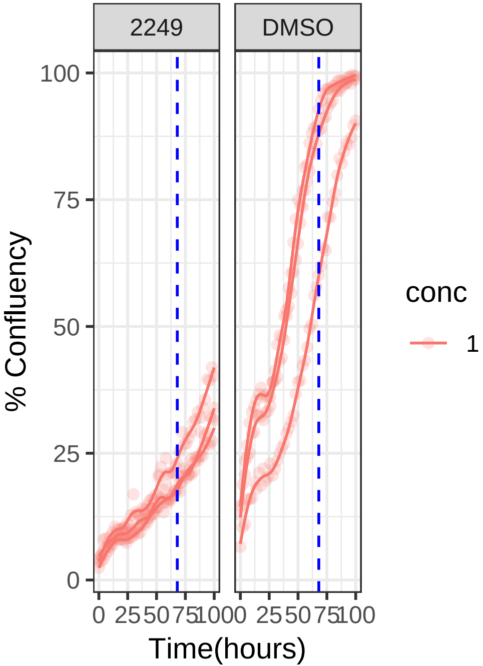

**1 uM 2249 RPE1**

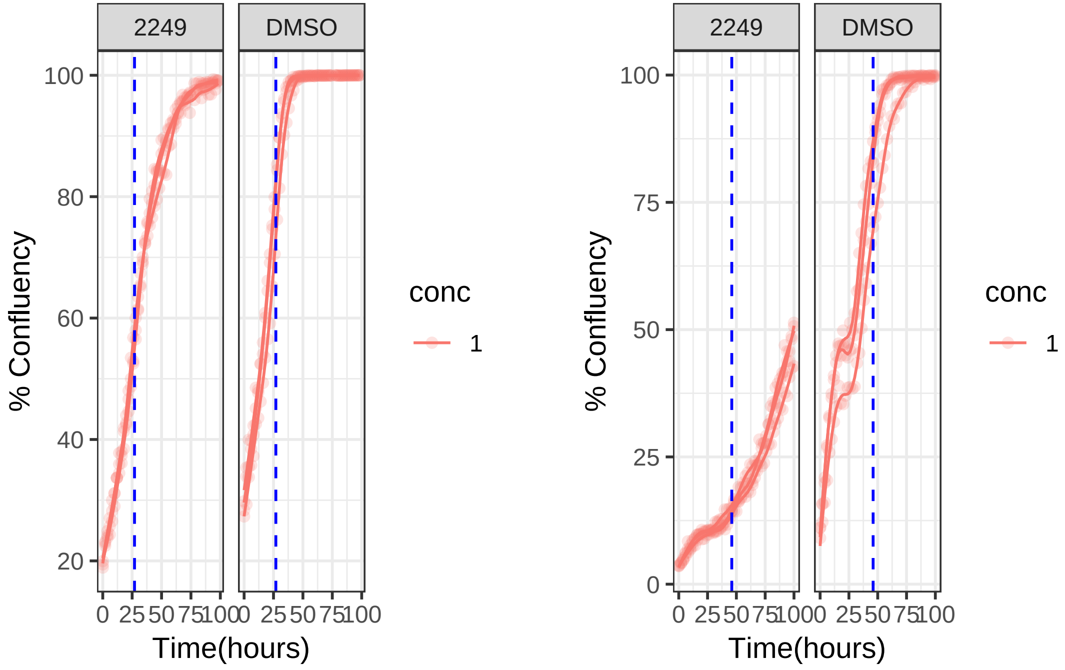

**0.1 uM 2250 RPE1 Mad2KD**

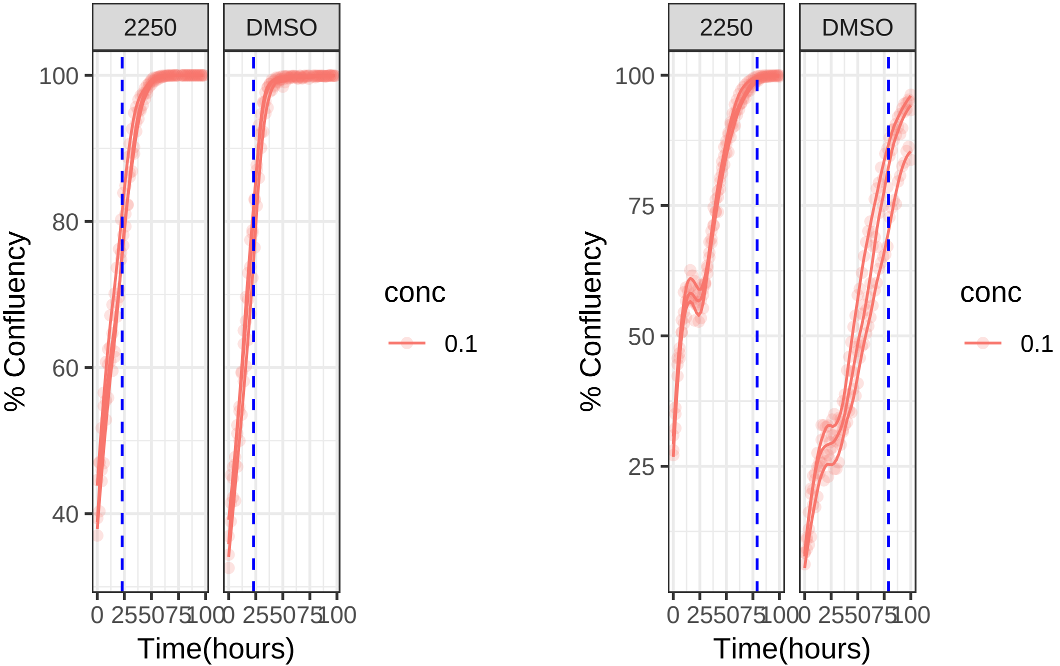

**0.1 uM 2250 RPE1**

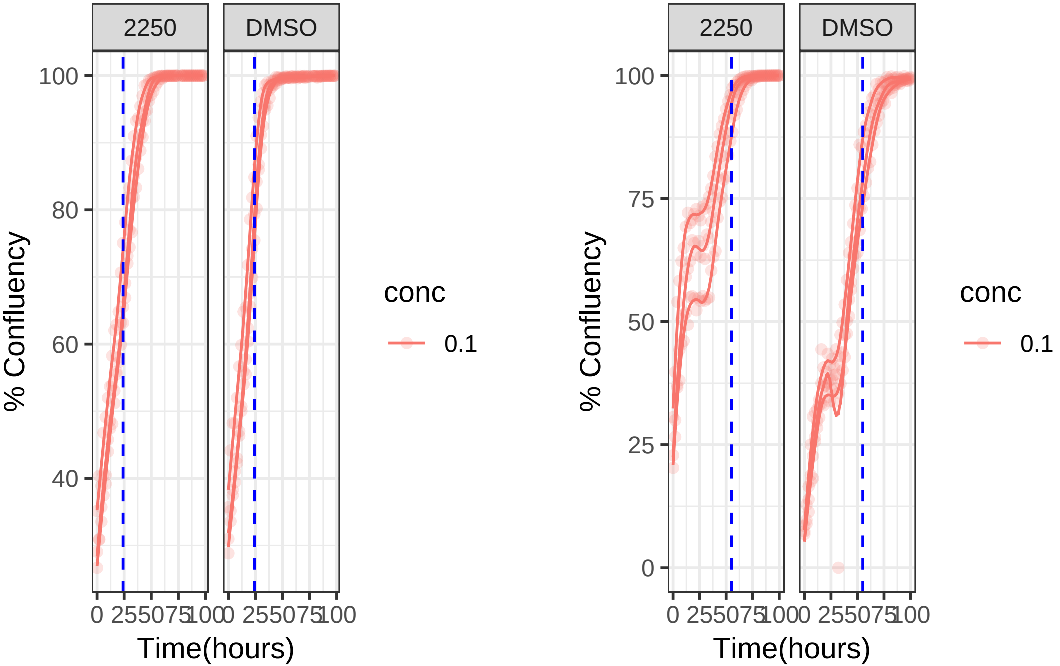

**10 uM 2262 RPE1 Mad2KD**

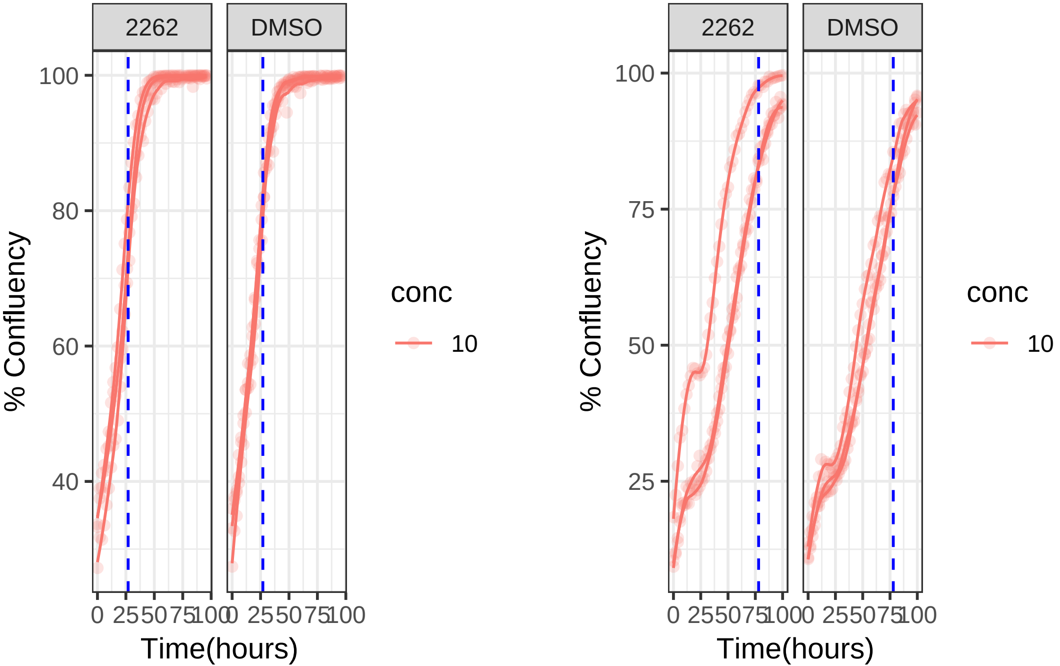

**10 uM 2262 RPE1**

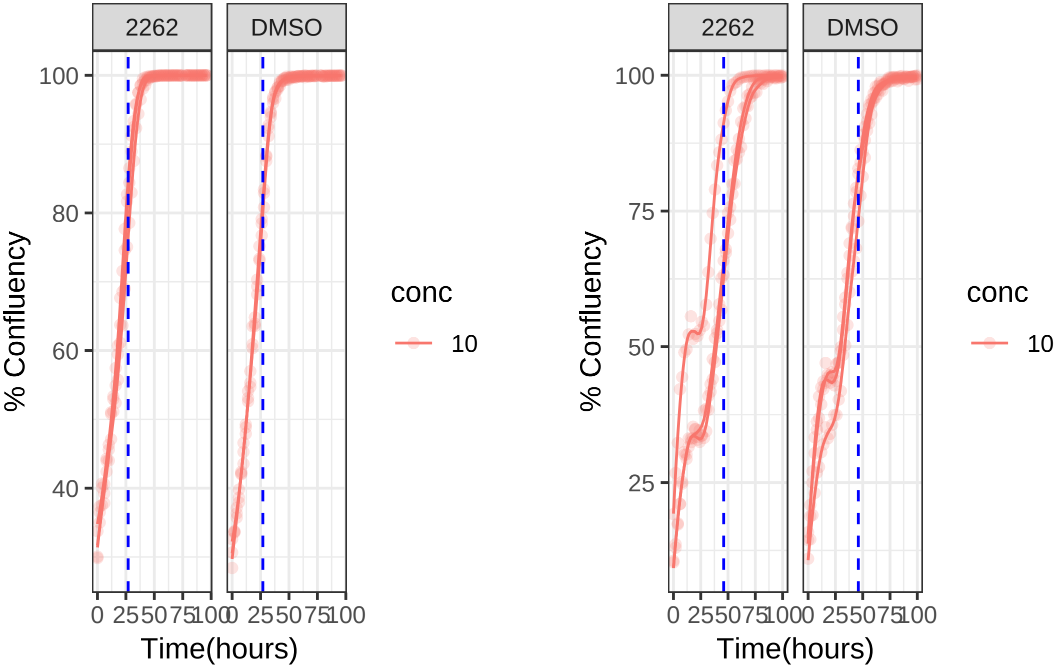

**10 uM 2268 RPE1 Mad2KD**

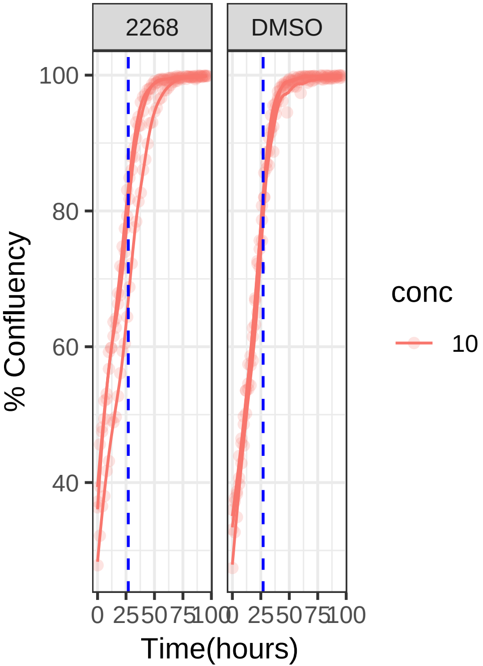


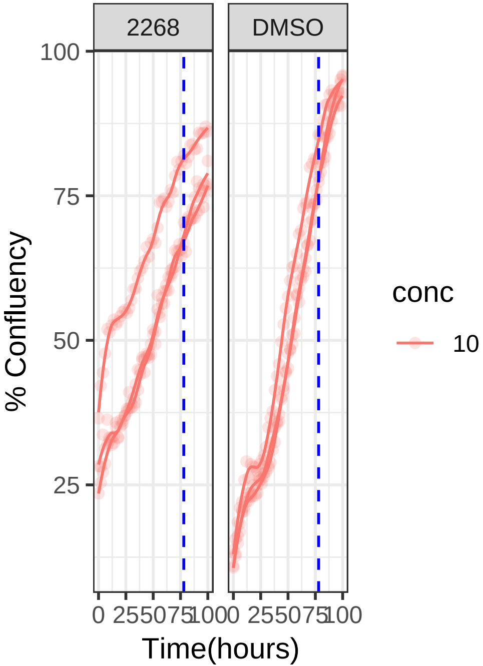

**10 uM 2268 RPE1**

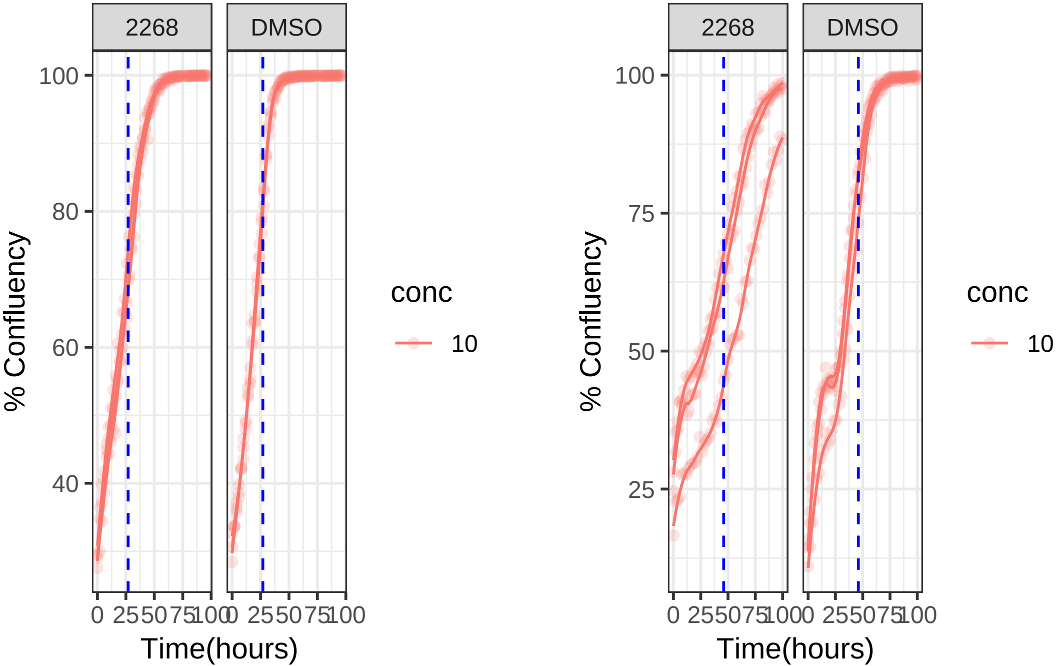

**1 uM 2300 RPE1 Mad2KD**

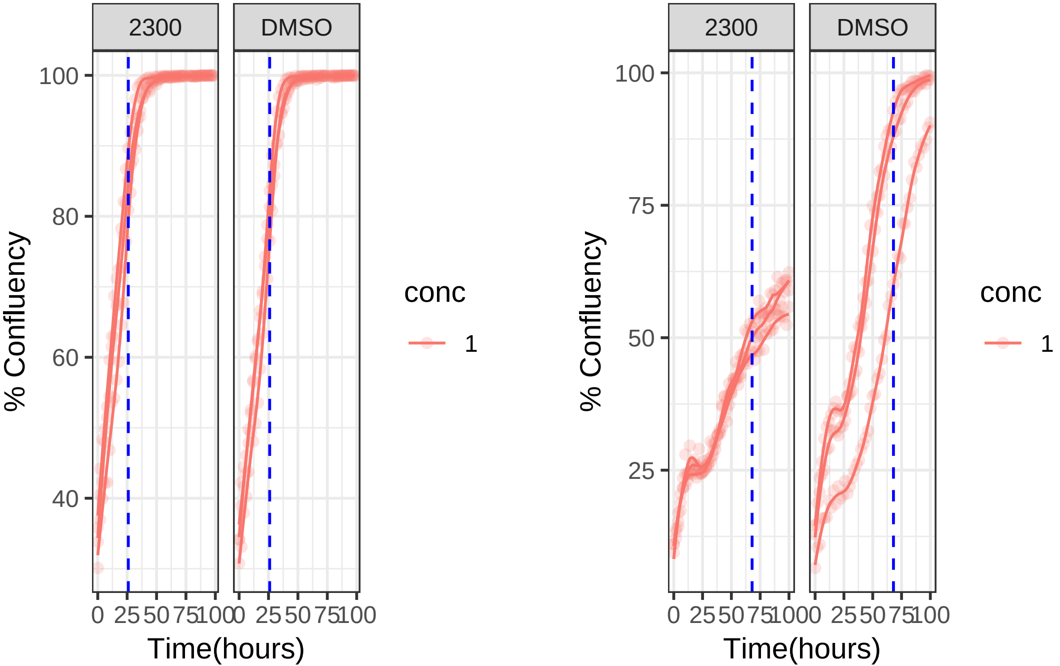

#### 1 uM 2300 RPE1

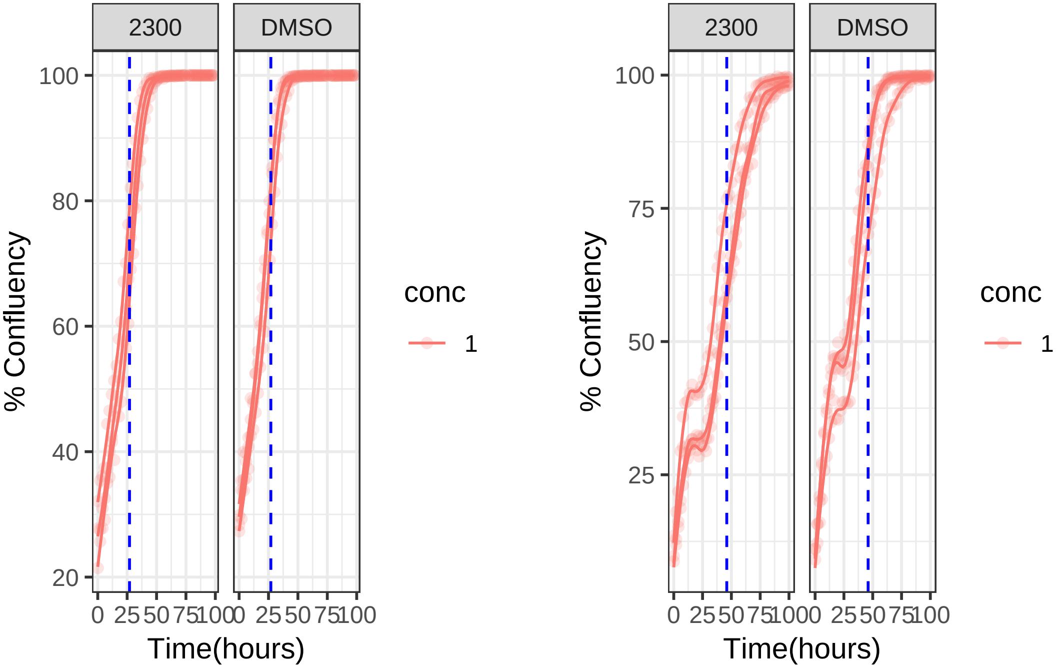

**1 uM 2323 RPE1 Mad2KD**

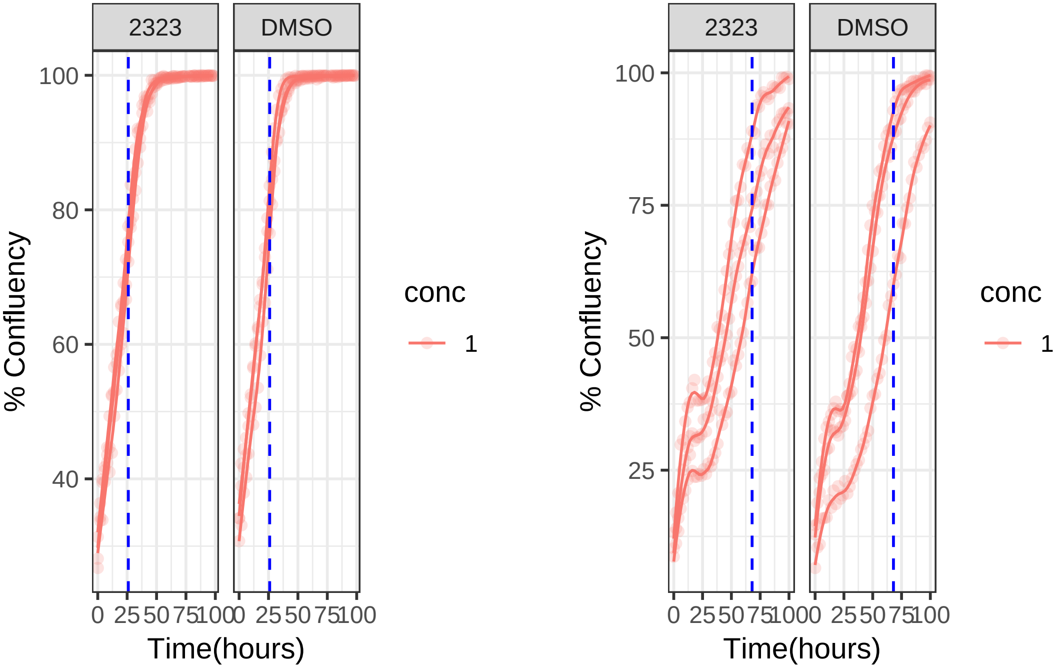

**1 uM 2323 RPE1**

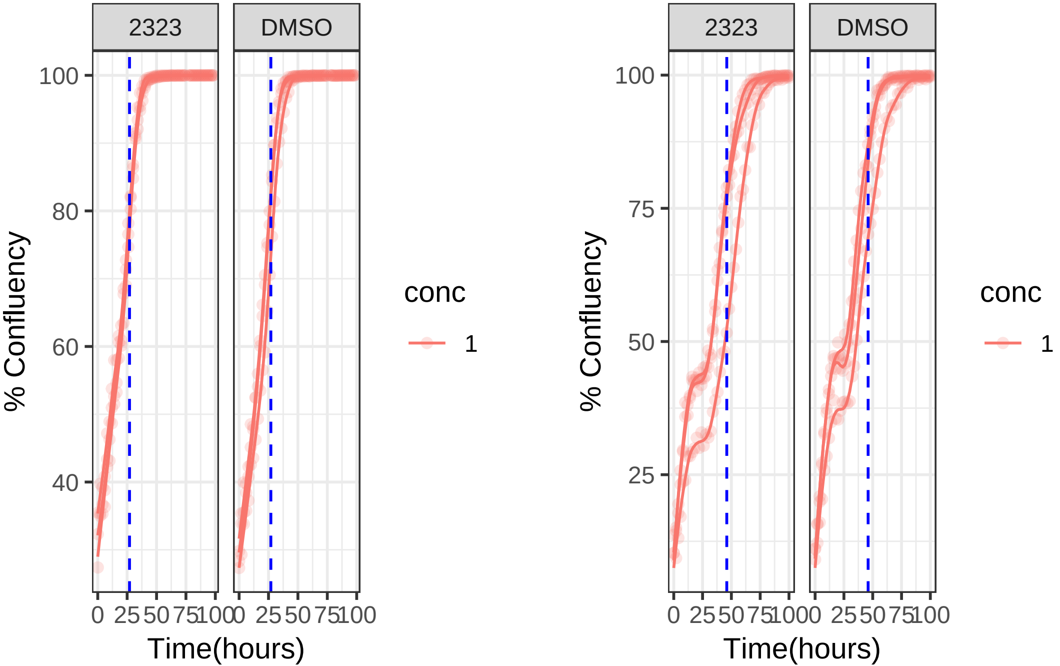

**1 uM 2347 RPE1 Mad2KD**

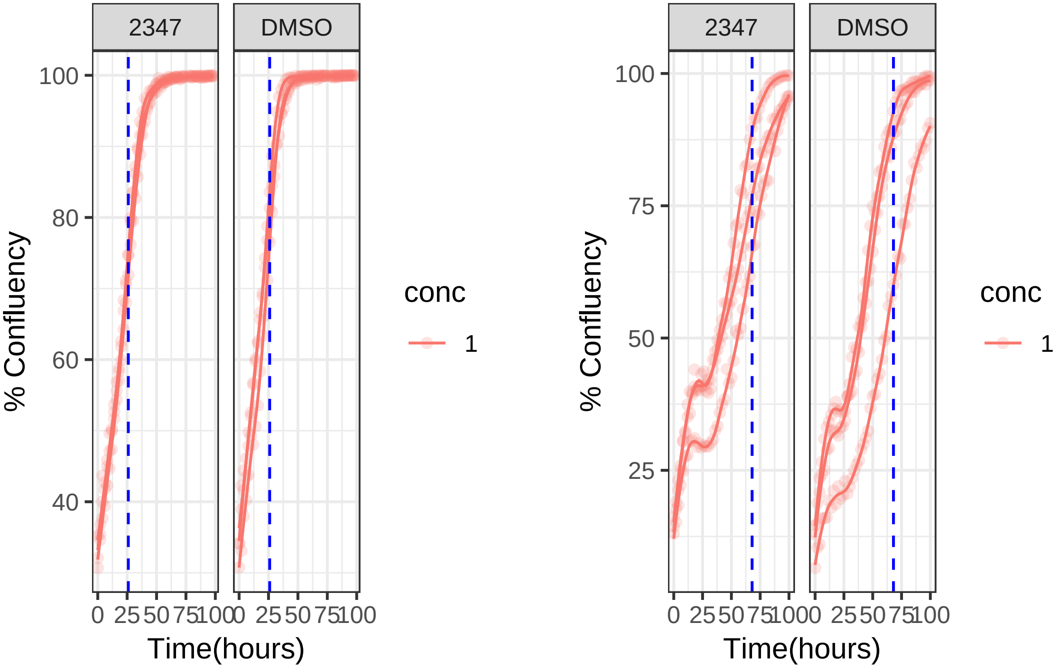

**1 uM 2347 RPE1**

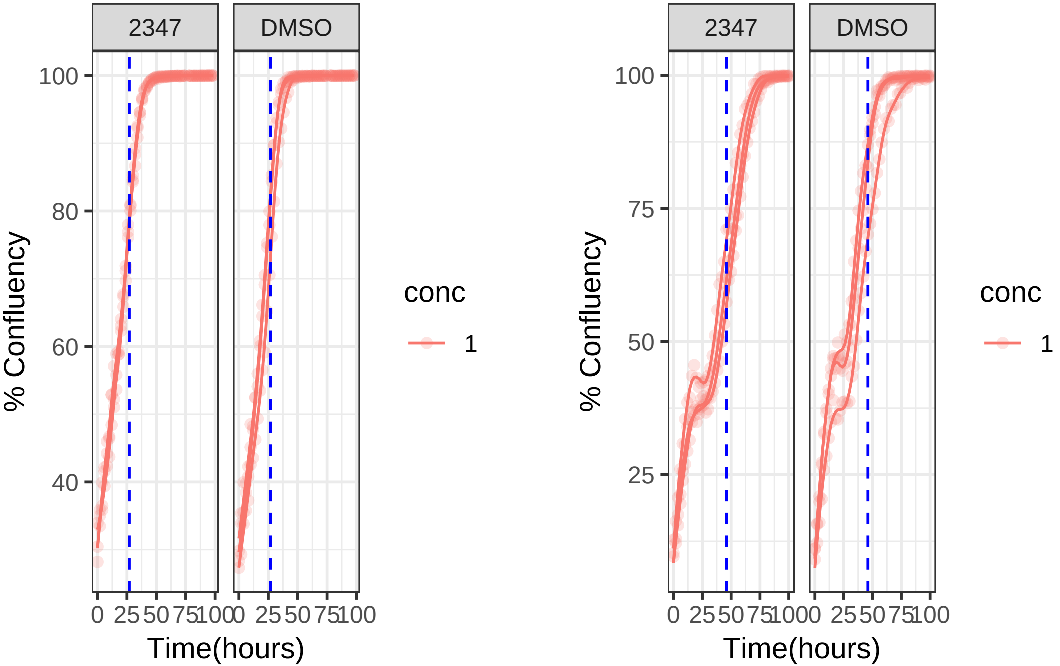

**10 uM 2358 RPE1 Mad2KD**

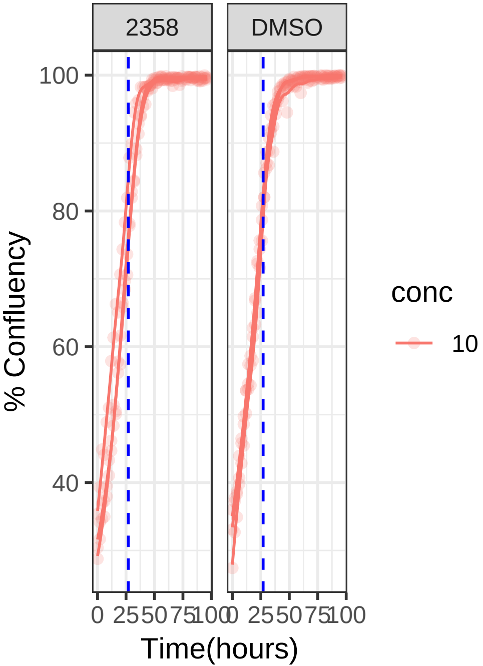


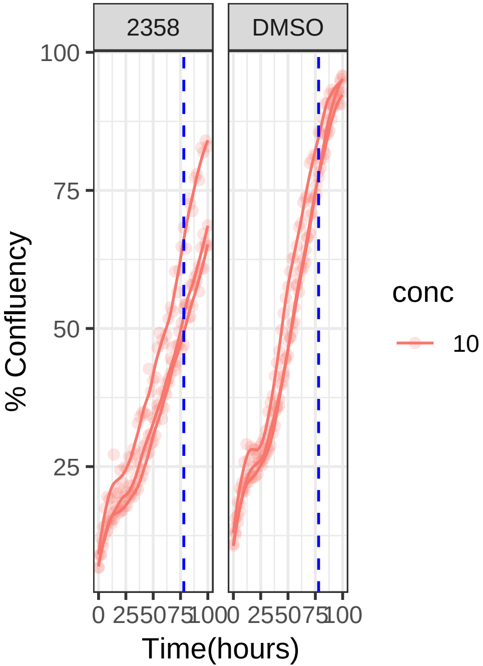

**10 uM 2358 RPE1**

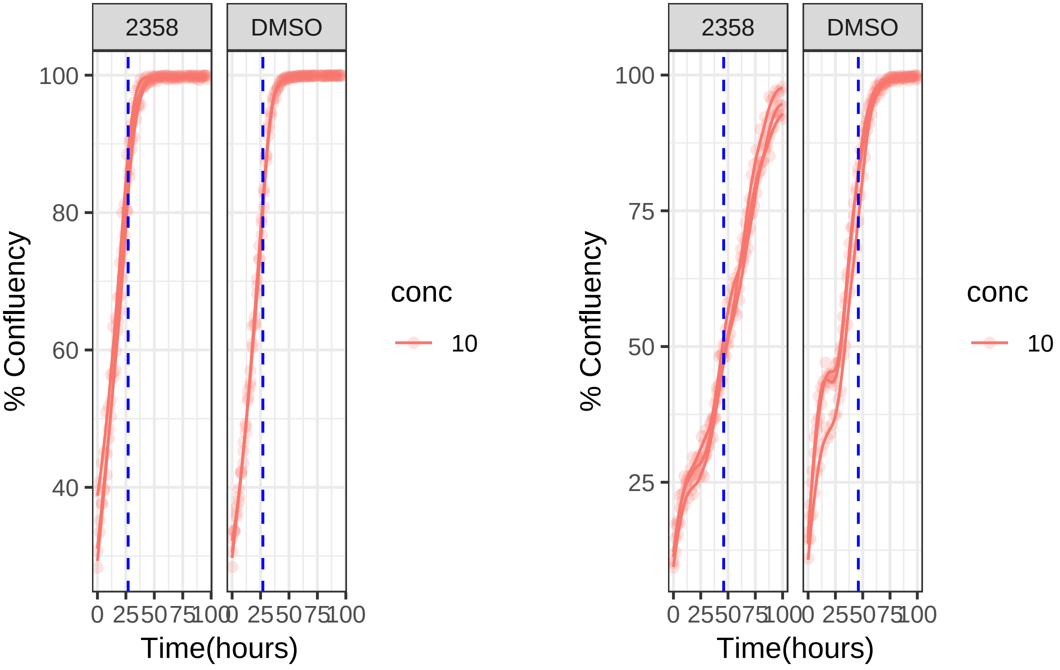

**10 uM 2369 RPE1 Mad2KD**

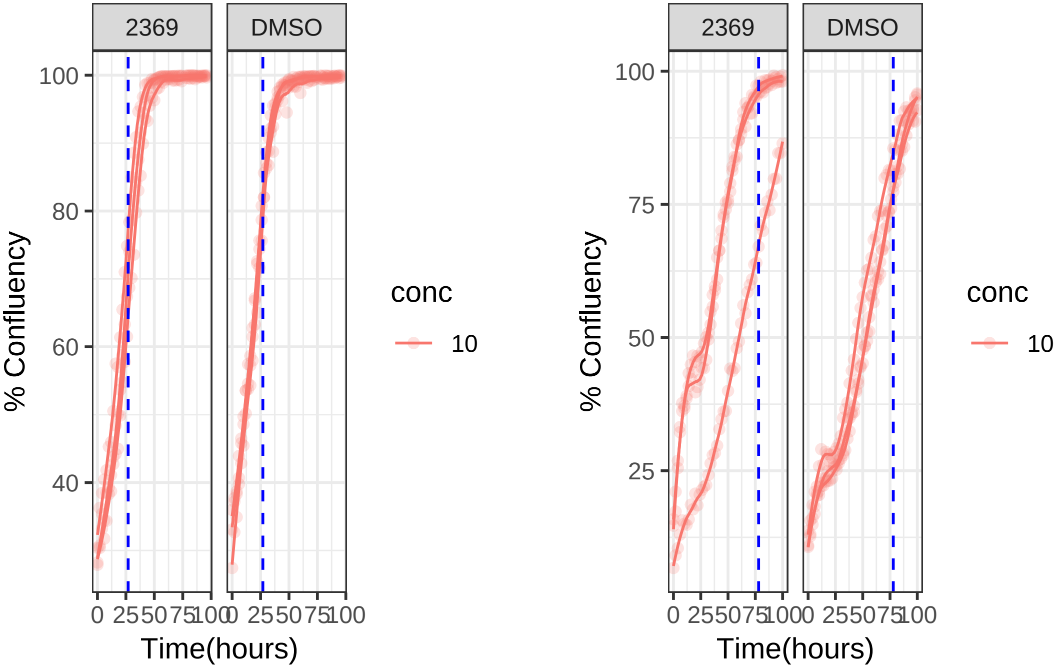

**10 uM 2369 RPE1**

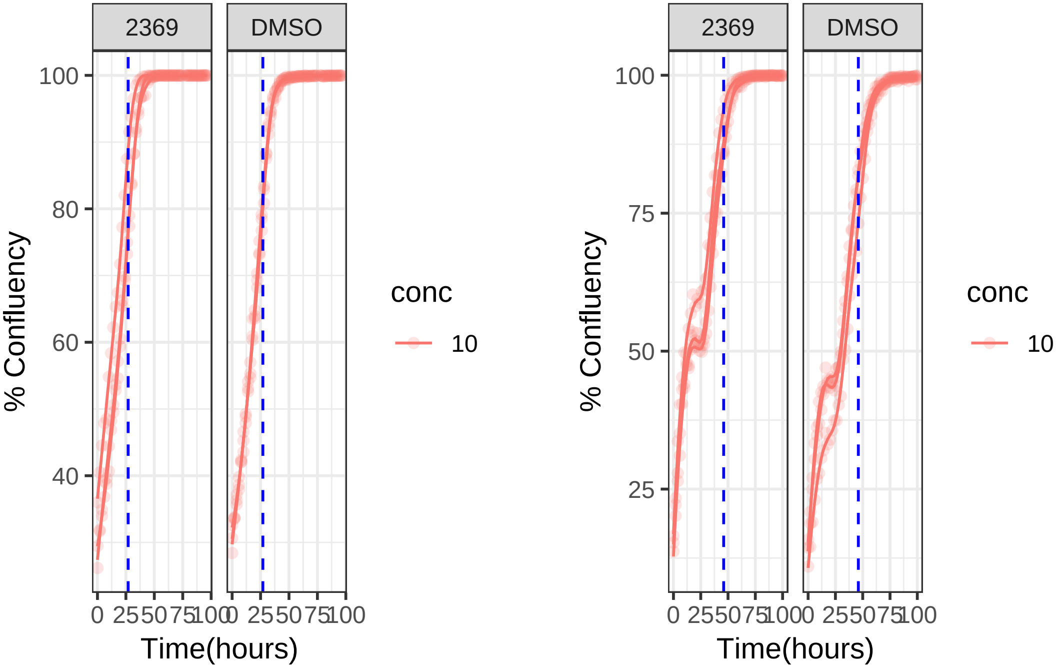

**10 uM 2375 RPE1 Mad2KD**

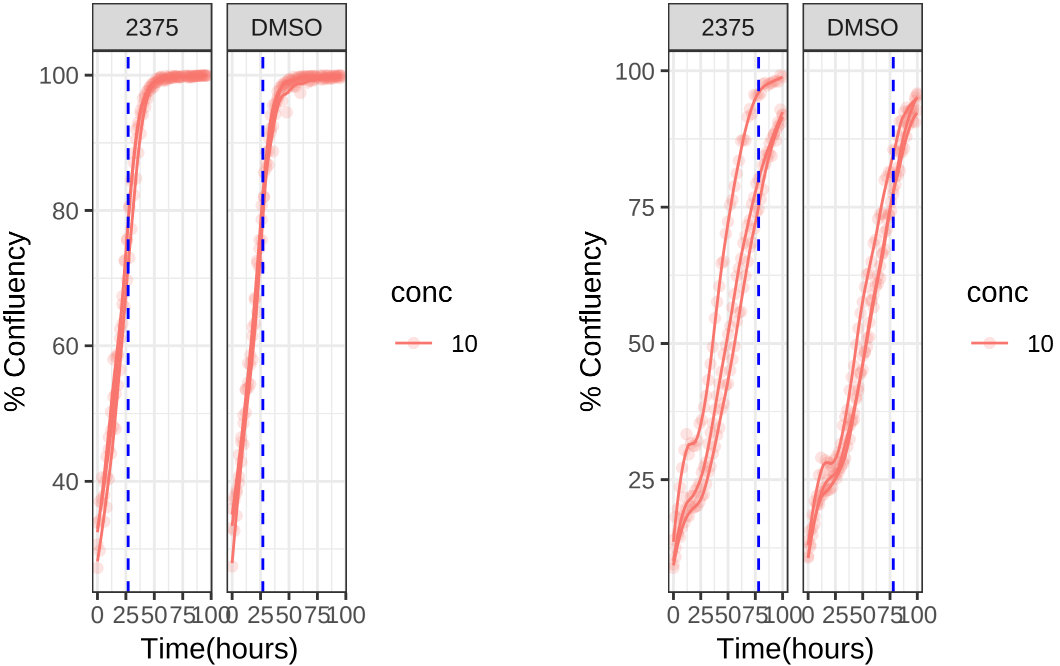

**10 uM 2375 RPE1**

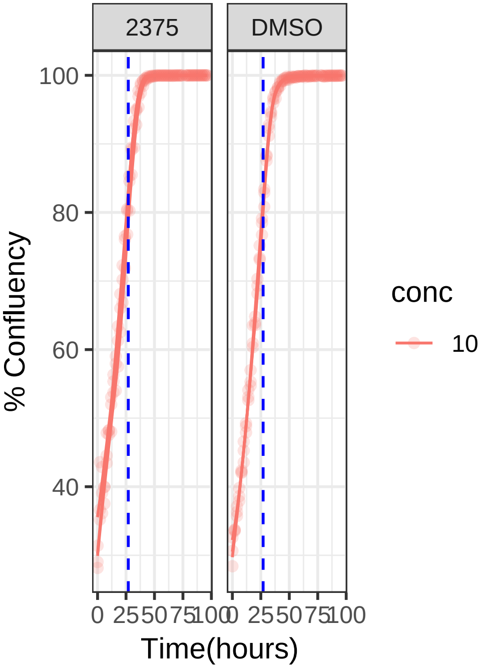


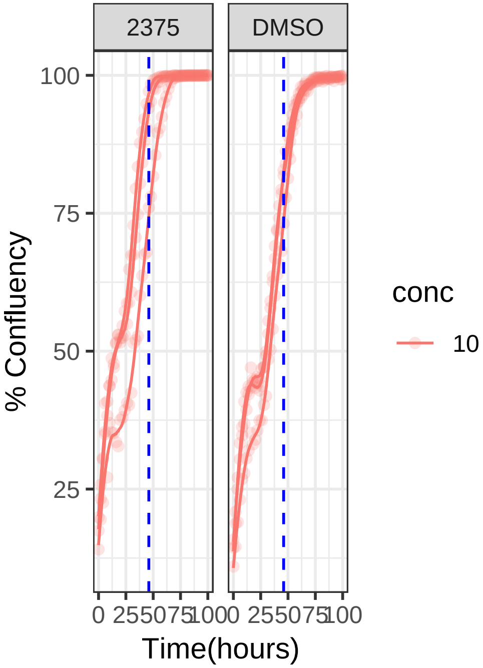

**10 uM 2402 RPE1 Mad2KD**

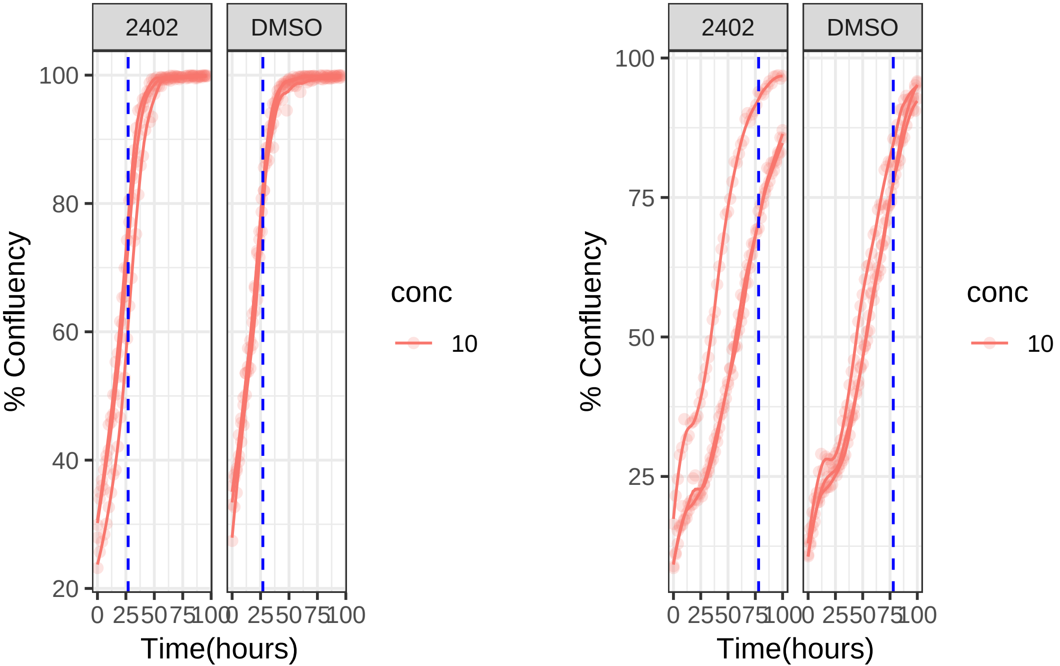

**10 uM 2402 RPE1**

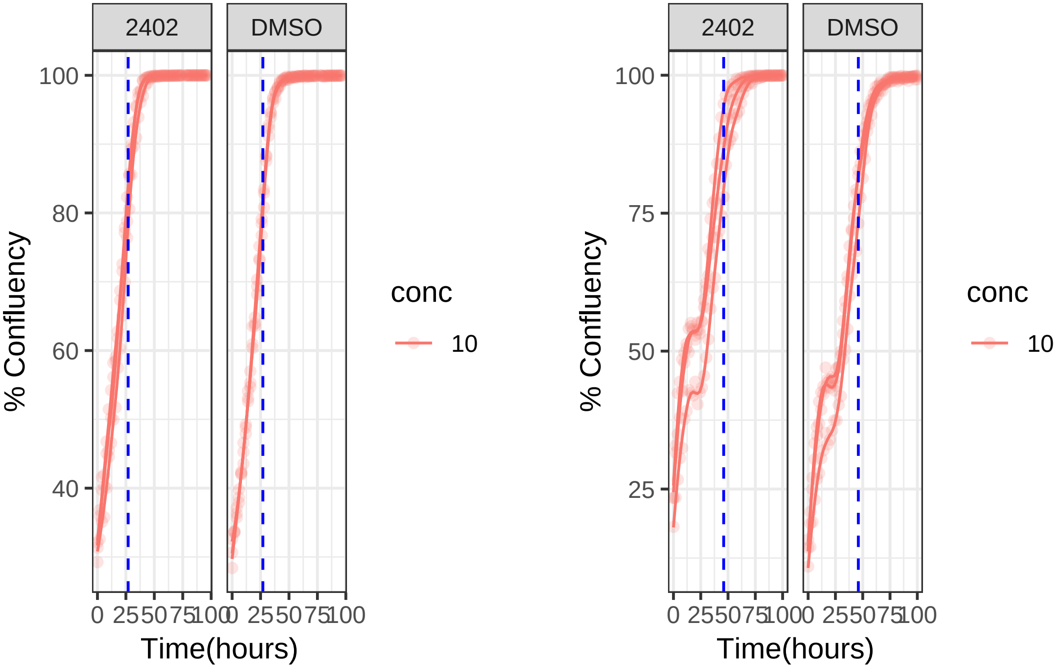

**1 uM 2831 RPE1 Mad2KD**

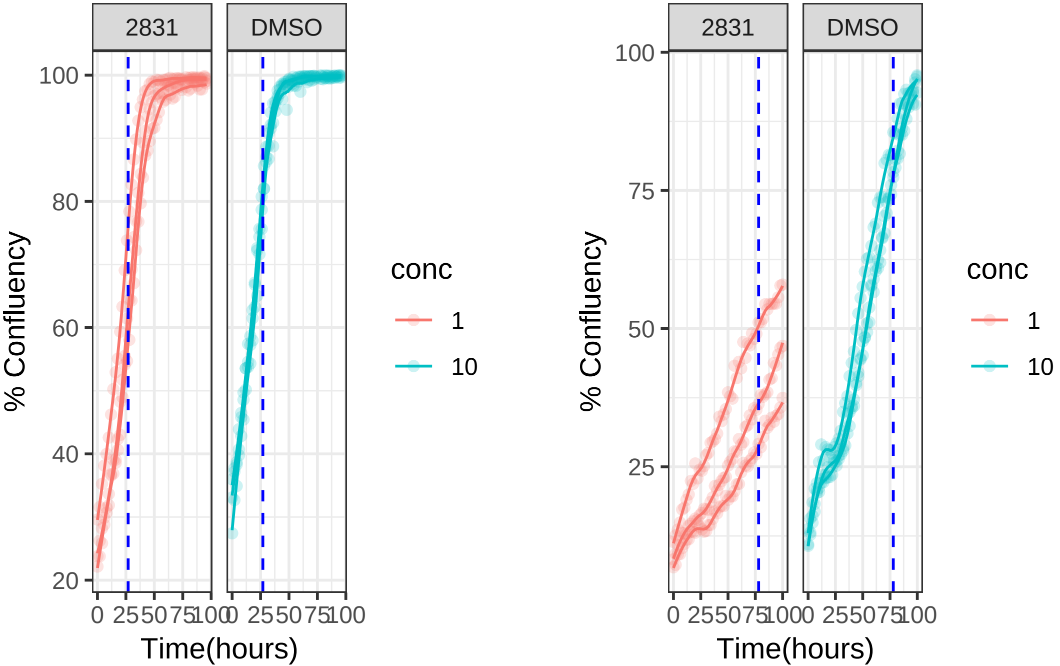

**10 uM 2831 RPE1 Mad2KD**

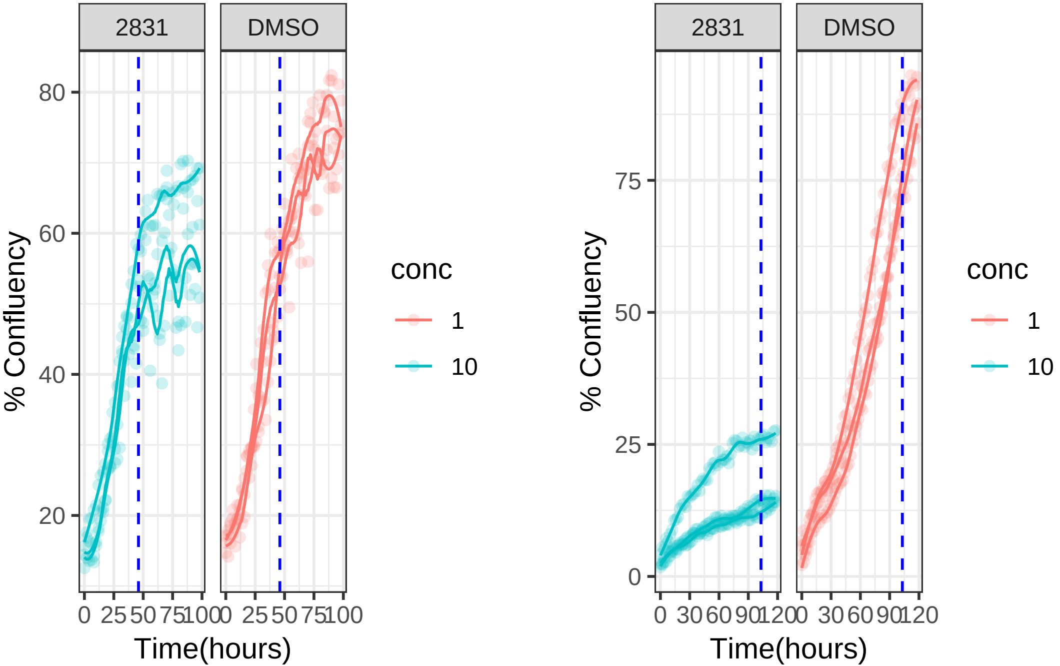

**1 uM 2831 RPE1**

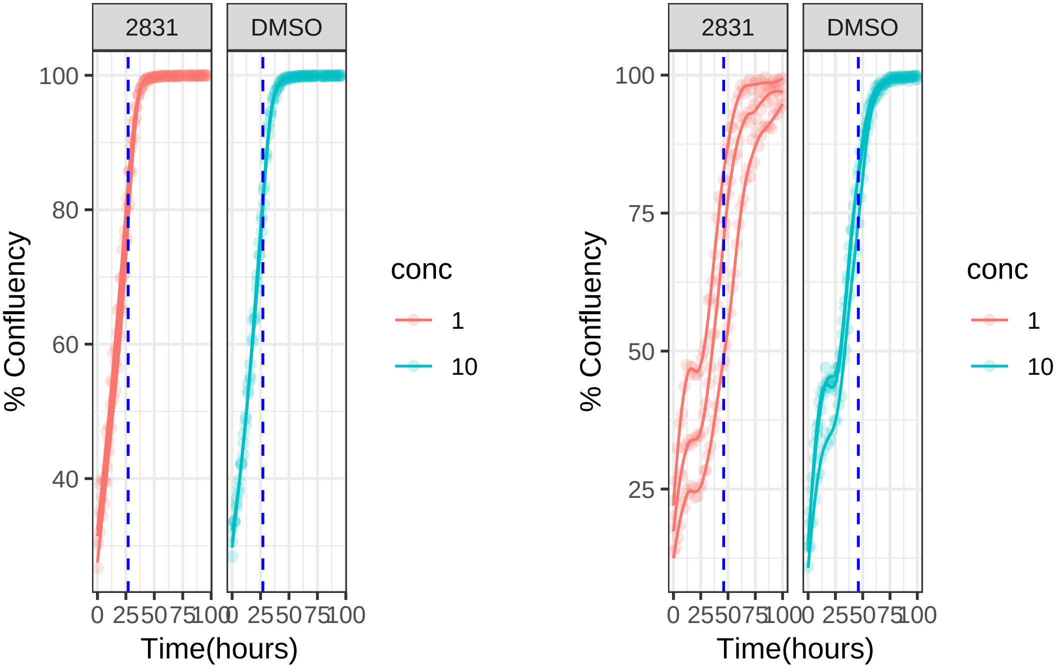

**10 uM 2831 RPE1**

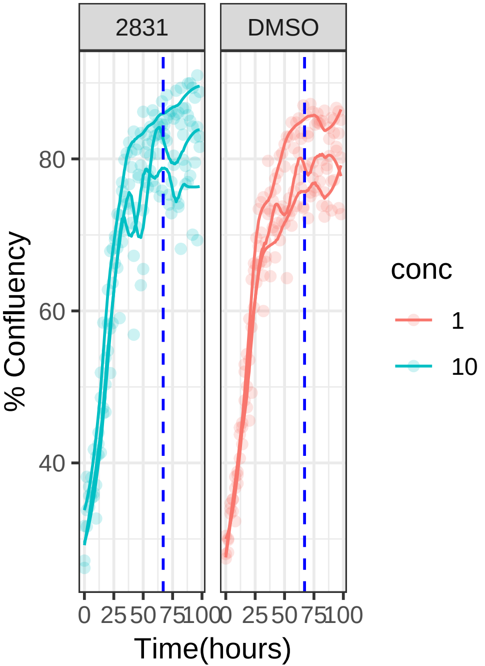


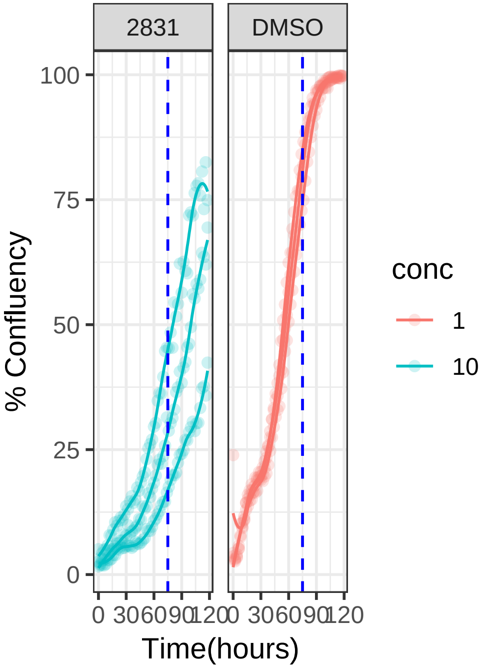

**0.1 uM 2nd.control RPE1 Mad2KD**

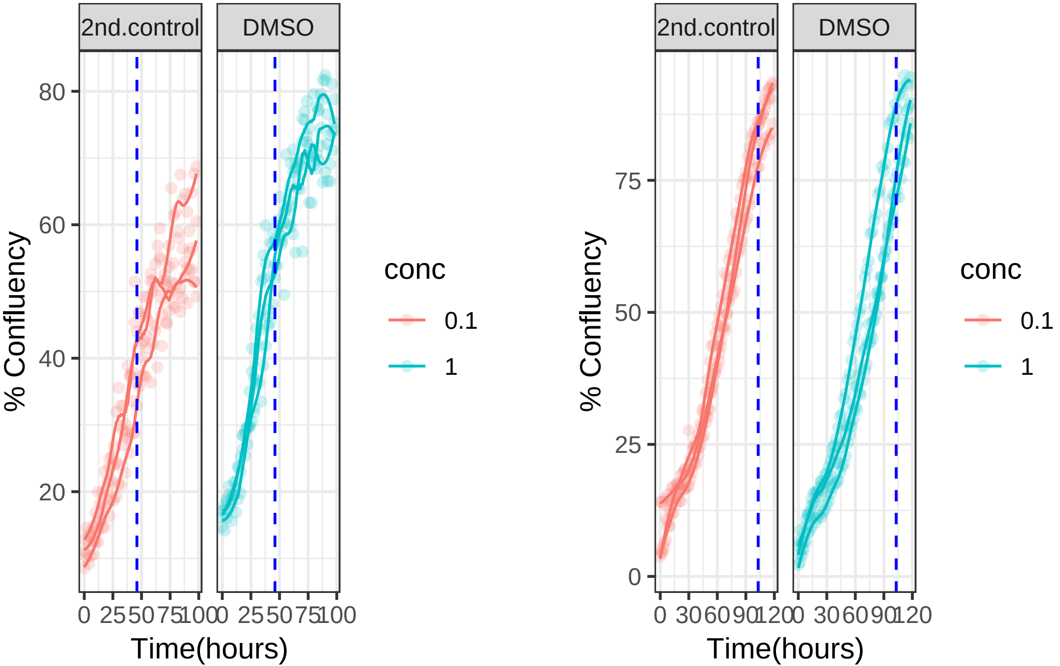

**0.1 uM 2nd.control RPE1**

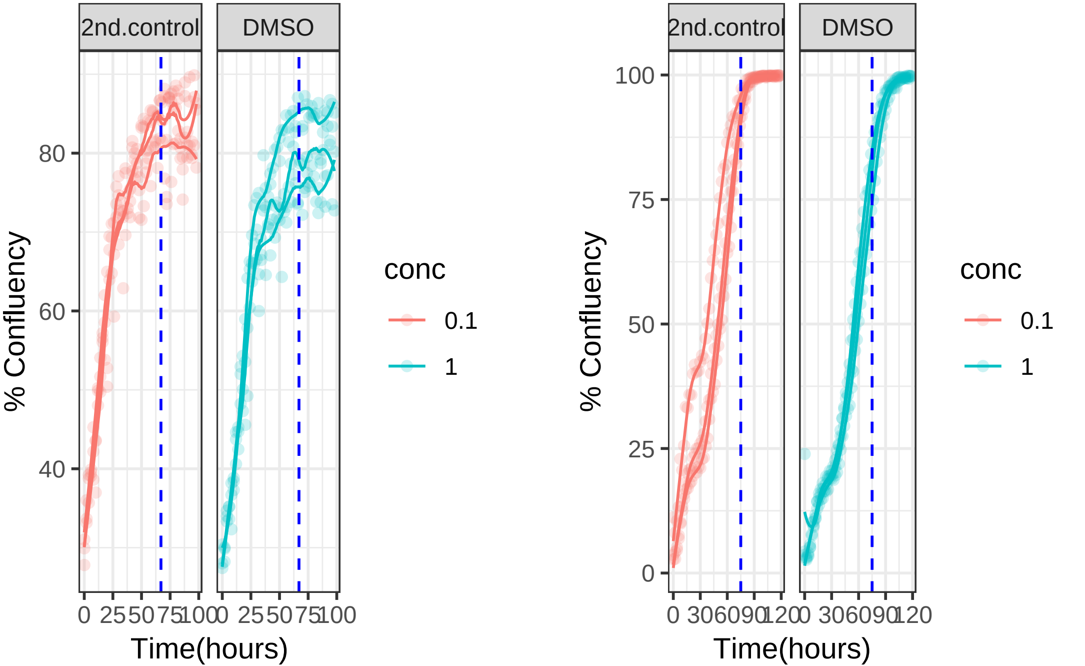
