## Supplementary material for "Altering microtubule dynamics is synergistically toxic with inhibition of the spindle checkpoint": Sup data 3

### **Supplementary data 3:** Drug screen concentration curves

This PDF file contains all growth curves for all 95 drugs used in the aneuploid and CIN screen. One page per drug. Each page contains one or more growth curves comparing RPE1 cells with DMSO and the drug. All drug concentrations start at 10uM. Drugs were screened until they were no longer toxic to RPE1 cells at that concentration, with 1:10 dilution intervals. All concentrations in micromolar (uM). All curves are triplicates.

Drug ID is listed top left corner.

Or

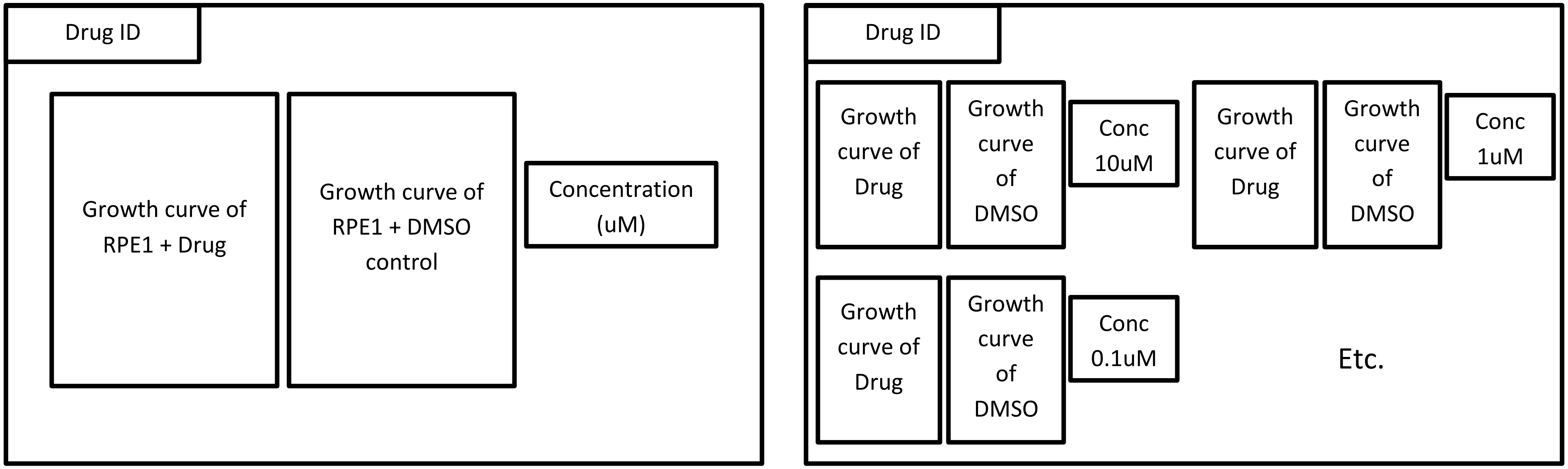

**1122**

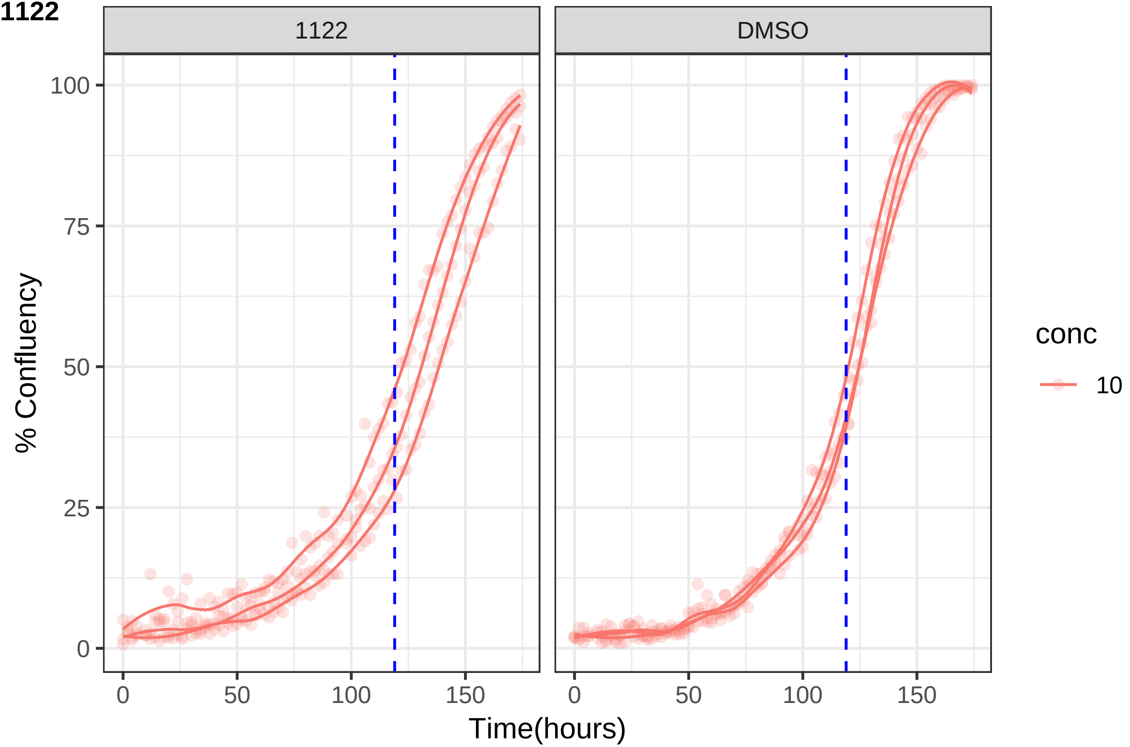

1122
DMSO
100
75
50
25
0
% Confluency
0
50
100
150
Time(hours)
conc
10

**1134**

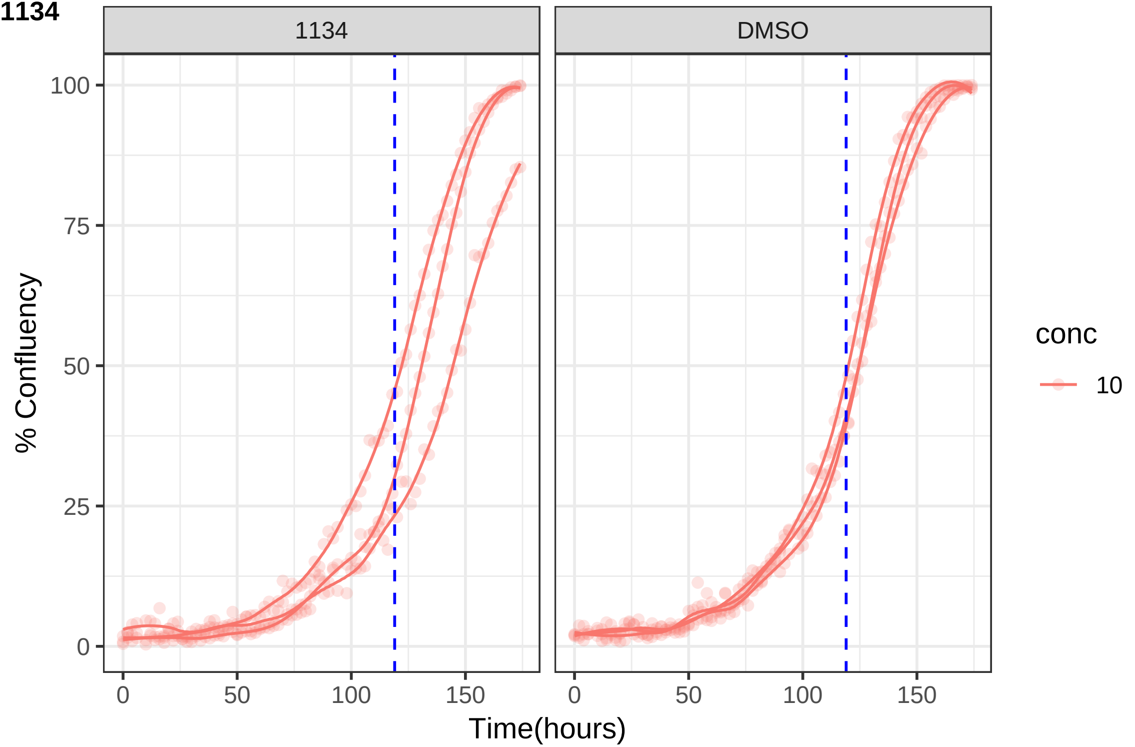

1134
DMSO
% Confluency
100
75
50
25
0
0
50
100
150
Time(hours)
conc
10

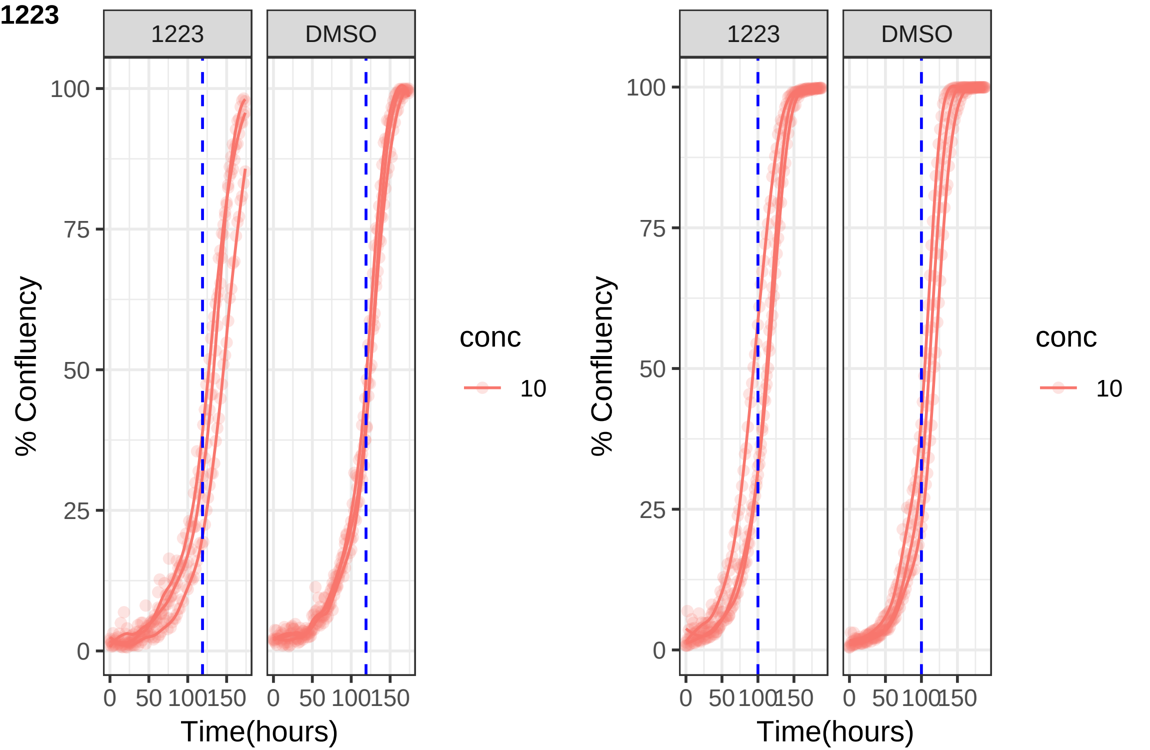

1223
1223
DMSO
% Confluency
100
75
50
25
0
0
50
100
150
Time(hours)
conc
10
1223
DMSO
% Confluency
Time(hours)
conc
10

**1233**

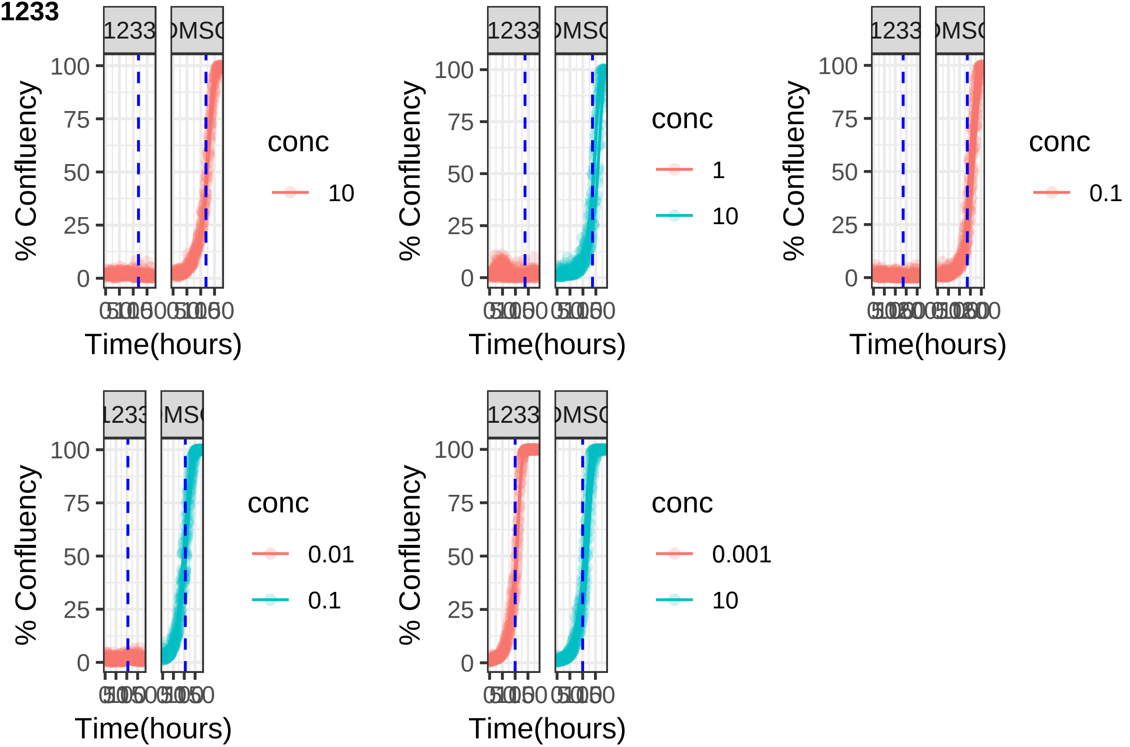

**1237**

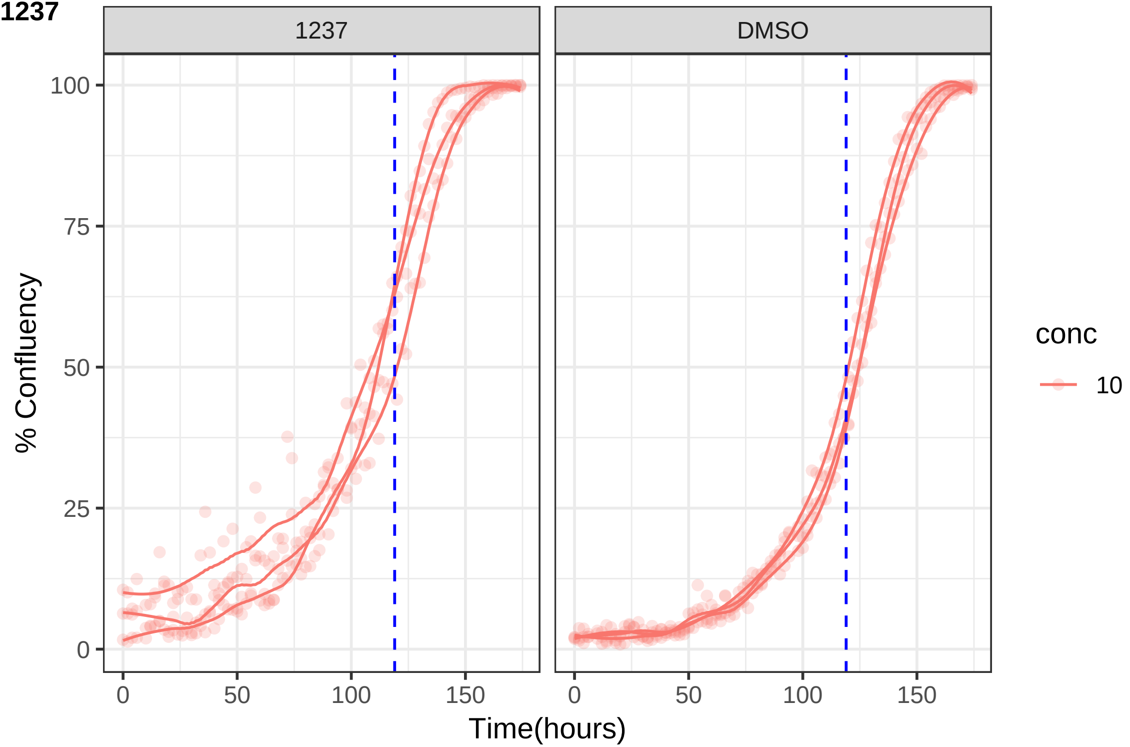
1237
DMSO
% Confluency
100
75
50
25
0
0
50
100
150
Time(hours)
conc
10

**1239**

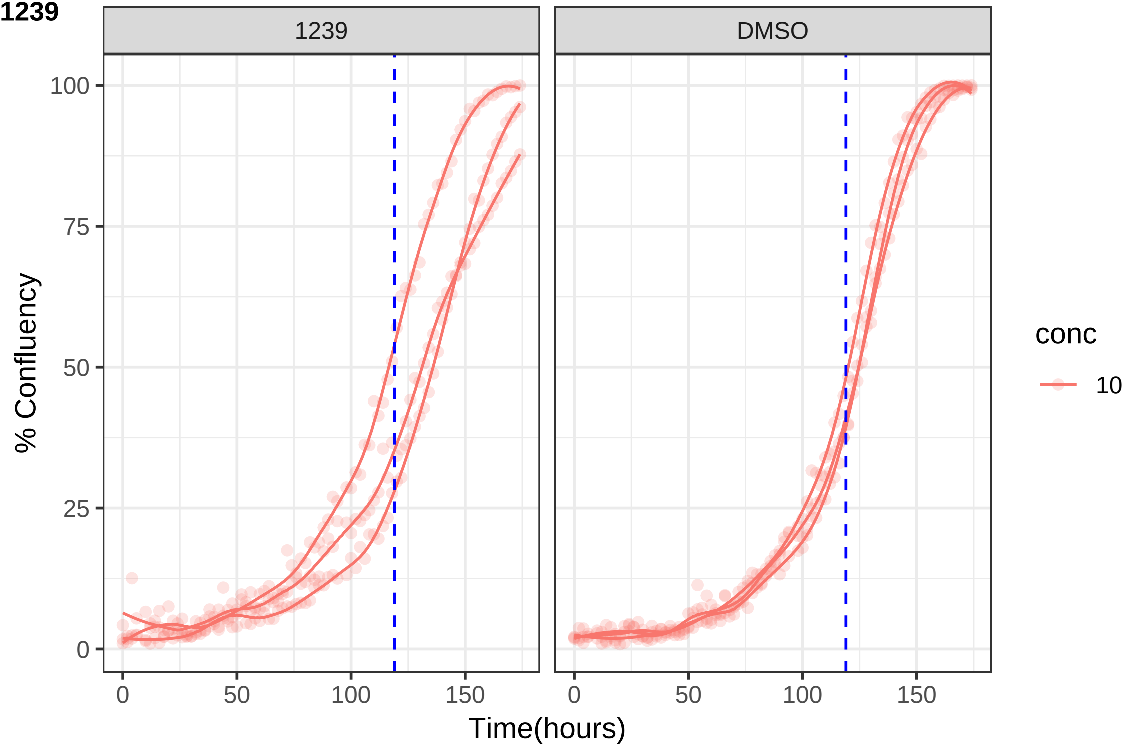

1239
DMSO
100
75
50
25
0
% Confluency
0
50
100
150
Time(hours)
conc
10

**1243**

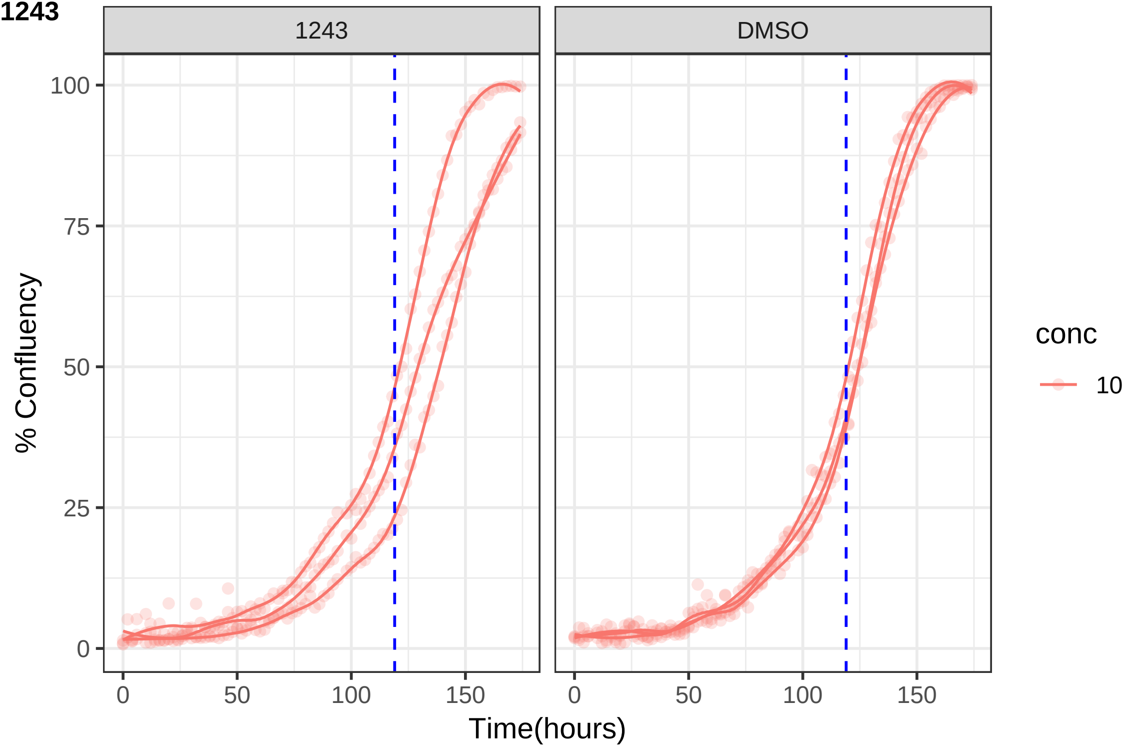

1243
DMSO
100
75
50
25
0
% Confluency
0
50
100
150
Time(hours)
conc
10

**1254**

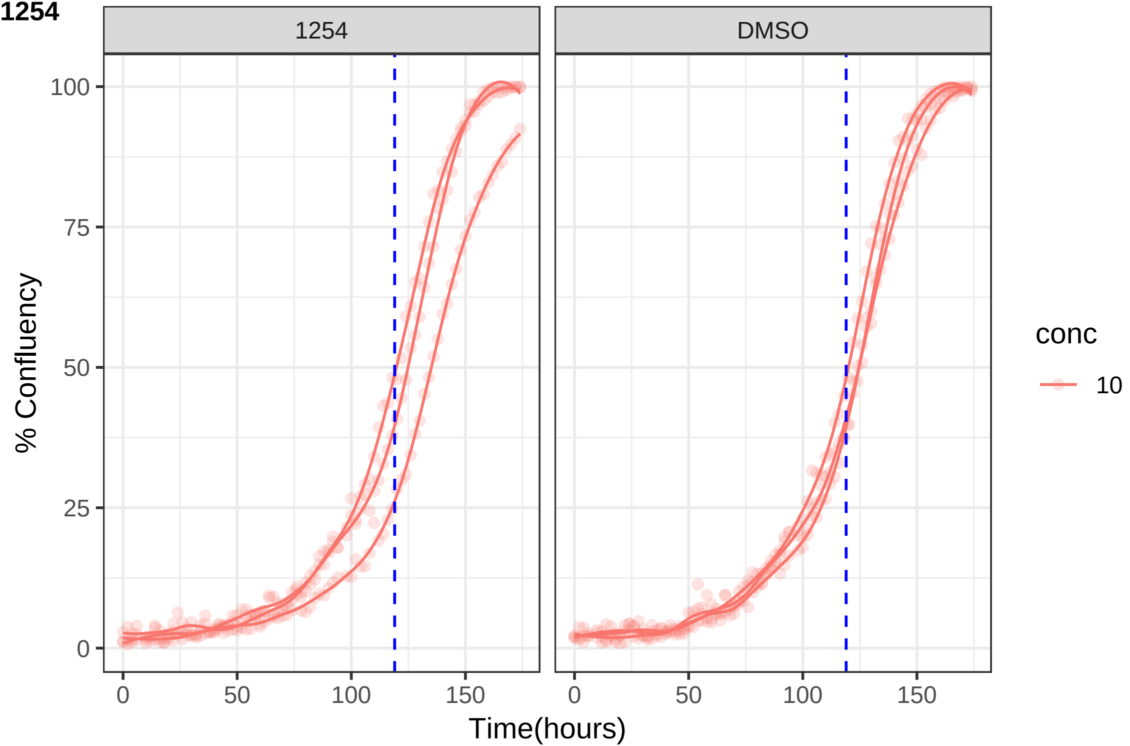

1254
DMSO
% Confluency
0
25
50
75
100
0
50
100
150
Time(hours)
conc
10

**1268**

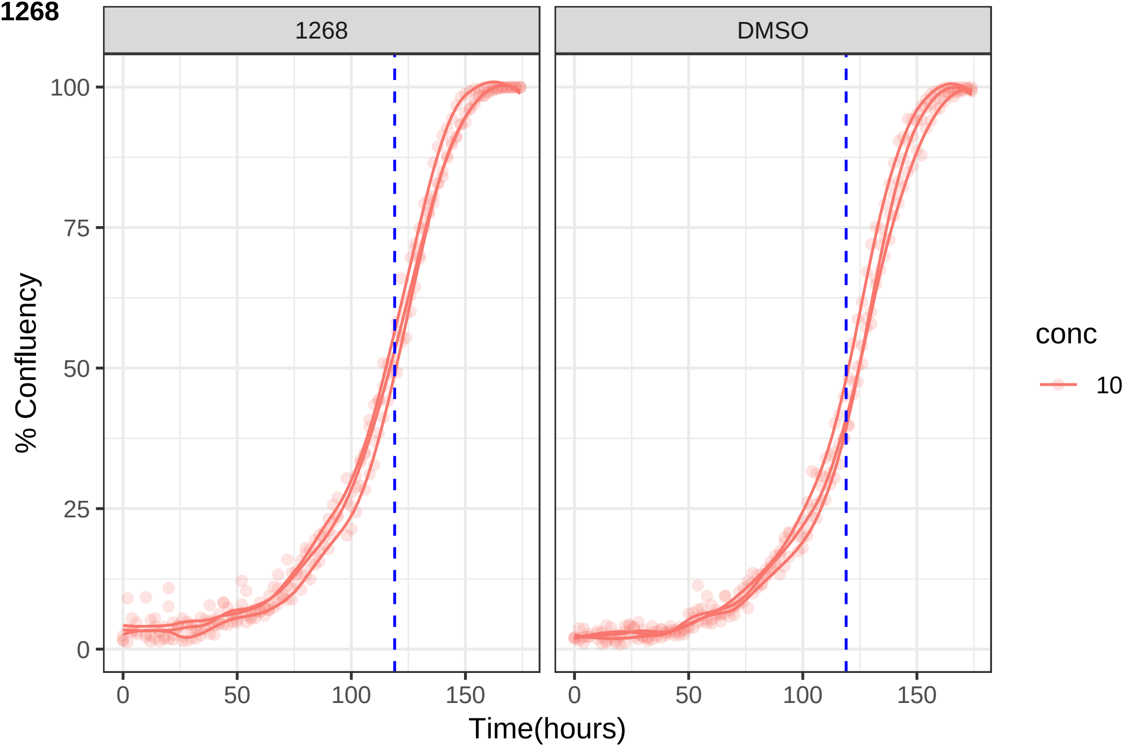
1268
DMSO
100
75
50
25
0
% Confluency
0
50
100
150
Time(hours)
conc
10

**1275**

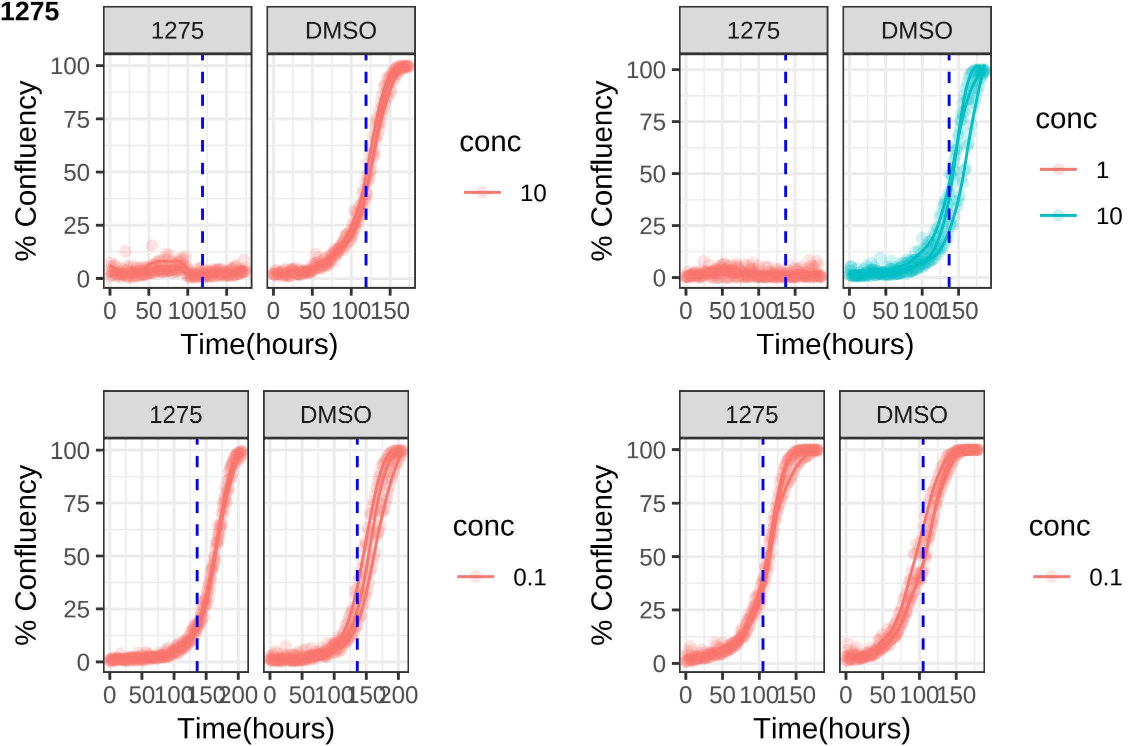

1275
DMSO
% Confluency
100
75
50
25
0
0 50 100 150
Time(hours)
conc
10
1275
DMSO
% Confluency
0 50 100 150
Time(hours)
conc
1
10
1275
DMSO
% Confluency
0 50 100 150 200
Time(hours)
conc
0.1
1275
DMSO
% Confluency
0 50 100 150
Time(hours)
conc
0.1

**1310**

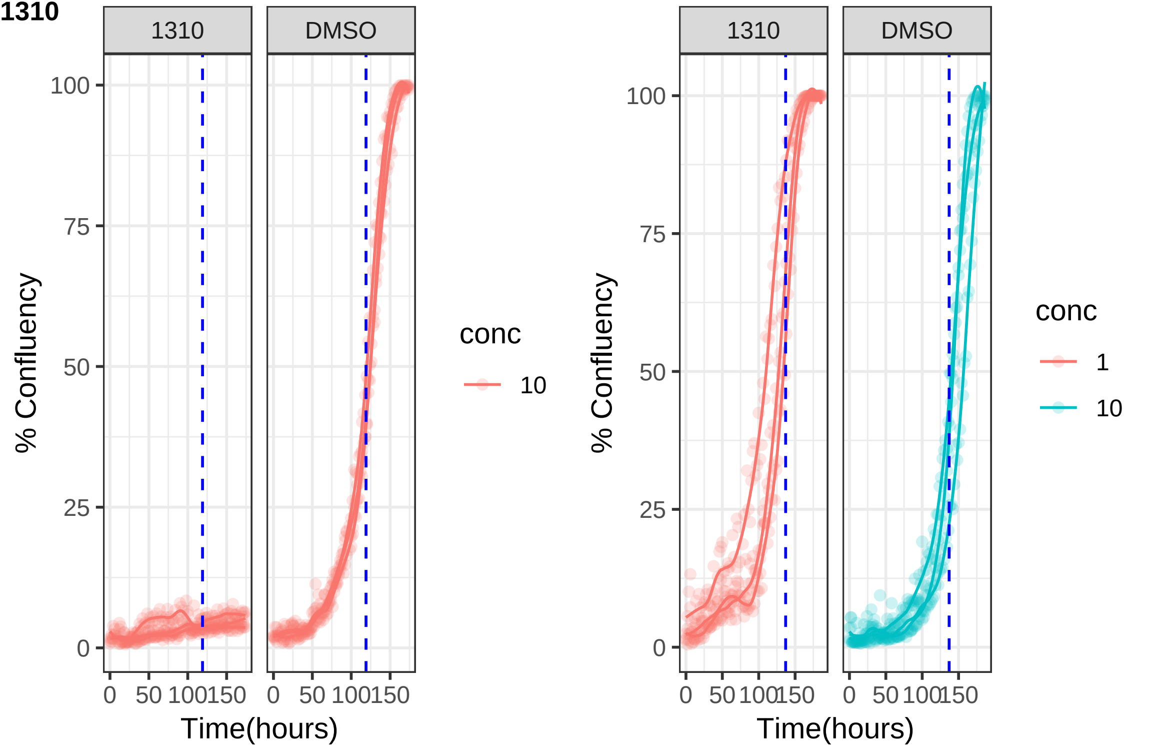

1310
DMSO
100
75
50
25
0
% Confluency
0
50
100
150
Time(hours)
conc
10
1310
DMSO
% Confluency
Time(hours)
conc
1
10

**1334**

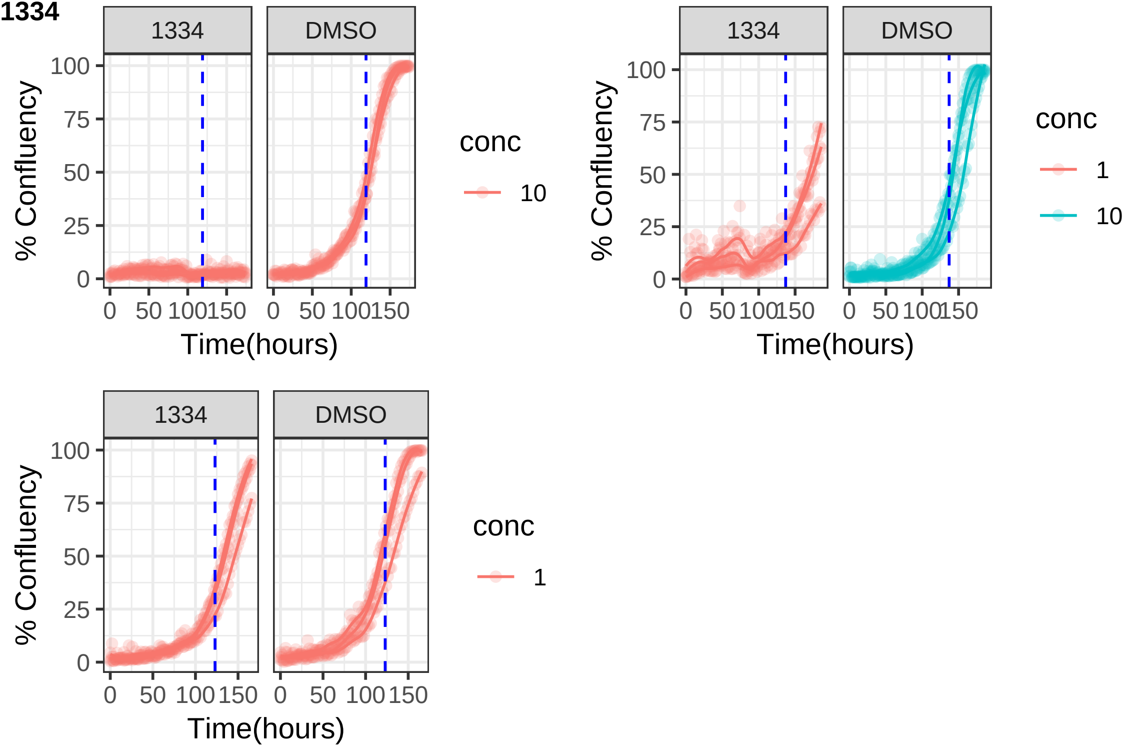
1334
DMSO
100
75
50
25
0
% Confluency
0 50 100 150
Time(hours)
conc
10
1

**1355**

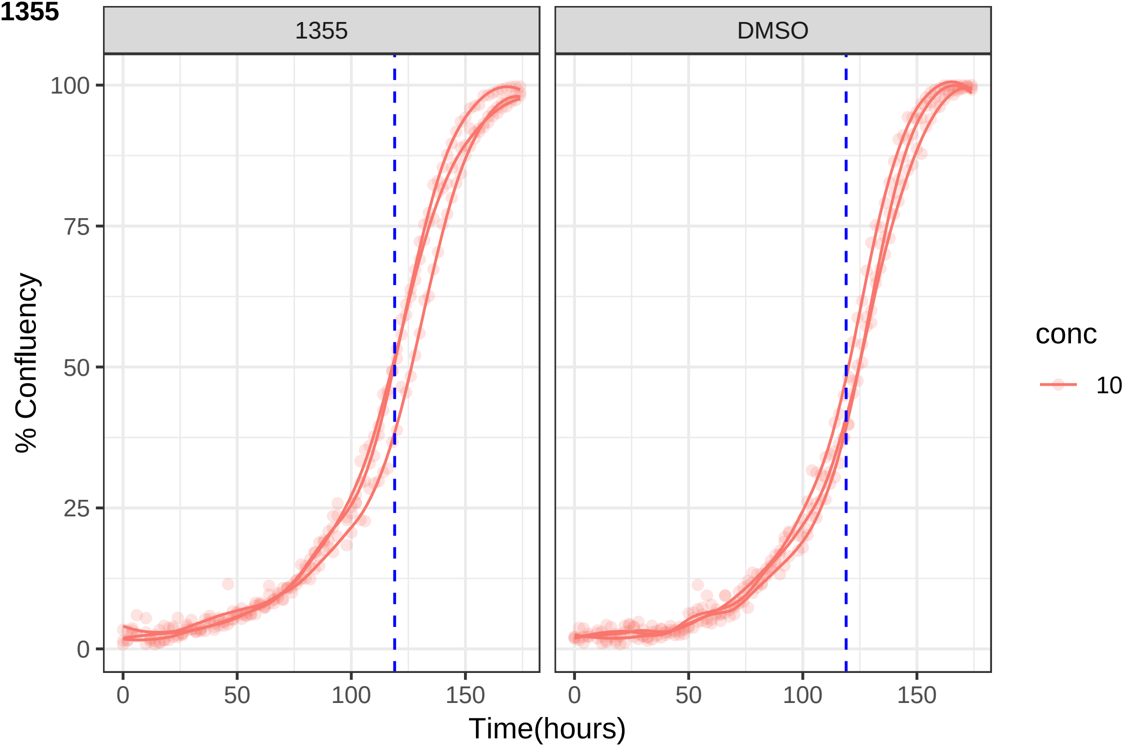
1355
DMSO
100
75
50
25
0
% Confluency
0
50
100
150
Time(hours)
conc
10

**1356**

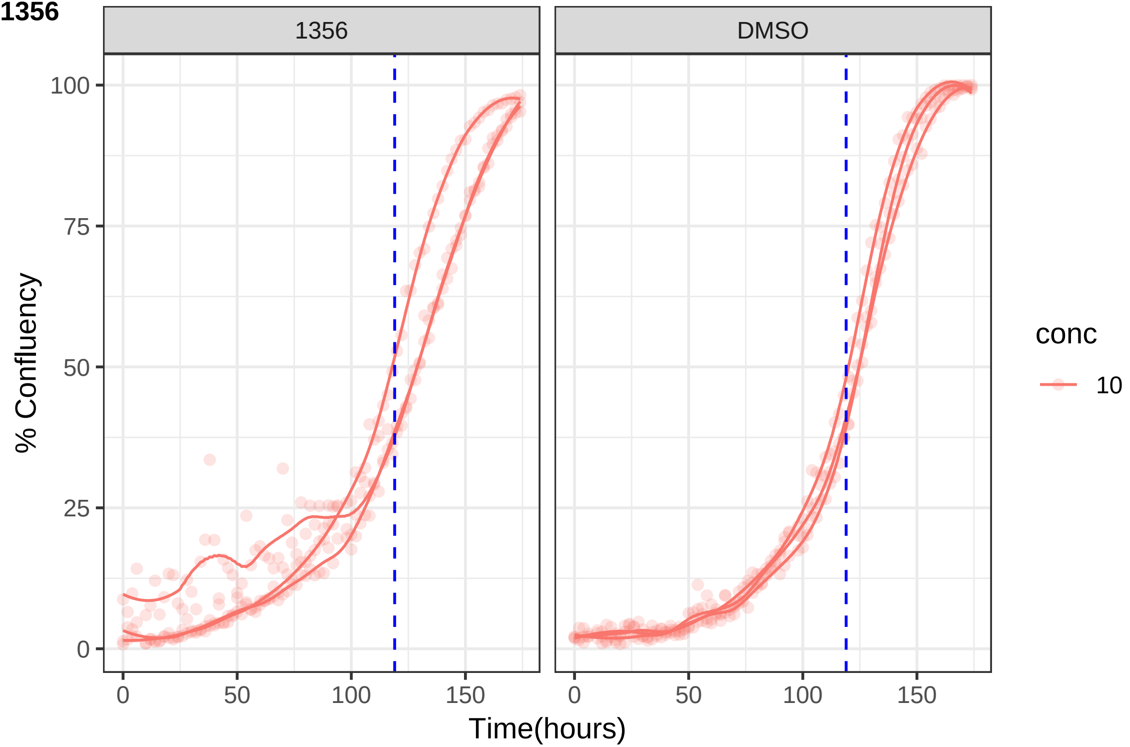
1356
DMSO
% Confluency
0
25
50
75
100
0
50
100
150
Time(hours)
conc
10

**1362**

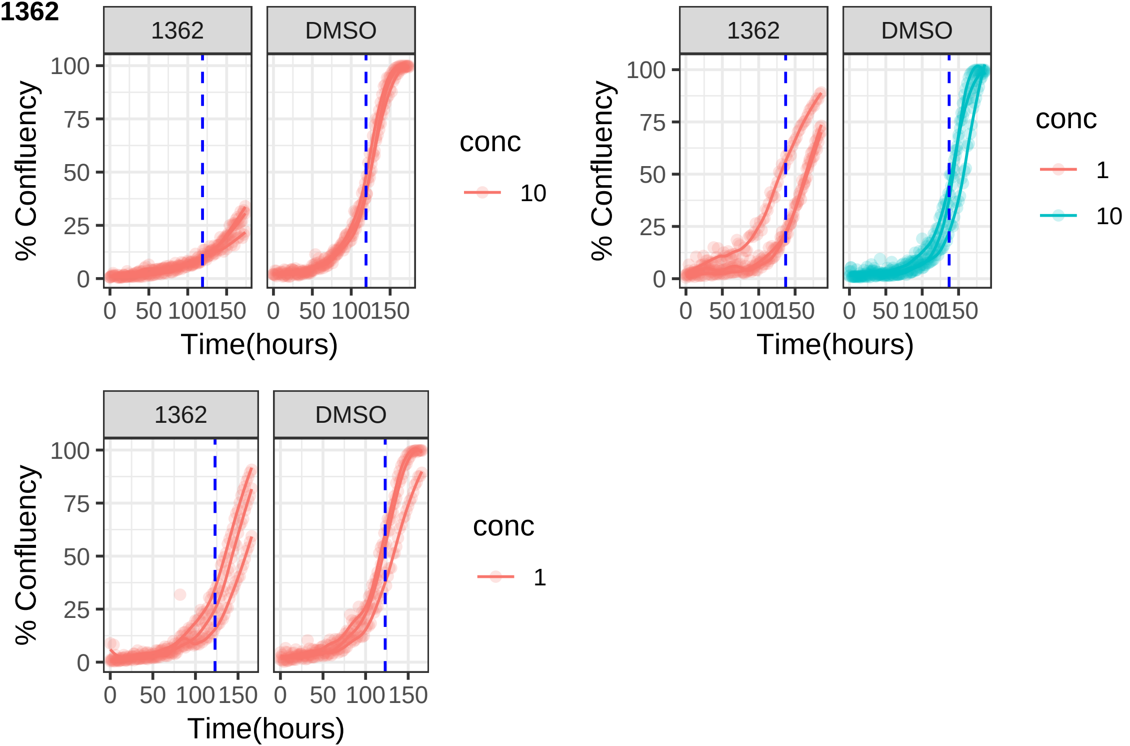
1362
DMSO
% Confluency
100
75
50
25
0
0
50
100
150
Time(hours)
conc
10
1

**1375**

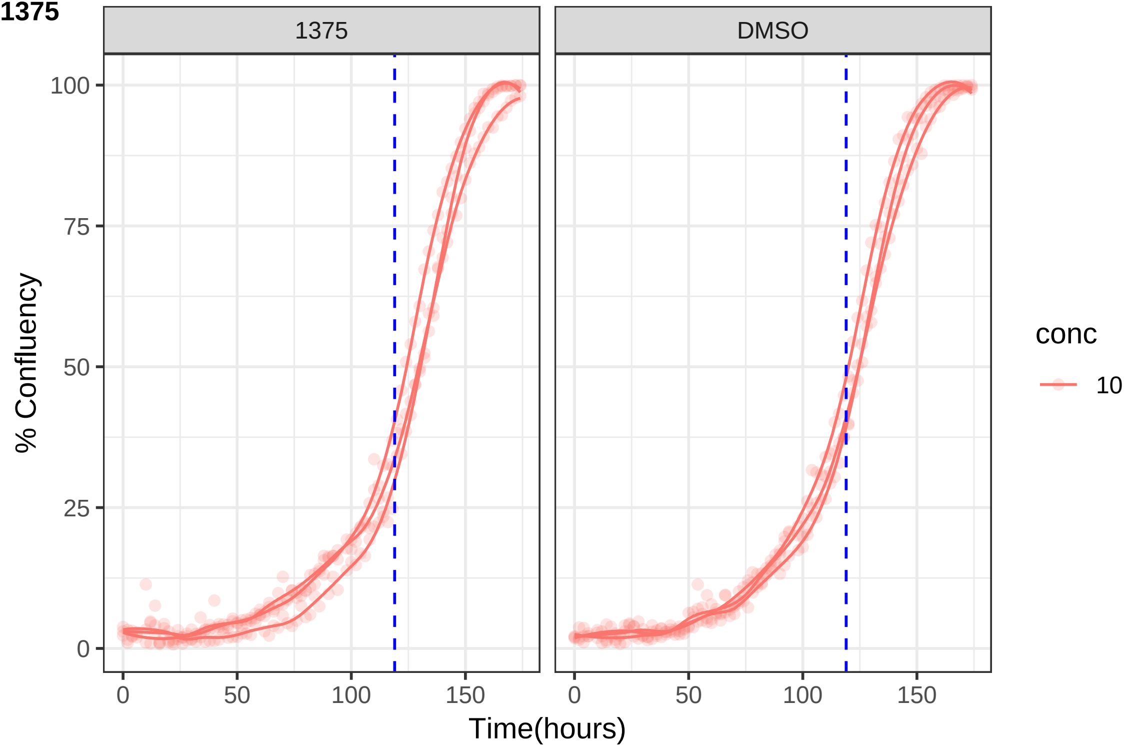

1375
DMSO
100
75
50
25
0
% Confluency
0
50
100
150
Time(hours)
conc
10

**1377**

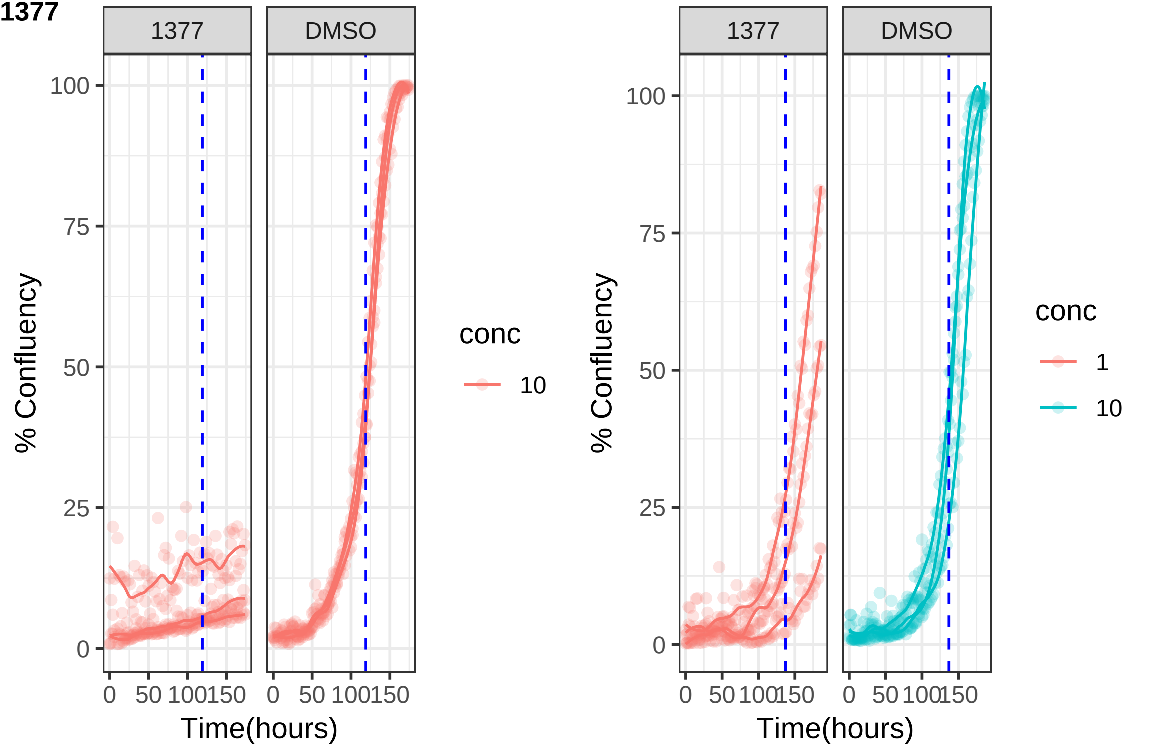
1377
DMSO
100
75
50
25
0
% Confluency
0 50 100 150
Time(hours)
conc
10
1

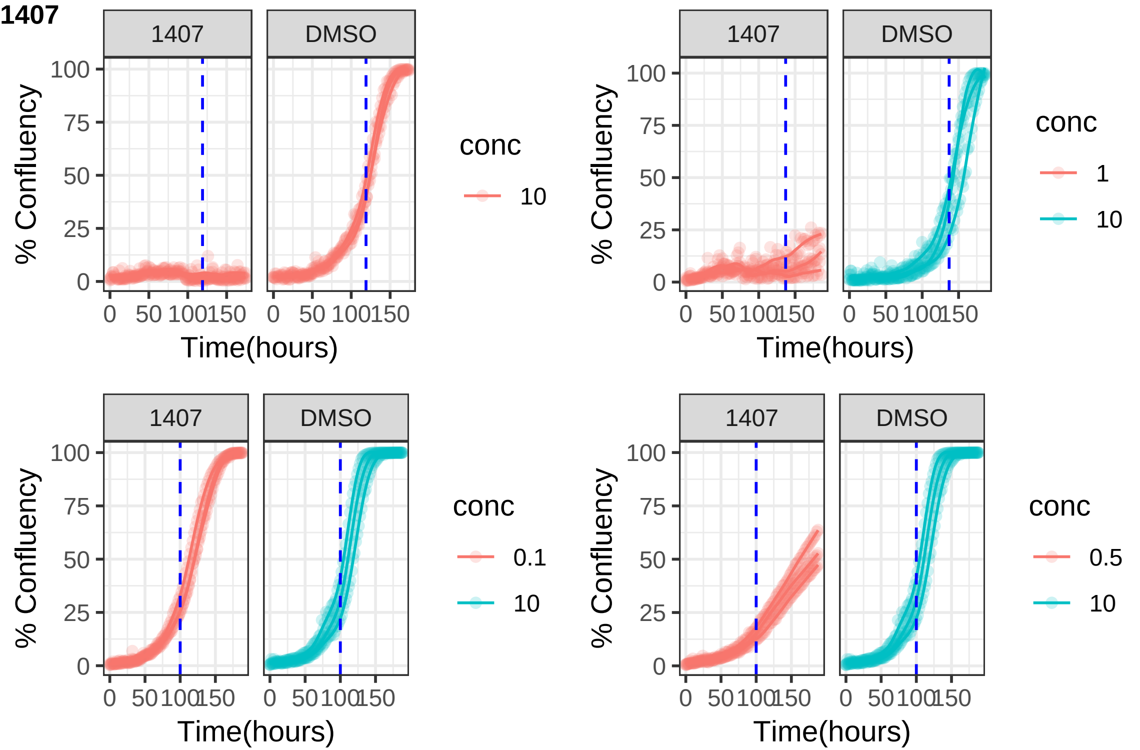
1407
1407
DMSO
% Confluency
100
75
50
25
0
0
50
100
150
Time(hours)
conc
10
1
0.1
0.5

**1424**

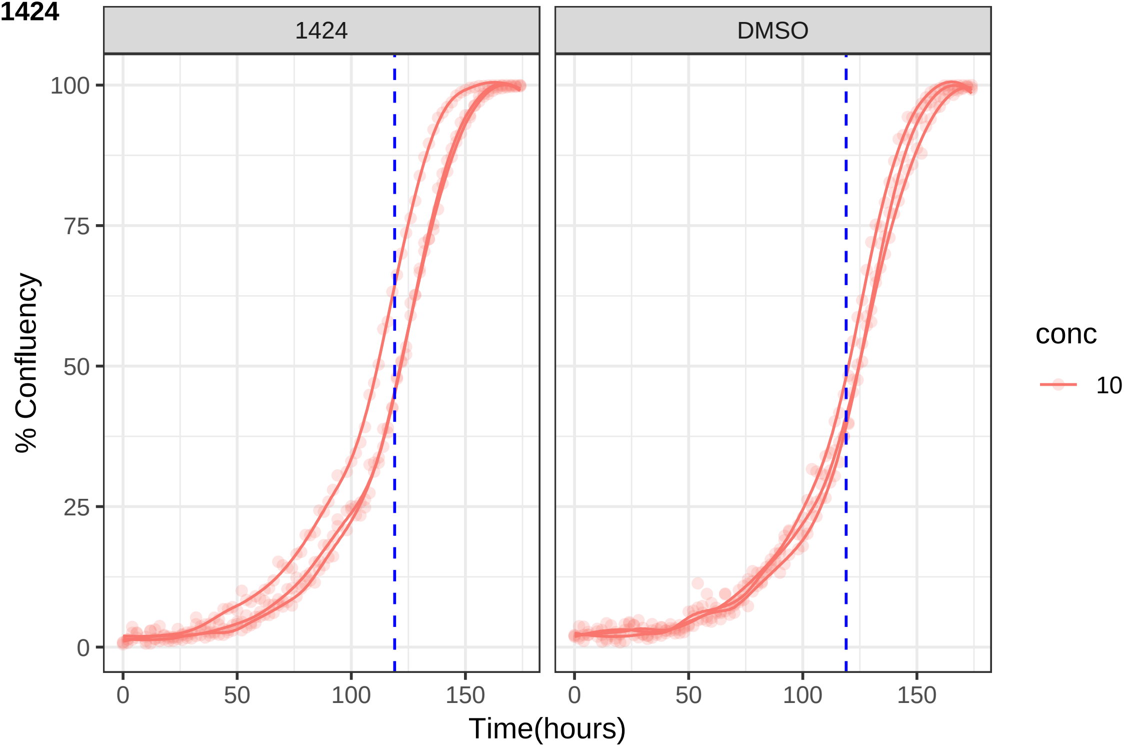

1424
DMSO
100
75
50
25
0
% Confluency
0
50
100
150
0
50
100
150
Time(hours)
conc
10

**1459**

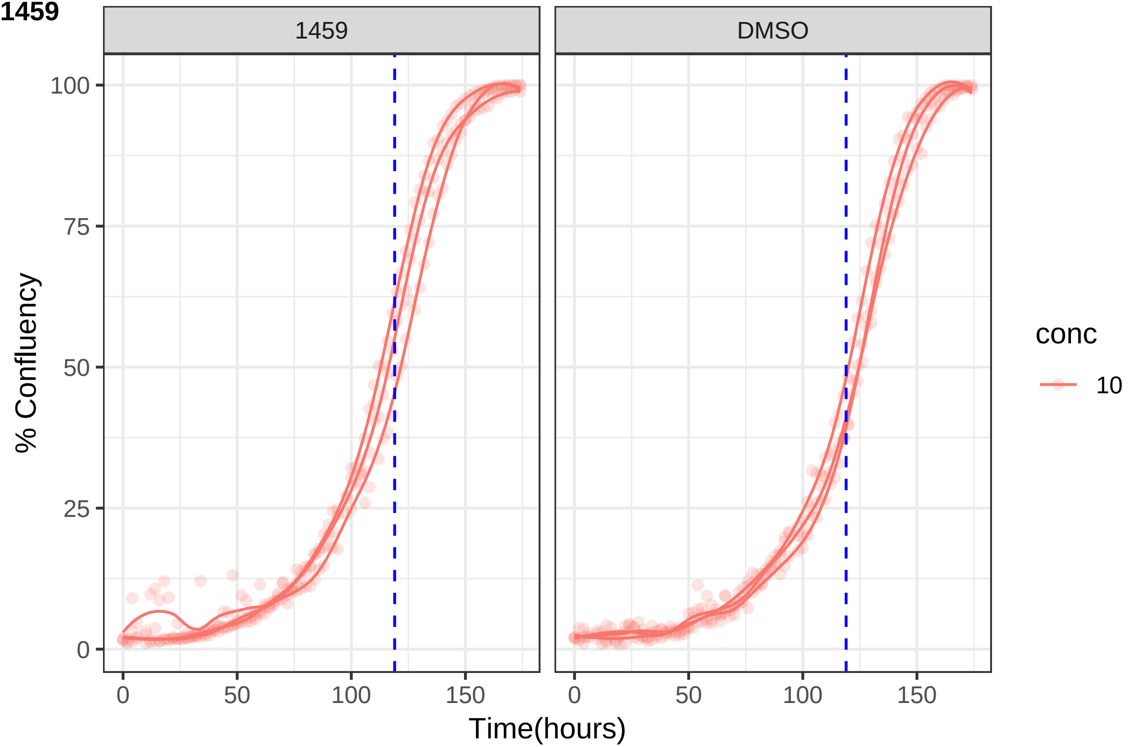

1459
DMSO
100
75
50
25
0
% Confluency
0
50
100
150
Time(hours)
conc
10

**1463**

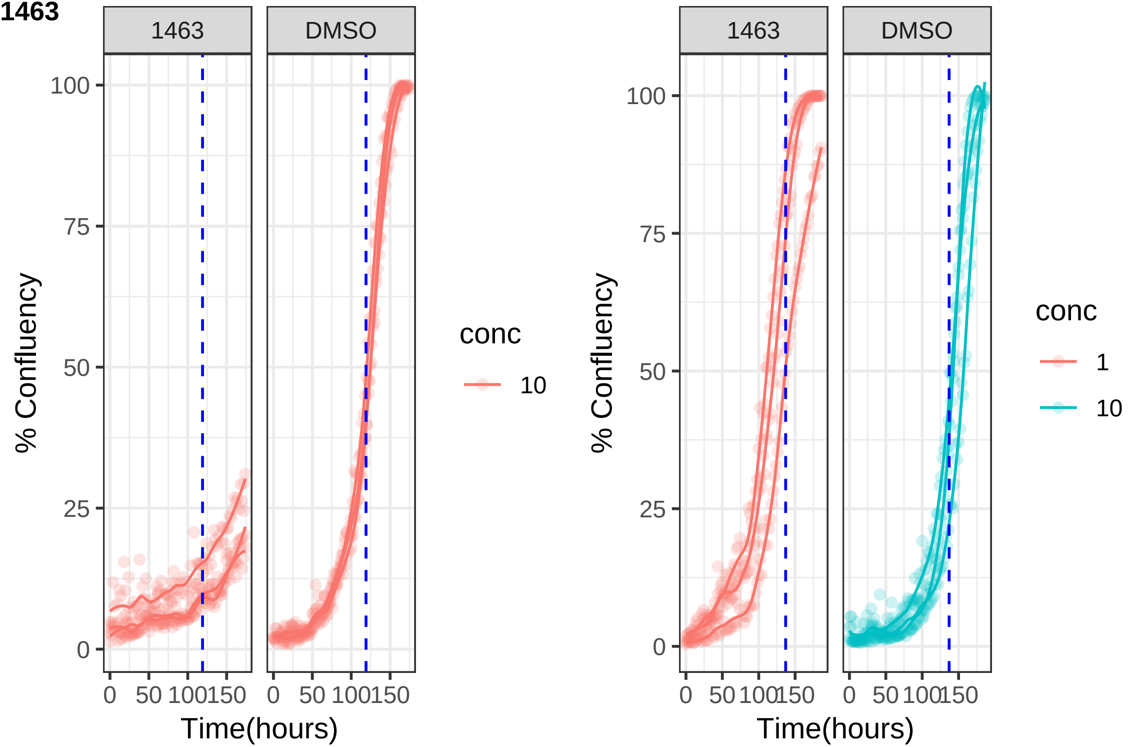
1463
DMSO
100
75
50
25
0
0
50
100
150
% Confluency
Time(hours)
conc
10
1

**1472**

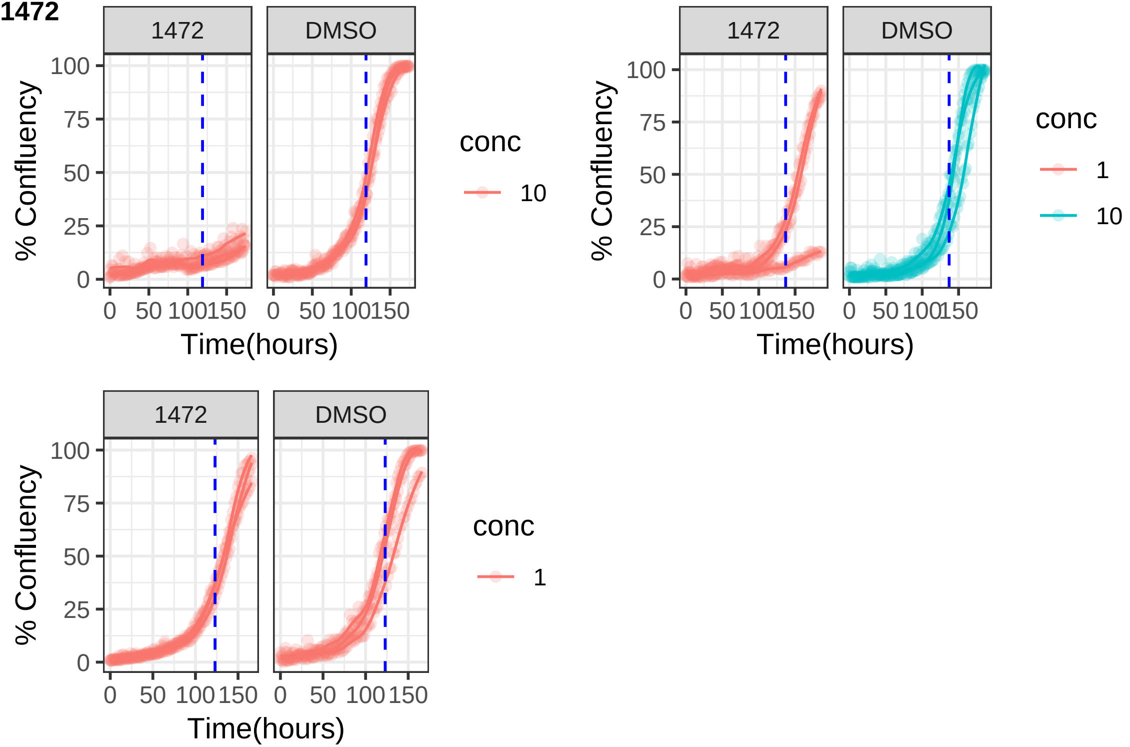

1472
DMSO
100
75
50
25
0
% Confluency
0
50
100
150
Time(hours)
conc
10
1472
DMSO
% Confluency
Time(hours)
conc
1
10
1472
DMSO
% Confluency
Time(hours)
conc
1

**1494**

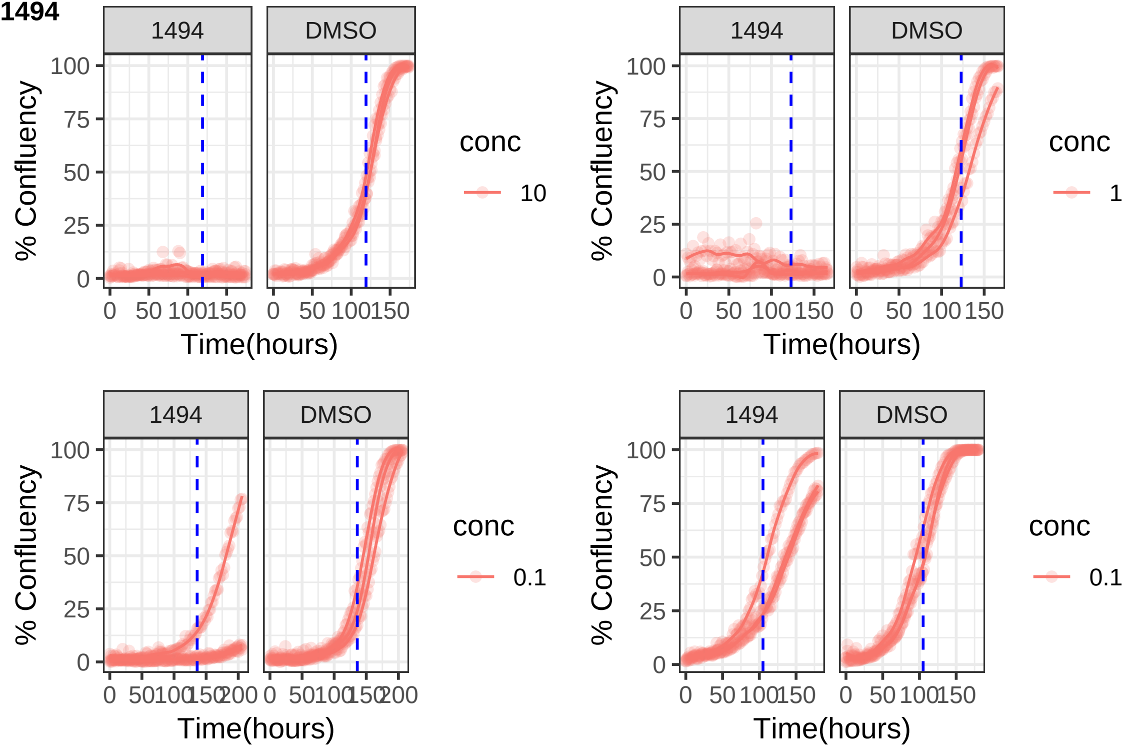
1494
DMSO
% Confluency
Time(hours)
conc
10
1
0.1
0
25
50
75
100
0
50
100
150
200

**1516**

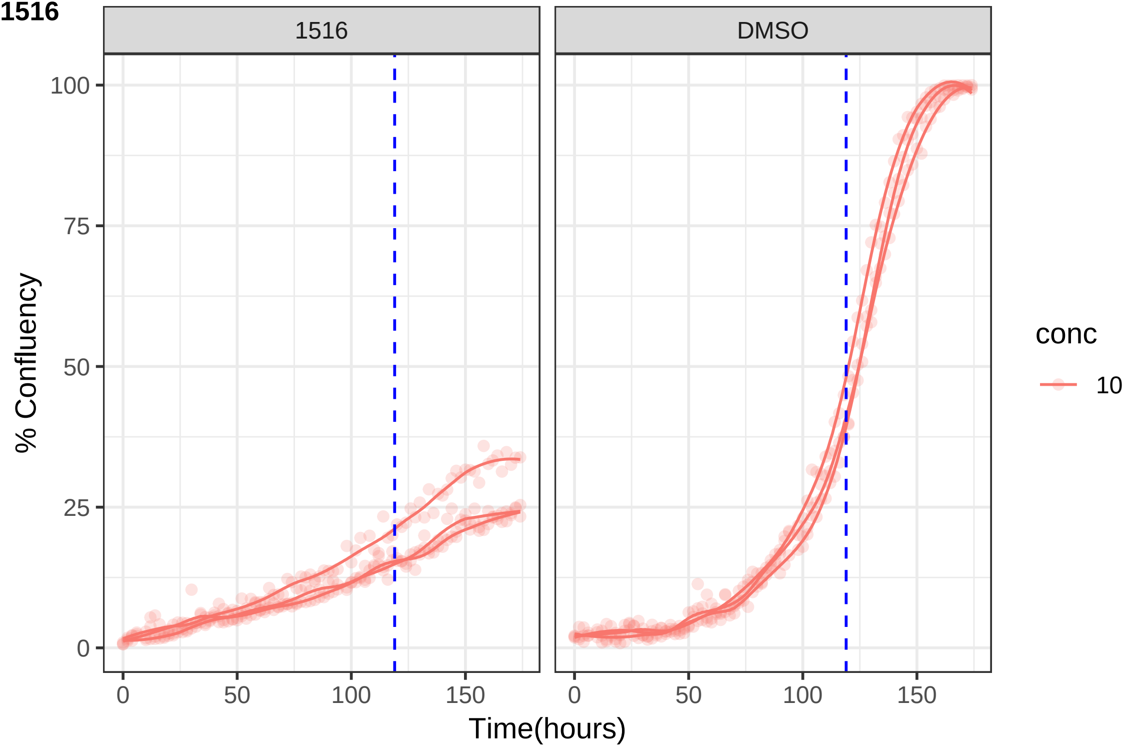

1516
DMSO
100
75
50
25
0
% Confluency
0
50
100
150
Time(hours)
conc
10

**1520**

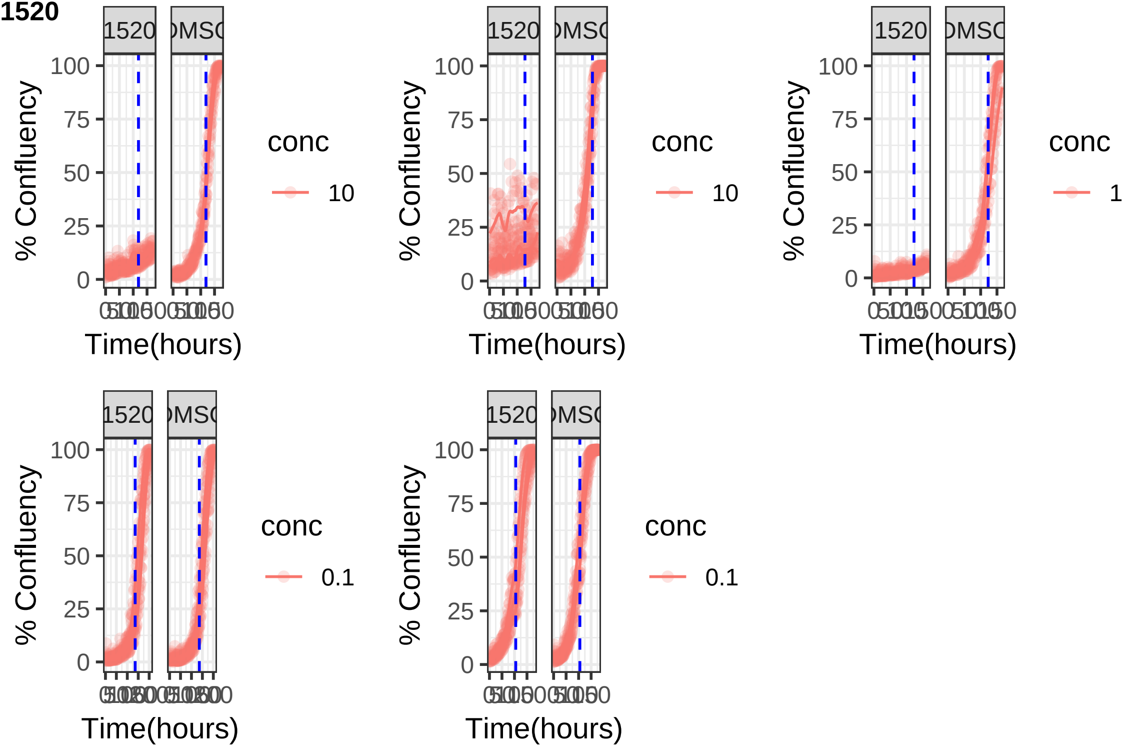

**1561**

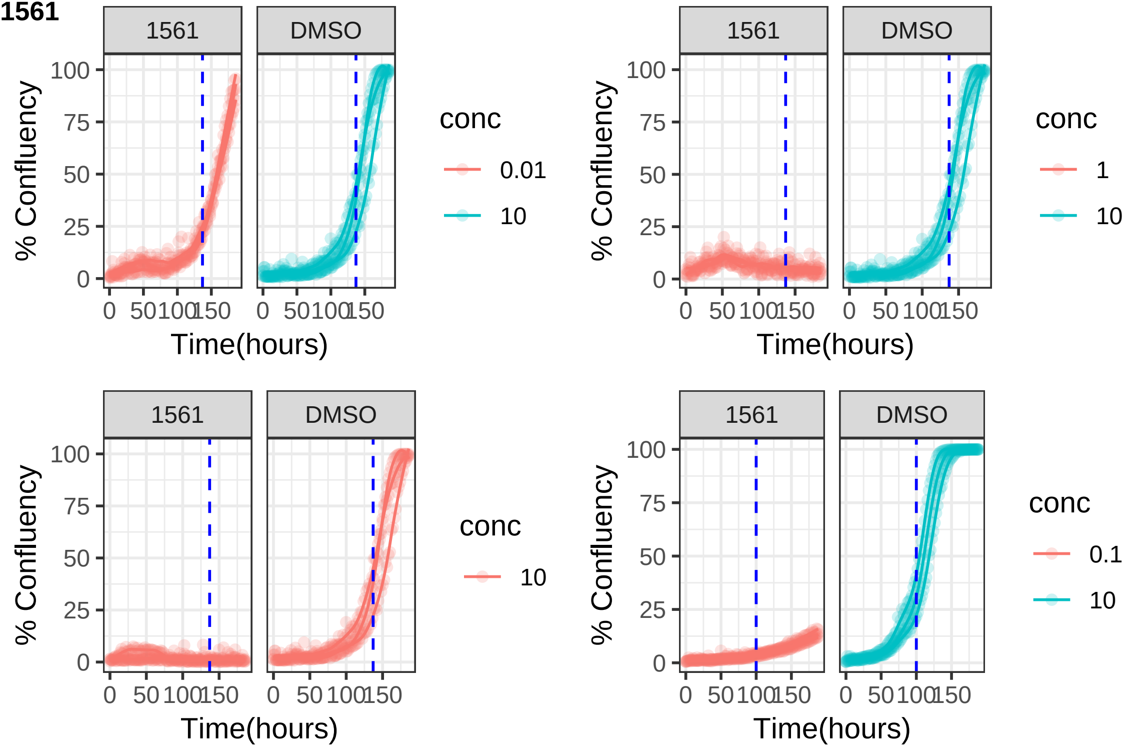
1561
DMSO
% Confluency
100
75
50
25
0
0
50
100
150
Time(hours)
conc
0.01
10
1561
DMSO
conc
1
10
1561
DMSO
conc
10
1561
DMSO
conc
0.1
10

**1566**

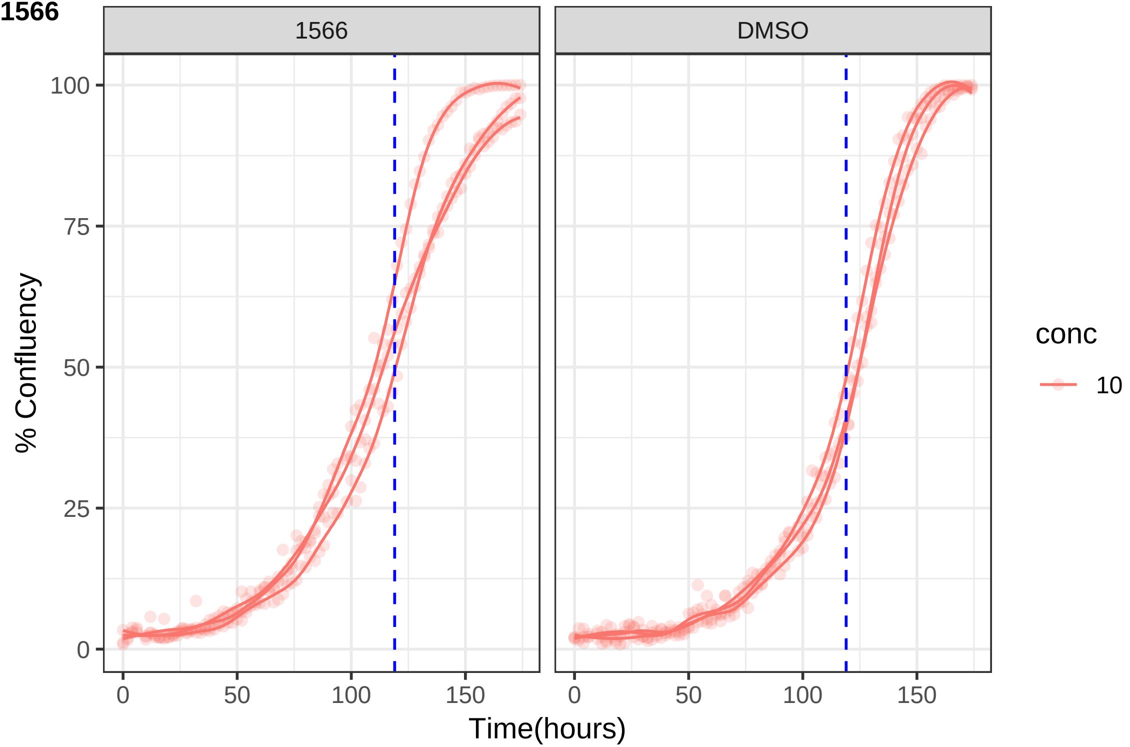

1566
DMSO
% Confluency
Time(hours)
0
25
50
75
100
0
50
100
150
conc
10

**1596**

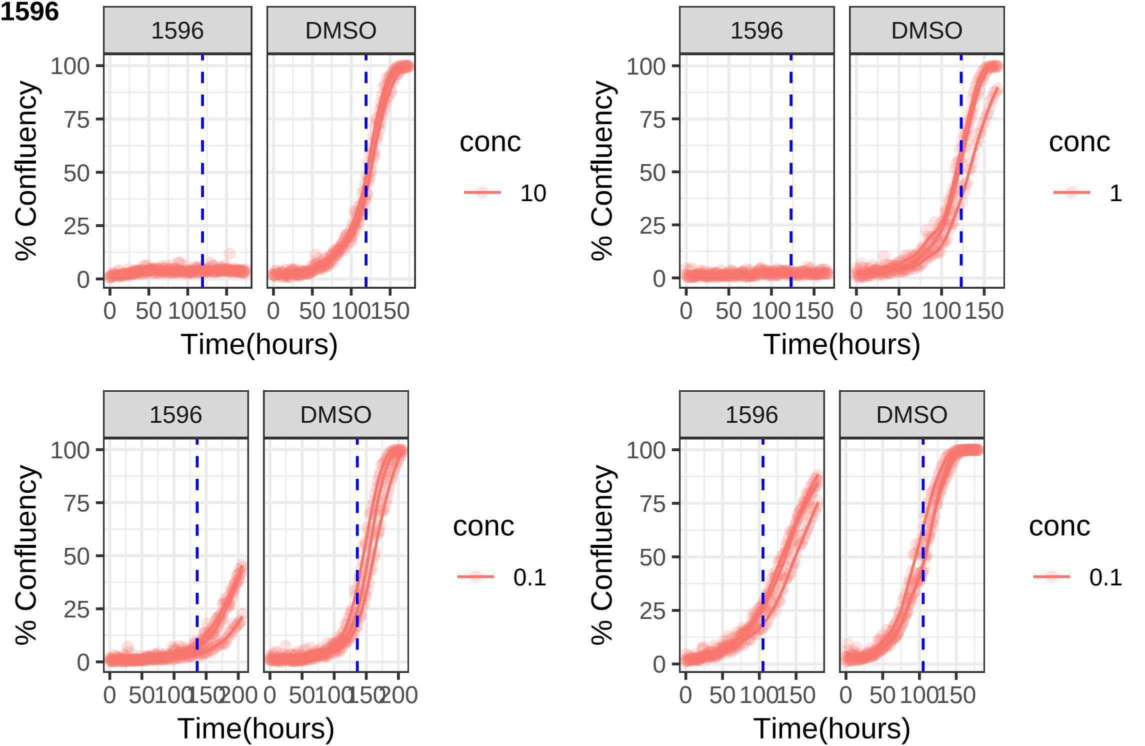
1596
DMSO
100
75
50
25
0
% Confluency
0 50 100 150
Time(hours)
conc
10
1
0.1
0 50 100 150 200

**1629**

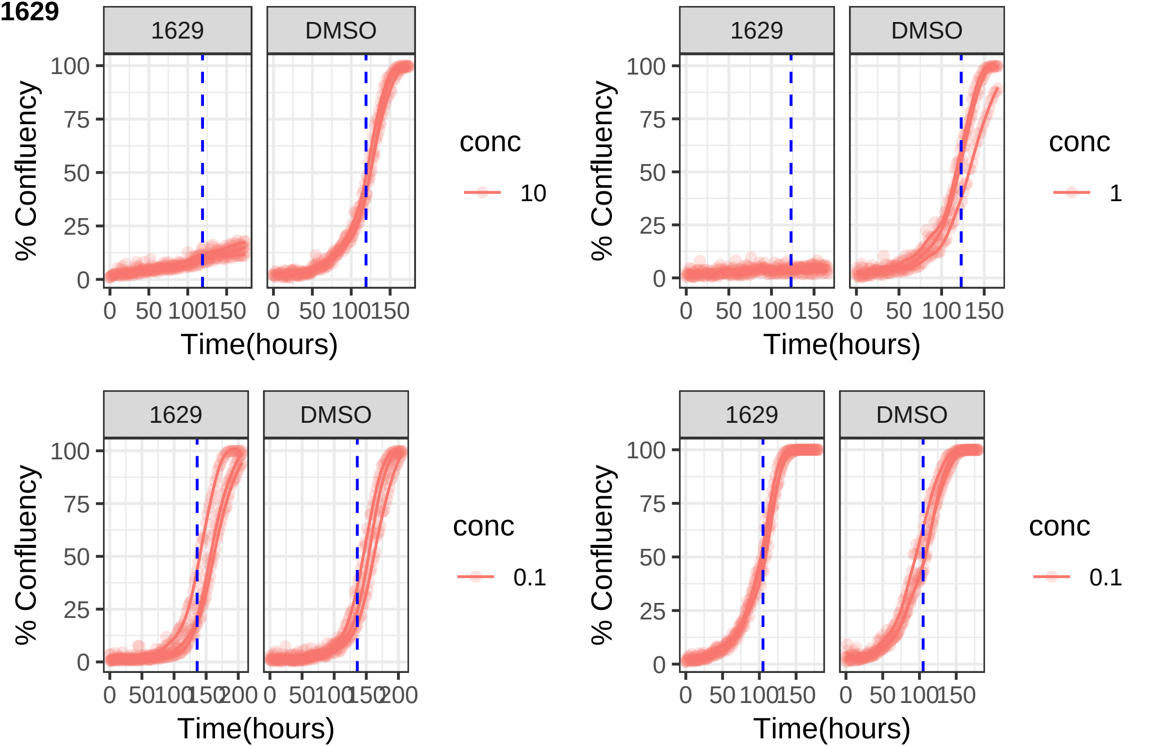
1629
DMSO
% Confluency
Time(hours)
conc
10
1
0.1
0.1
0
25
50
75
100
0
50
100
150
200

**1645**

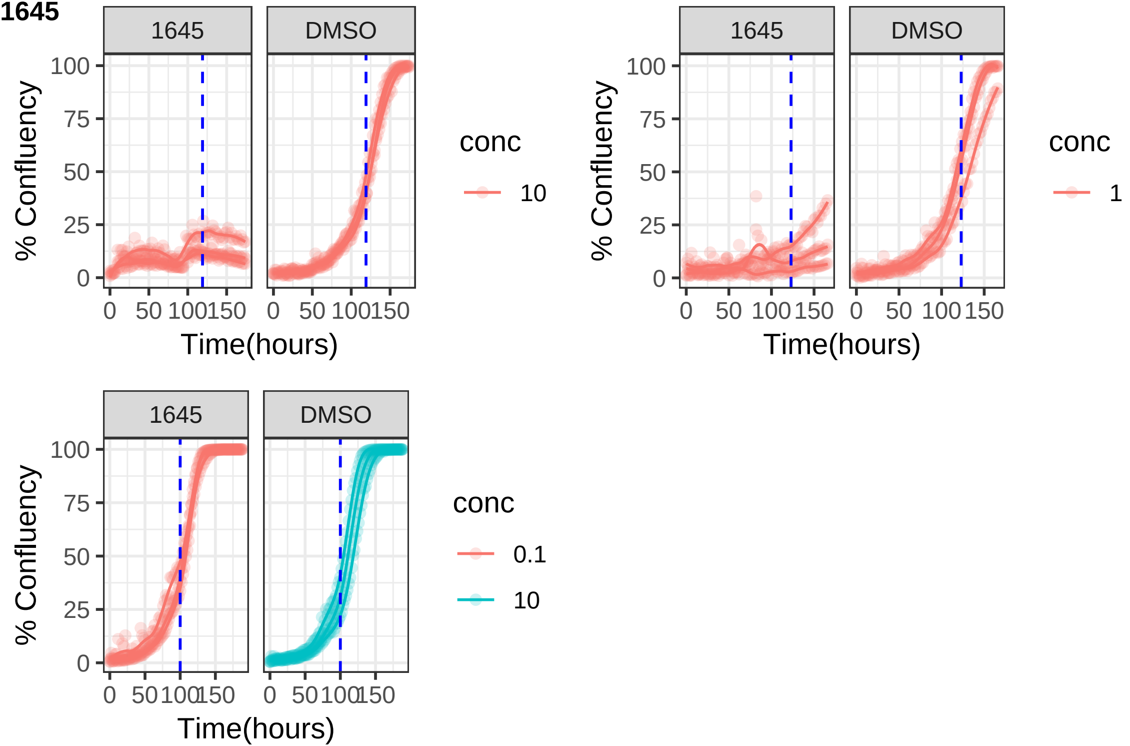
1645
DMSO
100
75
50
25
0
% Confluency
0
50
100
150
Time(hours)
conc
10
1
0.1

**1687**

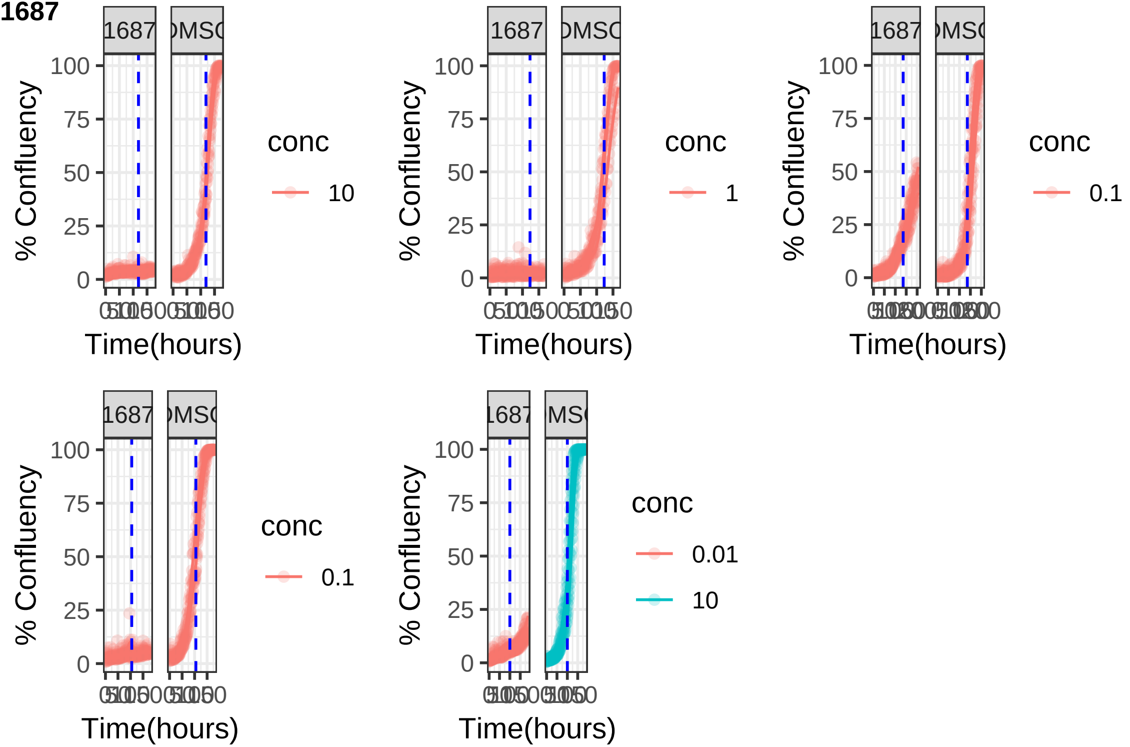

**1691**

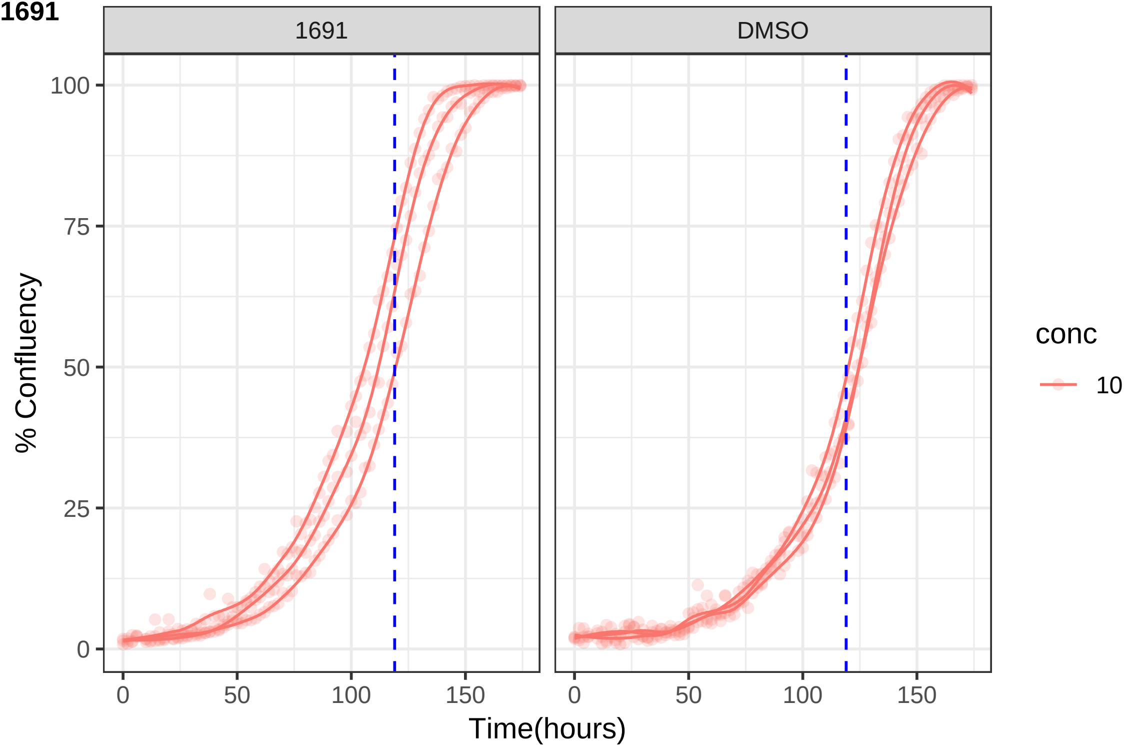
1691
DMSO
% Confluency
100
75
50
25
0
0
50
100
150
Time(hours)
conc
10

**1729**

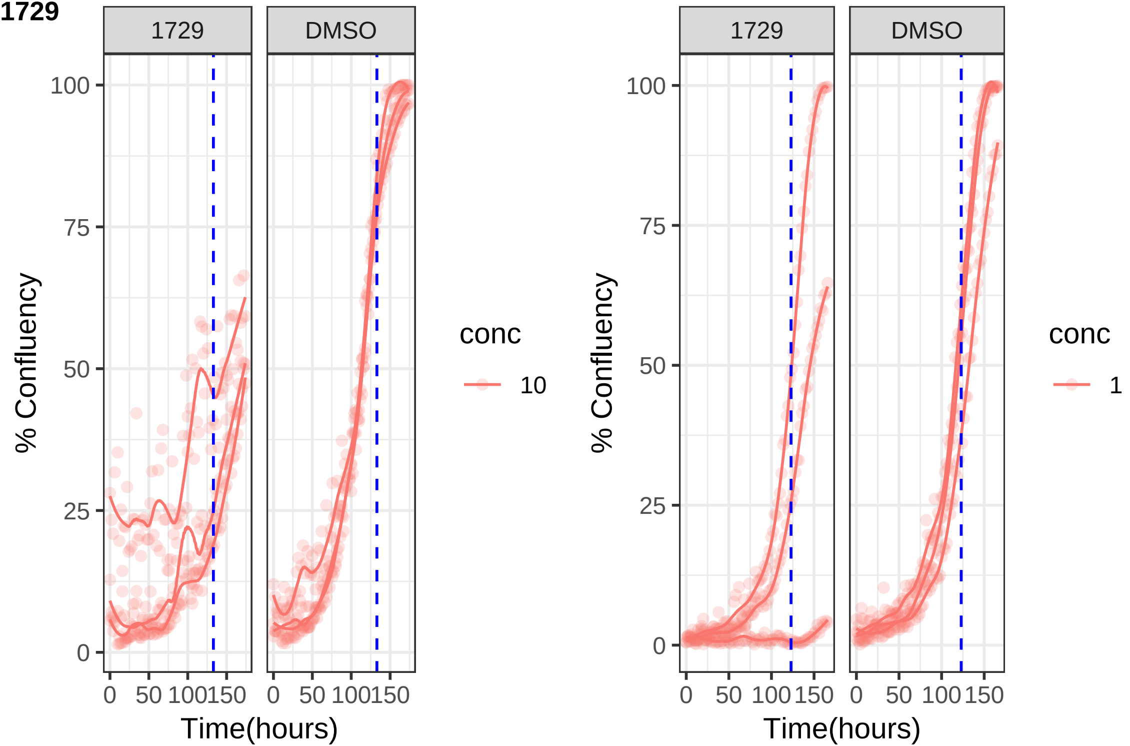
1729
DMSO
100
75
50
25
0
% Confluency
0 50 100 150
Time(hours)
conc
10
1729
DMSO
100
75
50
25
0
% Confluency
0 50 100 150
Time(hours)
conc
1

1743

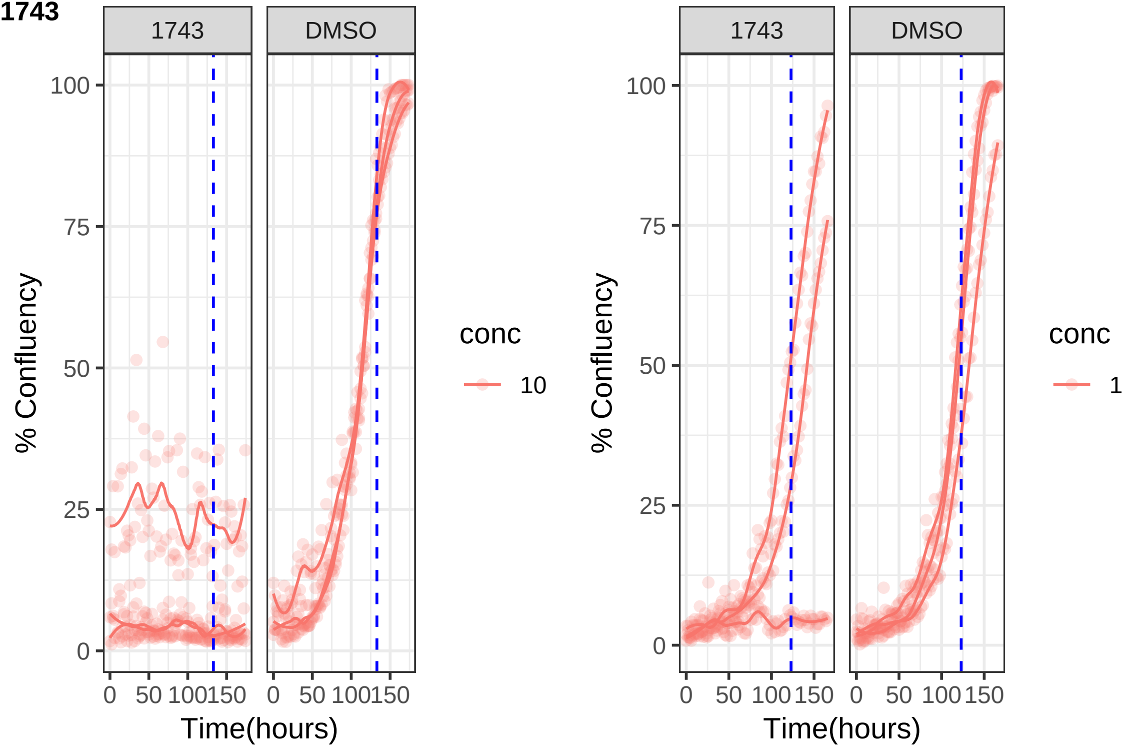

1743
DMSO
100
75
50
25
0
% Confluency
0
50
100
150
Time(hours)
conc
10
1

**1765**

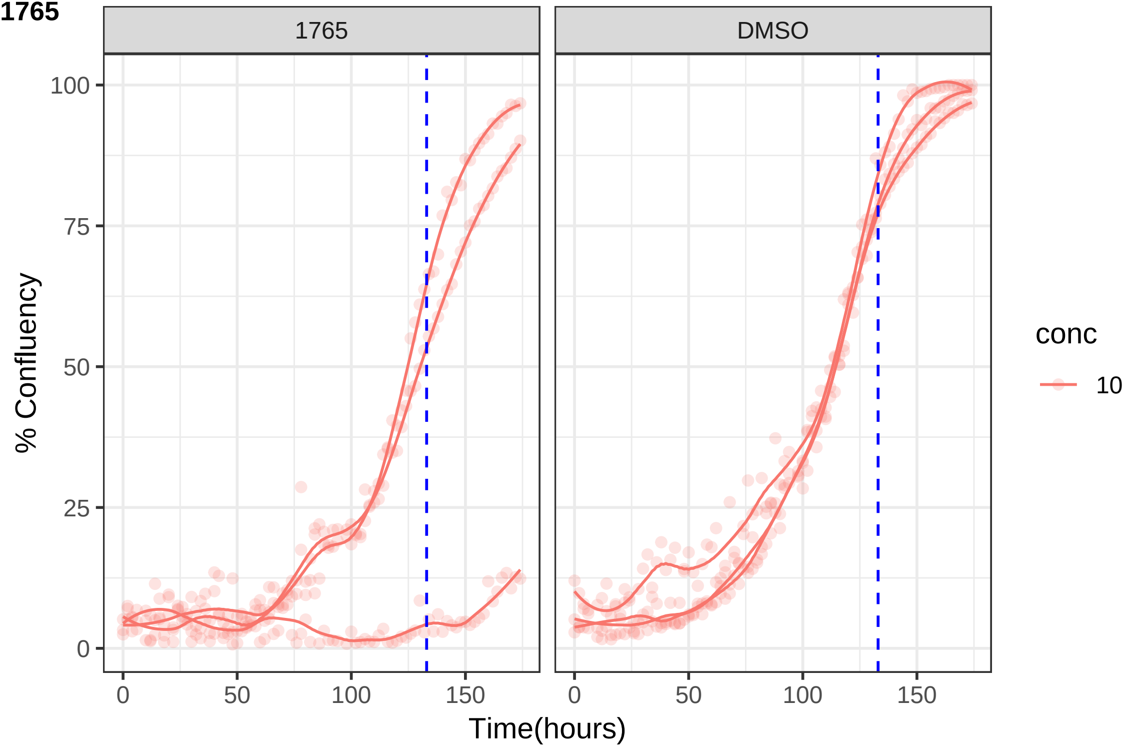

1765
DMSO
100
75
50
25
0
% Confluency
0
50
100
150
Time(hours)
conc
10

**1789**

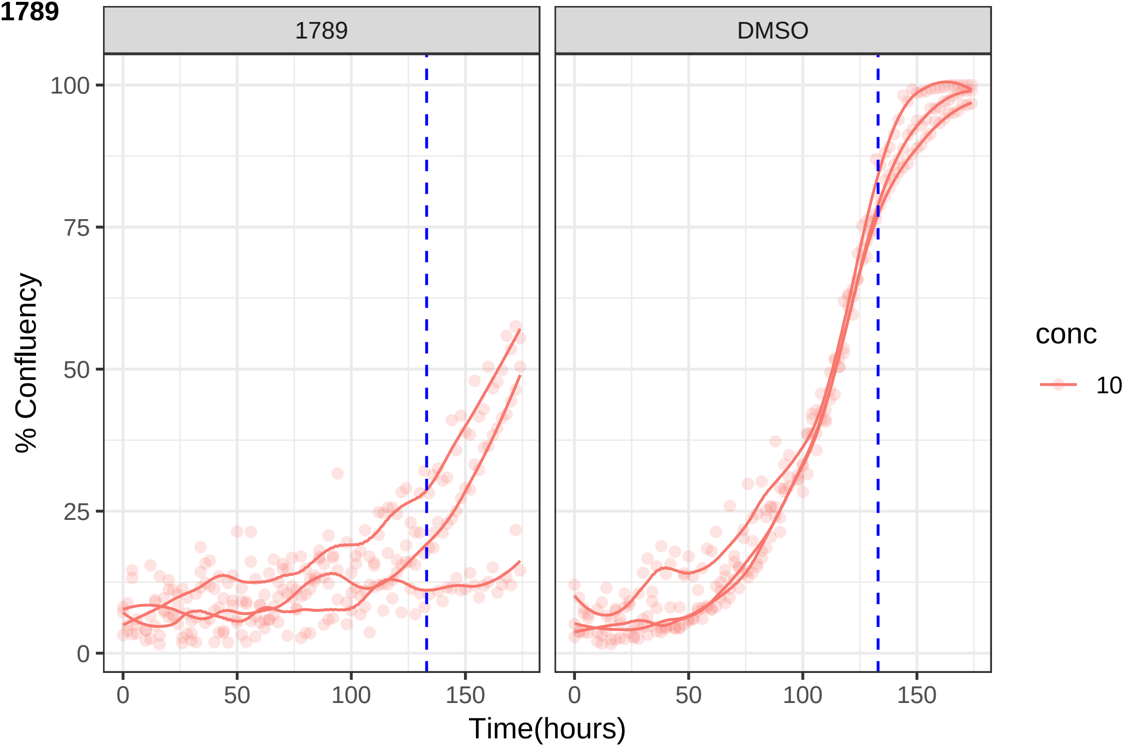

1789
DMSO
% Confluency
100
75
50
25
0
0
50
100
150
Time(hours)
conc
10

**1797**

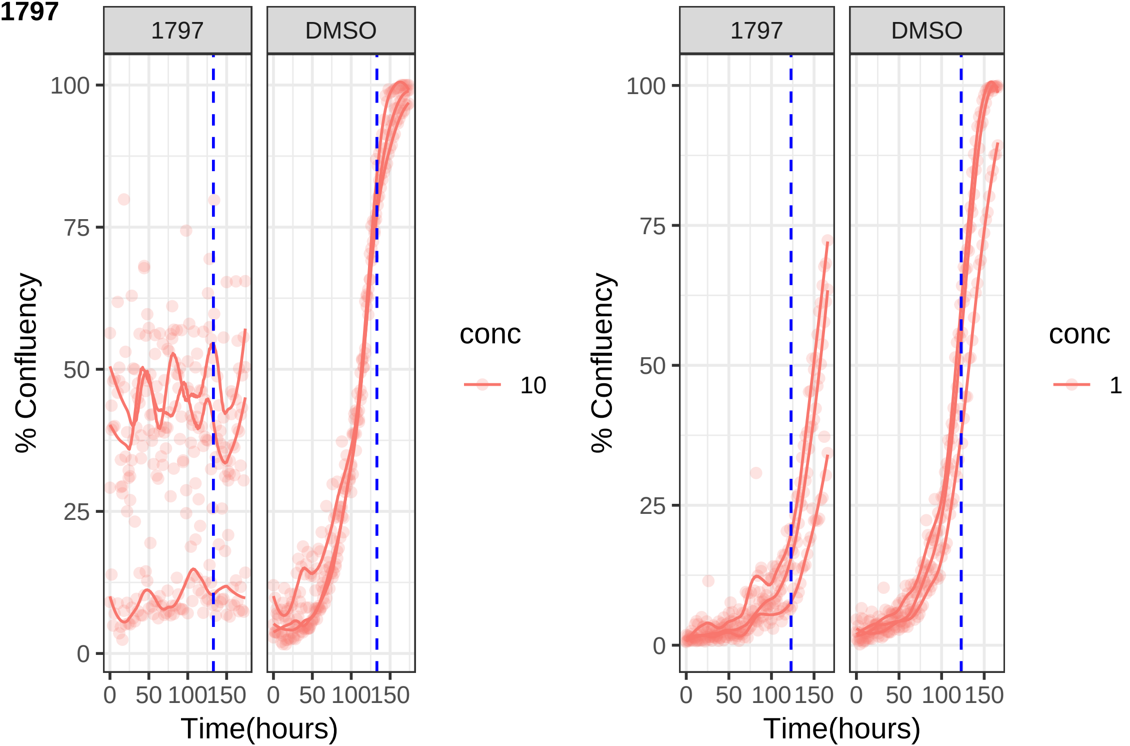

1797
DMSO
100
75
50
25
0
% Confluency
0
50
100
150
Time(hours)
conc
10
1797
DMSO
% Confluency
Time(hours)
conc
1

**1801**

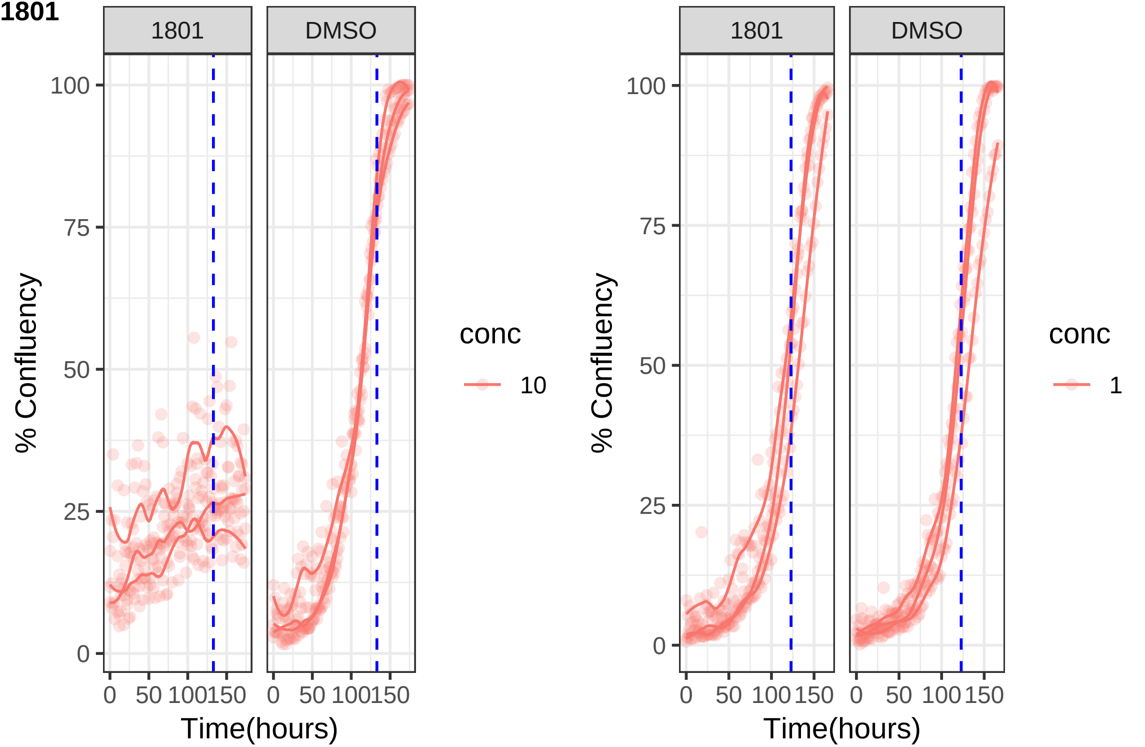

1801
DMSO
100
75
50
25
0
% Confluency
0
50
100
150
Time(hours)
conc
10
1801
DMSO
100
75
50
25
0
% Confluency
0
50
100
150
Time(hours)
conc
1

**1803**

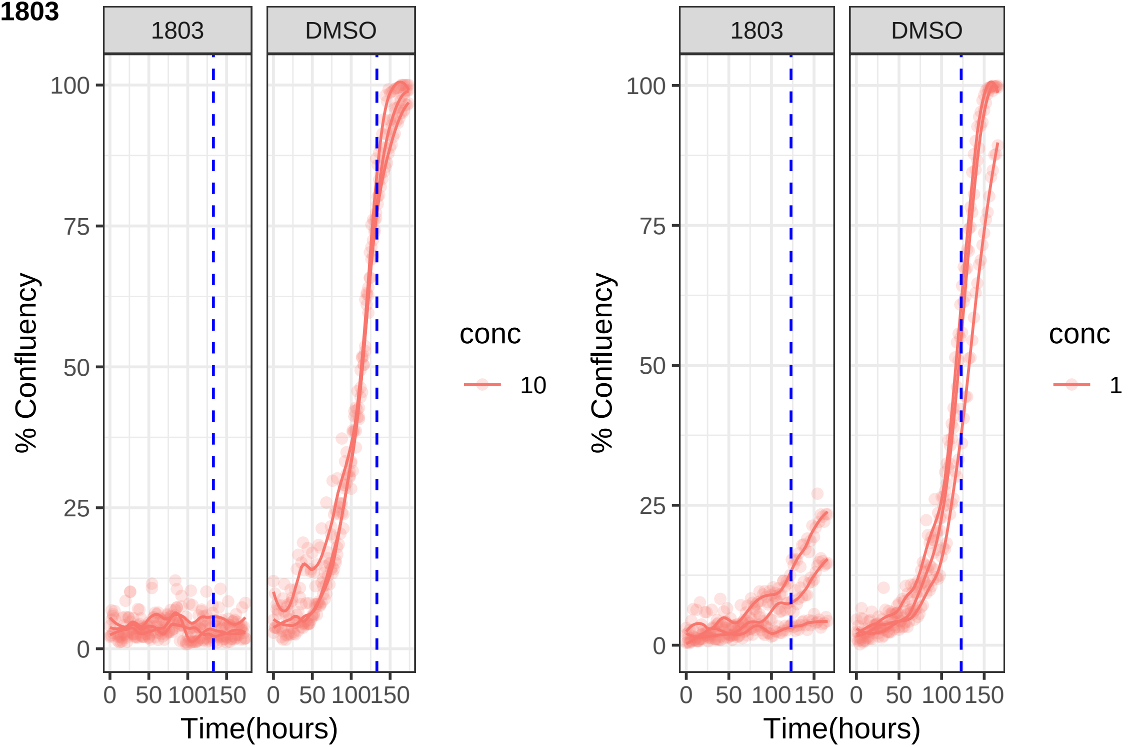

1803
DMSO
100
75
50
25
0
% Confluency
0
50
100
150
Time(hours)
conc
10
1803
DMSO
100
75
50
25
0
% Confluency
0
50
100
150
Time(hours)
conc
1

**1807**

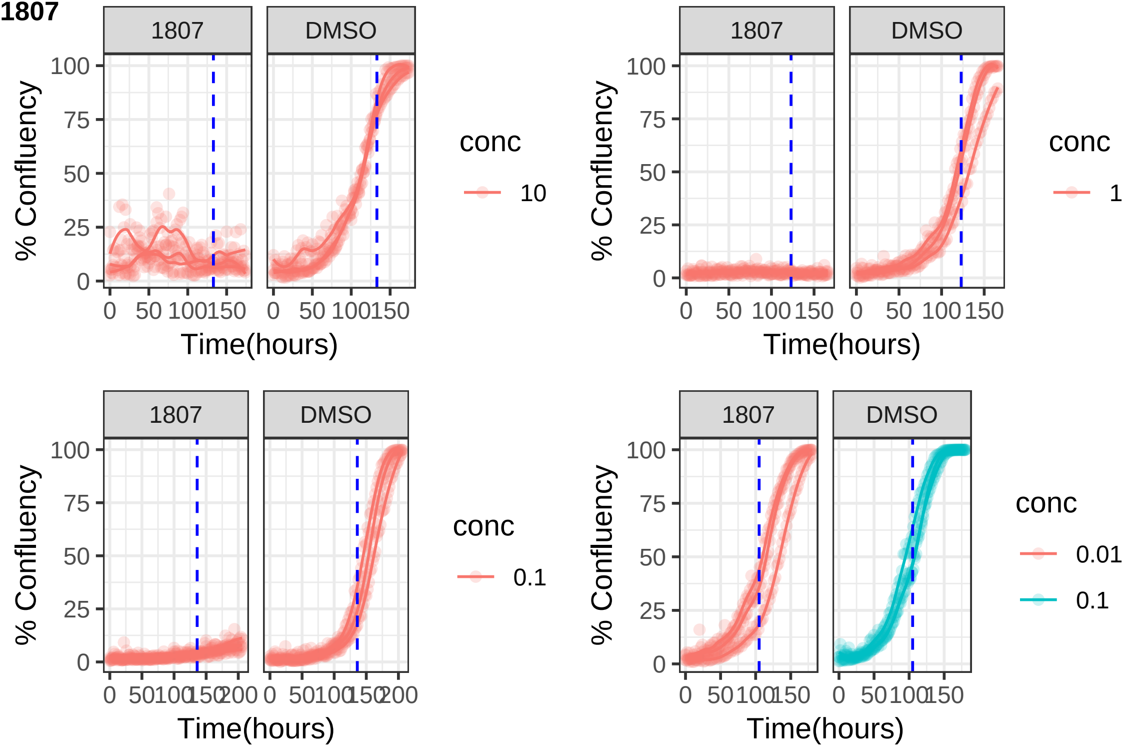
1807
DMSO
% Confluency
Time(hours)
conc
10
1
0.1
0.01
100
75
50
25
0
50
150
200

**1814**

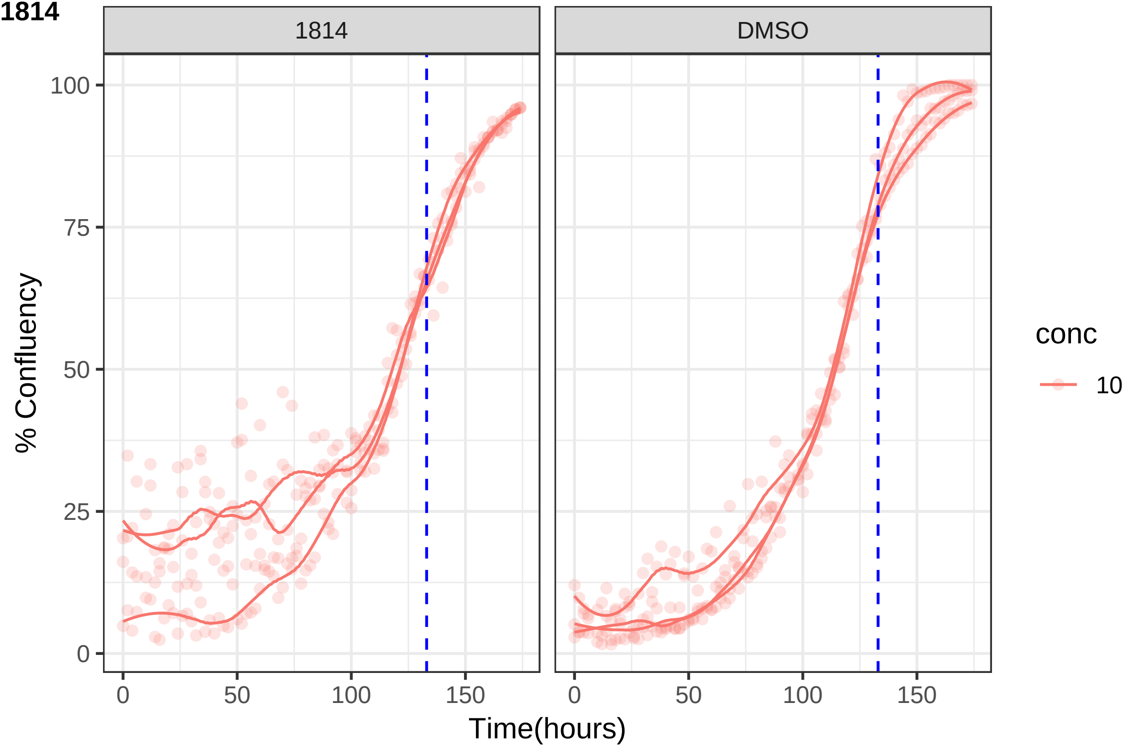
1814
DMSO
100
75
50
25
0
% Confluency
0
50
100
150
Time(hours)
conc
10

**1831**

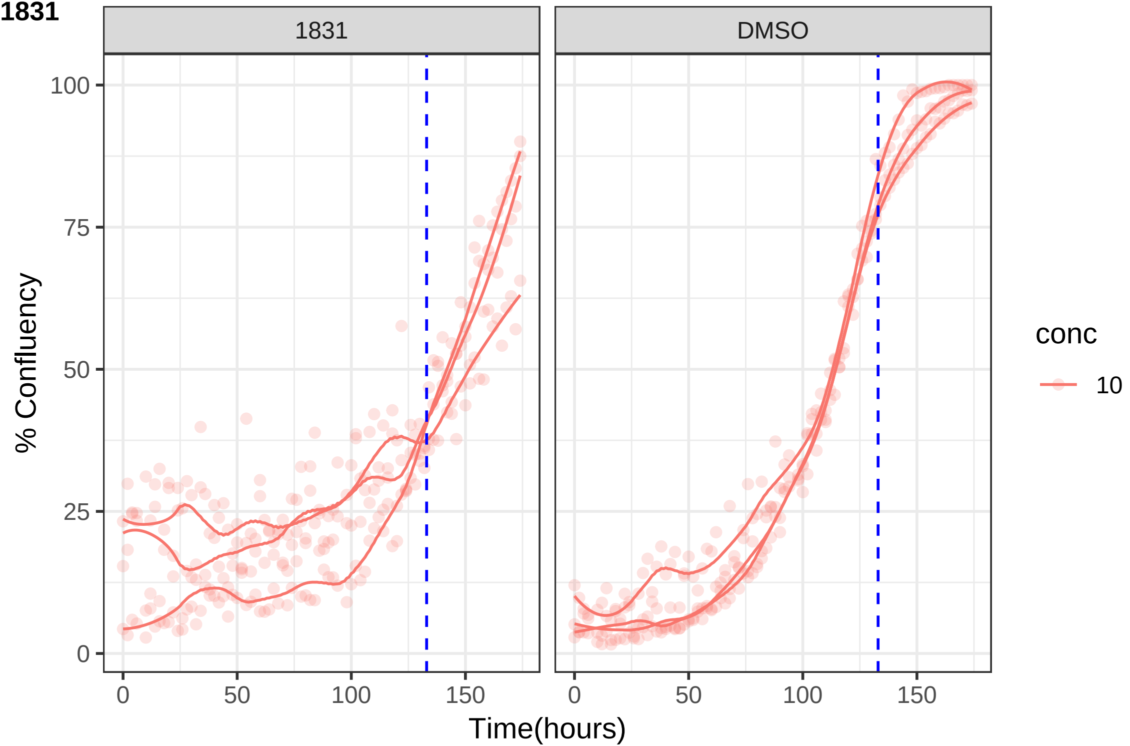

1831
DMSO
100
75
50
25
0
% Confluency
0
50
100
150
Time(hours)
conc
10

**1853**

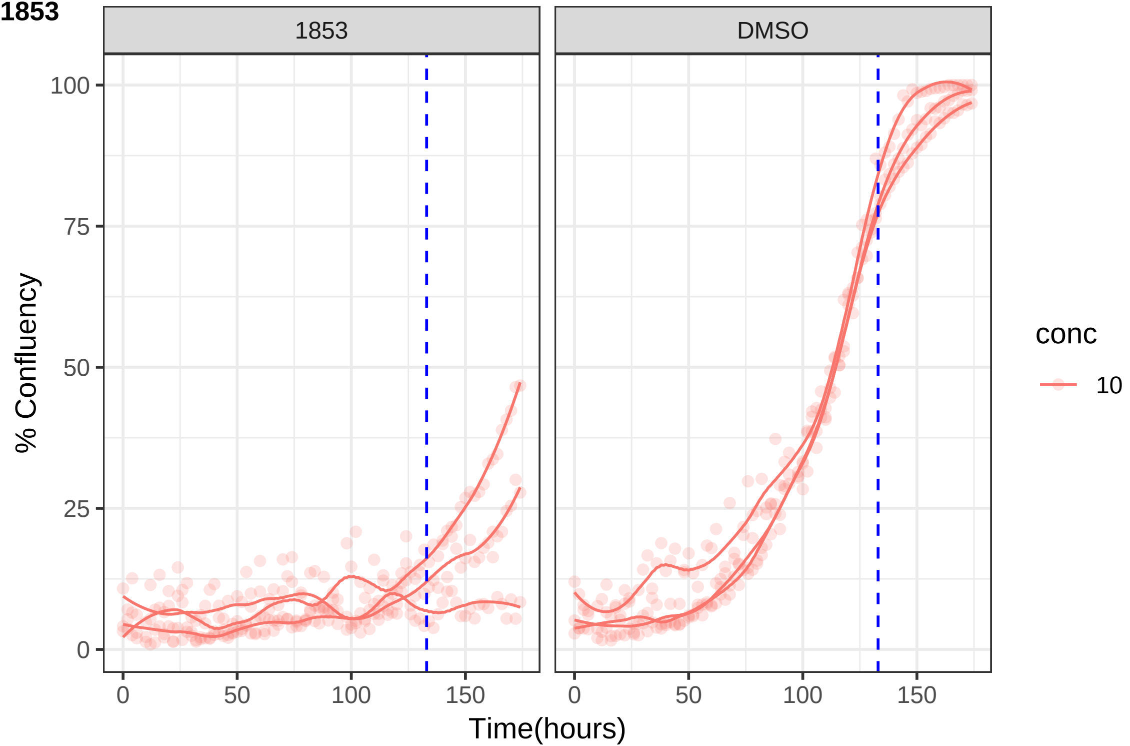

1853
DMSO
100
75
50
25
0
% Confluency
0
50
100
150
Time(hours)
conc
10

**1895**

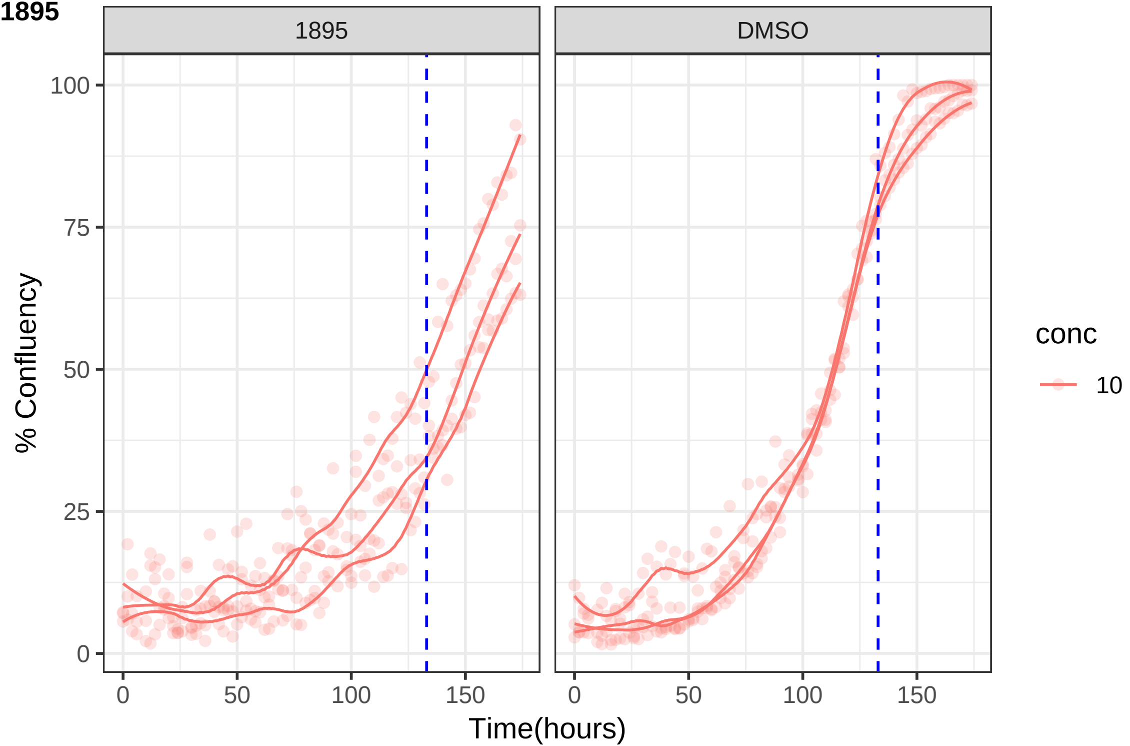

1895
DMSO
100
75
50
25
0
% Confluency
0
50
100
150
Time(hours)
conc
10

**1928**

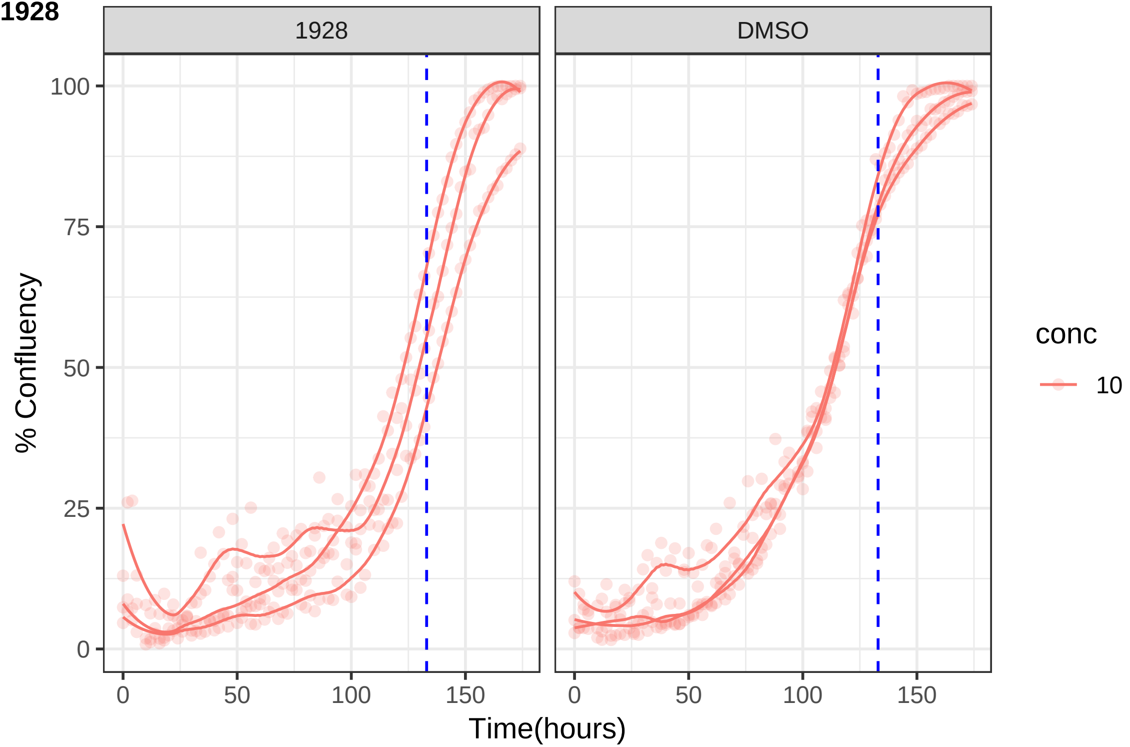
1928
DMSO
% Confluency
100
75
50
25
0
0
50
100
150
Time(hours)
conc
10

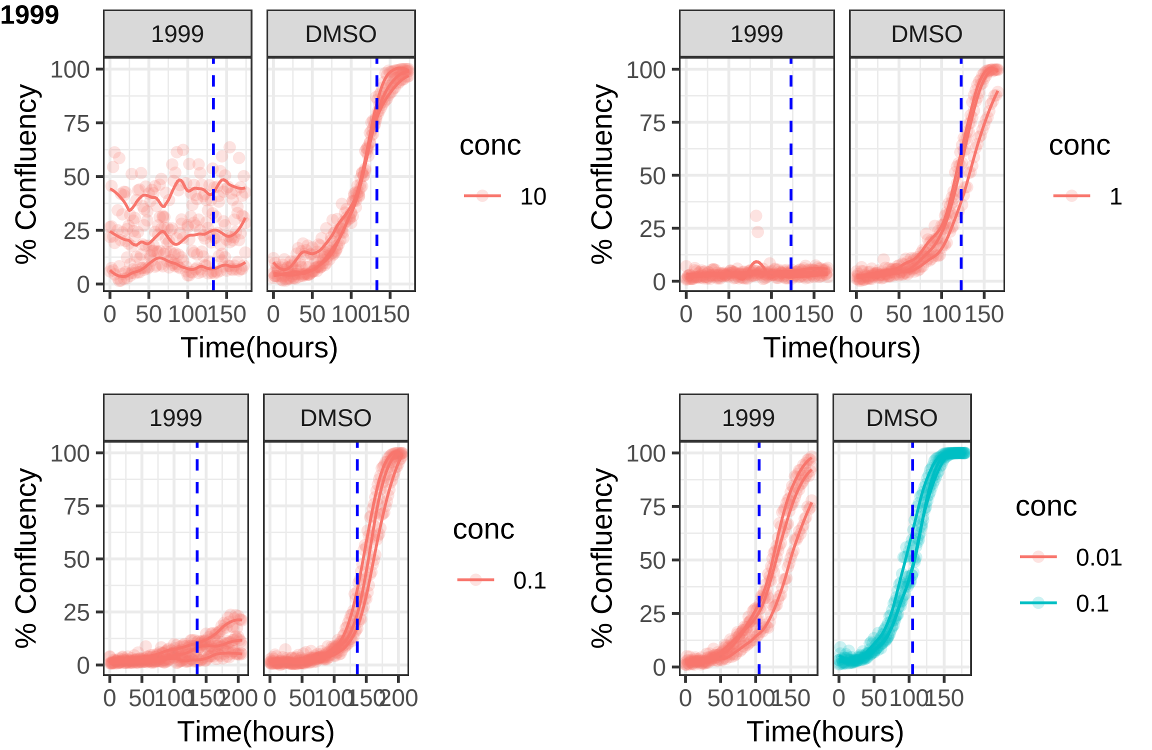

1999
1999
DMSO
% Confluency
100
75
50
25
0
0
50
100
150
Time(hours)
conc
10
1
0.1
0.01
200

**2008**

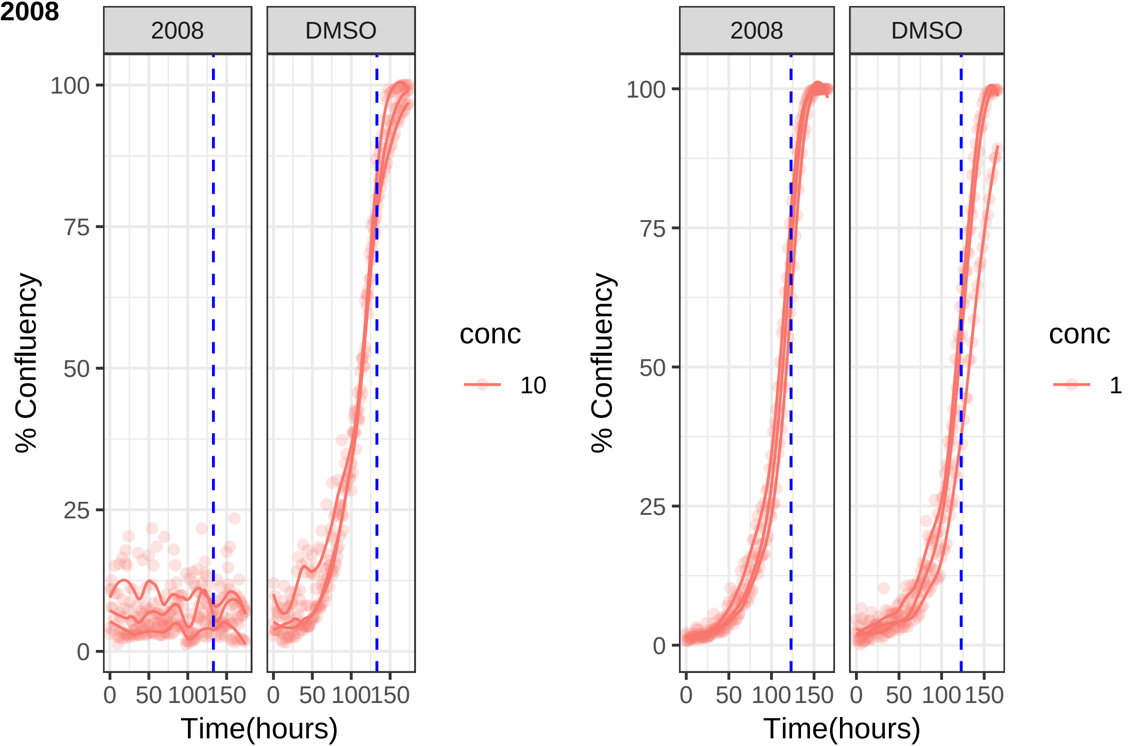

2008
DMSO
100
75
50
25
0
% Confluency
0
50
100
150
Time(hours)
conc
10
2008
DMSO
100
75
50
25
0
% Confluency
0
50
100
150
Time(hours)
conc
1

**2014**

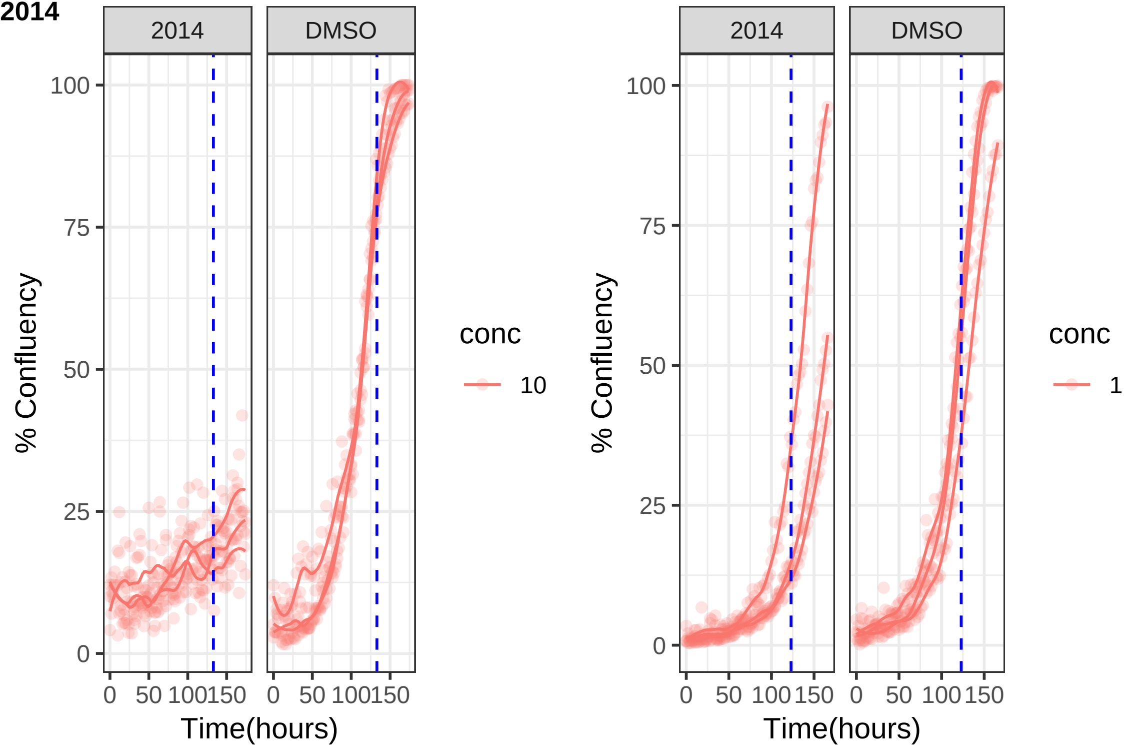

2014
DMSO
% Confluency
100
75
50
25
0
0
50
100
150
Time(hours)
conc
10
2014
DMSO
% Confluency
100
75
50
25
0
0
50
100
150
Time(hours)
conc
1

**2084**

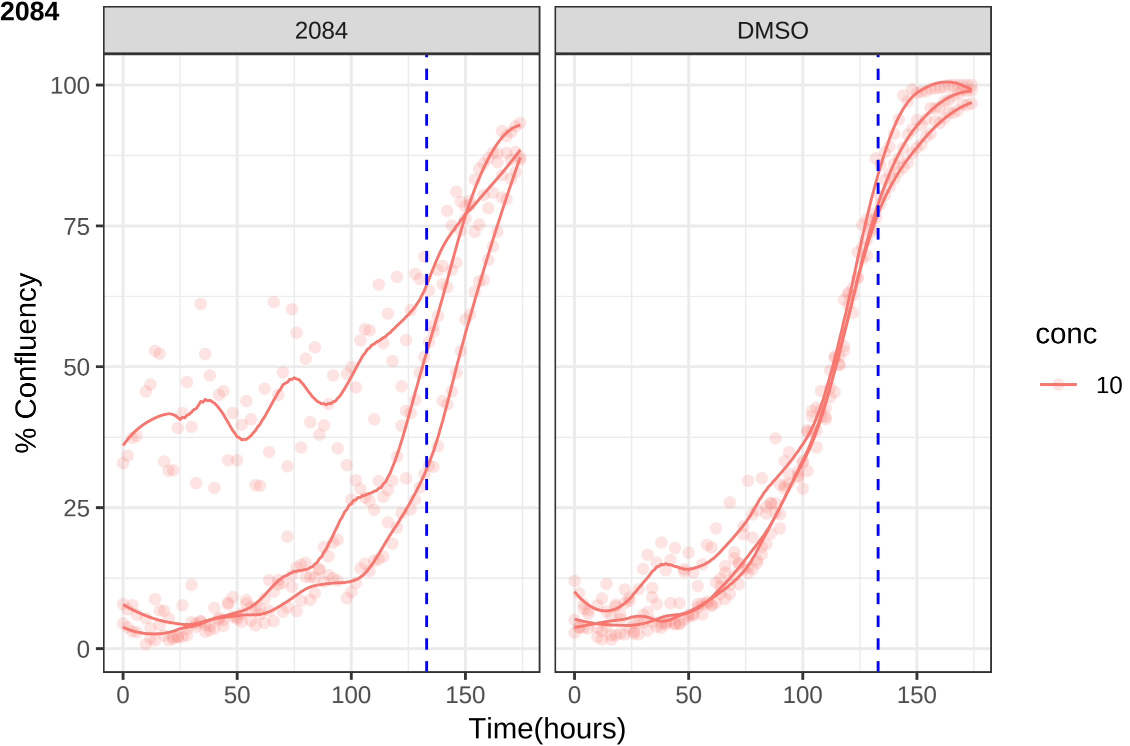
2084
DMSO
% Confluency
Time(hours)
0
25
50
75
100
0
50
100
150
conc
10

**2100**

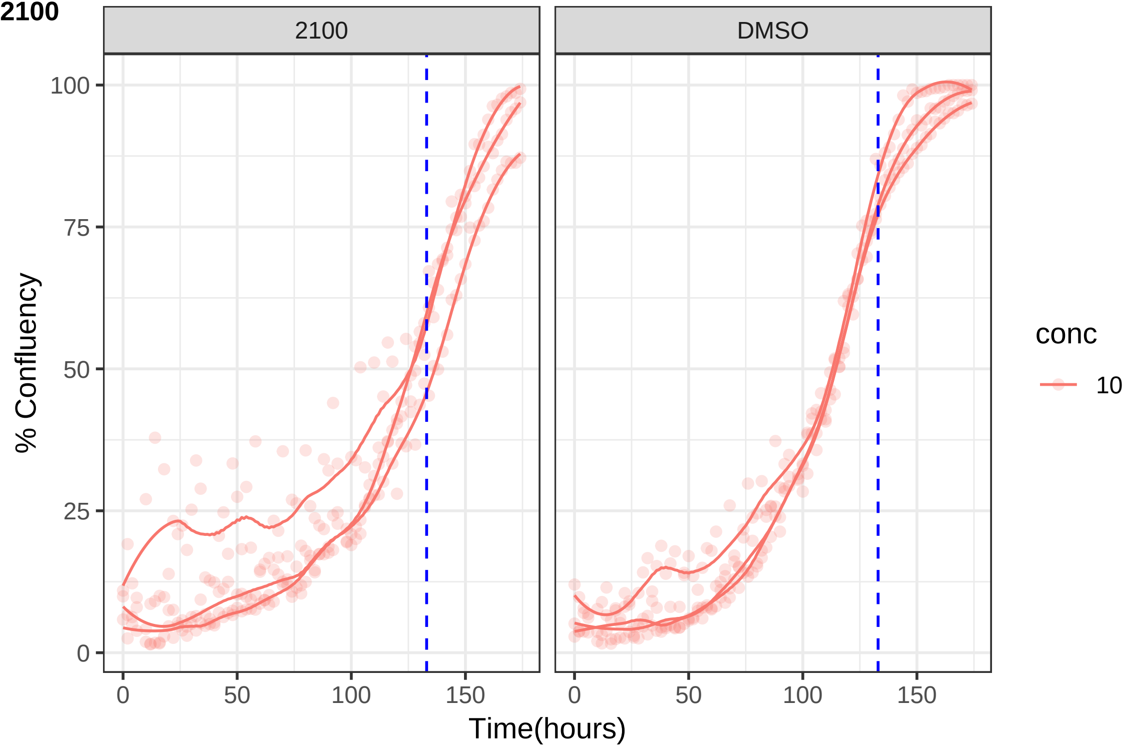

2100
DMSO
% Confluency
100
75
50
25
0
0
50
100
150
Time(hours)
conc
10

**2114**

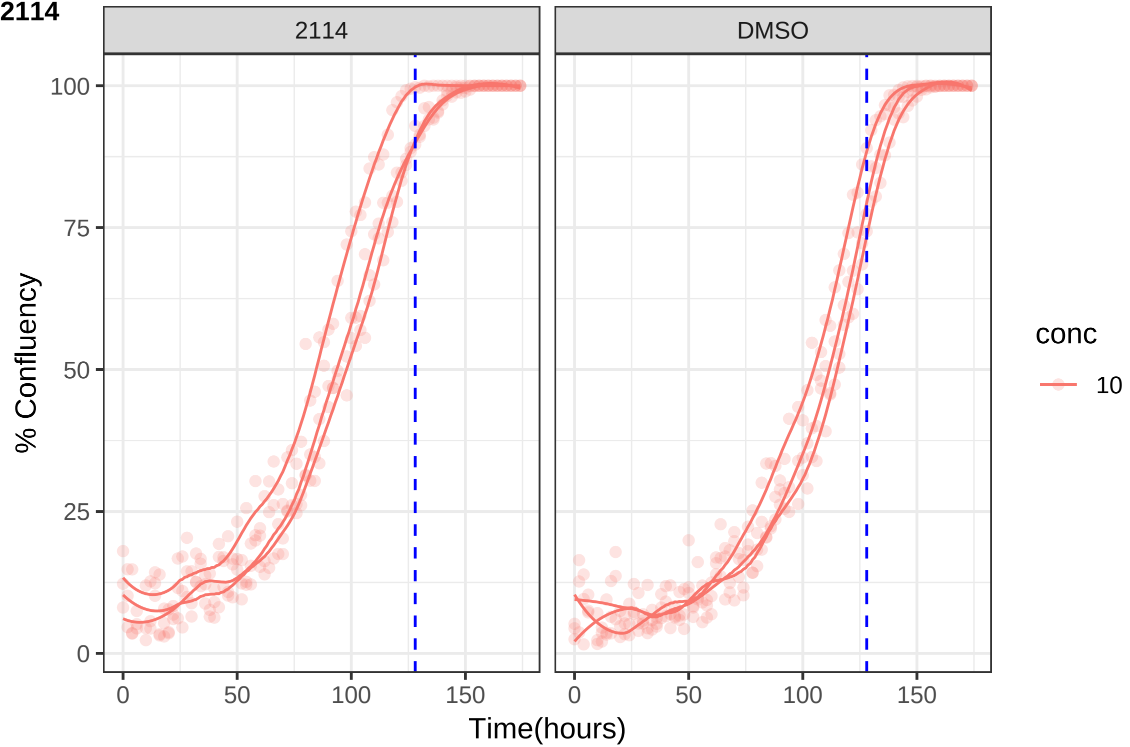
2114
DMSO
100
75
50
25
0
% Confluency
0
50
100
150
Time(hours)
conc
10

**2120**

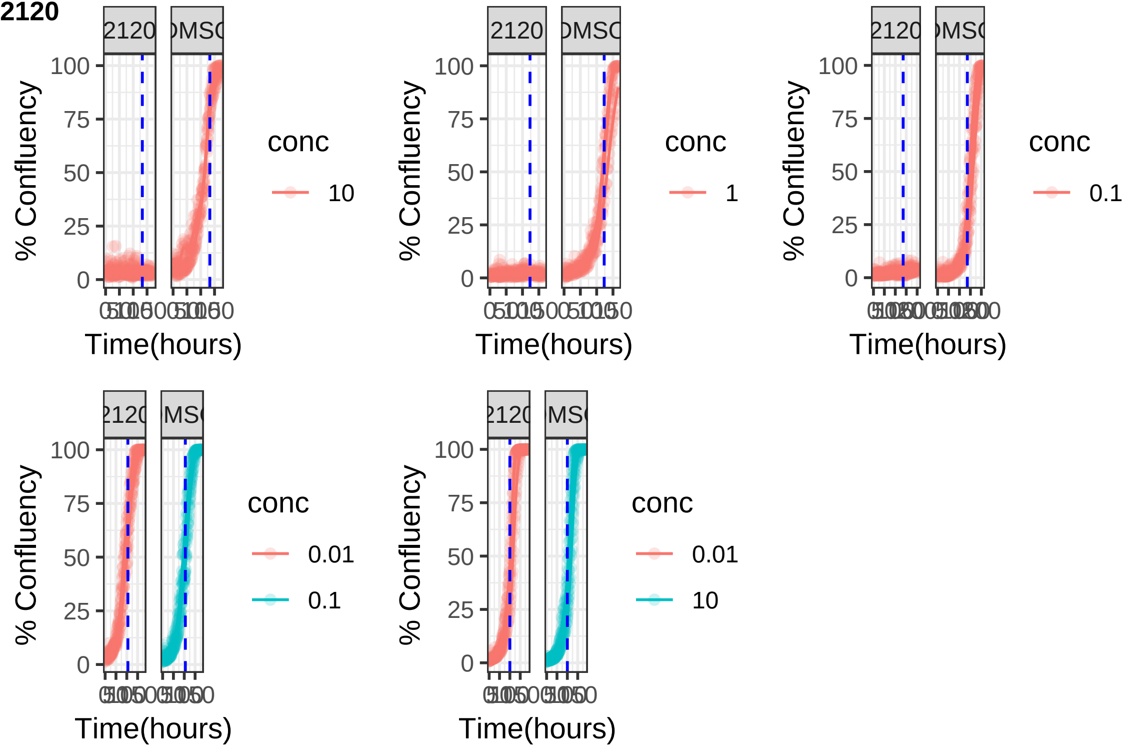

**2138**

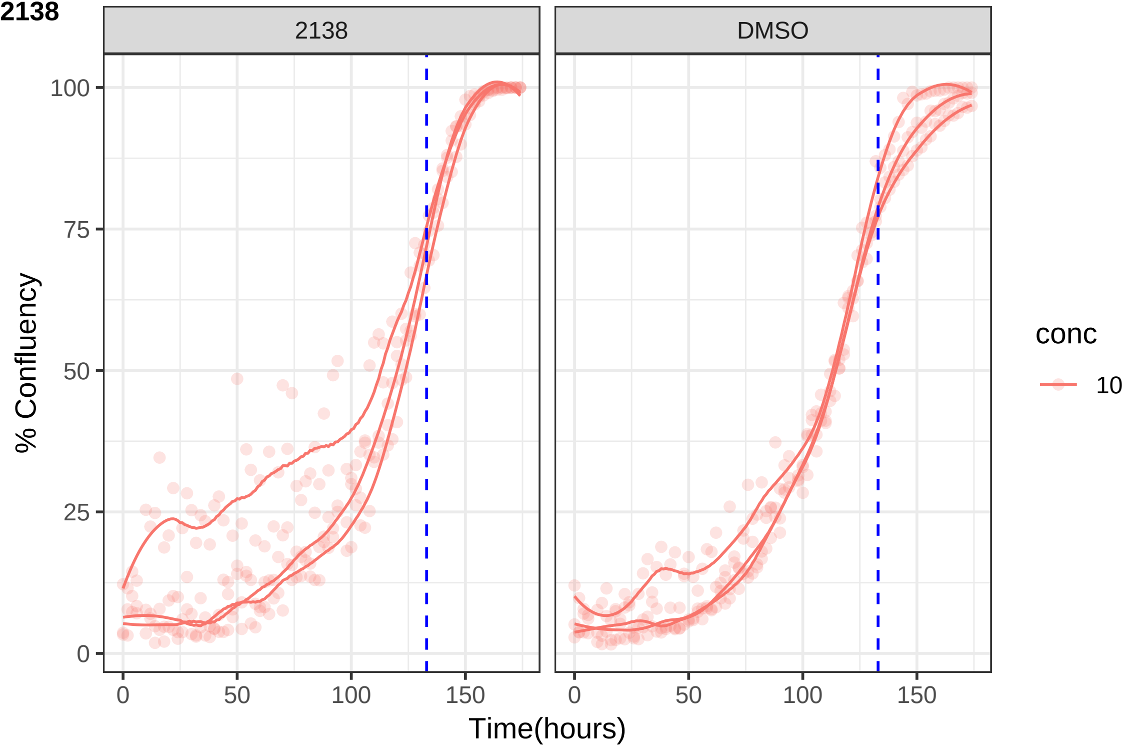

2138
DMSO
100
75
50
25
0
% Confluency
0
50
100
150
Time(hours)
conc
10

**2140**

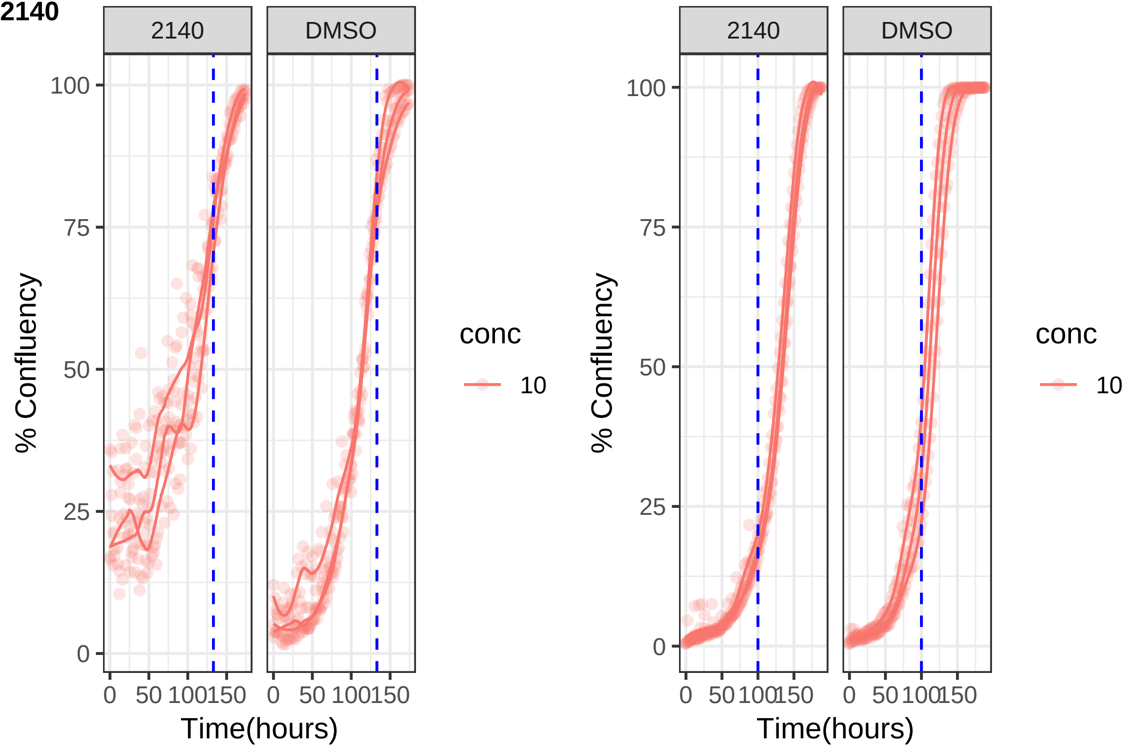

2140
DMSO
100
75
50
25
0
% Confluency
0 50 100 150
Time(hours)
conc
10
2140
DMSO
100
75
50
25
0
% Confluency
0 50 100 150
Time(hours)
conc
10

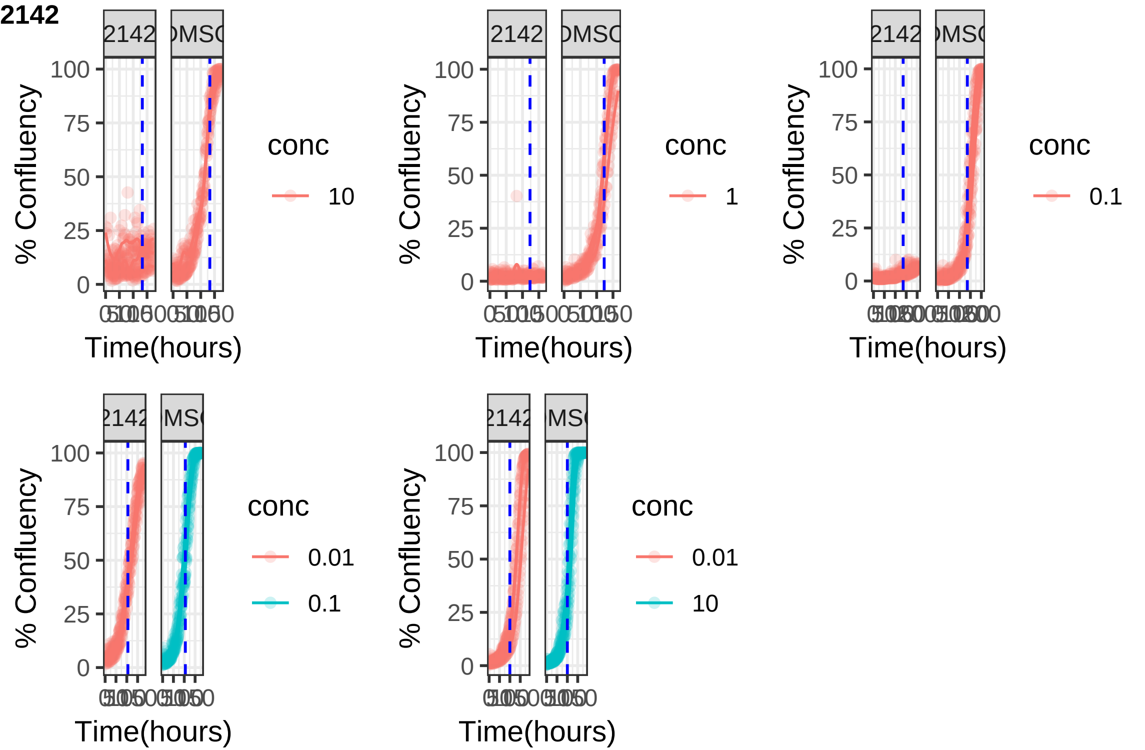
2142
2142
% Confluency
100
75
50
25
0
Time(hours)
conc
10
1
0.1
0.01
0.1
0.01
10

**2158**

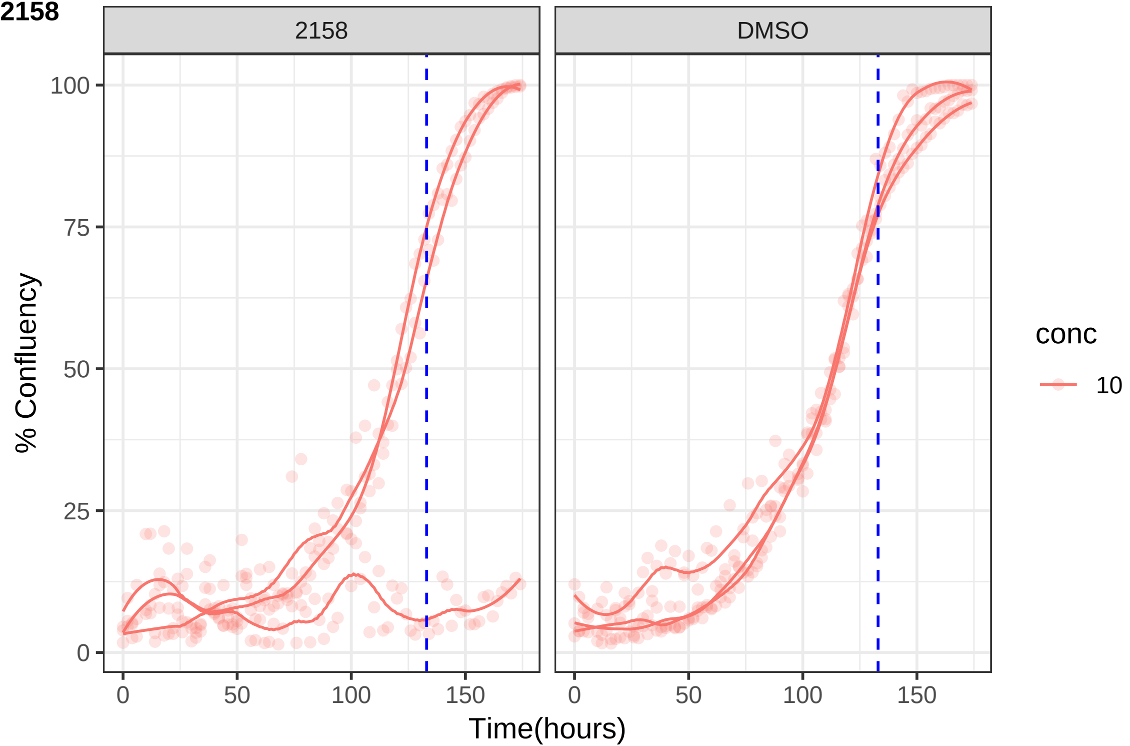
2158
DMSO
100
75
50
25
0
% Confluency
0
50
100
150
Time(hours)
conc
10

**2159**

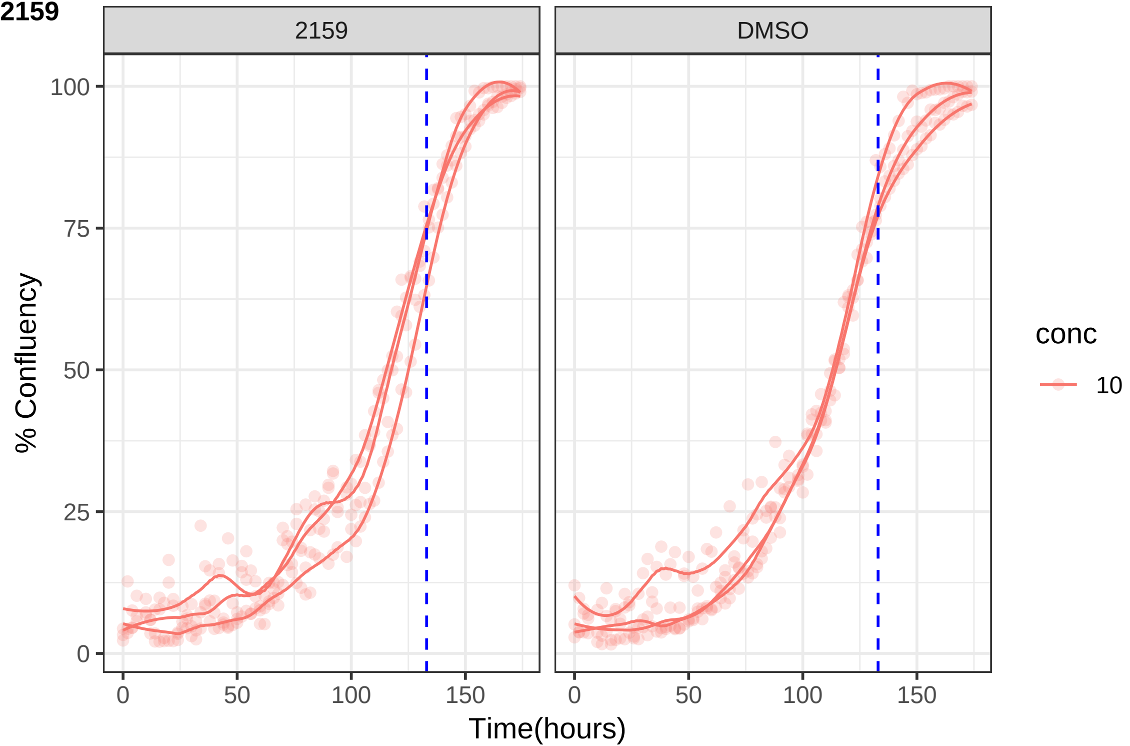
2159
DMSO
% Confluency
100
75
50
25
0
0
50
100
150
Time(hours)
conc
10

**2160**

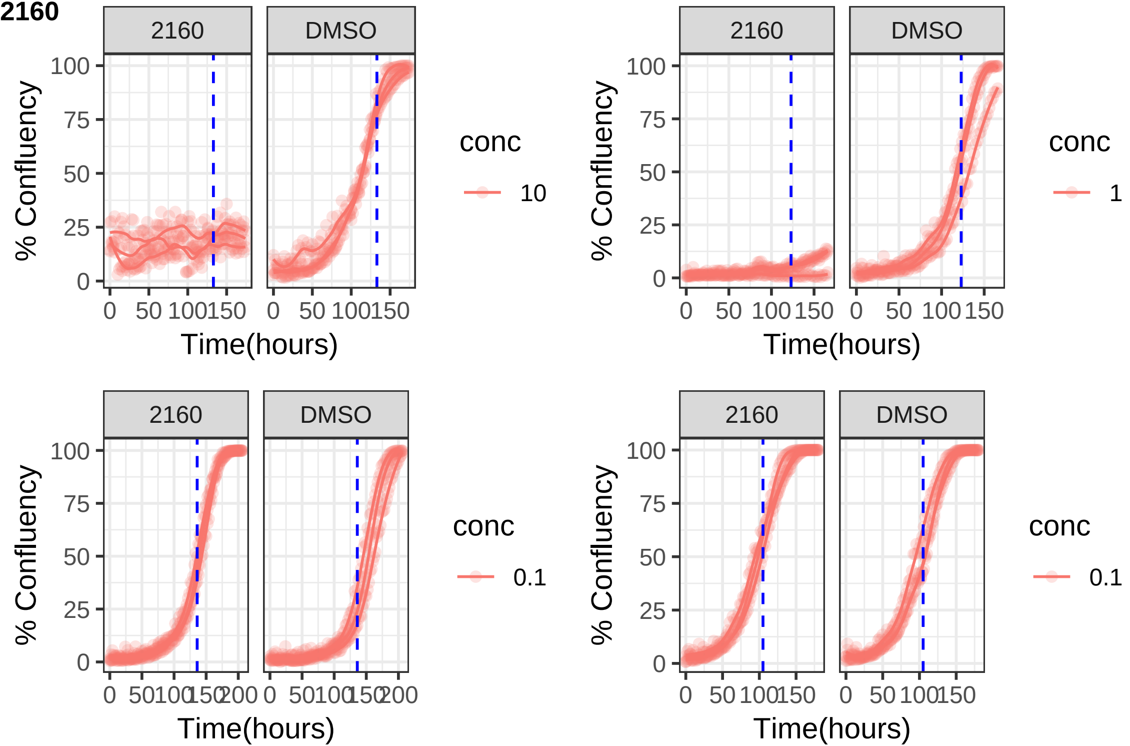
2160
DMSO
100
75
50
25
0
% Confluency
0
50
100
150
200
Time(hours)
conc
10
1
0.1

**2165**

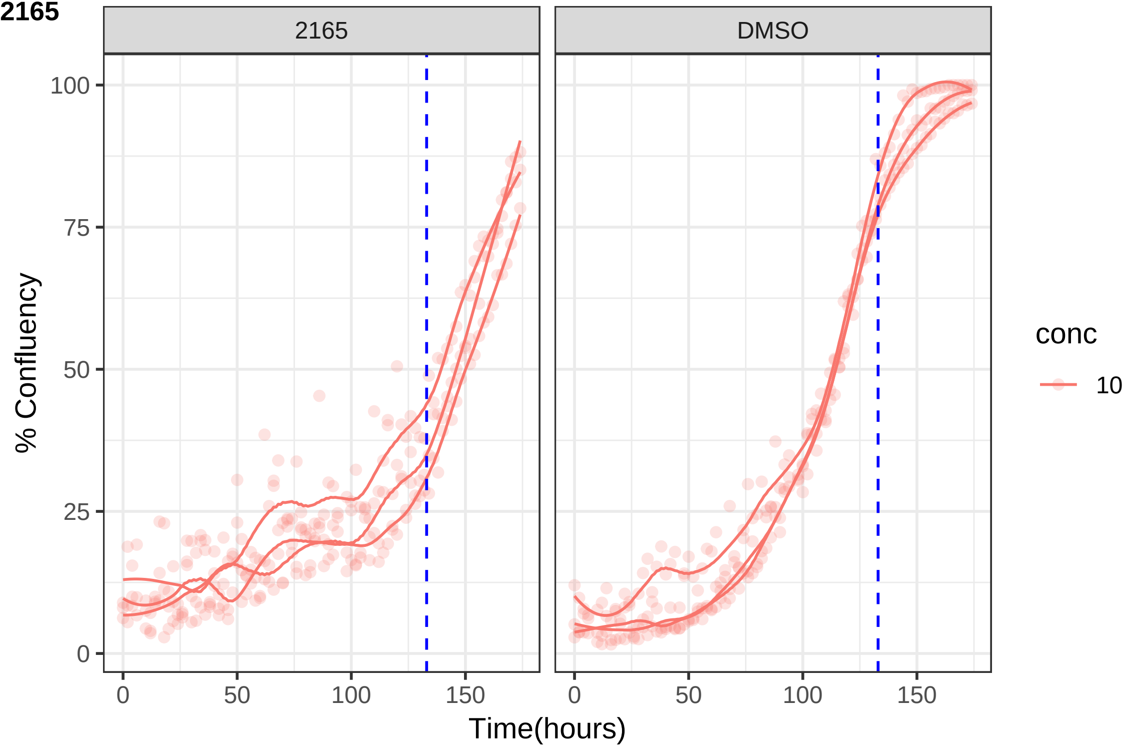

2165
DMSO
100
75
50
25
0
% Confluency
0
50
100
150
Time(hours)
conc
10

**2168**

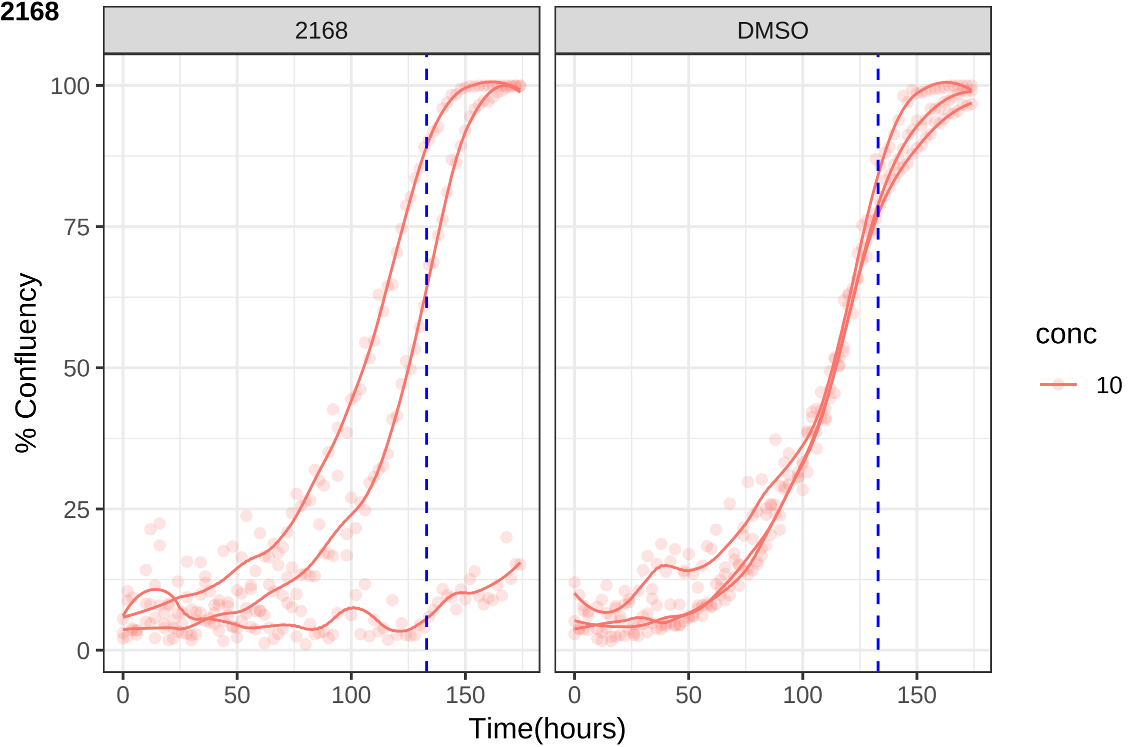
2168
DMSO
% Confluency
0
25
50
75
100
0
50
100
150
Time(hours)
conc
10

**2170**

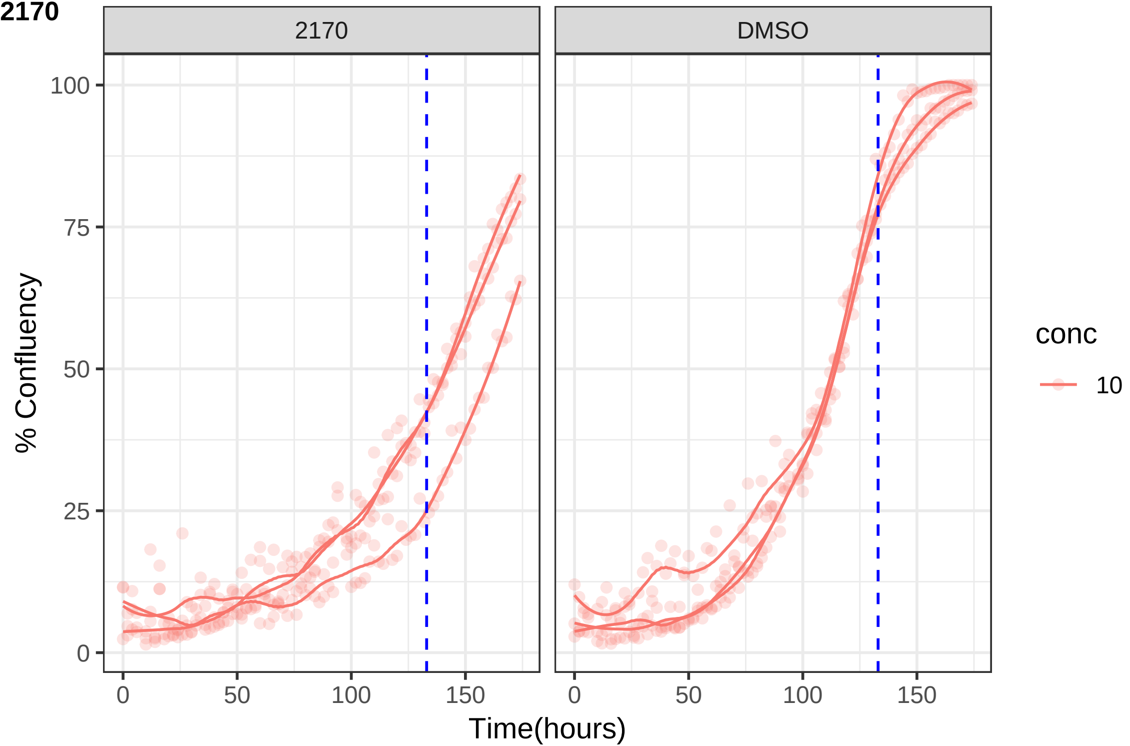

2170
DMSO
% Confluency
100
75
50
25
0
0
50
100
150
0
50
100
150
Time(hours)
conc
10

**2180**

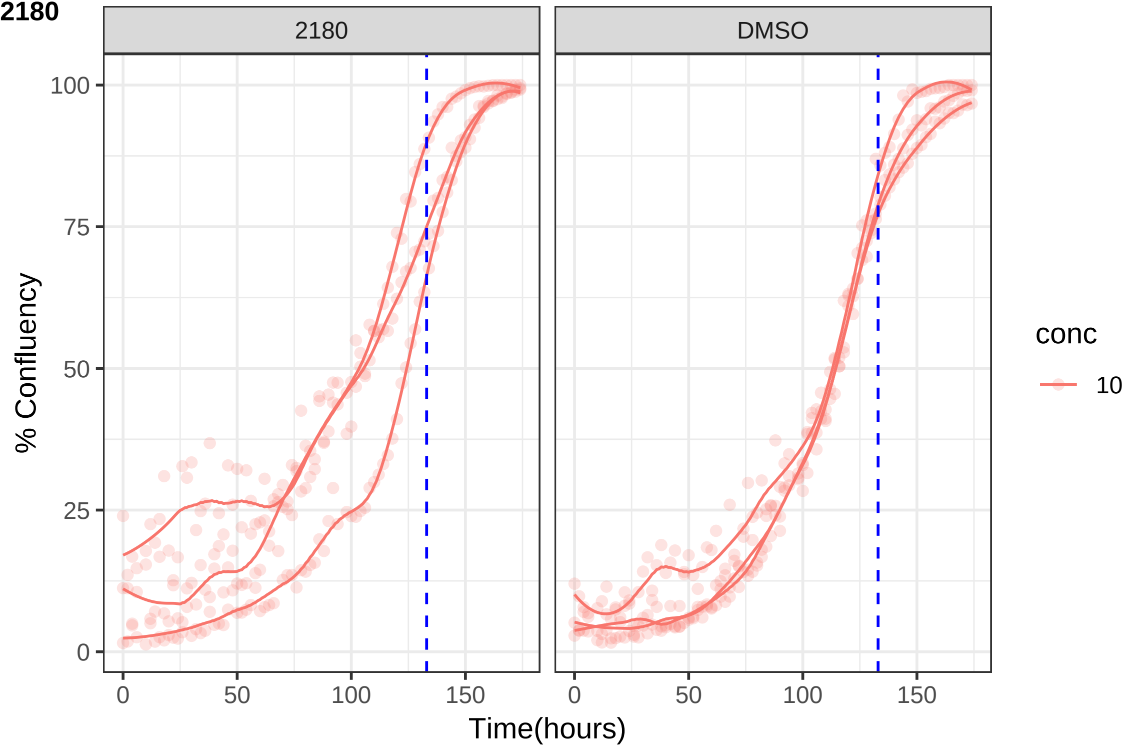
2180
DMSO
100
75
50
25
0
% Confluency
0
50
100
150
Time(hours)
conc
10

**2185**

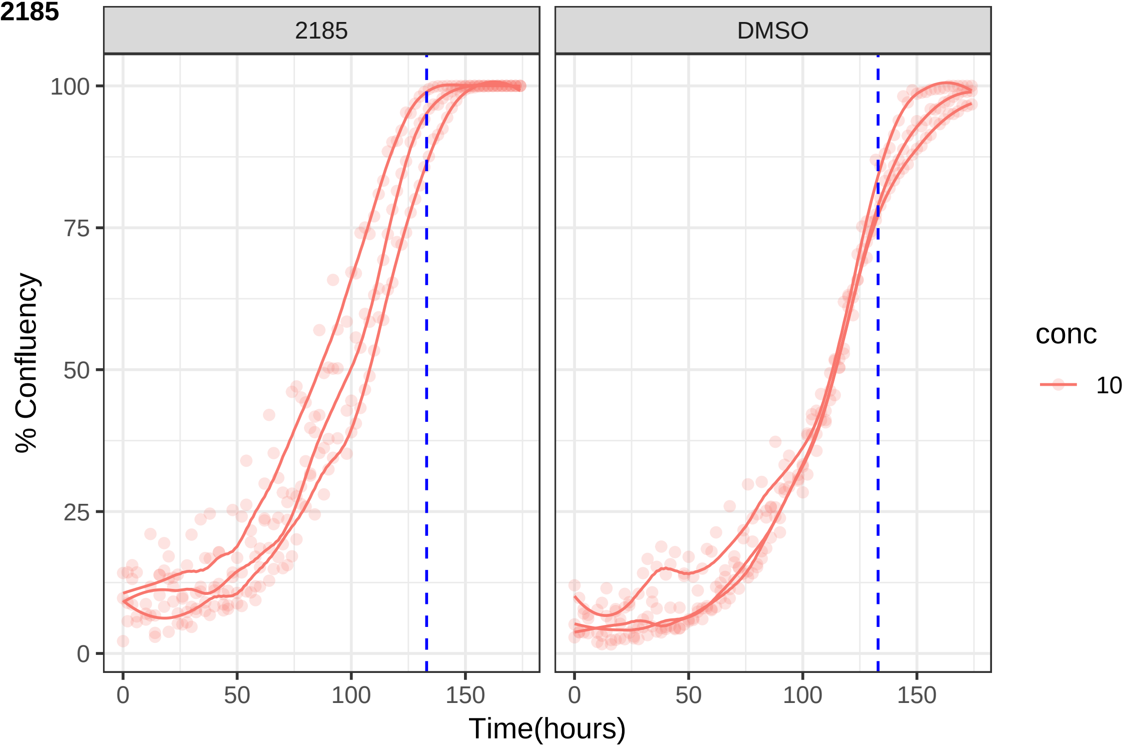
2185
DMSO
100
75
50
25
0
% Confluency
0
50
100
150
Time(hours)
conc
10

**2188**

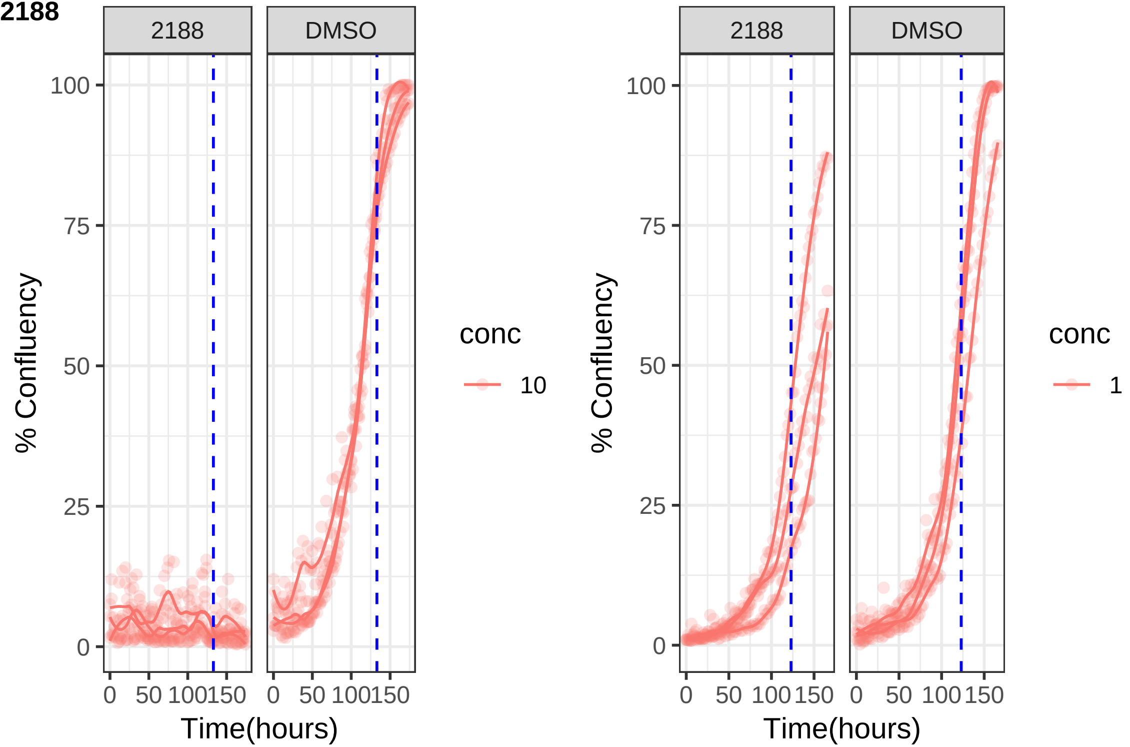

2188
DMSO
100
75
50
25
0
% Confluency
0
50
100
150
Time(hours)
conc
10
2188
DMSO
100
75
50
25
0
% Confluency
0
50
100
150
Time(hours)
conc
1

**2195**

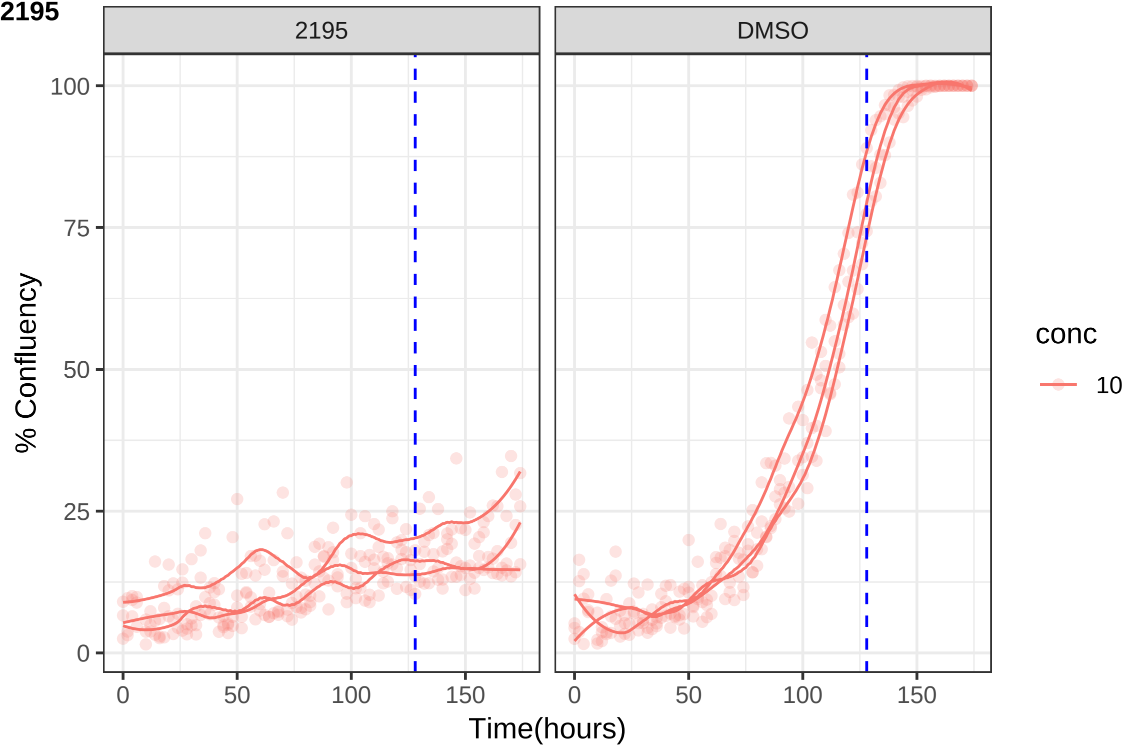
2195
DMSO
100
75
50
25
0
% Confluency
0
50
100
150
Time(hours)
conc
10

**2198**

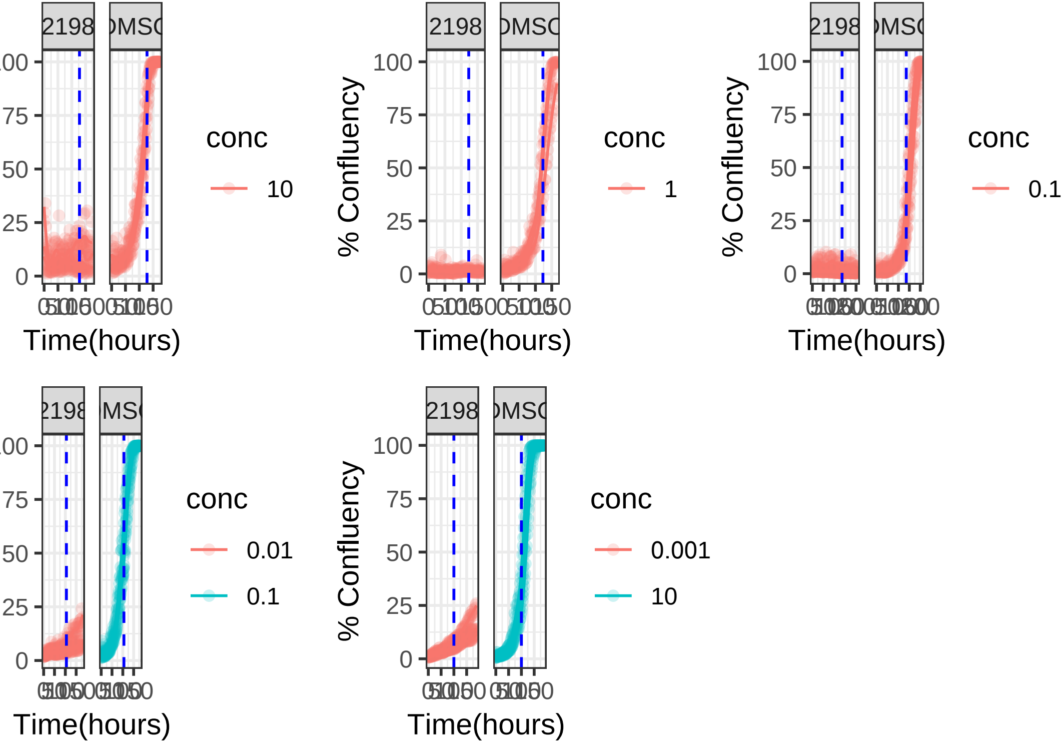
2198
DMSO
% Confluency
100
75
50
25
0
Time(hours)
conc
10
1
0.1
0.01
0.001

**2209**

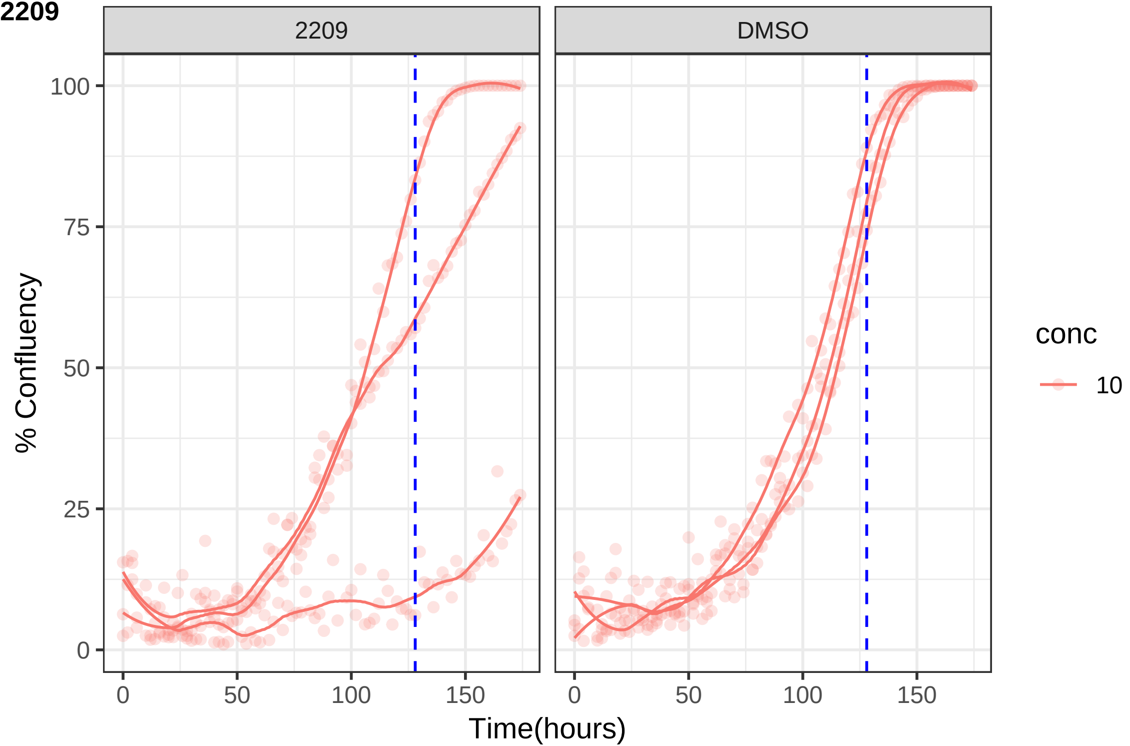
2209
DMSO
% Confluency
100
75
50
25
0
0
50
100
150
Time(hours)
conc
10

**2210**

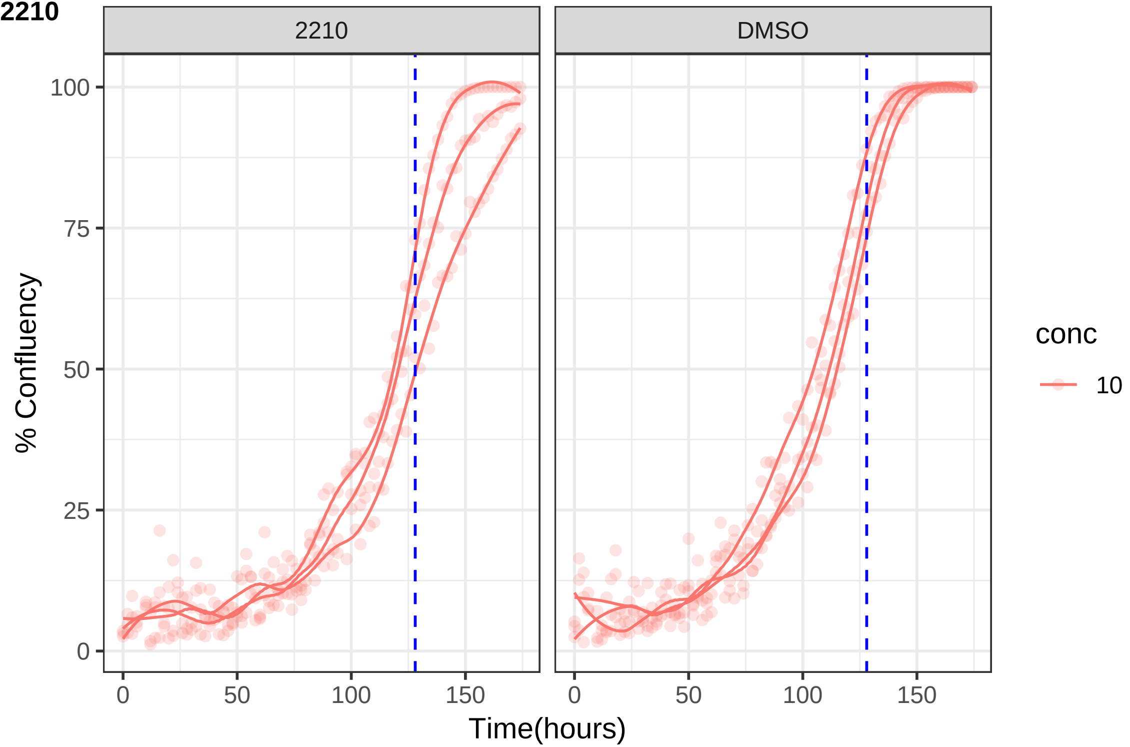
2210
DMSO
100
75
50
25
0
0
50
100
150
% Confluency
Time(hours)
conc
10

**2223**

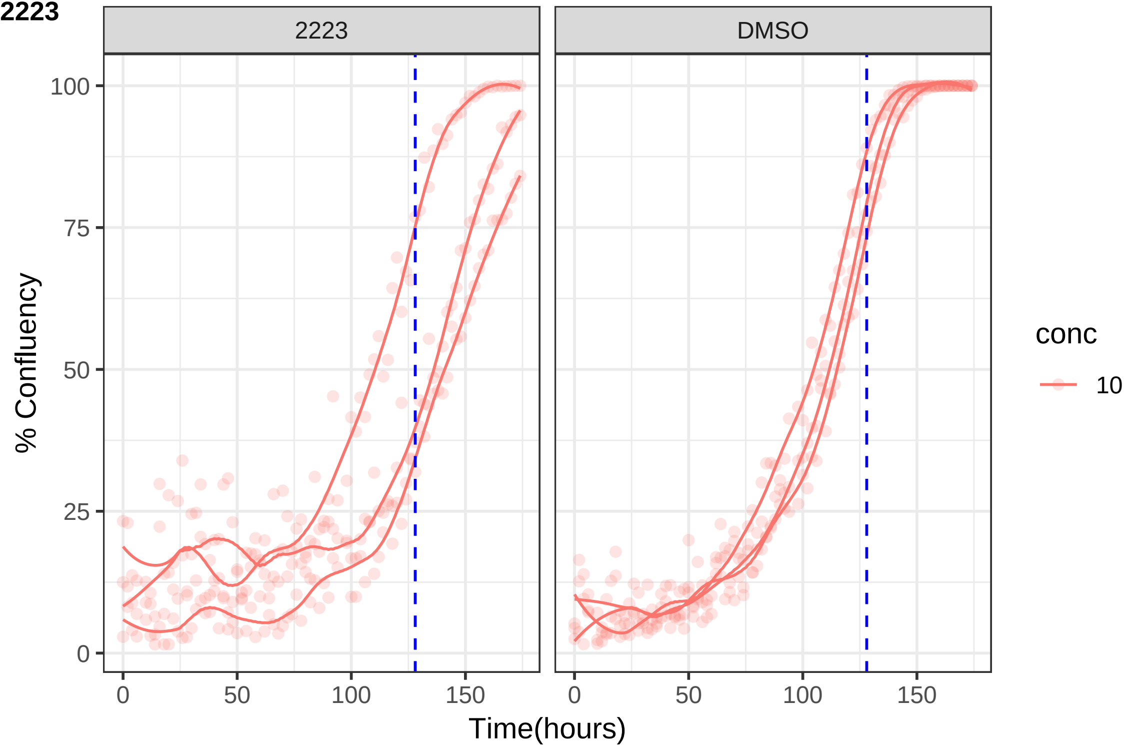
2223
DMSO
% Confluency
100
75
50
25
0
0
50
100
150
Time(hours)
conc
10

**2227**

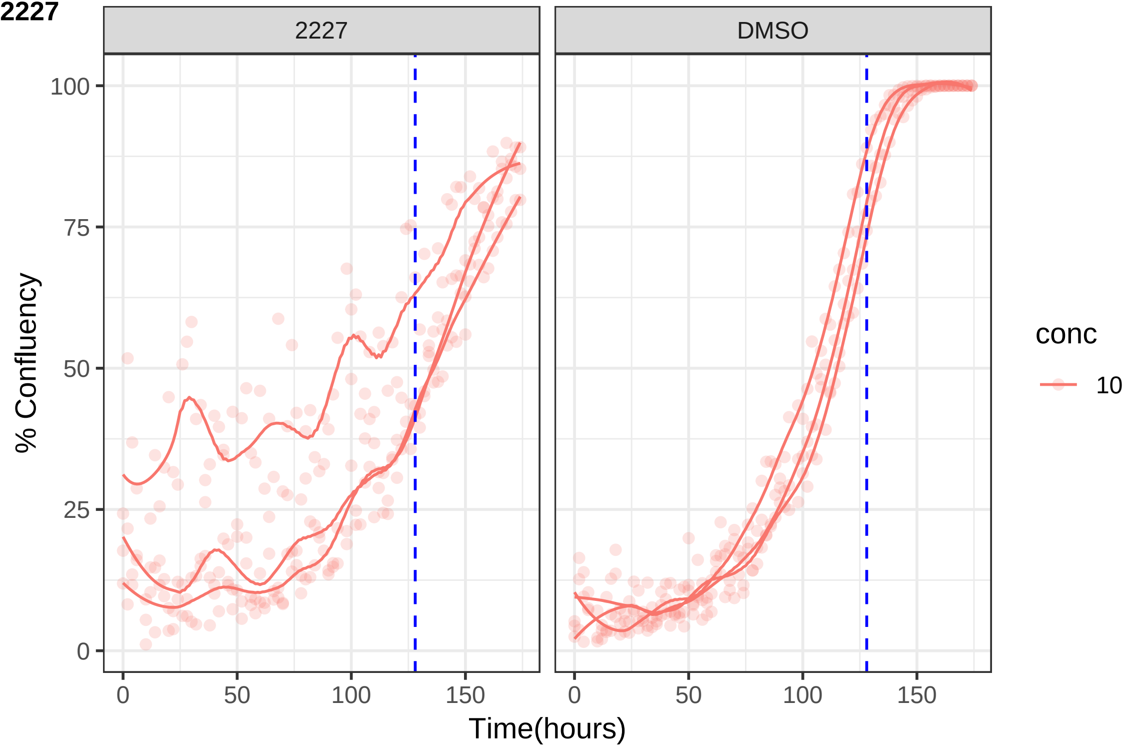

2227
DMSO
100
75
50
25
0
% Confluency
0
50
100
150
Time(hours)
conc
10

**2241**

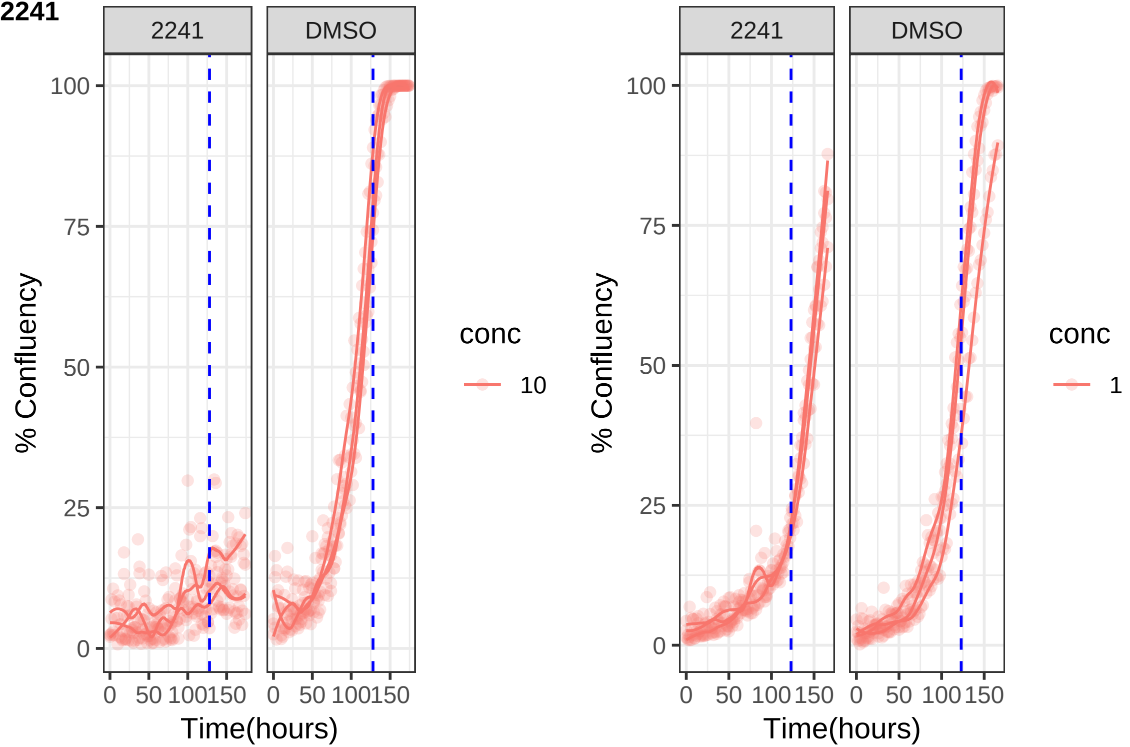
2241
DMSO
100
75
50
25
0
% Confluency
0 50 100 150
0 50 100 150
Time(hours)
conc
10
2241
DMSO
100
75
50
25
0
% Confluency
0 50 100 150
0 50 100 150
Time(hours)
conc
1

**2242**

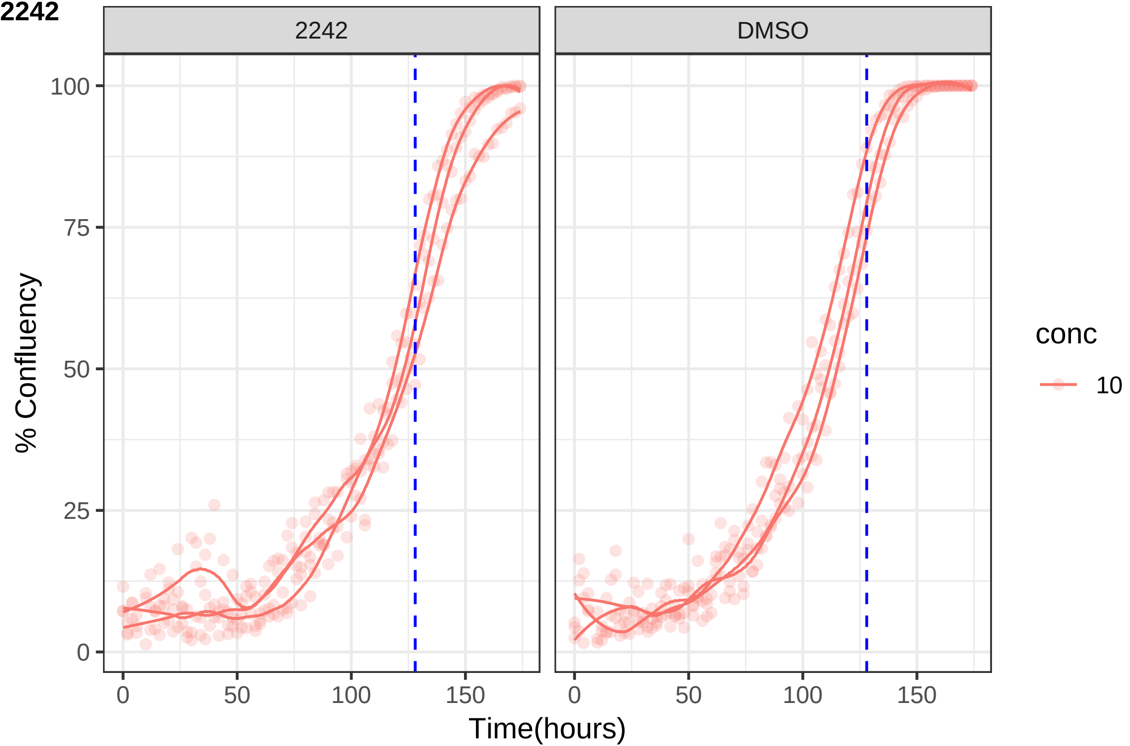

2242
DMSO
100
75
50
25
0
% Confluency
0
50
100
150
Time(hours)
conc
10

**2244**

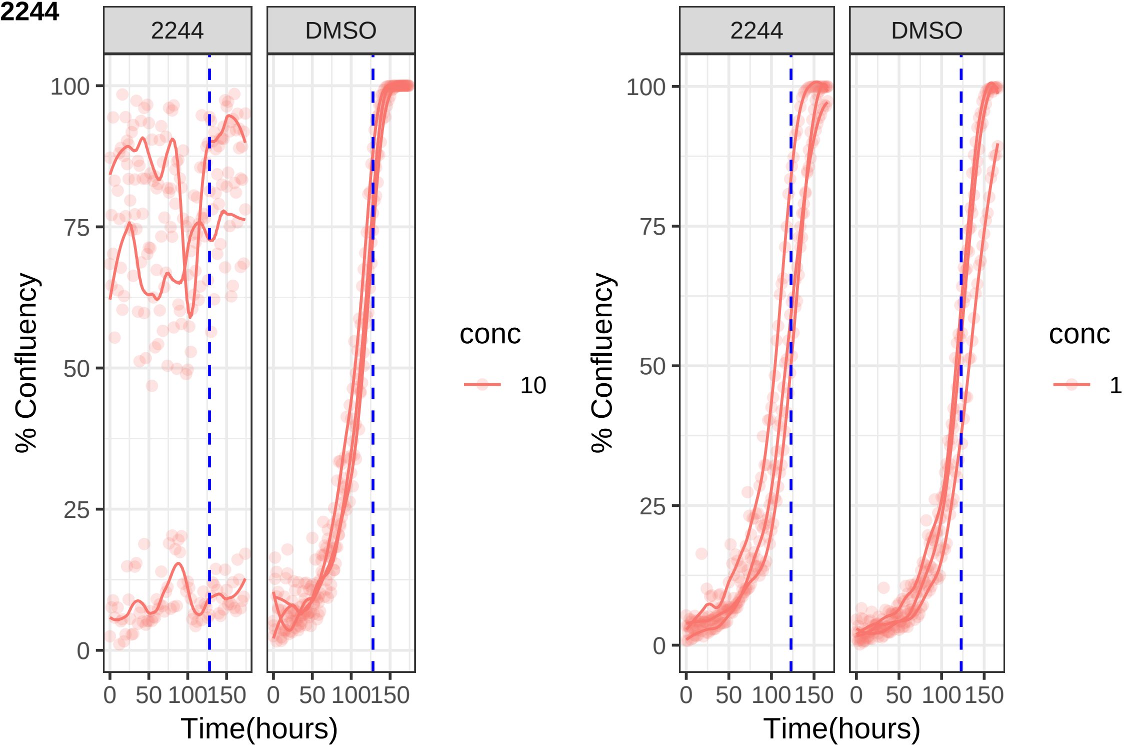

2244
DMSO
100
75
50
25
0
% Confluency
0
50
100
150
Time(hours)
conc
10
2244
DMSO
% Confluency
Time(hours)
conc
1

**2249**

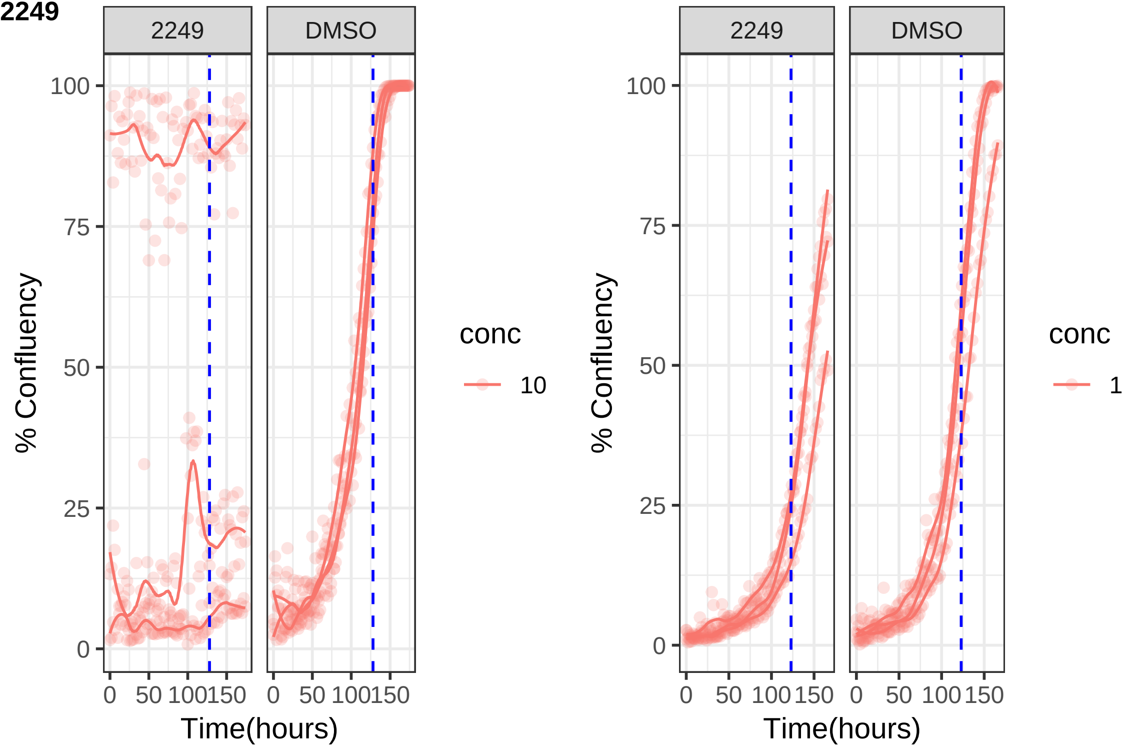
2249
DMSO
% Confluency
100
75
50
25
0
0
50
100
150
Time(hours)
conc
10
2249
DMSO
% Confluency
Time(hours)
conc
1

**2250**

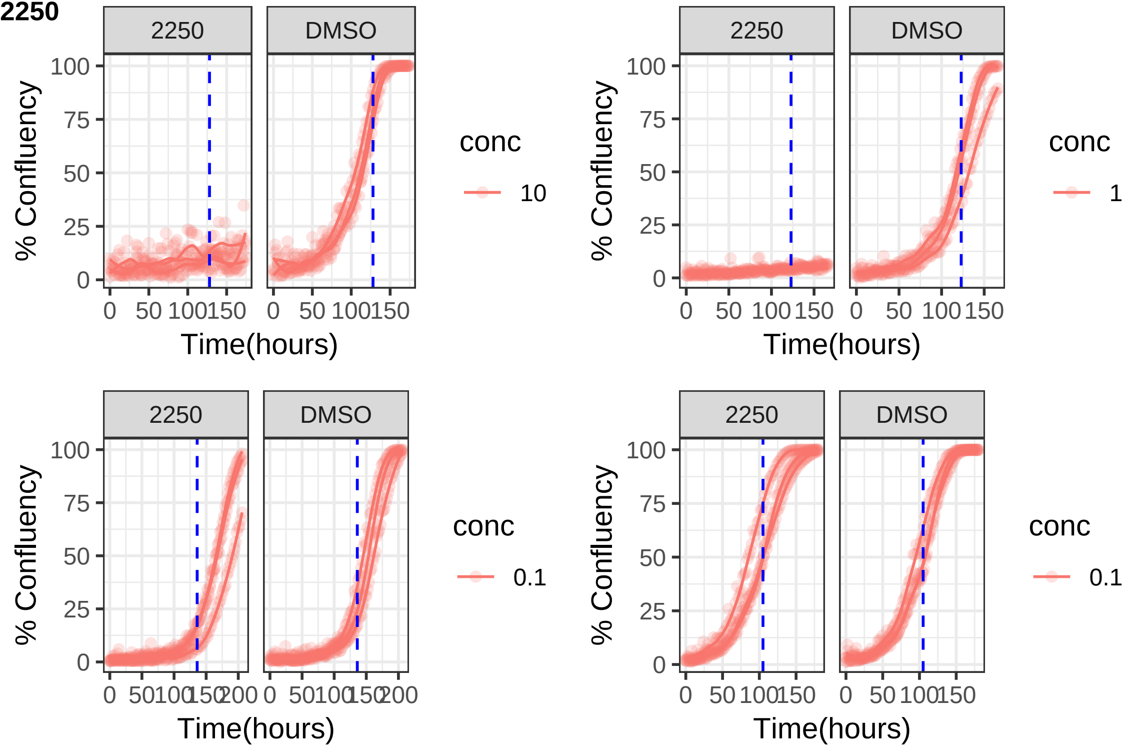

2250
DMSO
% Confluency
100
75
50
25
0
0 50 100 150
Time(hours)
conc
10
1
0.1
200

**2262**

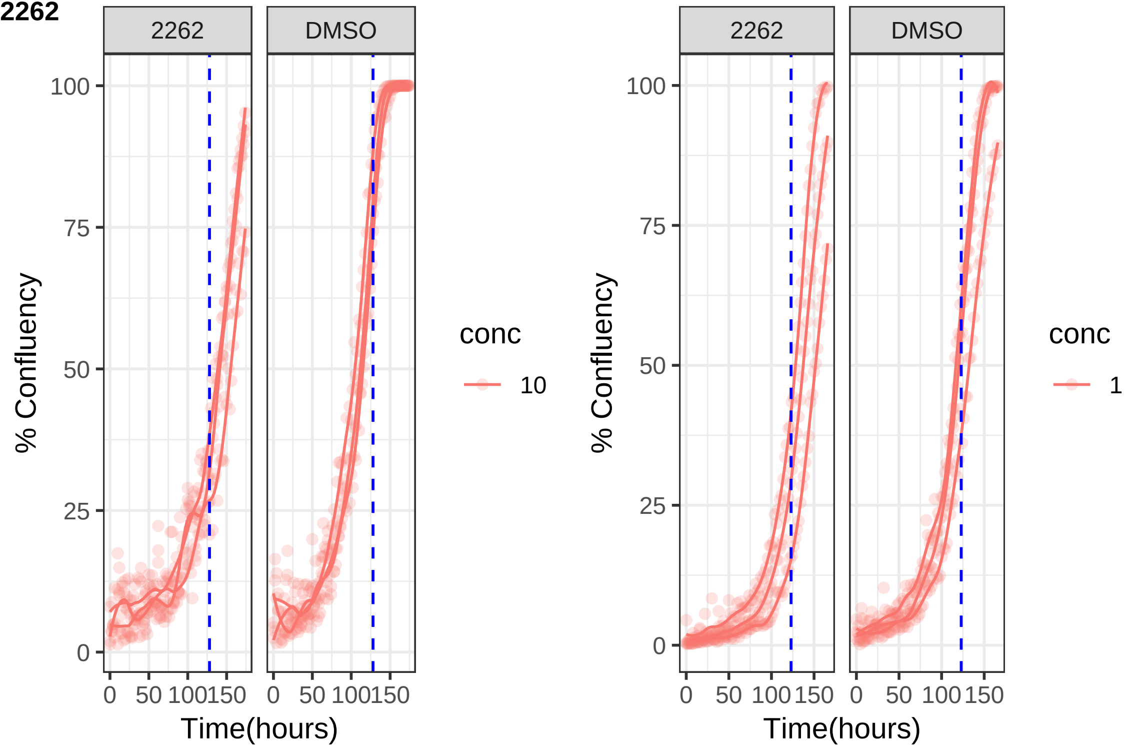
2262
DMSO
% Confluency
100
75
50
25
0
0
50
100
150
Time(hours)
conc
10
2262
DMSO
% Confluency
conc
1

**2268**

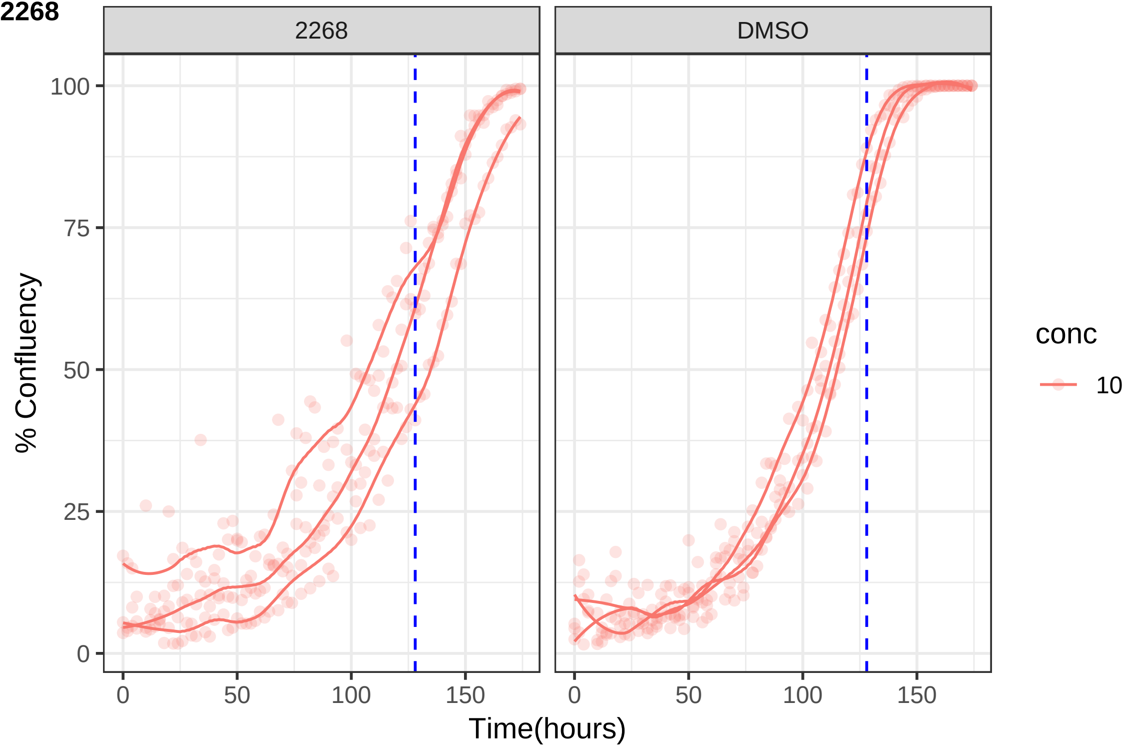

2268
DMSO
100
75
50
25
0
0
50
100
150
% Confluency
Time(hours)
conc
10

**2278**

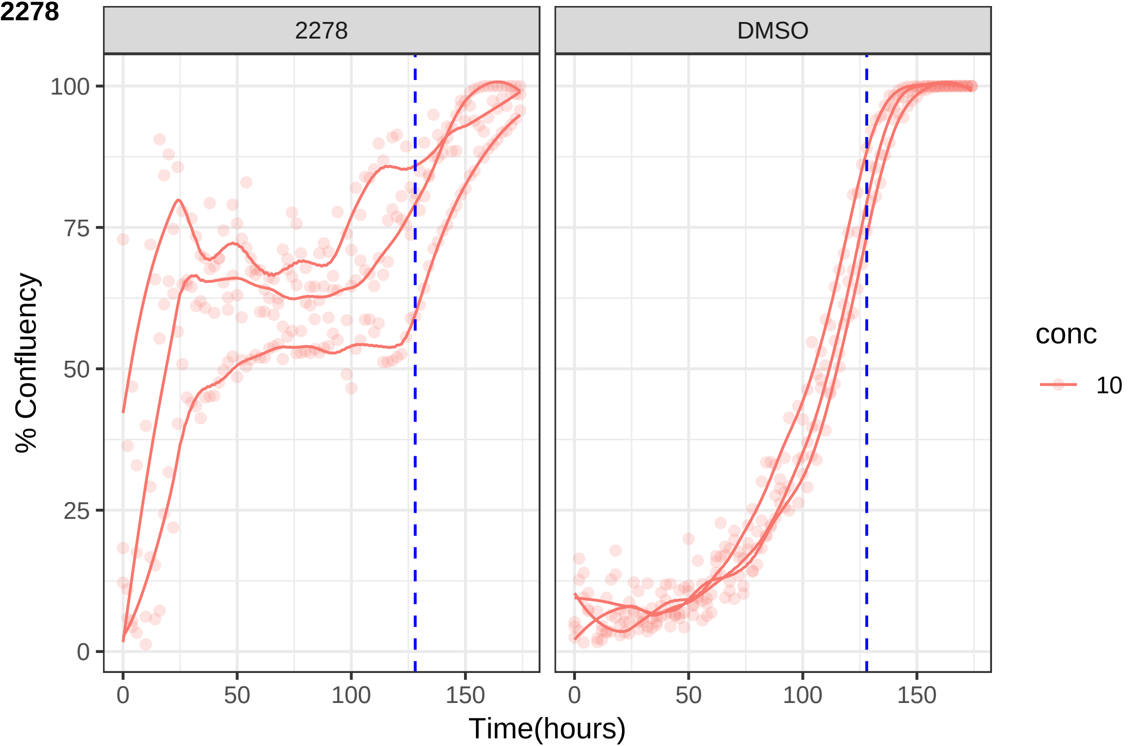

2278
DMSO
100
75
50
25
0
% Confluency
0
50
100
150
Time(hours)
conc
10

**2293**

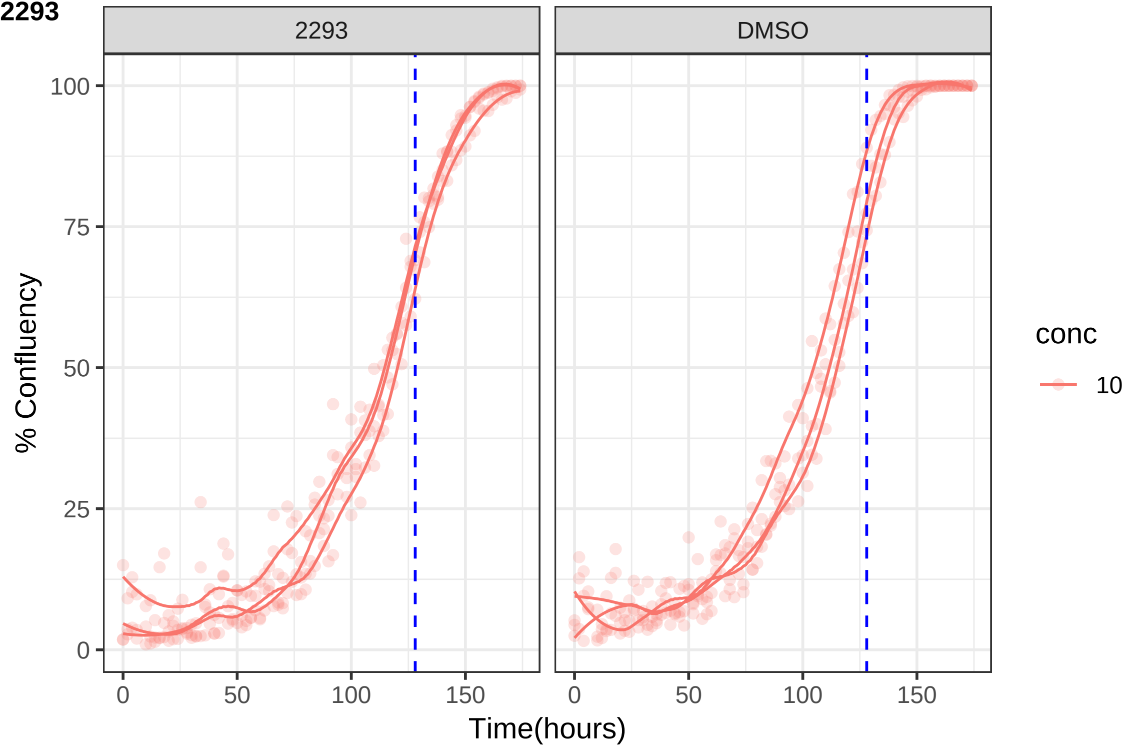
2293
DMSO
% Confluency
Time(hours)
0
25
50
75
100
0
50
100
150
conc
10

**2300**

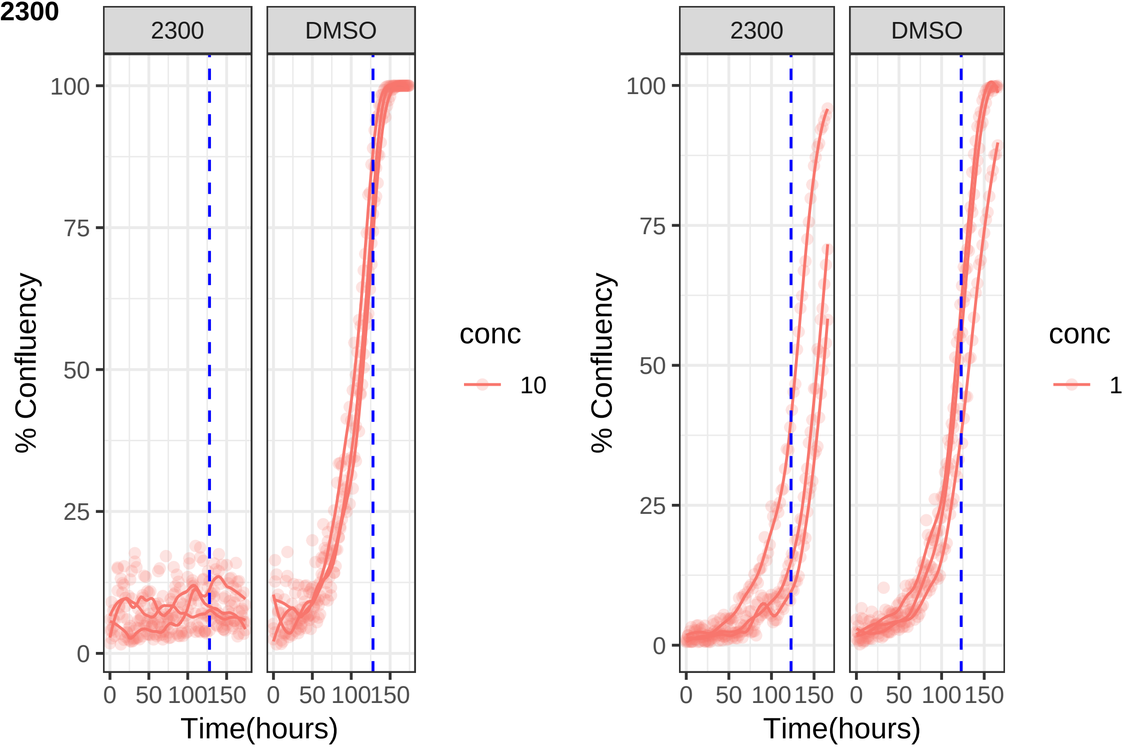
2300
DMSO
100
75
50
25
0
% Confluency
0
50
100
150
Time(hours)
conc
10
2300
DMSO
100
75
50
25
0
% Confluency
0
50
100
150
Time(hours)
conc
1

**2319**

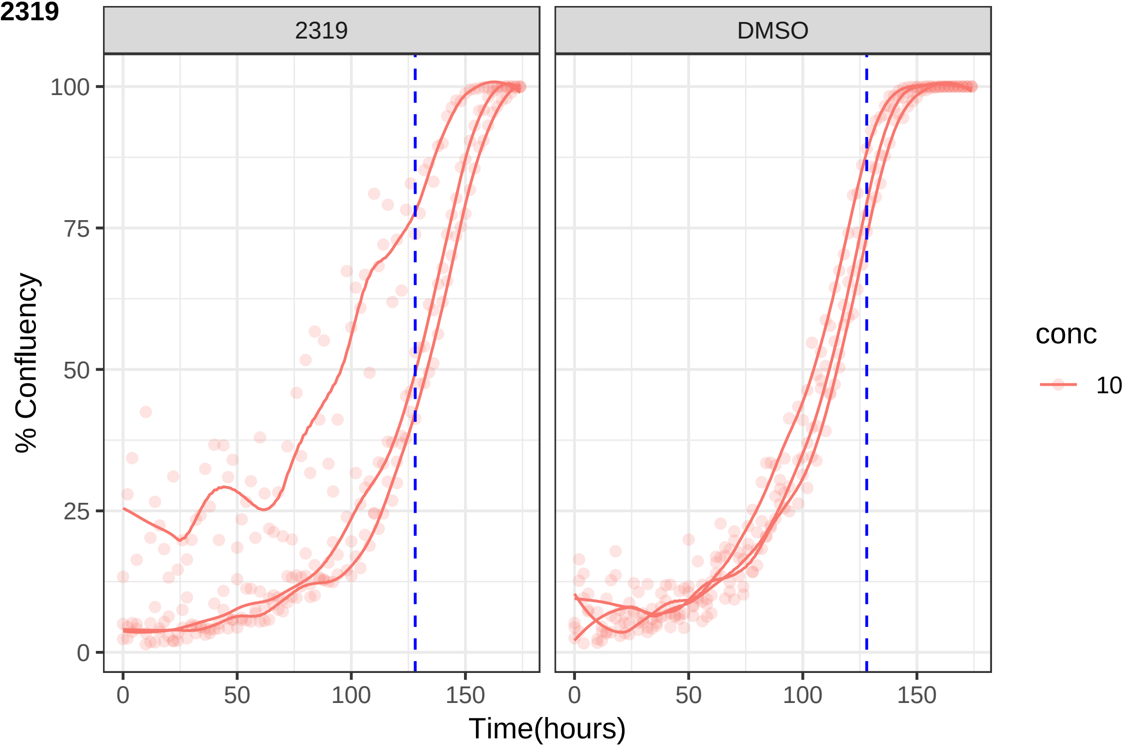
2319
DMSO
100
75
50
25
0
% Confluency
0
50
100
150
Time(hours)
conc
10

**2323**

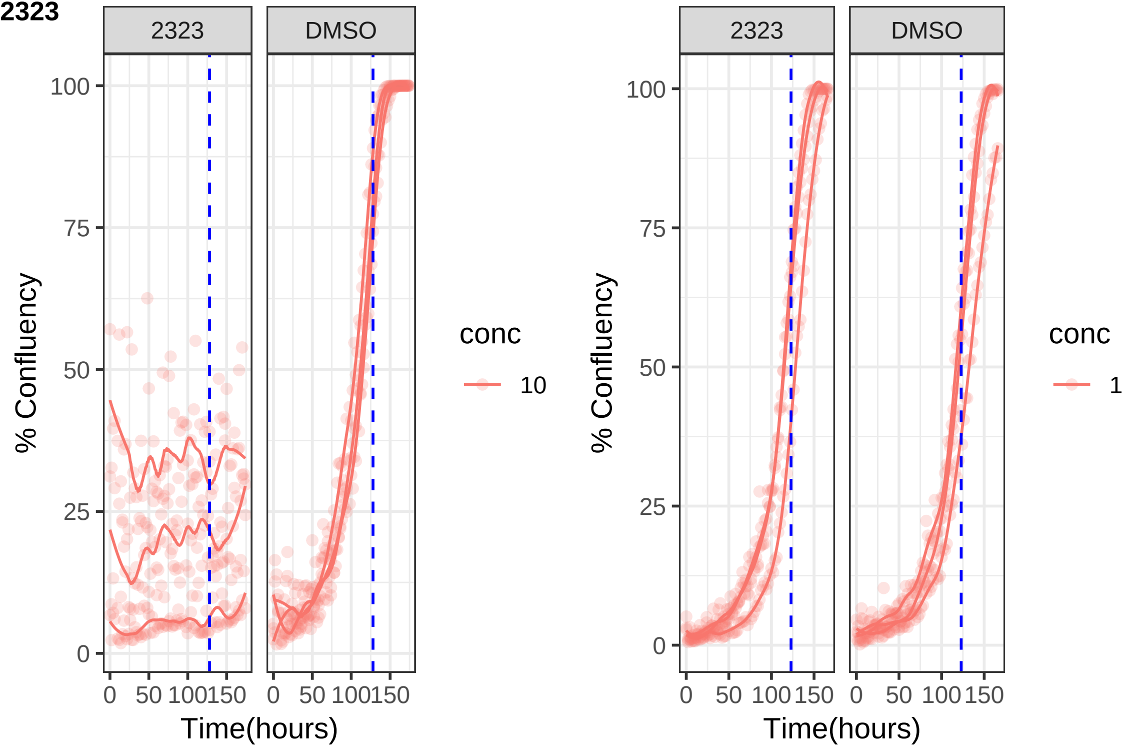
2323
DMSO
100
75
50
25
0
% Confluency
0
50
100
150
Time(hours)
conc
10
2323
DMSO
100
75
50
25
0
% Confluency
0
50
100
150
Time(hours)
conc
1

**2325**

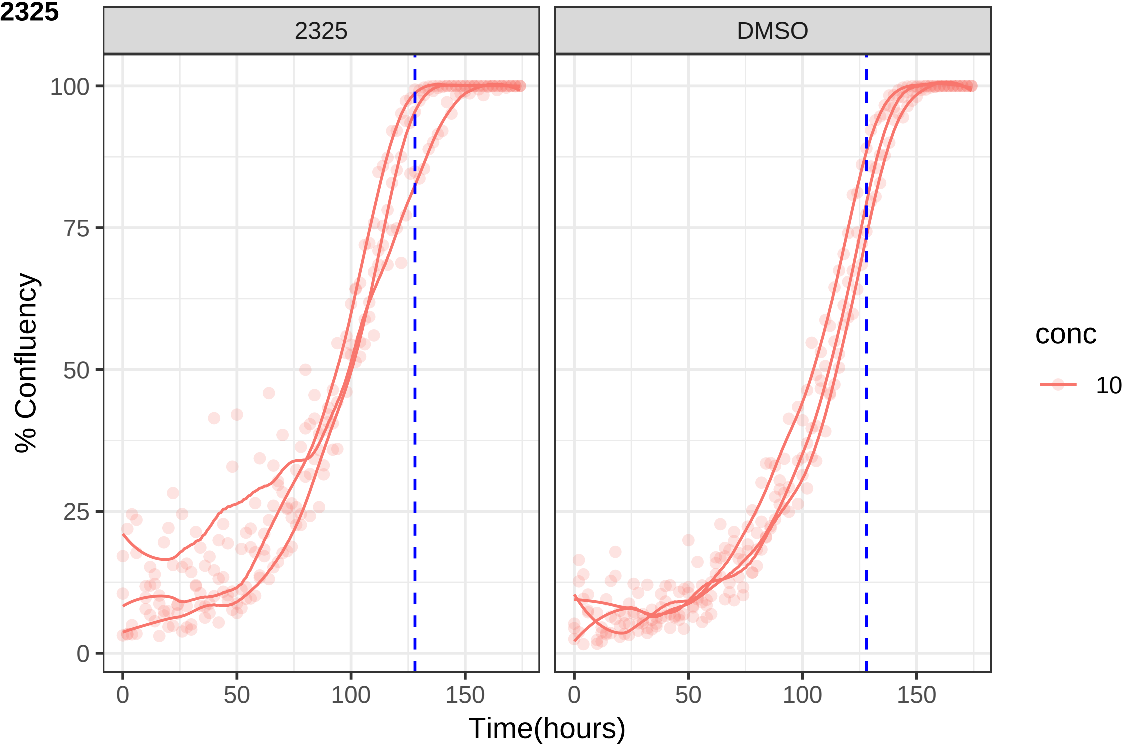

2325
DMSO
100
75
50
25
0
% Confluency
0
50
100
150
Time(hours)
conc
10

**2326**

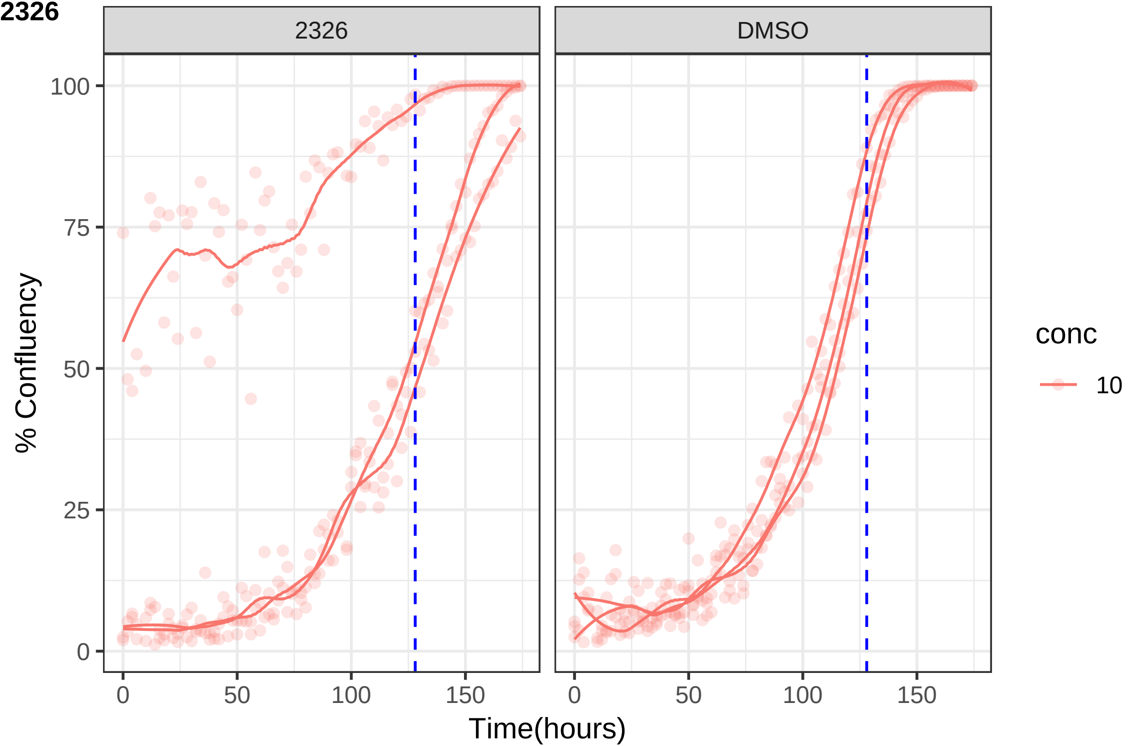

2326
DMSO
100
75
50
25
0
% Confluency
0
50
100
150
Time(hours)
conc
10

**2327**

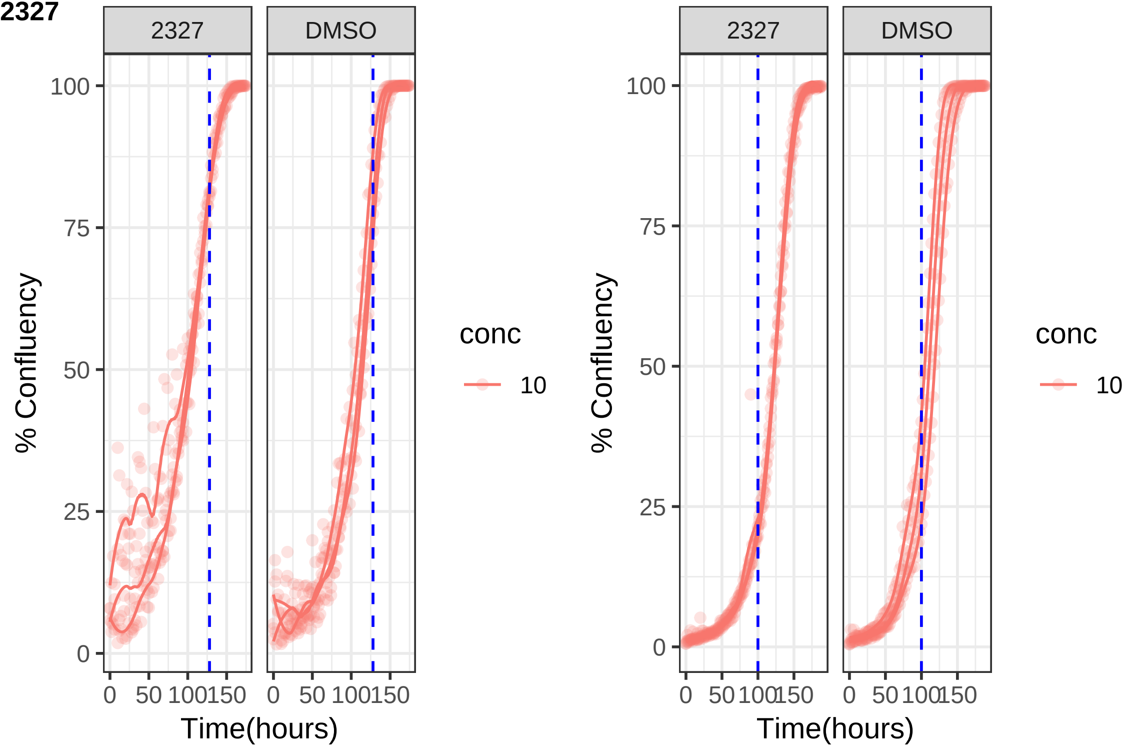

2327
DMSO
100
75
50
25
0
% Confluency
0
50
100
150
Time(hours)
conc
10

**2339**

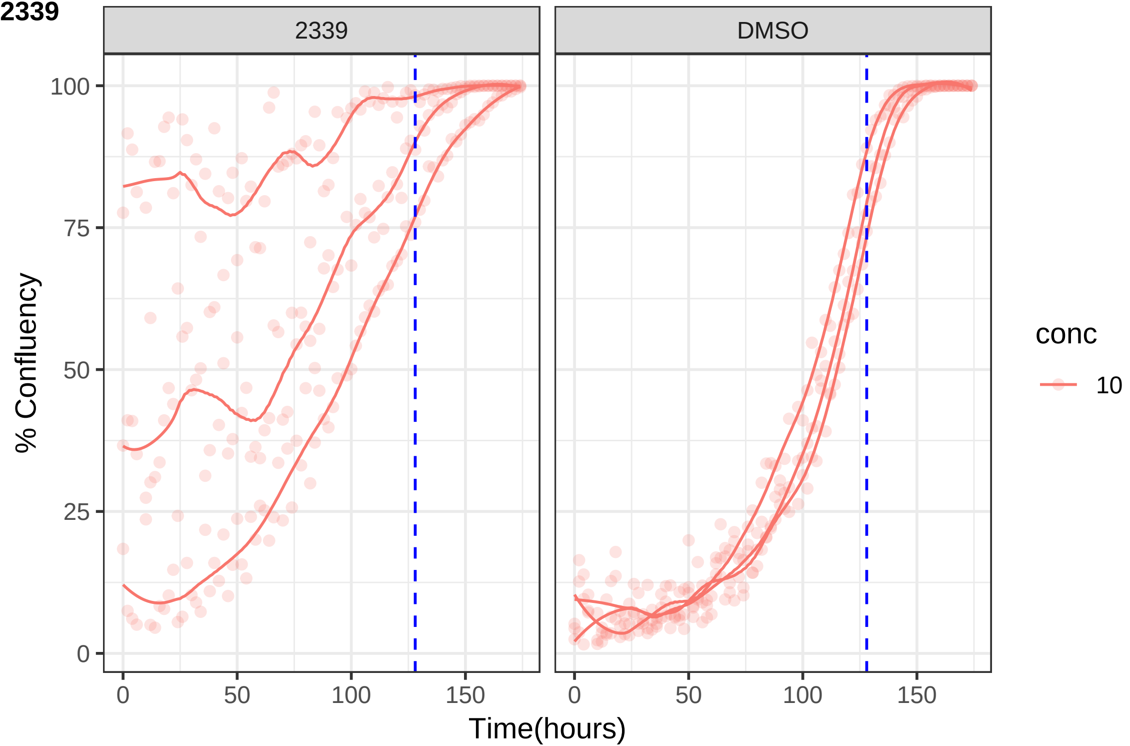

2339
DMSO
% Confluency
Time(hours)
100
75
50
25
0
0
50
100
150
conc
10

**2347**

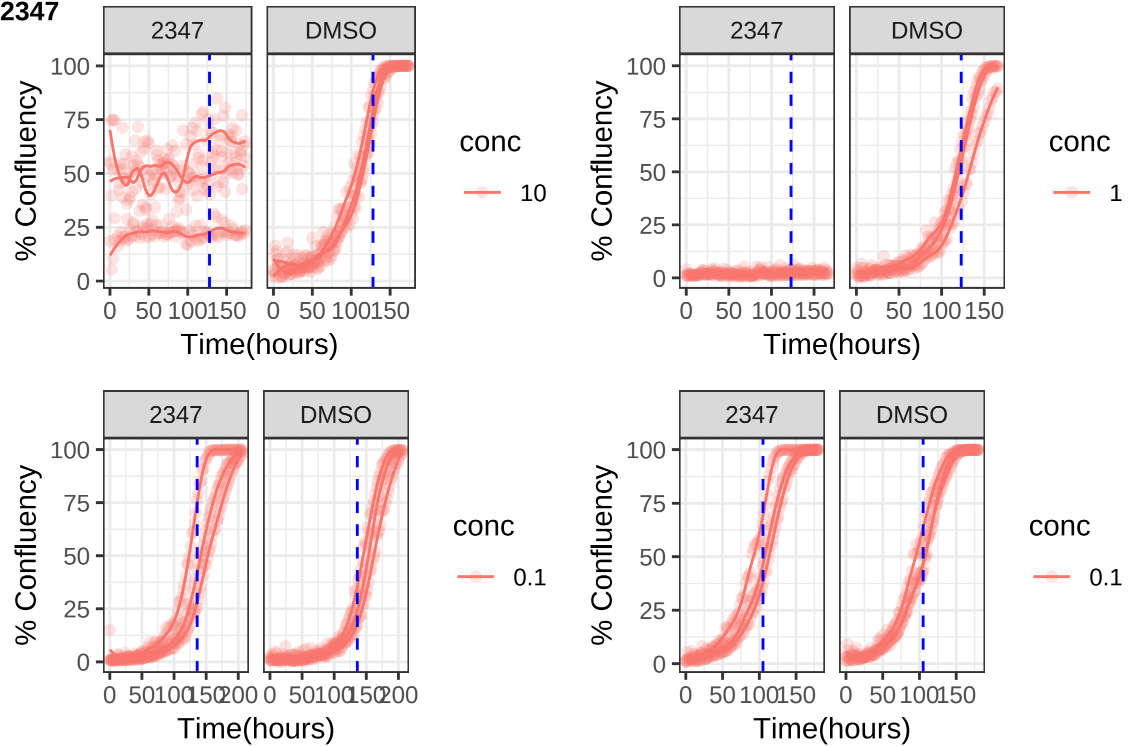

2347
DMSO
100
75
50
25
0
% Confluency
0 50 100 150
Time(hours)
conc
10
1
0.1
0 50 100 150 200

**2358**

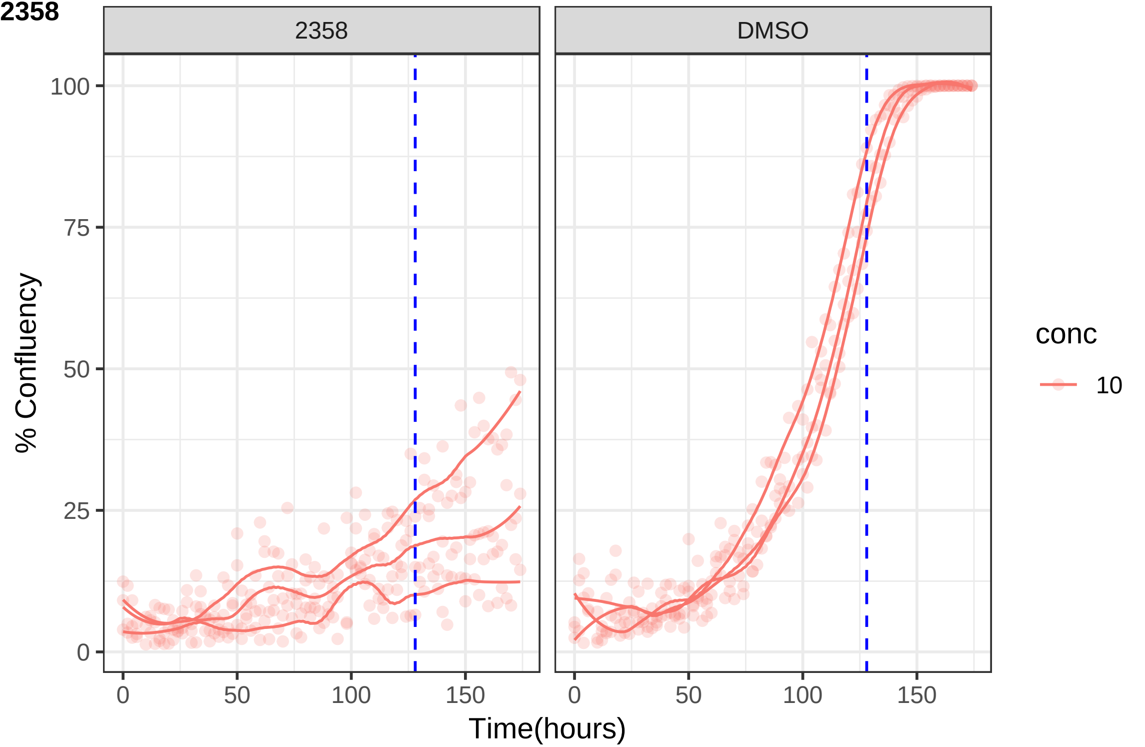

2358
DMSO
% Confluency
0
25
50
75
100
Time(hours)
0
50
100
150
conc
10

**2359**

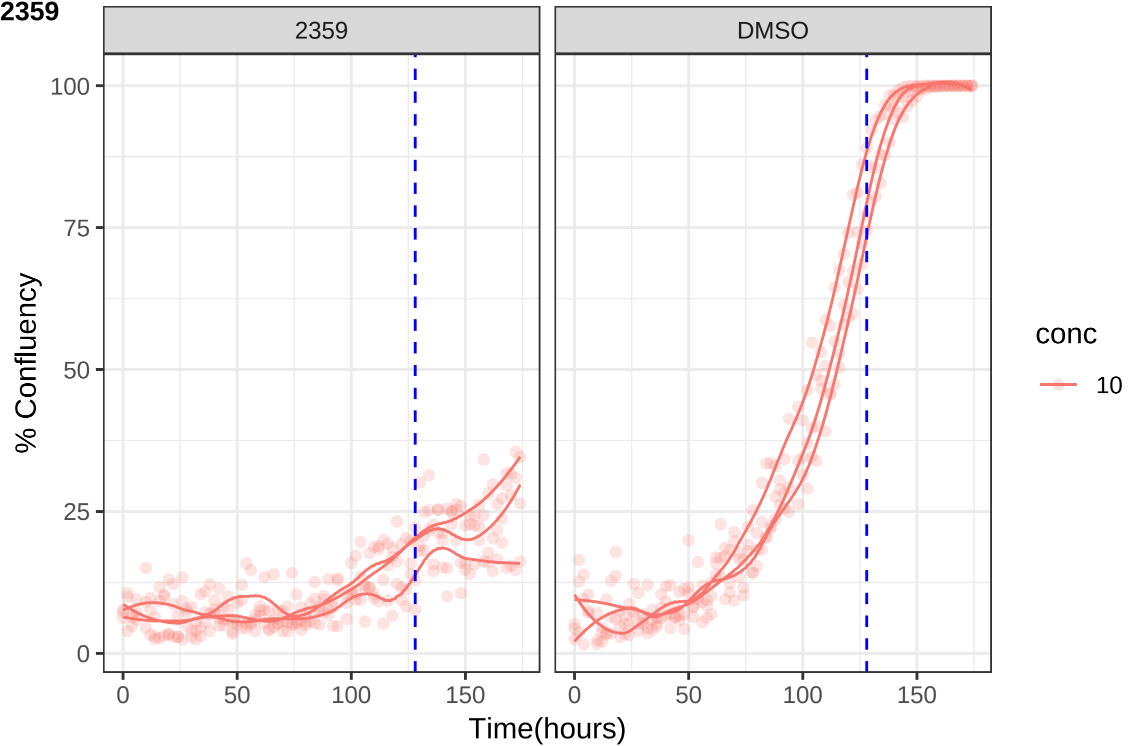

2359
DMSO
100
75
50
25
0
0
50
100
150
% Confluency
Time(hours)
conc
10

**2369**

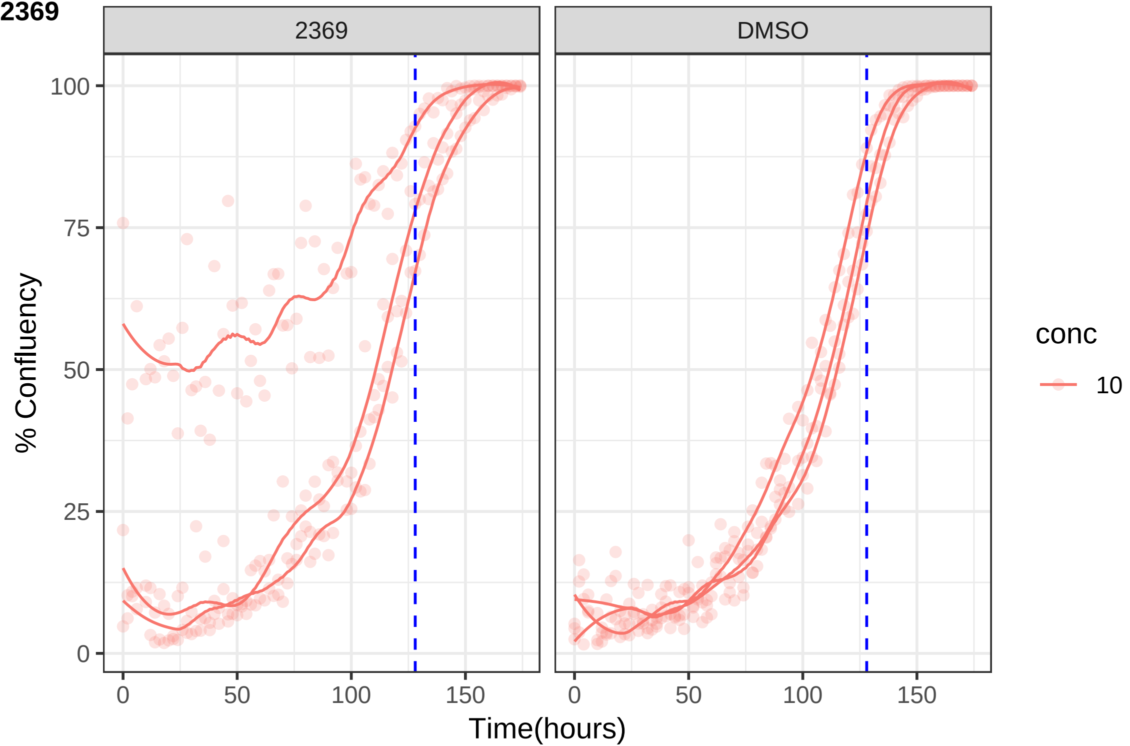
2369
DMSO
100
75
50
25
0
% Confluency
0
50
100
150
Time(hours)
conc
10

**2375**

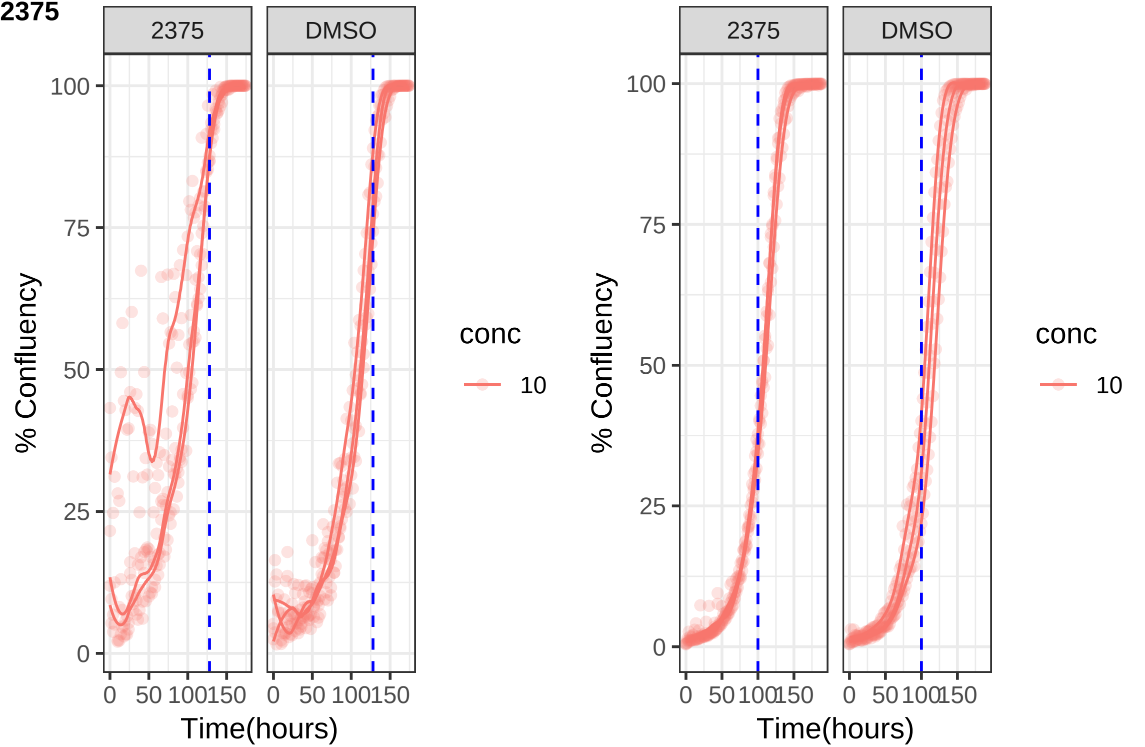

2375
DMSO
100
75
50
25
0
% Confluency
0 50 100 150
Time(hours)
conc
10
2375
DMSO
100
75
50
25
0
% Confluency
0 50 100 150
Time(hours)
conc
10

**2379**

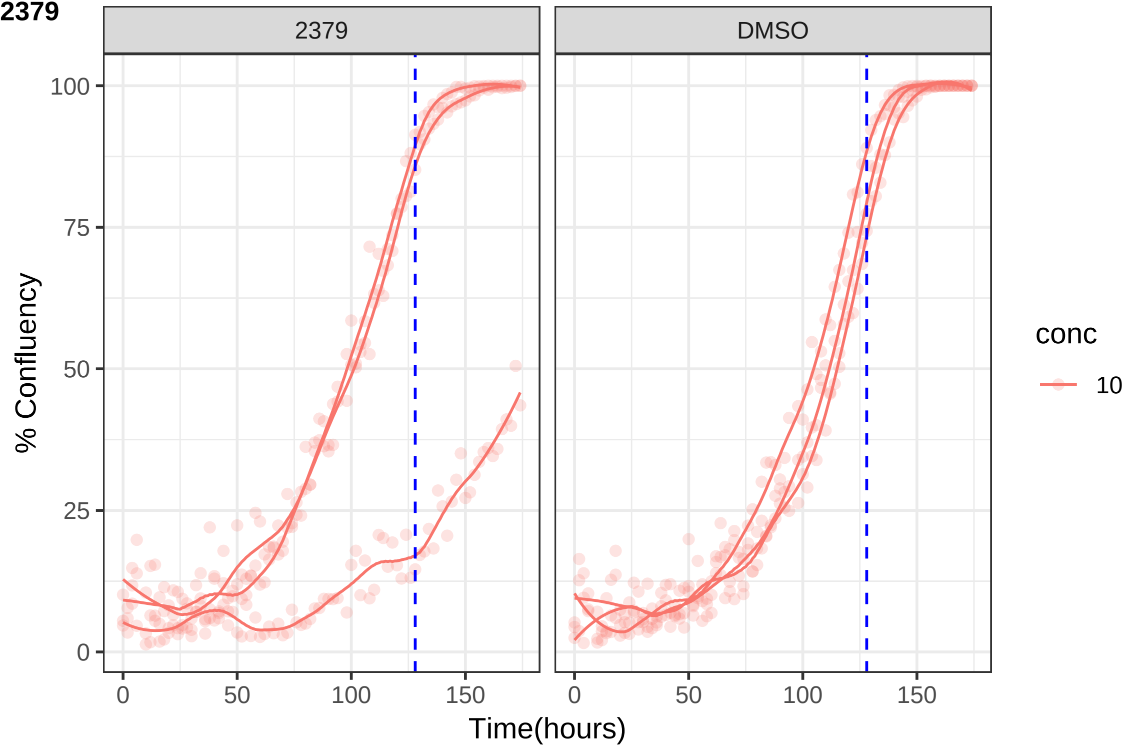
2379
DMSO
100
75
50
25
0
% Confluency
0
50
100
150
0
50
100
150
Time(hours)
conc
10

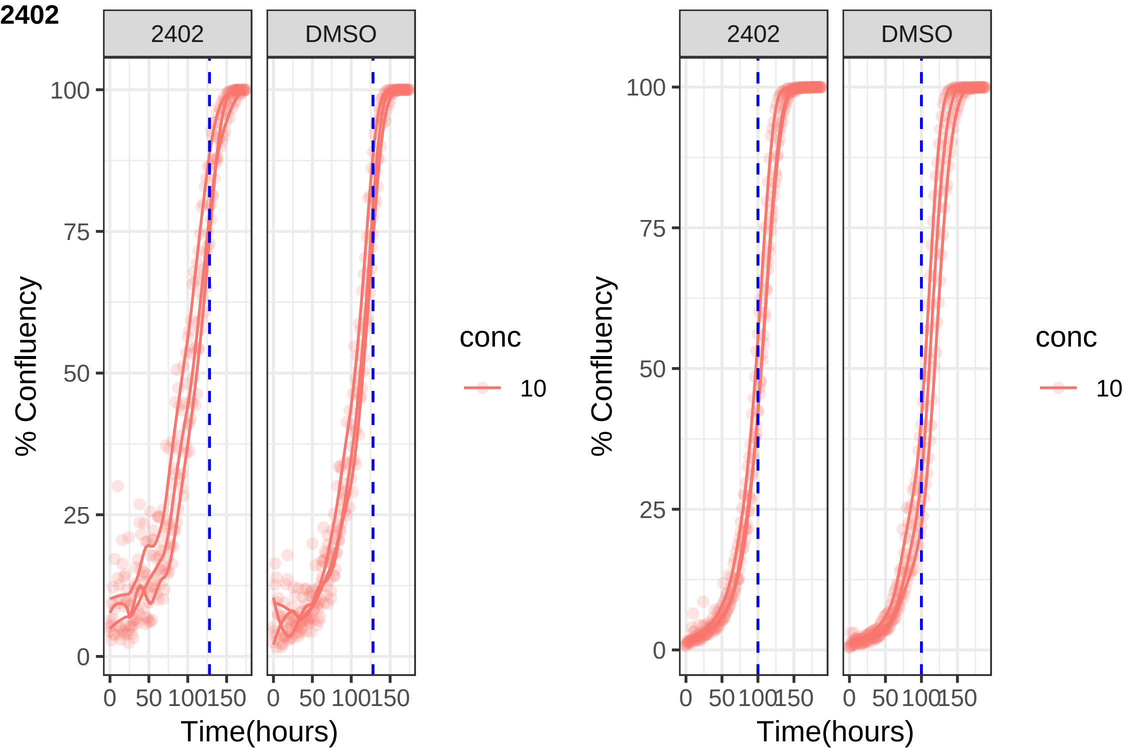

2402
2402
DMSO
100
75
50
25
0
% Confluency
0
50
100
150
Time(hours)
conc
10
2402
DMSO
100
75
50
25
0
% Confluency
0
50
100
150
Time(hours)
conc
10

**2831**

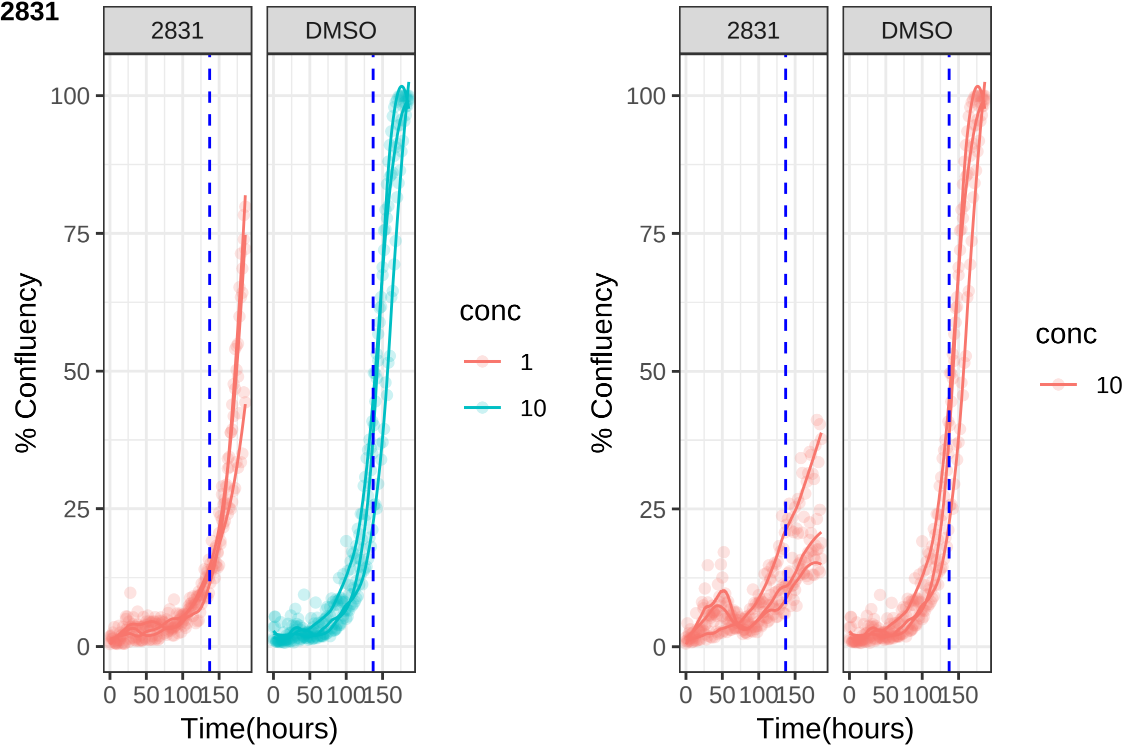
2831
DMSO
100
75
50
25
0
% Confluency
0
50
100
150
Time(hours)
conc
1
10
2831
DMSO
100
75
50
25
0
% Confluency
0
50
100
150
Time(hours)
conc
10

**2ndcontrol**

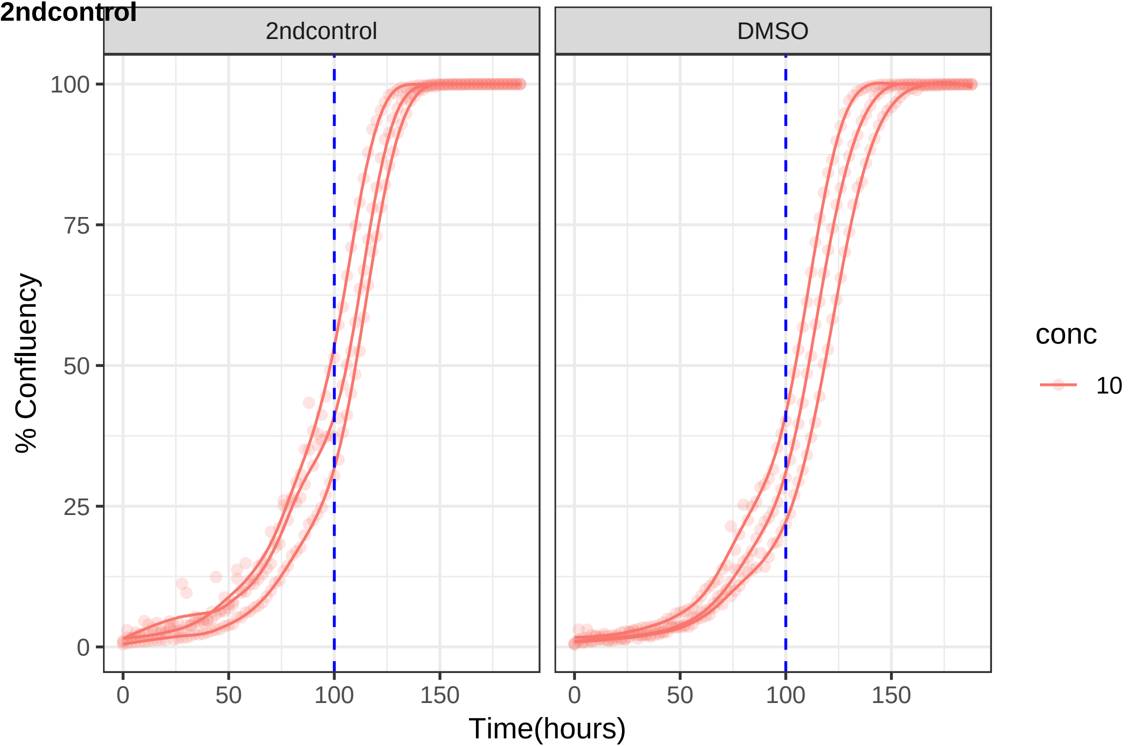

2ndcontrol
DMSO
% Confluency
0
25
50
75
100
0
50
100
150
0
50
100
150
Time(hours)
conc
10

2ndControl

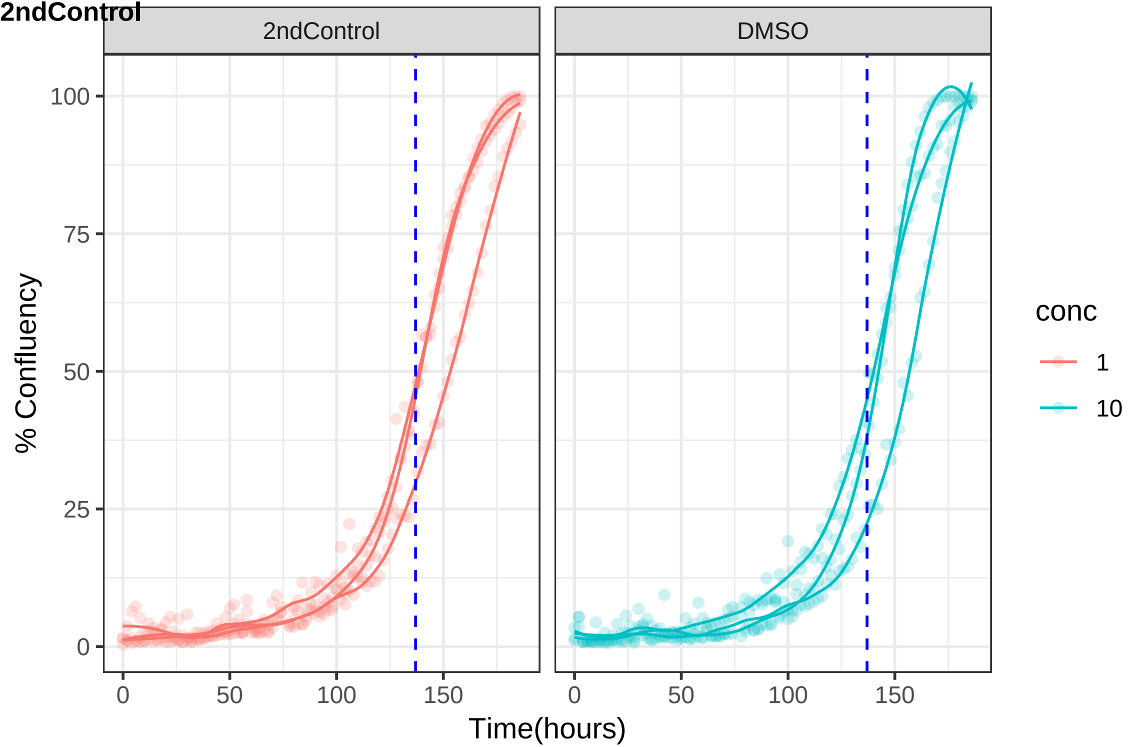
2ndControl
DMSO
% Confluency
Time(hours)
0
25
50
75
100
0
50
100
150
conc
1
10
